# The phenotypic variation of *widefins* medaka is due to the insertion of a giant transposon containing a viral genome within hoxca cluster

**DOI:** 10.1101/2025.04.28.651082

**Authors:** Rina Koita, Shunsuke Otake, Natsuki Fukaya, Kenji Yamamoto, Akiteru Maeno, Haruna Kanno, Masaru Matsuda, Akinori Kawamura

**Author notes:** Author to whom correspondence should be addressed: Akinori Kawamura, Ph.D., Division of Life Science, Graduate School of Science and Engineering, Saitama University, Saitama 338-8570, Japan, Masaru Matsuda, Ph.D., Center for Bioscience Research and Education, Utsunomiya University, Utsunomiya 321-8505, Japan.

## Abstract

Phenotypic variation in species arises from genetic differences and environmental influences on gene expression. Differences in epigenetic modifications, such as histone modifications and DNA methylation, can also contribute to phenotypic variations, even among individuals with identical genetic information. However, the underlying molecular mechanisms are not yet fully understood, particularly in vertebrates. The number of fin rays in teleosts, such as medaka, serves as a useful model for studying this variation. In a previous study, we demonstrated that the teleost Hox code plays a crucial role in determining the anterior-posterior identity necessary for the formation of dorsal and anal fins. In this study, we investigated *widefins* medaka, a spontaneous mutant displaying phenotypic variation in the number of dorsal and anal fin rays. Long-read whole-genome sequencing revealed that an extremely large transposon, *Teratorn*, containing a herpesvirus genome, was inserted into the *hoxc12a* 3’ UTR. This insertion decreased *hoxc12a* expression and, in some cases, also affected neighboring *hox* genes, resulting in variations in fin size and the presence or absence of dorsal fins. Additionally, *hoxc6a*, located 50 kb away from the insertion, was also downregulated in *widefins* medaka. These findings suggest that this large transposon insertion leads to a reduction in nearby *hox* gene expression, contributing to the phenotypic variation observed in *widefins* medaka. These results highlight the role of transposable elements and epigenetic regulation in generating phenotypic diversity in vertebrates.

## Introduction

Phenotypic variation refers to observable differences in traits among individuals of a species, such as physical appearance, physiological functions, and behavior. This diversity has been attributed to differences in genomic sequences and stochastic gene expression (Burrell et al., 2013; Raj and van Oudenaarden, 2008). However, evidence shows that genetically identical individuals can also exhibit substantial trait variation driven by epigenetic mechanisms such as DNA methylation, histone modifications, and chromatin remodeling (Jaenisch and Bird, 2003; Peaston and Whitelaw, 2006; Triantaphyllopoulos et al., 2016). These processes regulate developmental gene networks, shaping phenotypic diversity without altering the DNA sequence.

In vertebrates, the contribution of epigenetic regulation to morphological diversity is not yet fully understood. Recent studies suggest that context-dependent epigenetic states—shaped by intrinsic developmental cues and extrinsic environmental signals—can modulate developmental outcomes and generate phenotypic heterogeneity (Reik, 2007). The precise molecular pathways mediating these effects, particularly during early embryogenesis and tissue specification, are still under investigation.

The number of fin rays in the dorsal and anal fins of teleosts provides a useful model for studying phenotypic variation, as quantitative analysis is possible. In our previous study with zebrafish and medaka, we demonstrated that the teleost Hox code is essential for establishing the anterior-posterior identity required for the formation of dorsal and anal fins (Adachi et al., 2024). During animal development, Hox genes are crucial for providing positional information along the body axes and are organized into clusters on specific chromosomes (Hubert and Wellik, 2023). Medaka possess seven *hox* clusters (Kurosawa et al., 2006). In medaka, the loss of *hoxc11a* leads to the absence of the dorsal fin, while the loss of *hoxc12a* results in a posterior expansion of both the dorsal and anal fins. Additionally, *hoxc12a;hoxc13a* double mutants cause an even greater posterior expansion of these fins, highlighting the important roles of these *hoxca* genes in dorsal and anal fin formation.

Medaka serves as a valuable model organism for genetic analysis in vertebrates. Since the 1950s, the late Dr. Hideo Tomita at Nagoya University in Japan collected over 80 spontaneous mutants from wild-type populations, showcasing a variety of intriguing phenotypes (Ishikawa and Araki, 2000; Tomita, 1992). Previous studies using these medaka from the Tomita collection have greatly enhanced our understanding of active transposons, such as *Tol1* and *Tol2,* in vertebrates (Koga et al., 1995; Koga et al., 1996), as well as of vertebrate developmental mechanisms (Fukamachi et al., 2001; Inohaya et al., 2010; Moriyama et al., 2012). Additionally, large-scale screenings of artificially induced mutants, similar to those conducted with zebrafish, have produced a diverse array of mutants (Furutani-Seiki et al., 2004), further contributing to the elucidation of new molecular mechanisms (Omran et al., 2008; Porazinski et al., 2015). Moreover, medaka is a popular ornamental fish, particularly in Japanese households. Since the 2000s, the number of new and improved medaka species resulting from natural crossings among breeders for ornamental purposes has rapidly increased, with reports indicating as many as 1,000 new variants. Consequently, annual medaka fairs have been held in Japan to showcase these new traits. Thus, medaka is recognized as a unique model fish for genetic analysis, benefiting from both laboratory-generated and a significant number of spontaneous mutants. These spontaneous mutants arise from mutations induced under natural conditions, revealing unexpected genetic changes that are difficult to create using genome editing technology, potentially leading to the discovery of novel biological phenomena. Furthermore, recent advances in large-scale sequencing technology have simplified the identification of causative genomic regions in spontaneous mutants, and these mutants are expected to contribute significantly to our understanding of various biological phenomena.

In this study, we analyzed *widefins* medaka, a spontaneous mutant identified by a Japanese enthusiast. Our results indicate phenotypic variation among the *widefins*: while most individuals exhibit expanded dorsal and anal fins, a small subset either lacks a dorsal fin or possesses even larger dorsal and anal fins. Whole genome analysis (WGA) revealed that a giant transposon containing the herpesvirus genome was inserted into *hoxc12a* locus, leading to decreased expression not only of the inserted gene but also of neighboring genes. Our findings suggest that the insertion of this extraordinarily large transposon with a viral genome is responsible for the phenotypic differences observed among the *widefins*.

## Materials and methods

### Medaka Husbandry

Medaka (*Oryzias latipes*, Hd-rR strain), provided by the National BioResource Project for Medaka (NBRP Medaka), were maintained at 25°C under a 14-hour light and 10-hour dark cycle. The *widefins* medaka, a spontaneous mutant isolated by a Japanese breeding enthusiast in June 2016, have been continuously maintained outdoors through intercrosses over successive generations. For research purposes, the *widefins* were transferred to the laboratory at Utsunomiya University, where they were kept under the same conditions as the wild-type medaka. The *hoxc12a^MT1674^* and *hoxc12a;hoxc13a ^MT1675^* frameshift mutants used in this study were previously reported (Adachi et al., 2024). Embryos were collected from natural spawning, and developmental stages from embryo to adulthood were determined as previously described (Iwamatsu, 2004). All experiments involving medaka were conducted in accordance with the ARRIVE guidelines set forth by the Institutional Animal Care and Use Committee at Utsunomiya University and Saitama University.

### Alizarin Red S Staining

Adult medaka were anesthetized with 2-phenoxyethanol, euthanized, and fixed in 10% neutral formalin in PBS for 3 days. They were then rinsed with D.W. for approximately 3 days, with several changes of D.W. Next, the specimens were incubated in a 2.5% trypsin solution with saturated sodium tetraborate for 1-3 days until the entire body became soft. After several washes with D.W., the specimens were soaked in 0.05% Alizarin Red S in a 0.5% KOH solution for at least 1 day. Subsequently, the stained specimens were cleared in a 1% KOH/0.5% H_2_O_2_ solution for several hours, followed by treatment with a 0.5% KOH/0.5% H_2_O_2_ solution for another 3 days, during which the 0.5% KOH was gradually replaced with 100% glycerol. For the stained fish, images were captured using a fluorescence stereomicroscope (M205 FA, Leica).

### Ossified Bone Staining of Juvenile Medaka with Calcein

To visualize the calcified bones in larvae and juveniles, calcein staining was performed as previously described (Koita et al., 2024). After several washes, the images were captured using a fluorescence stereomicroscope (M205 FA, Leica).

### X-ray Micro-CT scan

Micro-CT scan of an adult medaka was carried out as previously described (Adachi et al., 2024; Akama et al., 2020; Maeno et al., 2024). Briefly, adult medaka were fixed with 4% paraformaldehyde in PBS at 4 °C overnight and transferred to 70% ethanol. Using an X-ray micro-CT (ScanXmate-E090S105; Comscantechno), the fixed specimens were scanned at a tube peak voltage of 85 kV and a tube current of 90 µA. For higher-resolution whole-body scanning of medaka, the whole body was scanned in four parts for each sample. For each part, the specimens were rotated 360° in 0.3° steps, generating 1200 projection images of 992 × 992 pixels. The micro-CT data of each part were reconstructed using coneCTexpress software (Comscantechno) and stored as a dataset with an isotropic resolution of 14.2 to 18.3 µm. Finally, data from four locations were combined to generate a dataset of the entire specimen. Three-dimensional image analysis was performed using OsiriX MD software (Pixmeo).

### Short-read Whole-Genome Sequencing of Widefins *Medaka*

After anesthesia with 2-phenoxyethanol, the adult male *widefins* medaka used for the complementation test was euthanized, and the entire body was fixed in 100% ethanol. Following fixation, the dissected testis was incubated overnight in a lysis buffer containing Proteinase K. Genomic DNA was then extracted using the Maxwell RSC Tissue DNA Kit (Promega) according to the manufacturer’s protocol. After the quality check of the extracted genomic DNA, whole-genome sequencing of the *widefins* medaka was performed by Gene Nex (Chemical Dojin). The raw sequence data obtained from Gene Nex were initially trimmed for adapter sequences using Trimmomatic software (Java version “21.0.1,” 2023-10-17 LTS) to extract paired-end data. A filtering process was subsequently applied to retain only high-quality sequences. Reference genomes, specifically the *Oryzias latipes* Hd-rR strain (Oryzias_latipes.ASM223467v1.dna.toplevel.fa) and the HNI strain (Oryzias_latipes_hni.ASM223471v1.dna.toplevel.fa), were downloaded along with their respective annotation files (Oryzias_latipes.ASM223467v1.113.gff3 for the Hd-rR strain and Oryzias_latipes_hni.ASM223471v1.113.gff3 for the HNI strain). Next, an index of the reference genome was created using BWA software (0.7.18-r1243-dirty), and the sequence data were mapped against the reference genome using BWA-MEM, which is suitable for Illumina paired-end reads. The mapping results were stored in SAM files, which were then converted to BAM files using Samtools (1.21) and sorted. Finally, the reference file and the BAM file of the sequence data were opened in Integrative Genomics Viewer (2.18.4) to visualize the sequences mapped by BWA, while the annotation files were used to map protein-coding genes on the genome.

### Long-read Nanopore Sequencing and Assembly of hoxca Cluster in Widefins Background

For the complementation test, a heterozygous *hoxc12a;hoxc13a ^MT1675^* mutant was crossed with *widefin*s medaka. High-molecular-weight genomic DNA was extracted from the testes of adult male progeny exhibiting elongated dorsal and anal fins, which were presumed to carry both the *hoxc12a;hoxc13a ^MT1675^* mutant allele and the *widefins* allele. The extracted genomic DNA was size-selected using the PacBio Short Read Eliminator XL kit (Pacific Biosciences) to enrich for long fragments. Library preparation was performed using the Ligation Sequencing Kit (SQK-LSK114; Oxford Nanopore Technologies), following the manufacturer’s instructions. Sequencing was carried out on a PromethION 24 platform using an R10.4.1 flow cell (FLO-PRO114M; Oxford Nanopore Technologies). Basecalling was performed using Dorado v1.0.1 with the basecalling model (dna_r10.4.1_e8.2_400bps_sup@v5.2.0). The sequencing run yielded approximately 60 Gb of total data, with a read length N50 of 6 kb. *De novo* assembly was performed using Hifiasm v0.25.0. Three contigs encompassing *hoxc11a*, *hoxc12a*, and *hoxc13a* regions were identified in the assembled genome. One contig, designated NPdata_hoxc11a_h1tg000051l, contained both *hoxc12a* and *hoxc13a* sequences, and the MT1675-specific mutations were confirmed. Another contig, NPdata_hoxc1213a_h2tg000324l, also included sequences corresponding to *hoxc12a* and *hoxc13a*, while *hoxc11a* was found in NPdata_hoxc11a_h1tg000083l. Due to evidence of misassembly in the region containing the *Teratorn* transposon, we performed a targeted reassembly using only the reads associated with the two aforementioned contigs. This reassembly yielded a 204 Mbp sequence that included a partial wild-type *hoxc11a*, a complete *hoxc12a* locus, and an approximately 184 kb *Teratorn* transposon sequence.

### Whole-mount In situ hybridization

Whole-mount *in situ* hybridization was performed as previously described (Adachi et al., 2024). For the stained fish, the images were captured using a fluorescence stereomicroscope (M205 FA, Leica).

## Results

The *widefins* medaka, a spontaneous mutant characterized by dorsal and anal fins that expand posteriorly, was isolated by a Japanese medaka breeder (Supplementary Fig.1). This phenotype resembles that of the medaka *hoxc12a* and *hoxc12a;c13a* frameshift-induced mutants that we previously created using CRISPR-Cas9 (Adachi et al., 2024). To compare the phenotypes of *widefins* with those of the *hoxc12a* and *hoxc12a;c13a* mutants, we first analyzed the skeletal structures of adult *widefins* obtained from intercrosses using alizarin red staining and micro-CT scans (Fig. 1a-h). The analysis of fin ray counts revealed that some individuals of *widefins* exhibited varying phenotypes, unlike the consistent similarities observed in *hoxc12a* or *hoxc12a;c13a* mutants. Generally, the *widefins* medaka displayed a greater number of dorsal fin rays compared to the wild-type (*n*=9/11); many individuals had a fin ray count comparable to that of *hoxc12a* mutants (Fig. 1d, h, i), while a few had more fin rays, though still fewer than those of *hoxc12a;c13a* mutants (Fig. 1f, i). Notably, one individual examined among the *widefins* lacked a dorsal fin (Fig. 1e, i), sharply contrasting with the typical phenotype of *widefins*. This dorsal fin-absent phenotype resembles that of the spontaneous *dorsalfinless* medaka, which encodes the hypomorphic allele of *hoxc11a* (Adachi et al., 2024). Additionally, the anal fin of the *widefins* exhibited a greater number of fin rays; typical individuals showed a comparable or slightly higher count than *hoxc12a* mutants (Fig. 1d, j). In some individuals, the number of fin rays was comparable to that of *hoxc12a;c13a* mutants (Fig. 1f, j). Furthermore, the number of vertebrae in *widefins* medaka also displayed variation similar to the phenotypes between *hoxc12a* mutants and *hoxc12a;c13a* mutants (Fig. 1k). These results indicate that *widefins* exhibit individual variations in the phenotypes of their dorsal and anal fins, as well as vertebrae. Overall, the phenotypes of *widefins* medaka are similar to those of *hoxc12a* mutants or *hoxc12a;c13a* mutants, showing a slightly stronger phenotype than *hoxc12a* mutants but weaker than *hoxc12a;c13a* mutants. Furthermore, in rare instances, *widefins* display a dorsalfinless phenotype resembling that of *hoxc11a* mutants.

**Figure 1.**
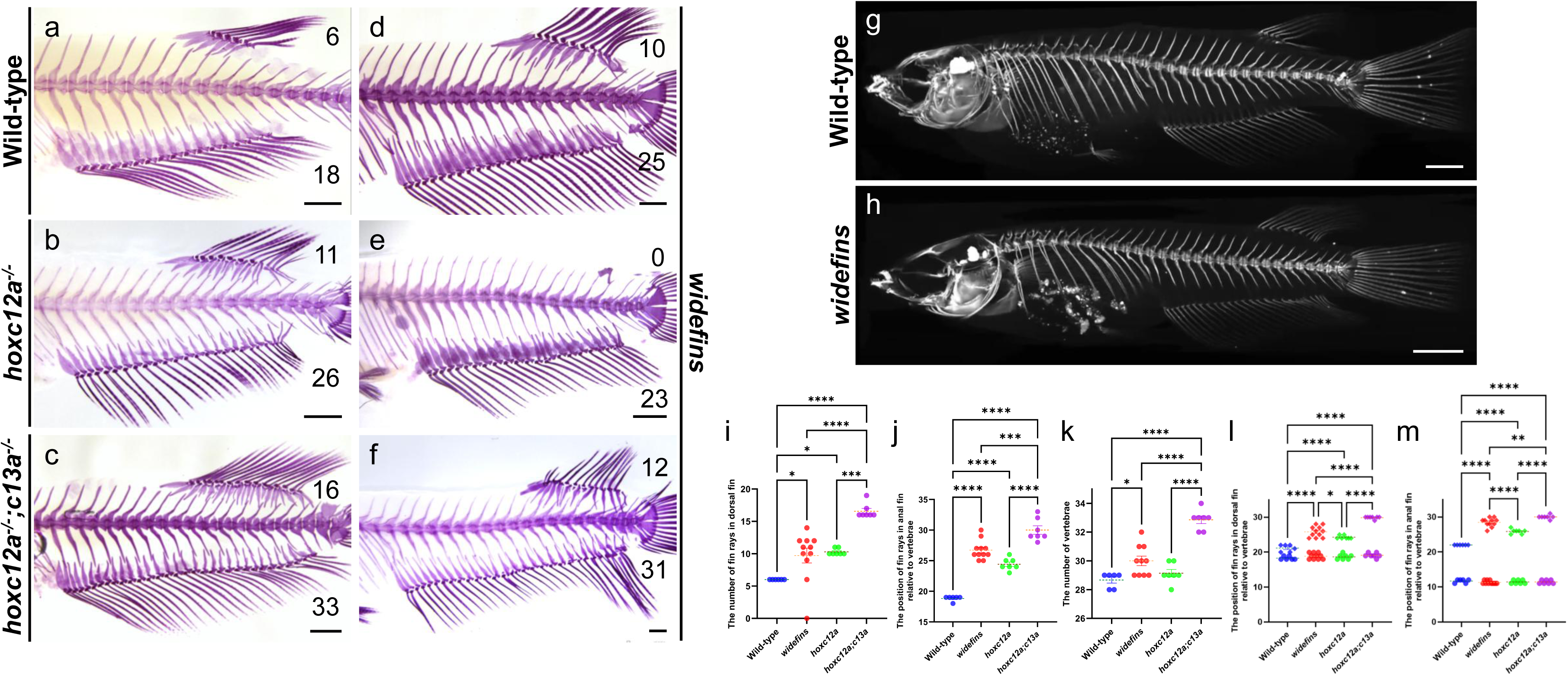
*widefins* medaka exhibit a phenotypic variation, particularly in the dorsal fin. (a-d) Skeletal structures of the posterior body in wild-type (a), *hoxc12a*^-/-^ (b), *hoxc12a*^-/-^ *;c13a*^-/-^ (c), and *widefins* (d-f) fish. The ossified bones were stained with Alizarin Red S. The number of dorsal fin rays for each individual is indicated in the upper right corner, while the number of anal fin rays is shown in the lower right corner. Three different phenotypes of *widefins* medaka are presented, illustrating individual variations in the number of dorsal and anal fin rays. Scale bar: 2 mm. (g, h) Micro-CT scan images of wild-type and *widefins* adult medaka. Scale bar: 2 mm. 3D movies are available in Supplementary Movies 1 and 2. (i-k) Comparison of the number of fin rays in the dorsal fin (i) and anal fin (j), as well as the number of vertebrae (k) among the mutants. (l, m) Comparison of the anterior and posterior ends of vertebrae attached to the fin rays of the dorsal fin (l) and anal fin (m) via the radials. The anterior end of the vertebra is indicated by a circle, while the posterior end is marked with a rhomboid. A single *widefins* fish with a missing dorsal fin (e) was excluded from the statistical analysis in (l). For all statistical analyses, the Tukey-Kramer test was performed on data obtained from Alizarin Red S-stained fish and micro-CT scans. \**P*□<□0.05, \*\**P* <□0.01, \*\*\**P* <□0.001, and \*\*\*\**P*□<□0.0001.

To determine whether the abnormalities in *widefins* are associated with the loss of function of *hoxc12a* and *hoxc13a*, we conducted a genetic complementation test by crossing *widefins* with frameshift-induced *hoxc12a;c13a* heterozygous medaka. Analysis of 21 juveniles from this cross revealed that the *widefins* and *hoxc12a;c13a* mutations do not complement each other. We observed some individuals with a normal number of fin rays (*n*=9) and others with an increased number of fin rays (*n*=12) (Fig. 2a-g). Genotyping for frameshift mutations showed that all juvenile fish with a normal number of fin rays were identified as *hoxc12a^+/wf^;c13a^+/wf^*, while all juvenile fish with an increased number of fin rays were identified as *hoxc12a^-/wf^;c13a^-/wf^*. These genetic results indicate that *widefins* encodes a recessive mutation associated with the loss of function of the hox genes, specifically *hoxc12a* and *hoxc13a*.

**Figure 2.**
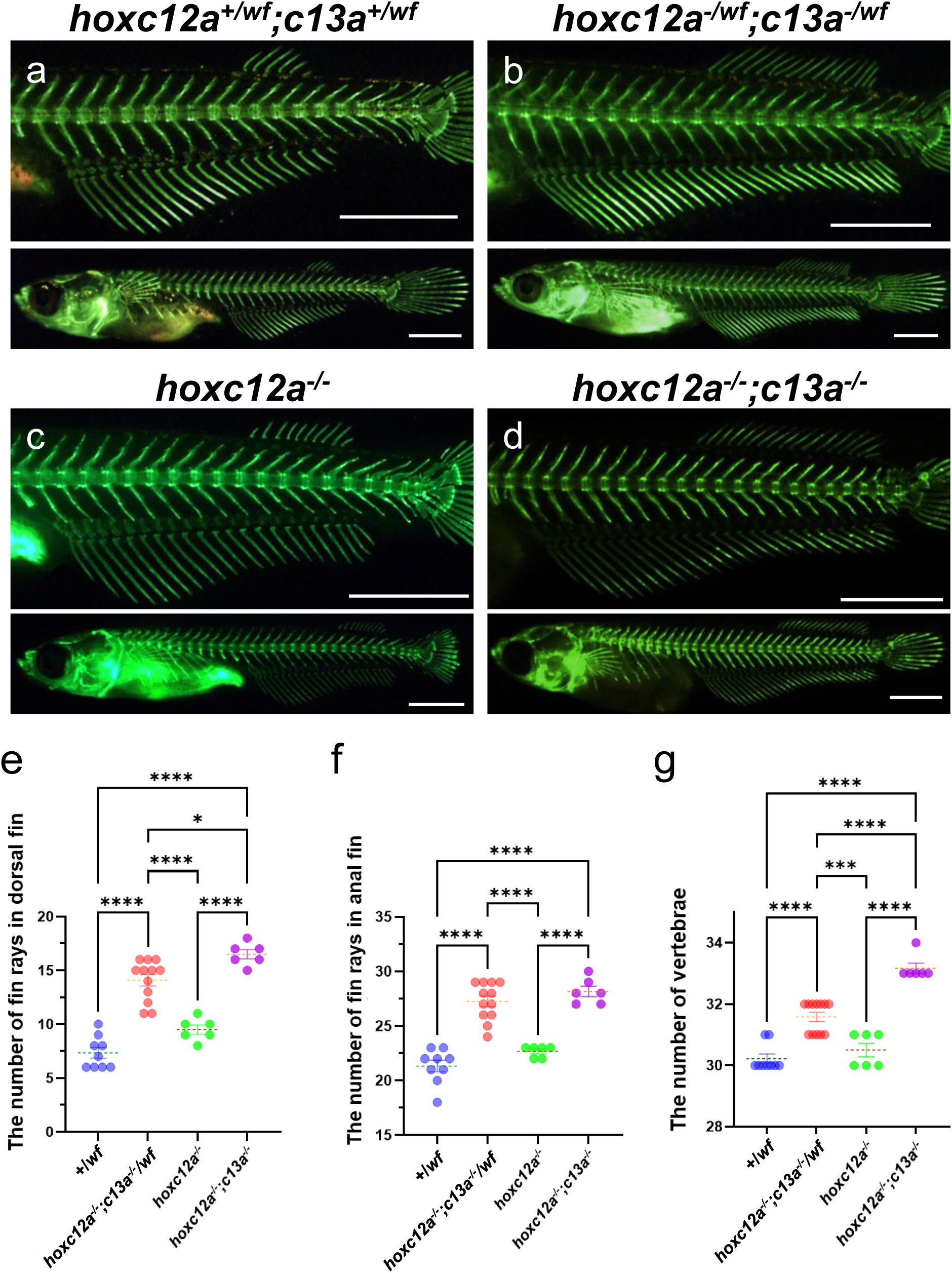
*widefins* medaka do not complement *hoxc12a;c13a* mutations. (a-d) The calcified bones of juvenile fish, obtained by crossing male *widefins* with female *hoxc12a;c13a* heterozygous mutants, were visualized using calcein staining. Enlarged views (top) and the entire body (bottom) of the same representative fish are shown. After capturing images, the presence or absence of frameshift mutations in *hoxc12a* and *hoxc13a*, introduced by CRISPR-Cas9, was examined by PCR as previously described (Adachi et al., 2024). Scale bar: 1 mm. (e-g) Comparison of the number of fin rays in the dorsal fin (e) and anal fin (f), as well as the number of vertebrae (g). The data, obtained from *hoxc12a^+/wf^;c13a^+/wf^*, *hoxc12a^-/wf^; c13a^-/wf^*, *hoxc12a^-/-^*, and *hoxc12a*^-/-^*;c13a*^-/-^ fish, were compared. Previously described data for *hoxc12a^-/-^*and *hoxc12a*^-/-^*;c13a*^-/-^ fish were also included for comparison (Adachi et al., 2024). The Tukey-Kramer test was performed. \**P*□<□0.05, \*\*\**P* <□0.001, and \*\*\*\**P*□<□0.0001.

Next, we performed short-read whole-genome sequencing (WGS) using Illumina to identify the genomic region responsible for the phenotypic features observed in *widefins* medaka. Our results suggested that the mutation is likely located in the genomic region encompassing *hoxc12a* and *hoxc13a* loci. The extensive sequence data obtained from WGS were mapped to the reference wild-type medaka genome, with the read sequences typically overlapping. However, in the 3’ UTR of *hoxc12a*, reads from both strands were joined to an unrelated sequence, indicating the insertion of a DNA element (Fig. 3a). Closer examination of the *mapped* reads revealed that the inserted sequence corresponded to both ends of a DNA transposon known as *Teratorn*. Originally identified in the medaka spontaneous mutant *Double analfin* (Moriyama et al., 2012), *Teratorn* is unique among DNA transposons in that it is exceptionally large, measuring 180 kb in length, and represents a fusion between a herpes virus genome and *piggyBac*-type transposon elements (Inoue et al., 2017). Using PCR, we confirmed that the 5’ and 3’ ends of the *Teratorn* transposon sequences were indeed inserted into the 3’UTR of *hoxc12a* in *widefins* (Fig. 3b). Furthermore, long-read nanopore sequencing confirmed that a DNA fragment of approximately 184 kb is inserted into *hoxc12a* locus of *widefins*. This fragment closely resembles the previously reported *Teratorn* sequence and similarly contains numerous open reading frames derived from a herpesvirus genome (Supplementary Fig.2 and Fig.3). Since the coding sequences of both *hoxc12a* and *hoxc13a* remain intact in *widefins*, our results suggest that the insertion of this giant transposon in *hoxc12a* locus is responsible for the *widefins* phenotype.

**Figure 3.**
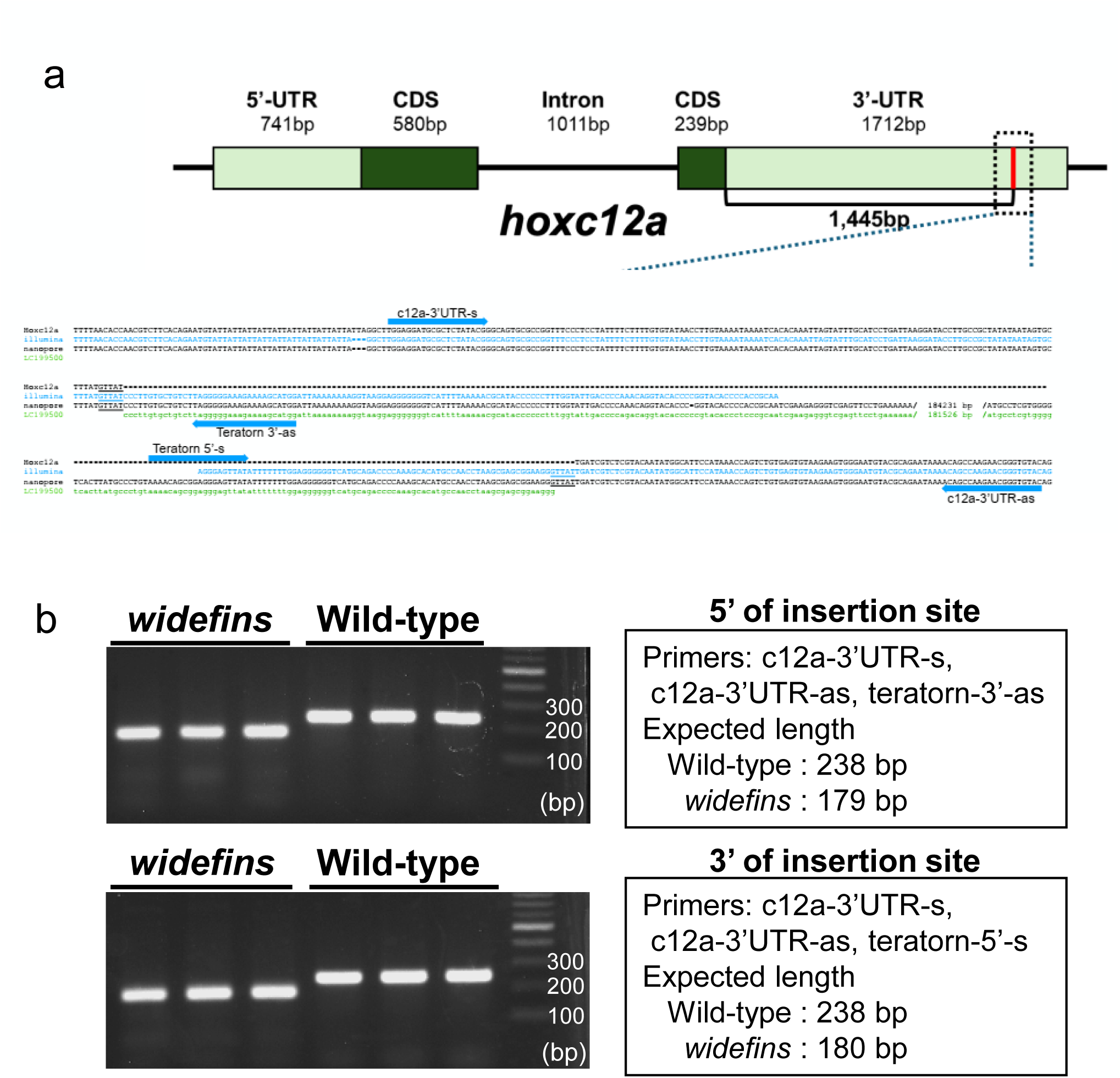
Insertion of the *Teratorn* transposon in the 3’ UTR of *hoxc12a*. (a) Schematic representation of the medaka *hoxc12a* genomic structure, which consists of two exons. Below the diagram are the alignment results of the normal *hoxc12a* sequence from Ensembl medaka, the Illumina short-read and Nanopore *hoxc12a* sequences from *widefins*, and the previously reported *Teratorn* transposon sequence (accession no. LC199500). The positions of the primers used for PCR-based genotyping are shown in arrows. (b) Genotyping of wild-type and *widefins* medaka was performed using three primers (indicated on the right side of the panel) for each near end of the *Teratorn* transposon insertion site. Genomic DNA from three individuals of each genotype was used as templates for PCR. PCR products were separated on a 2% agarose gel in 0.5x TBE buffer. In *widefins*, no PCR products were amplified with the c12a-3’UTR-s and c12a-3’UTR-as primers due to the *Teratorn* insertion.

To investigate whether the insertion of *Teratorn* affects the expression of *hoxc12a*, we next compared the expression patterns of hox genes in wild-type and *widefins* embryos using whole-mount *in situ* hybridization. We found that the expression of *hoxc12a* was significantly downregulated in *widefins* (Fig. 4c, d; *n*=8/8), which could lead to the expansion of dorsal and anal fins, as observed in *hoxc12a* mutants. Additionally, we examined the expression of neighboring hox genes. Although not to the same extent as *hoxc12a*, the expression patterns of *hoxc11a* were also reduced in *widefins* (Fig. 4a, b; *n*=7/8). If the reduction in *hoxc11a* expression were more pronounced, the loss of *hoxc11a* function would result in the absence of a dorsal fin in *widefins* (Fig. 1e), because *hoxc11a* mutants lack the dorsal fin. Furthermore, we analyzed the expression pattern of *hoxc13a*, but could not detect significant differences (Fig. 4e, f; *n*=9/9). These results suggest that the insertion of the *Teratorn* transposon in the 3’UTR of *hoxc12a* significantly reduces the expression of *hoxc12a* and contributes to the decreased expression of neighboring hox genes in the *hoxca* cluster.

**Figure 4.**
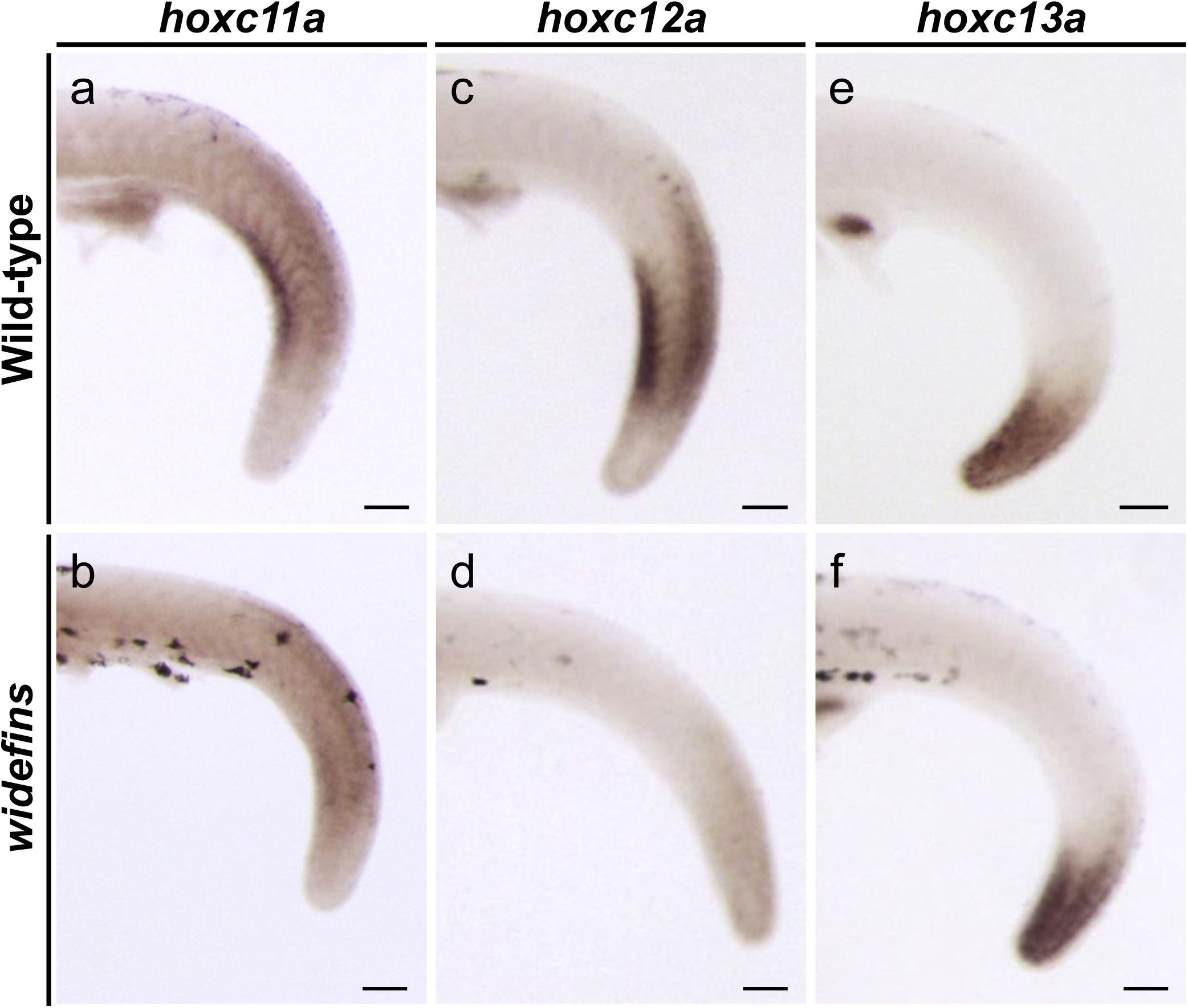
Significantly decreased expression of *hoxc12a* and reduced expression of *hoxc11a* in *widefins* medaka. (a-f) Expression patterns of *hoxc11a*, *hoxc12a*, and *hoxc13a* were analyzed between wild-type and *widefins*. Whole-mount *in situ* hybridization was performed on embryos at stages 29-30. For each staining, at least three different specimens were used to confirm reproducibility. Lateral view of the posterior body. Scale bar: 100 μm.

Since the insertion of the *Teratorn* transposon decreased the expression of *hoxc12a* and nearby *hoxc11a*, we investigated whether this effect extends further. In medaka, *hoxc6a* locus is located approximately 50 kb away from the insertion site (Fig. 5a). Loss of function of medaka *hoxc6a* results in an anteriorization phenotype characterized by the absence of pleural ribs in the second vertebra, where rib formation should occur; instead, rib formation begins at the third vertebra (Maeno et al., 2024). Following a detailed examination of *widefins* medaka, we found that the most anteriorly formed rib, which differed from that of wild-type, was formed from the third vertebra (Fig. 5b-g; *n*=5/5). This phenotype resembles that of *hoxc6a* mutants, so we compared the expression patterns of *hoxc6a* between wild-type and *widefins* embryos. Using whole-mount in *situ* hybridization, we found that the expression of *hoxc6a* was decreased in *widefins* compared to wild-type (Fig. 5h, i; *n*=4/5). Taken together, these results suggest that the insertion of a giant *Teratorn* transposon containing viral genes reduces the expression of *hoxc6a*, which is located 50 kb away from the insertion site.

**Figure 5.**
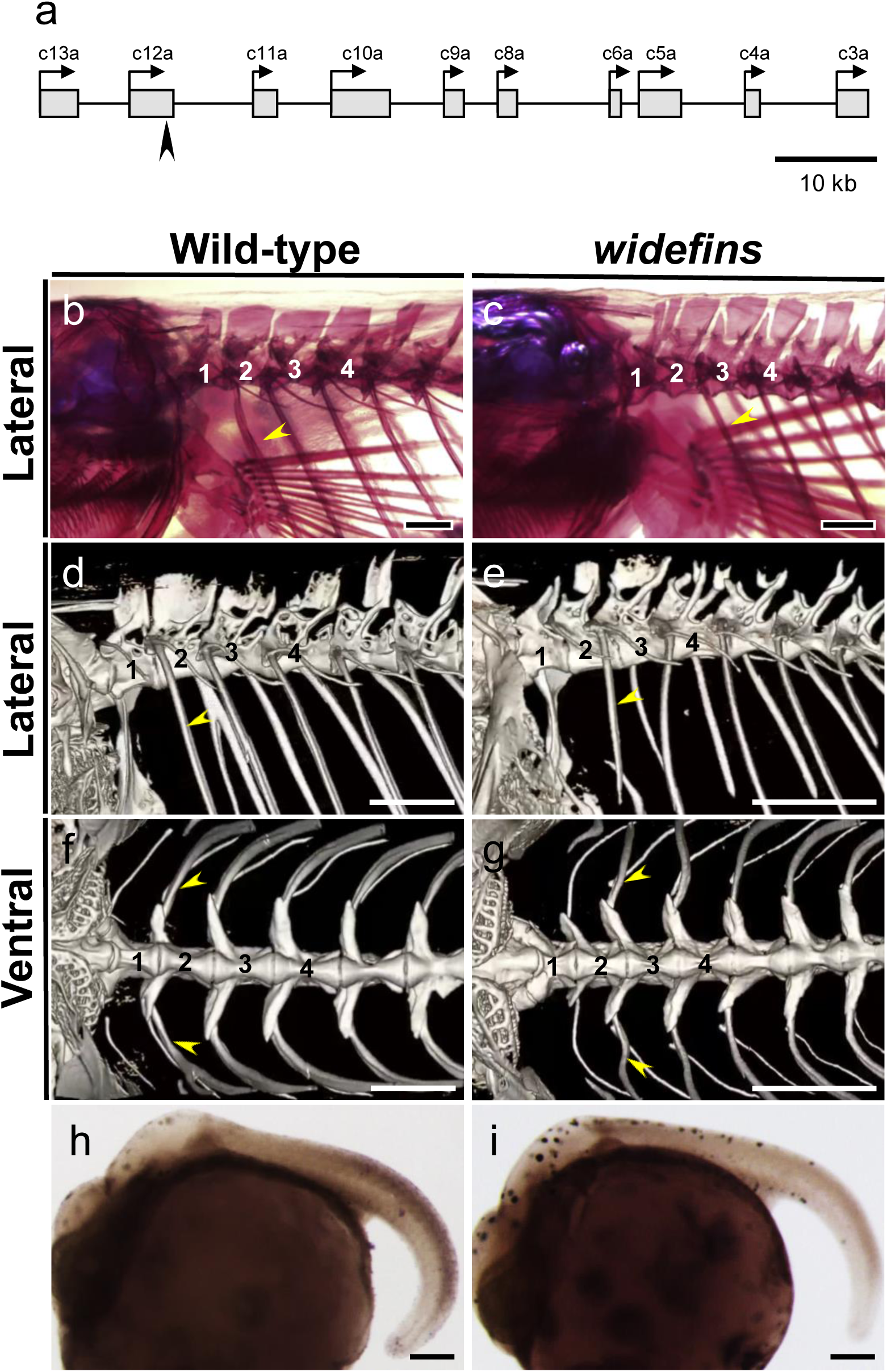
The expression of *hoxc6a* is decreased, resulting in anteriorizing vertebral patterning in *widefins* medaka. (a) Schematic representation of medaka *hoxca* cluster. Boxes represent the hox genes, and arrows indicate their transcriptional orientation. The arrowhead marks the insertion site of the *Teratorn* transposon in the 3’UTR of *hoxc12a*. (b, c) Skeletal structures of the anterior vertebrae in wild-type and *widefins*. The ossified bones were stained with Alizarin Red S. Numbers indicate the number from the most anterior vertebra. Yellow arrowheads point to the anteriormost pleural rib in both wild-type and *widefins.* Scale bar: 2 mm. (d-g) Micro-CT scanning of the anterior vertebrae in wild-type and *widefins.* Scale bar: 2 mm. (h, i) Expression patterns of *hoxc6a* in wild-type and *widefins* embryos. Whole-mount *in situ* hybridization was performed on embryos at stages 29-30. At least three different specimens were used for each staining to confirm reproducibility. Lateral view of the posterior body. Scale bar: 100 μm.

## Discussion

In this study, we analyzed *widefins*, a spontaneous medaka mutant isolated by a Japanese enthusiast for ornamental breeding. Our analysis revealed that the *widefins* phenotypes can be categorized into three groups based on differences in the number of dorsal and anal fin rays (Fig. 1). This variation is particularly pronounced in the dorsal fin, where the number of fin rays differed significantly among individuals, with some *widefins* even lacking a dorsal fin entirely. WGS analysis further identified the insertion of approximately 184 kb *Teratorn* transposon, containing a herpes virus genome, into the 3’ UTR of *hoxc12a*. Our results suggest that this insertion reduces the expression of *hoxc12a* and, in some cases, neighboring hox genes, thereby contributing to phenotypic variation observed in *widefins*.

One possible explanation for the phenotypic variation observed in *widefins* medaka is that the insertion of the *Teratorn* transposon might occasionally influence the expression of the inserted *hoxc12a* and, in some cases, nearby hox genes. The *Teratorn* transposon is unusually large, incorporating elements from a herpesvirus genome that comprise approximately 80% of its total sequence (Inoue et al., 2017). In other systems, the integration of foreign DNA, such as viral genomes and transposable elements, has often been associated with gene silencing through heterochromatin formation (Cabrera et al., 2022; Deniz et al., 2019; Slotkin and Martienssen, 2007). Epigenetic markers of heterochromatin, such as histone H3 trimethylation at lysine 9 (H3K9me3) and CpG methylation, can spread into adjacent regions via positive feedback of histone methyltransferase recruitment (Al-Sady et al., 2013; Grewal and Jia, 2007; Grewal, 2023). This process can, but does not necessarily, influence regions neighboring the insertion site. However, the mere insertion of a transposon does not necessarily impact the surrounding region. In our previous study of spontaneous *dorsalfinless* medaka mutants, which lack a dorsal fin, we demonstrated that a transposon of about 12 kb was inserted into the 3’ UTR of *hoxc11a* (Adachi et al., 2024), adjacent to *hoxc12a* where the *Teratorn* insertion was identified in *widefins* medaka in the current study. Interestingly, this 12 kb DNA transposon significantly reduces the expression of *hoxc11a*, but it does not appear to influence the expression of nearby hox genes, including *hoxc12a*. Given that *Teratorn* spans 180 kb (Inoue et al., 2017), it implies that around 150 kb of virus-derived long sequences have integrated into *hoxc12a*, suggesting a high capacity to form highly condensed heterochromatic regions. Since the 30-40 copies of *Teratorn* are distributed throughout the haploid medaka genome (Inoue et al., 2017) and cannot be individually distinguished, we cannot provide direct evidence that heterochromatin is indeed formed at the *Teratorn* insertion into *hoxc12a* in *widefins*. Nevertheless, it remains plausible that the insertion of *Teratorn* could contribute to heterochromatin formation at the insertion site, and, in some individuals, that this effect might extend into the surrounding region, thereby modulating the expression of neighboring hox genes.

Based on this model, we speculate that in most *widefins* individuals, the predominant effect of the *Teratorn* insertion is a reduction in the expression of the inserted *hoxc12a*. Whether this decrease reflects transcriptional repression and/or reduced mRNA stability remains unclear. Although expression of the adjacent *hoxc11a* also appears somewhat reduced, it may still be sufficient to permit dorsal fin development. Thus, *widefins* generally display a phenotype resembling *hoxc12a* mutants, characterized by wide dorsal and anal fins (Fig. 6a). In some individuals, however, heterochromatin associated with *Teratorn* could extend further, potentially affecting neighboring hox genes. If both *hoxc11a* and *hoxc12a* were impaired, medaka could exhibit both dorsal fin loss and anal fin expansion (Fig. 6c). Similarly, if *hoxc12a* and *hoxc13a* were both afected, this could result in even wider dorsal and anal fins, similar to *hoxc12a;hoxc13a* mutants (Fig. 6d). The variation in phenotypes among *widefins* individuals resembles position effect variation (PEV), which has been extensively studied in *Drosophila*. PEV arises from stochastic silencing of genes near heterochromatin as a result of chromosomal rearrangements (Elgin and Reuter, 2013). Our analysis, therefore, raises the possibility that the insertion of a giant transposon such as *Teratorn* may generate PEV-like phenomena in teleosts.

**Figure 6.**
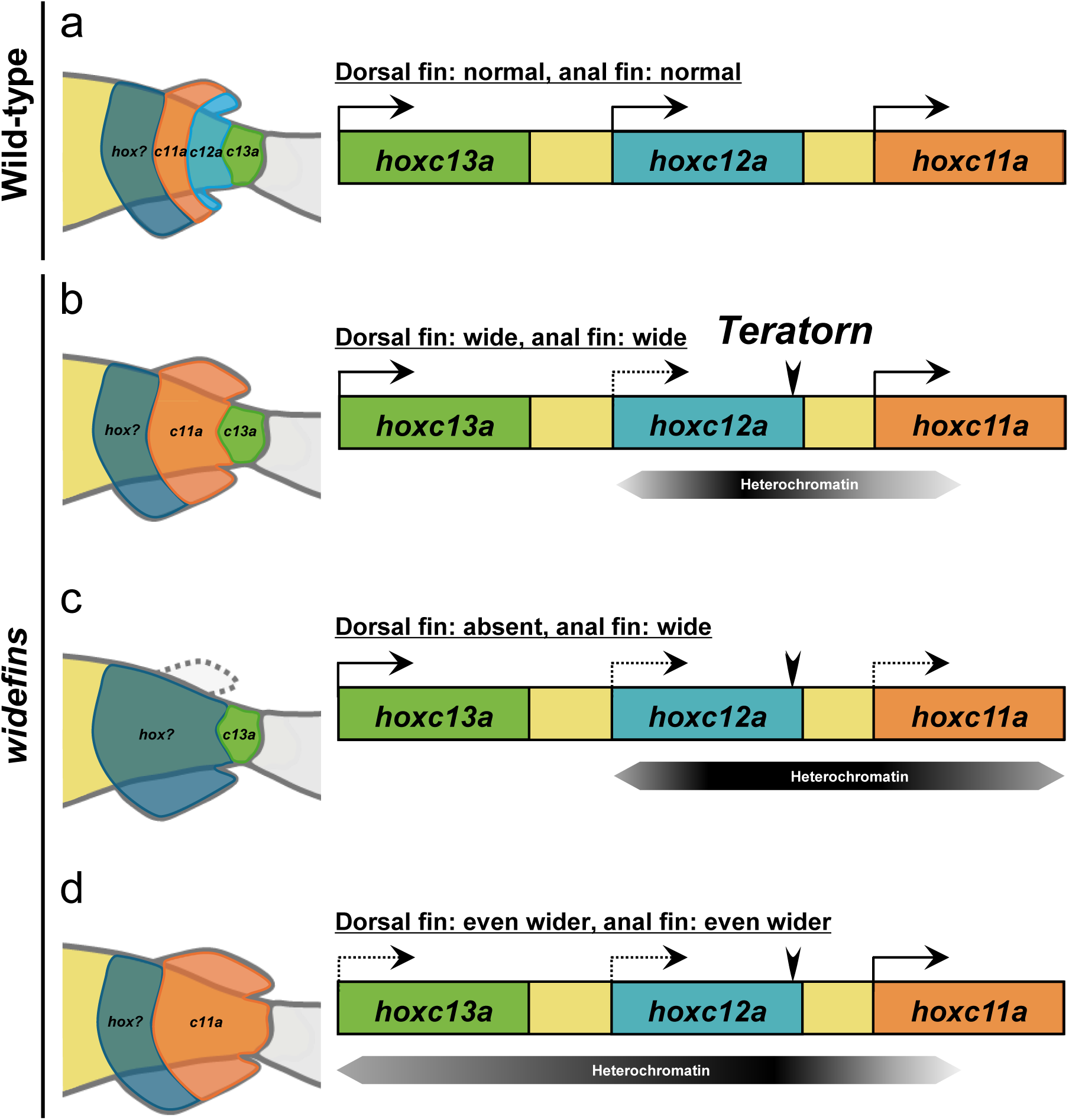
Proposed model illustrating phenotypic variation among *widefins* individuals. (a) In medaka, the genomic loci of *hoxc11a*, *hoxc12a*, and *hoxc13a* are aligned in the same transcriptional direction (arrow in right schematic). In the posterior body, these *hox* genes define regional identities competent for the formation of dorsal and anal fins (left schematic). (b) The insertion of a giant *Teratorn* transposon containing a viral genome within *hoxc12a* (arrowhead) leads to the formation of heterochromatin in the surrounding region, significantly reducing *hoxc12a* expression (dashed arrow). This reduction results in wide dorsal and anal fins, resembling the phenotype observed in *hoxc12a* mutants (Adachi et al., 2024). (c, d) In some instances, the insertion of the *Teratorn* transposon may expand the heterochromatin to a broader genomic region, affecting nearby *hox* genes. When the heterochromatin expands to the *hoxc11a* locus, *hoxc11a* expression is considerably reduced, resulting in a wide anal fin due to *hoxc12a* deficiency, as well as the absence of a dorsal fin seen in *hoxc11a* mutants (Adachi et al., 2024). If the heterochromatin extends to the *hoxc13a* locus, the expression of *hoxc13a* is also reduced, leading to fish with even wider dorsal and anal fins, as observed in *hoxc12a;hoxc13a* mutants (Adachi et al., 2024).

Additionally, the *hoxc6a* locus, located approximately 50 kb from the insertion site, was also found to be downregulated. While one possible explanation is the spreading of heterochromatin, this alone seems insufficient to account for the effect, as the *hoxc6a* locus is relatively distal from the insertion site. Instead, vertebrate Hox genes are known to be intricately regulated by enhancers both within and outside the clusters, and it has been well established that Hox clusters are organized within topologically associating domains (TADs) (Acemel et al., 2016; Afzal and Krumlauf, 2022; Andrey et al., 2013; Mallo and Alonso, 2013), which provide structural frameworks for long-range enhancer–promoter interactions (Dixon et al., 2012; Nora et al., 2012). Thus, a more plausible explanation is that the *Teratorn* insertion disrupted local TAD organization. Disruption of TAD structures is increasingly recognized as a cause of misregulation in vertebrate Hox clusters. For instance, mesomelic dysplasias associated with the human HOXD locus have been shown to result from regulatory reallocations due to altered TAD boundaries (Bolt et al., 2021). By analogy, the *Teratorn* insertion in *widefins* may similarly perturb local chromatin architecture and reassign enhancer–promoter contacts within the hoxca cluster. Such disruption could interfere with higher-order chromatin organization and alter enhancer–promoter communication, leading to the reduced expression of *hoxc6a*. In this scenario, the enhancer(s) normally required for *hoxc6a* activation may have been functionally insulated or miswired by the insertion.

*widefins* have also been selectively bred and maintained as ornamental medaka with a novel trait, deliberately excluding the dorsal fin deletion and increased vertebra phenotypes. According to the breeder, the dorsal-fin–absent phenotype was more frequently observed when *widefins* were first identified. After several generations of continuous selection for the widefin trait, this phenotype became more pronounced. While this study describes the phenotypic differences observed among *widefins* individuals, variations in the proportion of phenotypes across generations were also noted, although these changes were not systematically recorded. Epigenetic modifications have been shown to persist across generations in fish (Iwanami et al., 2020; Kelley et al., 2021), suggesting that a similar phenomenon may be occurring in *widefins* medaka.

## Data Availability

All data necessary to support the conclusions of this article are provided within the main text, figures, and supplementary data. *widefins* medaka are commercially available through aquarium shops. The frozen sperm of *hoxc12a* and *hoxc12a;c13a* mutant medaka used in this study have been deposited in the NBRP Medaka repository (https://shigen.nig.ac.jp/medaka/) and are available upon request. The induced mutations of *hoxc12a* and *hoxc12a;c13a* mutants, generated by CRISPR-Cas9, were previously described (Adachi et al., 2024). Raw sequence reads from short-read whole-genome sequencing using Illumina and long-read whole-genome sequencing (Oxford Nanopore) have been deposited in DDBJ under BioProject accession number PRJDB20724, with BioSample Accession number SAMD00916351 (Illumina) and SAMD01619948 (Nanopore). The sequence of the *Teratorn* in *widefins* allele has been deposited under accession number LC888077.

## Acknowledgments

We thank Dr. Ken Naito for performing the long-read Nanopore sequencing. We also acknowledge the NBRP medaka for providing fish and preserving the mutant lines used in this study. This work was supported by KAKENHI Grants-in-Aid for Scientific Research from the Ministry of Education, Culture, Sports, Science, and Technology, Japan (23K05790 to A.K.), and by the National Institute of Genetics under the Joint Research and Research Meeting (NIG-JOINT) program (31A2023, 26A2024) to A.K.).

## Author Contributions

A.K. and M.M. conceived and designed the experiments, R.K., S.O., N.F., and H.K. analyzed *widefins* medaka, K.Y. isolated *widefins* medaka, A.M. performed micro-CT scan, A.K. wrote the manuscript. All authors discussed the results and commented on the manuscript.

---

**Supplemental Figure 1. External appearance of the *widefins* medaka used in this study.**

Two *widefins* medaka swimming in a tank. The dorsal and anal fins are enlarged. To produce an ornamental strain, the fish have been crossed with a mutant exhibiting abnormal pigmentation.

**Supplemental Figure 2. Dot plot comparing *Teratorn* sequences in *widefins* and the previously identified *Teratorn*.**

The dot plot shows the similarity between the *Teratorn* sequence (184,498 bp) inserted into *hoxc12a* locus in *widefins* and the previously reported *Teratorn* sequence (181,795 bp; accession number LC199500), generated using YASS web tool (https://bioinfo.lifl.fr/yass/index.phpdotplot). Green dots indicate forward matches, where the two sequences align in the same orientation. Red dots indicate reverse matches, where one sequence is inverted relative to the other.

**Supplemental Figure 3. Comparisons of nucleotide sequences of *Teratorn* sequences identified in *widefins* and the previously identified *Teratorn*.**

The *Teratorn* sequence inserted at the *hoxc12a* locus in *widefins* was compared with the previously reported *Teratorn* sequence using ClustalW. In the previously reported *Teratorn* sequence, the nucleotide regions proposed to encode proteins were highlighted in bold.

**Supplementary Movie 1. Micro-CT scanning of the whole skeletal structure of wild-type medaka.**

Micro-CT scanning was performed on adult wild-type medaka to visualize the entire skeletal structure.

**Supplementary Movie 2. Micro-CT scanning of the whole skeletal structure of *widfeins* medaka.**

Micro-CT scanning was conducted on adult *widefins* medaka to show the entire skeletal structure. The most common phenotype of *widefins* medaka, which resembles that of *hoxc12a* mutants, is presented.

**Supplementary Movie 3. Micro-CT scanning of the anterior vertebrae of wild-type and *widefins* medaka.**

Micro-CT scanning of the anterior vertebrae of wild-type and *widefins* medaka was conducted using the same specimens featured in Supplementary Movies 1 and 2.

