## Supplemental Figure 1 and 2 for "The phenotypic variation of *widefins* medaka is due to the insertion of a giant transposon containing a viral genome within hoxca cluster"

### *widefins*

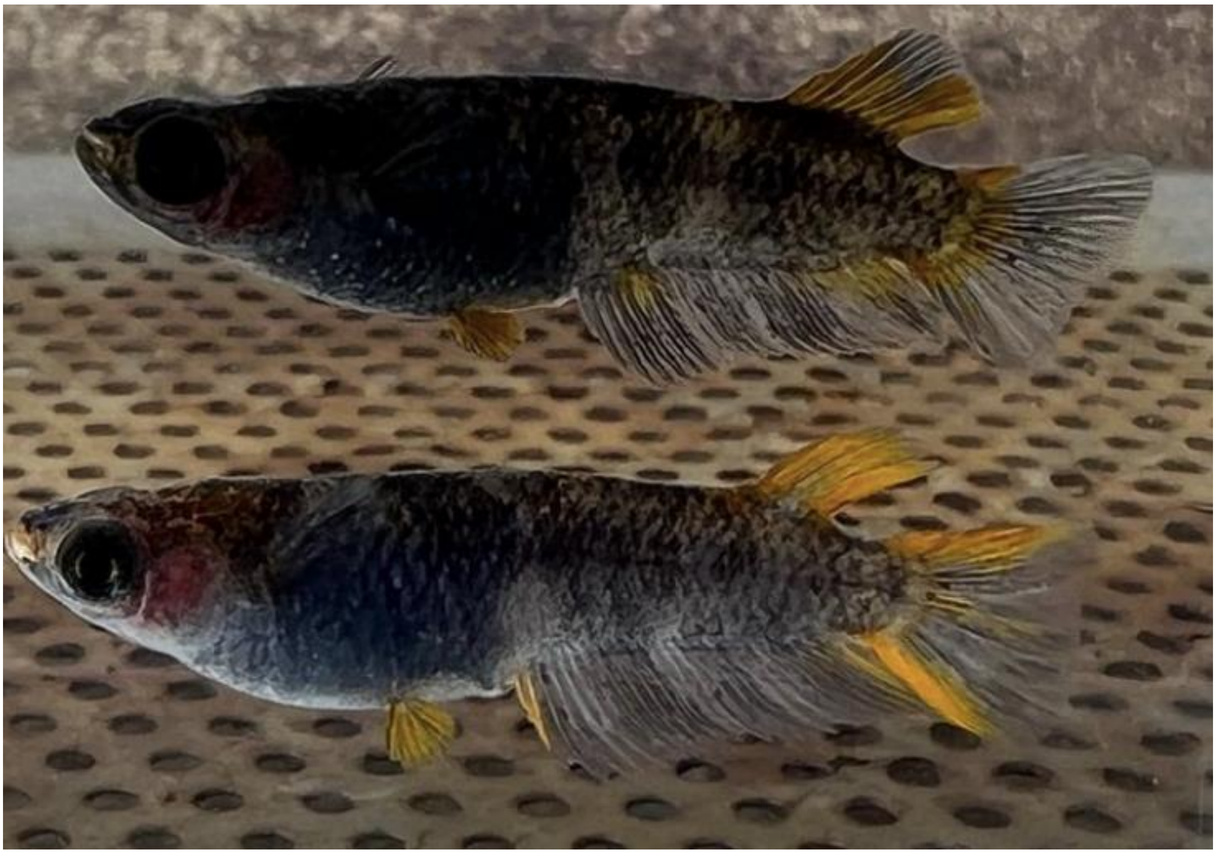

**Supplemental Figure 1. External appearance of the *widefins* medaka used in this study.**

Two *widefins* medaka swimming in a tank. The dorsal and anal fins are enlarged. To produce an ornamental strain, the fish have been crossed with a mutant exhibiting abnormal pigmentation.

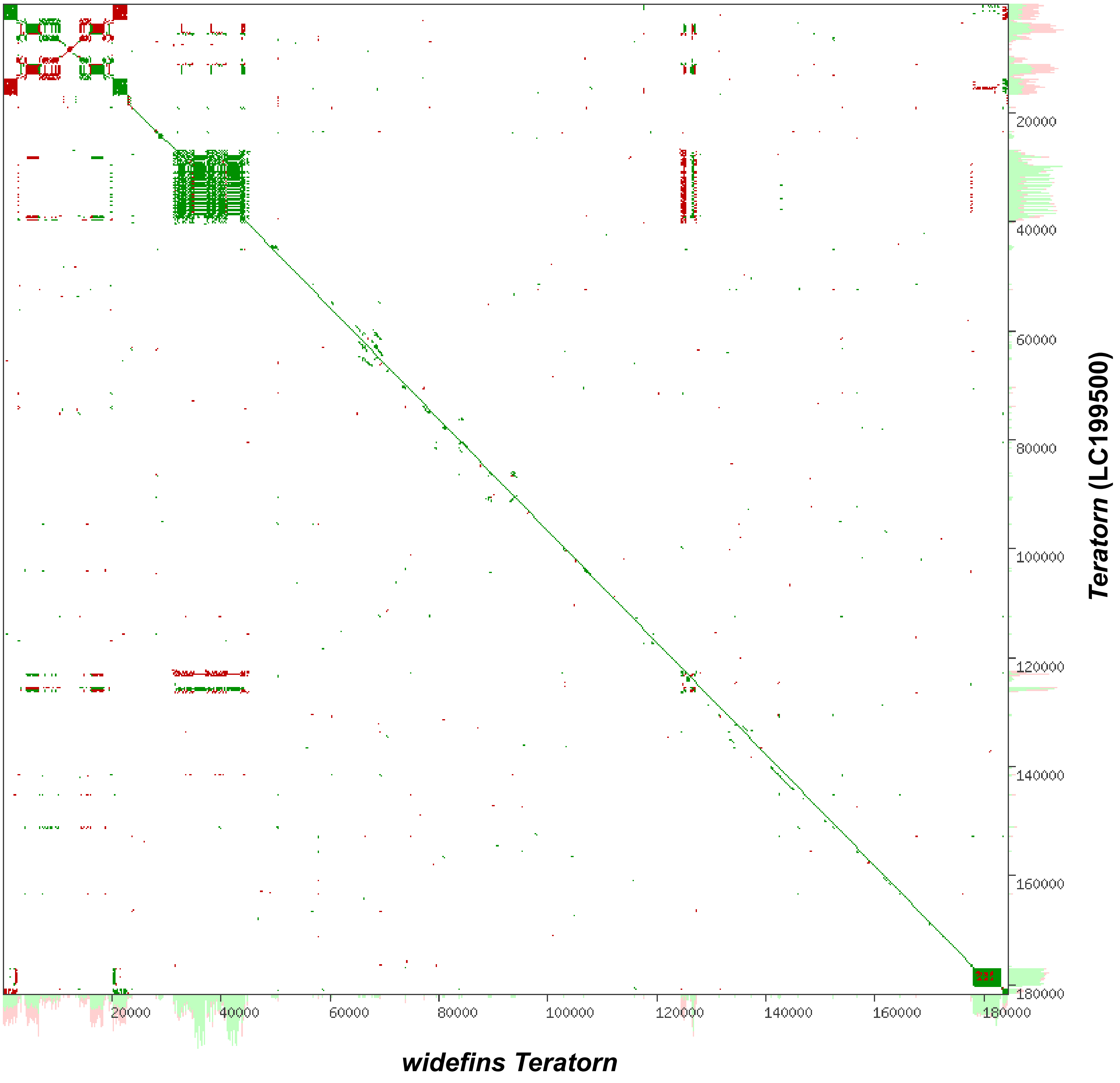


**Supplemental Figure 2. Dot plot comparing *Teratorn* sequences in *widefins* and the previously identified *Teratorn*.**

The dot plot shows the similarity between the *Teratorn* sequence (184,498 bp) inserted into *hoxc12a* locus in *widefins* and the previously reported *Teratorn* sequence (181,795 bp; accession number LC199500), generated using YASS web tool (https://bioinfo.lifl.fr/yass/index.phpdotplot). Green dots indicate forward matches, where the two sequences align in the same orientation. Red dots indicate reverse matches, where one sequence is inverted relative to the other.
