## Supplemental Figure 3 for "The phenotypic variation of *widefins* medaka is due to the insertion of a giant transposon containing a viral genome within hoxca cluster"

Method:  Clustal W

```
TeratornLC199500  CCCTTCCGCTCGCTTAGGTTGGCATGTGCTTTGGGGTCTGCATGACCCCCCTCCAAAAAAATATAACTCCCTCCGCTGTTTTACAGGGCATAAGTGACCCCACGAGGCATTTCTCGGTGG   120
WF-Teratorn       CCCTTCCGCTCGCTTAGGTTGGCATGTGCTTTGGGGTCTGCATGACCCCCCTCCAAAAAAATATAACTCCCTCCGCTGTTTTACAGGGCATAAGTGACCCCACGAGGCATTTCTCGGTGG   120

TeratornLC199500  CCAGCACGAAACCCATTCCCTCCGCTGTTTTACAGGGCATAAGTGACCCCACGAGACAATACTTTATTTTTTTTTCTCTCTCCATCCTCTCCGCTGTTTTACAGGGCATAAGTGACCCCA   240
WF-Teratorn       CCAGCACGAAACCCATTCCCTCCGCTGTTTTACAGGGCATAAGTGACCCCACGAGACAATACTTTATTTTTTTTTCTCTC----------------------------------------   200

TeratornLC199500  CAAGGCATTTCTCGATGGCCAGCACGAAACCCATTCCCTCCGCTGTTTTACAGGGCATAAGTGACCCCACGAGACAATACTTTATTTTTTTT--CTCTCTCCATCCTCTCTTCATTTTCT   358
WF-Teratorn       -------------------------------CATTCCCTCCGCTGTTTTACAGGGCAGAAGTGACCCCACGAGACAATACTTTATTTTTTTTTTTCTCTCTCCATCCTCTCTTCATTTTCC   289

TeratornLC199500  TTACACCCCAAGAC---CAGGGGGTGGAGAGGGAGGGAGGAGGG--------------TATCAATTTATTTTTATACCCCTCGCGCTGTGCTTTGGCGTGCATTCGACCCCGTATTATTA   461
WF-Teratorn       TTACACCCCAAGACCACCAGAGGGTGGAGAGGGAAGGAGGGGGGGTATAATTTTTTTTTTTTAATTTATTTTCATACCCCTCGCGCTGTGCTTTCGCTTGCATTCGACCCCGTATTATTA   409

TeratornLC199500  AAAAAAAAAAAGC-GTTCACTCCGCTGTTTTACGGAGCTCAAATGACCCCAAGAGACAATACTTTATTTTTTTTTTCTCCA-AAAAAAGCGTTCACTCCGCTGTTTTACGGAGTGCAAAT   579
WF-Teratorn       AAAAAAAAAAAACCATTCACTCCGCTGTTTTACGGAGCTCAAATGACCCCAAGAGACAATACTTT----TTTTTTTCTCCAGAAAAAAGCGTTCACTCCGCTGTTTTACGGAGTGCAAAT   525

TeratornLC199500  GACCCCACGAGACAATGCTTTTTTTTTTTCACCCACTATTCATTT-CCTTACAGCGCACGACCACCAGGGGGAGGGGAAGGGGGGG-------CATCAATTCATTTATTTATTTACATAC   691
WF-Teratorn       GACCCCACGAGACAATGCTTTTTTTTT--CATACACTCTTCATTTTCCTTACAGCGCACGACCACCAGGGGGAGGGGAAGGGGGGGGGGGGGGCATCAATTCATTTATTTATTTACATAC   643

TeratornLC199500  CCCTCGCGCTGTGCTTTCGCGTGCATTCGACCCCTGTATTATTTAAAAAAAAAGCGTTCACTCCGCTGTTTTATGGAGCTCAAATGACCCCAAGAGACAATACTTTATTTT--TTTTTCT   809
WF-Teratorn       CCCTCGCGCTGTGCTTTCGCGTGCATTCGACCCCTGTATTATTTAAAAAAAAAGCGTTCACTCCGCTGTTTTACGGAGCTAAAATGACCCCAAGAGACAATACTTTATTTTTATTTTTCT   763

TeratornLC199500  CCAAA--CAAAAGCGTTCACTCCGCTGTTTTACGGGGCTAAAATGACCCCAAGAGACAATACTTTATTTTTATTTTCCTCCAAAAACAAAAGCGTTCACTCCGCTGTTTTACGGAGCTAA   927
WF-Teratorn       CCAAAAACAAAAGCGTTCACTCCGCTGTTTTACGGAGTGCAAATGACCCCAAGAGACAATACAGTATTTTTATTTTTCTCCAAAAACAAAAGCGTTCACTCCGCTGTTTTACGGAGCTAA   883

TeratornLC199500  AATGACCCCAAGAGACAATACTTTATTTTTTTTTTTTCTCCAAAAACAAAAGCGTTCACTCCGCTGTTTTACGGAGCTCAAATGACCCCAAGAGACAATACTTTATTATTTTTTTCTCCA  1047
WF-Teratorn       AATGACCCCAAGAGACTATACAGTATTTTATTTTTTTCTCCAAAAACAAAAGCGTTCAATACGCTGTTTTACGGAGCTCAAATGACCCCAAGAGACAATACTTTATTATTTTTTACTCCA  1003

TeratornLC199500  AAAACAAAAGCGTTCACTCCGCTGTTTTACAGAGTGCAAATGACCCCACGAGACTATGCTTTTTT---TTCCTATCTTCATTTTCCTTACACCGCACGACCACCAGGGGGAAGGGGGGG-  1163
WF-Teratorn       AAAACAAAAGCGTTCACTCCGCTGTTTTACGGAGTGCAAATGACCCCACTAGACTATGCTTTTTTATTTTCCTATTTTCATTTTCCTTACACCGCACGACCACCAGGGGGAAGGGGGGGG  1123

TeratornLC199500  -----------------------------------------------GGGTATA------------ATTTTT------------------------------------------------  1176
WF-Teratorn       TATATTTTTTTTTAAATTTTTTGTCATACCCCTCGCGCTGTAGTTTGGGGTATATTTGACCCCCGTATTTTTAAAAAAAAAAAAAAAAAAATCCCATCCGCTGTTTTTCGGAGCACCTGTG  1243

TeratornLC199500  --------------TTTTTTGTTTATGGCATTTGGCAATTCCCTCGCGCTGTAGTTTGGGGTGCATTTGACCCCCGTATTTTTTTTCAAAAAAAAATAAAAATTCCAACCGCTGTTTATC  1282
WF-Teratorn       ACCCCACGAGGCATTTTTTTGTTTATGGCATTTGGCAATTCCCTCGCGCTGTAGTTTGGGGTGCATTTGACCCCCGTATTTTTAAAAAAAAAAAAA-AAAAATTCCAACCGCTGTTTTTC  1362

TeratornLC199500  GGAGCACGTGTGACCCCACGAGGCGTTTTTTTTTTTTTTTTT--TGGTGGATGGCATTTGGCAATTCCCTCGCGCTGTAGTTTGGGGTGCATTTGACCTCCGTATTTATATAACCCTCCC  1400
WF-Teratorn       GGAGCACGTGTGACCCCACGAGGCGTTTTTTTTTTTTTTTTTTTTGGTGGATGGCATTTGGCAATTCCCTCACGCTGTAGTTTGGGGTGCATTTGACCTCCGTATTTATATAACCCTCCC  1482

TeratornLC199500  ACTCGCTCAGGTGGGCATGTGCTTTGGGGTCTGCATGACCCCCCAAAAAATCCGCTGTTTTACGCAGCTCGTGTGACCCCACGAGAAATTTCTCAGTGGCCGGCATAAAAAACATTCAGT  1520
WF-Teratorn       ACTCGCTCAGGTGGGCATGTGCTTTGGGGTCTGCATGACCCCCCAAAAAATCCGCTGTTTTACGGAGCT-------------------------------------AAAAAACATTCAGT  1565

TeratornLC199500  CCGCTGTTTTACTGATCGTAAACGACCCCACGAGAAATTTCTCTGTGGCCGGCATAAAAAACATTCAGTCCGCTGTTTTACGGATCGTAAACGACCCCACGAGACAATACTTTTTTTATC  1640
WF-Teratorn       CCGCTGTTTTACGGATCGTAAACGACCCCACGAGAAATTTCTCTGTGGCCGGCATAAAAAACATTCAGTCCGCTGTTTTAGGGATCGTAAACGACCCCGCGAGACAATACTTTTTTTATC  1685

TeratornLC199500  ATCCTCTCTTCATTGTCCTTACACCCCACGACCACCAGGGGGTGGAGGGGGAGGAGGGGGGCATCAATTTATTTATTTATTTTCATACCTCTCGCGCTGTGCTTTGGCGTGCATTTGACC  1760
WF-Teratorn       ATCCTCTCTTCATTGTCCTTACACCCCACGACCACCAGGGGGTGGAGGGGGAGGAGGGGGGCATCAATTTATTTATTTATTTTCATACCTCTCGCGCTGTGCTTTGGCGTGCATTTGACC  1805

TeratornLC199500  CCATATTTTTTATCATCCTCTCTTCATTGTCCTTACACCCCACGACCACCAGGGGGTGGAGGGGGAGGAGGCGATGAGGGGGGGCATCAAACTATTTATTTATTTTCATACCCCTCGCGC  1880
WF-Teratorn       CCATATTTTT----TCATCTCTTCATTGTCCTTACACCCCACGACCACCAGGGGGTGGAGGGGGAGGAGG--------------------------------------------------  1872

TeratornLC199500  TGTCTTTTGGGTGCATTTGACCCCGTATTTTTAAAAAAAAAAACATTCACTCCGCTGGTTTACGGAACGTAAAGGACCCCATCCTCTCTTCATTGTACTTACACCCCACGACCACCAGGG  2000
WF-Teratorn       ------------------------------------------------------------------------------------------------------------------------  1872
```

```
TeratornLC199500  GGTGGAGGGGATGAGGGGGGGCATCAATCTATTTATTTATTTTCATACCCCTCGCGCTGTCTTTTGGGTGCATTTGACCCCGTATTTTTTTTTTCTTCATTGTCCTTACACCCCACGACC    2120
WF-Teratorn       --------GGATGAGGGGGGGCATCAATCTATTTATTTATTTTCATACC-----------------------------------------------------------------------    1913

TeratornLC199500  ACCAGGGGGTGGATGGGGGGAGAAGGGGGGCATCAATTTATTTATTTATTTTCATACCTCTCGCGCTGTGCTTTGGCGTGCATTTGACCCCATATTTTTTTCATCTCTTCATTGTCCTTA    2240
WF-Teratorn       ----------------------------------------------------------TCTCGCGCTGTGCTTTGGCGTGCATTTGACCCCATATTTTTTTCATCTCTTCATTGTCCTTA    1975

TeratornLC199500  CACCCCACGACCACCAGGGGGTGGAGGGAGAGGAGGGGATGAGGGGGGGCATCAATCTATTTATTTATTTTTATACCCCTCGCGCTGTCTTTTGGTTGCATTTGACCCCATATTTTTTTT    2360
WF-Teratorn       CACCCCACGACCACCAGGGGGTGGAGGGGGAGGAGGCGATGAGGGGGGGCATCAAACTATTTATTTATTTTCATACCCCTCGCGCTGTCTTTTGGGTGCATTTGACCCCGTATTTTTA--    2093

TeratornLC199500  AAAAAAAAAACATTCACTCCGCTGTTTTACGGAACGTAAAGGACCCCATCCTCTCTTCATTGTACTTACACCCCACGACCACCAGGGGGTGGAGGGGATGAGGGGGGGCATCAATCTATT    2480
WF-Teratorn       AAAAAAAAAACATTCACTCCGCTGGTTTACGGAACGTAAAGGACCCCATCCTCTCTTCATTGTACTTACACCCCATGACCACCAGGGGGTGGAGGGGATGAGGGGGGGCATCAATCTATT    2213

TeratornLC199500  TATTTATTTTCATACCCCTCGCGCTGTCTTTTGGGTGCATTTGACCCCGTATTTGTTTTCTTCATTGTCCTTACACCCCACGACCACCAGGGGGTGGAGGGGGGGAGAAGGGGGGCATCA    2600
WF-Teratorn       TATTTATTTTCATACCCCTCGCGCTGTCTTTTGGGTGCATTTGACCCCGTATTTTTTTTCTTCATTGTCCTTACACCCCACGACCACCAGGGGGTGGAGGGGGGGAGAAGGGGGGCATCA    2333

TeratornLC199500  ATTTATTTATTTATTTTCATACCTCTCGCGCTGTACTTTAGGGTGCATTTGACCCCGTACTTTAAAAAAAAAAAAAAACTTTCACTTTCACAAACTTTCATTTACACAGAGTATAAATGAC    2720
WF-Teratorn       ATTTATTTATTTATTTTCATACCTCTCGCGCTGTACTTTAGGGTGCATTTGACCCCGTACTTTTTAAAAAAAAAAAAACTTTCACTCC-----ACTGT--TTTACA--GAGTATAAATGAC    2444

TeratornLC199500  CCCACGAGACATTACTTGGTGGCCTGCATTTGGCAATTCCCTCGCGCTGTACTTTAGGGTGCATTTGACCCCTGTGTTTTTTTTTTTTTTTATCCATCCACTGTTTTACTGTGAGTGACC    2840
WF-Teratorn       CCCACGAGACATTACTTGGTGGCCGGTATTTGGCAATTCCCTCGCGCTGTACTTTAGGGTGCATTTGACCCCTGTGTTTTTTTTTTA----ATCCATCCACTGTTTTACTGTGAGTGACC    2560

TeratornLC199500  AACCAGACATTCTTTTTTTTTTCTCTCTCTTCATTTCCTTACCCCTCGCACTTGCTTAGGTTGGCGTGTGCTTTAGAAAAATCCCCTCCCCAGACACAGGGACACAGACACAGAAAGCTG    2960
WF-Teratorn       AACGAGACATTCCTTTTTTTTT-TCTCTCTTCATTTCCTTACCCCTCGCACTTGCTTAGGTTGGCGTGTGCTTTAGAAAAATCCCCTCCCCAGACACAGGGACACAGACACAGAAAGCTG    2679

TeratornLC199500  TAGAACGAGCTCAGAGAACACGCGCGAGAGAGATAGAGGGGGGAGGGAGGCATGCGGAGACCTCCGCTCTGTGAGGCCGACCGAGGCACGTTAGAGAAATCCCCTCCGCTGTCAAGCCAA    3080
WF-Teratorn       TCGAAGGAGCTCAGAGAACACGCGCGAGAGAGATAGAGGGGGGAGGGAGGCATGCGGAGGCCTCCGCTCTGTGAGGCCGACCGAGGCACGTTAGAGAAATCCCCTCCGCTGTCAAGCCAA    2799

TeratornLC199500  GAGAACAGCC--------------------------------------------------------------------------------------------------------------    3090
WF-Teratorn       GAGAACAGCCTGCATCGCTGTTTTTTTTTTACATGGCAGATGAAGGGAAACATAAATGACTCTCAACAAACAGAAACTTGGATGAGAGAGAAACAAAGAAAAGAAAAAAGAAAAAGACAA    2919

TeratornLC199500  ------------------------------------------------------------------------------------------------------------------------    3090
WF-Teratorn       CAGAAAACAAAAAACAATAACCAAATCCTGTCTTCGAAGGAGTTCTATTCGCGTAGAGCGAAGAGAAGACAGCCGCTGAAATCTCACATCCAAAGCTCACACCAAATACGTGTGCAGAGC    3039

TeratornLC199500  ------------------------------------------------------------------------------------------------------------------------    3090
WF-Teratorn       TCAGGTTTTGGGGCCAAGACAAAGGGAGCTACGGACACGGCTCATGAAAAGCTGCTCCAGTGGCCAGAAGGGAGGAGGGTGCGTGATGGTGCACTCAGGGGCTGCGGGCTTCTGGGACAG    3159

TeratornLC199500  ------------------------------------------------------------------------------------------------------------------------    3090
WF-Teratorn       CAACAGACTCCACCTTACAGCCGCGAGGCTGTGTAATTGTTGGCTCATTCGTAGAGCTCTGAGGCTGTGTGGCTCTGGTTGCGAAAGCCTTCAGCCTGCCCCCACACAGGGCTCCTGCGA    3279

TeratornLC199500  --------------------------------------AGAGAAACAGCAGGCCAGACGCAGAGGAGGGACCGTGCTGCCTGCAGCTGCGAGGAAAGACCAGAGTTTTTTTTTT---TTG    3169
WF-Teratorn       GACAGAGAACGCACTCTGCACACACTCACGAGCAGGCGAGAGAAACAGCAGGCCAGACGCAGAGGAGGGACCGTGCTGCCTGCAGCTGCGAGCAAAGACCAGAGTTTTTTTTTTTTTTTG    3399

TeratornLC199500  AGGAAGCAAGGGAACTCGGCCGATCAGCAGCTTTCTTCTGTGTTTCAAAGCAGGGAGGTTCTCAGCTCAGGGAGCGGGCAGGGCCCTGCAAGAAAAGCAGCGTCTCGCAGCTATGCCGAG    3289
WF-Teratorn       AGGAAGCAAGGGAACTCGGCCGATCAGCAGCTTTCTTCTGTGTTTCAAAGCAGGGAGGTTCTCAGCTCAGGGAGCGGGCAGGGCCCTGCAAGAAAAGCAGCGTCTCGCAGCTATGCCGAG    3519

TeratornLC199500  CAGGATTTTCAATGTCACTAATCACAAAGAGAAAATAAAAAGGATGCTCGTTGGAGC---------------------------------------------------------------    3346
WF-Teratorn       CAGGATTTTCAATGTCACTAATCACAAAGAGAAAATAAAAAGGATGCTCGTTGGAGCTGGGTTCATTTCAGAGGCACCCGAATGATTTCTTTGACTAGATGTACCGAAACTCTAAAAAAA    3639

TeratornLC199500  ------------------------------------------------------------------------------------------------------------------------    3346
WF-Teratorn       ACAACAAAAAGCATTCATCCTATGACAGAAACGGCGCTAAACAATGATAAAAAGCTGCTCCTCTGAGAAGGGAGAACGGTGCGTGATGGCGCAGTAAGAATCTGCAGGCCTCCTAGTTTC    3759

TeratornLC199500  ---------------------------ATTGCTGGTTTTACACGCCTGAGTGTCTGCAAGTCACGTCCAAAGCCGCTTGGCTTCGGCTCATTCGCAGAGCTC------------------    3421
WF-Teratorn       AGCTGGAGCCAGCTTGGCCATAGGAACATTGCTGGTTTTACACGCCTGAGTGTCTGCAAGTCACGTCCAAAGCCGCTTGGCTTCGGCTCATTCGCAGAGCTCTGAGAGCCTGCCCCCACA    3879

TeratornLC199500  ----------------------------------------------CCTCCCTGAAAACGACGTGTTTGCATTTTTACACACACCAAATACGTGTCCACAGCTCAGCTTTTTGTGGGGCC    3495
WF-Teratorn       GAGGGCCCCTGAGGAAGCAGCGGCAGGGAGGCCAGAGTGGAATTTCCCTCTCTGAAGACGACGTGTTTGCATTTTTACACACACCAAATACGTGTCCACAGCTCAGCTGTTTGTGGGGCC    3999
```

TeratornLC199500 AGATCAGAGATGAGGGCAGTGGGCAGACACAGAAAGCTTTCCGAAGGGGCTCAGAGAACGCTCGCGACAGAGAAACAGAGAGAAAGGGAAGAAGGAGGAGGTGGAGGAGGAGGAGGAGGA 3615
WF-Teratorn AGATCAGAGATGAGGGCAGTGGGCAGACACAGAAAGCTTTCCGAAGGGGCTCAGAGAACGCGCGCGACAGGGAAACAGAGAAACAGAGAGAAAGGGAA----GAAGGAGGAGGAGGAAGA 4115

TeratornLC199500 GAGAAGGAATGTG------------AGGCAGAGACTGGGGAGAGGGGGTTGGAGACAGAGATGATTTGGCCTCGGCTCTTGGAGA**ATGACTGACTCGGACCGCCCAGTGGAGCCTCAC**-- 3721
WF-Teratorn GAGAAGGAGTGTGTGTGACTGAAAGAGGCAGAGACTGGGGAGAGGGGGTTGGAGAAAGAGATGATTTGGCCTCGGCTCTTGGAGAATGACTGACTCGGACCGCCC--CAGAGCCACACAG 4233

TeratornLC199500 **TCACGAGCGGGCGG----------GAGAAACAGCAAG-GCAGACCCAGAG----GAGGAACTGTGGTGCCTGCAGCTGCGAG---CAGGGAGTTTTGAGGAAGGAGGGG--GCAGGGA--** 3819
WF-Teratorn CCTCG-GCGCTCTGTTTGCATTTTTACACACACCAAATACGTGTCCACAGCTCAGGAGAGAAGGAGTGTGTGTGACTGAAAGGGGCAGAGACTGGGGAGAGAGGGGGGGGAGACAGAGATG 4352

TeratornLC199500 **ACTCGGCC---GATC-----AGCAGTTTTCTTCTG-TGTTTTCAAAGCCGCACAGCCTCTGAGCTCTGCGAAGAC---AGGGGCCACGGCTCAGGGGGGAAGAATGCAGCCTTACTCTCA** 3927
WF-Teratorn ATTTGGCCTCGGCTCTTGGAGGCTGACTGACTCGGACCGCCCCAGAGCCACACAGCCTCGGCGCTCTGCTCTTTCTCGAGAGGCCA--GCTCAGGGGGGAAGAATGCAGCCTTACTCTCA 4470

TeratornLC199500 **GACGAGTCACACAGCCTCTGAGCTCTGCGAAGACAGGGGTCAGCCCAGGGGCAGGCCGCAGCCTTTCTTGGTGGGGGAAGAATGCAGCCTGTCGAACCAGAGCCAGCCGCACAGCCTCTG** 4047
WF-Teratorn GACGAGTCACACAGCCTCTGAGCTCTGCGAAGACAGGGGTCAGCCCAGGGGCAGGCCGCAGCCTTTCTTGGTGGGGGAAGAATGCAGCCTGTCGAACCAGAGCCAACCGCACAGCCTCTG 4590

TeratornLC199500 **A**GCTCTGCGAAGACAGGGGCCACGGCTCAGGGGGGAAGAATGCAGCCTTACTCTCAGACGAGTCACACAGCCTCTGAGCTCTGCGAAGACAGGGGTCAGCCCAGGGGCAGGCCGCAGCCT 4167
WF-Teratorn AGCTCTGCGAAGACAGGGGCCACGGCTCAGGGGGGAAGAATGCAGCCTTACTCTCAGACGAGTCACACAGCCTCTGAGCTCTGCGAAGACAGGGGTCAGCCCAGGGGCAGGCCGCAGCCT 4710

TeratornLC199500 TTCTTGGTGGGGGAAGAATGCAGCCTGTCGAAGCAGAGCCAGCCGCACAGCCTCTGAGCTCTGCGAAGACAGGGGCCACTGCTCAGAGGGGGGAAGAATGCAGCCTTACTCTCAGACGAGT 4287
WF-Teratorn TTCTTGGTGGGGGAAGAATGCAGCCTGTCGAACCAGAGCCAGCCGCACAGCCTCTGAGCTCTGCGAAGACAGGGGCCACGGCTCAGAGGGGGGAAGAATGCAGCCTTACTCTCAGACGAGT 4830

TeratornLC199500 CACACAGCCTCTGAGCTCTGCGAAGACAGGAGTCAGCCCAGGGGCAGGCCGCAGCCTTTCTTGGTGGGGGAAGAATGCAGCCTGTCGAAGTTGAGCCAGCCGCACAGCCTCTGAGCTCTG 4407
WF-Teratorn CACACAGCCTCTGAGCTCTGCGAAGACAGGAGTCAGCCCAGGGGCAGGCCGCAGCCTTTCTTGGTGGGGGAAGAATGCAGCCTGTCGAACCAGAGCCAGCCGCACAGCCTCTGAGCTCTG 4950

TeratornLC199500 CGAAGACAGGGGCCACGGCTCAGAGGGGGAAGAATGCAGCCTTACTCTCAGACGAGTCACACAGCCTCTGAGCTCTGCGAAGACAGGAGTCAGCCCAGGGGCAGGCCGCAGCCTTTCTTG 4527
WF-Teratorn CGAAGACAGGGGCCACGGCTCAGAGGGGGAAGAATGCAGCCTTACTCTCAGACGAGTCACACAGCCTCTGAGCTCTGCGAAGACAGGAGTCAGCCCAGGGGCAGGCCGCAGCCTTTCTTG 5070

TeratornLC199500 GTGGGGGAAGAATGCAGCCTGTCGAACCA----CAGCCGCACAGCCTCTGAGCTCTGCGAAGACAGGAGTCAGCCCAGGGGCAGGCCGCAGCCTTTCTTGGTGGGGGAAGAAAGCAGCCT 4643
WF-Teratorn GTGGGGGAAGAATGCAGCCTGTCGAAGCAGAGCCAGCCGCACAGCCTCTGAGCTCTGCGAAGACAGGGGTCAGCCCAGGGGCAGGCCGCAGCCTTTCTTGGTGGGGGAAGAATGCAGCCT 5190

TeratornLC199500 GTCGAAGCAGAGCCAGCCGCACAGCCTCTGAGCTCTGCGAAGACAGGGG----------------------------------------------------------------------- 4692
WF-Teratorn GTTGAAGCAGAGCCAGCCGCACAGCCTCTGAGCTCTGCGAAGACAGGGGTCAGCCCAGGGGCAGGCCGCAGCCTTTCTTGGTGGGGGAAGAATGCAATCTGTCGAACCAGAGCCAGCCGC 5310

TeratornLC199500 -----------------------------CCACGGCTCAGGGGGGAAGAATGCAGCCTTACTCTCTGACGAGTCACACAGCCTCTGAGCTCTGCGAAGACAGGGGTCAGCCCAGGGGCAG 4783
WF-Teratorn ACAGCCTCTGAGCTCTGCGAAGACAGGGGCCACGGCTCAGGGGAAAAAAATGCAGCCTTAATCTCAGACGAGTCACATAGCCTCTGAGCTCTGCGAAGACAGGGGTCAGCCCAGGGGCAG 5430

TeratornLC199500 GCCGCAGCCTTTCTTGGTGGGGGAAGAATGCAGCCTGTCGAAGCAGAGCCAGCCGCAC------------------------------------------------------------- 4841
WF-Teratorn GCCGCAGCCTTTCTTGGTGGGGGAAGAATGCAATCTGTCGAACCAGAGCCAGCCGCACAGCCTCTGAGCTCTGCGAAGACAGGGGTCAGCCCAGGGGCAGGCCGCAGCCTTTCTTGGTGG 5550

TeratornLC199500 ----------------------------------AGCCTCTGAGCTCTGCGAAGACAGGGGCCACGGCTCAGAGGGGGGAAGAATGCAGCCTTACTCTCAGACGAGTCACACAGCCTCTGA 4927
WF-Teratorn GGGAAGAATGCAGCCTGTCGAAGCAGAGCCGCACAGCCTCTGAGCTCTGCGAAGACAGGGGCCACGGCTCAGAGGGGGGAAGAATGCAGCCTTACTCTCAGACGAGTCACACAGCCTCTGA 5670

TeratornLC199500 GCTCTGCGAAGACAGGAGTCAGCCCAGGGGCAGGCCGCAGCCTTTCTTGGTGGGGGAAGAATGCAGCCTGTCGAACCAGAGCCAGCCGCACAGCCTCTGAGCTCTGCGAAGACAGGGGCC 5047
WF-Teratorn GCTCTGCGAAGACAGGGGTCAGCCCAGGGGCAGGCCGCAGCCTTTCTTGGTGGGGGAAGAATGCAGCCTGTCGAACCAGAGCCAACCGCACAGCCTCTGAGCTCTGCGAAGACAGGGGCC 5790

TeratornLC199500 ACGGCTCAGGGGGGAAGAATGCAGCCTTACTCTCAGACGAGTCACACAGCCTCTGAGCTCTGCGAAGACAGGGGTCAGCCCAGGGGCAGGCCGCAGCCTTTCTTGGTGGGGGAAGAATGC 5167
WF-Teratorn ACGGCTCAGGGGGGAAGAATGCAGCCTTACTCTCAGACGAGTCACACAGCCTCTGAGCTCTGCGAAGACAGGGGTCAGCCCAGGGGCAGGCCGCAGCCTTTCTTGGTGGGGGAAGAATGC 5910

TeratornLC199500 AGCCTGTCGAAGCAGAGCCAGCCGCACAGCCTCTGAGCTCTGCGAAGACAGGGGCCACTGCTCAGAGGGGGGAAGAATGCAGCCTTACTCTCAGACGAGTCACAGCAC-----GGCTCAG- 5281
WF-Teratorn AGCCTGTCGAACCAGAGCCAGCCGCACAGCCTCTGAGCTCTGCGAAGACAGGGGCCACGGCTCAGAGGGGGGAAGAATGCAGCCTTACTCTCAGACGAGTCACACAGCCTCTGAGCTCTGC 6030

TeratornLC199500 --AGGCAGGAGGCAGGCTGGAGGCGG----CAGCCTTTC-------------------GCC------ATCAGA----GACGCACAGCCTCTGAGCTCTGCGAAGACAGGGGTCAGCCCAG 5366
WF-Teratorn GAAGACAGGAGTCAGCCCAGGGGCAGGCCGCAGCCTTTCTTGGTGGGGGAAGAATGCAGCCTGTCGAAGCAGAGCCAGCCGCACAGCCTCTGAGCTCTGCGAAGACAGGGGTCAGCCCAG 6150

TeratornLC199500 GGGCAGGCCGCAGCCTTTC----------------GAAACCT---------GAGCCGCACAGCCTCTGAGCTCTGCGAA----------------------------------------- 5420
WF-Teratorn GGGCAGGCCGCAGCCTTTCTTGGTGGGGGAAGAATGCAGCCTGTTGAAGCAGAGCCGCACAGCCTCTGAGCTCTGCGAAGACAGGGGCCACGGCTCAGGGGGGAAGAATGCAGCCTTACT 6270

```
TeratornLC199500 ------------------------------------------------------------------------------------------------TGAGCCAAAGCCA-----------   5433
WF-Teratorn      CTCAGACGAGTCACACAGCCTCTGAGCTCTGCGAAGACAGGGGTCAGCCCAGGGGCAGGCCGCAGCCTTTCTTGGTGGGGGAAGAATGCAGCCTGTCGAACCAGAGCCAGCCGCACAGCC   6390

TeratornLC199500 ----AGCTCTTC------------------------------------------CTCTCAGAAGAGTCACAGAG----------------------------------------GCCACA   5467
WF-Teratorn      TCTGAGCTCTGCGAAGACAGGGGCCACGGCTCAGGGGGGAAGAATGCAGCCTTACTCTCAGACGAGTCACACAGCCTCTGAGCTCTGCGAAGACAGGGGTCAGCCCAGGGGCAGGCCGCA   6510

TeratornLC199500 ------------------AGA----------------CAGAGCCG---------TGGCCTATGTCTTCGCAGAGCTC-----------------CTTGGGA--AAGG-----TTTTTCTC   5520
WF-Teratorn      GCCTTTCTTGGTGGGGGAAGAATGCAGCCTGTCGAACCAGAGCCGCACAGCCTCTGACCCCTGTCTTCGCAGAGCTCAGAGGCTGTGTGACTCGTCTGAGAGTAAGGCTGCATTCTTCCC   6630

TeratornLC199500 T----AGCTGTA----------TAACAGAA----ATCAAAAATTAAAAAAAGTG-------ACAGAAAA---TCCAAAGCAGCACAGAAATTTCTT------------------------   5588
WF-Teratorn      CCCTGAGCCGTGGCCCCTGTCTTCGCAGAGCCAGATCAGAGAT-GAGGGCAGTGGGCAGACACAGAAAGCTTTCCGAAGGGGCTCAGAGAACGCGCGCGACAGAGAAACAGAGAGAAAGG   6749

TeratornLC199500 ---------------------------TGGTGCGCGATGACT------------TT-----------TTATTTATTTT----TTTT------TCATT-------ACAACCCAG-GGA---   5637
WF-Teratorn      GAAGAAGGAGGAGGAGGAGAGAAGGAATGTGAGGCAGAGACTGGGGAGAGGGGGTTGGAGACAGAGATAATTTGGCTTCGGCTCTTGGAGGCTGACTGACTCGGACCGCACAGTGGAGCC   6869

TeratornLC199500 TAGCTCA---GCGAAATGGAGAAAGAGT----------TGGAGCAGGATATTCAATGTCACTAATCACAAAGAGAAAAT----AAAAAGGATGCTCGTTGGAGCATTGCTGGT-------   5733
WF-Teratorn      TCGCTCACGAGCGGGCGGGAGAAACAGCAAGGCAGACCCAGAGGAGGAACCGTGGTGCCTGCAGCTGCGAGCAGGGAGTTTTGAGGAAGGAGGGGCAG-GGAACTCTGCCGATCAGCAGT   6988

TeratornLC199500 ---TTTACACGCCT---------GAGTGTCT--------GCAA------------GTCACGTCCAAAGCCGCTTGGCTTCGGCTCATTCGCAGAGCTCTGAGAGCCTGCCCCCACAGAGG   5821
WF-Teratorn      TTTCTTCTGTGTTTCAAAGCAGGGAGTTTCTCAGCTCAGGCAAAGACCAGAGTGTGTCACGTCCAAAGCCGCTTGGCTTCGGCTCATTCGCAGAGCTCTGAGAGCCTGCCCCCACAGAGG   7108

TeratornLC199500 GCCCCTGAGGAAGCAGCGGCAGGCAGGCCAGAGTGGAATTTTCCTCTCTGAAGACGACGTGTTTGCATTTTTACACACACCAAATACGTGTCCACAGCTCAGCTTTTTGTGGGGCCAGAT   5941
WF-Teratorn      GCCCCTGAGGAAGCAGCGGCAGGGAGGCCAGAGTGGAATTTCCCTCTCTGAAGACGACGTGTTTGCATTTTTACACACACCAAATACGTGTCCACAGCTCAGCTTTTTGTGGGGCCAGAT   7228

TeratornLC199500 CAGGGATGAGGGCAGCGGGCAGACACAGAAAGCTTTCCGAAGGTTCTCAGAGAACGCGCGCGACAGAGAAACAGAG-----------------------------------AGAAAAGGA   6026
WF-Teratorn      CAGAGATGAGGGCAGTGGGCAGACACAGAAAGCTTTCCGAAGGGGCTCAGAGAACGCGCGCGACAGAGAAACAGAGAAACAAAGAGAAAGGGAAGAAGGAGGAGGAGGAGGAGAAAAGGA   7348

TeratornLC199500 G-----------------------------------------------------------------------------------------------------------------------   6027
WF-Teratorn      GTGTGTGTGACTGAAAGAGGCAGAGACTGGGGAGAGGGGGTTGGAGACAGAGATTATTTGGCCTCGGCTCTTGGAGAATGACTGACTCGGACCGCCCCAGAGCCACACAGCCTCGGCGCT   7468

TeratornLC199500 ------------------------------------------------------------------------------------------------------------------------   6027
WF-Teratorn      CTGTTTGCATTTTTACACACACCAAATACGTGTCCACAGCTCAGCTTTTTGTGGGGCCAGATCAGGGATGAGGGCAGTGGGCAGACACAGAAAGCTTTCCGAAGGGGCTCAGAGAACGCG   7588

TeratornLC199500 ------------------------------------------------------------------------------------------------------------------------   6027
WF-Teratorn      CGCGACAGAGAAACAGAGAGAAATGGAAGAAGGAGGAGGAGGACAGAAGGAATGTGTGTGACTGCAAGAGGCACAGACTGGGGAGAGAGGGGGGAGAGAGACAGAGATGATTTGGCCTAG   7708

TeratornLC199500 ------------------------------------------------------------------------------------------------------------------------   6027
WF-Teratorn      GCTCTTGGAGGCTGACTGACTTGTGGGGCCAGATCAGGGCTCGAGAGGCCAGCTCAGGGGGGAAAAAATGTAGCCTTACTCTCAGACGAGTCACACCACAGCTCAGCTTTTTGTGGGGCC   7828

TeratornLC199500 ------------------------------------------------------------------------------------------------AGAGGAGGAGGAGGAGAGAAGGAA   6051
WF-Teratorn      AGATCAGGGATGAGGGCAGCGGGCAGACACAGAAAGCTTTCCGAAGGGGCTCAGAGAACGCGCGCGACAGAGAAACAGAGAGAAATGGAAGAAGCAGGAGGAGGAGGAGGACAGAAGGAA   7948

TeratornLC199500 TGTGTGTGACTGCAAGAGGCACAGACTGGGGAGAGAGGGGGGAGAGAGACAGAGATGATTTTGCCTAGGCTCTTGGAGGCTGACTGACTCGGACCGCCCCAGAGCCACACAGCCTCGTCG   6171
WF-Teratorn      TGTGTGTGACTGCAAGAGGCACAGACTGGGGAGAGAGGGGGGAGAGAGACAGAAATGATTTGGCCTAGGCTCTTGGAGGCTGACTGACTCGGACCGCCCCAGAGCCACACAGCCTCGGCG   8068

TeratornLC199500 CTCTGTTTGCATTTTTACACACACCAAATACGTGTCCACAGCTCAGCTTTTTGTGGGGCCAGATTAGGGATGACGGCAGCGGGCAGACACAGAAAGCTTTCCGAAGGGGCTCAGAGAACG   6291
WF-Teratorn      CTCTGTTTGCATTTTTACACACACCAAATACGTGTCCACAGCTCAGCTTTTTGTGGGGCCAGATCAGGGATGAGGGCAGTGGGCAGACACAGAAAGCTTTCCGAAGGGGCTCAGAGAACG   8188

TeratornLC199500 CGCGCGACAGAGAAACAGAGAGAAATGGAAGAAGGAGGAGGAGGAGAGAAGGAATGTGTGTGACTGAAAGAGGCACAGACTGGGGAGAGAGGGGGGAGAGAGACAGAGATGATTTTGCCT   6411
WF-Teratorn      CGCGCGACAGAGAAACAGAGAGAAATGGAAGAAGGAGGAGGAGGAGAGAAGGAATGTGTGTGACTGCAAGAGGCACAGACTGGGGAGAGAGGGGGGAGAGAGACAGAAATGATTTGGCCT   8308

TeratornLC199500 AGGCTCTTGGAGGCTGACTGACTCGGACCGCCCCAGAGCCACACAGCCTCGTCGCTCTGTTTGCATTTTTACACACACCAAATACGTGTCCACAGCTCAGCTTTTTGTGGGGCCAGATCA   6531
WF-Teratorn      AGGCTCTTGGAGGCTGACTGACTCGGACCGCCCCAGAGCCACACAGCCTCGGCGCTCTGTTTGCATTTTTACACACACCAAATACGTGTCCACAGCTCAGCTTTTTGTGGGGCCAGATCA   8428

TeratornLC199500 GGGATGAGGGCAGTGGGCAGACACAGAAAGCTTTCCGAAGGGGCTCAGAGAACGCGCGCGACAGAGAAACAGAGAGAAATGGAAGAAGGAGGAGGAGGA---------------------   6630
WF-Teratorn      GGGTTGAGGGCAGTGGGCAGACACAGAAAGCTTTCCGAAGGGGCTCAGAGAACGCGCGCGACAGAGAAACAGAGAGAAATGGAAGAAGGAGGAGGAGGACAGAAGGAATGTGTGTGACTG   8548
```

```
TeratornLC199500 ------------------------------------------------------------------------------------------------------------------------   6630
WF-Teratorn      CAAGAGGCACAGACTGGGGAGAGAGGGGGGAGAGAGACAGAGATGATTTGGCCTAGGCTCTTGGAGGCTGACTGACTTGTGGGGCCAGATCAGGGATGAGGGCAGTGGGCAGACACAGAA   8668

TeratornLC199500 ------------------------------------------------------------------------------------------------------------------------   6630
WF-Teratorn      AGCTTTCCGAAGGGGCTCAGAGAACGCGCGCGACAGAGAAACAGAGAGAAATGGAAGAAGGAGGAGGAGGAGAGAAGGAATGTGTGTGACTGCAAGAGGCACAGACTGGGGAGAGAGGGG   8788

TeratornLC199500 ------------------------------------------------------------------------------------------------------------------------   6630
WF-Teratorn      GGAGAGAGACAGAGATGATTTGGCCTAGGCTCTTGGAGGCTGACTGACTTGTGGGGCCAGATCAGGGCTCGAGAGGCCAGCTCAGGGGGGAAAAAATGTAGCCTTACTCTCAGACGAGTC   8908

TeratornLC199500 ------------------------------------------------------------------------------------------------------------------------   6630
WF-Teratorn      ACACCACAGCTCAGCTTTTTGTGGGGCCAGATCAGGGATGAGGGCAGCGGGCAGACACAGAAAGCTTTCCGAAGGGGCTCAGAGAACGCGCGCGACAGAGAAACAGAGAGAAATGGAAGA   9028

TeratornLC199500 -------------GGAGGAGAGAAGGAATGTGTGTGACTGCAAGAGGCAAAGACTGGGGAGAGAGGGGGGAGAAAGACAGAAATGATTTGGCCTCGGCTCTTGGAGGCTGACTGACTCGG   6737
WF-Teratorn      AGCAGGAGGAGGAGGAGGACAGAAGGAATGTGTGTGACTGCAAGAGGCACAGACTGGGGAGAGAGGGGGGAGAGAGACAGAAATGATTTGGCCTAGGCTCTTGGAGGCTGACTGACTCGG   9148

TeratornLC199500 ACCGCCCC------------------------------------------------------------------------GTGGAGCC---TCA--------------------------   6756
WF-Teratorn      ACCGCCCCAGAGCCACACAGCCTCGGCGCTCTGTTTGCATTTTTACACACACCAAATACGTGTCCACAGCTCAGCTTTTTGTGGGGCCAGATCAGGGATGAGGGCAGTGGGCAGACACAG   9268

TeratornLC199500 -----------------CTCAC--------------------------------------------------------------------------------------------------   6761
WF-Teratorn      AAAGCTTTCCGAAGGGGCTCAGAGAACGCGCGCGACAGAGAAACAGAGAGAAAAGGAGAGAGGAAGAGGAGGAGGAGGAGAGAAGGAATGTGTGTGACTGCAAGAGGCACAGACTGGGGA   9388

TeratornLC199500 --GAGCGGGCGGGAGACACAG---------------------------------------------------------------------------------------------------   6780
WF-Teratorn      GAGAGGGGGAGAGAGACACAGATGATTTGGCCTAGGCTCTTGGAGGCTGACTGACTTGTGGGGCCAGATCAGGGCTCGAGAGGCAGTGGGCAGACACAGAAAGCTTTCCGAAGGGGCTCA   9508

TeratornLC199500 -------------------------------------------------------------------------CAAG-------------------------------------------   6784
WF-Teratorn      GAGAACGCGCGCGACAGAGAAACAGAGAGAAATGGAAGAAGGAGGAGGAGGACAGAAGGAATGTGTGTGACTGCAAGAGGCACAGACTGGGGAGAGAGGGGGGAGAGAGACAGAGATGAT   9628

TeratornLC199500 ------------------------------------------------------------------------------------------------------------------------   6784
WF-Teratorn      TTGGCCTAGGCTCTTGGAGGCTGACTGACTTGTGGGGCCAGATCAGGGCTCGAGAGGCCAGCTCAGGGGGGAAAAAATGTAGCCTTACTCTCAGACGAGTCACACCACAGCTCAGCTTTT   9748

TeratornLC199500 -------------------------------GCAGACGCAGA---------------------------------------------------------------------GGAGGA---   6801
WF-Teratorn      TGTGGGGCCAGATCAGGGATGAGGGCAGTGGGCAGACACAGAAAGCTTTCCGAAGGGGCTCAGAGAACGCGCGCGACAGAGAAACAGAGAGAAATGGAAGAAGCAGGAGGAGGAGGAGGA   9868

TeratornLC199500 -----------------------------------------------------------------------------------------------------ACCG---------------   6805
WF-Teratorn      CAGAAGGAATGTGTGTGACTGCAAGAGGCACAGACTGGGGAGAGAGGGGGGAGAGAGACAGAAATGATTTGGCCTAGGCTCTTGGAGGCTGACTGACTCGGACCGCCCCAGAGCCACACA   9988

TeratornLC199500 ------------TG--------------------------GTGCCTGCAGCT-----------------------------------GCGAGCAGG------GAGTTTT-----------   6835
WF-Teratorn      GCCTCGGCGCTCTGTTTGCATTTTTACACACACCAAATACGTGTCCACAGCTCAGCTTTTTGTAGGGCCAGATCAGGGATGACGGCAGCGGGCAGACACAGAAAGCTTTCCGAAGGGGCT  10108

TeratornLC199500 ------------------------------------GAGGAAGGAGG---------------------------------GGCA------------------------------------   6850
WF-Teratorn      CAGAGAACGAGCGCGACAGAGAAACACAGAGAAAAGGAGAGAGGAGGAGGAGGAGAGAAGGAATGTGTGTGACTGAAAGAGGCACAGACTGGGGAGAGAGGGGGGAGAGAGACAGAGATG  10228

TeratornLC199500 ------------------------------------------------------------------------------------------------------------------------   6850
WF-Teratorn      ATTTGGCCTAGGCTCTTGGAGGCTGACTGACTCGGACCGCCCCAGAGCCACACAGCCTCGGCGCTCTGCTCTTACTCGAGAGGCCACGGCTCAGGGGGGGGGAAGAATGCAAAGCAGGGA  10348

TeratornLC199500 ---------------------------------------------------GGGAACTCTGCCGATCAGCAGTTTTCTTCTGTGTTTCAAAGCAGGGAGTTTCTCAGCTCAGGCAAAGAC   6919
WF-Teratorn      GTTTCTCAGCTCAGGCAAAGACCAGAGTGTTTTTTTTTTGTGAGGAAGCAAGGGAACTCGGCCGATCAGCAGCTTTCTTCTGTGTTTCAAAGCAGGGAGTTTCTCAGCTCAGGCAAAGAC  10468

TeratornLC199500 CAGAGTGTTACTTGTTTGTTTATATCTTTCTTTATTTTCCTTTACATATGTAAAATACATTTAA------AAAAAAAATACAGATTGCGAGAGGAGCAGGAGGGATAGACCAGCATTT--   7031
WF-Teratorn      CAGAGTGTTACTTGTTTGTTTATATCTTTCTTTATTTTCCTTTACATATGTAAAATACATTTAAAAAAATAAAAAAAAATACAGATTGCGAGAGGAGCAGGAGGGATAGACCAGCATTTGA  10588

TeratornLC199500 ------------------------------------------------------------------------------------------------------------------------   7031
WF-Teratorn      TGTATGTATGTATGAATGTATGTGTGTGTGTGTGTGTGTGTGTGTGTGTGTGTGTGTTGAAGGACGATAATCATTTTATTCCCATTTAGCTTATAATCTTGTTTATATTTACGCTGAACA  10708

TeratornLC199500 ------------------------------------------------------------------------------------------------------------------------   7031
WF-Teratorn      CTTTTATTGAAACTGCTGAAACTTTTATATAACTTTTGCATTATCTTAAACTAGATGTGGATTTTACTTAAATTTGTATGCTCATATTAGATCTGTTTTTATTAGTCACGTAGTGTATCG  10828
```

```
TeratornLC199500 ------------------------------------------------------------------------------------------------------------------------ 7031
WF-Teratorn      CTGAGTTTAATGTAGCATAACCCAAATGTGCACAATGAGAGCACGCAGGATGGAGCTGACAGGCTTCGATGAGCCAACTAGCGGAGACGGTGGCATGCTTCGGCTGAAACCTGGCCATCG 10948

TeratornLC199500 --------CAACAGTTTGATTTCTGCCTGAGGCCTCCAGCACAAGCCTTGTCTACAGAATCAATGTCTTTTCTCTTTCTTTTATCGCGAAGCCCTTTTCGCACCTTAAAAGAGTTGTAAA 7143
WF-Teratorn      GAGCATTTCAACAGTTTGATTTCTGCCTGAGGCCTCCAGCACAAGCCTTGTCTACAGAATCAATGTCTTTTCTCTTTCTTTTATCGCGAAGCCCTTTTCGCACCTTAAAAGAGTTGTAAA 11068

TeratornLC199500 GTGCTATATGTATGAACAAAGTGTCATTCGTTCAGGTTCCCTTCAATGAAATAACAATATTTATTAAAATATAACTGCCTCATCCAGGCCTGTGTTTGTTGAGAGTAATTTCTGTTTCCC 7263
WF-Teratorn      GTGCTATATGTATGAACAAAGTGTCATTCGTTCAGGTTCCCTTCAATGAAATAACAATATTTATTAAAATATAACTGCCTCATCCAGGCCTGTGTTTGTTGAGAGTAATTTATGTTTCCC 11188

TeratornLC199500 TTCATCTGCCGTATGAAAAACAGCGATGCAGGACAGCCTCCGGGCTACATGCTGAATTTGACCAACATGATTCCAGCTGGAGCTCCAGACTCTCCTCCTCCTCCAACATACTCCAGTTAG 7383
WF-Teratorn      TTCATCTGCCGTATGAAAAACAGCGATGCAGGACAGCCTCCGGGCTACATGCTGAATTTGACCAACATGATTCCAGCTGGAGCTCCAGACTCTCCTCCTCCTCCAACATACTCCAGTTAT 11308

TeratornLC199500 GTTTTTCTAGTAACCGTAACGGCGATGCTGCTCTTGGTTATTTTCATCCTCACCTTGTACCCCCTTTACGGTCCAAAATGCAACACAGAACCAACGATAATGTAAAGATTTGTCTGAAAT 7503
WF-Teratorn      GTTTTTCTAGTAACCGTAACGGCGATGCTGCTCTTGGTTATTTTCATCCTCACCTTGTACCCCCTTTACGGTCCAAAATGCAACACAGAACCAACGATAATGTAAAGATTTGTCTGAAAT 11428

TeratornLC199500 GTTCACTGATTTGTGCGTGCGTGTGGAGTGAGGAATTTGGAAAACAAATCTTCAGGCTTTGAAGCGATGGTCAGGGAACCAAGCGTCAGAGCTATTAGGCGTATTTTCTCCCCGGACACG 7623
WF-Teratorn      GTTCACTGATTTGTGCGTGCGTGTTGAGTGAGGAATTTGGAAAACAAATCTTCAGGCTTTGAAGCGA-----------------GCCGGACTTTCCTGGAGAGTTCTGTTGGTGGA-GCA 11530

TeratornLC199500 AAAC---GCTCCCGCTAGACACTGCATCCTACCACCACAGTTCGAGGGGCCAACGCGGAGAACGAGGCAGGCTGAGGCGGGCTGCGGCACGCAGTGCTTAACAG--CAGCCGGACTTTCC 7738
WF-Teratorn      ACACCAAGGTTCTGTCTGTC-CT-CATGCTGTTTTTA----TTAAG-----AACACA-AGAAGAAGGTAAA--------AATTTCAAAACACAGAA---AAAAGTTCATTGTGGGTGTCG 11627

TeratornLC199500 TGGAGAGTTCTGTTGGTGGAGCATCCCCCTCTGACTTACAGTCATAACGCCCGCTCGCACACACCCAAA---AGCGTG----CACACCCCCTGCAGCTGGTCACCGATTGGTCACCGATT 7851
WF-Teratorn      TATAATGACCCATT--TGGTGCAT-----TCTAACTT-CAACCA--------GTCCTTAT-CAGCTAAGGACAGTGTGTCGTCAAACCCCCT-CAGCCGTGCTGGGATGAGAAGCAGATG 11729

TeratornLC199500 GACCCTCA----CCCCTAAAAGCCCAGATATAAAAAACG--TCGAGTTGACAGT--AGCCTA---------GTCAAAAAGAAAA-TAAAACGTCTACTTAT---TAACGCCGGCAGAGAG 7950
WF-Teratorn      AACACACAAATCTTTCTAAAA----AGGTAAAAAGGTCAATCTGAGTTAACATTTAAGCATACACATTTAAACCAATGATAATACTAAAACAATTTCCTTTACATAAT-CCAACACA--- 11841

TeratornLC199500 CACAGTTCAAGGACACCTTAGAAAAGCTTAGGCGG-----TGCACTGCGTTCGACC----CCCACCAGAGGCCCACCCAGA---------TGGA-------------CATCCTCCTCAA- 8038
WF-Teratorn      CACACTTATCTTGCATGTCAGAACGAATGGGGGGAATGGCTGTAAAGAGTCCAGCCTGGTCTCATTAAAAAAATAAAATAAAAAAAAATAATGAAAAATTAAGCAAGGCATCGTTCTGAGC 11961

TeratornLC199500 ----GGCCATA-CGCTAC-GTGGAGG---CGCAGCAGCTGCAG-CAGGGGTTCAAGTCCTCTCTGTCATCTCAGCCATCA---CGCGAGGAAG--GAG--GGACG--CGGGCTCAGACTG 8139
WF-Teratorn      TGACGGTCCTAGTACTACCGAGAAGGTCTCTCTGCAGTTTTAGTCAGGCGC---AGTAAGAGCGACCGTCTGAGAAAGCGGGCTGCGCGGAGGCTGGGCCGGACGTCCGGTCCTGGACGG 12078

TeratornLC199500 CTTTCTTA-GACGGTCGTTCTTT-TGCTCGGAGCTCTGATGCCAGGGGCTCAGTCGAGATTTTGCCCTCCTCCTCGGCTTTCTCATCCGTCTCGCGTCCGCCCCAGCGGCTGAAA---AC 8254
WF-Teratorn      CCTCCGGGCGAAGGTTGTCCCACGTGGCTGAGTCCCCGGCGTCAGAGCTCCACTCGAAAG---ACCAACCGTCTGAGAT-----AGCTGGCT-GCG-------CGGAGGCTGACATGTAC 12182

TeratornLC199500 CGAGGACG-ACGAGGAGCAAA---AGCCCC--AAGGTGGAAGACGATGGGCTGAACGTTGTAAGAGAGCA-CGGTTCAGG--GACACCTTAGAAAAGCCGAG-GCGGTGCACTGCGTTCG 8364
WF-Teratorn      CCCTGTTGTAGGAGCAGTGCATCCAGCCCCTCAACGT---AGCTTATGG------CGTTG-AGGAGAATGTCCATATTGGTCGGCTTCTTGGGGGGGTCGAACGCAGTGCACCGC-CTCA 12291

TeratornLC199500 ACCCCCCCAAGAAGCCGACCAATATGGACATTCTCCT-CAACG------CCATAAGCT---ACGTTGAGGGGCTGGATGCACTGCTCCTACAACAGGGGTACATGTCAGCCTCCG----- 8469
WF-Teratorn      GCTTTTCTAAGGTGTC--CCTGAACCG-TGCTCTCTTACAACGTTCAGCCCATCGTCTTCCACCTTG--GGGCT---TTTGCTCCTCGT-CGTCCTCGGT---TTTCAGCCGCTGGGGCG 12399

TeratornLC199500 --CGC-AGCCAGCT-----ATCTCAGACGGTCGGTC---TTTCGAGTGGAGCTCTGACGCCGGGGACTCAGCCACGTGGGACAACCTTCGCCCGGAGGCCGTCCAGGACCGGACGTCCGG 8578
WF-Teratorn      GACGCGAGACGGATGAGAAAGCCGAGGAGGACGGCGAAATCTCGACTGAGCCCCTGGCATCAGAGCTCCGAGCAAA-AGAACGACCGTC-TAAGAAAGCAGTCTGAGCCCG--CGTCCCT 12515

TeratornLC199500 CCCGGCCTCCGCGCAGCCCGCTTTCTCAGA--------------CGGTTGCTCTTACTGCGCCT-------GACTAAAACTGCAGAGAGA----CCTTCTCGGTAGTACTAGGACCGTCA 8673
WF-Teratorn      CCTTCCTCCCGTGATGGCTGAGATGACAGAGAGGACTTGAACCCCTGCTGCAGCTGCTGCGCCTCCACGTAGCGTATGGCCTTGAGGAGGATGTCCATCTGGGTGGGCCTCTGGTGG--G 12633

TeratornLC199500 GCTCAGA-ACGATGCCTTGCTTAATTTTTCATTATTTTTTTTTTA-TTTTATTTTTTAATGAGACCAG--GCTGGACTCTTTACAGCCATTCCCCCCATTCGTTCTGAC-ATGCAA--GA 8786
WF-Teratorn      GTTCGAACGCAGTGCACCGCCTAAGCTTTTCTAAGGTGTCCTTGAACTGTGCTCTCTGCCGGCGTTAATAAGTAGACGTTTTA------TT-------TTCTTTTTGACTAGGCTACTGT 12740

TeratornLC199500 TAAGTGTGTGTGTTGGATTATGTAAAGGAAATTGTTTTAGTATTATCATTGGTTTAAATGTGTATGCTTAAATGTTAACTCAGATT---GACCTTTTTACCTTTTTAGAAAGATTTGTGT 8903
WF-Teratorn      CAACTCGACGTTTT---TTATATCTGGGC-----TTTTAGGGGTGA----GGGTCAATCG---GTGACCAATCGGTGAC-CAGCTGCAGGGGGTGTGCACGCTTTTGG--------GTGT 12836

TeratornLC199500 GTTTGTTCATCTGCTTCTCATCCCAGCACGGCTGAGGGGGTTTGACGACACACTGTCCTTAGCTGATAAGGACTGGTTGAAGTTAGAATGCACCAAATGGGTCATTATACGACACCCACA 9023
WF-Teratorn      GTGCGAGCGGGCGTT----ATGACTGTAAGTCAGAAGGGGATGCTCCACCAACAGAACTCTCCAGG-AAAGTCCGGCTGCTGTTAG---GCACTGCGTG--CCGCAGCCCG----CCTCA 12942
```

```
TeratornLC199500  ATGAACTTTTTTCTGTGTTTTGAAATTTTTACCTTCTTCTTGTGTTCTTAATAAAAACAGCACGAGGAGAGA------CAGAACCTTGGTGTTGCTCCACCAACAGAACTCTCCAGGAAA   9137
WF-Teratorn       GCCTGCCTCGTTCTCCGCGTTGGCCTCTCGAACT-------GTGGTGGTAG--GACGCAGTGTCTAGCGGGAGCGTTTCGTGTCCGGGGAGAAAATACGCCTA-AGAGCTCTGACGCTTG  13052

TeratornLC199500  GTCC---GGC--TCGCTTCAAAGCCTGAAGATTTGTTTTCCAAATTCCTCACTCCACACGCACGCACAAATCAGTGAACATTTCAGACAAATCTTTACATTATCGTTGGTTCTGTGTTGC   9252
WF-Teratorn       GTTCCCTGACCATCGCTTCAAAGCCTGAAGATTTGTTTTCCAAATTCCTCACTCAACACGCACGCACAAATCAGTGAACATTTCAGACAAATCTTTACATTATCGTTGGTTCTGTGTTGC  13172

TeratornLC199500  ATTTTGGACCGTAAAGGGGGTACAAGGTGAGGATGAAAATAACCAAGAGCAGCATCGCCGTTACGGTTACTAGAAAAACCTAACTGGAGTATGTTGGAGGAGGAGGAGAGTCTGGAGCTC   9372
WF-Teratorn       ATTTTGGACCGTAAAGGGGGTACAAGGTGAGGATGAAAATAACCAAGAGCAGCATCGCCGTTACGGTTACTAGAAAAACATAACTGGAGTATGTTGGAGGAGGAGGAGAGTCTGGAGCTC  13292

TeratornLC199500  CAGCTGGAATCATGTTGGTCAAATTCAGCATGTAGCCCGGAGGCTGTCCTGCATCGCTGTTTTTCATACGGCAGATGAAGGGAAACAGAAATTACTCTCAACAAACACAGGCCTGGATGA   9492
WF-Teratorn       CAGCTGGAATCATGTTGGTCAAATTCAGCATGTAGCCCGGAGGCTGTCCTGCATCGCTGTTTTTCATACGGCAGATGAAGGGAAACAGAAATTACTCTCAACAAACACAGGCCTGGATGA  13412

TeratornLC199500  GGCAGTTATATTTTAATAAATATTGTTATTTCATTGAAGGGAACCTGAACGAATGACACTTTGTTCATACATATAGCACTTTACAACTCTTTTAAGGTGCGAAAAGGGCTTCGCGATAAA   9612
WF-Teratorn       GGCAGTTATATTTTAATAAATATTGTTATTTCATTGAAGGGAACCTGAACGAATGACACTTTGTTCATACATATAGCACTTTACAACTCTTTTAAGGTGCGAAAAGGGCTTCGCGATAAA  13532

TeratornLC199500  AGAAAGAGAAAAGACATTGATTCTGTAGACAAGGCTTGTGCTGGAGGCCTCAGGCAGAAATCAAACTGTTGAAATGCT------------------------------------------   9690
WF-Teratorn       AGAAAGAGAAAAGACATTGATTCTGTAGACAAGGCTTGTGCTGGAGGCCTCAGGCAGAAATCAAACTGTTGAAATGCTCCGATGGCCAGGTTTCAGCCGAAGCATGCCACCGTCTCCGCT  13652

TeratornLC199500  ------------------------------------------------------------------------------------------------------------------------   9690
WF-Teratorn       AGTTGGCTCATCGAAGCCTGTCAGCTCCATCCTGCGTGCTCTCATTGTGCACATTTGGGTTATGCTACATTAAACTCAGCGATACACTACGTGACTAATAAAAACAGATCTAATATGAGC  13772

TeratornLC199500  ------------------------------------------------------------------------------------------------------------------------   9690
WF-Teratorn       ATACAAATTTAAGTAAAATCCACATCTAGTTTAAGATAATGCAAAAGTTATATAAAAGTTTCAGCAGTTTCAATAAAAGTGTTCAGCGTAAATATAAACAAGATTATAAGCTAAATGGGA  13892

TeratornLC199500  ------------------------------------------------------------------------------------GGTCTATCCCTCCTGCTCCTCTCGCAATCTGTATTT   9726
WF-Teratorn       ATAAAATGATTATCGTCCTTCAACACACACACACACACACACACACACACACATACATTCATACATACATACATCAAATGCTGGTCTATCCCTCCTGCTCCTCTCGCAATCTGTATTT  14012

TeratornLC199500  T------TTTTTTAAATGTATTTTACATATGTAAAGGAAAATAAAGAAAGATATAAACAAACAAGTAACACTCTGGTCTTTGCCTGAGCTGAGAAACTCCCTGCTTTGAAACACAGAAGA   9840
WF-Teratorn       TTTTTTATTTTTTAAATGTATTTTACATATGTAAAGGAAAATAAAGAAAGATATAAACAAACAAGTAACACTCTGGTCTTTGCCTGAGCTGAGAAACTCCCTGCTTTGAAACACAGAAGA  14132

TeratornLC199500  AAACTGCTGATCGGCAGAGTTCCCT-GCCCC----------------------TCCTT-CCTCA------AAACTCCCTGCT-----------------------CGCAGCT--------   9899
WF-Teratorn       AAGCTGCTGATCGGCCGAGTTCCCTTGCTTCCTCAAAAAAAAAAACACTCTGGTCTTTGCCTGAGCTGAGAAACTCCCTGCTTTGCATTCTTCCCCCCCTGAGCCGTGGCCTCTCGAGT  14252

TeratornLC199500  ----GCAG-GCACCACGGTT-------------------------------CCTCC---------------------TCTGC----------------------GTCTG-----------   9929
WF-Teratorn       AAGAGCAGAGCGCCGAGGCTGTGTGGCTCTGGGGCGGTCCGAGTCAGTCAGCCTCCAAGAGCCTAGGCCAAATCATCTCTGTCTCTCTCCCCCCTCTCTCCCCAGTCTGTGCCTCTTTCA  14372

TeratornLC199500  ------------------------------------CCTTG---CTGTGTCTCCC---------------------------------------------GCCCGCT-------------   9952
WF-Teratorn       GTCACACACATTCCTTCTCTCCTCCTCCTCCTCTCTCCTTTTCTCTGTGTTTCTCTGTCGCGCTCGTTCTCTGAGCCCCTTCGGAAAGCTTTCTGTGTCTGCCCGCTGCCGTCATCCCTG  14492

TeratornLC199500  -----------------------------------------CGTGAGT--------------------GAGGCT------CCACGGGGCGGTCCGAGTCAGTCAGCCTCCAAGAGCCGAG  10005
WF-Teratorn       ATCTGGCCCTACAAAAAGCTGAGCTGTGGACACGTATTTGGTGTGTGTAAAAATGCAAACAGAGCGCCGAGGCTGTGTGGCTCTGGGGCGGTCCGAGTCAGTCAGCCTCCAAGAGCCTAG  14612

TeratornLC199500  GCCAAATCATTTCTGTCTTTCTCCCCCCTCTCTCCCCAGTCTTTGCCTCTTGCAGTCACACACATTCCTTCTCTCCTCCTCCTCCTCCTCCTTCTTCCATTTCTCTCTGTTTCTCTGTCG  10125
WF-Teratorn       GCCAAATCATTTCTGTCTCTCTCCCCCCTCTCTCCCCAGTCTGTGCCTCTTGCAGTCACACACATTCCTTCTGTCCTCCTCCTCCTCCTGCTTCTTCCATTTCTCTCTGTTTCTCTGTCG  14732

TeratornLC199500  CGCGCGTTCTCTGAGCCCCTTCGGAAAGCTTTCTGTGTCTGCCCACTGCCCTCATCCCTGATCTGGCCCCACAAAAAGCTGAGCTGTGG----ACACGT---------------------  10220
WF-Teratorn       CGCGCGTTCTCTGAGCCCCTTCGGAAAGCTTTCTGTGTCTGCCCACTGCCCTCATCCCTGATCTGGCCCCACAAAAAGCTGAGCTGTGGTGTGACTCGTCTGAGAGTAAGGCTACATTTT  14852

TeratornLC199500  ------------------------------ATTTGGT---------------GTGTGTAAAAA-------------------------------------------------------TG  10240
WF-Teratorn       TTCCCCCCTGAGCTGGCCTCTCGAGCCCTGATCTGGCCCCACAAGTCAGTCAGCCTCCAAGAGCCTAGGCCAAATCATCTCTGTCTCTCTCCCCCCTCTCTCCCCAGTCTGTGCCTCTTG  14972

TeratornLC199500  CAAACA--------------------------------------------------------------------GAGCGCC----GAGG-----CTGTGT---------GGCTCTGGGGC  10274
WF-Teratorn       CAGTCACACACATTCCTTCTGTCCTCCTCCTCCTTCTTCCATTTCTCTCTGTTTCTCTGTCGCGCGCGTTCTCTGAGCCCCTTCGGAAAGCTTTCTGTGTCTGCCCACTGCCTCTCGAGC  15092

TeratornLC199500  ---GGTCCG--------AGTCAGTCAGCCTCCAAGAGCCTAGGCAAAATCATCTCTGTCTCTCTCCCCCCTCTCTCCCCAGTCTGTGCCTCTTTCAGTCACACACATTCCTTCTCTCCTC  10383
WF-Teratorn       CCTGATCTGGCCCCACAAGTCAGTCAGCCTCCAAGAGCCTAGGCCAAATCATCTGTGTCTCTCTCCCCCCTCTCTCCCCAGTCTGTGCCTCTTGCAGTCACACACATTCCTTCTCTCCTC  15212
```

TeratornLC199500 CTCCTCCTTCT------TCCATTTCTCTCTGTTTCTCTGTCGCGCGCGTTCTCTGAGCCCCTTCGGAAAGCTTTCTGTGTCTGCCCGCTGCCGTCATCCCTAATCTGGCCCCACAAAAAG 10497
WF-Teratorn CTCCTCCTCTTCCTCTCTCCTTTTCTCTCTGTTTCTCTGTCGCGCGCGTTCTCTGAGCCCCTTCGGAAAGCTTTCTGTGTCTGCCCACTGCCCTCATCCCTGATCTGGCCCCACAAAAAG 15332

TeratornLC199500 CTGAGCTGTGGACACGTATTTGGTGTGTGTAAAAATGCAAACAGAGCGACGAGGCTGTGTGGCTCTGGGGCGGTCCGAGTCAGTCAGCCTCCAAGAGCCTAGGCAAAATCATCTCTGTCT 10617
WF-Teratorn CTGAGCTGTGGACACGTATTTGGTGTGTGTAAAAATGCAAACAGAGCGCCGAGGCTGTGTGGCTCTGGGGCGGTCCGAGTCAGTCATTCTCCAAGAGCCGAGGCCAAATAATCTCTGTCT 15452

TeratornLC199500 CTCTCCCCCCTCTCTCCCCAGTCTGTGCCTCTTGCAGTCACACACATTCCTTCTCTCCTCCTCCTCCTC-----------TCTCCTTTTCTCTCTGTTTCTCTGTCGCGCGCGTTCTCTG 10726
WF-Teratorn CCAACCCCC---TCTCCCCAGTCTCTGCCTCTTTCAGTCACACACACTCCTTTTCTCCTCCTCCTCCTCCTTCTTCCCTTTCTCTTTGTTTCTCTGTTTCTCTGTCGCGCGCGTTCTCTG 15569

TeratornLC199500 AGAACCTTCGGAAAGCTTTCTGTGTCTGCCCGCTGCCCTCATCCCTGATCTGGCCCCACAAAAAGCTGAGCTGTGGACACGTATTTGGTGTGTGTAAAAATGCAAACACGTCGTCTTCAG 10846
WF-Teratorn AGCCCCTTCGGAAAGCTTTCTGTGTCTGCCCACTGCCCTCATCTCTGATCTGGCCCCACAAAAAGCTGAGCTGTGGACACGTATTTGGTGTGTGTAAAAATGCAAACACGTCGTCTTCAG 15689

TeratornLC199500 AGAGGAAAATTCCACTCTGGCCTGCCTGCCGCTGCTTCCTCAGGGGCCCTCTGTGGGGGCAGGCTCTCAGAGCTCTGCGAATGAGCCGAAGCCAAGCGGCTTTGGACGTGAC-------- 10958
WF-Teratorn AGAGGGAAATTCCACTCTGGCCTCCCTGCCGCTGCTTCCTCAGGGGCCCTCTGTGGGGGCAGGCTCTCAGAGCTCTGCGAATGAGCCGAAGCCAAGCGGCTTTGGACGTGACACACTCTG 15809

TeratornLC199500 ----TTGC--------AGACACTC---------AGGCGTGTAAAA----------CCAGCAATGCTCCAACGAGCATCCTTTTT----ATTTTCTCTTTGTGATTAGTGACATTGAATAT 11043
WF-Teratorn GTCTTTGCCTGAGCTGAGAAACTCCCTGCTTTGAAACACAGAAGAAAACTGCTGATCGGCAGAGTTCCC-TGCCCCTCCTTCCTCAAAACTCCCTGCTCGCAGCTGCAGGCACCACGGTT 15928

TeratornLC199500 CCTGCTCCAACTCT----------TTCTCCATTTCGCT---GAGCTA---TCC-CTG---GGTTGTAATGA----------AAAA----AAAATAAATAA-----------AAAGTCATC 11118
WF-Teratorn CCTCCTCTGGGTCTGCCTTGCTGTTTCTCCCGCCCGCTCGTGAGCGAGGCTCCACTGTGCGGTCCGAGTCAGTCAGCCTCCAAGAGCCGAAGCCAAATTATCTCTGTCTCCAACCCCCTC 16048

TeratornLC199500 GC----------GCACCAAA-------------------------------------------------------GAAATTTCTGTGCTGCTTTGGA---TTTTCTGT-------CACTTTT 11165
WF-Teratorn TCCCCAGTCTCTGCCTCACATTCCTTCTCTCCTCCTCCTCCTTCTTCCCTTTCTCTCTGTTTCTCTGTCGCGCGCGTTCTCTGAGCCCCTTCGGAAAGCTTTCTGTGTCTGCCCACTGCC 16168

TeratornLC199500 TTTAATTTTTGAT----TTCTGTTA----------TACAGCT----AGAGAAAAA-----CCTT--TCCCAAG-----------------GAGCTCTGCGAAGACATAGGCCA------- 11236
WF-Teratorn CTCA-TCTCTGATCTGGCTCTGCGAAGACAGGGGCCACGGCTCAGGGGGGAAGAATGCAGCCTTACTCTCAGACGAGTCACACAGCCTCTGAGCTCTGCGAAGACAGGGGTCAGAGGCTG 16287

TeratornLC199500 --CGGCTCTG----------------TCTT------------------GTGGCCTC----------------------------------------TGTGACTCTTCTGAGAGGAAG--- 11277
WF-Teratorn TGCGGCTCTGGTTCGACAGGCTGCATTCTTCCCCCACCAAGAAAGGCTGCGGCCTGCCCCTGGGCTGACCCCTGTCTTCGCAGAGCTCAGAGGCTGTGTGACTCGTCTGAGAGTAAGGCT 16407

TeratornLC199500 ----------------------------------------AGCT---------------TGGCTTTGGCTCA-------------------------------------------------- 11294
WF-Teratorn GCATTCTTCCCCCCTGAGCCGTGGCCCCTGTCTTCGCAGAGCTCAGAGGCTGTGCGGCTGGCTCTGGTTCGACAGGCTGCATTCTTCCCCCACCAAGAAAGGCTGCGGCCTGCCCCTGGG 16527

TeratornLC199500 --------------------------------------------------------------------------------------------TTCGCAGAGCTCAGAGGCTGTGCGGCTC---- 11322
WF-Teratorn CTGACCCCTGTCTTCGCAGAGCTCAGAGGCTGTGTGACTCGTCTGAGAGTAAGGCTGCATTCTTCCCCCCTGAGCCGTGGCCCCTGTCTTCGCAGAGCTCAGAGGCTGTGCGGCTCTGCT 16647

TeratornLC199500 -----AGGTTTC----------------GAAAGGCTGCGGCCTGCCCCTGGGCTGACCCCTGTCTTCGCAGAGCTCAGAGGCTGTGCGTCT----CTGAT------GGC----------- 11400
WF-Teratorn TCAACAGGCTGCATTCTTCCCCCACCAAGAAAGGCTGCGGCCTGCCCCTGGGCTGACCCCTGTCTTCGCAGAGCTCAGAGGCTGTGCGGCTGGCTCTGCTTCGACAGGCTGCATTCTTCC 16767

TeratornLC199500 --------GAAAGGCTG----CCGCCTCCAGCCTGCCTCCTGCCT---CTGAGCC-----GTGCTGTGACTCGTCTGAGAGTAAGGCTGCATTCTTCCCCCCTCTGAGCAGTGGCCCCTGT 11500
WF-Teratorn CCCACCAAGAAAGGCTGCGGCCTGCCCCTGGGCTGACTCCTGTCTTCGCAGAGCTCAGAGGCTGTGTGACTCGTCTGAGAGTAAGGCTGCATTCTTCCCCCCTCTGAGCCGTGGCCCCTGT 16887

TeratornLC199500 CTTCGCAGAGCTCAGAGGCTGTGCGGCTGGCTCTGCTTCGACAGGCTGCATTCTTCCCCCACCAAGAAAGGCTGCGGCCTGCCCCTGGGCTGACCCCTGTCTTCGCAGAGCTCAGAGGCT 11620
WF-Teratorn CTTCGCAGAGCTCAGAGGCTGTGCGGCTGGCTCTGGTTCGACAGGCTGCATTCTTCCCCCACCAAGAAAGGCTGCGGCCTGCCCCTGGGCTGACCCCTGTCTTCGCAGAGCTCAGAGGCT 17007

TeratornLC199500 GTGTGACTCGTCTGAGAGTAAGGCTGCATTCTTCCCCCCTGAGCCGTGGCCCCTGTCTTCGCAGAGCTCAGAGGCTGTGCGGCTGGCTCTGGTTCGACAGGCTGCATTCTTCCCCCACCA 11740
WF-Teratorn GTGTGACTCGTCTGAGAGTAAGGCTGCATTCTTCCCCCCTGAGCCGTGGCCCCTGTCTTCGCAGAGCTCAGAGGCTGTGCGGTTGGCTCTGGTTCGACAGGCTGCATTCTTCCCCCACCA 17127

TeratornLC199500 AGAAAGGCTGCGGCCTGCCCCTGGGCTGACTCCTGTCTTCGCAGAGCTCAGAGGCTGTGTGACTCGTCTGAGAGTAAGGCTGCATTCTTCCCCCCTCTGAGCCGTGGCCCCTGTCTTCGCA 11860
WF-Teratorn AGAAAGGCTGCGGCCTGCCCCTGGGCTGACCCCTGTCTTCGCAGAGCTCAGAGGCTGTGTGACTCGTCTGAGAGTAAGGCTGCATTCTTCCCCCCTCTGAGCCGTGGCCCCTGTCTTCGCA 17247

TeratornLC199500 GAGCTCAGAGGCTGTGCGGCT-----------------------------------------------------------------------------------------------GGC 11884
WF-Teratorn GAGCTCAGAGGCTGTGCGGCTCTGCTTCGACAGGCTGCATTCTTCCCCCACCAAGAAAGGCTGCGGCCTGCCCCTGGGCTGACCCCTGTCTTCGCAGAGCTCAGAGGCTGTGCGGCTGGC 17367

TeratornLC199500 TCTGCTTCGACAGGCTGCATTCTTCCCCCACCAAGAAAGGCTGCGGCCTGCCCCTGGGCTGACCCCTGTCTTCGCAGAGCTCAGAGGCTGTGTGACTCGTCAGAGAGTAAGGCTGCATTC 12004
WF-Teratorn TCTGGTTCGACAGATTGCATTCTTCCCCCACCAAGAAAGGCTGCGGCCTGCCCCTGGGCTGACCCCTGTCTTCGCAGAGCTCAGAGGCTATGTGACTCGTCTGAGATTAAGGCTGCATTT 17487

TeratornLC199500 TTCCCCCCTGAGCCGTGGCCCCTGTCTTCGCAGAGCTCAGAGGCTGTGCGGCTGGCTCTG------------------------------------------------------------ 12064
WF-Teratorn TTTTCCCCTGAGCCGTGGCCCCTGTCTTCGCAGAGCTCAGAGGCTGTGCGGCTGGCTCTGGTTCGACAGATTGCATTCTTCCCCCACCAAGAAAGGCTGCGGCCTGCCCCTGGGCTGACC 17607

TeratornLC199500 ----------------------------------------CTTCGACAGGCTGCTTTCTTCCCCCACCAAGAAAGGCTGCGGCCTGCCCCTGGGCTGACTCCTGTCTTCGCAGAGCTCAG 12144
WF-Teratorn CCTGTCTTCGCAGAGCTCAGAGGCTGTGCGGCTGGCTCTGCTTCAACAGGCTGCATTCTTCCCCCACCAAGAAAGGCTGCGGCCTGCCCCTGGGCTGACCCCTGTCTTCGCAGAGCTCAG 17727

TeratornLC199500 AGGCTGTGCGGCTG----TGGTTCGACAGGCTGCATTCTTCCCCCACCAAGAAAGGCTGCGGCCTGCCCCTGGGCTGACTCCTGTCTTCGCAGAGCTCAGAGGCTGTGTGACTCGTCTGA 12260
WF-Teratorn AGGCTGTGCGGCTGGCTCTGCTTCGACAGGCTGCATTCTTCCCCCACCAAGAAAGGCTGCGGCCTGCCCCTGGGCTGACTCCTGTCTTCGCAGAGCTCAGAGGCTGTGTGACTCGTCTGA 17847

TeratornLC199500 GAGTAAGGCTGCATTCTTCCCCCTCTGAGCCGTGGCCCCTGTCTTCGCAGAGCTCAGAGGCTGTGCGGCTGGCTCAACTTCGACAGGCTGCATTCTTCCCCCACCAAGAAAGGCTGCGGC 12380
WF-Teratorn GAGTAAGGCTGCATTCTTCCCCCTCTGAGCCGTGGCCCCTGTCTTCGCAGAGCTCAGAGGCTGTGCGGCTGGCTCTGGTTCGACAGGCTGCATTCTTCCCCCACCAAGAAAGGCTGCGGC 17967

TeratornLC199500 CTGCCCCTGGGCTGACTCCTGTCTTCGCAGAGCTCAGAGGCTGTGTGACTCGTCTGAGAGTAAGGCTGCATTCTTCCCCCTCTGAGCAGTGGCCCCTGTCTTCGCAGAGCTCAGAGGCTG 12500
WF-Teratorn CTGCCCCTGGGCTGACTCCTGTCTTCGCAGAGCTCAGAGGCTGTGTGACTCGTCTGAGAGTAAGGCTGCATTCTTCCCCCTCTGAGCCGTGGCCCCTGTCTTCGCAGAGCTCAGAGGCTG 18087

TeratornLC199500 TGCGGCTGGCTCTGCTTCGACAGGCTGCATTCTTCCCCCACCAAGAAAGGCTGCGGCCTGCCCCTGGGCTGACCCCTGTCTTCGCAGAGCTCAGAGGCTGTGTGACTCGTCTGAGAGTAA 12620
WF-Teratorn TGCGGCTGGCTCTGCTTCGACAGGCTGCATTCTTCCCCCACCAAGAAAGGCTGCGGCCTGCCCCTGGGCTGACCCCTGTCTTCGCAGAGCTCAGAGGCTGTGTGACTCGTCTGAGAGTAA 18207

TeratornLC199500 GGCTGCATTCTTCCCCCCTGAGCCGTGGCCCCTGTCTTCGCAGAGC**TCAGAGGCTGTGCGGCTGGCTCTGGTTCGACAGGCTGCATTCTTCCCCCACCAAGAAAGGCTGCGGCCTGCCCC** 12740
WF-Teratorn GGCTGCATTCTTCCCCCCTGAGCCGTGGCCCCTGTCTTCGCAGAGCTCAGAGGCTGTGCGGTTGGCTCTGGTTCGACAGGCTGCATTCTTCCCCCACCAAGAAAGGCTGCGGCCTGCCCC 18327

TeratornLC199500 **TGGGCTGACCCCTGTCTTCGCAGAGCTCAGAGGCTGTGTGACTCGTCTGAGAGTAAGGCTGCATTCTTCCCCCCTGAGCCGTGGCCCCT---GTCTTCGCAGAGCTCAGAGGCTGTGCGG** 12857
WF-Teratorn TGGGCTGACCCCTGTCTTCGCAGAGCTCAGAGGCTGTGTGACTCGTCTGAGAGTAAGGCTGCATTCTTCCCCCCTGAGC--TGGCCTCTCGAGAAAGAGCAGAGCGCCGAGGCTGTGTGG 18445

TeratornLC199500 **CTTTGAAA-ACACAGAAGAAAACTGCT-----GATC---GGCCGAGT--TCCCTGC--CCCCTCCTTCCTCAAAACTCCCTGCTCG---CAGCTGCAGGCACCACAGTTCCTC----CTC** 12957
WF-Teratorn CTCTGGGGCGGTCCGAGTCAGTCAGCCTCCAAGAGCCGAGGCCAAATCATCTCTGTCTCCCCCCCTCTCTCCCCAGTCTCTGCCCCTTTCAGTCACACACACTCCTTCTCTCCTGAGCTG 18565

TeratornLC199500 **TGGGTCTGC-CTTGCTGTTTCTC----------CCGCCCGCTCGTGA--GTGAGGCTCCACTGGGCGGTCCGAGTCAGTCAT**TCTCCAAGAGCCGAGGCCAAATCATCTCTGTCTCCAAC 13064
WF-Teratorn TGGACACGTATTTGGTGTGTGTAAAAATGCAAACAGAGCGC-CGAGGCTGTGTGGCTCTG--GGGCGGTCCGAGTCAGTCATTCTCCAAGAGCCGAGGCCAAATCATCTCTTTCTCCAAC 18682

TeratornLC199500 CCCCTCTCCCCAGTCTCTGCCTC------------ACATTCCTTCTCTCCTCCTCCTCCTCCTCCACCTCCTCCTTCTTCCCTTTCTCTCTGTTTCTCTGTCGCGAGCGTTCTCTGAGCC 13172
WF-Teratorn CCCCTCTCCCCAGTCTCTGCCTCTTTCAGTCACACACACTCCTTCTCTCTTCCTCCTCCTCCTTCTTC----CCTTTCTCTCTGTTTCTCTGTTTCCCTGTCGCGCGCGTTCTCTGAGCC 18798

TeratornLC199500 CCTTCGGAAAGCTTTCTGTGTCTGCCCACTGCCCTCATCTCTGATCTGGCCCCACAAAAAGCTGAGCTGTGGACACGTATTTGGTGTGTGTAAAAATGCAAACACGTCGTTTTCAGGGAG 13292
WF-Teratorn CCTTCGGAAAGCTTTCTGTGTCTGCCCACTGCCCTCATCTCTGATCTGGCCCCACAAACAGCTGAGCTGTGGACACGTATTTGGTGTGTGTAAAAATGCAAACACGTCGTCTTCAGAGAG 18918

TeratornLC199500 GGA----------------------------------------------------------------GCTCTGCGAATGAGCCGAAGCCAAGCGGCTTTGGACGTGACTTGCAGACACTC 13348
WF-Teratorn GGAAATTCCACTCTGGCCTCCCTGCCGCTGCTTCCTCAGGGGCCCTCTGTGGGGGCAGGCTCTCAGAGCTCTGCGAATGAGCCGAAGCCAAGCGGCTTTGGACGTGACTTGCAGACACTC 19038

TeratornLC199500 AGGCGTGTAAAACCAGCAATG--------------------------------------------------------------------------------------------------- 13369
WF-Teratorn AGGCGTGTAAAACCAGCAATGTTCCTATGGCCAAGCTGGCTCCAGCTGAAACTAGGAGGCCTGCAGATTCTTACTGCGCCATCACGCACCGTTCTCCCTTCTCAGAGGAGCAGCTTTTTA 19158

TeratornLC199500 ---------------------------------------------------------------------------------------------------------------CTCCAACGA 13378
WF-Teratorn TCATTGTTTAGCGCCGTTTCTGTCATAGGATGAATGCTTTTTGTTGTTTTTTTTAGAGTTTCGGTACATCTAGTCAAAGAAATCATTCGGGTGCCTCTGAAATGAACCCAGCTCCAACGA 19278

TeratornLC199500 GCATCCTTT**TTATTTTCTCTTTGTGATTAGTGACATTGAAAATCCTGCTCGGCATAGCTGCGAGACGCTGCTTTTCTTGCAGGGCCCTGCCCGCTCCCTGAGCTGAGAACCTCCCTGCTT** 13498
WF-Teratorn GCATCCTTTTTATTTTCTCTTTGTGATTAGTGACATTGAAAATCCTGCTCGGCATAGCTGCGAGACGCTGCTTTTCTTGCAGGGCCCTGCCCGCTCCCTGAGCTGAGAACCTCCCTGCTT 19398

TeratornLC199500 **TGAAACACAGAAGAAAGCTGCTGATCGGCCGAGTTCCCTTGCTTCCTCAAAAAAAA----AAACTCTGGTCTTTCCTCGCAGCTGCAGGCAGCACGGTCCCTCCTCTGCGTCTGGCCTGC** 13614
WF-Teratorn TGAAACACAGAAGAAAGCTGCTGATCGGCCGAGTTCCCTTGCTTCCTCAAAAAAAAAAAAAAACTCTGGTCTTTGCTCGCAGCTGCAGGCAGCACGGTCCCTCCTCTGCGTCTGGCCTGC 19518

TeratornLC199500 **TGTTTCTCT**--------------------------------------------------------------------------------------------------------------- 13623
WF-Teratorn TGTTTCTCTCGCCTGCTCGTGAGTGTGTGCAGAGTGCGTTCTCTGTCTCGCAGGAGCCCTGTGTGGGGGCAGGCTGAAGGCTTTCGCAACCAGAGCCACACAGCCTCAGAGCTCTACGAA 19638

TeratornLC199500 ------------------------------------------------------------------------------------------------------------------------ 13623
WF-Teratorn TGAGCCAACAATTACACAGCCTCGCGGCTGTAAGGTGGAGTCTGTTGCTGTCCCAGAAGCCCGCAGCCCCTGAGTGCACCATCACGCACCCTCCTCCCTTCTGGCCACTGGAGCAGCTTT 19758

TeratornLC199500 ------------------------------------------------------------------------------------------------------------------------ 13623
WF-Teratorn TCATGAGCCGTGTCCGTAGCTCCCTTTGTCTTGGCCCCAAAACCTGAGCTCTGCACACGTATTTGGTGTGAGCTTTGGATGTGAGATTTCAGCGGCTGTCTTCTCTTCGCTCTACGCGAA 19878

TeratornLC199500 ------------------------------------------------------------------------------------------------------------------------ 13623
WF-Teratorn TAGAACTCCTTCGAAGACAGGATTTGGTTATTGTTTTTTGTTTTCTGTTGTCTTTTTCTTTTTTCTTTTCTTTGTTTCTCTCTCATCCAAGTTTCTGTTTGTTGAGAGTCATTTATGTTT 19998

TeratornLC199500 -------------------------------------**GGCTGTTCTCTTGGCTTGACAGCGGAGGGGATTTCTCTAACGTGCCTCGGTCGGCCTCACAGAGCGGAGGTCTCCGCAT**GCCT 13706
WF-Teratorn CCCTTCATCTGCCATGTAAAAAAAAAAACAGCGATGCAGGCTGTTCTCTTGGCTTGACAGCGGAGGGGATTTCTCTAACGTGCCTCGGTCGGCCTCACAGAGCGGAGGCCTCCGCATGCCT 20118

TeratornLC199500 CCCTCCCCCCTCTATCTCTCTCGCGCGTGTTCTCTGAGCTCGTTCTACAGCTTTCTGTGTCTGTGTCCCTGTGTCTGGGGAGGGGATTTTTCTAAAGCACACGCCAACCTAAGCAAGTGC 13826
WF-Teratorn CCCTCCCCCCTCTATCTCTCTCGCGCGTGTTCTCTGAGCTCCTTCGACAGCTTTCTGTGTCTGTGTCCCTGTGTCTGGGGAGGGGATTTTTCTAAAGCACACGCCAACCTAAGCAAGTGC 20238

TeratornLC199500 GAGGGGTAAGGAAATGAAGAGAGAGAAAAAAAAAAAGAATGTCTGGTTGGTCACTCACAGTAAAACAGTGGATGGATAAAAAAAAAAAAAAACACAGGGGTCAAATGCACCCTAAAGTACA 13946
WF-Teratorn GAGGGGTAAGGAAATGAAGAGAGA-AAAAAAAAAGGAATGTCTCGTTGGTCACTCACAGTAAAACAGTGGATGGATTAAAAAAAAAA----CACAGGGGTCAAATGCACCCTAAAGTACA 20353

TeratornLC199500 GCGCGAGGGAATTGCCAAATGCAGGCCACCAAGTAATGTCTCGTGGGGTCATTTATACTCTGTGTAAATGAAAGTTTGTGAAAGTGAAAGTTTTTTTTTTTTTTAAAGTACGGGGTCAAA 14066
WF-Teratorn GCGCGAGGGAATTGCCAAATACCGGCCACCAAGTAATGTCTCGTGGGGTCATTTATACTCTGTA-AAACA-------GTGGA-GTGAAAGTTTTTTTTTTTTTAAAAAGTACGGGGTCAAA 20464

TeratornLC199500 TGCACCCTAAAGTACAGCGCGAGAGGTATGAAAATAAATAAATAAATTGATGCCCCCCTTCTCCCCCCCTCCACCCCCTGGTGGTCGTGGGGTGTAAGGACAATGAAGAAAACAAATACG 14186
WF-Teratorn TGCACCCTAAAGTACAGCGCGAGAGGTATGAAAATAAATAAATAAATTGATGCCCCCCTTCTCCCCCCCTCCACCCCCTGGTGGTCGTGGGGTGTAAGGACAATGAAGAAAAAAAAATACG 20584

TeratornLC199500 GGGTCAAATGCACCCAAAAGACAGCGCGAGGGGTATGAAAATAAATAAATAGATTGATGCCCCCCCTCATCCCCTCCACCCCCTGGTGGTCGTGGGGTGTAAGTACAATGAAGAGAGGAT 14306
WF-Teratorn GGGTCAAATGCACCCAAAAGACAGCGCGAGGGGTATGAAAATAAATAAATAGATTGATGCCCCCCCTCATCCCCTCCACCCCCTGGTGGTCATGGGGTGTAAGTACAATGAAGAGAGGAT 20704

TeratornLC199500 GGGGTCCTTTACGTTCCGTAAAACAGCGGAGTGAATGTTTTTTTTTTAAAAAAAATATGGGGTCAAATGCAACCAAAAGACAGCGCGAGGGGTATAAAAATAAATAAATAGATTGATGCC 14426
WF-Teratorn GGGGTCCTTTACGTTCCGTAAACCAGCGGAGTGAATGTTTTTTTTTTTAAAAA--TACGGGGTCAAATGCACCCAAAAGACAGCGCGAGGGGTATGAAAATAAATAAATAGTTTGATGCC 20822

TeratornLC199500 CCCCCTCATCCCCTCCTCTCCCTCCACCCCCTGGTGGTCGTGGGGTGTAAGGACAATGAAGAGATGAAAAAAATATGGGGTCAAATGCACGCCAAAGCACAGCGCGAGAGGTATGAAAAT 14546
WF-Teratorn CCCCCTCATCGCCTCCTCCCCCTCCACCCCCTGGTGGTCGTGGGGTGTAAGGACAATGAAGAGATGAAAAAAATATGGGGTCAAATGCACGCCAAAGCACAGCGCGAGAGGTATGAAAAT 20942

TeratornLC199500 AAATAAATAAATTGATGCCCCCCTTCTCCCCCCATCCACCCCCTGGTGGTCGTGGGGTGTAAGGACAATGAAGAAAAAAAAAAATACGGGGTCAAATGCACCCAAAAGACAGCGCGAGGGG 14666
WF-Teratorn AAATAAATA--------------------------------------------------------------------------------------------------------------- 20951

TeratornLC199500 TATGAAAATAAATAAATAGATTGATGCCCCCCCTCATCCCCTCCACCCCCTGGTGGTCGTGGGGTGTAAGTACAATGAAGAGAGGATGGGGTCCTTTACGTTCCGTAAACCAGCGGAGTG 14786
WF-Teratorn ------------------GATTGATGCCCCCCCTCATCCCCTCC---------------------------------------------------------------------------- 20977

TeratornLC199500 AATGTTTTTTTTTTTAAAAATACGGGGTCAAATGCACCCAAAAGACAGCGCGAGGGGTATGAAAATAAATAAATAGTTTGATGCCCCCCCTCATCGCCTCCTCCCCCTCCACCCCCTGGT 14906
WF-Teratorn -------------------------------------------------------------------------------------------------------TCCCCCTCCACCCCCTGGT 20996

TeratornLC199500 GGTCGTGGGGTGTAAGGACAATGAAGAGAGGATGATAAAAAATATGGGGTCAAATGCACGCCAAAGCACAGCGCGAGAGGTATGAAAATAAATAAATAAATTGATGCCCCCCTCCTCCCC 15026
WF-Teratorn GGTCGTGGGGTGTAAGGACAATGAAGAGATGA----AAAAAATATGGGGTCAAATGCACGCCAAAGCACAGCGCGAGAGGTATGAAAATAAATAAATAAATTGATGCCCCCCTCCTCCCC 21112

TeratornLC199500 CTCCACCCCCTGGTGGTCGTGGGGTGTAAGGACAATGAAGAGAGGATGATAAAAAAAGTATTGTCTCGTGGGGTCGTTTACGATCCGTAAAACAGCGGACTGAATGTTTTTTATGCCGGC 15146
WF-Teratorn CTCCACCCCCTGGTGGTCGTGGGGTGTAAGGACAATGAAGAGAGGATGATAAAAAAAGTATTGTCTCGCGGGGTCGTTTACGATCCCTAAAACAGCGGACTGAATGTTTTTTATGCCGGC 21232

TeratornLC199500 CACAGAGAAATTTCTCGTGGGGTCGTTTACGATCAGTAAAACAGCGGACTGAATGTTTTTTATGCCGGCCACTGAGAAATTTCTCGTGGGGTCACACGAGCTGCGTAAAACAGCGGATTT 15266
WF-Teratorn CACAGAGAAATTTCTCGTGGGGTCGTTTACGATCCGTAAAACAGCGGACTGAATGTTTTTTA------------------------------------GCTCCGTAAAACAGCGGATTT 21315

TeratornLC199500 TTTGGGGGGTCATGCAGACCCCAAAGCACATGCCCACCTGAGCGAGTGGGAGGGTTATATAAATACGGAGGTCAAATGCACCCCAAACTACAGCGCGAGGGAATTGCCAAATGCCATCCA 15386
WF-Teratorn TTTGGGGGGTCATGCAGACCCCAAAGCACATGCCCACCTGAGCGAGTGGGAGGGTTATATAAATACGGAGGTCAAATGCACCCCAAACTACAGCGTGAGGGAATTGCCAAATGCCATCCA 21435

TeratornLC199500 CCAAAAAAAAAAAAAAAAAA---CGCCTCGTGGGGTCACACGTGCTCCGATAAACAGCGGTTGGAATTTTTATTTTTTTTTGAAAAAAAAATACGGGGGTCAAATGCACCCCAAACTACAG 15503
WF-Teratorn CCAAAAAAAAAAAAAAAAAAAAACGCCTCGTGGGGTCACACGTGCTCCGAAAAACAGCGGTTGGAATTTTTTTTTTTTTTTTTTAAAAA-TACGGGGGTCAAATGCACCCCAAACTACAG 21554

TeratornLC199500 CGCGAGGGAATTGCCAAATGCCATAAACAAAAAAA---------------------------------------------------------AAAAT------------TATACC 15549
WF-Teratorn CGCGAGGGAATTGCCAAATGCCATAAACAAAAAAAATGCCTCGTGGGGTCACAGGTGCTCCGAAAAACAGCGGATGGGATTTTTTTTTTTTTTTTTAAAAATACGGGGTCAAATATACC 21674

```
TeratornLC199500  CC----------------------------------------------------CCCCCCTTCCCCCTGGTGGTCGTGCGGTGTAAGGAAAATGAAGATAGGAAA---AAAAAGCATAGTCTCG   15618
WF-Teratorn       CCAAACTACAGCGCGAGGGGTATGACAAAAAATTTAAAAAAAAATATACCCCCCCCTTCCCCCTGGTGGTCGTGCGGTGTAAGGAAAATGAAAATAGGAAAATAAAAAAGCATAGTCTAG   21794

TeratornLC199500  TGGGGTCATTTGCACTCTGTAAAACAGCGGAGTGAACGCTTTTGTTTTTGGAGAAAAAAATAATAAAGTATTGTCTCTTGGGGTCATTTGAGCTCCGTAAAACAGCGGAGTGAACGCTTT   15738
WF-Teratorn       TGGGGTCATTTGCACTCCGTAAAACAGCGGAGTGAACGCTTTTGTTTTTGGAGTAAAAAATAATAAAGTATTGTCTCTTGGGGTCATTTGAGCTCCGTAAAACAGCGTATTGAACGCTTT   21914

TeratornLC199500  TGTTTTTGGAGAAAAAAAAAAAAATAAAGTATTGTCTCTTGGGGTCATTTTAGCTCCGTAAAACAGCGGAGTGAACGCTTTTGTTTTTGGAGGAAAATAAAAATAAAGTATTGTCTCTTGG   15858
WF-Teratorn       TGTTTTTGGAGAAAAAAATAAAATACTGTATAGTCTCTTGGGGTCATTTTAGCTCCGTAAAACAGCGGAGTGAACGCTTTTGTTTTTGGAGAAAAATAAAAATACTGTATTGTCTCTTGG   22034

TeratornLC199500  GGTCATTTTAGCCCCGTAAAACAGCGGAGTGAACGCTTTTGTT--TGGAGAAAA--AAAAATAAAGTATTGTCTCTTGGGGTCATTTGAGCTCCATAAAACAGCGGAGTGAACGCTTTTT   15974
WF-Teratorn       GGTCATTTGCACTCCGTAAAACAGCGGAGTGAACGCTTTTGTTTTTGGAGAAAAATAAAAATAAAGTATTGTCTCTTGGGGTCATTTTAGCTCCGTAAAACAGCGGAGTGAACGCTTTTT   22154

TeratornLC199500  TTTTAAATAATACAGGGGTCGAATGCACGCGAAAGCACAGCGCGAGGGGTATGTAAATAAATAAATGAATTGATGCCC-------CCCCTTCCCCTCCCCCTGGTGGTCGTGCGCTGTAA   16087
WF-Teratorn       TTTTAAATAATACAGGGGTCGAATGCACGCGAAAGCACAGCGCGAGGGGTATGTAAATAAATAAATGAATTGATGCCCCCCCCCCCCCCTTCCCCTCCCCCTGGTGGTCGTGCGCTGTAA   22274

TeratornLC199500  GGAAA-TGAATAGTGGGTGAAAAAAAAAAAGCATTGTCTCGTGGGGTCATTTGCACTCCGTAAAACAGCGGAGTGAACGCTTTTTT-TGGAGAAAAAAAAAAATAAAGTATTGTCTCTTGG   16205
WF-Teratorn       GGAAAATGAAGAGTGTATGAAAAAAAAA--GCATTGTCTCGTGGGGTCATTTGCACTCCGTAAAACAGCGGAGTGAACGCTTTTTTCTGGAGAAAAAAAAAA----GTATTGTCTCTTGG   22388

TeratornLC199500  GGTCATTTGAGCTCCGTAAAACAGCGGAGTGAACG--CTTTTTTTTTTTAATAATACGGGGTCGAATGCACGCCAAAGCACAGCGCGAGGGGTATAAAAATAAATTGATA---------   16314
WF-Teratorn       GGTCATTTGAGCTCCGTAAAACAGCGGAGTGAATGGTTTTTTTTTTTTTAATAATACGGGGTCGAATGCAAGCGAAAGCACAGCGCGAGGGGTATGAAAATAAATTAAAAAAAAAAAAT   22508

TeratornLC199500  -----CCCTCCTCCCTCCCTCTCCACCCCCTG---GTCTTGGGGTGTAAAGAAAATGAAGAGAGGATGGAGAGAGAAAAAAAA--TAAAGTATTGTCTCGTGGGGTCACTTATGCCCTGT   16424
WF-Teratorn       TATACCCCCCCTCCTTCCCTCTCCACCCTCTGGTGGTCTTGGGGTGTAAGGAAAATGAAGAGAGGATGGAGAGAGAAAAAAAAAATAAAGTATTGTCTCGTGGGGTCACTTCTGCCCTGT   22628

TeratornLC199500  AAAACAGCGGAGGGAATGGGTTTCGTGCTGGCCATCGAGAAATGCCTTGTGGGGTCACTTATGCCCTGTAAAACAGCGGAGAGGATGGAGAGAGAAAAAAAAATAAAGTATTGTCTCGTG   16544
WF-Teratorn       AAAACAGCGGAGGGAATGG--------------------------------------------------------------------------AGAGAAAAAAAAATAAAGTATTGTCTCGTG   22677

TeratornLC199500  GGGTCACTTATGCCCTGTAAAACAGCGGAGGGAATGGGTTTCGTGCTGGCCACCGAGAAATGCCTCGTGGGGTCACTTATGCCCTGTAAAACAGCGGAGGGAGTTATATTTTTTTGGAGG   16664
WF-Teratorn       GGGTCACTTATGCCCTGTAAAACAGCGGAGGGAATGGGTTTCGTGCTGGCCACCGAGAAATGCCTCGTGGGGTCACTTATGCCCTGTAAAACAGCGGAGGGAGTTATATTTTTTTGGAGG   22797

TeratornLC199500  GGGGTCATGCAGACCCCAAAGCACATGCCAACCTAAGCGAGCGGAAGGGTTATATTGACAAACCAATACCTTTTTACATGCTTATCACAGGTAAGATTGATAACCCTTGTGCTGTCTTAG   16784
WF-Teratorn       GGGGTCATGCAGACCCCAAAGCACATGCCAACCTAAGCGAGCGGAAGGGTTATATTGACAAACCAATACCTTTTTACATGCTTATCACAGGTAAGATTGATAACCCTTGTGCTGTCTTAG   22917

TeratornLC199500  GGGGAAAGAAAAGCATGGATTAAAAAAAAAGGTAAGGAGGGGGGGTCATTTTAAAAACGCATACCCCCCTTTGGTATTGACCCCAGACAGGTACACCCCGGTCTGGCCTCCGGGAATCGAA   16904
WF-Teratorn       GGGGAAAGAAAAGCATGGATTAAAAAAAAAGGTAAGGAGGGGGGGTCATTTTAAAAACGCATACCCCCCTTTGGTATTGACCCCAAACAGGTACACCC-----------------------   23014

TeratornLC199500  GAGGGTGAAGTTCCTGAATTTATTTTCACTGTGGATTTACTCAGTCAAATTCAGTGGGGAAATCCAGCTTCTTTCAGTCAAAGCAGCTGGTTCAAGTCAAATTGTTTTAATCAATAAAGC   17024
WF-Teratorn       ------------------------------------------------------------------------------------------------------------------------   23014

TeratornLC199500  ACTTTGACACAGTAATCTGCGCCTTTGTTTCTATTCGACTGGATTACTGTAACTCTCTTTCCATCTCTCACTCGTCCTTATCCTTTCTCAAACTGGCAAAGATCGCTGCTGTATGTTTGT   17144
WF-Teratorn       ------------------------------------------------------------------------------------------------------------------------   23014

TeratornLC199500  TAACTGGCATGTGGAACAGAGAGCATATTACTCAAGTTCTGACTTCTCTACACTGGCTGACTGTTCATTTTAAAATTCTTTTATTTGTTTCTAAATCTTTAAACAGTCTGGCTCTGCCCT   17264
WF-Teratorn       ------------------------------------------------------------------------------------------------------------------------   23014

TeratornLC199500  GTCTCTCTGAGCTGCTACAGCTGCATTCCTCTTAGATCAGCTGACCAGCTGCTGCTGGATGTGCCAAGGACACAGGGGAAACTCAGAGTGTATGGGGCATTTTCCTTTTCTGGCCTCGGT   17384
WF-Teratorn       ------------------------------------------------------------------------------------------------------------------------   23014

TeratornLC199500  CTGGCCTCCGGGAATCGAAGAGGGTGAAGTTCCTGAATTTATTTTCACTGTGGATTTACTCAGTCAAATTCAGTGGGGAAATCCAGCTTCTTTCAGTCAAAGCAGCTGGTTCAAGTCAAA   17504
WF-Teratorn       ------------------------------------------------------------------------------------------------------------------------   23014

TeratornLC199500  TTGTTTTAATCAATAAAGCACTTTGACACAGTAATCTGCGCCTTTGTTTCTATTCGACTGGATTACTGTAACTCTCTTTCCATCTCTCACTCGTCCTTATCCTGTCTCAAACTGGCAAAG   17624
WF-Teratorn       ------------------------------------------------------------------------------------------------------------------------   23014

TeratornLC199500  ATCGCTGCTGTACGTTTGTTAACTTGCATGTGGAACAGAGAGCATATTACTCAAGTTCTGACTTCTCTACACTGGCTGACTGTTCATTTTAAAATTCTTTTATTTGTTTCTAAATCTTTA   17744
WF-Teratorn       ------------------------------------------------------------------------------------------------------------------------   23014
```

| | | |
|---|---|---|
| TeratornLC199500 | AACAGTCTGGCTCTGCCCTGTCTCTCTGAGCTGCTACAGCTGCATTCCTCTTAGATCAGCTGACCAGCTGCTGCTGGATGTGCCAAGGACACAGGGGAAACTCAGAGTGTATGGGGCATT | 17864 |
| WF-Teratorn | ------------------------------------------------------------------------------------------------------------------------ | 23014 |
| TeratornLC199500 | TTCCTTTTCTGGCCTCGGTCTGGCCTCCGGGAATCGAAGAGGGTGAAGTTCCTGAATTTATTTTCACTGTGGATTTACTCAGTCAAATTCAGTGGGGAAATCCAGCTTCTTTCAGTCAAA | 17984 |
| WF-Teratorn | ------------------------------------------------------------------------------------------------------------------------ | 23014 |
| TeratornLC199500 | GCAGCTGGTTCAAGTCAAATTGTTTTAATCAATAAAGCACTTTGACACAGTAATCTGCGCCTTTGTTTCTATTCGACTGGATTACTGTAACTCTCTGTCCATCTCTCACTCGTCCTTATC | 18104 |
| WF-Teratorn | ------------------------------------------------------------------------------------------------------------------------ | 23014 |
| TeratornLC199500 | CTGTCTCAAACTGGCAAAGATCGCTGCTGTACGTTTGTTAACTTGCATGTGGAACAGAGAGCATATTACTCAAGTTCTGACTTCTCTACACTGGCTGACTGTTCATTTTAAAATTCTTTT | 18224 |
| WF-Teratorn | ------------------------------------------------------------------------------------------------------------------------ | 23014 |
| TeratornLC199500 | ATTTGTTTCTAAATCTTTAAACAGTCTGGCTCTGCCCTGTCTCTCTGAGCTGCTACAGCTGCATTCCTCTTAGATCAGCTGACCAGCTGCTGCTGGATGTGCCAAGGACACAGGGGAAAC | 18344 |
| WF-Teratorn | ------------------------------------------------------------------------------------------------------------------------ | 23014 |
| TeratornLC199500 | TCAGAGTGTATGGGGCATTTTCCTTTTCTGGCCTCGGTCTGGCCTCCGGGAATCGAAGAGGGTGAAGTTCCTGAATTTATTTTCACTGTGGATTTACTCAGTCAAATTCAGTGGGGAAAT | 18464 |
| WF-Teratorn | -----------------------------------GGTCTGGCCTCCAGCAATCGAAGAGGGTCAAGTTCCTGAATTTATTTTCACTGTGGATTTACTCAGTCAAATTCAGTGGGGAAAT | 23099 |
| TeratornLC199500 | CCAGCTTCTTTCAGTCAAAGCAGCTGGTTAAAGTCAAATTGTTTTAATCAATAAAGCACTTTGACACAGTAATCTGCGCCTTTGTTTCTATTCGACTGGATTACTGTAACTCTCTTTCCA | 18584 |
| WF-Teratorn | CCAGCTTCTTTCAGTCAAAGCAGCTGGTTAAAGTCAAATTGTTTTTATCAATAAAGCACTTTGACACAGTAATCCGCGCCTTTATTTCTATTCGACTGGATCAGTGTAACTCTCTTTCCA | 23219 |
| TeratornLC199500 | TCTCTCACTCGTCCTTATCCTGTCTCAAACTGGCAAAGATCGCTGCTGTACGTTTGTTAACTTGCATGTGGAACAGAGAGCATATTACTCAAGTTCTGACTTCTCTACACTGGCTGACTG | 18704 |
| WF-Teratorn | TCTCTCACTCGTCCTTATCCTGTCTCAAACTGGCAAAGATCGCTGCTGTATGTTTGTTAACTGGCATGTGGAACAGAGAGCATATTACTCAAGTTCTGACTTCTCTACACTGGCTGACTG | 23339 |
| TeratornLC199500 | TTCATTTTAAAATTCTTTTATTTGTTTCTAAATCTTTAAACAGTCTGGCTCTGCCCTGTCTCTCTGAGCTGCTACAGCTGCATTCCTCTTAGATCAGCTGACCAGCTGCTGCTGGATGTG | 18824 |
| WF-Teratorn | TTCATTTTAAAATTCTTTTATTTGTTTCTAAATCTTTAAACAGTCTGGCTCTGCCCTGTCTCTCTGAGCTGCTACAGCTGCATTCCTCTTAGATCAGCTGACCAGCTGCTGCTGGATGTG | 23459 |
| TeratornLC199500 | CCAAGGACACAGGGGAAACTCAGAGTGTATGGGACATTTTCCTTTGCTGGCCTCAGCTTGTGAGATGCACTGCTCATGCAGGTTAGGCAGGCCCATTCTTCGAATGATTTTAAATCCAAG | 18944 |
| WF-Teratorn | CCAAGGACACAGGGAAAACTCAGAGTGTATGGGGCATTTTCCTTTGCTGGCCTCAGCTTGTGGGATGCACTGCTCATGCAGGTTAGGCAGGTCAATTCTTCGAATGATTTTAAATCCAAG | 23579 |
| TeratornLC199500 | TTGAAAACTCAGTTTTTCTCACTGGCTTTTAACTTAGTTTGAGTTGGTGATTGTACTGATTTTCATTACTTTTGTATTATGTTGTTCATTTTGGATCGTTGTCTTGTTTTATCTTTCATC | 19064 |
| WF-Teratorn | TTGAAAACTCAGTTTTTCTCACTGGCTTTTAACTTAGTTTGAGTTGGTGATTGTACTGATTTTTATTACTTTTGTATTATGTTGTTCATTTTGGATCGTTGTCTTGTTTTATCTTTCATC | 23699 |
| TeratornLC199500 | ATTAAGCACTTTGGTCAGCCTCTGCTGTCACAAAAGCGCTTTATAGATAAAGTTTACTTGACTCATGCCTTGCTGGAGTGAAAACATAGTTGAAAGAAAGAAACAAAAAACAACAAACAA | 19184 |
| WF-Teratorn | ATTAAGCACTTTGGTCAGCCTCTGCTGTCACAAAAGCGCTTTATAAATAAAGTTTACTTGACTCATGCCTTGCTGGAGTGAAAACATAGTTGAAAGAAAGAAACAAAAA-CAACAAACAA | 23818 |
| TeratornLC199500 | AACACATCAACCCCGTCATTGTTTTTATTGAGTTTCACAGGCAAGTTTACTTTAGCACAAGCAACAGTAAAAA---AGCAAAAGCATTTTTAAAACTAAAAAAAA-----ATATTTCACA | 19296 |
| WF-Teratorn | AACACATCAACCCCGTCATTGTTTTTATTGAGTTTCACAGGCAAGTTTACTGTAGCACAAGCAACAGTAAAAAAAAAGCAAAAGCATTTTTAAAACTAAAAAAAAAAACTATATTTCACA | 23938 |
| TeratornLC199500 | CACTCAAAGCGTCTGCCAGCACAGAGTCCTTTCACTGCTAGCATATT**TCACAGGAACAGGCTGGACTGTGAATAGTAGTCGCCATCCCCGTATGTTTTGGCAGCATTAAGACCGTCTCGA** | 19416 |
| WF-Teratorn | CACTCAAAGCGTCTGCCAGCACAGAGTCCTTTCACTGCTAGCGCATTTCACAGGAACAGGCTGGACTGTGAATAGTAGTCGCCATCCCCGTATGTTTTGGCAGCATTTAGACCGTCTCGA | 24058 |
| TeratornLC199500 | **TGAGCGTTGTGCGACATGAAGGCGTCTGTATTTTTCCCGTCGCCTTGCCTCTCTTTGAGCACTGTGGAGGTGAGCACCATGGTGATCACGTACATGGCGTATATGAGGATGTTGAACACT** | 19536 |
| WF-Teratorn | TGAGCGTTGTGCGACATGAAGGCGTCTGTATTTCTCCCGTCGCCTTGCCTCTCTTTGAGCACTGTGGAGGTGAGCACCATGGTGATCACGTACATGGCGTATATGAGAATGTTGAACACT | 24178 |
| TeratornLC199500 | **GCTATGGCGCACTCCACGTTGTGCATCCTGTGTTCGTCCCGAATCAGCGGCCTGTACAAGAGCTCCATGCAACCAGAGACGACTGCCATGAACAGCTCCAGATTCATCATGATCAGACAG** | 19656 |
| WF-Teratorn | GCTATGGCGCACTCCACGTTGTGCATCCTGTGTTCGTCACGAATCAGCGGCCTGTACAAGAGCTCCATGCAACCAGAGACGACTGCCATGAACAGCTCCAGATTCATCATGATCAGACAG | 24298 |
| TeratornLC199500 | **GCCAGCACCGTCCGGGTGCCTTTGTTGAGCCTCCTGGTTCCGTCCTCGGCGTCGTCCATCTTGCGCTTGCTCTTGGCCGCGAAAAAGCTCAGCACCACGCTCACGACCATCGCCAGACAG** | 19776 |
| WF-Teratorn | GCCAGCACCGTCCGGGTGCTTTTGTTGAGCCTTCTGGTTCCTTCCTCGGCGTCGTCCATCTTGCGCTTGCTCTTGGCCGCGAAAAAGCTCAGCACCACGCTCACGACCATCGCCAGACAG | 24418 |
| TeratornLC199500 | **CCCATGGTTATCACAGCTATGCCGCCGTGTATGTAAGCGTGATCGTAGCTACTCCTCTCCTGAGCCGTAGAGGAGCCGAAGGAGACGTCCCGCGCGCTCCCCCTCACGGCGTGACTGCTA** | 19896 |
| WF-Teratorn | CCCATGGTTATCACAGCTATGCCGCCGTGTATGTAAGCGTGATCGTAGCTACTCCTCTCCTGAGCCGTAGAGGAGCCGAAGGAGACGTCCCGAGCGCTCCCCCTCACGGCGTGACTGCTA | 24538 |
| TeratornLC199500 | **TCTATCAGGTAGAGGATGTTGTTGGGCAAGGCGAGGAACCCGACAATGAGAGAGGTGATGACCGCGATGGCGGCCCCCAACTTTCCCCTGAACCTGGCAACGCTCAGTACGGGGTTGGAC** | 20016 |
| WF-Teratorn | TCTATCAGGTAGAGGATGTTGTTGGGCAAGGCGAGGAACCCGACAATGAGAGAGGTGATGACCGCGATGGCGGCCCCCAACTTTCCCCTGAACCTGGCAACGCTCAGTACGGGGTTGGAC | 24658 |

| | | |
|---|---|---|
| TeratornLC199500 | **TTCCTCCCCATCAACATCTCTTTTCCGGACGAAATCAAAGAATCGAAGCCTCCGACGAGGCCGGACCTGGTCCGCTGCAT**CCCGCCCAGCAGAGGTGGGCAGACGGGGTCGCGTCCTGCG | 20136 |
| WF-Teratorn | TTCCTCCCCATCAACATCTCTTTTCCGGACGAAATCAAAGAATCGAAGCCTCCGACGAGGCCGGACCTGGTCCGCTGCATCCCGCCCAGCAGAGGTGGGCAGACGGGGTCGCGTCCTGCG | 24778 |
| TeratornLC199500 | CACGAGAGGGCCGAGCCAGCATAGGTTACCTACCCCCCTTTAACGGTTACTGCCGCACAGCGTCTTGGTCCACCCCGAGACATTTAATACG**ATGAAGCTCGGGAGGCCCGTGATCGAGTT** | 20256 |
| WF-Teratorn | CACGAGAGGGCCGAGTCAGCATAGGTTACCAACCCCCCTTTAACGGTTACTGCCGCACAGCGTCTTGGTCCACCCCGAGACATTTAATACGATGAAGCTCGGGAGGCCCGTGATCGAGTT | 24898 |
| TeratornLC199500 | **CAGAGGGTCGAGAAGAGACCTCCAGGCGAAGGCCGTGACGCAGGACCGGTGGTACAGGGTCCGCGGCATCACGAGGACCTTGAATTCGATCGCGCGCCGCAGGAGAGGCGTCCGATCTCA** | 20376 |
| WF-Teratorn | CAGAGGGTCGAGAAGAGACCTCCAGGCGAAGGCTGTGACGCAGGACTGGTGGTACAGGGTCCGCGGCATCACGAGGACCTTGGATTCGATCGCGCGCCGCAGGAGAGGCGTCCGATCTCA | 25018 |
| TeratornLC199500 | **CAACATGATATGTGTGGACCTATACGCTAGGTTCCTTAGCCACCTCGTCCGTGACCGACGAAACCCGCTCGGCTGGTTGTCTCATCACTTGGCCATCCTGAACGCGCCTGAGGACCCAGC** | 20496 |
| WF-Teratorn | CAACATGATATGTGTGGACCTATACGCTAGGTTCCTTAGCCACCTCGTCCGTGACCGACGAAACCCGCTCGGCTGGTTGTCTCATCACTTGGCCATCCTGAACGCGCCTGAGGACCCAGC | 25138 |
| TeratornLC199500 | **CCGTAAGTTTGAGGGCGAGTTGAGACGCCTGCTGGCATACGGCTCTCTCAACTTTGAGTCGTGCAGGCGCGCCTGGGACAGGAGCCGCATGTCCAGGGGCATAGGAAACGCAGTAGACGC** | 20616 |
| WF-Teratorn | CCGTAAGTTTGAGGGCGAGTTGAGACGCCTGCTGGCACACGGCTCTCTCAACTTTGAGTCGTGCAGGCGCGCCTGGGACAGGAGCCGCATGTCCAGGGGCATAGGAAACGCAGTAGACGC | 25258 |
| TeratornLC199500 | **TCAGTGCAAGGACGTGTTCAACGGAAAGAAGGCTGCTTCAGAGACCGGGCCCGTGTCTCGCCTCCTGCTCGCCCAGCTCGAGGCCCTGCGCCTCAAGCCGATCTGCGCGGGCCTCAAAGT** | 20736 |
| WF-Teratorn | TCAGTGCAAGGACGTGTTCAACGGAAAGAAGGCTGCTTCAGAGACCGGGCCCGTGTCTCGCCTCCTGCTCGCCCAGCTCGAGGCCCTGCGCCTCAAGCCGATCTGCGCGGGCCTCAAAGT | 25378 |
| TeratornLC199500 | **GGCCGGCGTGCCCACGCCCGACCGAGGCGGCAGCACGTACCACAACGGCTGCACCCGGGGCTGCGGGCCACTGCCTGGACGGAACACCTTCTACACGGAAGTGGACCTGCTGGCTCACGA** | 20856 |
| WF-Teratorn | GGCCGGCGTGCCCACGCCCGACCGAGGCGGCAGCACGTACCACAACGGCTGCACCCGGGGCTGCGGGCCACTGCCTGGACGGAACACCTTCTACACGGAAGTGGACCTGCTGGCTCACGA | 25498 |
| TeratornLC199500 | **TGCAACGACCGACACTGTGGCCCTCCTAGAGCTCAAGACCCGAAACAGCGACGTTTTGGATGCCCCTACCCTGTGGCGGTACAACGCCCAGCTGTGGCTGACCTGGACGATGTTCTCGCT** | 20976 |
| WF-Teratorn | TGCAACGACCGACACCGTGGCCCTCCTAGAGCTCAAGACCCGAAACAGCGACGTTTTGGATGCCCCTACCCTGTGGCGGTACAACGCCCAGCTGTGGCTGACCTGGACGATGTTCTCGCT | 25618 |
| TeratornLC199500 | **CACGTACCCGAGCGTGGCGGAGCGCTCAAACGCTTACCTGGTGATCGTGAGGCCAGGCACGAACACGGTCACCATAAGGAGCTGTCTGAGGCCCTCCGTGACCAAGACGCTGCGCACCAA** | 21096 |
| WF-Teratorn | CACGTACCCGAGCGTGGCGGAGCGCTCAAACGCTTACCTGGTGATCGTGAGGCCAGGCACGAACACAGTCACCATAAGGAGCTGTCTGAGGCCCTCCGTGACCAAGACGCTGCGCACCAA | 25738 |
| TeratornLC199500 | **GTTCCCCTGGATAAACTGCCTTTGTCCGCAGGTGCTCAACTGCCTCACGCCGACCTGTGTCAACGTGAGAGTGCAGGCCTATCCGAAGCCAGGCTCTGGCGGCAGGGCGAGCATCGACCC** | 21216 |
| WF-Teratorn | GTTCCCCTGGATAAACTGCCTTTGTCCGCAGGTGCTCAACTGCCTCACGCCGACCTGTGTCAACGTGAGAGTGCAGGCCTATCCGAAGCCAGGCTCTGGCGGCAGGGCGAGCATCGACCC | 25858 |
| TeratornLC199500 | **AAAGGACCTCTGTTACAGGAACGTCATGTTCAACCGAGAAAAGGCCAGAGTGCGTCAGAGACGTTACCCCGATCAGGGATCGCAAGGCCGTCCACCGCACCCAACGCCCTCCT---CCGA** | 21333 |
| WF-Teratorn | GAACGACCTCTGTTACAGGAACGTCATGTTCAACCGAGAAAAGGCCAGAGTGCGGCAGAGACGTTACCCCGATCAGGGATCGCAAGGCCGTCCACTGCACCCAACGCCCTCCTCCTCCGA | 25978 |
| TeratornLC199500 | **GACGAGCGGGGCCCCCCAGGGGTCCTGGGTCCGCATCACGTGGGAGCCGAACACGGCTCCCACGCAGGCATGA**CACTCGATAAAAAGCCCCGCATGTTCGAGCGCTGGATGGAAACTGAC | 21453 |
| WF-Teratorn | GACGAGCGGGGCCCCC-AGGGGTCCTGGGTCCGCATCACGTCGGAGCCAAACACGGCTCCCGCGCAGGCATGACACTCGGTGAAAAGCACCGCGTGTTCGAGCGCTGGATGGAAACTGAC | 26097 |
| TeratornLC199500 | GGACAACTAGAACTGAC**ATGGGTCTGAGGCACCAAGCGCTCCACGGCATGCTGGCCAACCCAGACGTGTCCCCTTTCAAAGTAATCGAGAAACCCAGCGAACAGCAGGACTGGATCCTGA** | 21573 |
| WF-Teratorn | GGACAACTAGAACTGACATGGGTCTGAGGCACCAAGCGCTCCACGGCATGCTGGCCAACCCAGACGTGTCCCCTTTCGAAGTAATCGAGAAACCCAGCGAACAGCAGGACTGGATCCTGA | 26217 |
| TeratornLC199500 | **TAGACCATCTCATTGTCACACACACAGTCAGAGAAAGCGATTACAGGGAGAGCGAACCGGAGATGTCGGTCGCGAGGATGGCGAGCTATTTCGCATCCTGGTACGTCAGAGAGAGGGCCC** | 21693 |
| WF-Teratorn | TAGACCATCTCATGGTCATACACACAGTCAGAGAAAGCGATTACAGGGAGAGCGAACCGGAGATGTCGGTCGCGAGGATGGCGAGCTATTTCGCATCCTGGTACGTCAGAGAGAGGGCCC | 26337 |
| TeratornLC199500 | **ACGGCCGATGCGTCGTGCTCGTAAAAGACGGTCCGGGGGCGCAGAAGCCTAGCGTGAGGGAAGAAAGGGCCAACAGGACGGTCCCTGGTCACTTGAAGTTCTGTAAGGCTCACCTGGCCG** | 21813 |
| WF-Teratorn | ACGGCCGATGCGTCGTGCTCGTAAAAGACGGTCCGGGGGCGCAGAAGCCTAGCGTGAGGGAAGAAAGGGCCAACAGGACGGTCCTTGGTCACTTGAAGTTCTGTAAGGCTCACCTGGCCG | 26457 |
| TeratornLC199500 | **AGATAGAGTACGGCACGATGAAACTGTTGGCCGACGTGGGCATGAGAGACAGCTTTTTCTTAGTCACAGGGGCGGTAGGCGAGTCGCGCTGGGCCCAGACCGCCGGCGAGCGCGCCAGCA** | 21933 |
| WF-Teratorn | AGATAGAGTACGGCACGATGAAACTGTTGGCCGACGTGGGCATGAGAGACAGCTTTTTCTTAGTCACGGGGGCGGTAGGCGAGTCGCGCTGGGCCGAGACCGCCGGCGAGCGTGCCAGCA | 26577 |
| TeratornLC199500 | **ACCTGGTCTCGTTCCACAACG---------AGCGCGGGGGAGAGGGGGCCAGTCCACTCACGGAGCCAGACCACTTCAAAATCGAGACCCTCCATTTTCCCGTCGTGGATCTGAGCTACT** | 22044 |
| WF-Teratorn | ACCTGGTCTCGTTCCACAACGAGCGCGAGGAGCGCGGGGGAGAGGGGGCCAGTCCACTCACGGAGCCAGACCACTTCAAAATCGAGACCCTCCATTTTCCCGTCGTGGATCTGAGCTACT | 26697 |
| TeratornLC199500 | **GCGACGGCAAGACCTACAGGCCTGAGGGGACGCCTGGAGTTTTCTTGACAGCTTTCGGGAGCCCGGAAGACGAGCGGCTCCAGGGCTTGCTAAAGGCTTGCGTCGAGTGTCGAGAAACGG** | 22164 |
| WF-Teratorn | GCGACGGCAAGACCTACAGGCCTGAGGGGACGCCTGGAGTTTTCTTGACAGCTTTCGGGAGCCCGGAAGACGAGCGGCTCCAGGGCTTGCTAAAGGCTTGCGCAGAGTGTCGAGAAACGG | 26817 |
| TeratornLC199500 | **AGGCCGACACGCTGATGGTGGAGTTGGCCAACGCCCTAACGGGCTCAGTGACAGTGTGCACAGGGGACACAGATCTGATAGCCGTGCTCGCTGCGAGCGGACGGCCCGGCCTGACACTGA** | 22284 |
| WF-Teratorn | AGGCCGACACGCTGATGGTGGAGTTGGCCAACGCCCTGACGGGCTCAGTGACAGTGTGCACAGGGGACACAGATCTGATAGCCGTGCTCGCTGCGAGCGGACGGCCCGGCCTGACACTGA | 26937 |

| | | |
|---|---|---|
| TeratornLC199500 | **GGTTGGACAACAAAAGCTACACTGAGGACAGAGCCATGCACGAATCGCCCTTCGGCGAGCTCCTTGTCGAGCCAGCGGTCCGACGGACCGCCAACAGAGCGCCGGGTGCCACAGACGACT** | 22404 |
| WF-Teratorn | GGTTGGACAACAAAAGCTACACTGAGGACAGAGCCATGCACGAATCGCCCTTCGGCGAGCTCCTTGTCGAGCCAGCGGTCCGACGGACCGCCAACAGAGCGCCGGGTGCCACAGACGACT | 27057 |
| TeratornLC199500 | **GGCTGAGCGCACTGTGCGACCTGGGCTCCGCAGAGCACCACCTGTTGCCCAACGGCGCGGCGGAAAGACACGCCGATCTGGGTCGCCTTCTGGAGAGCCACGGGGGCGAGGCGTTCAAAC** | 22524 |
| WF-Teratorn | GGCTGAGCGCACTGTGCGACCTGGGCTCCGCAGAGCACCACCTGTTGCCCAACGGCGCGGCGGAAAGACACGCCGATCTGGGTCGCCTTCTGGAGAGCCACGGGGGCGAGGCGTTCAAAC | 27177 |
| TeratornLC199500 | **TCTCAGTGTACGAGTTCTTGTACCGGGGCGGTGTTCGAGGCTCGGTTTACGTCGCTTTCCTGAGCGCGTTCTTCGAGTCTGACGCGAGGAGCAAACTGAGTGAACTGGCACTGGAGAGCC** | 22644 |
| WF-Teratorn | TCTCAGTGTACGAGTTCTTGTACCGGGGCGGTGTTCGAGGCTCGGTTTACGTCGCTTTCCTGAGCGCGTTCTTCGAGTCTGACGCGAGGAGCAAACTGAGTGAACTGGCACTGGAGAGCC | 27297 |
| TeratornLC199500 | **AGAAAACGTACCCCCCCGGCACACGCAGCGCAAACACGTTACGCTTCGCCTTCACGGTCCTGTGCGGCGCGGATGACCCGAAAGTCTTAGGCGCGGGGAACGCGCCCTCGTCGCAACTTT** | 22764 |
| WF-Teratorn | AGAAAACGTACCCCCCCGGCACACGCAGCGCAAACACGTTACGCTTCGCCTTCACGGTCCTGTGCGGCGCGGATGACCCGAAAGTCTTAGGCGCGGGGAACGCGCCCTCGTCGCAACTTT | 27417 |
| TeratornLC199500 | **TCGGTAACGCAGCACGCTCGGTCGAGCGGGGATCTAAGCGCCCCCGCGAGCCAGAGGTCGACCGGGAGGGCGAAGCAGAACCAGAGATCGGATCCGACCCCCAGAAGACCAAGGTAAAGA** | 22884 |
| WF-Teratorn | TCGGTAACGCAGCACGCTCGTTCGAGCGGGGATCTAAGCGCCCCCGCGAGCCAGAGGTCGACCGGGAGGGCGAAGCAGAACCAGAGATCGGATCCGACCCCCAGAAGACCAAGGTAAAGA | 27537 |
| TeratornLC199500 | **TCACGGCTGGGCTCAAGCGCACGGCCAAAGCCTACGCGGACGGAGCAATCCCGGAGTCCACTCACGCCCGTTACCTCAGACTGGCTCGCTCCGGAAGGCACCTGTTCCTGAGAGCCAAAG** | 23004 |
| WF-Teratorn | TCACGGCTGGGCTCAAGCGCACGGCCAAAGCCTACGCGGACGGAGCAATCCCGGAGTCCACTCACGCCCGTTACCTCAGACTGGCTCGCTCCGGAAGGCACCTGTTCCTGAGAGCCAAAG | 27657 |
| TeratornLC199500 | **CCAGAGCGCAGGAGGACCAGAGCGTCCGATCCAAACACTTGCTGTTCATGGCCCTCTGCGGAACAGACTACAGCTTCACTCCAGTGGGGCTGGGAATCAAAAGGTTGCTGAGTGGCGCGA** | 23124 |
| WF-Teratorn | CCAGAGCGCAGGAGGACCAGAGCGTCCGATCCAAACACTTGCTGTTCATGGCCCTCTGCGGAACAGACTACAGCTTCACTCCAGTGGGGCTGGGAATCAAAAGGTTGCTGAGTGGCGCGA | 27777 |
| TeratornLC199500 | **TACGCAACCGAGAGAAGTTCAGCTCTTGGTGCGCCAAGCTCGTGACCGCGCTGTGGTCCCAGGACGGAGAGGATGGCGGAGGGAGCGAAAGGTCCCTTGAGTTTCGCGCCATGGGCGAAC** | 23244 |
| WF-Teratorn | TACGCAACCGAGAGAAGTTCAGCTCTTGGTGCGCCAAGCTCGTGACCGCGCTGTGGTCCCGGGACGGAGAGGATGGCGGAGGGAGCGAAAGGTCCCTTGAGTTTCGCGCCATGGGCGAAC | 27897 |
| TeratornLC199500 | **AGTTGGCCAGCCTCACAGGAGTCCCCAAGTCTACGAGCTCAAAGCATTGGACTGGAAAAAAGTTTGAAATCATGTGCAGGACGCTCAAATACGTGTGCGACCTGTGGACCCTGAAGGGTC** | 23364 |
| WF-Teratorn | AGTTGGCCAGCCTCACAGGAGTCCCCAAGTCTACGAGCTCAAAGCATTGGACTGGAAAAAAGTTTGAAATCATGTGCAGGACGCTCAAATACGTGTGCGACCTGTGGACCCTGAAGGGTC | 28017 |
| TeratornLC199500 | **CCCGGATCGGTCCCCATTATGGGTTTCGTGTGCACGACGGAGTCGTGCGTTTCATAGAAGAGGAAGACCCAGAGTCTGTGTCTTGA**GAGTTGATACAGGGGCGAGATGTATTTCATTTCA | 23484 |
| WF-Teratorn | CCCGGATCGGTCCCCATTATGGGTTTCATGTGCACGACGGAGTCGTGCGTTTCATAGAAGAGGAAGACCCAGAGTCTGTGTCTTGAGAGTTGATACAGGGGCGAGATGTATTTCATTTCA | 28137 |
| TeratornLC199500 | TTTTCTGTGTAAAAAATGAAGAAGAAAAAAAAAAAAACGAGCGAACAAACACAATAAGAGGATGACCCAGAGTCTGTGTCTTGAAAGTTGATACAGGGGCGAGATGTATTTCATTTCAAT | 23604 |
| WF-Teratorn | TTTTCTGTGTAAAAAATGAAGAAGAAAAAAAAAAA--CGAGCGAACAAACACAATAAGAGGATGACCCAGAGTCTGTGTCTTGAAAGTTGATACAGGGGCGAGATGTATTTCATTTCAAT | 28255 |
| TeratornLC199500 | TCCAATTTCTGTGTAAAAAATGAAGAAGAAAAAAAAAAATAAGCGAACAAACACGATGGAAACAATAAAAGTAAATTCAAACACACCAGTGCAGTTTTCAGAGTTTTATTTCATTCGGAC | 23724 |
| WF-Teratorn | TCCAATTTCTGTGTAAAAAATGAAGAAGAAAAAAAA---TAAGCGAACAAACACGATGGAAACAATAAAAGTAAATTCAAACACACCAGTGCAGTTTTCAGAGTTTTATTTCATTCGGGC | 28372 |
| TeratornLC199500 | ACGTCTGTATTTACACCACACAATGTCGATTTCAACTCTCGAAGCCATCTCTAAATTACCACTGGAGTTCGTCGTGTGTCAACCGAAAGGATGATTGGCGTACTCGAGCTTGCTCATCCA | 23844 |
| WF-Teratorn | ACGTCTGTATTTACACCACACAATGTCGATTTCAACTCTCGAAGCCATCTCTAAATTACCGCTGGAGTTCGTCGTGTGTCAACCTAAAGGATGATTGGCGTACTCGAGCTTGCTCATCCA | 28492 |
| TeratornLC199500 | TGTACATTAAGGTCATAGGCGGCGCTAAGG**ATGAACAAAGGCCGCAAAAGAACACTAAGAAGCAGGTCGAGGCCCCTGGCCCACAAGCGGGGAACAAGAGCGGTCGAGCCAGAGGATCCT** | 23964 |
| WF-Teratorn | TGTACATTAAGGTCATAGGCGGCGCTAAGGATGAACAAAGGCCGCAAAAGAACACTAAGAAGCAGGTCGAGGCCCCTGGCCCACAAGCGGGGAACAAGAGCGGTCGAGCCAGAGGATCCT | 28612 |
| TeratornLC199500 | **CGGCCCGACTCGCAAGGGCAGCGCGGGGGGCCGAAGCCGGGCCGCTACGAACCCTACGACTGGGATTCGGACTCAGACGGGGGACACAGAGACGGAGGGGGTCACGGTCGAGTCGGCCGC** | 24084 |
| WF-Teratorn | CGGCCCGACTCGCAAGGGCAGCGCGGGGGGCCGAAGCCGGGCCGCTACGAACCCTACGACTGGGATTCGGACTCAGACGGGGGACACGGAGACGGAGGGGGTCACGGTCGAGTCGGCCGC | 28732 |
| TeratornLC199500 | **AAAACTCGCGCGGGCCGGCG---------------GGACAAAGCCCGTGAGCCTGATGCGGCAGCGGTGGAGCCCGTGGGTCCTCGGCACGACTCGGAAGGGCAGCGCGGGGCCCCGAAG** | 24189 |
| WF-Teratorn | AAAACTCGCGCGGGCCGGCGATCCCATCCGCCGGGGGACAAAGCCCGTGAACCTGATGCGGCAGCGGTGGAGCCCGTGGGTCCTCGGCACGACTCGGAAGGGCAGCGCGGGGCCCCGAAG | 28852 |
| TeratornLC199500 | **CCGGGCCGCTATGAACCCTACGACTGGGATTCGGACTCAGACGGGGGAGACGGAGGGGGCCACGGTCCGCCGCGGGACGAAGCCCTCGAGCCTGATGCGGCAGCGGTCGAGCCCGAGGGT** | 24309 |
| WF-Teratorn | CCGGGCCGCTATGAACCCTACGACTGGGATTCGGACTCAGACGGGGGAGACGGAGGGGGCCACGGTCCGCCGCGGGACGAAGCCCTCGAGCCTGATGCGGCAGCGGTCGAGCCCGAGGGT | 28972 |
| TeratornLC199500 | **CCTCGGCACGACTCGGAAGGGCAGGGCGGGGGAGACGGAGACAGAGGGGGCCACGGTCAGGTCGGCCGTGAAACTCTCGAGGACAGGCGATCCGATCCGCCGCGGGACGAAGCCCTCGAG** | 24429 |
| WF-Teratorn | CCTCGGCACGACTCGGAAGGGCAGGGCGGGGGAGACGGAGACAGAGGGGGCCACGGTCAGGTCGGCCGTGAAACTCTCGAGGACAGGCGATCCGATCCGCCGCGGGACGAAGCCCTCGAG | 29092 |
| TeratornLC199500 | **CCTGATGCGGCAGCGGTCGAGCCCGAGGGTCCTCGGCACGACTCGGAAGGGCAGGGCGGGGGAGACGGAGACAGAGGGGGCCACGGTCAGGTCGGCCGTGAAACTCTCGAGGACAGGCGA** | 24549 |
| WF-Teratorn | CCTGATGCGGCAGCGGTCGAGCCCG----------------------------------------------------------------------------------------------- | 29117 |

TeratornLC199500 **TCCGATCCGCCGCGGGACGAGGCCCACGAGCCTGACGATGGAGCGGTCGAGCCCGGGGGTCCTCAGCATGAATTGGAAGGGCAGGGCAGAGGCGACGAACCCTCCGATTGGGATTTGTAC** 24669
WF-Teratorn ---------------------------------------------------------GGGGTCCTCAGCATGAATTGGAAGGGCAGGGCAGAGGCGACGAACCCTCCGATTGGGATTTGTAC 29182

TeratornLC199500 **GAGTGTGAAACGGACGTGGACGAGCTCGATGGTTCTCAAAGTGAAACGGAAGAACAACACCGGGGCCCCAGGTCATCGGGCGAGGAAGACGTCGAGGAAGCCGTTGAGGAAGACGCCGAG** 24789
WF-Teratorn GAGTGTGAAACGGACGTGGACGAGCTCGATGGTTCTCAAAGTGAAACGGAAGAACAGCACCGGGGCCCCAGGTCATCGGGCGAGGAAGACGTCGAGGAAGCCGTTGAGGAAGACGCCGAG 29302

TeratornLC199500 **GAAGACATAGAGGAAGAACCAGCCGCAGAGAGCGCGAGCGTGTACCCGGATACATTTTCCTCCAAGAACGGTGAAATAAAGTGGTCAGCGAACGAGCGTGATCGGTCGAGCGTCCGGGCG** 24909
WF-Teratorn GAAGACATAGAGGAAGAACCAGCCGCAGAGAGCGCGAGCGTGTACCCGGATACATTTTCCTCCAAGAACGGTGAAATAAAGTGGTCAGCGAACGAGCGTGATCGGTCGAGCGTCCGGGCG 29422

TeratornLC199500 **TCCCCTCCTCACCAGGCCTCAGGGCCTACCACTGAGGCGAGGGAGGGGGCTCAAGACATCCTGTCTTCTTTTCTTTTGTTTTTCACTCCGAATGTAGAAAGACTGATTTTGATAGCTACC** 25029
WF-Teratorn TCCCCTCCTCACCAGGCCTCAGGGCCTACCACTGAGGCGAGGGAGGGGGCTCAAGACATCCTGTCTTCTTTTCTTTTGTTTTTCACTCCGAATGTAGAAAGACTGATTTTGATAGCTACC 29542

TeratornLC199500 **AACCGCCATAGCGCCGATAAGGCCCACGCCTTAAAGACTCCGAAACCCATGGACGAGATTGAACTGAGAGCCTACGTGGGACTGCTGATCTTGGCCGGGGTGTACAGGTCTCGAGGCGAA** 25149
WF-Teratorn AACCGCCATAGCGCCGATAAGGCCCACGCCTTAAAGACTCCGAAACCCATGGACGAGATTGAACTGAGAGCCTACGTGGGACTGCTGATCTTGGCCGGGGTGTACAGGTCTCGAGGCGAA 29662

TeratornLC199500 **GCGTTGGTGAGCCTGTGGGAGCCCGAGTGCGGACGACCGATCTTCGGGGCAACCATGTCCCTTAGAAGCTTTTACCAGCGCTCTGCGACGCTCAGGTTCGACGATCGAAGCACGAGAGCC** 25269
WF-Teratorn GCGTTGGTGAGCCTGTGGGAGCCTGAGTGCGGACGACCGATCTTCGGGGCAACCATGTCCCTTAGAAGCTTTTACCAGCGCTCTGCGACGCTCAGGTTCGACGATCGAAGCACGAGAGCC 29782

TeratornLC199500 **TCTAGGAGCGCTTCGGACAGATTGGCCGCAGTCAGAGAGGTGTGGGAAGAGTGGTGCGATCGGCTGCCCAGGCTTTACGACATCGGACCCGAGGTGACTGTGGACGAACGAATGGTTCCT** 25389
WF-Teratorn TCTAGGAGCGCTTCGGACAGATTGGCCGCAGTCAGAGAGGTGTGGGAAGAGTGGTGCGATCGGCTGCCCAGGCTTTACGACATCGGACCCGAGGTGACTGTGGACGAACGAATGGTTCCT 29902

TeratornLC199500 **TTCAAAG**GTGCGGTTTCAGCCTCCCTTTTTACTGTTCTCAGGGTATCCTTGACGTATTCGCGTCTGTCTTAACTCTTCCTTTTGTTTTTCTTTCTCTACGCAG**GACGTTGTTCTTTCAGA** 25509
WF-Teratorn TTCAAAGGTGCGGTTTCAGCCTCCCTTTTTACTTTTCTCAGGGTATCCTTGACGTA--CGCGTCT----TAACTCTTCCTTTTGTTTTTCTTTCTCTACGCAGGACGTTGTTCTTTCAGA 30016

TeratornLC199500 **CAGTACATTCCCAGCAAGCCTACCAAGTACGGCCTCAAACTATGGGTCGCTTGCGACGCAAGGACGAGCTACGCTTGGAGGGTCCAACCGTACATGGGCAAGCTGGCTTCGGGTGCTCGA** 25629
WF-Teratorn CAGTACATTCCCAGCAAGCCTACCAAGTACGGCCTCAAACTATGGGTCGCTTGCGACGCAAGGACGAGCTACGCTTGGAGGGTCCAACCGTACATGGGCAAGCTGGCTTCGGGTGCTCGA 30136

TeratornLC199500 **GAGACAAACCTTGCGTCCCGAGTGGTTTCGGATCTCACCCGGGGACTGGAGCACCGTAACGTCACGTGCGACAATTTCTTCACTTCGTACGCGCTCGCCACCGAGCTCCTGAAACGGAAC** 25749
WF-Teratorn GAGACAAACCTTGCGTCCCGAGTGGTTTCGGATCTCACCCGGGGACTGGAGCACCGTAACGTCACGTGCGACAATTTCTTCACTTCGTACGCGCTCGCCACCGAGCTCCTGAAACGGAAC 30256

TeratornLC199500 **ATCACCCTGCTCGGTACGATCAAAAGCAACAAACCCGAACTGCCTCCGGAGCTCGCGTCGGCCCGCAGACGAGAAGCACTCTCCTCCAGGTTCGCCTTTACGGACTGCGCCGCGGCCCTT** 25869
WF-Teratorn ATCACCCTGCTCGGTACGATCAAAAGCAACAAACCCGAACTGCCTCCGGAGCTCGCGTCGGCCCGCGGACGAGAAGCACTCTCCTCCAGGTTCGCCTTTACGGACTGCGCCGCGGCCCTT 30376

TeratornLC199500 **TCCTACGTGCCCAAAAAGAACAAAAACGTCCTGCTCCTTAGCACGAAGCATTCGAGGGTCGAAATATGCCACGAACGCAGGGACAAAAAACCAGAGATGATCTTGGATTACAATAAAACC** 25989
WF-Teratorn TCCTACGTGCCCAAAAAGAACAAAAACGTCCTGCTCCTTAGCACGAAGCATTCGAGGGTCGAAATATGCCACGAACGCAGGGACAAAAAACCAGAGATGATCTTGGATTACAATAGAACC 30496

TeratornLC199500 **AAGGGCGGCGTGGACAACCTGGACAAACTGCTGGCCACGTACAGCTGCAGGAGAATGACCAAGCGCTGGCCCTTGGCCATGTTCCACAACATGATAGACATGTCGGCGTACAACGCGTAC** 26109
WF-Teratorn AAGGGCGGCGTGGACAACCTGGACAAACTGCTGGCCACGTACAGCTGCAGGAGAATGACCAAGCGCTGGCCCTTGGCTATGTTCCACAACATGATAGACATGTCGGCGTACAACGCGTAC 30616

TeratornLC199500 **GTGATATGGAGGGAAATGCACCCGGACTGGATGCCGTGGAACCGCAACCGAAGGAGAATCTTCCTCGAGCAGCTGGGCAAAGATCTCGTCAGACCGCTCGTACTTCGCAGGAAGCGTGTT** 26229
WF-Teratorn GTGATATGGAGGGAAATGCACCCGGACTGGATGCCGTGGAACCGCAACCGAAGGAGAATCTTCCTCGAGCAGCTGGGCAAAGATCTCGTCAGACCGCTCGTACTTCGCAGGAAGCGTGTT 30736

TeratornLC199500 **CCTCGCGCAAGGCCGGCCTACGATCTAATGGTCAGCATGCAAGAAGCCGGCGGGACGAGCGCGCGCAGCGAGGGCAAAAGAAAACGATGTCAGATATGCCCGCGCTCCAAGGACACAAAG** 26349
WF-Teratorn CCTCGCGCAAGGCCGGCCTACGATCTAATGGTCAGCATGCAAAAAGCCGGCGGGACGAGCGCGCGCAGCGAGGGCAAAAGAAAACGATGTCAGATATGCCCGCGCTCCAAGGACACAAAG 30856

TeratornLC199500 **ACGCTGATCTCCTGCAAGAGCTGCGGAAAGAGCATCTGCAGAGGCTGCACTGTCCCCTTTTGCCCCCTTTGCTCCGAGCGCCAGGGAAGGGACGGCGACACAGACGCGTCCCAGTGA**AAG 26469
WF-Teratorn ACGCTGATCTCCTGCAAGAGCTGCGGAAAGAGCATCTGCAGAGGCTGCACTGTCCCCTTTTGCCCCCTTTGCTCCGAGCGCCAGGGAAGGGACGGCGACACAGACGCGTCCCAGTGAAAG 30976

TeratornLC199500 GGATCGAGCTGTCTGGTAGCGCGACCTGATGCCTCGTTTCCCACTCGAGTAACTGAAAATCTGTTCTGTTTTGATATCAGCGAGTCTGTCCTCTGTGCCGCCCCTGTATGTCCTCTGAAAA 26589
WF-Teratorn GGATCGAGCTGTCTGGTAGCGCGACCTGATGCCTCGTTTCCCACTCGAGTAACTGAAAATCTGTTCTGTTTTGATATCAGCGAGTCTGTCCTCTGTGCCGCCCCTGTATGTCCTCTGAAAA 31096

TeratornLC199500 TAAAACTGAGAACTGCTGTCGTACAAAGGTGTCTGTGTGCGCGTGTGTCTGTGTGAACATTCGAGAAGCAAAACTGTTGAAAGGGTTTGAGGGGTCTACGCGAAGGTACGGATATTTCGA 26709
WF-Teratorn TAAAACTGAGAACTGCTGTCGTACAAAGGTGTCTGTGTGCGCGTGTGTCTGTGTGAACATTCGAGAAGCAAAACTGTTGAAAGGGTTTGAGGGGTCTACGCGAAGGTACGGATATTTCGA 31216

TeratornLC199500 TCCAACGTGTTCGATTCACAAAGAAAGGCCTACAAACAGGGCTGGTTTGAGCTCGCAAAGCCTCTCAGCGCCCACAGGCCAACTCTGTCCTCGTTCGAGTGGATAAACAGAGTCAGCAGG 26829
WF-Teratorn CTTAACGTGTTCGATTCACAAAGAAAGGCCTACAAACAGGGCTGGTTTGAGCTCGCAAAGCCTCTCAGCGCCCACAGGCCAACTCTGTCCTCGTTCGAGTGGATAAACAGAGTCAGCAGG 31336

```
TeratornLC199500  GATTCTAAGCTTTAGGGGGAGAGAATTGGCCTGCAAGCTCTGAGGCTTGGCGTTCTGGAGGTATCTTCACATTCAGAATGCTTGAGTCCGGAAAGATTTTTTTTTTCTTCCCACTCGAG  26949
WF-Teratorn       GATTCTAAGCTTTACGGGGAGAGAATTGGCCTGCAAGCTCTGAGGCTTGGCGTTCTGGAGGTATCTTCACATTCAGAATGCTTGAGTCCGGAAAGATTTTTTTTTTCTTCCCACTCGAG  31456

TeratornLC199500  CTGTCAGAATTACATGTGGAGCCAGTTTGGGGAGGGAACCGCAACAGCCGAGAGCTCAGCGGAAGGGGAAAAAAATGAGTTGGCCTTGTAGCACCAAATCTCTTATGAATCCTATGAGCT  27069
WF-Teratorn       CTGTCAGAATTACATGTGGAGCCAGTTTGGGGAGGGAACCGCAACAGCCGAGAGCTCAGCGGAAGGGGAAAAAAATGAGTTGGCCTTGTAGCACCAAATCTCTTATGAATCCTATGAGCT  31576

TeratornLC199500  CTCCACCTAAACCTACTGTGAATGTTGTGTGAGATTTTCACTGGGTGTAAAATAAAATAAAAACAGAACAAATAATCGAAAAGAACCCAAACATTATAGTTTTTTTTTT-GTGTGATTCG  27188
WF-Teratorn       CTCCACCTAAACCTACTGTGAATGTTGTGTGAGATTTTCACTGGGTGTAAAATAAAATAAAAACAGAACAAATAATCGAAAAGAACCCAAACATTATAGTTTTTTTTTTTGTGTGATTCG  31696

TeratornLC199500  GGGATGTCTTCTGATGCAAAAGGCTCGTAGCGCTGGCCAGCCTTCGCGATCTGGTCTGCGCCCTGGAAACGAGCAGCGTGGGTTTGCGAGGGTAGCCGAGCTGCCACACGGCTGCTCGTT  27308
WF-Teratorn       GGGATGTCTTCTGATGCAAAAGGCTCGTAGCGCTGGCCAGCCTTCGCGATCTGGTCTGCGCCCTGGAAACGAGCAGCGTGGGTTTGCGAGGGTAGCCGAGCTGCCACACGGCTGCTCGTT  31816

TeratornLC199500  CTGGGATAGGATTGGCCTGCGAGCTCCTCTCATCAACGACACGCTTCAGTTTGCCTCTGATTCGGCTTCCCTGACACAGCGCGCTGTGATGTGACTCGCAGAGCCACACATCAAACCCTT  27428
WF-Teratorn       CTGGGATAGGATTGGCCTGCGAGCTCCTCTCATCAACGACACGCTTCAGTTTGCCTTTGATTCGGCTTCCCTGACACAGCGCGCTGTGATGTGACTCGCAGAGCCACACATCAAACCCTT  31936

TeratornLC199500  CCCATAGAGCTTCATGTGCTGTCAGAGGGGAGGCGATCGGCCTGCTCTGGGGCTGGGCTCCCCTTGGGGTTGCTTCACATTTGGAACATTTGGAGATACAAGCGGAGCCAGCCCGGAGAG  27548
WF-Teratorn       CCCATAGAGCTTCATGTGCTGTCAGAGTGGAGGCGATCGGCCTGCTCTGGGGCTGGGCTCCCCTTGGGGTTGCTTCACATTTGGAACATTTGGAGATACAAGCGGAGCCAGCCCGGAGAG  32056

TeratornLC199500  GGAACAGAAACGGACCACAGCTCAACGGAAAGGAGAGAATGTTGGTCTTTGAGCACTCCGCCTGAAGCTATCGCGAATATTCCGTGAGGTTTTCACTGTCTGTAAAATAAAAATAAAAAA  27668
WF-Teratorn       GGAACAGAAACGGACCACAGCTCAACGGAAAGGAGAGAATGTTGGTCTTTGAGCACTCCGCCTGAAGCTATCGCGAATATTCCGTGAGGTTTCCACTGTCTGTAAAATAAAAATAAAAAA  32176

TeratornLC199500  ACAGGACAGAAAATCCAAACGAACACAGAAATTGTTTTCATTTTGTAAGACTTTTTATTACAACATGACCTGGGTATCCAACCCAGGGTGGAACCGAAACGGTCAGAGCTCAGCGGAAT  27788
WF-Teratorn       -CAGGACAGAAAATCCAAACGAACACAGAAATTGTTTTTATTTTGTAAGACTTTTTATTACAACATGACCTGGGTATCCAACCCAGGGTGGAACCGAAACGGTCAGAGCTCAGCGGAAT  32295

TeratornLC199500  GGAGAAAGTCGACCCTCCCTGACACAGAGCGCTGCGGTGTGACTCGTGGCATCCAGCCTGTCTCTCAGTCAGCTCACTCTCAGGCGCAGGCTGCGTGTGTTTTTCAGACGCGAAAGGCCC  27908
WF-Teratorn       GGAGAAAGTCGGCCCTCCCTGACACAGAGCGCTGCGGTGTGACTCGTGGCATCCAGCCTGTCTCTCAGTCAGCTCACTCTCAGGCGCAGGCTGCGTGTGTTTTTCAGACGCGAAAGGCCC  32415

TeratornLC199500  CAGCCTGTCTCTGAGGATGCCCCCCTGCCCCCCGACACAGAGCTCTGTGGTGTGTGGATTGTGTTCTGAAAGGGTTTTCTCACTCGAGCGGTCAGAATTACATCCTGAGCCACGCAGAGC  28028
WF-Teratorn       CAGCCTGTCTCTGAGGATGCCCCCCTGCCCCCCGACACAGAGCTCTGTGGTGTGTGGATTGTGTTCTGAAAGGGTTTTCTCACTCGAGCGGTCAGAATTACATCCTGAGCCACGCAGAGC  32535

TeratornLC199500  CGCTCGTGCGGTTCCCTTCACCCAAACCTGAGCCTCTCCACGGAGCTTCATGTGCTGTCAGAGGGGAGAGGGTCTGCATGCGGGCTGCTCTCAGCCAATGCACGATCTAGCCTGCCCCTG  28148
WF-Teratorn       CGCTCGTGCGGTTCCCTTCACCCAAACCTGAGCCTCTCCACGGAGCTTCATGTGCTGTCAGAGGGGAGAGGGTCTGCATGCGGGCTGCTCTCAGCCAATGCACGATCTAGCCTGCCCCTG  32655

TeratornLC199500  CGCGGTCCAGCCTTCCTCTCATTCTCCTCCCCTGCAGGCGCGGGCTGCGTGTGTGCCTCAGACAGGAAAGGCTCCAGCCTTCCTCTGCCTCGGCCCCCCTTGAAACAGCGCTCAGTGGCT  28268
WF-Teratorn       CGCGGTCCAGCCTTCCTCTCATTCTCCTCCCCTGCAGGCGCGGGCTGCGTGTGTGCCTCAGACAGGAAAGGCTCCAGCCTTCCTCTGCCTCGGCCCCCCTTGAAACAGCGCTCAGTGGCT  32775

TeratornLC199500  TTACTTCACGCAGAGCCGCTCAGGATTTTTCCCTACACACACCCCAAGCCTTTCCGTCGAGCTTCATGTGCTGTCAGAGGGGACAGGCTCTGACTGCGGGCTGCGCTGCAGCTCGGCTTC  28388
WF-Teratorn       TTACTTCACGCAGAGCCGCTCAGGATTTTTCCCTACACACACCCCAAGCCTTTCCGTCGAGCTTCATGTGCTGTCAGAGGGGACAGGCTCTGACTGCGGGCTGCGCTGCAGCTCGGCTTC  32895

TeratornLC199500  CTTGAGGTTGCTTCAAAGTTAGAACGCTTGTGTTCGGAAAGGGTTTTCTCACTCCGGCTGTCAGAATTACATCCTGAGCCACCCTGGAGAGGGAACAGAAATGGCCCAGAGCTGAACAGA  28508
WF-Teratorn       CTTGAGGTTGCTTCAAAGTTAGAACGCTTGTGTTCGGAAAGGGTTTTCTCACTCCGGCTGTCAGAATTACATCCTGAGCCACCCTGGAGAGGGAACAGAAATGGCCCAGAGCTGAACAGA  33015

TeratornLC199500  CTGAAGAGAATGTCCTTCTTAGAGCTCTACGCCTGAATCTATCGCAAATGTTCCGTGAGGTTTTCACTGGCTGTAAAATAAAAATTAGAAAAAAACAGAACAGAAAATCCAAACGAATTT  28628
WF-Teratorn       CTGAAGAGAATGTCCTTCTTAGAGCTCTACGCCTGAATCTATCGCAAATGTTCCGTGAGGTTTTCACTGGCTGTAAAATAAAAATTAGAAAAAAACAGGACAGAAAATCCAAACGAATTT  33135

TeratornLC199500  CCCCCCGGGAGGGGCTGCGTGTGTTTCTCAGACGCGAAAGGCTCCGGCCTGCCTCTGAGAGTGCACCCTGAGCCTCTGCAGAGAGCCTCCGTAGAGAGCCTGCCTGCGAGTGGGGACTTC  28748
WF-Teratorn       CCCCCCGGGAGGGGCTGCGTGTGTTTCTCAGACGCGAAAGGCTCCGGCCTGCCTCTGAGAGTGCACCCTGAGCCTCTGCAGAGAGCCTCCGTAGAGAACCTGCCCGCGAGTGGGGTCTTC  33255

TeratornLC199500  CCGACAGAAGCGGAGCCCTGTGCCTTGTGACATCTGTTTCGAAAGACCGCAGCCTGCTTCTGC------------------TTCTGCAG--------------------------------  28819
WF-Teratorn       CCGACAGAAGCGGAGACCTTTGCTTTGTGACATCTGTTTCGAAAGACCGCAGCCTGCTTCTGCAGCTTGCTTGACACGGATTTCTGCGGAGCCTGGCTGGTGTTGGGAAAGATTTGTTTT  33375

TeratornLC199500  ------------------------------------------------------------------------------------------------------------------------  28819
WF-Teratorn       CTCACTCGAGCGGTCAGAATTAAGTGCAGGGCCAGCTCTGGGAGTGAACCGAAAGGGCCCGGAGCTCAGCGGAGTGAAGATAAAGTTTGCCACCGAGCTCTTTGCCCGAAGCTGTCTAGA  33495

TeratornLC199500  ------------------------------------------------------------------------------------------------------------------------  28819
WF-Teratorn       GTGTTCTGTGAAGATTTTGCTGGCTGTGAAATAAAAATAAAAACGAAATAAAAACCCAAAGGAACACAGAAATTTCTTTTGTGTGATGACTTTTTTATTACAACACGAGCTGGGTATCGA  33615
```

TeratornLC199500 ------------------------------------------------------------------------------------------------------------------------ 28819
WF-Teratorn ACCCAGGGGTGGAACAGAAACGGTCCGCAGTTCAGCCAAACGGAGAAAAAGTTGGCCTTCCAGGACCACATCTTCAGCTATCGCAGCCTGCTTAGGAGTGTGCACCTCTCAAACAGACAA 33735

TeratornLC199500 ------------------------------------------------------------------------------------------------------------------------ 28819
WF-Teratorn GGAGCCCTGTGGAGCGTGGTTCGCGTTGGGAACGCAGCAGCCTGCTTCTGAACCGCCCCCGGCTGATATAGAGCCCTGTGGAGCGTGGCTCGCGTTGCGAAAGGCTCCGGTCTGACTATG 33855

TeratornLC199500 ------------------------------------------------------------------------------------------------------------------------ 28819
WF-Teratorn AGTCTGTCAGCGGAATTCCCCTTCCAGCAGCACATCTGAATGCCCTCTCAGCCTGCTTATGAATCGGCCCTGCCTGACACACCGCTCTGCCGTGTGACTCGTGGCATCCTGCCTGTCTCT 33975

TeratornLC199500 ------------------------------------------------------------------------------------------------------------------------ 28819
WF-Teratorn CGGTCCGCTCACTCGCGGGCGGGGGCTGCCTTTGTTTCTCAGACGCGAAAGGCTCGGCATGCCTCTGAGGGTGCGGCCTGAGCCTCTGTAGAGATCCTCCGTATAGAGGCTCGAGCTGCA 34095

TeratornLC199500 --------------------------------------------------------------------------------------------CTTGCTTGACACGGATTTCTGCGGAGCG 28847
WF-Teratorn GCCTTCCAGCGAGTGGGGACTTCCCTACAGAAGCGGAGCCCTGTGCTTTGTGACTCGTGGTGCGAAAGACCGCAGCCTTCTTCTGAGCCGACCCTGCCTGACACAGGTTTCTGCGGAGCG 34215

TeratornLC199500 TGGCTGGTGTTGGGAAAGATTTGTTTTCTCACTCGAGCGGTCAGAATTAAGTGCAGGGCCAGCTCTGGGAGTGAACCGAACGGGCCCGGAGCTCAGCAGAGTGAAGAGAAAGTTTGCCAC 28967
WF-Teratorn TGGCTGGTGTTGGGAAATATTTGTTTTCTCACTCGAGCGGTCAGAATTAAGTGCAGGGCCAGCTCTGGGAGTGAACCGAACGGGCCCGGAGCTCAGCGGAGTGAAGAGAAAGTTTGCCAC 34335

TeratornLC199500 CGAGCTCTTTGCCCGTTGCTGTCTAGAGTGTTCTGTGAGGTTTTTGCTGGCTGTGAAATAAAAATGAAAACGAAGTAAAAACGGGACAGAAAACCCAAAAGAACACAGAAAATTTATTTT 29087
WF-Teratorn CGAGCTCTTTGCCCGTAGCTGTCTAGAGTGTTCTGTGAGGTTTTTGCTGGCTGTGAAATAAAAATGAAAACGAAGTAAAAACGGGACAGAAAACCCAAAAGAACACAGAAAATTTCTTTT 34455

TeratornLC199500 GTGTGATGACTTTTTTTATTACAACACGAGCTGGGTATCGAACCCAGGGGTGGAACAGAAACGGTCCGCAGCTCAGCCAACCCTGCCTGACACACAGCTCTGCCGTGTGACTCGTGGCAT 29207
WF-Teratorn GTGTGATGACTTTTTTTATTACAACACGAGCTGGGTATCGAACCCAGGGGTGGAACAGAAACGGTCCGCAGCTCAGCCAACCCTGCCTGCCACACAGCTCTGCCGTGTGACTCGTGGCAT 34575

TeratornLC199500 CCAGCCTGTCTCTCGCTCGGCTCACTCGCGGGCGGGGGCTGCCTGTGTTTCTCAGACGCGAAAGGCTCCGGCCTGCCTCTGAGGGTGCGGCCTGAGGAAAGGTTTTTTTTTTTTTTTTT 29327
WF-Teratorn CCAGCCTGTCTCTCGCTCGGCTCACTCGCGGGCGGGGGCTACCTGTGTTTCTCAGACGCGAAAGGCTCCGGCCTGCCTCTGAGGGTGCTGCCTGAGGAAAGGTTTTTTTTTTTTTTTTT 34695

TeratornLC199500 CCCTCTCGAGCTTTCAGAATTACATGTGGAGCCAGCTTGGGGAGGGAACCGCGACAGCCAAAAGCTCAGCGGAATGGAGAAAGAGTTGGCCTTCTAGCACTAAATCTCTTATTGTTTTTG 29447
WF-Teratorn CCCTCTCGAGCTTTCAGAATTACATGTGGAGCCAGCTTGGGGAGGGAACCGCGACAGCCAAAAGCTCAGCGGAATGGAGAAAGAGTTGGCCTTCTAGCACTAAATCTCTTATTGTTTTTG 34815

TeratornLC199500 CAAATGAGCTGTCCACCTCATGCTACTGTGAATGTCGTGTGAGGTTTTCACTAGATGTAAAATAAAAAACAGAGCAAAAAATCCAAAAGAACCCAAAAATTATTGTAATTGTTTTATTTT 29567
WF-Teratorn CAAATGAGCTGTCCACCTCATGCTACTGTGAATGTCGTGTGAGGTTTTCACTAGATGTAAAATAAAAAACAGAGCAAAAAATCCAAAAGAACCCAAAAATTATTGTAATTGTTTTATTTT 34935

TeratornLC199500 TTATTTTTTGCATGATGATTTTCTTTGTTGTGACTCGGGGATGCCTCCTGATGCAAAAGGCTCATCGCGTTGGCCAGAAAGAGTTGCATTCAGAAAGAGTTTTTCTCACTCGAGCTG**TCA** 29687
WF-Teratorn TTATTTTTTGCATGATGATTTTCTTTGTTGTGACTCGGGGATGCCTCCTGATGCAAAAGGCTCATCGCGTTGGCCAGAAAGAGTTGCATTCAGAAAGAGTTTTTCTCACTCGAGCTGTCA 35055

TeratornLC199500 **GAGATACAAGCGGATCCAGCCCGAAGAGGGAACAGAAACGGGCCACAGCTCAACGGTCTGCCTGCGGGCCGCTCTCAGACAAAGCACGATCCAGCCTGCCTTCAGTCGCTCCCCTGCACG** 29807
WF-Teratorn GAGATACAAGCGGAGCCAGCCCGAAGAGGGAACAGAAACGGGCCACAGCTCAACGGTCTGCCTGCGGGCCGCTCTCAGACAAAGCACGATCCAGCCTGCCTTCAGTCGCTCCCCTGCACG 35175

TeratornLC199500 **ATCCAGCCTGCCTCTCATTCCCTCCCCTGCAGGCTCGTGTGCCTCAGACGCAAAAGGCTCCAGCCTTCCTCTGCGTTGGCCCCCCCTTGACACGAGCCTCCGTGTCTCTACTCCACACAG** 29927
WF-Teratorn ATCCAGCCTGCCTCTCATTCCCTCCCCTGCAGGCTCGTGTGCCTCAGACGCAAAAGGCTCCAGCCTTCCTCTGCGTTGGCCCCCCCTTGACACGAGCCTCCGTGTCTCTACTCCACACAG 35295

TeratornLC199500 **AGCCTGCCTGCGGGCCGCTCTCAGACAAAGCACGATCCAGCCTGCCTTCAGTCGCTCCCC--TGCACGATCCAGCCTGCCTCTCATTCCCTCCCCTGCAGGCTCGTGTGCCTCAGACGCA** 30045
WF-Teratorn AGCCTGCCTGCGGGCCGCTCTCAGACAAAGCACGATCCAGCCTGCCTTCAGTCTCTCCCCCCTGCACGATCCAGCCTGCCTCTCATTCCCTCCCCTGCAGGCTC---------------- 35399

TeratornLC199500 **AAAGGCTCCAGCCTTCCTCTGCGTTGGCCCCCCCTTGACACGAGCCTCCGTGTCTCTACTCCACACAGAGCCTGCCTGCGGGCCGCTCTCAGACAAAGCACGATCCAGCCTGCCTTCAGT** 30165
WF-Teratorn -----------------------------CCCCCTTGACACGTGCCTCCGTGTCTCTACTCCACACAGAGCCTGCCTGCGGGCCGCTCTCAGACAAAGCACGATCCAGCCTGCCTTCAGT 35490

TeratornLC199500 **CGCTCCCCTGCACGATCCAGCCTGCCTCTCATTCCCTCCCCTGCAGGCTCGTGTGCCTCAGACGCAAAAGGCTCCAGCCTTCCTCTGCGTTGGCCCCCCCTTGACACGAGCCTCCGTGTC** 30285
WF-Teratorn CGCTCCCCTGCACGATCCAGCCTGCCTCTCATTCCCTCCCCTGCAGGCTCGTGTGCCTCAGACGCAAAAGGCTCCAGCCTTCCTCTGCGTTGGCCCCCCCTTGACACGAGCCTCCGTGTC 35610

TeratornLC199500 **TCTACTCCACACAGAGCCTGCCTGCGGGCCGCTCTCAGACAAAGCACGATCCAGCCTGCCTTCAGTCGCTCCCCTGCACGATCCAGCCTGCCTCTCATTCCCTCCCCTGCAGGCTCGTGT** 30405
WF-Teratorn TCTACTCCACACAGAGCCTGCCTGCGGGCCGCTCTCAGACAAAGCACGATCCAGCCTGCCTTCAGTCGCTCCCCTGCACGATCCAGCCTGCCTCTCATTCCCTCCCCTGCAGGCTCGTGT 35730

TeratornLC199500 **GCCTCAGACGCAAAAGGCTCCAGCCTTCCTCTGCGTTGGCCCCCCCTTGACACGAGCCTCCGTGTCTCTACTCCACACAGAGCCTGCCTGCGGGCCGCTCTCAGACAAAGCACGATCCAG** 30525
WF-Teratorn GCCTCAGACGCAAAAGGCTCCAGCCTTCCTCTGCGTTGGCCCCCCCTTGACACAAGCCTCCGTGTCTCTACTCCACACAGAGCCTGCCTGCGGGCCGCTCTCAGACAAAGCACGATCCAG 35850

TeratornLC199500 **CCTGCCTTCAGTCGCTCCCC--TGCGCGATCCAGCCTGCCTCTCAT**TCCTTCCCCTGCACGCGTGTGTGCCTCAGACGCAAAAGGCTCCAGCCTTCCTCTGCGTTGGGCCCCCCCTTGAC 30643
WF-Teratorn CCTGCCTTCAGTCTCTCCCCCCTGCACGATCCAGCCTGCCTCTCATTCCCTCCCCTGCAGGCTCGTGTGCCTCAGACGCAAAAGGCTCCAGCCTTCCTCTGAGTTGG-CCCCCCCTTGAC 35969

TeratornLC199500 ACGAGCTTCAGTGTCTTTACTCCACACAGAGCCTGCCTGCGGGCGGCTCTCAGACAAAGCACGATCCAGCCTGCCTTCAGTCGCTCCCC--TGCACAAATCCAGCCTGCCTCTCATTCCC 30761
WF-Teratorn ACGAGCCTCCGTGTCTCTACTCCACACAGAGCCTGCCTGCGGGCCGCTCTCAGACAAAGCACGATCCAGCCTGCCTTCAGTCTCTCCCCCCTGCACGA-TCCAGCCTGCCTCTCATTCCC 36088

TeratornLC199500 TCCCCTGCAGGCGTGTGTGCCTCAGACGCAAAAGGCTCCAGCCTTCCTCTGCGTTGGGCCCCCCCTTGACACGAGCCTCGCGGAATGGAGAAAGAGTTGGCCTTCTAGCACTAAATCTCT 30881
WF-Teratorn TCCCCTGCAGGC---TCCCCCCTTGACACGAG-------------CCTCCGTGT-----CTCTACTCCACAC--------------------AGAGCCTGCCTGCGGGCCG-----CTCT 36162

TeratornLC199500 CATTGTTTTTGCAAATGAGCTCTCCAACTCATGCTACTGTGAATGTCGTGTGAGGTTTTCACTAGATGTAAAATAAAAAACAGAGCAAAAAATCCAAAAGAACCCAAAAATTATTGTAAT 31001
WF-Teratorn CAG-------ACAAA-GCACGATCCAGC-----CTGCCTTCA--GTCGC--------TCCCCTGCACG----------------------ATCCAG----------------------- 36213

TeratornLC199500 TGTTTTATTTTTTATTTTTTGCATGATGATTTTCTTTGTTGAGACTCGGGGATGCCTTCTGATGCAAAAGGCTCATCGCATTGGCCAGAAAGAGTTGAATTCAGAAAGAGTTTTTCTCAC 31121
WF-Teratorn ---CCTGCCTCTCATTCCCT------------CCCCTGC--AGGCTCGTG--TGCCT-CAGACGCAAAAGGCTC-------CAGCC------------------------TTCCTCTGCG 36282

TeratornLC199500 TCGAGCTGTCAGAGATACAAGCGGAGCCA-GCCTGAAGAGGGAACAGAAACGGGCCACAGCTCA-ACGGTCTGCCTGCGGGCGGCTCTCAGACAAAGCACGATCCAGCCTGCCTTCAGTC 31239
WF-Teratorn TTGGCCCCCCCTTGACACGAGCCT--CCGTGTCT--------------------CTACTCCTCACAGAGCCTGCCTGCGGGCCGCTCTCAGACAAAGCACGATCCAGCCTGCCTTCAGTC 36380

TeratornLC199500 GCTCCCCTGCACAAATCCAGCCTGTCTCTCATTCCCTCCCCTGCAGGCGTGTGTGCCTCAGACGCAAAAGGCTCCAGCCTTCCTCTGCGTTGGCCCCCCCTTGACACGAGCTTCAGTGTC 31359
WF-Teratorn GCTCCCCTGCACGA-TCCAGCCTGCCTCTCATTCCCTCCCCTGCAGGCTCGTGTGCCTCAGAGGCAAAAGGCTCCAGCCTTCCTCTGCGTTGGCCCCCCCTTGACACGAGCCTCCGTGTC 36499

TeratornLC199500 TTTACTCCACACAGAGCCTGCCTGCGGGCCGCTCTCAGACAAAGCACGATCCAGCCTGCCTTCAGTCGCTCCCCTGCGCGATCCAGCCTGCCTCTCATTCCCTCCCCTGCACGCGTGTGT 31479
WF-Teratorn TCTACTCCACACAGAGCCTGCCTGCGGGCCGCTCTCAGACAAAGCACGTTCCAGCCTGCCTTCAGTCGCTCCCCTGCACGATCCAGCCTGCCTCTCATTCCCTCCCCTGCAGGCTCGTGT 36619

TeratornLC199500 GCCTCAGACGCAAAAGGCTCCAGCCTTCCTCTGCGTTGGGCCCCCCCTTGACACGAGCCTCCGTGTCTCTACTCCACACAGAGCCTGCCTGCGGGCGGCTCTCAGACAAAGCACGATCCA 31599
WF-Teratorn GCCTCAGACGCAAAAGGCTCCAGCCTTCCTCTGCGTTGG-CCCCCCCTTGACACGAGCCTCCGTGTCTCTACTCCACACAGAGCCTGCCTGCGGGCCGCTCTCAGACAAAGCACGATCCA 36738

TeratornLC199500 GGCTGCCTTCAGTCGCTCCCC--TGCACAAATCCAGCCTGCCTCTCATTCCCTCCCCTGCAGGCGTGTGTGCCTCAGACGCAAAAGGCTCCAGCCTTCCTCTGCGTTGGGCCCCCCCTTG 31717
WF-Teratorn GCCTGCCTTCAGTCTCTCCCCCCTGCACGA-TCCAGCCTGCCTCTCATTCCCTCCCCTGCAGGC---TCCCCCCTTGACACGAG-------------CCTCCGTGT-----CTCTACTCC 36836

TeratornLC199500 ACACGAGCCTCGCGGAATGGAGAAAGAGTTGGCCTTCTAGCACTAAATCTCTCATTGTTTTTGCAAATGAGCTCTCCAACTCATGCTACTGTGAATGTCGTGTGAGGTTTTCACTAGATG 31837
WF-Teratorn ACAC--------------------AGAGCCTGCCTGCGGGCCG-----CTCTCAG-------ACAAA-GCACGATCCAGC-----CTGCCTTCA--GTCGC--------TCCCCTGCACG 36908

TeratornLC199500 TAAAATAAAAAACAGAGCAAAAAATCCAAAAGAACCCAAAAATTATTGTAATTGTTTTATTTTTTATTTTTTGCATGATGATTTTCTTTGTTGAGACTCGGGGATGCCTTCTGATGCAAA 31957
WF-Teratorn ----------------------ATCCAG-----CC----------------TGCCTCTCATTCCCTCCCCTGCATG-------------------CTCGTG--TGCCT-CAGACGCAAA 36962

TeratornLC199500 AGGCTCATCGCGTTGGCCAGAAAGAGTTGAATTCAGAAAGAGTTTTTCTCACTCGAGCTGTCAGAGATACAAGCGGAGCCAGCCTGAAGAGGGAACAGAAACGGGCCACAGCTCAACG-- 32075
WF-Teratorn AGGCTC-------CAGCC--------TTCCTCT------GCGTTGCCCCCCCTTGA-----------CACGAGC--------CTC----------------CGTGTCTCTACTCCACACA 37026

TeratornLC199500 --GTCTGCCTGCGGGCGGCTCTCAGACAAAGCACGATCCAGCCTGCCTTCAGTCGCTCCCCTGCGCGATCCAGCCTGCCTCTCATTCCCTCCCCTGCACGCGTGTGTGCCTCAGACGCAA 32193
WF-Teratorn GAGCCTGCCTGCGGGCCGCTCTCAGACAAAGCACGATCCAGCCTGCCTTCAGTCGCTCCCCTGCACGATCCAGCCTGCCTCTCATTCCCTCCCCTGCAGGATCGTGTGCCTCAGAG--AA 37144

TeratornLC199500 AAGGCTC-------CAGCCTTCCTCTGCG-TTGGGCC--CCCCCTTGACACGAGCTTCAGTGTCTTTACT---------------CCACACGGAGCCT------------------GCCT 32270
WF-Teratorn AAGGCTCGTAGCGCTGGCCAGCCTTCGCGATCTGGTCTGCGCCCTGGAAACGAGCAGC-GTGGGTTTGCGAGGGTAGCCGAGCTGCCACACGGCTGCTCGTTCTGGGATAGGATTGGCCT 37263

TeratornLC199500 GCGGGCGGCTCTCAGACAAAGCACGATCCAGCCTGCCTTCAGTC--GCTCCCCTG-CACAA---------ATCCAGCCTGC-----CTCTCATT----CCCTCCCCTGCAGGCGTGTGTG 32369
WF-Teratorn GCGAGCTCCTCTCATCAACGACTCGCTTCAGTTTGCCTCTGATTCGGCTTCCCTGACACAGCGCGCTGTGATGTGACTCGCAGAGCCACACATCAAACCCTTCCCATAGAGCTTCATGTG 37383

TeratornLC199500 CC-TCAGACGCAAAAGGCTCCAGCCTTCCTCTGCGTTGGGCCCCCC---------CTTGACAC---GAGCCTCGCGGAATGGAGAAAGAGTTGGCCTTC--------TAGCACTAAATCT 32468
WF-Teratorn CTGTCAGA-GTGGAGGCGATCGGCCTGCTCTGGGGCTGGGCTCCCCTTGGGGTTGCTTCACATTTGGAACATTTGGAGATACAAGCGGAGCCAGCCCGGAGAGGGAACAGAAACGGACCA 37502

TeratornLC199500 ---CTCATTG------------TTTTTGCAAATGAGCTCTCCAACTCATGCTACTGTGAATGTCGTGTGAGGTTTTCACTAGATGTAAAATAAAAA------ACAGAGCAAAAAATCCAA 32567
WF-Teratorn CAGCTCAACGGAAAGGAGAGAATGTTGGTCTTTGAGCACTCCGCCTGAAGCTATCGCGAATATTCCGTGAGGTTTCCACTGTCTGTAAAATAAAAATAAAAAACAGGACAGAAAATCCAA 37622

TeratornLC199500 AAGAACCCAAAAATTATTGTAATTGTTTTA---TTTTTTATT-------------------------------------------------TTTTGCATGAT----------GATTTTCT 32625
WF-Teratorn ACGAACACAGAAATTGTTTTTATTTTGTAAGACTTTTTTATTACAACATGACCTGGGTATCCAACCCAGGGTGGAACCGAAACGGTCAGAGCTCAGCGGAATGGAGAAAGTCGGCCCTCC 37742

```
TeratornLC199500  TTGT--------------TGAGACTCG----------------------------------------GGGATGC-------CTTCTGATGCAAAAGGCTC-----------------  32664
WF-Teratorn       CTGACACAGAGCGCTGCGGTGTGACTCGTGGCATCCAGCCTGTCTCTCAGTCAGCTCACTCTCAGGCGCAGGCTGCGTGTGTTTTTCAGACGCGAAAGGCCCCAGCCTGTCTCTGAGGAT  37862

TeratornLC199500  ATCGCGTTGGCC----AGAAAGAG----------------TTGAATTCAGAAAGAGTTTTTCTCACTCGAGCTGTCAGAGATACAAGCGGAGCCAGCCTGAA------GAGGGA------  32752
WF-Teratorn       GCCCCCCTGCCCCCCGACACAGAGCTCTGTGGTGTGTGGATTGTGTTCTGAAAGGGTTTT-CTCACTCGAGCGGTCAGAATTACATCCTGAGCCACGCAGAGCCGCTCGTGCGGTTCCCT  37981

TeratornLC199500  --ACAGAAAC--GGGCCAC-------AGCTCAACG------------------GTCTGCCTGCGGGCGGCTCTCAGACAAAGCACGATCCAGCCTGCCTTCAGTCGCTCCCCTGCACAA  32842
WF-Teratorn       TCACCCAAACCTGAGCCTCTCCACGGAGCTTCATGTGCTGTCAGAGGGGAGAGGGTCTGCATGCGGGCTGCTCTCAGCCAATGCACGATCTAGCCTGCC------------CCTGCGCGG  38089

TeratornLC199500  ATCCAGCCTGCCTCTCATTC-CCTCCCCTGCAGGCG---------TGTGTGCCTCAGACGCAAAAGGCTCCAGCCTTCCTCTGCGTTGGGCCCCCCCTTGACACGAGCCTCGCGGAATGG  32952
WF-Teratorn       -TCCAGCCTTCCTCTCATTCTCCTCCCCTGCAGGCGCGGGCTGCGTGTGTGCCTCAGACAGGAAAGGCTCCAGCCTTCCTCTGCCTCGG--CCCCCCTTGAAACAGCGCTC----AGTGG  38202

TeratornLC199500  AGAAAGAGTTGGCCTTCTAGCACTAAATCTCTCATTGTTTTTGCAAATGAGCTCTCCAACTCATGCTACTGTGAATGTCGTGTGAGGTTTTCACTAGATGTAAAATAAAAAACAGAGCAA  33072
WF-Teratorn       C-------TTTACTTC---ACGCAGAGCCGCTCAGGATTTTTCCCTACACACACCCCAAGCCTTTCC---------GTCG----AGCTTT-------ATGTGCTGT------CAGAGGGG  38286

TeratornLC199500  AAAATCCAAAAGAACCCAAAAATTATTGTAATTGTTTTATTTTTTATTTTTTGCATGATGATTTTCTCTGTTGAGACTCGGGGATGCCTTCTGATGCAAAAGGCTCATCGCGTTGGCCAG  33192
WF-Teratorn       ACAG---------GCTC-----TGACTGCGG--GCTGCGCT-----------GCAGCTCGGCTTCCTT-----------GAGGTTGCTT-------CAAA-----------GTTAG----  38346

TeratornLC199500  AAAGAGTTGAATTCAGAAAGAGTTTTTCTCACTCGAGCTGTCAGAGATACAAGCGGAGCCAGCCTGAAGAGGGAACAGAAACGGGCCACAGCTCAACGGTCTGCCT-GCGGGCCGCTCTC  33311
WF-Teratorn       -AACGCTTGTGTTCGGAAAGGGTTTT-CTCACTCCGGCTGTCAGAATTACATCCTGAGCCACCCTGGAGAGGGAACAGAAATGGCCCAGAGCTGAACAGACTGAAGAGAATGTCCTTCTT  38464

TeratornLC199500  AGA---CAAAGCACGA-TCCA--GCCTGCCTTCAGTC--GCTCCC-----CTGCGCGATCCAGCCTG---------------------CCT--CTCATTCCCTCCCCTGCA----CGCGT  33391
WF-Teratorn       AGAGCTCTACGCCTGAATCTATCGCAAATGTTCCGTGAGGTTTTCACTGGCTGTAAAATAAAAATTAGAAAAAAACAGAACAGAAAATCCAAACGAATTTCCCCCCGGGAGGGGCTGCGT  38584

TeratornLC199500  GTGTGCCTCAGACGCAAAAGGCTCCAGCCTTCCTCTGCGTTGGGCCCCCCCTTGACACGAGCTTCAGTGTCTTTACTCCACACAGAGCCTGCCTGCGGGCGG---CTCTCAGACA-AAGC  33507
WF-Teratorn       GTGTTTCTCAGACGCGAAAGGCTCCGGCCTGCCTCTGAG---AGTGCACCCT------GAGCCTCTGCAGAGAGCCTCCGTAGAGAACCTGCCCGCGAGTGGGGTCTTCCCGACAGAAGC  38695

TeratornLC199500  ACGATCCAGCCTGCCTTCAGTCGCTCCCCTGCACAAATC-CAGCCTGCCTCT-CATTCCCTCCCCTGCAGGCGTGTGT---GCCT--CAGACGC---AAAAGGCTCCAGCCTTCCTCTGC  33617
WF-Teratorn       GGAGACC--TTTGCTTTGTGACATCTGTTTCGAAAGACCGCAGCCTGCTTCTGCAGCTTGCTTGACACGGATTTCTGCGGAGCCTGGCTGGTGTTGGGAAAGATTT--GTTTTCTCACTC  38811

TeratornLC199500  GTTGGGCCCCCCCTTGACACGAGCCTCGC----GGAATGGA--GAAAGAGTTGGCCTTCTAGC--ACTAAATCTCTCATTGTTTTTGCAAATGAGCTCTCCAACTCATGCTACTGTGAAT  33729
WF-Teratorn       GAGCGGTCAGAATTAAGTGCAGGGCCAGCTCTGGGAGTGAACCGAAAGGGCCCGGAGCTCAGCGGAGTGAAG-----ATAAAGTTTGCCACCGAGCTCTTTGCCCGAAGCTGTCTAGAGT  38926

TeratornLC199500  GTCGTGTGAGGTTTTCACTAGATGTAAAATAAAAAACAGAGCAAAA----AATCCAAAAGAACCCAAAAATTATTGTAATTGTTTTATTTTTTATTTTTTGCATGATGATTTTCTTTGTT  33845
WF-Teratorn       GTTCTGTGAGGATTTTGCTGGCTGTGAAATAAAAATAAAAACGAAATAAAAACCCAAAGGAACACAGAAATTTCTTT---TGTGTGATGACTTTTTATTACAACACGAGCTGGGT-ATC  39042

TeratornLC199500  GAGACTCGGGGATGCCTTCTGATGCAAAAGGCTCATCGCGTTGGCCA------GAAAGAGTTGA--ATTCAGAAAGAGTTTTTC------------TCACTCGAG---------CTGTCA  33936
WF-Teratorn       GA-ACCCAGGGGTGG-----AACAGAAACGGTCCGCAGT-TCAGCCAAACGGAGAAAAAGTTGGCCTTCCAGGACCACATCTTCAGCTATCGCAGCCTGCTTAGGAGTGTGCACCTCTCA  39155

TeratornLC199500  GAGATACAAG--------CGGAGC----------------------CAGCCTG------AAGAG---------GGAACAGAA----ACGGGCCACAGCTC---------AA-----CGGT  33993
WF-Teratorn       AACAGACAAGGAGCCCTGTGGAGCGTGGTTCGCGTTGGGAACGCAGCAGCCTGCTTCTGAACCGCCCCCGGCTGATATAGAGCCCTGTGGAGCGTGGCTCGCGTTGCGAAAGGCTCCGGT  39275

TeratornLC199500  CTGCCTGCGGG--------CGG-------CTCTCAG----ACA---AAGCACGATC-CAGCCTGCCT-----TCAGT--------------CGCTCCCCTGCACAA--------ATCCAG  34063
WF-Teratorn       CTGACTATGAGTCTGTCAGCGGAATTCCCCTTCCAGCAGCACATCTGAATGCCCTCTCAGCCTGCTTATGAATCGGCCCTGCCTGACACACCGCTCTGCCGTGTGACTCGTGGCATCCTG  39395

TeratornLC199500  CCTGCCTCTCATTCC-CTCCCCTGCAGGCGTG---------TGTGCCTCAGACGCAAAAGGCTCCAGCGTTCCTCTGCG--------TTGGGCCCCCCTTGACAC----------GAGCC  34155
WF-Teratorn       CCTGTCTCTCGGTCCGCTCACTCGCGGGCGGGGGCTGCCTTTGTTTCTCAGACGCGAAAGGCTC-GGCATGCCTCTGAGGGTGCGGCCTGAGCCTCTGTAGAGATCCTCCGTATAGAGGC  39514

TeratornLC199500  TCGCG---------------GAATGGAGA---------AAGAG--------------TTG-GCCTTCTAGCACTAAATCTCTCATTGTTTTTGCAAATGAGCTCT--CCAACTCATGCTA  34234
WF-Teratorn       TCGAGCTGCAGCCTTCCAGCGAGTGGGGACTTCCCTACAGAAGCGGAGCCCTGTGCTTTGTGACTCGTGGTGCGAAAGACCGCAGCCTTCTTCTGAGCCGACCCTGCCTGACACAGGTTT  39634

TeratornLC199500  CTGTGAA----------TGTCGTGTGAGGTTT------TCACT--AGATGT-AAAATAAAAAACAGAGCAAA------AAAT--CCAAAAAGAACCCAAAAATTATTGTAATTGT-----  34322
WF-Teratorn       CTGCGGAGCGTGGCTGGTGTTGGGAAAGATTTGTTTTCTCACTCGAGCGGTCAGAATTAAGTGCAGGGCCAGCTCTGGGAGTGAACCGAAAGGGCCCGGAGCTCAGCGGAGTGAAGAGAA  39754

TeratornLC199500  --TTTATTTTTTATTTTTTGCATGATGATTTT-------CTTTGTTGAGACTCGG--GGATGCCTTCTGATGCCAAAA---GGCTCATCGCGTTGGCCAGAAAG---AGTTGAATTCAG  34425
WF-Teratorn       AGTTTGCCACCGAGCTCTTTGCCCGAAGCTGTCTAGAGTGTTCTGTGAGGATTTTGCTGGCTGTGAAATAAAAATAAAAACGAAATAAAAACG--GGACAGAAAACCCAAAAGAACACAG  39872
```

```
TeratornLC199500  AAAGAGTTTTTT-----CTCACTCGAGCTGTCAGAGATACAAGCGGAG---CCAGCCTGAAGAGGGAACAGAAAACGGGCCACAGCTCAAC-GGTCCTGCCTG-CGGGCGGCTTC-TCAG   34534
WF-Teratorn       AAATTTTCTTTTGTGTGATGACTTTTTTTATTACA-ACACGAGCTGGGTATCGAACCCAGGGGTGGAACAGAAA-CGGTCCGCAGCTCCGCCAACCCTGCCTGACACACAGCTCTGCCGT   39990

TeratornLC199500  ACAAAGCACG--ATCCAGCCTGCCTTCAGTCGCTCCACCTGCGCGATCCNNNNNAGCCTGTCTCTCATTCCCTCCCCTGCAGGCGTGTGTGCCTCAGACGCAAAAGGCTCCAGCCTTCCT   34652
WF-Teratorn       GTGACTCGTGGCATCCAGCCTGTCTC---TCGCTTGGCTCACTCG---CGGAGGGGGCTGCCT-----------------------GTGTTTCTCAGACGCGAAAGGCTCCGGACTGCCT   40081

TeratornLC199500  CTGCGTTGGGCCCCCCCTTGACACGAGCTTCAGTGTCTTTACTCCACACAGAGCCTGCCTGCGGGCGG---CTCTCAGACA-AAGCACGATCCAGGCTGCCTTCAGTCG-CTCCCCTGCG   34767
WF-Teratorn       CTGAG---AGTGCACCCT------GAGCCTCTGCAGAGAGCCTCCGTAGAGAGCCTGCCTGCGAGTGGGGACTTCCCGACAGAAGCGGAGCCCTG--TGCCTTGTGACATCTGTTTCGAA   40190

TeratornLC199500  CGATC-CAGCCTGTCTCTCATTCC----CTCCCCTG---CAGGCGT-------GTGTGCCTCA-GACGCAAAAGGCTCCAGCCTTCCTCTGCGTTGGGCCCCCCCTTGACACGAGCCTCG   34871
WF-Teratorn       AGACCGCAGCCTGCTTCTGCTTCTGCAGCTTGCTTGACACGGATTTCTGCGGAGCGTGGCTGGTGTTGGGAAAGATTT--GTTTTCTCACTCGAGCGGTCAGAATTAAGTGCAGGGCCAG   40308

TeratornLC199500  C----GGAATGGA--GAAAGAGTTGGCCTTCTAGC--ACTAAATCTCTCATTGTTTTTGCAAATGAGCTCTCCAACTCAT-GCTACTGTGAATGTCGTGTGAGGTTTTCACTAGATGTAA   34982
WF-Teratorn       CTCTGGGAGTGAACCGAACGGGCCCGGAGCTCAGCGGAGTGAAG-----AGAAAGTTTGCCACCGAGCTCTTTG-CCCGTAGCTGTCTAGAGTGTTCTGTGAGGTTTTTGCTGGCTGTGA   40422

TeratornLC199500  AATAAAAA-----ACAGAGCAAAAA--ATCCAAAAGAACCCAAAAATTATTGTAATTGTTTTATTTTTTATTTTTTGCATGATGATTTTCTTTGTTGAGACTCGGGGATGCCTTCTGATG   35095
WF-Teratorn       AATAAAAATGAAAACGAAGTAAAAACGGGACAGAAAACCCAAAAGAACACAGAAA-------ATTTCTT-----TTGTGTGATGACTTTTTTTATTACAACACGAG--------CTGG-G   40521

TeratornLC199500  CAAAAGGCTCATCGCGTTGGCCAGAAAGAGTT--GAATTCAGAAAGAGTTTTTCTCACTCG-AGCTGT-CAGAGATACAAGCGG-AGCCAGCCTGAAGAGGGAACAGAAACGGGCCACAG   35210
WF-Teratorn       TATCGAACCCAGGG-GTGAAACAGAAACGGTCCGCAGCTCAGCCA-ACCCTGCCTGACACACAGCTCTGCCGTGTGACTCGTGGCATCCAGCCTGTCT------CTCGCTCGGCTCACTC   40633

TeratornLC199500  CTCAACGGT--CTGCCTGCGGGCCGCTCTCAGACAAAGCACGATCCAGCCTGCCTTCAGTCGCTCC-CCTGCACAAATCCAGCCTGCCTCTCATTCCCTCCCCTGCAGGCGTGTGTGCCT   35327
WF-Teratorn       GCGGGCGGGGGCTGCCTGTGT----TTCTCAGACGCGAAAGGCTCCGGCCTGCCTCTGAGGGTGCGGCCTGAGGAAA---GGTTTTTTTTTTTTTTCCCCTCTCG--AGCTT-------T   40737

TeratornLC199500  CAGA--CGCAAAAGGCTCCAGCCTTCCTCTGCGTTGGGCCCCCCCTTGACACGAGCCTCGCGGAATGGAGAAAGAGTTGGCCTTCTAGCACTAAATCTCTCATTGTTTTTGCAAATGAGC   35445
WF-Teratorn       CAGAATTACATGTGGAGCCAGCTTG-----GGGAGGGAACCGCGACAGCCAAAAGCTCAGCGGAATGGAGAAAGAGTTGGCCTTCTAGCACTAAATCTCTTATTGTTTTTGCAAATGAGC   40852

TeratornLC199500  TGTCCAACTCATGCTACTGTGAATGTCGTGTGAGGTTTTCACTAGATGTAAAATAAAAAACAGAGCAAAAAATCCAAAAGAACCCAAAAATTATTGTAATTGTTTTATTTTTTATTTTTT   35565
WF-Teratorn       TGTCCACCTCATGCTACTGTGAATGTCGTGTGAGGTTTTCACTAGATGTAAAATAAAAA-CAGAGCAAAAAATCCAAAAGAACCCAAAAATTATTGTAATTGTTTTATTTTTTATTTTTT   40971

TeratornLC199500  GCATGATGATTTTCTTTGTTGAGACTCGGGGATGCCTTCTGATGCAAAAGGCTCATCGCGTTGGCCAGAAAGAGTTGAATTCAGAAAGAGTTTTTCTCACTCGAGCTGTCAGAGATACAA   35685
WF-Teratorn       GCATGATGATTTTCTTTGTTGTGACTCGGGTATGCCTCCTGATGCAAAAGGCTCTTCGCGTTGGCCAGAAAGAGTTGCATTCAGAAAGAGTTTTTCTCACTCGAGCTGTCAGAGATACAA   41091

TeratornLC199500  GCGGAGCCAGCCTGAAGAGGGAACAGAAACGGGCCACAGCTCAACGGTCTGCCTGCGGGCCGCTCTCAGACAAAGCACGATCCAGCCTGCCTTCAGTCGCTCCCCTGCGCGATCCAGCCT   35805
WF-Teratorn       GCGGAGCCAGCCCGAAGAGGGAACAGAAACGGGCCACAGCTCAACGGTCTGCCTGCGGGCCGCTCTCAGACAAAGCACGATCCAGCCTGCCTTCAGTCGCTCCCCTGCACGATCCAGCCT   41211

TeratornLC199500  GCCTCTCATTCCCTCCCCTGCACGCGTGTGTGCCTCAGACGCAAAAGGCTCCAGCCTTCCTCTGCGTTGGGCCCCCCCTTGACACGAGCTTCAGTGTCTTTACTCCACACGGAGCCTGCC   35925
WF-Teratorn       GCCTCTCATTCCCTCCCCTGCAGGCTCGTGTGCATCAGACGCAAAAGGCTCCAGCCTTCCTCTGCGTTGG-CCCCCCCTTGACACGAGCCTCCGTGTCTCTACTCCACACAGAGCCTGCC   41330

TeratornLC199500  TGCGGGCGGCTCTCAGACAAAGCACGATCCAGCCTGCCTTCAGTCGCTCCCCTGCACAAATCCAGCCTGCCTCTCATTCCCTCCCCTGCAGGCGTGTGTGCCTCAGACGCAAAAGGCTCC   36045
WF-Teratorn       TGCGGGCCGCTCTCAGACAAAGCAAGATCCAGCCTGCCTTCAGTCGCTCCCCTGCACGA-TCCAGCCTGCCTCTCATTCCCTCCCCTGCAGGCTCGTGTGCCTCAGACGCAAAAGGCTCC   41449

TeratornLC199500  AGCCTTCCTCTGCGTTGGGCCCCCCCTTGACACGAGCCTCGCGGAATGGAGAAAGAGTTGGCCTTCTAGCACTAAATCTCTCATTGTTTTTGCAAATGAGCTCTCCAACTCATGCTACTG   36165
WF-Teratorn       AGCCTTCCTCTGCGTTGG-CCCCCCCTTGACACGAGCCTC-CG---------------TGTC--TCTA-------------------CTCCACACA-GAGCC----------TGC--CTG   41518

TeratornLC199500  TGAATGTCGTGTGAGGTTTTCACTAGATGTAAAATAAAAAACAGAGCAAAAAATCCAAAAGAACCCAAAAATTATTGTAATTGTTTTATTTTTTATTTTTTGCATGAT--GATTTTCTTT   36283
WF-Teratorn       CGG--GCCGC-------TCTCA-----------------GACAAAGCACGA--TCCAG-----CCTGC------CTTCAGTCGCTCC-----------CCTGCACGATCCAGCCTGCCT-   41587

TeratornLC199500  GTTGAGACTCGGGGATGCCTTCTGATGCAAAAGGCTCATCGCGTTGGCCAGAAAGAGTTGAATTCAGAAAGAGTTTTTCTCACTCGAGCTGTCAGAGATACAAGCGGAGCCAGCCTGAAG   36403
WF-Teratorn       -------CTC----ATTCCCTCCCCTGCA---GGCTCCCCCC-TTGAC----ACGAGCC---TCC-------GTGTCTCT-ACTC---CT------------------------------   41644

TeratornLC199500  AGGGAACAGAAACGGGCCACAGCTCAACGGTCTGCCTGCGGGCGGCTCTCAGACAAAGCACGATCCAGCCTGCCTTCAGTCGCTCCCCTGCGCGATCCAGCCTGCCTCTCATTCCCTCCC   36523
WF-Teratorn       ------------------CACAG------AGCCTGCCTGCGGGCCGCTCTCAGACAAAGCACGATCCAGCCTGCCTTCAGTCGCTCCCCTGCACGATCCAGCCTGCCTCTCATTCCCTCCC   41741

TeratornLC199500  CTGCACGCGTGTGTGCCTCAGACGCAAAAGGCTCCAGCCTTCCTCTGCGTTGGGCCCCCCCTTGACACGAGCTTCAGTGTCTTTACTCCACACGGAGCCTGCCTGCGGGCGGCTCTCAGA   36643
WF-Teratorn       CTGCAGGCTCGTGTGCCTCAGAGGCAAAAGGCTCCAGCCTTCCTCTGCGTTGG-CCCCCCCTTGACACGAGCCTCCGTGTCTCTACTCCACACAGAGCCTGCCTGCGGGCCGCTCTCAGA   41860
```

```
TeratornLC199500  CAAAGCACGATCCAGCCTGCCTTCAGTCGCTCCCCTGCACAAATCCAGCCTGCCTCTCATTCCCTCCCCTGCAGGCGTGTGTGCCTCAGACGCAAAAGGCTCCAGCCTTCCTCTGCGTTG  36763
WF-Teratorn       CAAAGCACGATCCAGCCTGCCTTCAGTCGCTCCCCTGCACGA-TCCAGCCTGCCTCTCATTCCCTCCCCTGCAGGCTCGTGTGCCTCAGACGCAAAAGGCTCCAGCCTTCCTCTGCGTTG  41979

TeratornLC199500  GGCCCCCCCTTGACACGAGCCTC-GCGG----AATGGAGAAAGAGTTGGCCTTCTAGCACTAAATCTCTCATTGTTTTTGCAAATGAGCTCTCCAACTCATGCTACTGTGAATGTCGTGT  36878
WF-Teratorn       GCCCCCCCCTTGACACGAGCCTCCGTGTCTCTACTCCACACAGAGCCTGCCTGCGGGCCG-----CTCTCAG-------ACAAA-GCACGATCCAGC-----CTGCCTTCA--GTCGC--  42077

TeratornLC199500  GAGGTTTTCACTAGATGTAAAATAAAAAACAGAGCAAAAAATCCAAAAGAACCCAAAAATTATTGTAATTGTTTTATTTTTTATTTTTTGCATGATGATTTTCTTTGTTGAGACTCGGGG  36998
WF-Teratorn       ------TCCCCTGCACG----------------------ATCCAG-------------------------CTTGCCTCTCATTCCCT------------CCCCTGC--AGGCTCGTG-  42127

TeratornLC199500  ATGCCTTCTGATGCAAAAGGCTCATCGCGTTGGCCAGAAAGAGTTGAATTCAGAAAGAGTTTTTCTCACTCGAGCTGTCAGAGATACAAGCGGAGCCA-GCCTGAAGAGGGAACAGAAAC  37117
WF-Teratorn       -TGCCT-CAGACGCAAAAGGCTC-------CAGCC-----------------------TTCCTCTGCGTTGGCCCCCCCTTGACACGAGCCT--CCGTGTCT----------------  42195

TeratornLC199500  GGGCCACAGCTCA-ACGGTCTGCCTGCGGGCGGCTCTCAGACAAAGCACGATCCAGCCTGCCTTCAGTCGCTCCCCTGCACAAATCCAGCCTGCCTCTCATTCCCTCCCCTGCAGGCGTG  37236
WF-Teratorn       ---CTACTCCTCACAGAGCCTGCCTGCGGGCCGCTCTCAGACAAAGCACGATCCAGCCTGCCTTCAGTCGCTCCCCTGCACGA-TCCAGCCTGCCTCTCATTCCCTCCCCTGCAGGC---  42308

TeratornLC199500  TGTGCCTCAGACGCAAAAGGCTCCAGCCTTCCTCTGCGTTGGGCCCCCCCTTGACACGAGCCTCGCGGAATGGAGAAAGAGTTGGCCTTCTAGCACTAAATCTCTCATTGTTTTTGCAAA  37356
WF-Teratorn       TCCCCCCTTGACACGAG-------------CCTCCGTGTC---TCTACTCCTCACA-GAGCCT---------------------GCCTGCGGGCCG-----CTCTCAG-------ACAAA  42378

TeratornLC199500  TGAGCTCTCCAACTCATGCTACTGTGAATGTCGTGTGAGGTTTTCACTAGATGTAAAATAAAAAACAGAGCAAAAAATCCAAAAGAACCCAAAAATTATTGTAATTGTTTTATTTTTTAT  37476
WF-Teratorn       -GCACGATCCAGC-----CTGCCTTCA--GTCGC--------TCCCCTGCACG----------------------ATCCAG-------------------------CCTGCCTCTCAT  42433

TeratornLC199500  TTTTTGCATGATGATTTTCTCTGTTGAGACTCGGGGATGCCTTCTGATGCAAAAGGCTCATCGCGTTGGCCAGAAAGAGTTGAATTCAGAAAGAGTTTTTCTCACTCGAGCTGTCAGAGA  37596
WF-Teratorn       TCCCT------------CCCCTGC--AGGCTCGTG--TGCCT-CAGAGGCAAAAGGCTC-CAGCCTTCCTCTGC----GTTGGCCCC----------CCCTTGACACGAGCCTCC-----  42516

TeratornLC199500  TACAAGCGGAGCCAGCCTGAAGAGGGAACAGAAACGGGCCACAGCTCAACGGTCTGCCTGCGGGCCGCTCTCAGACAAAGCACGATCCAGCCTGCCTTCAGTCGCTCCCCTGCGCGATCC  37716
WF-Teratorn       -----GTGTCTCTACTCCA--------------------CACAG------AGCCTGCCTGCGGGCCGCTCTCAGACAAAGCACGATCCAGCCTGCCTTCAGTCGCTCCCCTGCACGATCC  42605

TeratornLC199500  AGCCTGCCTCTCATTCCCTCCCCTGCACGCGTGTGTGCCTCAGACGCAAAAGGCTCCAGCCTTCCTCTGCGTTGGGCCCCCCCTTGACACGAGCTTCAGTGTCTTTACTCCACACGGAGC  37836
WF-Teratorn       AGCCTGCCTCTCATTCCCTCCCCTGCAGGCTCGTGTGCCTCAGACGCAAAAGGCTCCAGCCTTCCTCTGCGTTGG-CCCCCCCTTGACACGAGCCTCCGTGTCTCTACTCCACACAGAGC  42724

TeratornLC199500  CTGCCTGCGGGCGGCTCTCAGACAAAGCACGATCCAGCCTGCCTTCAGTCGCTCCCCTGCACAAATCCAGCCTGCCTCTCATTCCCTCCCCTGCAGGCGTGTGTGCCTCAGACGCAAAAG  37956
WF-Teratorn       CTGCCTGCGGGCCGCTCTCAGACAAAGCACGATCCAGCCTGCCTTCAGTCGCTCCCCTGCACGA-TCCAGCCTGCCTCTCATTCCCTCCCCTGCAGGCTCGTGTGCCTCAGACGCAAAAG  42843

TeratornLC199500  GCTCCAGCCTTCCTCTGCGTTGGGCCCCCCCTTGACACGAGCCTC-GCGG----AATGGAGAAAGAGTTGGCCTTCTAGCACTAAATCTCTCATTGTTTTTGCAAATGAGCTCTCCAACT  38071
WF-Teratorn       GCTCCAGCCTTCCTCTGCGTTGG-CCCCCCCTTGACACGAGCCTCCGTGTCTCTACTCCACACAGAGCCTGCCTGCGGGCCG-----CTCTCAG-------ACAAA-GCACGATCCAGC-  42948

TeratornLC199500  CATGCTACTGTGAATGTCGTGTGAGGTTTTCACTAGATGTAAAATAAAAAACAGAGCAAAAAATCCAAAAGAACCCAAAAATTATTGTAATTGTTTTATTTTTTATTTTTTGCATGATGA  38191
WF-Teratorn       ----CTGCCTTCA--GTCGC--------TCCCCTGCACG----------------------ATCCAG-------------------------CCTGCCTCTCATTCCCT---------  42996

TeratornLC199500  TTTTCTTTGTTGAGACTCGGGGATGCCTTCTGATGCAAAAGGCTCATCGCGTTGGCCAGAAAGAGTTGAATTCAGAAAGAGTTTTTCTCACTCGAGCTGTCAGAGATACAAGCGGAGCCA  38311
WF-Teratorn       ---CCCCTGC--AGGCTCGTG--TGCCT-CAGACGCAAAAGGCTC-------CAGCC-----------------------TTCCTCTGCGTTGGCCCCCCCTTGACACGAGCCT--CCG  43075

TeratornLC199500  -GCCTGAAGAGGGAACAGAAACGGGCCACAGCTCA-ACGGTCTGCCTGCGGGCGGCTCTCAGACAAAGCGCGATCCAGCCTGCCTCTCATTCCCTCCCCTGCAGG--CGTGTGTGCCTCA  38427
WF-Teratorn       TGTCT--------------------CTACTCCTCACAGAGCCTGCCTGCGGGCCGCTCTCAGACAAAGCACGATCCAGCCTGCCT-TCAGTCGCTCCCCTGCACGATCCAGCCTGCCTC-  43173

TeratornLC199500  GACGCAAAAGGCTCCAGCCTTCCTCTGCGTTGGGCCCCCCCTTGACACGAGCTTCAGTGTCTTTACTCCACACGGAGCCTGCCTGCGGGCGGCTCTCAGACAAAGCACGATCCAGCCTGC  38547
WF-Teratorn       ------------TCATTCCCTCCCCTGCA--GGCTCCCCCCTTGACACGAGCCTCCGTGTCTCTACTCCTCACAGAGCCTGCCTGCGGGCCGCTCTCAGACAAAGCACGATCCAGCCTGC  43279

TeratornLC199500  CTTCTGTCGCTCCCCTGCACGATCCAGCCTGCCTCTCATTCCCTCCCCTGCAGGCTCGTGTGCCTCAGACGCAAAAGGCTCCAGCCTTCCTCTGCGTTGGCCCCCCCTTGACACGAGCCT  38667
WF-Teratorn       CTTCAGTCGCTCCCCTGCACGATCCAGCCTGCCTCTCATTCCCTCCCCTGCAGGCTCGTGTGCCTCAGACGCAAAAGGCTCCAGCCTTCCTCTGCGTTGGCCCCCCCTTGACACGAGCCT  43399

TeratornLC199500  CCGTGTCTCTACTCCACACAGAGCCGCTCGTGATTTTTGCTTCTCACAAACCGAGCCTTTCCGTCAAGCTTCATGTGCTGACGGAGGGGACAGGGTCCGCCTGCGGGCTGCGCTGCAGCT  38787
WF-Teratorn       CCGTGTCTCTACTCCACACAGAGCCGCTCGTGATTTTTGCTTCACACAAACCGAGCCTTTCTGTCAAGCTTCATGTGCTGACGGAGGGGACAGGGTCCGCCTGCGGGCTGCGCTGCAGCT  43519

TeratornLC199500  CCCGTGAATTTGCTTCAAAGTCAGAACGCTTGTGTTCGGAAAGGGTTTTCTCAATCCGGCTGTCAGAATTACATCCCGAGGCAGCCCGGAGAGGGAACAGAAATGGCCCAGAGCTCAGCA  38907
WF-Teratorn       CCCGTGAAGTTGCTTCAAAGTCAGAACGCTTGTGTTCGGAAAGGGTTTTCTCAATCCGGCTGTCAGAATTACATCCCGAGGCAGCCCGGAGAGGGAACAGAAATGGCCCAGAGCTCAGCA  43639
```

TeratornLC199500 GACTGAATAGAATGTCGGTCTTAGAGCTCTCCGCCTGAAGCTATTGGGAATGTTCTTTGAGGTTTTCAATGAGTGTGAAATAAAAAAATTAAAAAAAACAAACAGGACAGAAAATCCGAA 39027
WF-Teratorn GACTGAATAGAATGTCGGTCTTAGAGCTCTCCGCCTGAAGCTATTGGGAATGTTCTTTGAGGTTTTCAATGAGTGTGAAATAAAAAAAT-AAAAAAAACAAACAGGACAGAAAATCCAAA 43758

TeratornLC199500 TGAAAACAGAAATTGTTTGTTTTTTGTATGACTTCTCCAGCCTGCCTCCTGCCTCTGAGTCGCCCCTCCCTGACACAGAGCTCTGTGGCGTGTGACTCGTCTGAGAGGAAGTCTCC---- 39143
WF-Teratorn TGAAAACAGAAATTGTTTGTTTTT-GTATGACTTCTCCAGCCTGCCTCCTGCCTCTGAGTCGCCCCTCCCTGACACAGAGCTCTGTGGCGTGTGACTCGTCTGAGAGGAAGTCTCCCGCC 43877

TeratornLC199500 ------------------------------------------------------------------------AGCCCGCCTCTGAGCGGGCTCCCCTCGACAGGACGCACAGCTCCGTGG 39191
WF-Teratorn TGCCTCCTGAGCGGGCTTCTCCTGACAGAAGAAGAGCTCCGAGGCTGTGCGGCTCTGGTTGCTAAAGGCTCCAGCCCGCCTCTGAGCGGGCTCCCCTCGACAGGACGCACAGCTCCGTGG 43997

TeratornLC199500 CGTGTGACTCGTCTCGACCAAAGGTTGCAGCCTGCCCCTGAGTCTGCCTCTCCTGACAGACGCAGGGCTGCCTGGCGTGTGGGTCGTTTTTGGAAAGGTTTTTTCTCTCTCGAGCGGCCA 39311
WF-Teratorn CGTGTGACTCGTCTCGACCAAAGGTTGCAGCCTGCCCCTGAGTCTGCCTCTCCTGACAGACGCAGGGCTGCCTGGCGTATGGGTCGTTTTTGGAAAGGTTTTTTCTCTCTCGAGCGGCCA 44117

TeratornLC199500 GAATTACATGCGGGGCCAGATCGGGGAGTGAACCGAAACGGCCCAGAGCTCAGCAGAACGAAGAGAAAGAGTTGGCCTTCAAGCAACAAATTTAAAGCTATTCTGAATGTTCTGTGAGGT 39431
WF-Teratorn GAATTACATGCGGGGCCAGATCGGGGAGTGAACCGAAACGGCCCAGAGCTCAGCAGAACGAAGAGAAAGAGTTGGCCTTCAAGCAACAAATTTAAAGCTATTCTGAATGTTCTGTGAGGT 44237

TeratornLC199500 TTGCGATGGGTGTGAAATAAAAGAAAAAAAAA-TGAAACGGGACAGAAAATCCAAACGAACACAGAAATTGTTTGTTTTGGTATGACTTTTTTATTTCAACCCAGGGATAGCTCAGCGAA 39550
WF-Teratorn TTGCGATGGGTGTGAAATAAAAGAAAAAAAAAATCAAACGGGACAGAAAATCCAAACGAACACAGAAATTGTTTGTTTTGGTATGACTTTTTTATTTCAACCCAGGGATAGCTCAGCGAA 44357

TeratornLC199500 ATGGAGAAAGAGTTGGCTTTCCAGCCCCACACCTGAAGCAATTGCAGCCCGCAGAGCTCAGTGGAGTGTGACTCTTCTGAGAAGTACCCAGATTTTTGTTTTGTTCTTGTGGGCATACAA 39670
WF-Teratorn ATGGAGAAAGAGTTGGCTTTCCAGCCCCACACCTGAAGCAATTGCAGCCCGCAGAGCTCAGTGGAGTGTGACTCTTCTGAGAAGAACCCAGATTTTTGTTTTGTTCTTGTGGGCATACAA 44477

TeratornLC199500 CGGGGAACAGAGATGCTTCAGAGCAGGCTCAGAGACCCCGCCGAGCTTTGGAGAGCGATAAAAAGAAATGAAAAGCAAAAGAAGCTGTGCTTTTTTTTTTAAAAAAAAAA-CGGTGCTTA 39789
WF-Teratorn CGGGGAACAGAGATGCTTCAGAGCAGGCTCAGAGACCCCGCCGAGCTTTGGAGAGTGATAAAAAGAAATGAAAAGCAAAAGAAGCTGTGCTTTTTTTTTAAAAAAAAAAAAACGGTGCTTA 44597

TeratornLC199500 GAAAGGCGACTGGTAGTCGTGCGAAACACTTTCGGTGCTGCTGTTAGATACCGCACTCTTCATTCTGTTTCAGAGTCTGCCCATCGGAGTCCTCGCTGTCCCCAGCCACGCATTTGAGAG 39909
WF-Teratorn GAAAGGCGACTGGTAGTCGTGCGAAACACTTTCGGTGCTGCTGTTAGATACCGCACTCTTCATTCTGTTTCAGAGTCTGCCCATCGGAGTCCTCGCTGTCCCCAGCCACGCATTTGAGAG 44717

TeratornLC199500 GGAAATACCGCCGCCCCTCACTACGGCACGCTGCCCCACGAGTAAATCCCACGCTCGAACAGTCAGGGGAAGAGCCTCGGGATTTACATTTAGGGCTGACCCAGGCTGGAAACCAAAAGG 40029
WF-Teratorn GGAAATACCGCCGCCCCTCACTACGGCACGCTGCCCCACGAGTAAATCCCACGCTCGAACAGTCAGGGGAAGAGCCTCGGGATTTACATTTAGGGCTGACCCAGGCTGGAAACCAAAAGG 44837

TeratornLC199500 GCCCAGAACTCCGTCACGAGAGGAGAGGGTTGGCGTTCGAGCAATAGGTTCAGAAGCTATTGCAAACATTGTGTGTGGTTTTCACTGGCTGTGGAATAAAACAATAACAAAACGAATAAA 40149
WF-Teratorn GCCCAGAACTCCGTCACGAGAGGAGAGGGTTGGCGTTCGAGCAATAGGTTCAGAAGCTATTGCAAACATTGTGTGTGGTTTTCACTGGCTGTGGAATAAAACAATAACAAAACGAATAAA 44957

TeratornLC199500 AAAGCCACAAAGACACCCAGAATTGGTTTTGCGTGGTGATTTTTTTTTT-ATTACAGCATGACTTGGAGACGCTCTCATAGACAATAAAATGGAAAAATACAAAAGATTTGTCCAGCGAA 40268
WF-Teratorn AAAGCCACAAAGACACCCAGAATTGGTTTTGCGTGGTGATTTTTTTTTTTATTACAGCATGACTTGGAGACGCTCTCATAGACAATAAAATGGAAAAATACAAAAGATTTGTCCAGCGAA 45077

TeratornLC199500 TCGCTCAGATGTTCCTAG**TCAACGTCTCCGTCGCTTGTCAAGGCTCTCAGAGCAGACTGACCTGGAGGTCAGCCGTTTGCGCGAACGGCCCATCCCTCTCAGTCCAGAAGTCCTCACTTC** 40388
WF-Teratorn TCGCTCAGATGTTCCTAGTCAACGTCTCCGTCGCTTGTCAAGGCTCTCAGAGCAGATTGACCTGGAGGTCAGCCGTTTGCGCGAACGGCCCATCCCTCCCAGTCCAGAAGTCCTCACTTC 45197

TeratornLC199500 **GTGCCGAGTTTCACCTGCGCGAGACTGGGTCGAAGGCCACGTTCTCATCTCCAGAGCGGCGCGACCGTCGGCCTTGCGCAAGACGCTCACCGCGCGGGCCAACCTTCGCGATCTCATAGG** 40508
WF-Teratorn GTGCCGAGTTTCACCTGCGCGAGACTGGGTCGAAGGCCACGTTCTCATCTCCAGAGCGGCGCGACCGTCGGCCTTGCGCAAGACGCTCACCGCGCGGGCCAACCTTCGCGATCTCATAGG 45317

TeratornLC199500 **CGCCCTGGACGCGTCCTCCCCCTTGACCACAGTCATGGCCACGGCGTAGTCCCGTTTCAGGCTTTCGACGGTGAACGCATCATGACCGTGCAGTCTGTTGAACACCTCGCAGGCTTCCAG** 40628
WF-Teratorn CGCCCTGGACGCGTCCTCCCCCTTGACCACAGTCATGGCCACGGCGTAGTCCCGTTTCAGGCTTTCGACGGTGAACGCATCATGACCGTGCAGTCTGTTGAACACCTCGCAGGCTTCCAG 45437

TeratornLC199500 **CCACTCCCAATCCGGTTCCAGCTCGACGACGGGCCGGCGGGTGCCGTCTTCCTCCCTCAGCAGAATTCGCTCCTGGTGGTACACGGTTCGCGTGGCGCAGAAGGAGCTCAGGTTTTTCAC** 40748
WF-Teratorn CCACTCCCAATCCGGTTCCAGCTCGACGACGGGCCGGCGGGTGCCGTCTTCCTCCCTCAGCAGAATTCGCTCCTGGTGGTACACGGTTCGCGTGGCGCAGAAGGAGCTCGCGTTTTTCAC 45557

TeratornLC199500 **CGGCAGGCCCGAGTACTCCGGCGAGTAGACGCTCAAGCGAGGCCCGTGAAAAGTGTCCGCCGGGCCTCTCTCGGACTCCCTCTCGTGCAGAGGGTCCAGAGATCCCCGCGGCGGGTCGTC** 40868
WF-Teratorn CGGCAGGCCCGAGTACTCCGGCGAGTAGACGCTCAAGCGAGGCCCGTGAAAAGTGTCCGCCGGGCCTCTCTCGGACTCCCTCTCGTGCAGAGGGTCCAGAGATCCCCGCGGCGGGTCGTC 45677

TeratornLC199500 **TCCCCTGAGCCTGAAGGCTCTCGCCACCAGCCCTCCGTAGTACAAGGGCGCAGTGCAGCGAAAGAGCTGTCCCCCGATGGCGCATTTGTTGAACGAGAGCTCGTTCAGAGGATCCACTCC** 40988
WF-Teratorn TCCCCTGAGCCTGAAGGCTCTCGCCACCAGCCCTCCGTAGTACAAGGGCGCAGTGCAGCGAAAGAGCTGTCCCCCGATGGCGCATTTGTTGAACGAGAGCTCGTTCAGAGGATCCACTCC 45797

TeratornLC199500 **CGCCGTCGCCTCGAGTATCAGGGAGCGAACCGGGTCGTCGCCGAAGTCCAGGGAAAACACCAGTCTAGCGCTCCGATCCATCTCCTCGTCCCCGTCGGCCTGTGCGGTCCCGAACGCGTT** 41108
WF-Teratorn CGCCGTCGCCTCGAGTATCAGGGAGCGAACCGGGTCGTCGCCGAAGTCCAGGGAAAACACCAGTCTAGCGCTCCGATCCATCTCCTCGTCCCCGTCGGCCTGTGCGGTCCCGAACGCGTT 45917

TeratornLC199500 **GGTCTTTTCGACCGTTTCGTTGTCATAGGCGTCCATCATCGCTTCGCCCAGCTGGTCCCAGAAGCCCCGGTGCGCCTCGGGCGCGTACAGAGCCAGCATGTCGTAGACCGCTTCGAGAGA** 41228
WF-Teratorn GGTCTTTTCGACCGTTTCGTTGTCATAGGCGTCCATCATCGCTTCGCCCAGCTGGTCCCAGAAGCCCCGGTGCGCCTCGGGCGCGTACAGAGCCAGCATGTCGTAGACCGCTTCGAGAGA 46037

TeratornLC199500 **AAGGGTGACGGAGCTCACCTTTTTGGGCCGAGCGTGTCCACGCCTGCCCTGCTGCGTGTTCTGAGAGGGGGCGCATTTGTACGAGCAGAGCCTGCGGCTCCTGGGACCCGTCCCGCCACG** 41348
WF-Teratorn AAGGGTGACGGAGCTCACCTTTTTGGGCCGAGCGTGTCCACGCCTGCCCTGCTGCGTGTTCTGAGAGGGGGCGCATTTGTACGAGCAGAGCCTGCGGCTCCTGGGACCCGTCCCGCCACG 46157

TeratornLC199500 **TTCGGGAAACGTGCAGTTCGAGCAAAGCTGGAGCATTTCGTGAGCGGCGTCGTTGCCGCCTCTAAATTCGGGGCCCGAAAAACTCAAAGAATCGACGCCCGAAATGCACATCTGACCGTC** 41468
WF-Teratorn TTCGGGAAACGTGCAGTTCGAGCAAAGCTGGAGCATTTCGTGAGCGGCGTCGTTGCCGCCTCTAAATTCGGGGCCCGAAAAACTCAAAGAATCGACGCCCGAAATGCACATCTGACCGTC 46277

TeratornLC199500 **CTTGACGGCGGTCTTCAGGCCCTTTTCCCGCACCTGACAAAGAAGGCGGCCCAGCGCCTTGCACGCGAAGCGTCCCTGCGCGTCCAGCACGCGGGGGCAAATGTCCGGCAGGCAGGCGGC** 41588
WF-Teratorn CTTGACGGCGGTCTTCAGGCCCTTTTCCCGCACCTGACAAAGAAGGCGGCCCAGCGCCTTGCACGCGAAGCGTCCCTGCGCGTCCAGCACGCGGGGGCAAATGTCCGGCAGGCAGGCGGC 46397

TeratornLC199500 **CGCGCAGAGCTTGAACGCCGCGTTGGTCTCCAGGCAGATCTGTCCGAAAGACAGCCTGTCCAGGGCCCTCAGTCTGTTCCTGTTCAGCTGCGTCCAGAAGCCGCGGAGGTCCGGGAAGCT** 41708
WF-Teratorn CGCGCAGAGCTTGAACGCCGCGTTGGTCTCCAGGCAGATCTGTCCGAAAGACAGCCTGTCCAGGGCCCTCAGTCTGTTCCTGTTCAGCTGCGTCCAGAAGCCGCGGAGGTCCGGGAAGCT 46517

TeratornLC199500 **CCCCTCGTCTGGCACCTCCGTCCACGGAGGGACCCACACGTCGAACACAAAGTGGCAGGCCAGTTTCATGGCCTCGGCGAAGACCAGCTGGTTTATGCAGAAGGACACCCGTCTGCTCAA** 41828
WF-Teratorn CCCCTCGTCTGGCACCTCCGTCCACGGAGGGACCCACACGTCGAACACAAAGTGGCAGGCCAGTTTCATGGCCTCGGCGAAGACCAGCTGGTTTATGCAGAAGGACACCCGTCTGCTCAA 46637

TeratornLC199500 **GGCTCCCGCGCACACGACGTGACTGGCGTACATGGCCTCGACGCAGGCGTCTCTGGCCGCGTCGAACCTCCTGTCGAACTGCATGATCTCGGACTCGGCGATCATGGTGGCCGACTGCAC** 41948
WF-Teratorn GGCTCCCGCGCACACGACGTGACTGGCGTACATGGCCTCGACGCAGGCGTCTCTGGCCGCGTCGAACCTCCTGTCGAACTGCATGATCTCGGACTCGGCGATCATGGTGGCCGACTGCAC 46757

TeratornLC199500 **CAGGTTCTGCCTGTACAGGCAGACGGCAGCCACCTTGTCCATGGATCGCTTCTCCACTGTGTCCTGCAGCTGTTTCGAGCTGCGCGTCCGAGTCACGTGCGCCGAGAACTCGGATTCCAT** 42068
WF-Teratorn CAGGTTCTGCCTGTACAGGCAGACGGCAGCCACCTTGTCCATGGATCGCTTCTCCACTGTGTCCTGCAGCTGTTTCGAGCTGCGCGTCCGAGTCACGTGCGCCGAGAACTCGGATTCCAT 46877

TeratornLC199500 **GATCACGCAGCGGTCGGCCCAGGCGTCGCTCACGAACCCCAGGGTGTTGGTGTTCCACACGAACCCGCAGTTGTGCACCGCGTGATACCTCACGGTGTCCACCTTTCCGCTGCACGCGTC** 42188
WF-Teratorn GATCACGCAGCGGTCGGCCCAGGCGTCGCTCACGAACCCCAGGGTGTTGGTGTTCCACACGAACCCGCAGTTGTGCACCGCGTGATACCTCACGGTGTCCACCTTTCCGCTGCACGCGTC 46997

TeratornLC199500 **CTTGCCCACCACGTCGCTCTCCAAGACGCTGGTGTCCAGCAGGTTCTTGAACGTGTTGGGTATCACGCTGGACTCCTTGTCCAGGGCCTTTAGGTTTTTCTCGTGGGTGATGTGAGCGTC** 42308
WF-Teratorn CTTGCCCACCACGTCGCTCTCCAAGACGCTGGTGTCCAGCAGGTTCTTGAACGTGTTGGGTATCACGCTGGACTCCTTGTCCAGGGCCTTTAGGTTTTTCTCGTGGGTGATGTGAGCGTC 47117

TeratornLC199500 **GTCTATCATCACCACGCAAGCTTTCCTCTGTTGCTTGTACTTGAAAGCTTGCGGGGTGAAGGACGTGAGCGTCTCTATGCAGCCCTTCACCTGCCTGAACTGGCAGTTCAGCACGTTGTT** 42428
WF-Teratorn GTCTATCATCACCACGCAAGCTTTCCTCTGTTGCTTGTACTTGAAAGCTTGCGGGGTGAAGGACGTGAGCGTCTCTATGCAGCCCTTCACCTGCCTGAACTGGCAGTTCAGCACGTTGTT 47237

TeratornLC199500 **GGCGTAGCTCTTGCCCTCCCCGGGCCCGGTGACGTTGAACAGCATGATCCGTTCGATCGTGTACCTGGGCGCGGTGTTGGTCACCACGTAGTTGAAGGTGGTGGAGGCCTGAAATAGCCT** 42548
WF-Teratorn GGCGTAGCTCTTGCCCTCCCCGGGCCCGGTGACGTTGAACAGCATGATCCGTTCGATCGTGTACCTGGGCGCGGTGTTGGTCACCACGTAGTTGAAGGTGGTGGAGGCCTGAAATAGCCT 47357

TeratornLC199500 GCCCACCTGGAGGCTGTCAGAGAACATGTAGCGCACCAGCGCGTGGTCGTAGACCTTCACCGGGTTCAGCGAGCGCGGGTCCTCGGCGTAGTCGTCGTAGTAGTCCTCAGCCGGGAACTC 42668
WF-Teratorn **GCCCACCTGGAGGCTGTCAGAGAACATGTAGCGCACCAGCGCGTGGTCGTAGACCTTCACCGGGTTCAGCGAGCGCGGGTCCTCGGCGTAGTCGTCGTAGTAGTCCTCAGCCGGGAACTC** 47477

TeratornLC199500 **CAGGCACAGCTCCTCGTGTATGTCGGGGAGAGCGCGGCCGAGAGACAGGCGCGAGTTGAGCATCCCTTCTCCCATCAGGTCCCGGAGCCCCGCGTACAGCGCGTCGGCGTGCGAGGAACG** 42788
WF-Teratorn CAGGCACAGCTCCTCGTGTATGTCGGGGAGAGCGCGGCCGAGAGACAGGCGCGAGTTGAGCATCCCTTCTCCCATCAGGTCCCGGAGCCCTGCGTACAGCGCGTCGGCGTGCGAGGAACG 47597

TeratornLC199500 **GCGCTTCGCGAGGATGTCGAAGACCATGTTGGCTATGCTCTCGAAGTGCCTGTGTCCTCCGTATTTCGTCTTCATGTCCTTGCTCAGGCGTTCGAGTTTTCCCAGGTGAGGGTACAGGTG** 42908
WF-Teratorn GCGCTTCGCGAGGATGTCGAAGACCATGTTGGCTATGCTCTCGAAGTGCCTGTGTCCTCCGTATTTCGTCTTCATGTCCTTGCTCAGGCGTTCGAGTTTTCCCAGGTGAGGGTACAGGTG 47717

TeratornLC199500 **CGCAAAGTAATGGTTGTAGTCGCACCTGCTCACGGAGAAGGGGTGCTGCTCGCACACGTTGTAGAAGCACGCGTACGCAGTCTGACAGTCTTCCCGGCACCTCCTCCCTCGACGATCCTG** 43028
WF-Teratorn CGCAAAGTAATGGTTGTAGTCGCACCTGCTCACGGAGAAGGGGTGCTGCTCGCACACGTTGTAGAAGCACGCGTACGCAGTCTGACAGTCTTCCCGGCACCTCCTCCCTCGACGATCCTG 47837

TeratornLC199500 **GTCGACGGCCGGGCCGGGCCGGGGGCAGGGGCAGTGGCAGCGGCCGCCCGTCAACAGCCGAGCGAGGTGCCGGGTCCTGCGGTTGGCCCTGATCTCTCGCAGCGCTCTCGAGATGCGAGG** 43148
WF-Teratorn GTCGACGGCCGGGCCGGGCCGGGGGCAGGGGCAGTGGCAGCGGCCGCCCGTCAACAGCCGAGCGAGGTGCCGGGTCCTGCGGTTGGCCCTGATCTCTCGCAGCGCTCTCGAGATGCGAGG 47957

TeratornLC199500 **CTTCGTCACCGGCATGTGCCTGTGGAGCATGGAGTTCAGGAAGGCGTCGGGTTTGTCCCTGTCTATAAGCCTTTCGATCGAGTACCTAAACACCAGGAGCCTACCCACGCAGACTTGGGG** 43268
WF-Teratorn CTTCGTCACCGGCATGTGCCTGTGGAGCATGGAGTTCAGGAAGGCGTCGGGTTTGTCCCTGTCTATAAGCCTTTCGATCGAGTACCTAAACACCAGGAGCCTACCCACGCAGACTTGGGG 48077

TeratornLC199500 **CGCTGTCCCGCCCGCGAGGGCTCGCGTGAAGGCTCTCACCTCCTCCAAGAATCTTGCGTCGTTTGGGATGCTCGAGCTGTACCTCTGCACCACTTTGTGCCTTCTCCAGTTGTCCTCTGC** 43388
WF-Teratorn CGCTGTCCCGCCCGCGAGGGCTCGCGTGAAGGCTCTCACCTCCTCCAAGAATCTTGCGTCGTTTGGGATGCTCGAGCTGTACCTCTGCACCACTTTGTGCCTTCTCCAGTTGTCCTCTGC 48197

TeratornLC199500 **GAGCCTCGAGACCAGGCGCTCGAAGAGTTCCTTGCCTTGGTCCCGACTCTCGGTGTCCGTGATCCTCCCTATCGCCTCGGCCAGGGACTTCAGCGTCAGACCGGTGATCGCCTCGGGCAG** 43508
WF-Teratorn GAGCCTCGAGACCAGGCGCTCGAAGAGTTCCTTGCCTTGGTCCCGACTCTCGGTGTCCGTGATCCTCCCTATCGCCTCGGCCAGGGACTTCAGCGTCAGACCGGTGATCGCCTCAGGCAG 48317

TeratornLC199500 **CGCGGTCGTCTGCTTTCCGCCCTGGCAGGCGGGCACGTACATCGAGCCCTGCAGTTCCTTGACCTTTTTCCGATCTATCTTGTCGGGGTTCAGCGTATCGCTGCGAATGTCTAACCACTC** 43628
WF-Teratorn CGCGGTCGTCTGCTTTCCGCCCTGGCAGGCGGGCACGTACATCGAGCCCTGCAGTTCCTTGACCTTTTTCCGATCTATCTTGTCGGGGTTCAGCGTATCGCTGCGAATGTCTAACCACTC 48437

TeratornLC199500 **GAACGCAGGGTTCAGTTCTCCCTTGGGGTCTCTCATGACGATCATAGCGTTGTTCAGCGACACAGCTTTCTGCCTGACGAGCTGAAGTCTCATGAAGGCGTTGTAGGCCTCCTCTCCAAG** 43748
WF-Teratorn GAACGCAGGGTTCAGTTCTCCCTTGGGGTCTCTCATGACGATCATAGCGTTGTTCAGCGACACAGCTTTCTGCCTGACGAGCTGAAGTCTCATGAAGGCGTTGTAGGCCTCCTCTCCAAG 48557

TeratornLC199500 **CTCGTAGTCGATGGCGGTCCAGTCGTGTGGCTCCTCGTCGCCGTCATCGCCCTCGTCAGGCTCGCTGTCCACACTGGTGTCGGAATGCTCGTCTCCGCTCGCCCCGGGACTGCGGAACAC** 43868
WF-Teratorn CTCGTAGTCGATGGCGGTCCAGTCGTGTGGCTCCTCGTCGCCGTCATCGCCCTCGTCAGGCTCGCTGTCCACACTGGTGTCGGAATGCTCGTCTCCGCTCGCCCCGGGACTGCGGAACAC 48677

TeratornLC199500 **GCTCGCGTCCGCCTCCACGAAACGCAGTAGACCACCGCCGTAAACGCACTCGGCCGGGTCGCCGTCCTCGCGACCGCGACGTTTACCCGACTCGCGTCCAGCGGCCACAAAGCTAGAAAT** 43988
WF-Teratorn GCTCGCGTCCGCCTCCACGAAACGCAGTAGACCACCGCCGTAAACGCACTCGGCCGGGTCGCCGTCCTCGCGACCGCGACGTTTACCCGACTCGCGTCCAGCGGCCACAAAGCTAGAAAT 48797

TeratornLC199500 **TCCGGGAGAGTCGCCCCCGTTCATGTCCAACACGAAACAGAAAAGCCTTCCGTGCGCAAAGTCCTCCCTGACGGGGGCCGACATGCTCGAAAAGTCCGGGTCCTCCAAGGTCAACTGGAG** 44108
WF-Teratorn TCCGGGAGAGTCGCCCCCGTTCATGTCCAACACGAAACAGAAAAGCCTTCCGTGCGCAAAGTCCTCCCTGACGGGGGCCGACATGCTCGAAAAGTCCGGGTCCTCCAAGGTCAACTGGAG 48917

TeratornLC199500 **GACTTGTCCTTTGCAGCAATAGGTGAGGTAGAGGAGCAGCTGCACCAAAGACCTTTCGCAGGGCGGCAGAGCCCGAAAAGCCAGCTCGACCGTCTCGTCGTCGTGCAGCATTTCGGTGTC** 44228
WF-Teratorn GACTTGTCCTTTGCAGCAATAGGTGAGGTAGAGGAGCAGCTGCACCAAAGACCTTTCGCAGGGCGGCAGAGCCCGAAAAGCCAGCTCGACCGTCTCGTCGTCGTGCAGCATTTCGGTGTC 49037

TeratornLC199500 **CAACGCGAGGAACAGAGCTCTCCTGTCGGTGGGGAGTTCCACCATTCGCACCACTGCGGTGTTAGATTCCCTGTGGGGCAACGCCGCTCGCTCCTTCTGCTGGGCCTTTTTGTCAGACTC** 44348
WF-Teratorn CAACGCGAGGAACAGAGCTCTCCTGTCGGTGGGGAGTTCCACCATTCGCACCACGGCGGTGTTAGATTCCCTGTGGGGCAACGCCGCTCGCTCCTTCTGCGGGGCCTTTTTGTCAGACTC 49157

TeratornLC199500 **CAT**GCTCGTGGTGTGCGCGAACCTGCTGCCAACACTACATCTGGATCTGCGTGTGGCTCACTCACTCACGAGCTCATCGGTAGGCTGGCCAAGTGAGCAGTCTGACTTCAACTGAGGCGG 44468
WF-Teratorn CATGCTCGTGGTGTGCGCGAACCTGCTGCCAACACTACATCTGGATCTGCGTGTGGCTCACTCACTCACGAGCTCATCGGTAGGCTGGCCAAGTGAGCAGTCTGACTTCAACTGAGGCGG 49277

TeratornLC199500 TGGGAAACGCTCCGAGTCTGGAGGCAGGGCATCCCCGTCAGACTTTGCTGTGGGTTGGACACTTTCAGAGGCGGCCTCTGGCTCACTCACTCACGAGCTCATCGGTAGGCTGGCCAAGTG 44588
WF-Teratorn TGGGAAACGCTCCGAGTCTGGAGGCAGGGCATCCCCGGCAGATTTTGCTGTGGGTTGGACACTTTCAGAGGCGGCCTCTGGCTCACTCACTCACGAGCTCATCGGTAGGCTGGCCAAGTG 49397

TeratornLC199500 AGCAGTCTGACTTCAACTGAGGCGGTGGGAAACGCTCAGAGTCTGGAGGCAGGGCATCCCCG---------------------------------------------------------- 44650
WF-Teratorn AGCAGTCTGACTTCAACTGAGGCGGTGGGAAACGCTCCGAGTCTGGAGGCAGGGCATCCCCGGCAGATTTTGCTGTGGGTTGGACACTTTCAGAGGCGGCCTCTGGCTCACTCACTCACG 49517

TeratornLC199500 ------------------------------------------------------------------------------------------------------------------------ 44650
WF-Teratorn AGCTCATCGGTAGGCTGGCCAAGTGAGCAGTCTGACTTCAACTGAGGCGGTGGGAAACGCTCAGAGTCTGGAGGCAGGGCATCCCCGGCAGATTTTGCTGTGGGTTGGACACTTTCAGAG 49637

TeratornLC199500 ------------------------------------------------------------------------------------------------------------------------ 44650
WF-Teratorn GCGGCCTCTGGCTCACTCACTCACGAGCTCATCGGTAGGCTGGCCAAGTGAGCAGTCTGACTTCAACTGAGGCGGTGGGAAACGCTCAGAGTCTGGAGGCAGGGCATCCCCGGCAGATTT 49757

TeratornLC199500 ------------------------------------------------------------------------------------------------------------------------ 44650
WF-Teratorn TGCTGTGGGTTGGACACTTTCAGAGGCGGCCTCTGGCTCACTCACTCACGAGCTCATCGGTAGGCTGGCCAAGTGAACAGTCTGACTTCAACTGAGGCGGTGGGAAACGCTCAGAGTCTG 49877

TeratornLC199500 ------------------------------------------------------------------------------------------------------------------------ 44650
WF-Teratorn GAGGCAGGTCATCCCCGTCAGTTTTTTGCTGTGGGTTGGACACTTTCAGAGGCGGCCTCTGGCTCACTCACTCACGAGCTCATCGGTAGGCTGGCCAAGTGAACAGTCTGACTTCAACTG 49997

TeratornLC199500 -------------------------------------------TCAGTTTTTTGCTGTGGGTTGGACACTTTCAGAGGCGCTCACAGCCGGCCTCTTTTCGATCAGATTTTGCTGTGGGT 44727
WF-Teratorn AGGCGGTGGGAAACGCTCAGAGTCTGGAGGCAGGTCATCCCCGTCAGTTTTTTGCTGTGGGTTGGACACTTTCAGAGGCGCCCACAGCCAGCCTCTTTTCGATCAGATTTTGCTGTGGGT 50117

TeratornLC199500 TGGACACTTTCAGAGGCGGCCTCTGGCTCACTCACTCACGAGCTCATCGGTAGGCTGGCCAAGTGAGCAGTCTGACTTCAACTGAGGCGGTGGGAAACGCTCAGAGTCTGGAGGCAGGGC 44847
WF-Teratorn TGGACACTTTCAGAGGCGGCCTCTGGCTCACTCACTCACGAGCTCATCGGTAGGCTGGCCAAGTGAGCAGTCTGACTTCAACTGAGGCGGTGGGAAACGCTCAGAGTCTGGAGGCAGGGC 50237

TeratornLC199500 ATCCCCGTCAGTTTTTTGCTGTGGGTTGGACACTTTCAGAGGCTGCCTCTTTTCGATCGGATCGTTTTGCTGGCGGTCGGGGCTGGTCTTTGCTACGTCGGAGATTTTAGATCGTGGCCG 44967
WF-Teratorn ATCCCCGTCAGTTTTTTGCTGTGGGTTGGACACTTTCAGAGGCTGCCTCTTTTCGATCGGATCGTTTTGCTGGCGGTCGGGGCTGGTCTTTGCTACGTCGGAGATTTTAGATCGTGGCCG 50357

TeratornLC199500 GACCCTTTCAGCGGCGGTCGGAGTCCTTCTGTGTAGCAAGGGGAACCTGTGTTGAGAAGAAGAAAGAAAAGAGGTGAATAAGAAAGAAGAAGAAGAAAATAGAAGAGAAAAGGAATTTAA 45087
WF-Teratorn GACCCTTTCAGCGGCGGTCGGAGTCCTTCTGTGTAGCAAGGGGAACCTGTGTTGAGAAGAAGAAAGAAAAGAGGTGAATAAGAAAGAAGAAGAAGAAAATAGAAGAGAAAAGGAATTTAA 50477

TeratornLC199500 AAATAAATAAAATAAAATAAAAATAAACCAAACACCTTGTGTCCAGTCTAAGTTCACTGTACCCCCCATTTTTTTTTTTTTCTGAGGAGAACG**ATGGATAGGCCCATTCCTTTCGTGAAT** 45207
WF-Teratorn AAATAAATAAAATAAAA-----ATAAACCAAACACCTTGTGTCCAGTCTAAGTTCACTGTACCCCCCATTTTTTTTTT---CTGAGGAGAACGATGGATAGGCCCATTCCTTTCGTGAAT 50589

TeratornLC199500 **TTGTTCCCCGACATCTTTCCGAAACGTAGAGCAGTATCGCCCTCGGAAAGAGGTTTGATCGGGCTCATGGTCAAGTCTGTGATGTTTCACAGGTTGGAGGAGGCACTCTTGTGCAGGGGC** 45327
WF-Teratorn TTGTTCCCCGACATCTTTCCGAAACGTAGAGCAGTATCGCCCTCGGAAAGAGGTTTGATCGGGCTCATGGTCAAGTCTGTGATGTTTCACAGGTTGGAGGAGGCACTCTTGTGCAGGGGC 50709

TeratornLC199500 **TTGGTGCTGCAGGCTCACAACGGCACCACGGTGTACACCACAGTGTGTTTCAAAGTGGTGGGCATAATGCCCGTGGTGGGCTATGACTTGGAATTGGACTCTGACCAGGGCGACGCCTGT** 45447
WF-Teratorn TTGGTGCTGCAGGCTCACAACGGCACCACGGTGTACACCACAGTGTGTTTCAAAGTGGTGGGCATAATGCCCGTGGTGGGCTATGACTTGGAATTGGACTCTGACCAGGGCGACGCCTGT 50829

TeratornLC199500 **GAGAGCTGGGAGCGGGTGAGTCGAGAGTGGCGCAGACACTTTGCTGCGGAGCCGGCTCACGTGAGAGTGGTGCACGACGACGTGAAAAGGGAAGTGAAGGCCGCTGCGTTCCCGCTGGGG** 45567
WF-Teratorn GAGAGCTGGGAGCGGGTGAGTCGAGAGTGGCGCAAACACTTTGCTGCGGAGCCGGCTCACGTGAGAGTGGTGCACGACGACGTGAAAAGGGAAGTGAAGGCCGCTGCGTTCCCGCTGGGG 50949

TeratornLC199500 **AGCATAGCGCTGGCCAAAAAATATATGGAGACCGTAAAAGGAGCTCTCACGTTTGGGGATTTCCACCCAGAGAGCGGCTACTACGTTGCACGGGACGGCAACACGCCGGCCCTGTGCTGC** 45687
WF-Teratorn AGCATAGCGCTGGCCAAAAAATATATGGAGACCGTAAAAGGAGCTCTCACGTTTGGGGATTTCCACCCAGAGAGCGGCTACTACGTTGCACGGGACGGCAACACGCCGGCCCTGTGCTGC 51069

TeratornLC199500 **GATGTCATGTACGGACGAGTGTATTTCTTGCCCGAGGAAAAGTGTGGCGGGACCCTGGTCCCAAGCGAGCTCGTGTACGACCACGAGAGGTCCGAAGAGGAGGCCAAGCTGTTTGACACC** 45807
WF-Teratorn GATGTCATGTACGGACGAGTGTATTTCTTGCCCGAGGAAAAGTGTGGCGGGACCCTGGTCCCAAGCGAGCTCGTGTACGACCACGAGAGGTCCGAAGAGGAGGCCAAGCTGTTTGACACC 51189

TeratornLC199500 **GTGCACGCGCCGCACGTCTACAGGGCCGCCTGTCTTGACATCGAGACCGTGGTGGACGAGGCCTTCCGCGATCGCTCCCTGAGGTGCGAGTTTTTCGCCCACAGGTTCCCCTACTGCACG** 45927
WF-Teratorn GTGCACGCGCCGCACGTCTACAGGGCCGCCTGTCTTGACATCGAGACCGTGGTGGACGAGGCCTTCCGCGATCGCTCCCTGAGGTGCGAGTTTTTCGCCCACAGGTTCCCCTACTGCACG 51309

TeratornLC199500 **GAGCGCACGGTTTCCGATTTGACGGCGTACAGGAGCAAACTGGTCAAGATCCTGAGCGACTTGAAGAAAGGCCCGCTCAGAGGGGCTTGTCGGGTGTCCATACCGGCCCTGGCCAGTGAC** 46047
WF-Teratorn GAGCGCACGGTTTCCGATTTGACGGCGTACAGGAGCAAACTGGTCAAGATCCTGAGCGACTTGAAGAAAGGCCCGCTCAGAGGGGCTTGTCGGGTGTCCATACCGGCCCTGGCCAGTGAC 51429

TeratornLC199500 **ATGCCGGGACAGGAGCACGAGATCACATGCGTGTCCTTGGTGCTGATGAACTCGCACCTCCCAAAGCCCGAGCCGGGGAGTCACAGGAAACAACTGATAGTGGTCTACAACAAACTGAAG** 46167
WF-Teratorn ATGCCGGGACAGGAGCACGAGATCACATGCGTGTCCTTGGTGCTGATGAACTCGCACCTCCCAAAGCCCGAGCCGAGGAGTCACAGGAAACAACTGATAGTGGTCTACAACAAACTGAAG 51549

TeratornLC199500 **GTAAAGGGCCAGGGACGACCGAAACCCGATGGCGCGCTCGTGGAGGCCGTGGGACTGGAGTCCGCGTCCAGGATCCTTTTCTACCCCTGCGCCGGAGAGCTCGCGCTCCTAGAGCAGACC** 46287
WF-Teratorn GTAAAGGGCAAGGGACGCCCGAAACCCGATGGCGCGCTCGTGGAGGCCGTGGGACTGGAGTCCGCGTCCAGGATCCTTTTCTACCCCTGCGCCGGAGAGCTCGCGCTCCTAGAGCAGACC 51669

TeratornLC199500 **CTGCGGCTGCTCGATCGCCACGGCGTGGAGCTCCTGTACGTGTACAATGCGGAATTCGACCTGAGAGTCATCGACCAGCGCGTGCATTTCTACGCGGACTCCAACTACGCAGAGCGATCG** 46407
WF-Teratorn CTGCGGCTGCTCGATCGCCACGGCGTGGAGCTCCTGTACGTGTACAATGCGGAATTCGACCTGAGAGTCATCGACCAGCGCGTGCATTTCTACGCGCACTCCAACTACGCAGAGCGATCG 51789

TeratornLC199500 **GACGAGGATACCCGCAGAAGGTGCGCTGAAATCCTCCGGTCTTGGCACGGGCTCTTTGTCACCAGGCACCTCTCGGAAGACCTGCCTCCCGTTTTCCGTTTCGAGTCGGTGCGCCACTTG** 46527
WF-Teratorn GACGAGGATACCCGCAGAAGGTGCGCTGAAATCCTCCGGTCTTGGCACGGGCTCTTTGTCACCAGGCACCTCTCGGAAGACCTGCCTCCCGTTTTCCGTTTCGAGTCGGTGCGCCACTTG 51909

TeratornLC199500 **GAAATCTACAAGGAGATGCTGACCGCCGTGGGCTCGAGCCTGACCGGAGGCAAGCTCACCGAGCGTGAGGTCGAAGCGGCGTGTCGACACGTGGACAAGTTCAACAAGGACAAGGGCAAG** 46647
WF-Teratorn GAAATCTACAAGGAGATGCTGACCGCCGTGGGCTCGAGCCTGACCGGAGGCAAGCTCACCGAGCGCGAGGTCGAAGCGGCGTGTCGACACGTGGACAAGTTCAACAAGGACAAGGGCAAG 52029

TeratornLC199500 **CTGGGCCACTTTAAGATGATCGGCTGCGGCATGACAGTCGTGGACCTGTACAGGATGGCGGGCACCAGGGAGATCAGATTCGCCTGCACGAGCATGAAGCTGAACGACGTGGCGCCCTTC** 46767
WF-Teratorn CTGGGCCACTTTAAGATGATCGGCTGCGGCATGACAGTCGTGGACCTGTACAGGATGGCGGGCACCAGGGAGATCAGATTCGCCTGCACGAGCATGAAGCTGAACGACGTGGCGCCCTTC 52149

TeratornLC199500 **GTGATTTCCAGAGACCGAGAGCTGCGCGGACTGCCTCCCAAAGACCCACGTAAGCTGCGGAAGCTGACCGACGTGCCCTACTCGATGATGGACGGCATGATCCACCGCGGCGGGACCGAA** 46887
WF-Teratorn GTGATTTCCAGAGACCGAGAGCTGCGCGGACTGCCTCCCAAAGACCCACGTAAGCTGCGGAAGCTGACCGACGTGCCCTACTCGATGATGGACGGCATGATCCACCGCGGCGGGACCGAA 52269

TeratornLC199500 **CTGTTCGCCGTGCTGCTGTACAACTTGGTGGACTCGCAGCTGTGTTCGAGGATGGCCAAGGCGCTCAAGCCCGCGTCCGCGCTGTTCCACAGGTGCAGGATCACGCTCAACATAGACGTG** 47007
WF-Teratorn CTGTTCGCTGTGCTGCTGTACAACTTGGTGGACTCGCAGCTGTGTTCGAGGATGGCCAAGGCGCTCAAGCCCGCGTCCGCGCTGTTCCACAGGTGCAGGATCACGCTCAACATAGACGTG 52389

TeratornLC199500 **GTGGTGCACGGGCGGGGAGACAACTTTGGCGGTTTCGTTCAGAGCATCCACTCGGTCCAGATGCCCCAGCTCAAGTACGTGCTCGACACTCTGAGGGTCAGGGCGGGACCCTCGGGCAGA** 47127
WF-Teratorn GTGGTGCACGGGCGGGGAGACAACTTTGGCGGTTTCGTGCAGAGCATCCACTCGGTCCAGATGCCCCAGCTCAAGTACGTGCTCGACACTCTGAGGGTCAGGGCGGGACCCTCGGGCAGA 52509

TeratornLC199500 **GAGCTGAGCAGCCGCGAGCCCTGGGACGCCTCACCGCCGGGGCTCGACGGAGCGGGCGAGGACACGTGGAAGGGAGGGTCCGTGTGCGAACCCCTCTCGGGCCTCCACTACTCTGGGCCC** 47247
WF-Teratorn GAGCTGAGCAGCCGCGAGCCCTGGGACGCCTCACCGCCGGGGCTCGACGGAGCGGGCGAGGACACGTGGAAGGGAGGGTCTGTGTGCGAACCCCTCTCGGGCCTCCACTACTCTGGGCCC 52629

TeratornLC199500 **GGCATGGGCTTGGAGCTGGCCTTCGATTTCGCCAGCATGTACCCGTCCATCATGTGCGCGCTCAACATTTCTCCAGAGACCACCGTGCCCTGGCCTCCGCGCTCCTTCCCCCACGATCTC** 47367
WF-Teratorn GGCATGGGCTTGGAGCTGGCCTTCGATTTCGCCAGCATGTACCCGTCCATCATGTGCGCGCTCAACATTTCTCCAGAGACCACCGTGCCCTGGCCCCCGCGCTCCTTCCCCCACGATCTC 52749

```
TeratornLC199500 TCCGGATGGGTCGTGTACAAGTGGGAGGCCGAAGGCTTCGAGTACGCGACGCTGATCCTCCGGTACGACGCGGACCGAGCCGTCTTCGAGCGGGCGCCGGCCGTCTTCTCCTCCTCCGTG 47487
WF-Teratorn      TCCGGATGGGTCGTGTACAAGTGGGAGGCCGAAGGCTTCGAGTACGCGACGCTGATCCTCCGGTACGACGCGGACCGAGCCGTCTTCGAGCGGGCGCCGGCCGTCTTCTCCTCCTCCGTG 52869

TeratornLC199500 GAGCATTACCTCAACAGGCGGGACGAGTTCAAGGCCAAGCTTAAGGACCCGGCCCTCTGCGACGCGGAGCGAGCGTACTACAAGGTGCAGGAGGCAGAGTGCAAAGTCATGGCCAATTCG 47607
WF-Teratorn      GAGCATTACCTCAACAGGCGGGACGAGTTCAAGGCCAAGCTTAAGGACCCGGCCCTCTGCGACGCAGAGCGAGCGTACTACAAGGTGCAGGAGGCAGAGTGCAAAGTCATGGCCAATTCG 52989

TeratornLC199500 TTTTACGGGACCGCTCCCCCACCCTGCGGGTCCCTCATATCGGGACACGGCAGGAGACAGATATCGGTGGTGAACGACTGCGTGTCGACTTTTTACAAAAACCGCTGCCCGGTCATGTAC 47727
WF-Teratorn      TTTTACGGGACCGCTCCCCCACCCTGCGGGTCCCTCATATCGGGACACGGCAGGAGACAGATATCGGTGGTGAACGACTGCGTGTCGACTTTTTACAAAAACCGCTGCCCGGTCATGTAC 53109

TeratornLC199500 GGGGACACGGACAGCGTGATGGTGTCCGCGGGGTACGGGCCGGACGACCTGCCCGAGAGCGACGTGTCGGACGAACAGCCGGGCGACGGGGAGTCCCCGGAGCTCGAGACCGAGCGGCTT 47847
WF-Teratorn      GGGGACACGGACAGCGTGATGGTGTCCGCGGGGTACGGGCCGGACGACCTGCCCGAGAGCGACGTGTCGGACGAACAGCCGGGCGACGGGGAGTCCCCGGAGCTCGAGACCGAGCGGCTT 53229

TeratornLC199500 CGGAACTTCGCGGCGGCCTCCCTGGCGGCCTTGGGGGCGAAATTCGAACGCGCTGCCTCCGAGGTACCGGGTTTCCTCCGATACGTGCACCTGGCGCTGGTGGAGGACACTCTGAAGAGG 47967
WF-Teratorn      CGGAACTTCGCGGCGGCCTCCCTGGCGGCCTTGGGGGCGAAATTCGAACGCGCTGCCTCCGAGGTACCGGGTTTCCTCCGATACGTGCACCTGGCGCTGGTGGAGGACACTCTGAAGAGG 53349

TeratornLC199500 ATGTACACCATAGAGAGGGGCAACGAGCCGGTGGCGGTGATCAGAGACCCCGAGGGCGCCTTCACGGAGGAGGGCTACCCCGTGTACGTGGCGCAAACTCCGGACTCCCGGAGCCTCCTG 48087
WF-Teratorn      ATGTACACCATAGAGAGGGGCAACGAGCCGGTGGCGGTGATCAGAGACCCCGAGGGCGCGTTCACGGAGGAGGGCTACCCCGTGTACGTGGCGCAAACTCCGGACTCCCGGAGCCTCCTG 53469

TeratornLC199500 GACGTGACGAGGCCCTTCGTGAAGGACCGGCGGGTGAAGCTGGAATACGAGTACAGCAGCTCGGTGTACTGCCACGTGGCGAAGAAGAGCTACGTGTCTCTCACTCACAACCTTGATGGG 48207
WF-Teratorn      GACGTGACGAGGCCCTTCGTGAAGGACCGGCGGGTGAAGCTGGAATACGAGTACAGCAGCTCGGTGTACTGCCACGTGGCGAAGAAGAGCTACGTGTCTCTCACTCACAACCTTGATGGG 53589

TeratornLC199500 AGGGGCCGCATGTCCTCCTCGACGGTCAAGATCAGAGGACTCTCCGCTGTAAAAAGCCTGCGCTCGCCCTGCGATTCCGCCGTCGCGGACACTTTCGTGGCCTGCGTCATGAGGGGAGAC 48327
WF-Teratorn      AGGGGCCGCATGTCCTCCTCGACGGTCAAGATCAGAGGACTCTCCGCTGTAAAAAGCCTGCGCTCGCCCTGCGATTCCGCCGTCGCGGACACTTTCGTGGCCTGCGTCATGAGGGGAGAC 53709

TeratornLC199500 TGCGTCAAGCTGGAGCCCCACGGACTCTCCTGCTTCTCCACCCTACCGTGGCATCGGCTATCGGAGGGGGATCTCGTACTGTACCCGGAGAGCGAGCTCACCGTGGACGGCCGCGGGCGG 48447
WF-Teratorn      TGCGTCAAGCTGGAGCCCCACGGACTCTCCTGCTTCTCCACCCTACCGTGGCATCGGCTATCGGAGGGGGACCTCGTGCTGTACCCGGAGAGCGAGCTCACCGTGGACGGCCGCGGGCGG 53829

TeratornLC199500 TGGGAGCAGCTGGATCGAGCGCAAGCCCTCCTCGTAGGCCACAGGGTGCTGCGCGTAACAGAGCTGGAACTGGACGGCGGGTTTTTGGCCGTCTCCGTCACGTTGGCGAGGCCGGGTTCC 48567
WF-Teratorn      TGGGAGCAGCTGGATCGAGCGCAAGCCCTCCTCGTAGGCCACAGGGTGCTGCGCGTAACGGAGCTGGAACTGGACGGCGGATTTTCGGCCGTCTCCGTCGCGTTGGCGAGGCCGGGTTCC 53949

TeratornLC199500 GGCGTCGTCACGGCAAGAGCCCTGTACAAGGACGGAACGTACTGCATGAACCACCTGTTCTCGAGCAGACACGGAGTGCTCAGGGACCTGCTCGCGTGCCGAGCCTCCGAGCTCATCGCG 48687
WF-Teratorn      GGCGTCGTCACGGCGAGAGCCCTGTACAAGGACGGAGCGTACTGCATGAACCACCTGTTCTCGAGCAGACACGGAGTGCTCAGGGACCTGCTCGCGTGCCGAGCCTCCGAGTTCATCGCG 54069

TeratornLC199500 TCCAAGATGGGAGCGGGCTTCTTTCCCTGGAGCGCCCTCGTCAAATCGGCGAAAAACAAGCACTTCAGCCAACAGACGCTGGAACGTTTCAAAGCGAGCAAGTCCGGCGTGAAGACGACG 48807
WF-Teratorn      TCCAAGATGGGAGCGGGCTTCTTTCCCTGGAGCGCCCTCGTCAAATCGGCGAAAAACAAGCACTTCAGCCAACAGACTCTGGAACGTTTCAAAGCGAGCAAGTCCGGCGTGAAGACGACG 54189

TeratornLC199500 TACGTGGAGATCGTCCGCACGTGGCTGCAGAGGATAACCGGGCTCTCTGTGAAACCCGTGGAGTGCGAGGAGTTCTACAAGTGCAGCCCCATCGAAGCCGCCTTCTACAGGTACCCGCTC 48927
WF-Teratorn      TACATGGAGATCGTCCGCACGTGGCTGCAGAGGATAACCGGGCTCTCTGTGAAACCCGTGGAGTGCGAGGAGTTCTACAAGTGCAGCCCCATCGAAGCCACCTTCTACAGGTACCCGCTC 54309

TeratornLC199500 AACCTGCAGGCCACCTCGGGGTGCACGACCGCGATGTTCGGAGGTTCCCTCGCCTGCCTCTCGGACAAACACGCCCGTGGCTGCGACGGCGAGGCCGGCGCCCGGAGCGCCTCGGCGGGC 49047
WF-Teratorn      AACCTGCAGGCCACCTCGGGGTGCACGACCGCGATGTTCGGAGGGTCCCTCGCCTGCCTCTCGGACAAACACGCCCGAGGCGGCGACGGCGAAGCCGGCGCCCGGAGCGCCTCGGCGGGC 54429

TeratornLC199500 CGTTTGGTCCCTCACCCCTGTCAGCCTCAGAGCCAGTGCTTCGCCGACGCCAGGGTCTTGACGGCGCACAAGGCAGACCGTGAGCTCGTCGGCCGATCCGGGAAGCTGGTGGCCCACTGC 49167
WF-Teratorn      CGTTTGGTCCCTCACCCCTGTCAGCCTCAGAGCCAGTGCTTCGCCGACGCCAGGGTCTTGACGGCGCACAAGGCAGACCGCGAGCTCGTCGGCCGATCCGGGAAGCTGGTGGCCCACTGC 54549

TeratornLC199500 ATCGTGGACTACTGCCTTCCGCGACGCATGTACGGGAGCGGCGCGGACGAGCTCGAGATCCTGGCAGACTGCAGGAGGCACCTGATCCTGTCATTCTCCGCCGTCCGAGAGCGCCTGAGC 49287
WF-Teratorn      ATCGTGGACTACTGCCTTCCGCGACGCATGTACGGGAGCGGCGCGGACGAGCTCGAGATCCTGGCAGACTGCAGGAGGCACCTGATCCTGTCATTCTCCGCCGTCCGAGAGCGCCTGAGC 54669

TeratornLC199500 AGAGCTACGGCCACGTTCAACGCGCTGATGAACGTCTGTCCGGAACACATGGAGCCGGCGTGCGGCCTCCGCGAGCTCGAGCGCCTGAGGGCTCTCCGTCGGGCTGCGCCTCGCCTCACG 49407
WF-Teratorn      AGAGCTACGGCCGCGTTCAACGCGCTGATGAACGTCTGTCCGGAACACATGGAGCCGGCGTGCGGCCTCCGCGAGCTCGAGCGCCTGAGGGCTCTCCGTCGGGCTGCGCCTCGCCTCACG 54789

TeratornLC199500 GGGGACAAACTGGAAGCCTGTCGCAGGGTCCTGGAGGAAGATCTCGCGCTGGTGCCCCGTGAGGTGTACGCCGGCCTGATCGGAGAGCTCGGCCGCTCGCTGGCCTTGCTGACTCGCGAA 49527
WF-Teratorn      GGGGACAAACTGGAAGCCTGTCGCAGGGTCCTGGAGGACGATCTCGCGCTGGTGCCCCGCGAGGTGTACGCCGGCCTGATCGGAGAGCTCGGCCGCTCGCTGGCCTTGCTGACTCGCGAA 54909

TeratornLC199500 GGCGTCGTGCGCGTGGGAACCTCGGACGAAGGCGAGGGGCTCTCGAGCGAGCGCATAGCCCTCCGCCTGCCGACGTGCCTGTTGGTCGATGCGGGCCTCGCCGGCTCGGAGGGATCGCTG 49647
WF-Teratorn      GGCGTCGTGCGCGTGGGAACCTCGGACGAAGGCGAGGGGCTCTCGGGCGAGCGCATAGCCCTCCGCCTGCCGACGTGCCTGTTGGTCGACGCGGGCCTCGCCGGCTCGGAGGGATCGCTG 55029
```

TeratornLC199500 **TCCATCTCGAAGAGCGAAGCCTCGGAGAGGGCCTTGGTGGACCTCGCGGGGCTTCTCTACCAGACGTTGCTCGCGCTGCACGTGAGATCGCCCGACGCGTTTTCCTCTGTCGTCGCCAGG** 49767
WF-Teratorn TCCATCTCGAAGAGCGAAGCCTCGGAGAGGGCCCTGGTGGACCTCGCGGGGCTTCTCTACCAGACATTGCTCGCGCTGCACGTGAGATCGCCCGACGCGTTTTCCTCTGTCGTCGCCAGG 55149

TeratornLC199500 **GAGCCCGATTCGACGCTGCGCATCATGCCGTACACGACAGAAGACCCCGATGCCCTGAGAGCCTGGATAGAGGACTTCAAGAGCAAAAAGTGGCTCGTGCGTCCAGCGCCGCCAACGGGA** 49887
WF-Teratorn GAGCCCGATTCGACGCTGCGCATCATGCCGTACACGACAGAAGACCCCGATGCCCTGAGAGCCTGGATAGAGGACTTCAAGAGCAAAAAGTGGCTCGTGCGTCCAGCGCCGCCAACGGGA 55269

TeratornLC199500 **CAGAAAAGGACGGCCCGCGACAGGGCCTCTCTCGGAGGGACCGCGGGAGTCACGTATCACTGCGCCGGTTGCAAAGACTTTTTCAGAGACCTGTGCGCAAACAAGGTGTCGGGCCGGACG** 50007
WF-Teratorn CAGAAAAGGACGGCCCGCGACAGGGCCTCTCTCGGAGGGACTGCGGGAGTCACGTATCACTGCGCCGGTTGCAAAGACTTTTTCAGAGACCTGTGCGCAAACAAGGTGTCGGGCCGGACG 55389

TeratornLC199500 **CTCCTGCGCAGTGTGTACGGAGACAACGGTAAGAGGAAGCTGGGGCGTTGCGTCAACGAAGCCTGGTGGCTCCGTTGA**GAAAGGGGACGGCGCCCGAGCGGCAAACGGGTCGCGGTCCGG 50127
WF-Teratorn CTCCTGCGCAGTGTGTACGGAGACAATGGTAAGAGGAAGCTGGGGCGTTGCGTCAACGAAGCCTGGTGGCTCCGTTGAGAAAGGGGACGGCGCCCGAGCGGCAAACGGGTCGCGGTCCGG 55509

TeratornLC199500 GGACGTGAACTGGGCAGACTAACTGAACCGAGGTGCGGAGGACCCCGAGCGAGCGAGCGACCAGGAGCGGACGAG**ATGGCCTTCAGTCAGAAGACCTACGTCTTTTGTCACGACGTGGCT** 50247
WF-Teratorn GGACGTGAACTGGGCAGACTAACTGAACCGAGGTGCGGAGGACCCCGAGCGAGCGAGCGACCAGGAGCGGACGAGATGGCCTTCAGTCAGAAGACCTACGTCTTTTGTCACGACGTGGCT 55629

TeratornLC199500 **AAGGACAGCGTGTCGAAACAGCCCCTCCTGTTCGCCGTCGACCTGCGGGATCTCATCTCGCCTTTCGAAGGATGCAGCCCCATGTTTGTGATCGAACAGATGAGCTTGCGGGCGTACAAG** 50367
WF-Teratorn AAGGACAGCGTGTCGAAACAGCCCCTCCTGTTCGCCGTCGACCTGCGGGATCTCATCTCGCCTTTCGAAGGATGCAGCCCCATGTTTGTGATCGAACAGATGAGCTTGCGGGCGTACAAG 55749

TeratornLC199500 **AGAGACACCTTCGAAAGGATACCGGCAAAGTTTGCGGTGCTCAACTACGCCAAGAGCAGGGTGGTCACCAACGACCCCTCCAGACTCTCGTCCGCGTGGGAAGTGGAGTTGAACGGAAAG** 50487
WF-Teratorn AGAGACACCTTCGAAAGGATACCGGCAAAGTTTGCGGTGCTCAACTACGCCAAGAGCAGGGTGGTCACCAACGACCCCTCGAGACTCTCGTCCGCGTGGGAAGTGGAGTTGAACGGAAAG 55869

TeratornLC199500 **GATGTCCGGATCAGGGCGGCGGGGACAGACACTGATCCTTTTTGCCGCACGCCGTCCACGCACGACGACGACAACAGACAGGCTCTCTTGAACGTTCTGGCCTTCATGATGTCGGGCGGG** 50607
WF-Teratorn GATGTCCGGATCAGGGCGGCGGGGACAGACACTGATCCTTTTTGCCGCACGCCGTCCACGCACGACGACGACAACAGACAGGCTCTCTTGAACGTTCTGGCCTTCATGATGTCGGGCGGG 55989

TeratornLC199500 **GTGCGAGCCTCGGATGGGGCGAGGCGCTCCCTCGACGCCATGAAAGGCTGTTTGTCCTCGGAACACTTCACCTACGCCTGCGAGCTGTGCAAGGGACAATGTGAGGCGACGGTCATCGTG** 50727
WF-Teratorn GTGCGAGCCTCGGATGGGGCGAGGCGCTCCCTCGACGCCATGAAAGGTTGTTTGTCCTCGGAACACTTCACCTACGCCTGCGAGCTGTGCAAAGGACAATGTGAGGCGACGGTCATCGTG 56109

TeratornLC199500 **ACCCAGCCCGGCTACCCGTCCCTGTCGTTGCCCTTGTGCGTCGACGGCTACGTGTGCGTGCCCGAGCGGTCGGGCTCGACGAGTCCGAAGTCTCAAACGTACAGGAGTGTGATTTCGCTC** 50847
WF-Teratorn ACCCAGCCCGGCTACCCGTCCCTGTCGTTGCCCTTGTGCGTCGATGGCTACGTGTGCGTGCCCGAGCGGTCGGGCTCGACGAGTCCGAAGTCTCAAACGTACAGGAGTGTGATTTCGCTC 56229

TeratornLC199500 **CTCAGGACTGCGTACGAAGACAACACGAGCCTCGTGGACCGAGCGATCCAGGCCGCGTTTGCGAACTCGGGGTTCGGGCGTTATTTGGAACGGGGCAACGGAGAAGTGGAACTGTTTCCC** 50967
WF-Teratorn CTCAGGACTGCGTACGAAGACAACACGAGCCTCGTGGACCGAGCGATCCAGGCCGCGTTTGCGAACTCGGGGGTCGGGCGTTATTTGAAACGGGGCAACAGAGAAGTGGAACTGTTTCCC 56349

TeratornLC199500 **AACCCGCTGATAAACTTTGTCGTTTCCCTGTATGACACGTTGAGCGACCAGGCCGGCAATTTCGTGGCGGTCGACAAACACGAGCCCGGTCTGAAGGTGCTGGTCGAGAGCCCCGCGGAG** 51087
WF-Teratorn AACCCGCTGATAAACTTTGTCGTTTCCCTGTATGACACGTTGAGCGACCAGGCCGGCAATTTCGTGGCGGTCGACAAACACGAGCCCGGTCTGAAGGTGCTGGTCGAGAGCCCCGCGGAG 56469

TeratornLC199500 **AAGGTGGGACACGTGACGGGCTCCCTCACTCTCACACTGCTGTACGGTCAGAGCGTGCTGGACGTGTACAGCAACGTGAAGGCACGGCACCGAGGCACCGACTGCCCACAGGACCTGCTG** 51207
WF-Teratorn AAGGTGGGACACGTGACGGGATCCCTCACTCTCACACTGCTGTACGGTCAGAGCGTGCTGGACGTGTACAGCAACGTGAAGGCACGGCACCGAGGCACCGACTGCCCACAGGACCTGCTG 56589

TeratornLC199500 **GACATGCACACGGCCGTGACTCGGCTCGGCTGCGACGCCTTGCTCAGACGATTCCCCTTGGACTTCTGCTCTAAGAAACGCCACCTCACCACAGTGGATTTGTGCAATTTTGACAACGAC** 51327
WF-Teratorn GACATGCACACGGCCGTGACTCGGCTCGGCTGCGACGCCTTGCTCAGACGATTCCCCTTGGACTTCTGCTCTAAGAAACGCCACCTCACCACAGTGGATTTGTGCAATTTTGACAACGAC 56709

TeratornLC199500 **TAA**GTGTGTCCCGTGGTTTATTTGAGGGTTATGCTTTATTGTGTGTTACCGGTTGCACTGTGCTCTAAACGCCAAGGTGACAATAAAAATCAAAGACAGAAAGAATATGTTTGGTCTCAC 51447
WF-Teratorn TAAGTGTGTCCCGTGGTTTATTTGAGGGTTATGCTTTATTGTGTGTTACCGGTTGCACTGTGCTCTAAACGCCAAGGTGACAATAAAAATCAAAGACAGAAAGAATATGTTTGGTCTCAC 56829

TeratornLC199500 GCACTCATTTATTTTGTGCACACAACCACATGGACAGAGAAATGAAAACGAGAAAAAAAAATAATAACGCAGCCTCTCGCGCTGGGCAGAGCTGACCCACGGAGGCATCTAAAGGTGAAGG 51567
WF-Teratorn GCACTCATTTATTTTGTGCACACAACCACATGGACAGAGAAAAGAAAACGAGAAAAAAAAA-AATAACGCAGCCTCTCGCGCTGGGCAGAGCTGACCCACGGAGGCATCTAAAGGTGAAGG 56948

TeratornLC199500 TGACAGCACAATTGTCTAATATCATGTGCAGACAT**TCACTTGGAAAGAAAAATCAGCCTTTTTAGCCGGCTGAGCCCGTACACCAGGGCCTCCATGTCCATGTGATAAGCGAGGCACTTT** 51687
WF-Teratorn TGACAGCACAATTGTCTAATATCATGTGCAGACATTCACTTGGAAAGAAAAATCAGCCTTTTTAGCCGGCTGAGCCCGTACACCAGGGCCTCCATGTCCATGTGATAAGCGAGGCACTTT 57068

TeratornLC199500 **TTGTCCCTCGGTCCGCACACCACGGTAAACCCACGGATGCCGAACACGACCTTGCACTTGTTGATGAGCAGTCCGTAGCCCCCGTTTCGTTTGCTCAAGTACATGATCTCGAACAAGTAC** 51807
WF-Teratorn **TTGTCCCTCGGTCCGCACACCACGGTAAACCCACGGATGCCGAACACGACCTTGCACTTGTTGATGAGCAGTCCGTAGCCCCCGTTTCGTTTGCTCAAGTACATGATTTCGAACAAGTAC** 57188

TeratornLC199500 **CCGGGACCCCTTTCTTCCTTCACGTCCGTCAGAGCGATCATGGCGCTCGGCCTCCTCTGTCGCTGCAGAGCCGTGAGGGGATACGTCAGCTTTGGCACTTGGGCCAAGTTGGTGTACGTA** 51927
WF-Teratorn CCGGGACCCCTTTCTTCCTTCACGTCCGTCAGCGCGATCATGGCGCTCGGCCTCCTCTGTCGCTGCAGAGCCGTGAGGGGATACGTCAGCTTTGGCACTTGGGCCAAGTTGGTGTACGTA 57308

TeratornLC199500 **CCCCACCAAACCTTGCCCATGGGGCCTCGTATCAGCCTGTGCAGCCGCTCTGGTATCCTCGCGTCGGCGAAAAAGCTCCGTTGCGAGGACGTCACGTTGGAGGGCTTGCGAGCCACCGTT** 52047
WF-Teratorn CCCCACCAAACCTTGCCCATGGGGCCTCGTATCAGCCTGTGCAGCCGCTCTGGTATCCTCGCGTCGGCGAAAAAGCTCCGTTGCGAGGACGTTACGTTGGAGGGCTTGCGAGCCACCGTT 57428

TeratornLC199500 **CGCTGGCATCCCAGTAGCAGCACGATGTCCCTCGGGTCATCGTAGTAGCCCTTGAGCTCCTTGCACACGTAAAGTCCCCTTTCCATTGCGATCTCCGTCGTCAGCACTGCGTCCGAACTG** 52167
WF-Teratorn CGCTCGCATCCCAGTAGCAGCACGATGTCCCTCGGGTCATCGTAGTAGCCCTTGAGCTCCTTGCACACGTAAAGTCCCCTTTCCATCGCGATCTCCGTCGTCAGCACTGCGTCCGAACTG 57548

TeratornLC199500 **CCCTGCATCGTCACGGAGCACAGCGCTCCGATCTTCGGCTCCAGGGGCTCCGAGACCACGACGGCGCATCGCCCCTTCTCGCTCCTCAAGAACGTAGCTATGTCATCTCGAGACACGGAG** 52287
WF-Teratorn CCCTGCTTCGTCACGGAGCACAGCGCTCCGATCTTCGGCTCCAGGGGCTCCGAGACCACGACGGCGCATCGCCCCTTCTCGCTCCTCAAGAACGTAGCTATGTCATCTCGAGACACGGAG 57668

TeratornLC199500 **CCCCACCAGCGATCGTCCAGCCAGTTCAGGAAACCCTTCCAAGAACGCATGACGCTCAACCAGTTCTGGACGAGCGTGTCGGAGTCAGAGTAGAGCTCTTCCATGTTCACTTCAACTGCT** 52407
WF-Teratorn CCCCACCAGCGATCGTCCAGCCAGTTCAGGAAACCCTTCCAAGAACGCATGACGCTCAACCAGTTCTGGACGAGCGTGTCAGAGTCAGAGTAGAGCTCTTCCATGTTCACTTCAACTGCT 57788

TeratornLC199500 **CTCCTCGAGAACATGGTGCTAGGTACGAGCAGCTGGGAGCCATCCTCCCCGTTGATGATACACTCTTCTTCTTTATCCATTTCTACATCTTCATATTCTCTCTCTTCTTTTTCTTTCCTT** 52527
WF-Teratorn CTCCTCGAGAACATGGTGCTAGGTACGAGCAGCTGGGAGCCATCCTCCCCGTTGATGATACACTCTTCTTCTTTATCCATTTCTACATCTTCATATTCTTCCTCTTCTTTTTCTTTCCTC 57908

TeratornLC199500 **CCTTCCCCTACTTTCTCTTCGTCCGCCTCTCCGGGTTTCTGAGACGTGGCGAACTTTGAATCCCGGAGCCCTCTCGCGATGCAGCGGAGCGGCTCTGAGACTGCCTGACGGCACTCGCCG** 52647
WF-Teratorn CCTTCCCCTACTTTCTCTTCGTCCGCCTCTCCGGGTTTCTGAGACGTGGCGAACTTTGAATCCCGGAGCCCTCTCGCGATGCAGCGGAGCGGCTCTGAGACTGCCTGACGGCACTCGCCG 58028

TeratornLC199500 **ACAAAGGTGTGGCGGCAGAACCCATAGCAACCCGAGCCGCTCGGGTCCTTCACGCACACGATGGGTTCGTAGTGGCCCAACTGGTTCTGCACGTATATCCCGGGCAGCTCGTTCGACCGA** 52767
WF-Teratorn ACAAAGGTGTGGCGGCAGAACCCGTAGCAACCCGAGCCGCTCGGGTCCTTCACGCACACGATGGGTTCGTAGTGGCCCAACTGGTTCTGCACGTATATCCCGGGCAGCTCGTTCGACCGA 58148

TeratornLC199500 **GGCCTGCCGTCTGGACTGTAGACTACCCACCCTCCGTGGTTGTGGGTGTACACGTTCACGTCCCAGCCGTTGGCCAGCGCTGCGATTTCCACCTCGGTGGCCCAGACACCCCGATCCTTC** 52887
WF-Teratorn GGCCTGCCGTCCGGACTGTAGACTACCCACCCTCCGTGGTTGTGCGTGTACACGTTCACGTCCCAGCCGTTGGCCAGCGCTGCGATTTCCACCTCGGTGGCCCAGACACCCCGATCCTTC 58268

TeratornLC199500 **GGCCTCCTTTCGAGAATGTACTGCTCCACGCTCGAGTACTGCGAGCGCAGCAAATCTCTGTATTTTTCTTTATTCGAACTCATTTCCCTGACGCACCAGCTTCTGATCTTGTAGTGGTAG** 53007
WF-Teratorn GGCCTCCTTTCGAGAATGTACTGCTCCACGCTCGAGTACTGCGAGCGCAGCAAATCTCTGTATTTTTCTTTATTCGAACTCATTTCCCTGACGCACCAGCTTCTGATCTTGTAGTGGTAG 58388

TeratornLC199500 **TCGTCCCTGCCGCAGACCAAGTAGCTGACGGCCCTGAAGAAACAGTTGCCGTCTCCCATCACCGGCACCGTCGCGCAGGGCCGTCCCAGAGGCACGGGTGAAAAATTCGGCCCGCGGCAC** 53127
WF-Teratorn TCGTCCCTGCCGCAGACCAAGTAGCTGACGGCCCTGAAGAAACAGTTGCCGTCTCCCATCACCGGCACCGTCGCGCAGGGCCGTCCCAGAGGCACGGGTGAAAAATTCGGCCCGCGGCAC 58508

TeratornLC199500 **AGAGCTCTCGGCTTCTCAGGAGCTGCCACAACGTACGCAGCCAGGTCGAGCAGCGTCGCTCCCCTCACCGGATCGAAGGTCATTTTGTTCCCCGGGGCAGATTCGATGTACATGACATCC** 53247
WF-Teratorn AGAGCTCTCGGCTTCTCAGGAGCTGCCACAACGTACGCAGCCAGGTCGAGCAGCGTCGCTCCCCTCACCGGATCGAAGGTCATTTTGTTCCCCGGGGCAGATTCGATGTACATGACATCC 58628

TeratornLC199500 **AAGTCCGAGTCGCTCCCAGCCCTCCCTCCAGATACCGGGGTGCTGTCCAT**TCTAACGAAGAGAGTGTTTCGGTCTGGAAACTCGAGAGGAGCAAATGAGGAAAACGAAA-AAAACAAAAA 53366
WF-Teratorn GAGTCCGAGTCGCTCCCAGCGCTCCCTCCAGATACCGGGGTGATGTCCATTCTAACGAAGAGAGTGTTTCGGTCTGGAAACTCGAGAGAAGCAAATGAGGAAAACGAAACAAAAAACAAA 58748

TeratornLC199500 AAAAAACAAAAAGAATACTACTTAGAAACTTTCCAAGCGACAGAGATACAAGGTACAACAGAGACAGTGCAACAGGTTTGCTCTCATCTGCAGAGCCTGTCAAGTGCCTGCCCCTGAAAG 53486
WF-Teratorn ACAAAACAAAAAGAATACTACTTAGAAACTTTCCAAGCGACAGAGATACAAGGTACAACAGAGACAGTGCAACAGGTTTGCTCTCATCTGCAGAGCCTGTCAAGTGCCTGCCCCTGAAAG 58868

TeratornLC199500 GGTGTGGGTGGGGGAAGAGGTTTCTTAGGAAAGGTGTGCCTGTTAAATATTTAAGCGGAGGGGCTGGCTTCTCTCCTCCTATCGTCTTGCAAACAGCCAGGAATTCAGAGACATGCAGTG 53606
WF-Teratorn GGTGTGGGTGGGGGAAGAGGTTTCTTAGGAAAGGTGTGCCTGTTAAATATTTAAGCGGAGGGGCTGGCTTCTCTCCTCCTATCGTCTTGCAAACAGCCAGGAATTCAGAGACATGCAGTG 58988

TeratornLC199500 TTAGGAATAATTAGAAACAGAATAGTTAGAAACTCAATCTTTTCTTTTATTACATTCACTTTTACCTTTTACTACACAAATGGTTTAAACTTGATTCAGTATTATCATCTCAAATTTATA 53726
WF-Teratorn TTAGGAATAATTAGAAACAGAATAGTTAGAAACTCAATCTTTACTTTTATTACATTCACTTTTACCTTTTACTACACAAATGGTTTAAACTTGATTCAGTATTATCATCTCAAAATTATA 59108

TeratornLC199500 TTACATCCTAAGACAGCACAGCAGAGTTATTGTTGTCACAGTTTCTCCTCATCCTCCCTTTTTCACTTTTTGTGTGTCCT---------------------------------------- 53806
WF-Teratorn TTACATCCTAAGACAGCACAGCAGAGATATTGTTGTCACAGTTTCTCCTCATCCTCCCTTTTTCACTTTTTGTGTGTCCTGGTGTTTGTGCTCTGGCAAATGTCTATGAGAACATCAGAG 59228

TeratornLC199500 ----------------------------------------------------CGTGTGGGT------------CCATAGACCCTCGGACCATCTTCTGAATAAGTAAATGCTAGGTATATAA 53864
WF-Teratorn AGATCTGGCACTTGAAGCTTTATGACCTTCGAGTGTCTAGCCTGCAACAGCATGTGGGTGGGTTTTTCACACCATAGACCCTCGGACCATCTTCTGAATAAGTAAATGCTAGGTATATAA 59348

TeratornLC199500 ACCGAACCTTTTTATGAGTGTCTTTGTTCAGACTTTGAGATCATGTTCCTCTGTCTGTAACCCTGCGTACAAATCTGTGCTGTTTTGTGCGGGGTAAATCTTCACTGTAGCTATGTATTG 53984
WF-Teratorn ACCGAACCTTTTTATGAGTGTCTTTGTTCAGACTTTGAGATCATGTTCCTCTGTCTGTAACCCTGCGTACAAATCTGTGCTGTTTTGTGCGGGGTAAATCTTCACTGTAGCTATGTATTG 59468

TeratornLC199500 TCTCTGAGTCTTATTCATTCATTTCTGTGTGCAGGAAACTGAAGGGAACGAACCAACCGAACGTTTTTATTAAAACAGTCCTTTCTAGAAAATCCAATTTTCTCACAGGCAGC**ATGGACG** 54104
WF-Teratorn TCTCTGAGTCTTATTCATTCATTTCTGTGTGCAGGAAACTGAAGGGAACGAACCAACCGAACGTTTTTATTAAAACAGTCCTTTCTAGAAAATCCAATTTTCTCACAGGCAGCATGGACG 59588

TeratornLC199500 **TCGACTCGGATGAGGAAGTCAACGCAGGACGGCGGTTTGTCCCAAGACGCCGGACCATCGGGACCAATACTGGGACCTCGCCCGAAGGGCCGCAGGCCAGCGTGGAGGATTCCGAGTCAG** 54224
WF-Teratorn TCGACTCGGATGAGGAAGTCAACGCAGGACGGCGGTTTGTCCCAAGACGCCGGACCATCGGGACCAATACTGGGACCTCGCCCGAAGGGCCGCAGGCCAGCGTGGAGGATTCCGAGTCAG 59708

TeratornLC199500 **ATTCTGACAACGATCGGGCCATGATGGCTTACTTGAAAAAACGGATCAAAGAGTCCCGTCAGTCCGGGAGCGACAGCAGCGAGCTGGAGTCTGGCGGAGAAAGCGTGGAGTCCTACGACG** 54344
WF-Teratorn ATTCTGACAACGATCGGGCCATGATGGCTTACTTGAAAAAACGGATCAAAGAGTCTCGTCAGTCCGGGAGCGACAGCAGCGAGCTGGAGTCTGGCGGAGAAAGCGTGGAGTCCTACGACG 59828

TeratornLC199500 **ACAAAGAAATTTTCCCGGCCCACCTTTGGGGTTTCGGGGACCATGAAATCGACTGCGAATGCTGCAACTGCGTCTTCTGGCGCGCAAAGTACCGTCGCAATTACTACCAGAGTAAGCACG** 54464
WF-Teratorn ACAAAGAAATTTTCCCGGCCCACCTTTGGGGTTTCGGGAACCATGAAATCGACTGCGAATGCTGCAACTGCGTCTTCTGGCGCGCAAAGTACCGTCGCAATTACTACCAGAGTAAGCACG 59948

TeratornLC199500 **GTAGGATCTATTACGATCGCACCACGCAGACTGGAGCAACGGAATCCACGAGCGCGCCAGCTCAACCCGAGAAATCTGAGCCGGCACGCGTGGATCGCGGCACGAGGCCCATAACCTCAA** 54584
WF-Teratorn GCAGGATCTATTACGATCGCACCACGCAGACGGGAGCAACGGAATCCACGAGCGCGCCAGCTCAACCCGAGAAATCTGAGCCGGCATGCGTGGATCGCGGCACGAGGCCCATAACCTCAA 60068

TeratornLC199500 **GATTCACCAGCAAGAAGATGCGACTCGCCCAAGCCAAGTCACATTTCAGGCACACGGGAGACATATACGTCGACAGCACGTCTGACGAGGATGAGCCCAGAAAACCCTAG**TTTACAGTC 54704
WF-Teratorn GATTCACCAGCAAGAAGATGCGACTCGCCCAAGCCAAGTCACATTTCAGGCGCACGGGAGACATATACGTCGACAGCACGTCTGACGAGGATGAGCCCAGAAAACCCTAGTTTACAGTC 60188

TeratornLC199500 ACGGGCTGTGCACTCGTTTTACCAAACGATGTGAACCTGGTTTTGTTTGAATTGAATTCTACGTTCCTGTGTTCCATTACAAAATAAAGACTCCTTGACTGGGGTGTTGCTCCTTGACTG 54824
WF-Teratorn ACGGGCTGTGCACTCGTTTTACCAAACGATGTGAACCTGGTTTTGTTTGAATTGAATTCTACGTTCCTGTGTTCCATTACAAAATAAAGACTCCTTGACTGGGGTGTTGCTCCTTGACTG 60308

TeratornLC199500 ATTTTTTGTTTGTTTTGTTTTTTCTGTCTCGACCCTATCTTTAACTCGATTGAATACTGAGGAGCAAGTTACACAC----ACACATGCACAAAAGACAAGCACACAAAAACACACACAAC 54940
WF-Teratorn ATTTTTT----------TTTTTTCTGTCTCGACCCTATCTTTAACTCGATTGAATACAGAGGAGCAAGTTGCACACAGACACACATGCACAAAAGACAAGCACACAAAAACACACGCAAC 60418

TeratornLC199500 ATAAAGGTGTGCACTGTTGGACAAGCATGAAGGCTTTTGGTCAGAGAGGTAAAGTATGTCCCTAGACCTTGAGAAAGCTCATGTGCCCCTTCCACCGTATTTTCTGTGTGCAGAAGAGAT 55060
WF-Teratorn ATAAAGGTGTGCACTGTTGGACAAGCATGGAGGCTTTTAGTCAGAGAGGTAAAGTATGTCCCTAGACCTTGAGAAAGCTCATGTGCCCCTTCCACCGTATTTTCTGTGTGCAGAAGAGAT 60538

TeratornLC199500 CTGCTCAAACAGACGTTGCATCCTACCTGCCATAACGGCCAGGCAGCGCGGAAAGTGTAAGAAGTAGAGACGAAC--------------------------------------------- 55135
WF-Teratorn CTGCTCAAACAGACGTTGCATCCTACCTGCCATAACGGCCAGGCAGCGCGGAAAGTGTAAGAAGTAGAGACGAACCGAGGAGAGTGCACAGAAATGTTATTTTATTTCATTTTTTTCCAT 60658

TeratornLC199500 ----------------------------------------------------GAGGAGAGTGCACAGAAATTTTATTTTATTTCATTTTTT--CCATACATAACAGACAGCAGAGGGCAAT 55202
WF-Teratorn ACATAACAGACAGCAGAGGGCAATAAATCTACAAATCCGTAGAGACGAACCGAGGAGAGTGCACAGAAATGTTATTTTATTTCATTTTTTTTCCATACATAACAGACAGCAGAGGGCAAT 60778

TeratornLC199500 AAATCTACAAATCCGGTCGCCACT**TCAACATTCCCGCCCGCTGTCGACTGTCGAGTCACGATCGCTGTCTTCGCGGTAGCCCCTGCCGCCTTGTTTCACCTCCCTTCCCTGACCGTGCCT** 55322
WF-Teratorn AAATCTACAAATCCGGTCGCCACTTCAACATTCCTGCCCGCTGTTGACTGTCGAGTCACGATCGCTGTCTTCGCGGTAGCCCCTGCCGCCTTGTTTCACCTCCCTTCCCTGACCGTGCCT 60898

TeratornLC199500 **TGTCGAGGAGAACTCCTTCGCACTCGAGCGACATCCATTGCGAACAGGCACTCTCCCTCGCTCTGTTCTGTCAAACCTCGATTCGTCCGCTTCACGCCATCCGCTTTCCACAGCTTCGCT** 55442
WF-Teratorn TGTCGAGGAGAACTCCTTCGCACTCGAGCGACATCCGTTGCGAACAGGCACTCTCCCTCGCTCTGTTCTGTCAAACCTCGATTCGTCCGCTTCACGCCATCCGCTTTCCACAGCTTCGCT 61018

TeratornLC199500 **CGGGTCAACCCTTTCGGCGTGACGACCGGGCCCTCTCGTCTGCGCCCGGAGCTTGCCCGCTCGAGAAAGCTGCGCAAAGTCAGATTCCTCCGGAGGAGCTGACGGGGCGGGCGACTGGAA** 55562
WF-Teratorn CGGGTCAACCCTTTCGGCGTGACGACCGGGCCCTCTCGTCTGCGCCCGGAGCTTGCCCGCTCGAGAAGGCTGCGCAAAGTCAGATTCCTCCGGAGGAGCTGACGGGGCGGGCGACTGGAA 61138

TeratornLC199500 **CGGAGGCAACCAAGCGTGGGGGATGTCTCTCAGGGGTTCGACCGCTCCGGCGCCAGCACCCCACGCTCGAAACCTGTTGTAAATGAAGCCCCCTCGATCGGACGATGCGAAGCTCACGCT** 55682
WF-Teratorn CGGAGGCAACCAAGCGTGGGGGATGTCTCTCAGGGGTTCGACCGCTCCGGCGCCAGCACCCCACGCTCGAAACCTGTTGTAAATGAAGCCCCCTCGTTCGGACGATGCGAAGCTCACGCT 61258

TeratornLC199500 **GTAGCGAAACAGCAGGTACACGTAGTACCTGGACAACTTGCGATGGTCCACGGCGATGCATTTCGGGACCTTGGCGCCGGCGTGCCCGGCGCTCTCGCACTCCATCCGATGGAGCGAGAT** 55802
WF-Teratorn GTAGCGAAACAGCAGGTACACGTAGTACCTGGACAACTTGCGATGGTCCACAGCGATGCATTTCGGGACCTTAGCGCCGGCGTGCCCGGCGCTCTCGCACTCCATCCGATGGAGCGAGAT 61378

TeratornLC199500 **CAGGTCGTCCTGACACACGTGCAGCGCGTCAGACCTCGTCCACATGTCCCTCAGAAGCTCGAGGCTCTTTTGAAGCGTGGGTCTGTGCTTGCTGACCACTTCGTTCTGCAGCAGGCCAGT** 55922
WF-Teratorn CAGGTCGTCCTGACACACGTGCAGCGCGTCAGACCTCGTCCACATGTCCCTCAGAAGCTCGAGGCTCTTTTGAAGCGTGGGTCTGTGCTTGCTGACCACTTCGTTCTGCAGCAGGCCAGT 61498

TeratornLC199500 **GGCGGGCGCCCCTGCCACTATCAGGCACTCGAAAAGGGCTACCGAGGCGTGGGGAGAAAACGCGATTTTCTGTCTCATTTCTGAGTTCATTTTATAAACTGACCGAGAGCCCCCTTCGGG** 56042
WF-Teratorn GGCGGGCGCCCCTGCCACTATCAGGCACTCGAAAAGGGCTACCGAGGCGTGGGGAGAAAACGCGATTTTCTGTCTCATTTCTGAGTTCATTTTATAAACTGACCGAGAGCCCCCTTCGGG 61618

TeratornLC199500 **CGCGCAAAGCATGGAGGACGGCGGACGGTGCACTCGGAGGCTTCTCGGACTGCTGGGAGAGCACCGCTTGTATCATGTTAAGACTCTTTGTTGCAGCGACGCTCCTGGAGCGCTGGAGTG** 56162
WF-Teratorn CGCGCAAAGCATGGAGGACAGCGGACGGTGCACTCGGAGGCTTCTCGGACTGCTGGGAGAGCACCGCTTGTATCATGTTAAGACTCTTTGTTGCAGCGACGCTCCTGGAGCGCTGGAGTG 61738

TeratornLC199500 **GCTGGGAGCTAAAAGCGTCGATCTGTGCGAACCTCGCCGGGGAAGAGAAAACCTAGTCGCCTGGCTCACGAGACTGTGCGGGCTGTGCGGCGTGGACACGGACAAGGTAGAGATATGCAG** 56282
WF-Teratorn GCTGGGAGCTAAAAGCGTCGATCTGTGCGAACCTCGCCGGGGAGGAGAAAACCTCGTCGCCTGGCTCACGAGACTGTGCGGGCTGTGCGGCGTGGACACGGACAAGGTAGAGATATGCAG 61858

TeratornLC199500 **CGTGGACGCGAAACCTCGCGTCCGAGAGACCACGGAGCGTTTGGTGGTGTGCGTAGAGAGGACCTGGTTCCTGGGTCGCGACGGGTCTCTGGTCCTGTTTTCGACTGTAGGGCGAAACGC** 56402
WF-Teratorn CGTGGACGCGAAACCTCGCGTCCGAGAGACCACGGAGCGTTTGGTGGTGTGCGTAGAGAGGACCTGGTTCCTGGGTCGCGACGGGTCTCTGGTCCTGTTTTCGACTGTAGGGCGAAACGC 61978

TeratornLC199500 **CCCATCTCAGAAAAACGACGAGGTGAGGTCCTTGTGCGCGGCCGCAAACGCGATCTTGGGCGCGGGAGGCTTCGAGTATGTCGCTCGCCGCGTTGCCCGGGGCTGCGAGTGCTCAATAAG** 56522
WF-Teratorn CCCATCTCAGAAAAACGACGGGGTGAGGTCCTTGTGCGCGGCCGCAAACGCGATCTTGGGCGCGGGAGGCTTCGAGTATGTCGCTCGCCGCGTTGCCCGGGGCTGCGAGTGCTCAATAAG 62098

TeratornLC199500 **ACCCAAGGAGCAGGGGAGCGCTCCCGTGCTGAGGAGCCTGAGACCAGAGAGGTACGACACCGTGCCTCCAACGACGCCGATTGTGACAGCTGACAGTTTCGGACGGCGTAGCGAGCCAGA** 56642
WF-Teratorn ACCCAAGGAGCAGGGGAGCGCTCCCGTGCTGAGGAGCCTGAGACCAGAGAGGTACGACACCGTGCCTCCAACGACGCCGATTGTGACAGCTGATTGTTTCGGACGGCGTAGCGAGCCAGA 62218

TeratornLC199500 **GGCGCCCACCCGCTCGTCCGAAAAATGGACGTTACGCGCCGTCCCGGAGACGGCGTCCGCGACGGCGGTCGACCTCGAGTCCCTGCTGCGCGACGGGAAGTCCGTGCTGGGCCGCGGCGG** 56762
WF-Teratorn GGCGCCCACCCGCTCGTCCGAAAAATGGACGTTACGCGCCGTCCCGGAGACGGCGTCCGCGACGGCGGTCGACCTCGAGTCCCTGCTTCGCGACGGGAAGTCCGTGCTGGGCCGCGGCGG 62338

TeratornLC199500 **GTTCGGCGTCACGGTGCGCATCGAGGAGGGGCTGGTTGCCAAGGTCAACCTCTTCCCCGAGATGCTGAACTGGAGCCTGCCCGTGGTGGAGGACGAGTTCAAGCGCTATGCGCACGTCGC** 56882
WF-Teratorn GTTCGGCGTCACGGTGCGCATCGAGGAGGGGCTGGTTGCCAAGGTCAACCTCTTCCCCGAGATGCTGAACTGGAGCCTGCCCGTGGTGGAGGACGAGTTCAAGCGCTATGCGCACGTCGC 62458

TeratornLC199500 **GACCCAGATGGAGGAGGTCCTGATCGGAGCCTCCATCAAGCACCCGAACGTGCTGCGGACATTCGGGGGCTTGTGGTGCGAGGCCCCCGGGTACCCTTTGGGTGGCAGGGTCGTATCCCT** 57002
WF-Teratorn GACCCAGATGGAGGAGGTCCTGATCGGAGCCTCCATCAAGCACCCGAACGTGCTGCGGACATTCGGGGGCTTGTGGTGCGAGGCCCCCGGGTACCCTTTGGGTGGCAGGGTCGTATCCCT 62578

TeratornLC199500 **GATGGAGGAGGCGGCGTGCTCCCTGCACGAGTTTTCGGCGCACATGGGGACCGAGACGAGGTACGTGCCCCTGGTCGAACTGGACGTTCTGCGCGCCCTGGACCACCTGCGGTCGAGAGC** 57122
WF-Teratorn GATGGAGGAGGCGGCGTGCTCCCTGCACGAGTTTTCGGCGCACATGGGGACCGAGACGAGGTACGTGCCCCTGGTCGAACTGGACGTTCTGCGCGCCCTGGACCACCTGCGGTCGAGAGC 62698

TeratornLC199500 **CATCCAGCACAGGGACATCACGCACAGAAACGTCCTGGTGTGTCGCGCCGAAAGCAGGAGGCCCCTGCCACTGGTGTTCAAGATCAGCGACTTTGGCACCGCGTGCAACTTCTCCACTCC** 57242
WF-Teratorn CATCCAGCACAGGGACATCACGCACAGAAACGTCCTGGTGTGTCGCGCCGAAAGCAGGAGGCCCCTGCCACTGGTGTTCAAGATCAGCGACTTTGGCACCGCGTGCAACTTCTCCACTCC 62818

TeratornLC199500 **CGATCAGCCCAGGGGCACCCGTGCGAACAGAGCTCCCGAGGTGCTGTGGTGCCTGGAGTCCGACACTCAGGCGGACGTGTTCAGCTGGTACTGCATGATCTGGGAGCTGTACAGCGGTGC** 57362
WF-Teratorn CGATCAGCCCAGGGGCACCCGTGCGAACAGAGCTCCCGAGGTGCTGTGGTGCCTGGAGTCCGACACTCAGGCGGACGTGTTCAGCTGGTACTGCATGATCTGGGAGCTGTACAGCGGCGC 62938

TeratornLC199500 **GCCCCTGATCGGCCACAGGGCGTCCGGTCAGCGGGAAGGTTTTTGCAGCAGGACCTACGCGCGGAATTTGTCCGAGCTGCTAGGCACGTACGAGCCTACCTCAGAAGCCGCCTTCGATTC** 57482
WF-Teratorn GCCCCTGATCGGCCACAGGGCGTCCGGTCAGCGGGAAGGTTTTTGCAGCAGGACCTACGCGCGGAATTTGTCCGAGCTGCTAGGCACGTACGAGCCTACCTCGGAAGCCGCCTTCGATTC 63058

TeratornLC199500 **GTATTACATGAAGAGCTTGGACGCCATGGAGCTGCGGCGGCAATTCCGCTCGAGGCCGCGGACTAGAGACGTCGCGGAGAAGCTGGAAGCCGCGGCCGCGAGGGCGTCCGGTCGAGCGGG** 57602
WF-Teratorn GTATTACATGAAGAGCTTGGACGCCATGGAGCTGCGGCGGCAATTCCGCTCGAGGCCGCGGACTAGAGACGTCGCGAAGAAGCTGGAAGCCGCGGCCGCGAGGGCGTCCGGTCGAGCGGG 63178

TeratornLC199500 **CCCAGATCCGGAGTTCGTCGCCATCGGGGCGGCGTGCGTCACGTTGTTCCCCCAGGAAAGGTGGTCGGCGTCTGAGCTGCTCGAACTCCCGAGGTATCGGGACCTGGGCGCGGATGCCGT** 57722
WF-Teratorn CCCAGATCCGGAGTTCGTCGCCATCGGGGCGGCGTGCGTCACGTTGTTCCCCCAGGAAAGGTGGTCGGCGTCTGAGCTGCTCGAACTCCCGAGGTATCGGGACCTGGGCGCGGATGCCGT 63298

TeratornLC199500 **CTCGGTTCCGGCGAGGGAGATCCCGACGAACATCGGCGCCGGCTGCTACAGAGCCTGTGACGCGGTGGCTTGCGAGGACTGCGAACCCCCCGCCTTCCGGGTGCCAGACACCTCTTCTCT** 57842
WF-Teratorn CTCGGTTCCGGCGAGGGAGATCCCGACGAACATCGGCGCCGGCTGCTACAGAGCCTGTGACGCGGTGGCTTGCGAGGACTGCGAACCCCCCGCCTTCCGGGTGCCAGACACCTCTTCTCT 63418

TeratornLC199500 **CGCTACCACATTCCGATCCGAGAGTAAGATCTTCTACGGGGTCGACGTGATAAGACTGGCTCCGGGTTACATACAGCCTTACGTGTGGTACTCGAGAAAGGCCAGCGCTTCATCCCAGGT** 57962
WF-Teratorn CGCTACCACATTCCGATCCGAGAGTAAGATCTTCTACGGGGTCGACGTGATAAGACTGGCTCCGGGTTACATACAGCCTTACGTGTGGTACTCGAGAAAGGCCAGCGCTTCATCCCAGGT 63538

TeratornLC199500 **CGCCAAGGAAGACTCGGGCCGTGTCGGGCGGTCTCCGGAGAGACTCACCTTCGCGCAGATGCGGATCGGGTCTCCAGAGCCAGACCCGTGCTCCGCTTCGGCCAGCTCGCCCGCGGGAGG** 58082
WF-Teratorn CGCCAAGGAAGACTCGGGCCGTGTCGGGCGGTCTCCGGAGAGACTCACCTTCGCGCGGATGCGGATCGGGTCTCCAGAGCCAGACCCGTGCTCCGCTTCGGCCAGCTCGCCCGCGGGAGG 63658

TeratornLC199500 **TCCCCGGGTCATCGTGAAAGGCAAAAGCGGATGCTGCGAGGGAGAGGTCGACACAAAGATGGCCATCCTCACCCAGGCATCTCCGGAGGACGCCGAGAGGTTCAGAGAGCACGTCGTCGA** 58202
WF-Teratorn TCCCCGGGTCATCGTGAAAGGCAAAAGCGGATGCTGCGAGGGAGAGGTCGACACAAAGATGGCCATCCTCACCCAGGCATCTCCGGAGGACGCCGAGAGGTTCAGAGAGCACGTCGTCGA 63778

TeratornLC199500 **GCTCTCGATTGCTGTCACGGAGCAGCGACCGGACTTGTTCGCGGGCCCGATCCGAGGCCTGTGCAGGAGCTCGGAGAGCTGCGACGGCGTTGTGGTGTTCCAGTACGCTCCCATGTTCGC** 58322
WF-Teratorn GCTCTCGATTGCTGTCACGGAGCAGCGACCGGACTTGTTCGCGGGCCCGATCCGAGGCCTGTGCAGGAGCTCGGAGAGCTGCGACGGCGTTGTGGTGTTCCAGTACGCTCCCATGTTCGC 63898

TeratornLC199500 **TGACTGCAAGAAAGCATCCGAGTGGACTCTCGAGAACGCGCCTTTCTCCGCTGACGCCTTCGCCGCTCAGGT-------ACACTTCACCAGGCCTTGGAGGCAAAGAAGCTCCCCACGCG** 58435
WF-Teratorn TGACTGCAAGAAAGCATCCGAGTGGACTCTCGAGAACGCGCCTTTCTCCGCTGACGCCTTCGCCGCTCAGGTGCTCCTGACACTTCACCAGGCCTTGGAGGCAGAGAAGCTCCCCACGCG 64018

TeratornLC199500 **CGCGTTAAATCTGGACGACTTGCTGATCTGCCGAGGCTGCGCGCTGATCGACCCGGTGGCTTACCTCGCCAAGCACTTTGGTCGCGGTGGTCCTTTTTTCGGGGTGCGAAACGAGGGCCT** 58555
WF-Teratorn CGCGTTAAATCTGGACGACTTGCTGATCTGCCGAGGCTGCGCGCTGATCGACCCGGTGGCTTACCTCGCCAAGCACTTTGGTCGCGGTGGTCCTTTTTTCGGGGTGGGAAACGAGGGCCT 64138

TeratornLC199500 **GCTCACGTTCTGTATCGAGGTTTTGGGCAGATGCGGCTTTCGCGAGGCGGATTCCATAAGCCTCCTCTCCTCAAGGGACCCCGTCGGGATCCTCAGCGGGGCCCTGGGGATCCTGAGTGA** 58675
WF-Teratorn GCTCACGTTCTGTATCGAGGTTTTGGGCAGATGCGGCTTTCGCGAGGCGGATTCCATAAGCCTCCTCTCCTCAAGGGACCCCGTCGGGATCCTCGGCGGGGCCCTGGGGATCCTGAGTGA 64258

TeratornLC199500 **GAGAGCGGGCCGGGGGAGAGCCGTCGTACCCAGCAGGCTCACCCTGGACAGCGCCTACTTCGCGTTCGAGGAGTACACGGCCGAGAGGTCCACTTGGCGGGCCGAGCCGAGCCTCACGGG** 58795
WF-Teratorn GAGAGCGGGCCGGGGGAGAGCCGTCGTACCCAGCAGGCTCACCCTGGACAGCGCCTACTTCGCGTTCGAGGAGTACACGGCCGAGAGGTCCACTTGGCGGGCCGAGCCGAGCCTCACGGG 64378

TeratornLC199500 **GACGAAATACGAGTCGGCCCGCGGGCTCGGCCTGCTGGTGTTCGGCAACCCTCCGCCGCGAAGGGGAAAACAGACGTTGCCGGGGCCTCCATCTGATTCCAAGTCGCTTTCTTCCCTGAT** 58915
WF-Teratorn GACGAAATACGAGTCGGGCCGCGGGCTCGGCCTGCTGGTGTTCGGCAACCCTCCGCCGCGAAGGGGAAAACAGACGTTGCCGGGGCCTCCATCTGATTCCAAGTCGCTTTCTTCCCTGAT 64498

TeratornLC199500 **ACGAAACCTGACGCTGAGACTGAAGGTCAGAGTGTTCGAGAGCCACAAGGTCGAAGTGGTCGCGTTCAACTGCACGAGAGCCCCGCTGACCGTGGACCTCAGCAGGGCCGGGCTGCGCGA** 59035
WF-Teratorn ACGAAACCTGACGCTGAGACTGAAGGTCAGAGTGTTCGAGAGCCACAAGGTCGAAGTGGTCGCGTTCAACTGCACGAGAGCCCCGCTGACCGTGGACCTCAGCAGGGCCGGGCTGCGCGA 64618

TeratornLC199500 **CGTCGCGGGGGTCGACCTACTGAGCCCCGACAGAGTGCAGCTGTCCCGAGACGGCAGGTGCTACACTCACGACGCAGACCCTCGGGTCGCCTTGGGTCTATTGAAATGGGCCCCCGTCGT** 59155
WF-Teratorn CGTCGCGGGGGTCGACCTACTGAGCCCCGACAGAGTGCAGCTGTCCCGAGACGGCAGGTGCTACACTCACGACGCAGACCCTCGGGTCGCCTTGGGTCTATTGAAATGGGCCCCCGTCGT 64738

TeratornLC199500 **AATGATCTCCAAACACGCAGCACGGAGCCCCGTGCCCGAACTCACCAAAGTGTATTACAGGATCGTCATCGCGCTCGTACGCCTGTGCGGTAACGGCGCCTCCGAGAGCTCGCTGGTCGA** 59275
WF-Teratorn AATGATCTCCAAACACGCAGCACGGAGCCCCGTGCCCGAACTCACCAAAGTGTATTACAGGATCGTCATCGCCCTCGTACGCCTGTGCGGTAACGGCGCCTCCGAGAGCTCGCTGGTCGA 64858

TeratornLC199500 **CCTCTTTGCCAGACTTTCGCCCCTCATCGTCTCCACTCACCTGCGGTGCTAG**TGAGGAGCCCTTCGATCAGAGGCCCAGCGTACATGCTCATGTATGTGTGTCGATGAATGTCCCAGAGA 59395
WF-Teratorn CCTCTTTGCCAGACTTTCGCCCCTCATCGTCTCCACTCACCTGCGGTGCTAGTGAGGAGCCCTTCGATCAGAGGCCCAGCGTACATGCTCATGTATGTGTGTCGATGAATGTCCCAGAGA 64978

TeratornLC199500 CACGGCTACGCTACACTGAGATCTGCTTCGGCTGGCTTTGTTTCTGTACTTTTACTTGACTTTTTCACGTTTGTTTGATTGTTTTCAGGTGTCCTTCGAGGGACCTACACATTGTTTGTT 59515
WF-Teratorn CACGGCTACGCTACACTGAGATCTGCTTCGGCTGGCTTTGTTTCTGTACTTTTACTTGACTTTTTCACGTTTGTTTGATTGTTTTCAGGTGTCCTTCGAGGGACCTACACATTGTTTGTT 65098

TeratornLC199500 TTTTTGTTGTATCGTCGAGTGAAAATAAAATGTATTGTGTTCAGGTGTCTGGTTCTCCTTTATTTTCAGAGGTGAGATAAACTGTGCACTCGCTCCACCGGGGGTACGGGGTACGGTAGT 59635
WF-Teratorn TTTTTGTTGTATCGTCGAGTGAAAATAAAATGTATTGTGTTCAGGTGTCTGGTTCTCCTTTATTTTCAGAGGTGAGATAAACTGTGCACTCGCTCCACCGGGGGTACGGCGTACGGTAGT 65218

TeratornLC199500 GTAACCGTCCGTTGGCTCCAGTGGTACTAGGGCCCGCCAGTTCTTATTAAAATATTGCGTTCCCTTTCGGACTGCTGACAGATAGTTTCGTGTATCCCGTCAGAAAAAAAAAAA-CCATG 59754
WF-Teratorn GTAACCGTCCGTTGGCTCCAGTGGTACTAGGGCCCGCCAGTTCTTATTAAAATCTCGCGTTCCCTTTCGGACTGCTGACAGATACTTTCGTGTATCCCGTCAGAAAAAAAAAAAAACCATG 65338

TeratornLC199500 AGAGGTCTCCCATTCGGACCGTGGCTACTTCTGACTCTCACATGGACCTTGAGCGCAGCGGGTGAGTCCTCTGTAGGCTTGCTGATACTGTTCCTAGCACGAGTACGATCGATGAGCCTG 59874
WF-Teratorn AGAGGTCTCCCATTCGGACCGTGGCTACTTCTGACTCTCACATGGACCTTGAGCGCAGCGGGTGAGTCCTCTGTAGGCTTGCTGATACTGTTCCTAGCACGAGTACGATCGATGAGCCTG 65458

TeratornLC199500 TGTTTTGAGAGGCAGGCCGTGACTATTCGATGAGTCTGTGTAGTCAGGTTACAAAGTCGCAGAAGGACTCGGGAGGGTTGCCGGCTAAATGTTTTTTTCTGCTTTTCTTTCTTGTGTCGC 59994
WF-Teratorn TGTTTTGAGAGGCAGGCCGTGACTATTCGATGAGTCTGTGTAGTCAGGTTACAAAGTCGCAGAAGGACTCGGGAGGGTTGCCGGCTAAAT-TTTTTTTATGCTTTCTTTTTTTGTGTCGC 65577

TeratornLC199500 TGTCACGTTGTTACAGACACCAACATCACCTGTCTCATCATGGAAACCTGCATCTTACCCTGCAG**TTTTCAGAGAGGAGACAATCCGATAGTGCACTGGATTAAGGATAACGAGGGCACC** 60114
WF-Teratorn TGTCACGTTGTTACAGACACCAACATCACCTGTCTCATCATGGAAACCTGCATCTTACCCTGCAGTTTTCAGAGAGGAGACGGTCCGATAGTGCACTGGATTAAGGATAACGAGGGCAAC 65697

TeratornLC199500 **CCCTTTCTTTCATATTACAACGCGAACCCTCAGTTCAGGAATGTCAAGGGCAGAGGTTTCTTGTTCTCGGATCAGGTCTCTAAAGGTAATGCCTCACTACGGTTGTCCTGGGTGCAGCTT** 60234
WF-Teratorn CCCTTTCTTTCATATTACAAAGCGAACCCTCAGTTCAGGAATGTCAAGGGCAGAGGTTCCTTGTTCTCGGATCAGGTCTCTAAAGGTAATGCCTCACTACGGTTGTCCTGGGTGCAGCTT 65817

TeratornLC199500 **CACGATCAGGGAAGATACACCTGCTACGTTACAACCCGTAAGAATGAGCGATACACTTATGTCAACGTCAACGTGCAAG**GTAAGAGCGGTGAGTGACTTAGGTGACCGTTTAGCGAAAAT 60354
WF-Teratorn CACGATCAGGGAAGATACACCTGCTACGTTACAACCCGTAAGAATGAGCGATACACTTATGTCAACGTCAACGTGCAAGGTAGGAGCGGTGAGTGACTTAGGTGACCGTTTAGCGAAAAT 65937

TeratornLC199500 GCGGCTCTAAATTCTAGTTTGTCTCGCAG**CGGCTCCGGTCCTCGACAGTTCGGAGGGGATCAGCCGGGAACGCGAACCGGTTCGGTCGACGGAGCCCACTACGGCCAGCAGGACTTTGAA** 60474
WF-Teratorn GCGGCTCTAAATTCTAGTTTGTCTCGCAGCGGCTCCGGTCCTCGACAGTTCGGAGGGGATCAGCCGGGAACGCGAACCGGTTCGGTCGACGGAGCCCACTACGGCCAGCAGGACTTTGAA 66057

TeratornLC199500 **CGCCGCGATCACCCCCCGGTGGTCCACGCAGCCGTTTCCCAACAAGGGCGGCTACCACGGTCCAACAACAGAGAACGTCGACGCGGACCGTGACCGCACAGCGGCGACTCCCGAACGAGA** 60594
WF-Teratorn CGCCGCGACCACCCCCCGGTGGTCCACGCAGCCGTTTCACAACAAGGGAGGCTACCACGGTCCAACAACAGAGAACGTCGGCGCGGACCGTGACCGCACAGCGGCGACTCGCGAACGAGA 66177

TeratornLC199500 **CGATGGGGCGACGAAAGAGGCTGAAG**GTGAGAGGGGCCGACGGTCACTGTCGAGAGGGAACGCTTGGCCCGCCAAGCAAACGGGTTGCGCTGAGCCCAGGATTCGTTTCTCGCAG**CGGCT** 60714
WF-Teratorn CGATGGGGCGACGAAAGAGGCTGAAGGTGAGAGGGGCCGACGGTCACTGTCGAGAGGGAACGCTTGGCCCGCCAAGCAAACAGGTTGCGCTGAGCTCAGGATTCGTTTCTCGCAGCGGCT 66297

TeratornLC199500 **CCGGTCCTCGAAGTCCGCATGGAGGAGGTGGACGACACTGTCGTCTGCTGCTCGAAAGGGATCTACCCGAAACCCGAGCTAACCTGGTCCACCGAGCCTCCGTCCGGTGGGGCTCTGAAC** 60834
WF-Teratorn CAGGTCCTCGAAGTCCGCATGGAGGAGGTGGACGACACTGTCGTCTGCTGCTCGAAAGGGATCTACCCGAAACCCGAGCTAACCTGGTCCACCGAGCCTCCGTCCGGTCGGGCTCTGAAC 66417

TeratornLC199500 **GCCACGACCGCCGTCTGGCGGACCTCGCTGCTGCTCTACGACATCCGCAGCTCCTTGTTACTGTCGCACGAGGACGCTGTCATGGTTTACAGCTGCGTTGTAACCTCTGGCAAGCACAGC** 60954
WF-Teratorn GCCACGACCGCCGTCTGGCGGACCTCGCTGCTGCTCTACGACATCCGCAGCTCCCTGTTACTGTCGCACGAGGACGCTGTCATGGTTTACAGCTGCGTCGTAACCTCTGGCAAGCACAGC 66537

TeratornLC199500 **AAGAAAGCAACTTTGATGGTGGAAGGTGTGCGAAGCAAGTGGTTCATTTGTTTAGTTTCTTTTT**-----**TTTTGCACCTACTGTTGGACTTCAGATTATGCACGAACGCCAGAGGCGTCA** 61069
WF-Teratorn AAGAAAGCAACTTTGATGGTGGAAGGTGTGCGAAGCAAGTGGTTCATTTGTTTAGTTTCTTTTTTTTTTTTTTGCACCTACTGTTGGACTTCAGATTATGCACGAACGGCAGAGGCGTCA 66657

TeratornLC199500 **CTATCCTCAATCTGTTCTCGCTTTCACGGCTTATTTCTTTTCGTTTTCCTCAGGCAGACTCCGAAAAGTTCTGA**CGATCTATGTCCTGATCCCCGTCGGTGTCGCGGCTCTCGCAATTTG 61189
WF-Teratorn CTATCCTCAATCTGTTCTCGCTTTCACGGCTTATTTCTTTTCGTTTTCCTCAGGCAGACTCCAAAAGCTTCTGACGATCTATGTCCTGATCCCCGTCGGTGTCGCGGCTCTCGCAATTTG 66777

TeratornLC199500 CCTGGCTCTCGGGATATGGAGCATGCTAGGGAGAACCAGGAGATGCGCTCCCGCCTGACCGTGGGACTCTCGGCGACGGACCTTTCCAATCTGCCTGCTCATGCATTGTTCCAACTTCAC 61309
WF-Teratorn CCTGGCTCTCGGGATATGGAGCATGCTAGGGAGAACCAGGAGATGCGCTCCTGCCTGACCGTGGGACTCTCGACGACGGACCTTTCCAATCTGCCCGC--ATGCATTGTTCCAACCTCAC 66895

TeratornLC199500 AGGCTTACTTTTGCTACGTGAACCCTTGTGTGTGTTCGTTTGATTGCCGTAACAATAAAGACCCTAGAGTCCACCTCTGAAAGATGTCTGTATTTTGTTTTTCTTTTTGCTTTTTCAGTA 61429
WF-Teratorn AGGCTTACTTTTGCTACGTGAACCCTTGTGTGTGTTCGTTTGATTGCCGTAACAATAAAGACCCTAGAGTCCACCTCTGAAAGATGTCT------------------------------- 66984

TeratornLC199500 ATGAGAAAGTGTGGTTACGAAAAAATAAAAAGGATGGGGCATGCTCCATGAGCCTTCGAGCGCTTTGCGGCAACCCAGGGTGACCGTCCTACGCTTAACGGGGTTTCACCACGAGGTCGG 61549
WF-Teratorn ------------------------------------------------------------------------------------------------------------------------ 66984

TeratornLC199500 TAGGGTCCTCCGAGCCCATCGCTTCACACTGCGCAAAAAACAAAAGAAAGGAAACCCAGGTTAAACGTCTGACTAGATTTTACTTTGTGTATCATCTCACACGATTACGTGTGAAAGTAG 61669
WF-Teratorn ------------------------------------------------------------------------------------------------------------------------ 66984

TeratornLC199500 ATAGAAGACAGACAAAGATGGCAGAGAGGTTCAGCCATTGAATCGAGCGGCCTTCACTGTATGAGCATGCCCTTTCGATTCTCGCCTTACGAGTTCTCTGGAATATGTAGCTCGAGAAAC 61789
WF-Teratorn ------------------------------------------------------------------------------------------------------------------------ 66984

TeratornLC199500 TCGGATGCGCCCCGAACTCGCACTTTCCTGACCTGCCTTCAGGGAATTCCCCATCTAGCGTCGGAACGGATTTGGGCGGGTCGAACGAAGCCCTTATAACTCCGCCGTGAATGCAGTGTT 61909
WF-Teratorn ------------------------------------------------------------------------------------------------------------------------ 66984

TeratornLC199500 TGACTGAATCACCACCATCTGCTACGGTTTTACCAGGCTCCCCGAGCTCCCATTAAACTTCAGCGCCCCCTCGGACGAAAGCGAAGCGGACAGCGAGCGACGACGGTCGAAAGGTCCTTC 62029
WF-Teratorn ------------------------------------------------------------------------------------------------------------------------ 66984

TeratornLC199500 TGTGTCTCCCACCGCCGACTCCCTCGACCCCAACCGTACTCTATCTGGCCTCAGTCATCAGAGCGGATAAACC**ATGAGAGGGTTCCTATTGGGACCGTGGATCCTTCTGACTCTCGCGTG** 62149
WF-Teratorn ------------------------------------------------------------------------------------------------------------------------ 66984

TeratornLC199500 **GACACTGAGAGCTGAGG**GTGAGTGAACTCGTTGTGTTGTGGAAGGCAGATGGTGAAACTTTGATGAATCTTTGTGAAAATGTGTGGGGTTTTGTTTTTCTTGTTTTTTTTTCTTAAGCCG 62269
WF-Teratorn ------------------------------------------------------------------------------------------------------------------------ 66984

TeratornLC199500 CTGTCATGTAG**TTACAGACACCGACGTTACCTGCCTCGTCATGGAAACCTGCATCTTACCCTGCAGTGTTCCGAGGGAGGGCTCCATAACAGTGCGCTGGTTTAAGGATCCAGAGATAGT** 62389
WF-Teratorn ------------------------------------------------------------------------------------------------------------------------ 66984

TeratornLC199500 **TCCCTTCCTCACATATGGTATAACAAACCCTGACAGGGGGAATGTCAACAACAGAACCTCCTTGTTCGAGGATCAGATCTTTGAAGGAAATGCTTCAATATTGCTGTCAAAGGTGCAGCT** 62509
WF-Teratorn ------------------------------------------------------------------------------------------------------------------------ 66984

TeratornLC199500 **TCAAGACCAGGGATCATACCGTTGTGTCGTTGTCAGCTCGAGAAATGGGCATACCAGAAACGTCAAAGTAATGGTGCAAG**GTAAGAGTCGCAAGTGTTGTGATTAAGTCAACCCCTGTGA 62629
WF-Teratorn ------------------------------------------------------------------------------------------------------------------------ 66984

TeratornLC199500 CCGTTAAGCGAAGACGCGGCCCTAAGTCCGAGTTTTGTCTCGCGGCAG**CTCCTGTCACCGACGGTCACGCCAAAGAGGGGTTCGACCGGGATCCGAGTCTGGCCCGCCCGACGCCTCCTC** 62749
WF-Teratorn ------------------------------------------------------------------------------------------------------------------------ 66984

TeratornLC199500 **CTCCACCTGGCTGGTCGAACGGCGGCGGCGCAGCCGGGGAAGAGGGGTCCAGCCGGGATCCGAGTCTGGCCAGCCCGACGCCTCCTCCTCGGACACACACCATGACCACCCCCCAGCAGT** 62869
WF-Teratorn ------------------------------------------------------------------------------------------------------------------------ 66984

TeratornLC199500 **CCACGCAGACTGTCCACGCAATCAGCGACTCCCGGGTTCGACCGACCGAGGTTGTCGACGTGGACCCCAACGAAACCCAGGCCACTGGCAAAGGAGACGATAAAGCGTCGAGGGAGGTGG** 62989
WF-Teratorn ------------------------------------------------------------------------------------------------------------------------ 66984

TeratornLC199500 **AAG**GTGAGAGAGGCTTTCCATTTTTTATTTATTTCAATTTTTTTTTTTTTAAATTTGTTTGTTTCTTTCTTCTTTTAG**TTTCTGCAGCATCTCCGGTCACCAAAGTCCGTCTGGAGCAGGT** 63109
WF-Teratorn ------------------------------------------------------------------------------------------------------------------------ 66984

TeratornLC199500 **TGACAACACCGTTGTGTGCATTTCGGAGGGCATCTACCCGGAACCCGAGCTGACTTGGTCCACCCAGCCTCCGTCGGGCAGGACTCTGAACGCCACTACCGCCGTACGACGGACCTCACG** 63229
WF-Teratorn ------------------------------------------------------------------------------------------------------------------------ 66984

TeratornLC199500 **GATGCTCTATGACATCCGCAGCACCGTGGAACTGTCTCCTGAGGACGCTGGCTTGGTTTACAGCTGCGCCGTAGCCTCTGGCATAAACAGCAAAAAAGCGACAGTGAAAATTCAGTTTT**G 63349
WF-Teratorn ------------------------------------------------------------------------------------------------------------------------ 66984

TeratornLC199500 TGAGTGGTGCTCTTATTCATTTATTCATTTATTTATTTATTTATTTATTTATTTATTTATTTATCTTTAACATTTGCATTTAATATCAGTTACGTGAGTGGTGCTCTTATTTATTTATTT 63469
WF-Teratorn ------------------------------------------------------------------------------------------------------------------------ 66984

TeratornLC199500 TTAACAATTGAATTTATTATTTGCAAAAGCTGATGGACACCATCAACTCTCGGTTTAACGGAACATCTCCCTTGTTCTTCTCAG**GCCGACTTCGAAATCGTGTTTTGTTTGTCCTTATCC** 63589
WF-Teratorn ------------------------------------------------------------------------------------------------------------------------ 66984

TeratornLC199500 **CCATCGGTCTCGCTGCTCTCGCGATTTGCGCGATCGTCTGGAGGATGCGACGAAAGATGAAAAGGGGATTGGCTCCTGCATCTATCAACCTCAGTGACTTACTTCATAAGACATAA**ACCC 63709
WF-Teratorn ------------------------------------------------------------------------------------------------------------------------ 66984

TeratornLC199500 ATGTGTGTGTTCATTTGATCACCGTTACAATAAAGACCATCGAGTCCACCTCTGAAAGATGTCTCTGTTTTGTTTTCGTTATTATCCTTTTCAGTGCTGAAAAAGTGTGGTTAGGACAAA 63829
WF-Teratorn ----------------------------------------------------------------CTGTTTTGTTTTCGTTATTATCCTTTTCAGTGCTGAAAAAGTGTGGTTAGGACAAA 67040

TeratornLC199500 AACTATATAAAGGACAAGAAAATGCTATGAGGCACTTTGAACAATGTGGCTACGCGTAGCAGGGCCCCAATGATTCTGCTTGAGGTCACTCGGGTCCTCCGAGCGCAAACCCGACGCGCC 63949
WF-Teratorn AACTATATAAAGGACAAGAAAATGCTATGAGGCACTTTGCACAATGTGGCTACGCCTAGCAGGGCCCCAGAGATTCTGCTTGAGGTCACTCGGGTCCTCCGAGCGCGAACCTGACGCGCC 67160

TeratornLC199500 GATCGCTTCAGACGGTGCAGACAAGGAAAAGAGACACAGGGCAAACGTCTGACATGATTTTATTTTGTATCTCA**TCACACGTGATCACACTGAGATGCTGCATCGGGCACTACAGCAGTC** 64069
WF-Teratorn GATCGCTTCAGACGGTGCAGACAAGGAAAAGAGACACAGGGCAAACGTCTGACATGATTTTATTTTGTATCTCATCACACGTGATCACACTGAGATGCTGCATCGGGCATTACAGCCGTC 67280

TeratornLC199500 **GAGGCGGAGCACAGGTCAGAGTCTGTGCTCTCGTCAACCGAGGGAGTGTGGAAGCGTAAGCACAAGCTTTCGGTCTTCGTCCGCTTCGCCCCGTCGTCAGCACGCCCACGATGCGAGTAG** 64189
WF-Teratorn GAAGCGGAGCACAGGTCAGAGTCTGTGCTCTCGTCAACCGAGGGAGTGTGGAAGCGTAAGCACAAGCTTTCGGTCTTCGTCCGCTTCGCCCCGTCGTCAGCACGCCCACGATGCGAGTAG 67400

TeratornLC199500 **ACGCTCACAAGCCCGACGTCTCGGTTTCCACTCGATGAGGAGGCCTCCCCCGAGCGAGGCACTTCAACGTGGTCGGCCAGCTCTTCCAAATAGGTCATCATGCGAGAGCTCTGTGCGCCG** 64309
WF-Teratorn ACGCTCACAAGCCCGACGTCTCGGTTTCCACTCGATGAGGAGGCCTCCCCCGAGCGAGGCACTTCAACGTGGTCAGCCAGCTCTTCCAAATAGGTCATCATGCGAGAGCTCTGTGCGCCG 67520

TeratornLC199500 **TGCGAGCGGAGAGTCTTCGAGGGCCCTGGTCGCCGGGACTCCAT**AGGCCCTGCCTGAAGGGAGATAGAAGACAGAGGAGTGCAACGGAAAGGTTAAAGGGGTTGAGTCTTCTTCGTCTGC 64429
WF-Teratorn TGCGAGCGGAGAGTCTTCGAGGGCCCTGGTCGCCGGGACTCCATAGGCCCTGCCTGAAGGGAGATAGAAGACAGAGGAGTGCAACGGAAAGGTTAAAGGGGTTGAGTCTTCTTCGTCTGC 67640

TeratornLC199500 CTCCAACGGTGTGAGCTTGTCGCCCCTCTGCCATTTATAAAAGACACCGGGTAACCCGACTTTGTGACCTGCCTTCGGGGAATTCTGCGTGACGTGTCGGAACGGTCTTCGGGTGTGATC 64549
WF-Teratorn CTCCAACGGTGTGAGCTTGTCGCCCCTCTGCCATTTATAAAAGACACCGGGTAACCCGACTTTGCGACCTGCCTTCGGGGAATTCTGCGTGACGTGTCGGAACGGTCTTCGGGTGTGATC 67760

TeratornLC199500 GAGTGGTGCAAAGTGGAATAGTGTTTGCCTCAATCAACGAACGTCTGCTGCTGCTTCACCGGGCTCTGCGAGCGTCCCTTCGGACGAAAGTGAAGCAGACGGCGAGCGACGACGGTCGAA 64669
WF-Teratorn GAGTGGTGCAAAGTGGAATAGTGTTTGCCTCAATCAACGAACGTCTGCTGCTGCTTCACCGGGCTCTGCGAGCGTCCCTTCGGACGAAAGTGAAGCAGACGGCGAGCGACAACGGTCGAA 67880

TeratornLC199500 AGGTCCTTCTGCGTCTCCCGCAGCAGACTCCGCGACCCCAACCCTACTCGATCGGGCCTCGGCCATCTGAGCAGCTAAACC**ATGACAGGGCTCCCATTCTTACCGTGGCTACTTCTGACT** 64789
WF-Teratorn AGGTCCTTCTGCGTCTCCCGCAGCAGACTCCGCGACCCCAACCCTACTCGATCGGGCCTCGGCCATCTGAGCAGCTAAACCATGACAGGGCTCCCATTCTTACCGTGGCTACTTCTGACT 68000

TeratornLC199500 **GTCGCGCGGACCCAGAGCGTGGAGG**GTGAGTGCTCTGTAAGCTTGCTGACATTGTTCCGAGTACGAGCGCGATCGTCATAACCCAAACAGTCAGCGCAACGCGGCTAGAGCAAAGCGAAA 64909
WF-Teratorn GTCGCGCGGACCCAGAGCGTGGCGGGTGAGTGCTCTGTAAGCTTGCTGACATTGTTCCGAGTACGAGCGCGATCGTCATAACCCAAACAGTCAGCGCAACGCGGCTGGAGCAAAGCGAAA 68120

TeratornLC199500 ACAGGGGATGGCGAATTCCACACTACGGGTTGAGATGCAGTATCGTGACGTTGACTAAAAGTTCTTTTTTGTTTTTGTTCACCGCTCACACTAAAAG**TCAGCGAGGAGGTGGTGTACTAC** 65029
WF-Teratorn ACAGGGAATGGCGAATTCCACACTACGGGTTGAGATGCAGTATCGTGACGTTGACTAAAAGTTCTTTTTTTTTTTTGTTCACCGCTCACACAAAAAGTCAACGAGGAGGTGGTGTACTAC 68240

TeratornLC199500 **GACATCTATTACGACCCAGACTACGCCTTCAATTACAGTTATGAGGGGATCAATCCGGAACCCGATCTGACCCGGCCAACGGAGCGTCCTCCATCCAGGTGGACATTGGACGACACCCAG** 65149
WF-Teratorn GACATCTATTACGACCCAGACTACGCCTTCAATTACAGTTATGAGGGGATCAATCCGGAACCCGATCTGACCCGGCCAACGGAGCGTCCTCCATCCAGCTGGACATTGGACGACACCCAG 68360

TeratornLC199500 **GCCACTCACGAAAGAGACAATGATGCGACGAGGGATGCTGAAG**GTGAGAGGGGCCGGCGGTCCCTTTCGAGGGGGCCGCTTGGGCCTAGAAGAAAACAGGTTGAGCTGAGCTCTGTCTTC 65269
WF-Teratorn GCCACTCACGAAAGAGACAATGATGCGACGAGGGATGCTGAAGGTGAGAGGGGCCGGCGGTCCCTTTCGAGGGGGCCGCTTGGGCCTCGAAGAAAACAGGTTGAGCTGAGCTCTGTCTTC 68480

TeratornLC199500 GTTTCAAAGCAG**CTCCGGTCACCGAAGTCCACCTGGAGCAGGTGGGTGACACCGCCGTGTGCAGTTCGGTGGGGATCTACCCGGAACCCGAGCTGACCTGGTCCACCGAGCCTCCGTCGG** 65389
WF-Teratorn GTTTCAAAGCAGCTCCGGTCACCGAAGTCCACCTGGAGCAGGTGGGCGACACCGCCGTGTGCAGTTCGGTGGGGATCTACCCGGAACCCGAGCTGACCTGGTCCACCGAGCCTCCGTCGG 68600

TeratornLC199500 **GCAGGAATCTGAGCGCCACGACCGCCGTCTGGCGGACCTCGCAGCTGCTCTACGACATCCGCAGCTCCTTGTTACTGTCGCCCGAGGACGCAGGCTTGGTTTACGTCTGCACCGTAGTCT** 65509<br>
WF-Teratorn GCAGGACTCTGAGCGCCAC------TGTCTGGCGGACCTCGCAGCTGCTCTACGACATCCGCAGCTCCTTGTTACTGTCGCCCGAGGACGCAGGCTTGGTTTACGTCTGCACCGTAGTCT 68714

TeratornLC199500 **CTGGCGAAAGCAGCAAGAAAGCGACCGTGAAAGTTAAGCGAGGCGAGTG**GTATTTTTTTTT--AAAAAAAAACAAACAATATTTTAATTTGGTTTTTGGACCCATTTGTTGACTGTTTCT 65627<br>
WF-Teratorn CTGGCAAAGGCAGCAAGAAAGCGACCGTGAAAGTTAAGCGAGGCGAGTGGTGTTTTTTTTTTTAAAAAAAAAACAACAATATTTTAATTTGGTTTTTGGACCCATTTGTTGACTGTTTCT 68834

TeratornLC199500 ATCCCC-GCTCTCTTTCAG**TCTTCTCAGGAGGACTGTTTGTCATGATATCCATCGGTGTCACGGCTCTGACTGTTTGCACGGCTCTCTGGATCTGGAGAACGCGAAGTGGAAACAGGAGA** 65746<br>
WF-Teratorn ATCCCCCGCTCTCTTTCAGTCTTCTCAGGAGGACTGTTTGTCATGATATCCATCGGTGTCGCGGCTCTGACTGTTTGCACGGCTCTCTGGATCTGGAGAACGCGAGGTGGAAACAGGAGA 68954

TeratornLC199500 **CAAGCTCCCGCCTGGGCGTAGGCTTATTCACGATGTGTCTTTCTGCCAGTGCGCCATAG**GGTGATGCAAACCCATGACTGTGTTCATTTGATCATTGTGACAATAAAGACCCTCGTGTAG 65866<br>
WF-Teratorn CAAGCTCCAGCCTGGGCGTAGTCTTATTCGCGATGTGTCTTTCTGCCAGTGCGCCATAGGGCGATGCAAACCCATGACTGTGTTCATTTGATCATTGTGACAATAAAGACCCTCGTGTAG 69074

TeratornLC199500 ACTCCGAATGATGTCTGTATTTTGTTTGCTTCATTTTTTTTTAATTTTCATTTTCTGTGCTAAGAAGAGTTAGATCAGAAAAGGTTATGTTTAGAAAAA-TAAATGTAAAAGCGGAGAGG 65985<br>
WF-Teratorn ACTCTGAATGATGTCTGTATTTTGTTTGCTTCATTTTTTTTTAATTTTCATTTTCTGTGCTAAGAAGAGTTAGATCAGAAAAGGTTATGTTTAGAAAAAATAAATGTAAAAGCGGAGAGG 69194

TeratornLC199500 TATACTCTGGTATTCCTCGAGGCATTTTCCGCGTTGCTGCAACGCAGTGTGACGTTCTACGCTTATCGGAGCTGCAGAGTTTTACCGTGAGACCGCACAAGAAAAAAAAAAAAGAAAGAAC 66105<br>
WF-Teratorn TATACTCTGGTAGCCCTCCAGGCGCTTTCCGCGTTGCTGCAACGCAGTGTGACGTTCTACGCTTATCGGAGCTGCAGAGTTTTACCGTGAGACCGCACAAGAAAAAAAAAAAAGAAAGAAC 69314

TeratornLC199500 ACTCAAAGCAAAGCGTCTTACTTTGATTTTTATTTCATCACACATAATTACACAGATCCCACGGATGTAGAAGCGGTGCACCAGACAGTCTGCGGTCGCATCGAACTAATGTCCCAAACG 66225<br>
WF-Teratorn ACTCAAAGCAAAGCGTCTTACTTTGATTTTTATTTCATCACACATAATTACACAGATCCCACGGATGTAGAAGCGGTGCACCAGACAGTCTGCGGTCGCATCGAACTAATGTCCCAAACG 69434

TeratornLC199500 AGAGCCCCAGTTTTGTCCACCGTGACTCCAATGCGTTCTGTCCGAAGGTTGGCTGAAGACAGGAGTGCAACGGAGAGGTCGAAGAGGTCGAGTCGGGTTCGTCTTCTTCGTCCGCCCCCA 66345<br>
WF-Teratorn AGAGCCCTAGTTTTGTCCACCGTGACTCCAATGCGTTCTGTCCGAAGGTTGGCTGAAGACAGGAGTGCAACGGAGAGGTCGAAGAGGTCGAGTCGGGTTCGTCTTCTTCGTCCGCCCCCA 69554

TeratornLC199500 CCGCTCGAGCTAGCCACTCCTCGACCTTTT**CTACATCCACACCGTGTACGGGGCGCTTCCCTCTCCATCGCTTTCGTCTTGCGAGAGCCGCCGCTGAGACGCCTTGCGCTCGTCCTTGCC** 66465<br>
WF-Teratorn CCGCTCGAGCTAGCCACTCCTCGACCTTTTCTACATCCACACCGTGTACGCGGCGCTTCCCTCTCCATCGCTTTCGTCTTGCGAGAGCCGCCGCTGAGACGCCTTGCGCTCGTCCTTGCC 69674

TeratornLC199500 **ATCTGTGCAGAGGCGACCGGGTCCCTCTCCGAACCCTCCTGCGAGCAAATCGGCTCCAGGCTCTCTGTTCTCTGTGGGCTCTGGGTCCTCTGTGAGCTGCGGGTGCTCGGCGTGAGACCC** 66585<br>
WF-Teratorn ATCGGTGCAGAGGCGACCGGGTCCCTCTCCGAACCCTCCTGCGAGCAAATCTGCTCCAGGCTCTCTGTTCTCTGCGGGTTCTGGGTCCTCTGTGAGCTGCGGGTGCTCGGCGTGAGACCC 69794

TeratornLC199500 **CGCGCGCTCCGCGCTCCTCTTTCTCTGCGGGTCGCTCGCTTCCGAGCTGCCCGGCGAACAGCTCAGTTTAACCGGCGTGATCGGCTTTGGTTTTGCCGGCGACGTCACAATTCGACCGTC** 66705<br>
WF-Teratorn CGCGCGCTCCGCGCTCCTCTTTCTCTGCGGGTCGCTCGCTTCCGAGCTGCCCGGCGAACAGCTCAGTTTAACCGGCGTGATCGGCTTTGGTTTTGCCGGCGACGTCACAATTCGACCGTC 69914

TeratornLC199500 **GGTCTCTCGCCTTTCGCCGCGAGACCTGACGGTTTGGCACCGCCTGCCCAGGTCCGTCGACTTCAGGGAGCTCCTGGCCTCACGCTCCCTGCGTGAACGGAGGCCTGCGGTCGATTTGTC** 66825<br>
WF-Teratorn GGTCTCTCGCCTTTCGCCGCGAGACCTGACGGTTTGGCACCGCCTGCCCAGGTCCGTCGACTTCAGAGAGCTCCTGGGCTCGCGCTCCCTGCGCGAACTGAGGCCTGCGGTCGATTTGTC 70034

TeratornLC199500 **AGAAAGGTCGGCCCCTTCGGATTCCCAACCGCTGCCTTCAGACGCGGACCCTCGACTCTCCGAACTGGAGGAGCGCGAACGAGCGATCCGAGCGCCGGACCTTTTCGAACCTCGACCGTC** 66945<br>
WF-Teratorn CGAAAGGTCGGCCCCTTCGGATTCCCAACCGCTGCCTTCAGACGCGGACCCTCGACTCTCCGAACTGGAGGAGCGCGAACGAGCGATCCGAGCGCCGGACCTTTTCGAACCTCGACCGTC 70154

TeratornLC199500 **GGCCTCTCTGCCTTTCTCGCCAGAGAAACGCAGCGGTTTCCCGGAGGCTGATTTTTGCAGCCCTCTCTCTGGGCGTCTTACGTCTTTGCGGCCGCTGGAGTAAGTCTTGGAGTGGTCTTC** 67065<br>
WF-Teratorn GACCTCTCTGCCTTTCTCGCCAGAGAAACGCAGCGGTTTCCTGGAGGCTGATTTTTGCAGCCCTCCCTCTGGGCGTCTTACGTCTTTGCGGCCGCTGGAGTAAGTCTTGGAGTGGTCTTC 70274

TeratornLC199500 **GGTGTCGTTGCGTTGTCGCCGCGTGAGCGCGGTTTCGAGCCCCGGCCTCTTTTCCCGCTCGACCGAGGGTCTCGACTTCGACCCTCTGCCTTTCGGCCTCGCGGAGACTCGCATCGCCCA** 67185<br>
WF-Teratorn GGTGTCTTTGCGTTGTCGCTGCGTGAGCGCGGTTTCGAGCCCCGGCCTCTTTTCCCGCTCGACCGAGGGTCTCGACTTCGACCCTCTGCCTTTCGGCCTCGTGGAGACTCGCATCGCCCA 70394

TeratornLC199500 **TGGGCCTTCCGTCTCCGAGCTGTCCGAGCTGAGTGCGCGCTTGTAGCCCCGCCGGTCGCTTTTGCACGAGGGCGAGGACGGCTGCGACCCGCAATACGGACGGCCGACCCCGTCCTCGTT** 67305<br>
WF-Teratorn TGGGCCTTCCGTCTCCGAGCTGTCCGAGCTGAGTGCGCGCTTGTAGCCCCGCCGGTCGCTTTTGCACGAGGGCGAGGACGGCTGCGACCCGCAATACGGACGGCCGACCCCGTCCTCGTT 70514

TeratornLC199500 **CCACCTGGCCTTGGAGCCCGCTTCCTCGCCAGCTCTTTGTCTGTACGCGCTACTGGGCGAG------GAAGGTCGCGACACGCAGTACGGACGGTCAACTCCGTCCTCGCGTCTCTTGGC** 67419<br>
WF-Teratorn CCTCCTGGCCTTGGAGCCCGCTTCCTCGCCAGCTCTTTGTCTGTACGCGCTACTGGGCGAGGGCGAGGAAGGTCGCGACACGCAGTACGGACGGTCAACTCCGTCCTCGCGTCTCTTGGC 70634

TeratornLC199500 **CCTGGCGACGTTGTCCTCTCCCGCCTCGATGTCTTCTTCGCTGGACGACGATGCGTCGTCTATGAAACTGTCGCTGCGATCGTAGTCGTCTGAGCTTTTGTCGCAATCCGAAGAGCTCTC** 67539<br>
WF-Teratorn CCTGGCGACGTTGTCCTCTCCCGCCTCGATGTCTTCTTCGCTGGACGACGATGCGTCGTCTATGAAACTGTCGCTGCGATCGTAGTCGTCTGAGCTTTTGTCGCAATCCGAAGAGCTCTC 70754

TeratornLC199500 **CGTGTCCCTGTCGCTCAAATACTCCGGCTCTTCCTCGCTTTCGGAGCTGAGAGGTTCCAGGCCGTTGTCTTCCTCGGCTTCGCTCAGTAAAAAACTGTTTCTCTGGCGGCGCCCGACCTT** 67659<br>
WF-Teratorn CGTGTCCCTGTCGCTCAAATACTCCGGCTCTTCCTCGCTTTCGGAGCTGAGAAGTTCCAGGCCGTTGTCTTCCTCGGCTTCGCTCAGTAAAAAACTGTTTCTCTGGCGGCGCCCGACCTT 70874

TeratornLC199500 **TCTCCTCTTGGTCGCCGGAGACCGCGAGGCTCGGACGCTCTCGGCCCAGGGGGACAACCGACGTCGAGCTCTGCCGACGGTCTGGCGGCAACTCCTCTCGGGCTTCTTGACTGGCGGCGG** 67779
WF-Teratorn TCTCCTCTTGGTCGCCGGAGACCGCGAGGCTCGGACGCTCTCGGCCCAGGGGGACAACCGACGTCGAGCTCTGCCGACGGTCTGGCGGCAGCTCCTCTCGGGCTTCTTGACTGGCGGCGG 70994

TeratornLC199500 **CGGGGAGAGGTGCCTTCGGGGAACCCGCCTGGACGCCGCCTTGCGCTGGAGTGCCGCGAAGGACCGGGCCACCTTGTCGCTCCACCTCTGCGTGGATTGGGACTCCAAGCATTGGATCAA** 67899
WF-Teratorn CGGGGAGAGGTGCCTTCGGGGAACCCGCCTGGACGCCGCCTTGCGCTGGAGTGCCGCGAAGGACCGGGCCACCTTGTCGCTCCACCTCTGCGTGGATTGGGACTCCAAGCATTGGATCAA 71114

TeratornLC199500 **CGCTCGGCTCAGGCGATCCTTGATGCCGCGCCTGCGGAGCTCGTTGACCGTGCCCTGGTTGTACGCCACCCATTCTCTCAAGAGTTCCCTGTTCGCCTCGCCGAGGAGAGCGCTGTCCAG** 68019
WF-Teratorn CGCTCGGCTCAGGCGATCCTTGATGCCGCGCCTGCGGAGCTCGTTGACCGTGCCCTGGTTGTACGCCACCCATTCTCTCAAGAGTTCCCTGTTCGCCTCGCCGAGGAGAGCGCTGTCCAG 71234

TeratornLC199500 **GTTCAGGAGCTTGCACACGGCCTCTCTCTGGTACCCCAAGGAGTCGTGCAACCAAGGGTAGCTCAGGAGCAACACGGACAAGTTTTTCATCCATGTCGAAACCGGGACGGGCTCTGGGAT** 68139
WF-Teratorn GTTCAGGAGCTTGCACACGGCCTCTCTCTGGTACCCCAAGGAGTCGTGCAACCAAGGGTAGCTCAGGAGCAACACGGACAAGTTTTTCATCCATGTCGAAACCGGGACGGGCTCTGGGAT 71354

TeratornLC199500 **GGGGTTGAAAAGCTCCGAGCGGTCGTTCGGGGGCTGTCTACATCCCTCGCCTGGCTCGACCGGCTCTCCGCAAGCGGGATTTCTGCCGCCCATGTCACCGTCAAAGCGGGCTGCGTTGAT** 68259
WF-Teratorn GGGGTTGAAAAGCTCCGAGCGGTCGTTCGGGGGCTGTCTACATCCCTCGCCTGGCTCGACCGGCTCTCCGCAAGCGGGATTTCTGCCGCCCATGTCACCGTCAAAGCGGGCTGCGTTGAT 71474

TeratornLC199500 **CCCCAGTTTCTTGAGCTTGCTGCAGTAAACTTTGAGCTTCAGGAACGCCACTTTGGCGGCCGCGTTCACCTCCTTGGCGTGGGCTTGCGCTTCCTTGTCTTCCTTGTTGCACTGGTCGCG** 68379
WF-Teratorn CCCCAGTTTCTTGAGCTTGCTGCAGTAAACTTTGAGCTTCAGGAACGCCACTTTGGCGGCCGCGTTCACCTCCTTGGCGTGGGCTTGCGCTTCCTTGTCTTCCTTGTTGCACTGGTCGCG 71594

TeratornLC199500 **TTTGCACATGGTGTACCAGTCTTTCTGGATCTTGTTCACTTCCCAGGGCTCTCTGTCCACCATTTCGGCGCAGGCTTTCGCTCGGCGGGCCGTCTCGGGATCGGCGTCTTTCCGGTTTTG** 68499
WF-Teratorn TTTGCACATGGTGTACCAGTCTTTCTGGATCTTGTTCACTTCCCAGGGCTCTCTGTCCACCATTTCGGCGCAGGCTTTCGCTCGGCGGGCCGTCTCGGGATCGGCGTCTTTCCGGTTTTG 71714

TeratornLC199500 **GCTGTAGCAGCGCAACTGCCGAGCGAGCGAACCGCACTCCTTCCTGATCAGCTCCATGACCGAGGCAGCGAACCTATCCCGCTCGCTTTTTCTGAACGTAGCGTCCGAGGCGTACCTGCT** 68619
WF-Teratorn GCTGTAGCAGCGCAACTGCCGAGCGAGCGAACCGCACTCCTTCCTGATCAGCTCCATGACCGAGGCAGCGAACCTATCCCGCTCGCTTTTTCTGAACGTAGCGTCCGAGGCGTACCTGCT 71834

TeratornLC199500 **AAGGAACTCGCAGTACCTCTGCAGCTTCCCAAGTGGACTGTGCAGCAGCTCGGCCACGGCGAATGCGCCTGGGATCTTGGCGTTCATGAACGTGATGGCGAGTTCAGCGATTTCGATCAT** 68739
WF-Teratorn AAGGAACTCGCAGTACCTCTGCAGCTTCCCAAGTGGACTGTGCAGCAGCTCGGCCACGGCGAATGCGCCGGGGATCTTGGCGTTCATGAACGTGATGGCGAGTTCAGCGATTTCGATCAT 71954

TeratornLC199500 **CCTGGGGTCCGCCAACTTGCTCTCCACGTGTCTGGGTATGCGGTCGCACCACCCGTTCGGATCGACGCGCTTGGTGAACGCAGACAGCGCCCATTGGTTGATTTCTTGGAACGTGAGCTG** 68859
WF-Teratorn CCTGGGGTCCGCCAACTTGCTCTCCACGTGTCTGGGTATGCGGTCGCACCACCCGTTCGGATCGACGCGCTTGGTGAACGCAGACAGCGCCCATTGGTTGATTTCTTGGAACGTGAGCTG 72074

TeratornLC199500 **CCCCGACAGATACGTCTCCCGGGCCATCTCCTGGAGCGTCTCGTTCCAGCAGCTCGCGAGCTCTCCGTTGCCTTTCACGCAGCTGCGTTTCAAGAGCCTGTAGAACCGGAGCGTGGTGGA** 68979
WF-Teratorn CCCCGACAGATACGTCTCCCGGGCCATCTCCTGGAGCGTCTCGTTCCAGCAGCTCGCGAGCTCTCCGTTGCCTTTCACGCAGCTGCGTTTCAAGAGCCTGTAGAACCGAAGCGTGGTGGA 72194

TeratornLC199500 **CACGTACAGCGTGGGCAAGTGTCTCTGCTCCGCTCTCACGGCCACCGTCGGCACCAGGTTGCTTCGCCTCCCATCTGACGCTTTCCCTCCTGACCTGAAGTACACCATGCGGGGTTCGCT** 69099
WF-Teratorn CACGTACAGCGTGGGCAAGTGTCTCTGCTCCGCTCTCACGGCCACCGTCGGCACCAGGTTGCTTCGCCTCCCATCTGACGCTTTCCCTCCTGACCTGAAGTACACCATGCGGGGTTCGCT 72314

TeratornLC199500 **GGCTTTGTACGTGAAACGACTCAGGTCGCATTTCACTGCGTCCTCCCCGAAACAGTCAACGCCGGCGTCCCCCTCGTTCTTCCTGTCGCTCGAGGACTCCCATGCGTGGCTAACTTCTGG** 69219
WF-Teratorn GGCTTTGTACGTGAAACGACTCAGGTCGCATTTCACTGCGTCCTCCCCGAAACAGTCAACGCCGGCGTCCCCCTCGTTCTTCCTGTCGCTCGAGGACTCCCATGCGTGGCTAACCTCTGG 72434

TeratornLC199500 **CGCGCAGCACATCGGCTCCACATCTTCCAGGTGAGAGGCCAACGCAGCCTGGCTGCTCTGATCCCTGCCGGGGGACCCGGGTGTGTTCGATATCAGCAGTTGACAAGGCTGAATCTGCTG** 69339
WF-Teratorn CGCGCAGCACATCGGCTCCACATCTTCCAGGTGAGAGGCCAACGCAGCCTGGCTGCTCTGATCCCTGCCGGGGGACCCGGGTGTGTTCGATATCAGCAGTTGACAAGGCTGAATCTGCTG 72554

TeratornLC199500 **GAAGGACGGACAGAGCGCCTCCACGTGCGTGGGCATCTCCAGCGACATGTCCCAAAAGTCGAACGCGTGGCCCGCGACCTTCCTCAGCAGCTTCAGGCTGTCCAGGGTCGGGGCGAACAG** 69459
WF-Teratorn GAAGGACGGACAGAGCGCCTCCACGTGCGTGGGCATCTCCAGCGACATGTCCCAAAAGTCGAACGCGTGGCCCGCGACCTTCCTCAGCAGCTTCAGGCTGTCCAGGGTCGGGGCGAACAG 72674

TeratornLC199500 **GACCTCTTGCACGAAGTAAGACCCCTTGCTTTCGCCGTCGTTTCCGGGGATGTCTGTCATGTAGGTCCCGACCATGGACCTCGCGGGGTCGTAACGCAGCAGGTTCTCCTTCGTCGAGTG** 69579
WF-Teratorn GACCTCTTGCACGAAGTAAGACCCCTTGCTTTCGCCGTCGTTTCCGGGGATGTCTGTCATGTAGGTCCCGACCATGGACCTCGCGGGGTCGTAACGCAGCAGGTTCTCCTTCGTCGAGTG 72794

TeratornLC199500 **CTGCGTCACCACGGGCACCACCGTCCGATCCTTTCCCCTCAGCTCCTCGGCGGGGAGGTAGTACGCCGTGTAATTTCCGCACGTCAGCAACACTTGCTTGAGGCCGGCTCTGTCCGAAGT** 69699
WF-Teratorn CTGCGTCACCACGGGCACCACCGTCCGATCCTTTCCCCTCAGCTCCTCGGCGGGGAGGTAGTACGCCGTGTAATTTCCGCACGTCAGCAACACTTGCTTGAGGCCGGCTCTGTCCGAGGT 72914

TeratornLC199500 **GACGATGGATCCGAAGTCGCTGCCTCCCAGAAGGTCCACCAGGTCCATAAGGGCGAGCAGGAGAGATCGAGAGGGACGCTGGGGTATCCTACCAGAGTCCGCTATGACCGCGTCTCTGAT** 69819
WF-Teratorn GACGATGGATCCGAAGTCGCTGCCTCCCAGAAGGTCCACCAGGTCCATAAGGGCGAGCAGGAGAGATCGAGAGGGACGCTGGGGTATCCTACCAGAGTCCGCTATGACCGCGTCTCTGAT 73034

TeratornLC199500 **CATCTTGTTCAGAACCCTCGGCCTGTAACGCAGGTGCTCTTCGTTGCTCAGGCGCCGTTCCTTTTGGGTCATCATGCTTACCACTGCATCGAAAACGCCACGTGGGTGCGAGCACGTGAA** 69939
WF-Teratorn CATCTTGTTCAGAACCCTCGGCCTGTAACGCAGGTGCTCTTCGTTGCTCAGGCGCCGTTCCTTTTGGGTCATCATGCTTACCACTGCATCGAAAACGCCACGTGGGTGCGAGCACGTGAA 73154

TeratornLC199500 **GACCTCGTCGTGGTCCACAGTATCGAAGTACCACGGAGGGCTGCTGGAAACGACGAAGGTTTGATTTGTAGACGCGGCCGCCTCAGTCGCCGATACCGTAGCCAT**TTCGAAAGCAAGACC 70059
WF-Teratorn GACCTCGTCGTGGTCCACAGTATCGAAGTACCACGGAGGGCTGCTGGAAACGACGAAGGTTTGATTTGTAGACGCGGCCGCCTCAGTCGCCGATACCGTAGCCATTTCGAAAGCAAGACC 73274

TeratornLC199500 TGTTGAATGCGCGTACGGACAAAGTGGAAACAGCAGCAACAGCTCTCCTAATCCACTCTGGTGAATTCAGCCTAAGTCCTGCGAGATCCTTGCGCTCTCTCTAGCTCTTAGAGGGGAGGT 70179
WF-Teratorn TGTTGAATGCGCGTACGGACAAAGTGGAAACAGCAGCAACAGCTCTCCTAATCCACTCTGGTGAATTCAGCCTAAGTCCTGCGAGATCCTTGCGCTCTCTCTAGCTCTTAGAGGGGATGT 73394

TeratornLC199500 TCGAAGGCCACTTATATCCTCCCAGTTGGGGTTTCCCACCTCCTCGATGCTTTGAACAGCCCAACCCTGATCCTTGCGCTCTCTCTAGCTCTTAGAGGGGAGGTTCGAAGGCCACTTACA 70299
WF-Teratorn TCGAAGGCCACTTATATCCTCCCAGTTGGGGTTTCCCACCTCCTCGATGCTTTGAACAGCCCAACCCTGATCCTTGCGCTCTCTCTAGCTCTTAGAGGGGAGGTTCGAAGGCCACTTATA 73514

TeratornLC199500 TCCTCCCAGTTGGGGTTTCCCACCTCCTCGATGCTTTGAACAGCCCAACCCTGAGGCTCGGAGGGATAGGGCGGAGAAATCTGCTCACCACCACTGCACACAGGCCCAACCCT------- 70412
WF-Teratorn TCCTCCCAGTTGGGGTTTCCCACCTCCTCGATGCTTTGAACAGCCCAACCCTGAGGCTCGGAGGGATAGGGCGGAGAAATCTGCTCACCACCACTGCACACAGGCCCAACCCTGAGGCTC 73634

TeratornLC199500 ---------------------------------------------------------TAGGCTAGGAGGGATAGGGTGGAGAAATCTGCTCACCACCACTGCACACACACTCGCAAGCAACTT 70478
WF-Teratorn GGAGGGATAGGGCGGAGAAATCTGCTCACCACCACTGCACACAGGCCCAACCCTTAGGCTAGGAGGGATAGGGTGGAGAAATCTGCTCACCACCACTGCACACACACTCGCAAGCAACTT 73754

TeratornLC199500 GCACGGAGGCCAGTCATGAGACAAACAATCCGACTGCTCTTTGCAGTGTCCGCCGGCCTTTCACAGTGTCCGACCGTCGCGCAGGGAGTCCGACCGCTGCTCAGAACATCCGACCGCCGC 70598
WF-Teratorn GCACGGAGGCCAGTCATGAGACAAACAATCCGACTGCTCTTTGCAGTGTCCGCCGGCCTTTCACAGTGTCCGACCGTCGCGCAGGGAGTCCGACCGCTGCTCAGAACATCCGACCGCCGC 73874

TeratornLC199500 TCAGTGACTCTGACCGCCGTTCAGTGACTCCGACCGTCGCTCAGTGACTCCGACCGATAGTCAGAGTACGACCGCCGATCCCTGTGTACACTGAATTCAGGGAGTCCCAGTGACACGGAG 70718
WF-Teratorn TCACTGACTCTGACCGCCGTTCAGTGACTCCGACCGTCGCTCAGTGACTCCGACCGATAGTCAGAGTACGACCGCCGATCCCTGTGTACACTGAATTCGGCGAGTCCCAGTGACACGGAG 73994

TeratornLC199500 CCTACAGAGAGAGCCTTTGTGCACGGACAGTGATGAAAATGAAATATCACAGTCACCTAGGTTGAATTCAAAGTCACCTAGGCTTGCTTGTAGCTTTGTTGTTTGTTTGGTTTTCTCCTC 70838
WF-Teratorn CCTACAGAGAGAGCCTTTGTGCACGGACAGTGATGAAAATGAAATATCACAGTCACCTAGGTTGAATTCAAAGTCACCTAGGCTTGCTTGTAGCTTTGTTGTTTGTTTGGTTTTCTCCTC 74114

TeratornLC199500 ATACAGCTTTTTATTTACCCCCCTCTGTGCTCAAAGAGAAAATCGTCGGGCTTTGCGATTTGAGGAGCAGAGCTCAGC**ATGGATGAGGAAATTAGAATTTTGATCAAAGGTGTTCTGGGG** 70958
WF-Teratorn ATACAGCTTTTTATTTACCCCCCTCTGTGCTCAAAGAGAAAATCGTCGGGCTTTGCGATTTGAGGAGCAGAGCTCAGCATGGATGAGGAAATTAGAATTTTGATCAAAGGTGTTCTGGGG 74234

TeratornLC199500 **CACTTCGTTGGACGACGCCAAGACTTTGAAGTCGAAAGGCCGCGTTGTCGGGAAGCTGTAAATGCTATGACGAGAGCTGTGTGTGAAGTGCTGGAGGTTCACAGACCGAAATACACAG**GT 71078
WF-Teratorn CACTTCGTTGGACGACGCCAAGACTTTGAAGTCGAAAGGCCGCGTTGTCGGGAAGCTGTAAATGCTATGACGAGAGCTGTGTGTGAAGTGCTGGAGGTTCACAGACCGAAATACACAGGT 74354

TeratornLC199500 GAGCAGCACAAACAGATCTGTTTTTTCAGAGTTATCTGTACGCTCAGCGTGATCTCTCCTTGTCCGGTGTCCCACTTTAG**AGATGGTTTCAGATCTAGGTGATTCATTTCCCGATGCCGC** 71198
WF-Teratorn GAGCAGCACAAACAGATCTGTTTTTTCAGAGTTATCTGTACGCTCAGCGTGATCTCTCGTTGTCCGGTGTCCCACTTTAGAGATGGTTTCAGATCTAGGTGATTCATTTCCCGATGCCGC 74474

TeratornLC199500 **GATGGTCAACGTCGCGAAAAGCCTTTTCGAGGACGGGATCATCAACTGGGGCCGTGTGGTCAGTCTGATATGTTTGGGGGCCGCGGTGTGCCAACATCAGGACAGGTTCAGTGGCAGCCC** 71318
WF-Teratorn GATGGTCAACGTCGCGAAAAGCCTTTTCGAGGACGGGATCATCAACTGGGGCCGTGTGGTCAGTCTGATATGTTTGGGGGCCGCGGTGTGCCAACGTCAGGACAGGTTCAGTGGCAGCCC 74594

TeratornLC199500 **TGGCTCCCTCTTGGCTGACACAGTGTCAGAGGCGATCGCCTCGTACTTGCTCACAGAGCACAGAGGCTGGCTGGAAGAGATGTGTTGGGTGAGTGGCCTGTACTCTAGGGGGGTTGAACTT** 71438
WF-Teratorn TGGCTCCCTCCTGGCTGACGCAGTGTCAGAAGCGATCGCCTCGTACTTGCTCACAGAGCACAGAGGCTGGCTGGAAGAGATGTGTTGGGTGAGTGGCCTGTACTCTAGGGGGGTTGAACTT 74714

TeratornLC199500 **CATGCTATCGCTGTTATGA**AACAATTTGTCTTCATGTTTTGCTTTTTTTTTTTTCTGTTCCTCTTTCTGTTCTTTTTTTAACGCTTTCTTCATGACCAGGATGACTTCGCTGACTTTTTC 71558
WF-Teratorn CACGCTATCGCTGTTATGAAACAATTTGTCTTCATGTTTTGCTTTTTTTTTT--CTGTTCCTCTTTCTGTTCTTTTTTTAACGCTTTCTTCATGACCAGGATGACTTCGCTGACTTTTTC 74832

TeratornLC199500 CCTTCACCTGAAGCCACGAAGTGCCATGACCTTGTAGGACCGGCACTTGGACTCTTTGTTTTTGCGGTCGGAACTGCACTCTTGTGCACGCACCTTGGGATGTGACCCGGTGGTTGTTTT 71678
WF-Teratorn CCTTCACCTGAAGCCACGAAGTGCCATGACCTTGTAGGACCGGCACTTGGACTCTTTGTTTTTGCGGTCGGAACTGCACTCTTGTGCACGTACCTTGGGATGTGACCCGGTGGTT-TTTT 74951

TeratornLC199500 CTGACAGCCTCCGGGGGCTTTGCCCTCAGAGTCCTCTAAAGTCTGGAGGGCCAGTGTCAGACACTGAAACAACCGGTGTGGACGAGAGGACGCTTCTTAGACACGCACTTTAAGGCATTG 71798
WF-Teratorn TTGACAGCCTCCAGAGGCTTTGCCCTCAGAGTGCTTTGAAGTCTGGAGGGCAAGTGTCAGACACTGAAACAACCGGTGTGGACGAGAGGACACTTCTTAGACACGCACTTTAAGGCATTG 75071

TeratornLC199500 GGAACGAGGAGACGTTCCTCAGGAGCCCGTTTGATCCTTTTCTTGTTTTTCTTTTGCTCCCGC**ATGGAGAAGTACTACAAAGACCCAGTGCGACCAGTGATCGGGCAAGGCACCTACGGA** 71918
WF-Teratorn GGAACGAGGAGACGTTCCTCAGAAGCCCGTTTGATCCTTTTATTGTTTTTCTTTTGCTTCCGCATGGAGAAGTACTACAAAGACCCAGTGCGACCAGTGATCGGGCAAGGTACCTACGGA 75191

TeratornLC199500 **GTTGTCTACAGGGTGCGAGATGTGAGAAACGGCCAAGAATATGCCCTCAAATGTCACAAGCAGGCAGACGTTATAGAGGACGCGACTGTGAGAGAGTTGTCCTGCCTGACAGCGCTGAAA** 72038
WF-Teratorn GTTGTCTACAGGGTGCGAGATGTAAGAAACGGCCAAGAATATGCCCTCAAATGTCACAAGCAGGCAGACGTTATAGAGGACGCGACTGTGAGAGAGTTGTCCTGCCTGACAGCGCTGAAA 75311

TeratornLC199500 **GGTCATCCCAACATAATACAGCTCCTGGACTGTTTCATGGACGAAGATCGGGTGGCGATGCTCATGCCTCACGTGCCCTTCGAGCTGACCAAAGCTATTCACCGCCAGCCACCGGCTCCG** 72158
WF-Teratorn GGTCATCCCAACATAATACAGCTCCTGGACTGTTTCATGGACGAGGATCGGGTGGCGATGCTCATGCCTCACGTGCCCTTCGAGCTGACCAAAGCTATTCACCGCCAGCCACCGGCTCCG 75431

TeratornLC199500 **CTCAGCTTCGTGGCCAGCTTCTCGCTTCAGACGGCGAACGCCCTGGAGTACATGCACGGACTGAACATCTTACACAGGGACCTGGTTCCGAGTAACGTGCTTCTCGAGGAAGATCTGACG** 72278
WF-Teratorn CTCAGCTTCGTGGCCAGCTTCTCGCTTCAGACGGCGAATGCCCTGGAGCACATGCACGGACTGAACATCTTACACAGGGACCTGGTTCCGAGTAACGTGCTTCTCGAGGAAGATCTGACG 75551

TeratornLC199500 **GTGAAGGTGGGCGACATGGGCCTAGCCAGACAGTACTCCCGACGCATGAGCCTTTGCGTGGTGACGGAGCCCTACAGGGCTCCGGAGTTGTTTGTCCGCGGGAACGCCGCGGAGTACACG** 72398
WF-Teratorn GTGAAGGTGGGCGACATGGGCCTAGCCAGACAGTACTCCAGACGCATGAGCCTTTGCGTGGTGACGGAGCCCTACAGGGCTCCGGAGTTGTTTGTCCGCGGGAACGCCGCGGAGTACACG 75671

TeratornLC199500 **TGCGCCGTAGACATGTGGAGTTTGGGAGCTATGATACTGGACGCAAAGGAAGGCGCCGTGGTGTTCGCTGCAACGCGTGGGTCCAGCACTCTGAGCAAAGTCATGGCACTGGCGGCCTCG** 72518
WF-Teratorn TGCGCCATAGACATGTGGAGTTTGGGAGCTATGATACTGGACGCAAAGGAAGGCGCCGTGGTGTTCGCTGCAACGCGTGGGTCCAGCACTCTGAGCAAAGTCATGGCACTGGCGGCCTCG 75791

TeratornLC199500 **GATCCCGACGGGGGAGCGGAGGAATCGACGCCCGCCAGGCAGGAACTGATACCCAACACTATGAAGTGTGAGCACACCAAACGCATCGCCTTCCGCCTAATCGCTCTCAGGCACGACAAG** 72638
WF-Teratorn GATCCCGACGGGGGAGCGGAGGAATCGACGCCCGCCAGGCAGGAACTGATACCCAACACTATGAAGTGTGAGCACACCAAACGCATCGCCTTCCGCCTAATCGCTCTCAGGCACGACAAG 75911

TeratornLC199500 **AGGCTTACGGCTCGCCAGCTGTTGGACGATGAGGAATGGAAGCGTTTGTCCCACATGACAGACGGAGAAAAGGCTGAGGTCTTGCAACAATGCGTCACGGGTGTCATGTAA**AAATACACA 72758
WF-Teratorn AGGCTTACGGCTCGCCAGCTGTTGGACGATGAGGAATGGAAGCGTTTGTCCCACATGACAGACGGAGAAAAGGCTGAGGTCTTGCAACAATGCGTCACGGGTGTCATGTAAAAATACACA 76031

TeratornLC199500 GTTCAAGAGGCTTTGCAATGTCTGAGTTTATCATTTTTCAATAAAAAAAAAATGAGCCAATGACCACAGACTCTATGTATCGTACAGGTGTAAATTTTGTTTATTATTTTGAATACGTTT 72878
WF-Teratorn GTTCAAGAGGCTTTGCAATGTCTGAGTTTATCAATTTTCAATAAAAAAAA--TGAGCCAATGACCACAGACTCTATGTATCGTACAGGTGTAAATTTTGTTTATTATTTTGAATACGTTT 76149

TeratornLC199500 GACCATAATAGCAGATGTATCATTCAGCACCACTGGCCCGCTCTGGGAGCACGCTTACACGTTTGTAGCGATGGAGATGAAAAGGGATGTAGCACTCAACGTGGAACACCACTTGGCGTA 72998
WF-Teratorn GACCATAATAGCAGATGTATCATTCAGCACCACTGGCCCGCTCTGGGAGCACGCTTACACGTTTGTAGCGATGGAGATGAAAAGGGATGTAGCACTCAACGTGGAACACCACTTGGCGTA 76269

TeratornLC199500 GTTTGTTTCGTAACTGCGCGAGGGCCCTCGGGTCTTTTGAGGGCTTCGGGCTCAACTCAGCTTCCGCTCTTCGTCTCAGAGCGCGGCGAAAACAACAGTGGCAGCCGCTGCGACGAGAGA 73118
WF-Teratorn GTTTGTTTCGTAACTGCGCGAGGGCCCTCGGGTCTTTTGAGGGCTTCGGGCTCAACTCAGCTTCCGCTCTTCGTCTCAGAGCGCGGCGAAAACAACAGTGGCAACCGCTGCGACGAGAGA 76389

TeratornLC199500 CAGGGCTAGAGGTTTGGCTCCATTTCTCTGACTAGGCCATCTATCCAGTTCCTCCTGAGCCTTTCTCACGGTTTCCTTCAACTTGCTGAGCACTGCAAACAGAAGACAGCGAACACTGGT 73238
WF-Teratorn CAGGGCTAGAGGTTTGGCTCCATTTCTCTGACTAGGCCATCTATCCAGTTCCTCCTGAGCCTTTCTCACGGTTTCCTTCAACTTGCTGAGCACTGCAAACAGAAGACAGCGAACACTGGT 76509

TeratornLC199500 TGGGAGAGAAGCGCTTTGCGATAGGCGGGTAGCTAAAATGCACACATACATACATACAGTCTGTCGTGTTGTGGAGTTTTCCGAAAGTCATCACGGGCTCCATTGGAACCAGGTCAGGCA 73358
WF-Teratorn TGGGAGAGAAGCGCTTTGCGATAGGCGGGTAGCTAAAATGCACACATACATACATACAGTCTGTCGTGTTGTGGAGTTTTCCGAAAGTCATCACGGGCTCCATTGGAACCAGGTCAGGCA 76629

TeratornLC199500 TGAGAATCTCAGGCCGACGAACGCCGTCCGAGTAAGAAGGCTCGTTGCTGTGATGTAAAACACAAGAGTCCACTGATGTCAACAGCACGAACACGAACAGGGTCAGATGCAGCATCGAAG 73478
WF-Teratorn TGAGAATCTCAGGCCGACGAACGCCGTCCGAGCAAGAAGGCTCGTTGCTGTGATGTAAAACACAAGAGTCCACTGATGTCAACAGCACGAACACGAACAGGGTCAGATGCAGCATCGAAG 76749

TeratornLC199500 TGGTAGTCATTGTGTCTTCAAGAGAAAAAAAAC-CCCACAAAGAACAGTTTTCTTACTGCAGTGCGCAAACACAACTCACGCCTGAAGAGGTTGCCTCTGTCAGCTCTCCACTCTTGCCT 73597
WF-Teratorn TGGTAGTCATTGTGTCTTCAAGAGAAAAAAAAACCCCACAAAGAACAGTTTTCTTACTGCAGTGCGCAAACACAACTCACGCCTGAAGAGGTTGCCTCTGTCAGCTCTCCACTCTTGCCT 76869

TeratornLC199500 TTTATACTTGCAGAAGAAAACACTCGCTGTGCAGATTCTAATCCCCCCTGCCCAAAAAAAAAAAAAAAAGTCCCTGGCCAAATCCCAACCACAAGCCCGAAAGAGAGGCCGCGACCGGTCA 73717
WF-Teratorn TTTATACTTGCAGAAGAAAACACTCGCTGTGCAGATTCTAATCCCCCCTGCCCAAAAAAAAAAAAAAAAGTCCCTGGCCAAATCCCAACCACAAGCCCGAAAGAGAGGCCGCGACCGGTCA 76989

TeratornLC199500 GCTCGCGTAAACCCAGCACGCCCCTCGCTCACAGGCAGGAGAGATCACGTAACAGGGTGATTCAGGGGAGAGGAACGCAGAGCATGGCGTCAGAGTCTGCCTGCCTCGGTCGGCCTCTCA 73837
WF-Teratorn GCTCGCGTAAACCCAGCACGCCCCTCGCTCACAGGCAGGAGAGATCACGTAACAGGGTGATTCAGGGGAGAGGAACGCAGAGCATGGCGTCAGAGTCTGCCTGCCTCGGTCGGCCTCTCA 77109

TeratornLC199500 TGGCGGAGGACGGATTCTCAGAGTGGAAATCGGTGTGAGAGAGCTGAGGTCGGAGCCAGCGTTTCTCAGTCCCTCCCCTACATCTCGGTTTGGTAGCGTAAGATGCCATCTGACTCAGTG 73957
WF-Teratorn TGGCGGAGGACGGATTCTCAGAGTGGAAATCGGTGTGAGAGAGCTGAGGTCGGAGCCAGCGTTTCTCAGTCCCTCCCCTACATCTCGGTTTGGTAGCGTAAGATGCCATCTGACTCAGTG 77229

TeratornLC199500 ATGTGTGGTCGGACTCCCTAAGGGTCGGTCGGACAGAGTGAATGACTGTCGGACTCCCTTAGCGCCGGTCGGACAGAGTCAATTGCGGTCAGACAGAGTGAATAGAAAAGGTCGGGCTGT 74077
WF-Teratorn ATGTGTGGTCGGACTCCCTAAGGGTCGGTCGGACAGAGTGAATGACTGTCGGACTCCCTGAGCGCCGGTCGGACAGAGTCAATTGCGGTCAGACAGAGTGAATAGAAAAGGTCGGGCTGT 77349

TeratornLC199500 CTGGACGCTGACCTGGCGAGGTGAAAAGCTGGCCCCATCTCTCGTCTGCAGCTTTGGAACTAGGAAATTTTAGCGGCTGTGTTCTCTGTGTTCTATGACACAGCTCCTACGTAGAAAGGA 74197
WF-Teratorn CTGGACGCTGACCTGGCGAGGTGAAAAGCTGACCCCATCTCTCGTCTGCAGCTTTGGAACTAGAAAATTTTAACGGCTGTGTTCTCTGTGTTCTATGACACAGCTCCTACGTAGAAAGGA 77469

TeratornLC199500 TTTAGTTGGGGATTTTTTATTCTTTTTTTTTT-CTTTCCTTCATATCCAACATCGACATCACAGCCCTTACCAACAAA**ATGGTTGGATTTATCTACAATTCCAATAAGCTGCTCCCCGCT** 74316
WF-Teratorn TTTAGTTGGGGATTTTTTATTCTTTTTTTTTTTCTTTCCTTCATATCCAACATCGACATCGCAGCCCTTACCAACAAAATGGTTGGATTTATTTACAATTCCCATAAGCTGCTCCCCGCT 77589

TeratornLC199500 **CTGGCCCCATCAGAGCCCCGAGCCGCTCTGAACACCCAACAGCTCTCACAGAGCCGGACATCCTCTCGCTTCATCAACCATCACGGATCCCCCACGTCTGCCCTTACCAACAAAATGAGC** 74436
WF-Teratorn CTGGCCCCATCAGAGCCCCGAGCCGCTCTGAACACCCAACAGCTCTCACAGAGCCGGACATCCTCTCGCTTCATCAACCATCACGGATCCCCCACGTCTGCCCTTACCAACAAAATGAGC 77709

TeratornLC199500 **AAATCGGGCTGCTACAGAGACGGCGGCATGCCGCAGCAGAAACCTGACTGTCTGAACATTTCCGCCAATTACAAGACTAAATTGTGCTGGTTCTTCGCGAAGAACGGCGTCTGCAAGTAC** 74556
WF-Teratorn AAATCGGGCTGCTACAGAGACGGCGGCATGCCGCAGCAGAAACCTGAATGTCTGAACATTTCCGCCAATTACAAGACTAAATTGTGCTGGTTCTTTGCGAAGAACGGCGTCTGCAAGTAC 77829

TeratornLC199500 **GGAGGGAAATGTCGGTACGCCCATGGCTACGGAGACCTGCGGAGCCTGCAGCTCAAAATGAACCTTTGCCACCATTTCTATAAGGGGCGCCCATGCCTCCACGGCGGAAGTTGTGGATTT** 74676
WF-Teratorn GGAGGGAAATGTCGGCACGCCCACGGCTACGGAGACCTGCG---------------AACGAACCTTTGCCACCATTTCTATAAGGGGCGCCCATGCCTCTACGGCGGAAGTTGTGGATTT 77934

TeratornLC199500 **ATTCACAATTCCCATCAGCTGCTCCCCGCTCAGCCCACATCAGAGCCCCGAGCCGCTCTGAACAAGGACGGCTACCGACAGATGGACGTTTCCCCACCCCAGCAGCTCTCACAGAGCCTG** 74796
WF-Teratorn ATTCACAATTCCCATCAGCTGCTCCCCGCTCAGCCCACATCAGAGCCCAGAGCCGCTCTGAACAAGGACGGCTACCGACAGATGGACGTTTCCCCACCCCAGCAGCTCTCACAGAGCCTG 78054

TeratornLC199500 **ACATCCTCTGGCTTCTTCAACCATCGCGGACCCTCCACAGTCAATGCCGGGGCCATCACCTGCTGCATCCCCACCGTCCCCGCCGTCGTCTTCTCGAAAGTCCCAAATCCCTGTAACCCC** 74916
WF-Teratorn ACATCCTCTGGCTTCTTCAACCATCACGGACCCTCCACAGTCATTGCCGGCGCCATCACCTGCTGCATCCCCACCGTCCCCGCCGTCGTCTTCTCGAAAGTCCCAAATCCCTGTAACCCC 78174

TeratornLC199500 **TGCCCCGGCCTTATCCCCACCGTCCCCGCCGTCGTCTTCTCGAAAGTCCCAAATCCCTGTCACCCCTGCCTCGGCCTTATCCCCACCGTCCCCGCCGTCGTCTTCTCGACAGTCCCAAAT** 75036
WF-Teratorn TGCC--------------------------------------------------------------TCGGCCTTATCCCCACCGTCCCCGCCGTCGTCTTCTCGACAGTCCCAAAT 78228

TeratornLC199500 **CCCTGTAACCTCTGCCCCGGCCTTCTTCTGCTTCACTAG**CAAAGAGGGGCTTTGTCCAAAATTTGTGACGCGCTGGCCTGAACGGTCTACCCGGTTTCTCTCGGGGCCCTGTGGTCATGC 75156
WF-Teratorn CCCTGTAACCTCTGCCCCGGCCTTCTTCTGCTTCACTAGCAAAGAGGGGCTTTGT-----------GACGCGCTGGCCTGAACGGTCTACCCGGGTTCTCTCGGGGTCCTGTGGTCATGC 78337

TeratornLC199500 GCAACAAGGCTTATGGCATCGAGGTCTTTATTTGTTTTTTTTGTGGTGTTTGTTTAAAATAAAAAATAAAACAAGACAAGAACAACGTGTTTTTACTACATTGTATCTGAAAAGATTCAA 75276
WF-Teratorn GCAACAAGGCTTATGGCATCGAGGTCTTTATTTGTTTTTTT-GTGGTGTTTGTTTAAAATAAAAAATAAAACAAGACAACAACAACGTGTTTTTACTACATTGTATCTGATAAGAGTCAA 78456

TeratornLC199500 AGTGTGTTATTCCGTGTTGTTTCGCTGACCCTGATATCGTTGAGCTTCTTAAGCAAATAAAAAGACTCAGACACATTCACTTTTTCTGCTCTTTATTCGTTTGCCGTGGACATTCACAGA 75396
WF-Teratorn AATGTGTT-TTCCGTGTTGTTTCGCTGACCCTGATATCGTTGAGCTTCTTAAGCAAATAAAAAGACTCAGACACATTCACTTTTTCTGCTCTTTATTCGTTTGCCGTGGACATTCACAGA 78575

TeratornLC199500 GTTGTCTTGGCGTCGCAGTATTCTGTCTGAAAATCTTTGGTGCAGACTTATACGCACTTTCCAGCCGTGC**TCACTTGCGTTTTACGCAACGTATCTCCTCCAAAGAAACAGGCTGTATTC** 75516
WF-Teratorn GTTGTCTTGGCGTCGCAGTATTCTGTCTGAAAATCTTTGGTGCAGACTTT--CGCACTTTCCAGCCGTGCTCACTTGCGTTTTACGCAACGTATCTCCTCCAAAGAAACAGGCCGTATTC 78693

TeratornLC199500 **TTGTGCCCCAGTGCTTGTCCCTGTCAGGGGGGACGTGCAGCAAAGTCCAGGTGGGTGTTTGTTCGGCAATGCCCTCCAAGAGTCTTCTTTCACAGCAGCGGATGACGCCGTGCATTCCGT** 75636
WF-Teratorn TCGTGCCCCAGTGCTTGTCCCTGTCAGGGGGGACGTGCAGCAAAGTCCAGGTGGGTGTTTGTTCGGCAATGCCCTCCAAGAGTCTTCTTTCACAGCAGCGGATGACGCCGTGCATTCCGT 78813

TeratornLC199500 **TGGCAAGTATTGTCCCTTCGCCGGAGTAATTCGTGCAGAGCCACGCCACCCCCATGCGCCGCAAATCCGGGTCTCCGTTCTTCTCTGTTGCAATCGCGTACTCCACGAACAAGTGAGTCT** 75756
WF-Teratorn TGGCAAGTATTGTCCCTTCGCCGGAGTAATTCGTGCAGAGCCACGCCACCCCCGTGCGCCGCAAATCCGGGTCTCCGTTCTTCTCTGTTGCAATCGCGTACTCCACGAACAAGTGAGTCT 78933

TeratornLC199500 **TTCCCACGCGCCTCACCCGTACGGGCAACGCTGCGATGCCGGCCGCTATCAAAGCATTCGCTCCGCTCAGAGCATCAACGTCAGAGTCCCAACCGGCGAAATCTTCGTCTGGACAAGCGC** 75876
WF-Teratorn TTCCCACGCGCCTCACCCGTACGGGCAACGCTGCGATGCCGGCCGCGATCAAAGCATTCGCTCCGCTCAGAGCATCAACGTCAGAGTCCCAACCGGCGAAATCTTCGTCTGGACAAGCGC 79053

TeratornLC199500 **AGCGGCCAAGGGGGTAGCGACCATCTTCCGTTCTACCACCCATTCCTTCGAGCTCCGCGTAAGTCGTCTCCTCGGATGAGGTTCTGACGATCAGCGTTCTGCGACCGTTGGTTTCACCCA** 75996
WF-Teratorn AGCGGCCAAGGGGGTAGCGACCATCTTCCGTTCTACCACCCATTCCTTCGAGCTCTGCGTAAGTCGTCTCCTCGGATGAGGTTCTGACGATGAGCGTTCTGCGACCGTTGGTTTCACCCA 79173

TeratornLC199500 **ACCGGAGCTTGTGCAT**TTTGAGGGTCTCCGTGAGTCTCCGTGATCTCAGAACGGTCAGGTTGCCTTTCTTCTTCTCGGCCCCTTGGGTCGTCCTCCCGTTGCCGCAGCCTCATTAGTATG 76116
WF-Teratorn ACCGGAGCTTGTGCATTTTGAGGGTCTCCGTGAGTCTCCGTGATCTCAGAACGGTCAGGTTGCCTTTCTTCTTCTCGGCCCCTTGGGTCGTCCTCCCGTTGCCGCAGCCTCATTAGTATG 79293

TeratornLC199500 TCAAAGGAAAAAGGCCCTGACGGGAGAGCTTTCCGTCTGATCAAAAAGTGAATTTCATGATCGCATCTGGAGCAAACTCGAAATGCAAGGGCACGCAATAAAGGCACAGAACCGCTTGAC 76236
WF-Teratorn TCAAAGGAAAAAGGCCCTGACGGGAGAGCTTTCCGTCTGATCAAAAAGTGAATTTCATGATCGCATCTGGAGCAAACTCGAAATGCAAGGGCACGCAATAAAGGCACAGAACCGCTTGAC 79413

TeratornLC199500 ACAAGGACGTCGGTATGAAATCGATTTAAACACCACACAGACTACAGCATTTTATCTTT--TATTTGACTTCAACCATCCAAAACATGTTCCAAAAATAAAAGGATACAGAAGCAGCACA 76354
WF-Teratorn ACAAGGACGTTGGTATGAGATCGATTTAAACACCACACAGACTACAGCATTTTATCTTTTTTATTTGACTTCAATCATCCAAAACATGTTCCAAAAATAAAAGGATACAGAAGCAGCACA 79533

TeratornLC199500 CAATGAACCTAATGGCAAACATAATGTCTCAGGGCGGCAGCCCCACATCCTTCTCTCCCAAACAAGCGTTCCCGAGCCCAGGCTCCTATGCTCTGAACTGGTCCCCTCATGTGACAGCGG 76474
WF-Teratorn CAATGAACCTAATGGCAAACATGATGTCTCAGGGCGGCAGCCCCACATCCTTCTCTACCAAACAGGCGTTCCCGAGCCCAGGCTCCTATGCTCTGAACTGGTCCCCTCATGTGACAGCGG 79653

TeratornLC199500 C**TCAATTTCCGATCAGGGTGATTTCCGAAACCTCCACCACCGGCGCCCTGCCGCTAGCGTTGCTCGCCTTATCTACGTGATCCCAATGCGGCGGATCTGAACAGTTGACCGCGGCGGGTT** 76594
WF-Teratorn CTCAATTTCCGATCAGGGTGATTTCCGAAACCTCCACCACGGGCGCCCTGCCGCTAGCGTCGCTCGCCTTATCTACGTGATCCCAATGCGGCGGATCTGAACAGTTGACCGCGGCGGGTT 79773

TeratornLC199500 **GTGCAGATTTGACCTTTGTCAGCTTCGCCAAATTGTACAAGTACGGCTCCTCTGGGTTTGCGCCTCCCTCACGCGTCGCTACCATGTAGCACCTCGTCTCGCGGGCCGGATCGGGGTCTT** 76714
WF-Teratorn GTGCAGATTTGACCTTTGTCAGCTTCGCCAAATTGTACAAGTACGGCTCGTCTGGGTTTGCGCCTCCCTGACGCGTCGCTACCATGTAGCACCTCGTCTCGCGGGCCGGATCGGGGTCTT 79893

TeratornLC199500 **TACTCAGCACGATGGCGTACAGGTGCTCCGTGGACGCCCCGTGGATCTCGTATCCTTCGCAAGGCGACCAGATAGCGACCAGCTTGCAGCTGAGAAGTTGCGATAGCTGTAGGAACGCTT** 76834
WF-Teratorn TACTCAGCACGATGGCATACAGGTGCTCCGTGGACGCCCCGTGGATCTCGTATCCTTCGCAAGGCGACCAGATAGCGACCAACTTGCAGCTGAGAAGTTGCGATAGCTGTAGGAACGCTT 80013

TeratornLC199500 **TCGAGCACCGTAGATCCACGGCGCTGCATTCCCTGGCGACATCATCTGTGGTCTGGAGCACCCCGCAAAGGGTCTTCGGAGCGCTTTTGCTCTCACCCAAGTGTATGAGCCTCACGTCTT** 76954
WF-Teratorn TCGAGCACCGTAGATCCACGGCGCTGCATTCCCTGGCGACATCATCTGTGGTCTGGAGCACCCCGCAAAGGGTCTTCGGAGCGCTTTCGCTCTCACCCAAGTGTATGAGCCTCACGTCTT 80133

TeratornLC199500 **GATGGGTGTAATGCCGCACAATGTTTGGCAGCGATACGGAGGCCGAGCTCTCCCCGAGTCGCACGCAGTGGACACGTAGGTGCTCCGATCTTTCTGACTTGGCCTCCGTTTTCAAAAAAT** 77074
WF-Teratorn GATGGGTGTAATGCCGCACAATGTTTGGCAGCGATACGGAGGCCGAGCTCTCCCCGAGTCGCACGCAGTGGACACGTAGGTGCTCCGATCTTTCTGACTTGGACTCCGTTTTCAAAAAAT 80253

TeratornLC199500 **CAGGGCGCTTGGAAGAAGCGCAGTTTGATCGGAACGCGTCTCCGCCTGACGAGGCGGCCCTGTGTACTTTCGTCGGCGTGGACGCACCGGTTTTCGAGCCTTCGTCCACCTCCGTCAATT** 77194
WF-Teratorn CAGGGCGCTTGGGAGAAGCGCAGTTTGATCGGAACGCGTCTCCGCCTGACGAGGCGGCCCTGTGTACTTTCGTCGGCGTGGACGCACCGGTTTTCGAGCCTTCGTCCACCTCCGTCAATT 80373

TeratornLC199500 **CGCTTGTGGGGCTGGCCTGCCGGGAGCCCAGGTTCAGCTTTCGCTTGGCCACCCTTCTCCAGGTGTCGCTGCAGGACGCTTCTGTGTGCTCCATCGGAGTTTGCTCCGTACACAGAGATG** 77314
WF-Teratorn CGCTTGTGGGGCTGACCTGCCGGGAGCCCAGGTTCAGCTTTCGCTTGGCCACCCTTCTCCAGGTGTCGCTGCAGGACGCTTCTGTGTGCTCCATCGGAGTTTGCTCCGTACACAGAGATG 80493

TeratornLC199500 **ACACGGTTGTCCTTATGTAATTTTCATAGTCCTGGAACAGCGAGCCTCCGGGGGATCCCATCGCTTCATCTTCGCTGCGGTCGAGTCCTTTTAGCCTTCTGACGCAGTCGTGCCGGAGCA** 77434
WF-Teratorn ACACGGTTGTCCTTATGTAATTTTCATAGTCCTGGAACAGCGAGCCTCCGGGGGATCCCATCGCTTCATCTTCGCTGCGGTCGAGTCCTTTTAGCCTTCTGACGCAGTCGTGTCGGAGCA 80613

TeratornLC199500 **GCCGTGACACTCTCTTCGCAGAGCACAGCGCGGCACACCGGATTCTTTCGATCAGCTCCGTCTCCACTGCGCTTTTCTCCACGCCTATCAGTTGGCACACAACGGCCACGATACCTTTCA** 77554
WF-Teratorn GCCGTGACACTCTCTTCGCAGAGCACAGCGCGGCACACCGGATTCTTTCGATCAGCTCCGTCTCCACTGCGCTTTTCTCCACGCCTATCAGTTGGCACACAACGGCCACGATACCTTTCA 80733

TeratornLC199500 **GAATCTCCGTTTCGTACGTGAGCTTTTCCGTGAGGCACTTTTGTCCGGCTATCGTCTCATCGGCCCGTTTCAGACGTGTCAGGAGGGCCTTGGTCTGGCTCTGACTTTCGTCTGTCGCGG** 77674
WF-Teratorn GAATCTCCGTTTCGTACGTGAGCTTTTCCGTGAGGCACTTTTGTCCGGCTATCGTCTCATCGGCCCGTTTCAGACGTGTCAGGAGGGCCTTGGTCTGGCTCTGACTTTCGTCTGTCGCGG 80853

TeratornLC199500 **TTGCGTTCTCGGCGGCCAGCTCTCGACTCCGCTTCAGTTCGGCCTCCAGCTCGTTGACGCGTTTCGACCGGGACTCGTTTTCGGCCTCCGCTTGCTTCTCTCGGTCCTGCAGCTGGGCGA** 77794
WF-Teratorn CTGCGTTCTCGGCGGCCAGCTCTCGACTCCGCTTCAGTTCGGCCTCCAGCTCGTTGACGCGTTTCGACCGGGACTCGTTTTCGGCCTCCGCTTGCTTCTCTCGGTCCTGCAGCTTGGCGA 80973

TeratornLC199500 **TGAGCTTTGATGCGCGCTCCTCCTCCACCTTGCTTCGATCTAAGTGTTCTGCGGCCAGCTCTCGGCTCTGCTTCAGATGGGCCTCCAGCTCAATTACGCGCTTTGACAGGGACTTCTTCT** 77914
WF-Teratorn TGAGCTTTGATGCGCGCTCCTCCTCCACCTTGCTTCGATCTAAGTGTTCTGCGGCCAGCTCTCGGCTCTGCTTCAGATGGGCCTCCAGCTCAATTACGCGCTTTGACAGGGACTTCTTCT 81093

TeratornLC199500 **CGGCTTCGTTTCGCTTCATTTCGGCCTCTGATTTCTTCGAGAGCATCAAGGCTGTCTCCTTTTCGGCTTTAATTTCCTTCACTTCGTTCTGCAGCTTCTCCAGGACTCGCCCCTTCTCGC** 78034
WF-Teratorn CGGCCTCATTTCGCTTCATTTCGGCATCCGATTTCTTCGAGAGCATCAAGGCTGTCTCCTTTTCGGCTTTCATTTCCTTCACTTCGTTCTGCAGCTTCTCCAGGACTCGCCCCTTCTCGC 81213

TeratornLC199500 **TTAAGTCCTTCTTCGATGAGGCTATGACATTTTCAGCGAAACTTAACTCTTGCCGCAGCTCAGCCAAGTCCAAGTTTGCTTTTCTAACTTTGGCTTCCATCTCCATGTGCTCTTTTGCGA** 78154
WF-Teratorn TTAAGTCCTTCTTCGATGAGGCTATGACATTTTCAGCGAAACTTAACTCTTGCCGCAGCTCAGCCAAGTCCAAGTTTGCTTTTCTAACTTTGGCTTCCATCTCCATGTGCTCTTTTGCGA 81333

TeratornLC199500 **GGTCTATTTTCATCTGCTGATGGGCTGATTTGTCCGTGGTGTACTTCAGGCTAACCTGCCCCAGTTGATCGCTCAGCTCAGCCAAGCGGGACTCGTATGTTTTCTGGTCGCGAGCCTCTC** 78274
WF-Teratorn GGTCTATTTTCATCTGCTGATGGGCTGATTTGTCCGTGGTGTACTTCAGGCTAACCTGCCCCAGTTGATCGCTCAGCTCAGCCAAGCGGGACTCGTATGTTTTCTGGTCGCGAGCCTCTC 81453

TeratornLC199500 **TTGACGCGACTTGGAGACGCTCCGCTAAACTCTGGACTTTGTCATGCAGGACCCAGCCAACACTTACCACACTTTTCCAACAGGCGCGAACGTGTGAGGGACTGAGGCGGCCCTGTGGCT** 78394
WF-Teratorn TTGACGCGACTTGGAGACGCTCCGCTAAACTCTGGACTTTGTCATGCAGGACCCAGCCAACACTTACCACACTTCTCCAACAGGCGCGAACGTGTGAGGGACTGAGGCGGCCCTGTGGCT 81573

TeratornLC199500 **CAGCTTGAGGCCAGTGAAATGTACCACAGCCCGGGGACGCGGACTCAAGTGATTCCAGGGCACGCATCAGGGCCCGCTGCCCACGTTCGTCCTCCGCTTCATTTCCCCCGGCGAGTATCT** 78514
WF-Teratorn CAGCTTGAGGCCAGTGAAATGTACCACAGCCCGGGGACGCGGACTCAAGTGATTCCAGGGCACGCATCAGGGCCCGCTGCCCACGTTCGGCCTCCGCTTCATTTCCCCCGGCGAGTATCT 81693

TeratornLC199500 **CGGACACGACTCCCAGGGTTTCGTCCGCGAGTGCCAGTCCCCCAGGTTCCCTCACGAATGTCCGGTCGCGGAACATATCCTGCACTGCTCCCGTCAACCGGTGAAGCGCCGCGTCGTAGC** 78634
WF-Teratorn CGGACACGACTCCCAGGGTTTCGTCCGCGAGTGCCAGTCCCCCAGGTTCCCTCACGAATGTCCGGTCGCGGAACATCTCCTGCACGGCTCCCGTCAACCGGTGAAGCGCCGCGTCGTAGC 81813

TeratornLC199500 **TGACGCAGAGTTCGGATGTCAGCTCGAATCCCAGCATCGCCTCCTGGCCCTCGGCGTAGGCCAGATTCTTGTGGAACCATTTGGCCAGGGCGTCGAATTGCCTATGGAACGCATGGGCTT** 78754
WF-Teratorn TGACGCAGAGTTCGGATGTCAGCTCGAATCCCAGCACCGCCTCTTGGCCCTCGGCGTAGGCCAGATTCTTGTGGAACCATTTGGCCAGGGCGTCGAATTGCCTATGGAACGCATGGGCTT 81933

TeratornLC199500 **CGTCAAGGGCTTCCTCCAAATAGCTGTTCCTCGTCGACGGGCGGTCTGCTGAGACGGCGGGATCGGTCCTCGATGTCGAGCAGGTCGCAGTAGAGCCCGAACGGACAAGTCCCGATATGA** 78874
WF-Teratorn CGTCAAGGGCTTCCTCCAAATAGCTGTTCCTCGTCGACGGGCGGTCTGCTGAGACGGCGGGATCGGTCCTCGATGTCGAGCAGGTCGCAGTAGAGCCCGAACGGACAAGTCCCGATATGA 82053

TeratornLC199500 **TACCCAAGAACTCCGCAAGACAGTCCTTTCCAATAGTCTGACGAGAGAGAAGCCCCCCGAGCTGCGCTCGCAGCTCGTCGGCCCTCCCGTTCGAGTAGTAGCTGCTCCACAACCTGAGG** 78994
WF-Teratorn TACCCAAGAACTCCGCAAGACTGTCCTTTCCAATAGTCTGACGAGAGAGAAGGCCCCCCGAGCTGCGCTCGGAGCTCGTCGGCCCTCCCGTTCGAGTAGTAGCTGCTCCACAACCTGAGG 82173

TeratornLC199500 **TAGGAGGGATCTTCTCCAACTGAGCGTCCGAGCACCTTCTCCCTGAGTTCGAGTCGAGCCCGTTTCTGCCAGATCCACCAGCCAGCGTCCGGCTCCTCCTCGGCCTCCGCGCGACGGTGA** 79114
WF-Teratorn TAGGAGGGATCGTCTCCAACTGAGCGTCCGAGCACCTTCTCCCTGAGTTCGAGTCGAGCCCGTTTCTGCCAGATCCACCAGCCAGCGTCCGGCTCCTCCTCGGCCTCCGCGTGACGGTGA 82293

TeratornLC199500 **AGTTCTTTGGCTTCGCTTCTCGGCTCATCCCCCCTGCTGCCTTCCAGCCCGCCGCCGTGCCTCCCGCCTTCCCTCAGCTCTTCCGCACTGACTCCGAATGTCCTACGGTGATTCAGGCAG** 79234
WF-Teratorn AGCTCTTTGGCTTCGCTTCTCGGCTCATCCCCCTTGCTGCCTTCCAGCCCGCCGCCGTGCCTCCCGCCTTCCCTCAGCTCTTCCGCACTGACTCCAAATGTCCTACGGTGATTCAGGCAG 82413

TeratornLC199500 **AGCCTCCCGCCTTGCGCCGGCACCGTTGCGCACAGGGTCGTGCAGATTTTTCCCAAGAGCAGGCCGTCGATCTGAGACACAGCCAGAGTCCTTAGGGCCGCTGGCTCGCAGGCCGAGGGG** 79354
WF-Teratorn AGCCTCCCGCCTTGCGCCGGCACCGTTGCGCACAGGGTCGTGCAGATTTTTCCCAAGAGCAGGCCGTCGATCTGAGACACAGCCAGAATCCTTAGGGCCGCTGGCTCGCAGGCCGAGGGG 82533

TeratornLC199500 **CAGACGCCGTTCGCGAAACCGCACCGTTGGCAGATTCGCCCCCAGGAAATCCCAGCGCTCCGCAGCTCTCGCGAGAGCAGGTCCAGTTCGCGAGACCAACGCGAAAGCGCTCCGCAGCAC** 79474
WF-Teratorn CAGACGCCGTTCGTGAAACCGCACCGTTGGCAGATTCGCCCCCAGGAAATCCCAGCGCTCCGCAGCTCTCGCGAGAGCAGGTCCAGTTCGCGAGACCAACGCGAAAGCGCTCCGCAGCAC 82653

TeratornLC199500 **AGTGCGGTCAACACCACGGCCCAGTACGGCGCGCGGACCTGGCGGTCCGCTGAAATGACGTCTGGGCCTTCGGGCTCGACCACCTCGATCCCCGAGCGAGTGAGATGCGCTAGCAGCCAC** 79594
WF-Teratorn AGTGCGGTCAACACCACGGCCCAGTACGGCGCGCGGACCTGGCGGTCCGCTGAAATGACGTCTGGGCCTTCGGGCTCGACCACCTCGATCCCCGAGCGAGTGAGATGCGCTAGCAGCCAC 82773

TeratornLC199500 **ACGCCGGAGCTCGGTGGTGAAGGAGGCCTCAGGGGACTGTTGACCACCCAGCGGGACAGGGCTCCGGGGTCTGCGTCACCCTCTCGTCGCCACAGCACACTGGTGCCCCAGACGGCTTGC** 79714
WF-Teratorn ACGCCGGAGCTCGGTGGTGAAGGAGGCCTCAGGGGACTGTTGACCACCCAGCGGGACAGGGCTCCGGGGTCTGCGTCACCCTCTCGTCGCCACAGCACACTGGTGCCCCAGACGGCTTGC 82893

TeratornLC199500 **GTGGGGTCGGGGAGCCAGCTCGAGTCTGACGCTCTGCCGTCAGTACGAAGCGCAGCGCGGGCCACGGTCAGCCGACTTTGTTCGTGCAAGATTTTAAACGCGAGCGGGGCTCCCAGCGTG** 79834
WF-Teratorn GTGGGGTCGGGGAGCCAGCTCGAGTCTGACGCTCTGCCGTCAGTACGAAGCGCAGCGCGGGCCACGGTCAGCCGACTTTGTTCGTGCAAGATTTTAAACGCGAGCGGGGCTCCCAGCGTG 83013

TeratornLC199500 **GCCAGCGTCGCGACCGTGACCTCCCGCGCAACTCTCTCCCTTTCAGACAGGTCCCCGTCGCGAGACGCCGCGACCGTGTAGTAGGAGAAGCGGTTCAAGCTGGTATATATCGAGGACACT** 79954
WF-Teratorn GCCAGCGTCGCGACCGTGACCTCTCGCGCAACTCTCTCCCTTTCAGACAGGTCCCCGTCGCGAGACGCCGCGACCGTGTAGTAGGAGAAGCGGTTCAAGCTGGTATATATCGAGGACACT 83133

TeratornLC199500 **AGCTTCTCAGCCTTATCCGTTCCATGTCCGCAGGCCAGACTCGCTCCAGGTTCTGCTGTGTCGTAGTGCTCGGAGTGGCGCAGCTCAACCCAGGCCAGGGGTGCGCTGGCCAAAACGGAG** 80074
WF-Teratorn AGCTTCTCAGCCTTATCCGTTCCATGTCCGCAGGCCAGACTCGCTCCAGGTTCTGCTGTGTCGTAGTGCTCGGAGTGGCGCAGCTCAACCCAGGCCAGGGGTGCGCTGGCCAAAACGGAG 83253

TeratornLC199500 **ATCCTTTTCACCGGGTAGCCCTTGGCATGTCCCTTCGCGGCTCTGAAATGCCTGTCCGTCACCAGACCATAACTCTCACTCCCCTGCGAGTACACCACCCTCAGGACCATGCAGCGCGCG** 80194
WF-Teratorn ATCCTTTTCACCGGGTAGCCCTTGGCATGTCGCTTCGCGGCTCTGAAATGCCTGTCCGTCACCAGACCATAACTCTCACTCCCCTGCGAGTACACCACCCTCAGGACCATGCAGCGCGCG 83373

TeratornLC199500 **CCTCGCAGAGCGCCGGGCCCGACTGGGATGAAGACGGGACCTCCTAGTTTCGGGCCGGTGAGCGTCACGCTCGAGGGCAACCTTTTCTCTATTTCGTAAAGCTGCACCCACGTGTACTCC** 80314
WF-Teratorn CCTCGCAGAGCGCCGGGCCCGACTGGGATGAAGACGGGACCTCCTAGTTTCGGGCCGGTGAGCGTCACGCTCGAAGGCAACCTTTTCTCTATTTCGTAAAGCTGCACCCACGTGTACTCC 83493

TeratornLC199500 **AGAAGGACCGGAGCATCGGACCCCTCCAGCTCCACAACCAGTTGAGGCTCGGAAGGCCCCGTCGCGCACATGCCGTCTAAACCCGGATCAACTTGAGAGCTTAAGTTCAATTTAAACACC** 80434
WF-Teratorn AGAAGGACCGGAGCATCGGACCCCTCCAGCTCCACAACCAGTTGAGGCTCGGAAGGCCCCGTCGCGCACATGCCGTCTAAACCCGGATCAACTTGAGAGCTTAAGTTCAATTTAAACACC 83613

TeratornLC199500 **ACACCAACCTTTTTTT----TTTGTTCATTTATTTGTCTCAT**ACCGTCAAAAACATGTGCCAAAGGAAAAAAAAA--AGTTACACAAGCAGCACACACGAACATAATGGCACCGCCACAC 80548
WF-Teratorn ACACCAACCTTTTTTTTTTTTTTGTTCATTTATTTGTCTCATACCATCAAAAACATGTGCCAAAGGAAAAAAAAAAAAGTTACACGAGCAGCACACACGAACATAATGGCACCGCCACAC 83733

TeratornLC199500 CGTTATAGCAGGGTGGCCTCCCGATGTCCTTTCCCCCC---------CCCCCCTGAAATCCTCCCTCGTATGCGGAGTGCCGAGCTCACGTTCCTCTCCTCTGCGGAGTGCCGAGCTCAC 80659
WF-Teratorn CGTTATAGCAGGGTGGCCTCCCGATGTCCTTCCCCCCCCCCCCCCCCCCCCCCCTGAAATCCTCCCTCGTATGCGGAGTGCCGAGCTCACGTTCCTCTCCTCTGCGGAGTGCCGAGCTCAC 83853

TeratornLC199500 GTTCCTCTCCTCGGACTCGGCTGTCTCGCACGACCCCCGAAATTCGGGCTGTTCTTTGTGTGGGAGCCATTCTTCTGTAGATCAGTGGTCTACGACGCCGAGTTCTTTGGCTTCTCCGCA 80779
WF-Teratorn GTTCCTCTCCTCGGACTCGGCTGTCTCGCACGACCCCCGAAATTCAGGCTGTTCTTTGTGTGGGAGCCATTCTTCTGTAGATCAGTGGTCTACGACGCCGAGTTCTTTGGCT-------- 83965

TeratornLC199500 CCATTATTGCATCGCCTAGGATCTCCGATCTTCTCCTAAGCGGTCTGTGGCGTGCCGCGGATCGAGGAACACTCTGAGCCCTCCTGCCGCGGACTCCTCTTCTGCGTTCCCCTTCGTCGG 80899
WF-Teratorn ------------------------------------------------------------------------------------------------------------------------ 83965

TeratornLC199500 TCGTATCGCCCGGAGAGGGGTCCTCTTCCATGACCGCTAAACTCTTGCGAGGGTCCACTATTTCCTCGATCACGCTTTCGTTCCTTCCTCTCCGAGATGGGGCGAGGCAGGTCGCGATC 81019
WF-Teratorn ------------------------------------------------------------------------------------------------------------------------ 83965

TeratornLC199500 ATTCTGTCGTGATACCTCATGACGTACGGCTCGTAGTAGCCACGAACCCCGAACTCCACAAACCGCTGTCCCCATCCGCCTTCGGGCGCTGAAACGGCTCGCTCTATCCGGAGGGTTCGG 81139
WF-Teratorn ---------------------------------------------------------------------------------------------------CGCTCTATCCGGAGGGTTCGG 83986

TeratornLC199500 AGAGGAGGTTGCGAAGACATCTCAGATGCCGATTCGTTTCCATGAACAGAACACGCAGCGTGTGTAGAGCGTGTCTAGGTCCACTAGTGAATGGTCTATTTGCAAGCGGGGCAGACGGTG 81259
WF-Teratorn AGAGGAGGTTGCGAAGACATCTCAGATGCCGATTCGTTTCCATGAACAGAACACGCAGCGTGTGTAGAGCGTGTCTAGGTCCACTAGTGAATGGTCTATTTGCAAGCGGGGCAGACGGTG 84106

TeratornLC199500 TACGGTTTCGGGTCCGGGAAGGTCAGCGTACTAAGTGGCACGTATTTTACCTAAAAAGGGGCCTAACGCGACGTGGTGACCTCACTCTGTGCGAGGAAACCTGCCACTTTGAATGGCTGA 81379
WF-Teratorn TACGGTTTCGGGTCCGGGAAGGTCAGCGTACTAAGTGGCACGTATTTTACCTAAAAAGGGGCCTAACGCGGCGTGGTGACCTCACTCTGTGCGAGGAAACCTGCCACTTTGAATGGCTGA 84226

TeratornLC199500 TATTATGAGAGCCCCTGGAGCATACTCGAAACGCACTGAACTATACTGAAGTGAACTACACACAGACTGTAATGTTGTGTCTTTTATTTGACTCGAACCATCAAAAGCGTGTTCCAAAAC 81499
WF-Teratorn GTTTATGAGAGCCCCTGGAGCATACTCGAAACGCACTGAACTATACTGAAGTGAACTACACACAGACTGTAATGTTGTGTCTTTTATTTGACTCGAACCATCAAAAGCGTGTTCCAAAAC 84346

TeratornLC199500 CAACCAAAAATTACACAAGCAGCACACAGAAACACAATGACACCAACGCGGCGTCGTGATATCGCAGGGCGTCAGCCTGACGTCTTTCTCCCCCTGAGATCGGGAGTCCCGAGCCCAGGC 81619
WF-Teratorn CAACCAAAAATTACACAAGCAGCACACAGAAACACAATGAAACCAACGCGGCGTCGTGATATCGCAGGGCGTCAGCCTGACGTCTTTCTCCCCCTGAGATCGGGAGTCCCGAGCCCAGGC 84466

TeratornLC199500 TC**TCATGCCTCACACCCTCCCGGGCGGTCCTCGTCCCAACGCTCCCCTCCGATTTCCGCGTAGAAGCTGAGAGTGGTGTTGCTCGCCACGCCGTCCAAGTACTGCGAATCGTGCAGAGGA** 81739
WF-Teratorn TCTCATGCCTCACACCCTCCCGGGCGGTCCTCGTCCCAACGCTCCCCTCCGATTTCCGCGTAGAAGCTGAGAGTGGTGTTGCTCGCCACGCCGTCCAAGTACTGCGAATCGTGCAGAGGA 84586

TeratornLC199500 **TCGACAAGAGCAACGGGCGCTCCTGCCGAGGACGCCTCCTCCGGGATCTTGCCTTCGTTCGGAGCTGGGTCCTCTTCGACGACCGCCAAGCTCTTGTGAGCTCGGACTATCTCCTCGATC** 81859
WF-Teratorn TCGACGAGAGCAACGGGCGCTCCTGCCGAGGACGCCTCCTCCGGGATCTTGCCTTCGTTCGGTGCTGGGTCCTCTTCGACGACCGCCAAGCTCTTGTGAGCTCGGACTATCTCCTCGATC 84706

TeratornLC199500 **ACGGGCTCCCTCCTTCCCAGCACATACACGGTGTCCAGGTACGCGGACAGGTTGTCGTACTTGCCCGTCTTCAGAGCGGGCCCTCCGCGAGGTCTTCGAGAAGAGGCGAAGCGGGCCTCG** 81979
WF-Teratorn ACGCGCTCCCTCCTTCCCAGCACATACACGGTGTCCAGGTACGCGGACAGGTTGTCGTACTTGCCCGTCTTCAGAGCGGGCCCTCCGCGAGGTCTTTGAGAAGAGGCGAAGCGGGCCTCG 84826

TeratornLC199500 **CTCCCGTAGTGTCTGTCCATTATGTCCCGAAACGTCGGGTCGGGCAGCTCGCAGACGCTTCGAAACACTGACGCCTCGGTCCCCGGTTCCCGGTCCAT**CTGGAGGGTCAAGGAGGGAGGG 82099
WF-Teratorn CTCCCGTAGTGTCTGTCCATTATGTCCCGAAACGTCGGGTCGGGCAGCTCGCAGACGCTTCGAAACACTGACGCCTCGGTCCCCGGTTCCCGGTCCATCTGGAGGGTCAAGGAGGGAGGG 84946

TeratornLC199500 GCAGGAGGACACGACGGGATC**TCAGGTGCCGATTCGGTTTGATGCGCAAAACACGCAGCGCGTGTGGAGCGTTTCCAAATCCACTAGTACTTCTGAGAAAATGGTCTGTTTGCAAGCGGG** 82219
WF-Teratorn ACAGGAGGACTCGAGGGGATCTCAGGTGCCGATTCGGTTTGATGCGCAAAACACGCAGCGCGTGTGGAGCGTTTCCAAGTCCACTAGTACTTCGGAGAAAATGGTTTGTTTGCAAGCGGG 85066

TeratornLC199500 **GCACACGGTGTACATCTGCTTCAGGTCGGTGAAGGTGCACGTGTTCACGGTCGACGTGCTGAGCGGCACGTTCGCCACCGGGGCCGCGCAGCAGGACGAGACGAATTTCATTTGTTCGAA** 82339
WF-Teratorn GCACACGGTGTACATCTGCTTCAGGTCGGTGAAGGTGCACGTGTTCACGGTCGACGTGCTGAGCGGCACGTTCGCCACCGGGGCCGCGCAGCAGGACGAGACGAATTTCATTTGTTCGAA 85186

TeratornLC199500 **ATCGAACGCCACGTCGATGTTCCTGGGGTTGCCTTTCACTTCGGACGATCGGAACGTCTGGTTCTTCAGACACCTGACGCACCTCAGGTCTCCCTCCGCGGCCCGAGCCCGGTCCCGACC** 82459
WF-Teratorn ATCGAACGCCACGTCGATGTTCCTGGGGTTGCCTTTCACTTCGGACGATCGGAACGTCTGGTTCTTCAGACACCTGACGCACCTCAGGTCTCCCTCCGCGGCCCGAGCCCGGTCCCGGCC 85306

TeratornLC199500 **CGGAAGCCCCACGGGTGCCAGCAGGATCTGTTTGTGGTTGTAGTCTATCCAGTTTATGTGACAGCTCAACACGGCGTACAGCAGCATCCGTCGTTCGGGGCTCTGGCTTCGCAGCTGCCC** 82579
WF-Teratorn CGGAAGCCCCACGGGTGCCAGCAGTATCTGTTTGTGGTTGTAGTCTATCCAGTTTATGTGACAGCTCAACACGGCGTACAGCAGCATCCGTCGTTCGGGGCTCTGGCTTCGCAGCTGCCC 85426

TeratornLC199500 **GAACTGCTCCCTGCTGATCATGTCCGCTGCTTCCCTGAGCTGCGGGATGCACGCCGCGGCCATCTCGAGCGGCCGGTCCGTCGTGTAAGCGGCGGCCAGCCTGTAATAGCAGCTCAAGGG** 82699
WF-Teratorn GAACTGCTCCCTGCTGACCATGTCCGCTGCTTCCCTGAGCTGCGGGATGCACGCCGCGGCCATCTCGAGCGGCCGGTCCGTCGTGTAAGCGGCGGCCAGCCTGTAATAGCAGCTCAAGGG 85546

TeratornLC199500 **CCACTCCGAATTTAGCGCGAGCATCGAGTGGCCCGAGGCCTTCACGCACTGAAATGCTCTGACGAGTTTGCAGTGGTACGACCGTCTCCTGGGGGCAAAGCAGAAGCGAGACCACTTCAC** 82819
WF-Teratorn CCACTCCGAATTTAGCGCGAGCATCGAGTGGCCCGAGGCCTTCACGCACTGAAATGCTCTGACGAGTTTGCAGTGGTACGACCGTCTCCTGGGGGCAAAGCAGAAGCGAGACCACTTCAC 85666

TeratornLC199500 **CGGGTCTTCGCTCGACCCGTCGCCTTCCGGGCGGCTGCTCAACGGAGCCGAGGTCGTCCGGAGGGTGGCGAAAGGAGACGCTGTGCGCAGGACACGTGTCACCCGTCTCCCGCGTTCGTT** 82939
WF-Teratorn GGGGTCTCCGCTCGACCCGTCGCCTTCCGGGCGGCTGCTCAATGGAGCAGAGGTCGTCCGGAGGGTGGCGAAAGGAGACACTGAGCGCAGTACACGTGTCACCCGTCTCCCGCATTCGTT 85786

TeratornLC199500 **CGCTCCGAGACCCGCTCGGCTCCGCCGCGACTTGGCCTGGGGGCCCGCTTCGCCGTCGAGAGGCGCTCCCGATTTCTCCCGAGAGGGAGGTCGGACGACGGAGCTGCCCTCTGCCCGCTG** 83059
WF-Teratorn CGCTCCGAGAGCCGCTCGGCTCCGCCGCGACATGGCCTGGGGGCCCGCTCCGCCGTCGAGAGGCGCTCCCGATTTCTCCCGAGAGGGAGGTCGGACGACGGAGCTGCCCTCTGCCCGCTG 85906

TeratornLC199500 **GTCCGAGATGAGCCTAAAAGCCTCCAGTCTGAGGGCCAGGGCCTTGTGCGAAGAGTGCGACTCGTCAAAGAACAGATCTTCTGCTTGGAACCTGTAGAAGTCAACAAGCAGAGCCACCCT** 83179
WF-Teratorn GTCCGAGATGAGCCTAAAAGCCTCCAGTCTGAGGGCCAGGGCCTTGTGCGAAGAGTGCGACTCGTCAAAAAACAGATCTTCTGCTTGGAACCTGTAGAAGTCAGCAAGCAGGGCCACCCT 86026

TeratornLC199500 **GAGCATTGCCTCGCGCCCCCTCAGCTTGGACAGCTCGTCGCAGTCGGGCACAGTCCCCTCTCCAAACAGACCGTCTGAGCCGACCAGCGCGGGGCGCAGAGTCGAATCCCAAAGGTTAGA** 83299
WF-Teratorn GAGCATTGCCTCGCGCCCCCTCAGCTTGGACAGCTCGTCGCGGTCGGGCACAGTCCCCTCTCCAAACAGACCGTCTGAGCCGACCAGCACGGGGCGCAGAGTCGAATCCCAAAGGTTAGA 86146

TeratornLC199500 **GCACAGCGAGTAGGCTTCGCCGTAATCCTCCACCGGAGCTTTGGCCTTGCTCGTCAGCAGCCTGACCAGCTGAGCTCGAACGCTCGAACTGTCCCGAGAACCCTGTCTAATTTTGTTATG** 83419
WF-Teratorn GCACAGCGAGTAGGCTTCGCCGTAATCCTCGACCGGAGCTTTGGCCTTGCTCGTCAGCAGCCTGACCAGCTGAGCTCGAACGCTCGAACTGTCCCGAGAACCCTGTCTAATTTTGTTATG 86266

TeratornLC199500 **GCACACGGCGTGGTTGTGGAGGCACTCAAGGTAAGCCTGGACGAGGGCTGCTTTTGCGTGGCAACCGGACCGCACGACTCCGCCGTCGCTTACCCGTGCCCCGTGGAAGCTTTCGCACCC** 83539
WF-Teratorn GCACACGGCGTGGTTGTGGAGGCACTCAAGGTAAGCCTGGACGAGGGCTGCTTTTGCGTGGCAACCGGACCGCACGACTCCGCCGTCGCTTACCCGTGCCCCGTGGAAGCTTTCGCGCCC 86386

TeratornLC199500 **GTGCTCCTCGGATCGCCCGTCACGCTGCATGTCTACTGCCCGGCCCTGTTCCCCGACAGGGTGTTGAGAGTGCACGCCGAACACCTGGGTCACTTCACCGGGATGAGGAACGTGCCTCGC** 83659
WF-Teratorn GTGCTCCTCGGATCGCCCGTCACGCTGCATGTCTACTGCCCGGCCCTGTTCCCCGACAGGGTGTTGAGAGTGCACGCCGAACACCTGGGTCACTTCACCGGGATGAGGAACGTGCCTCGC 86506

TeratornLC199500 **GTGGACCCCGAGCAGGAACGAGCGCTCTTCTCAGGTCTGTGTGGACTGTTCGGCACAGAGCGAGTCCTCGGCCTTTACCGGGAGGCCCTGAGCGTCGGAGACGGCGGCGAGGGAGCCCCG** 83779
WF-Teratorn GTGGACCCCCAGCAGGAACGAGCGCTCTTCTCAGGTCTGTGTGGACTGTTCGGCACAGAGCGAGTCCTCGGCCTTTACCGGGAGGCCCTGAACGTCGGAGACGGCGGCGAGGGAGCCCCG 86626

TeratornLC199500 **AGCCAGGATGGCGGAGCAATCGAAAAGCCGCCGATGAGCGACGGGGAGCTCTGCCCCTCCGTGTTCGTCAAGTTTCGCAGAGAAAACACGCGGGAGGCCAGGCGGATCGCATCTCTGGCT** 83899
WF-Teratorn AGCCAGGACGGCGGAGCAATCGAAAAGCCGCCGATGAGCGACGGGGAGCTCTGCCCCTCCGTGTTCGTCAAGTTTCGCAGAGAAAACACGCGGGAGGCCAGGCGGATCGCATCTCTGGCT 86746

TeratornLC199500 **AGCCAACTCACGGGGTTGTTCGACGTGGAGGATCTCAGAGAGAAAGCGATCCCCCACGTCAAGGGAGAGGAGCCGGCTCACATGTTGTTTCTGTTTCTCGTCCAGCTTATGGCTGTGTTC** 84019
WF-Teratorn AGCCAACTGACGGGGTTGTTCGACGTGGAGGATCTCGGAGAGAAAGCGATCCCCCACGTCAAGGGAGAGGAGCCGGCTCACATGTTGTTTCTGTTTCTCGTCCAGCTTACGGCCGTGTTC 86866

TeratornLC199500 **CTCAGCTCCTGCCTGGACGAAGCTAGCCTCGCCACGCTCATGAGCGTGGCTCTCTCCCCGTTCCTGGAGGGCAACTTCTGCGACCCCCTGTGCGCAGCCACCCTGCTGAAGCCCATGTTC** 84139
WF-Teratorn CTCAGCTCCTGCCTGGACGAAGCTAGCCTCGCCACGCTCATGAGCGTGGCTCTCTCCCCGTTCCTGGAGGGCAACTTCTGCGACCCCCTGTGCGCCGCCACCCTGCTGAAGCCCATGTTC 86986

TeratornLC199500 **ATCCCGTGGTTCGAAGCGTTCCCCAGGAAGGCGGACGGCCCGGAAGCAAAAGCAGACCCCGAAGCCAAGCTGAAGTACTACAGGCTGTACAAGGGCTACGAGATCACGAACGAGGGCGCG** 84259
WF-Teratorn ATCCCGTGGTTCGAAGCGTTCCCCAGGAAGGCGGACGGCCCGGAGGCGAAAGCAGACCCCGAAGCCAAGCTGAAGTACTACAGGCTGTACAAGGGCTACGAGATCACGAACGATGGCGCG 87106

TeratornLC199500 **GACGTCTCTTCCGTGCTGGAGGACGACCTCACGTGGACCAGCGCCAGCTTGCAGCTGCGCACTCTGGGCTACAAGATCGAGGAGAGCCTTAGCAAAACGCTCTGGTTCGTGGCCCCCCAA** 84379
WF-Teratorn GACGTCTCTTCCGTGCTGGAGGACGACCTCACGTGGACCAGCGCCAGCTTGCAGCTGCGCACTCTGGGCTACAAGATCGAGGAGAGCCTTAGCAAAACGCTCTGGTTCGTGGCCCCCCAG 87226

TeratornLC199500 **TTCGGAGACAGGCTGCGCATGACTCGCGACTCTTGCATCGGGAAAATGGCTGTGTCCCCGGGCGAGCACGTCGACCTCATCCGCTCCTGCCTGCTCTGCGACAACGTGATGGCCTACGTG** 84499
WF-Teratorn TTCGGAGACAGGCTGCGCATGACTCGCGACTCTTGCATCGGGAAAATGGCCGTGTCCCCGGGCGAGCACGTCAACCTCATCCGCTCCTGCCTGCTCTGCGACAACGTGATGGCCTACGTG 87346

TeratornLC199500 **CAGTGCGCGCTCTTCGGCACGATCCGCAACATGCTGCCCTGCCACGCCAACAGCGAGTTCTGTGAGACCGTGATGGCGGCCCTGAACGGCTCCATGAGGAGGTTTGCGGCCGAGGCGCGT** 84619
WF-Teratorn CAGTGCGCGCTCTTCGGCACGATCCGCAACATGCTGCCCTGCCACGCCAACAGCGAGTTCTGTGAGACCGTGATGGCGGCCCTGAACGGCTCCATGAGGAGGTTTGCGGCCGAGGCGCGT 87466

TeratornLC199500 **CACCAGAGCAGAGAGCTGTTCTCTCGCCTGGAAGGGTGGAGCGGCCCCTGCGCCGGCAACTGCCTGTACACGTCCGGGGTCAGTTTCGCTGGTCAGTCCGAGGGCAGGAGCCTGTTCTTC** 84739
WF-Teratorn CACCAGAGCAGAGAGCTGTTCTCTCGCCTGGAAGGGTGGAGCGGCCCCTGCGCCGGCAACTGCCTGTACACGTCCGGCGTCAGTTTCGCTGGTCAGTCCGAGGGCAGGAGCCTGTTCTTC 87586

TeratornLC199500 **GGCAAGATCTTCACCGGCCCCGAGGACAGGAAGCCTCCGGGCAAGTACAAGACGGCATTTGGAGACTTTTCCAGCGTCACACTGGGCCTTTCGCTCAAGACGGGCTTGCTGGAGGCGCTC** 84859
WF-Teratorn GGCAAGATCTTCACCGGCCCCGAGGACAGGAAGCCTCCGGGCAAGTACAAGACGGCATTTGGAGACTTTTCCAGCGTCACTCTGGGCCTTTCGCTCAAGACGGGCTTGCTGGAGGCGCTC 87706

TeratornLC199500 **AGGGTCGCGCTGGAACGGGACGAGGCGCCCAAGGCGGGCTGCGGATTGAGGCGCGAAGCCGGCGGCCGCATGCAAAAGATGCTCTCCACGGCCCAGGAACCCGTGTGCGCGGAGAGGGTG** 84979
WF-Teratorn AGGGTCGCGCTGGAACGGGACGAGGCGCCCAAGGCGGGCTGCGGATTGAGGCGCGAAGCCGGCGGCCGCATGCAAAAGATGCTCTCCACGGCCCAGGAACCCGTGTGCGCGGAGAGGGTG 87826

TeratornLC199500 **GTGACCGAAGTCGCTTTCTTCGCTCGCTATCACCACAGGTTCAACTCCGGAGAGTTCACCGAGGTGGGTTGCGTGAAAGCCATGAGCGAGCCCGGCTTCAACGAGCAGATACACAAGGAC** 85099
WF-Teratorn GTGACCGAGGTCGCTTTCTTCGCTCGCTATCACCACAGGTTCAACTCCGGAGAGTTCACCGAGGTGGGTTGCGTGAAAGCCATGAGCGAGCCCGGCTTCAACGAGCAGATACACAAGGAC 87946

TeratornLC199500 **GTGCTGGACGACAAGAGCGAGTACCGCCTGTCCCTCGCCCACGGCGCACTGCCGATGTGCGCGCTCACGCTTTCCATGACCATGAGACAGACCGTCATGATGACTTTGGGGTTCTTCTGC** 85219
WF-Teratorn GTGCTGGACGACAAGAGCGAGTACCGCCTGTCCCTCGCCCACGGCGCACTGCCGATGTGCGCGCTCACGCTTTCCATGACCATGAGACAGGCCGTCATGATGACTTTGGGGTTCTTCTGC 88066

TeratornLC199500 **GTGGAGCTCGCACACGGAGCTTACGCCGGGGCCAAACTCAACTACAGTACGCTGTGCTCCCGCTTGGTCTCCCTCTCCACCGTGGCCAAATTCAGGCACGGGAAAGACAGCGACAACACG** 85339
WF-Teratorn GTGGAGCTCGCGCACGGAGCTTACGCCGGGGCCAAACTCAACTACAGTACGCTGTGCTCCCGCTTGGTCTCCCTCTCCACCGTGGCCAAATTCAGGCACGGGAAAGACAGCGACAACACG 88186

TeratornLC199500 **AACGTCTTCCAGGCGTTGTACGGCAACTGCCTGAAGCTGACGTGCGCCATGTTCAGCAAACCGAATCCGCACGCACCGATAAACTATCACGATCTAGATCTAGTCGAGTGGCTGCACCCC** 85459
WF-Teratorn AACGTCTTCCAGGCGTTGTACGGCAACTGCCTGAAGCTGACGTGCGCCATGTTCAGCAAACCGAATCCGCACGCACCGATAAACTATCACGATCTAGATCTAGTCGAGTGGCTGCACCCC 88306

TeratornLC199500 **GACAACCCCCGCCTCCCCTGCTACCTGCGCAAATCGCTGGCCATGATAGGGGGCACCAAGGCTCGAGTTAGAGACATAATGCGGCAGCACGGACTCTGCTACAACAGAGTTATGTTCAGC** 85579
WF-Teratorn GACAACCCTCGCCTCCCCTGCTACCTGCGCAAATCGCTGGCCATGATAGGAGGCACCAAGGCTCGAGTTAGAGACATAATGCGGCAGCACGGACTCTGCTACAACAGAGTTATGTTCAGC 88426

TeratornLC199500 **GATGGCCAGCGCCGGTTGCTGTTCTCCCACGCTTTCGCCAACGCGGACTGCCTGATCCACGCACCGGTTACGATCTGCTCCAGCCTCAGGCTGTTTTCGAACACCCAGAAGTCGTTCGGG** 85699
WF-Teratorn GATGGCCAGCGCCGGTTGCTGTTCTCCCACGCTTTCGCCAACGCGGACTGCCTGATCCACGCACCGGTTACGATCTGCTCCAGCCTCAGGCTGTTTTCGAACACCCAGAAGTCGTTCGGG 88546

TeratornLC199500 **TCTCTCAAAGCCTGCAGGTGGGGCTGCTACTACCGAGTGACGATCATGAGCAACGGCTGCGTGAACACGGCCCTGTATACGAACCGCGGATACGGGGGCTTTGACAGCATCGCCTCTCAA** 85819
WF-Teratorn TCGCTCAAGGCCTGCAGGTGGGGCTGCTACTACCGAGTGACGATCATGAGCAACGGCTGCGTGAACACGGCCCTGTATACGAACCGCGGATACGGGGGCTTTGACGGCATCGCCTCTCAA 88666

TeratornLC199500 **GGCAGCGCCACCGGGGACGAGGTGTTGGAGAGGATCAAGCAAGAGGTGTCTCGCGTGTTGCCCAGGATCGACGTGGGCCGTATCAGGATGATGGCCCCGGAGATGAAGGAGACAGAGTTT** 85939
WF-Teratorn GGCAGCGCCACCGGGGACGATGTGTTGGAGAGGATCAAGCGAGAGGTGTCTCGCGTGTTGCCCAGGATCGACGTGGGCCGTATCAGGATGATGGCCCCGGAGATGAAGGAGACGGAGTTT 88786

TeratornLC199500 **GCAGCCCTGGTGTCCGCGGTGAAAGACAGCAACATCGAGGTCACCGGGCAAATGGACGGCGACGCTGAGCTTGAGCCGTGGCAGGAGGATGGCGCGGGGGAAGAAAATGCGAAACGCCCC** 86059
WF-Teratorn GCAGCCCTGGTGTCCGCGGTGAAAGACAGCAACATCGAGGTCACCGGACAAATGGACGGCGACGCTGAGCTTGAGCCGTGGCAGGAGGATGGCGCGGGGGAAGAAAATGCGAAACGCCCC 88906

TeratornLC199500 **TTGGACCACTCTG---ACGCCGACGACGCCAAGAGAGGTCGCTGGGATGCCTGTGAGACTGGTCGGGTTTCCGAGGACGAAGAATTTTTCTTTTTGTAG**TGAGTCGATGGTGGGGTGTGT 86176
WF-Teratorn TTGGACCACTCTGTCGGCGCCGACGACGCCAAGAGAGGTCGCTGGGATGCCTGTGAGACTGGTCGGGTGTCCGAGGACGAAGAATTTTTCTTTTTGGAGTGAGTCGATGGTGGGGTGTGT 89026

TeratornLC199500 GGGCTGTGATAGATGGAAAGGAAGAAAAAAAAAAAAC------TTTGCACTTGGAATGTGGTTGAAAGCAAACATGTGCGAATAAAAATGTTGTGAGAGAACAAACACAGAGTCGTCTGG 86290
WF-Teratorn GGGCTGTGATAGATGGAAAGGAAGAAAAAAAAAAAACAACAACTTTGCACTTGGAATGTGGTTGAAAGCAAACATGTGCGAATAAAAATGTTGTGAGAGAACAAACACAGAGTCGTCTGG 89146

TeratornLC199500 TCATTTTCTTGTCTCCATCCTTCACAAACGTACGAAATGAAACACGTTTGGCCGCGACGGCTCTACCTCGGCCGCGACACCGAGCTCAAGTGACGACAAGAGGCCCAATGCTGAGGAGAG 86410
WF-Teratorn TCATTTTCTTGTCTCCAGCCTTCACAAACTTACGAAATGAAACGCGTTTGGCCGCGACGGCTCTACCTCAGCCGCGACACCGAGCTCAAGTGACAACCAGAGGCCCAGTGCTGAGGAGAG 89266

TeratornLC199500 GACCGACGGCGAGAGGTGTCGCTCGGGCTGCCTCAGAGTACAGAGACTTTTTTTTTTTCATTTAGAGAGTGACACAGAGTGTGTACAAACCGACGGCGAGCGGCCGGGGAGGAGAGAGC 86530
WF-Teratorn GGCCGACGGCGAGAGGTGTCGCTCGGGCTGCCTCAGAGTACAGAGACTTTTTTTTTT-CATTTAGAGAGTGACACAGAGTGTGTACAAACCGACGGCGAGCGGCCGGGGAGGAGAGAGC 89385

TeratornLC199500 AGGAGAAAAGGCTGGGGCGTCAAGAAAGCATGCCGCAGACCGCTCCGTCGTCCGTTTAGATGATTT----------------------GTGTGCGGTTGGAAAGTGATTGAAAACAAGCC 86628
WF-Teratorn AGGGGAAGAGGCTGGGGCGTCGCGAAAGCATGCCGCAGACCGCTCCGTCGTCCGTTTAGATGATTTTGCGAAAGAGGAGACAACACTTGTGTGCGGTTGGAAAGTGATTGAAAGCAAGCC 89505

TeratornLC199500 TACGA--AAAGAATAAAGTGACATGACAAACACAGCCGTTTTCATTGTTTTATTGTTTCGCTGCCTCGAGTCT**TCACGAATGTAAGAAATGAAGTATGTTCAGGTGTAACAGCTCAACCG** 86746
WF-Teratorn TACGAATAAAGAATAAAGTGACATGACAAACACAGCCGTTTTCATTGTTTTATTGTTTCGCTGCCTCGAGTCTTCACGAGTGTAAGAAATGAAGTATGTTCAGGTGTAACAGCTCCACCG 89625

TeratornLC199500 **TGAGCATCAGCTGCCGGTAGGCGCAGCCCAGAGCGTGCAGATCCCGTCGCACGCAGAAAGCGAATCGGGACTCCATGCTCCCGATGGTGTTTCCCAAGCGGCCCTCCTGCGTCGTCCCGA** 86866
WF-Teratorn TGAGCATCAGCTGCCGGTAGGCGCAGCCCAGAGCGTGCAGATCCAGTCGCACGCAGAAAGCGAATCGGGACTCCATGCTCCCGATGGTGTTTCCCAAGCGGCCCTCCTGCGTCGTCCCGA 89745

TeratornLC199500 **CAGGCCTTGTTTTGCGCCGGGCCTCCAGTCTGCGCAGGGATTCCTCGGCCCTGCTGTAATCGAGCTCCCCGCAGCGGGTCAGCTGCAGCTCTCCGGGCTGCACGCGCGCGCTCACCAGAG** 86986
WF-Teratorn CGGGCCTCGTTTTGCGACGGGCCTCCAGTCTGCGCAGGGATTCCTCGGCCCTGCTGTAATCGAGCTCCCCGCAGCGGGTCAGCTGCAGCTCTCCGGGCTGCACGCGCGCGCTCACCAGAG 89865

TeratornLC199500 **GGCTGAGGGCCCCGCAGACCGACACGAGCATTTCCGAGTTTGCCGTCCGCTCGCGGGCCACGGGACACGCGAGGTTGGTGAACTTGCACCTAGGCGTTTCTTCCCTGCCGTGGTGTCGAT** 87106
WF-Teratorn GGCTGAGGGCCCCGCAGACCGACACGAGCATTTCCGAGTTTGCCGTCCGCTCGCGGGCCACGGGACACGCGAGGTTGGTGAACTTGCACCTAGGCGTTTCTTCCCTGCCGTGGTGTCGAT 89985

TeratornLC199500 **CGCGCGGGTCGCGGGGCTCTTCGAAGGAGCACAGCCCGTACATGTTCATGAACTGGTACGAGGTCGAGGCCGTGCTTTGCCTTTCCACGTGCTCGCGCACCATGTCTTTTATGTCAGAGA** 87226
WF-Teratorn CGCGCTGGTCGCGGGGCTCTTCGAAGGAGCACAGCCCGTACATGTTCATGAACTGGTACGAGGTCGAGGCCGTGCTTTGCCTTTCCACGTGCTCGCGCACCATGTCTTTTATGTCAGAGA 90105

TeratornLC199500 **GGGCTTTGGCCCTCAGTTCGCCGTACAGCCTCCTGTATTCCACGTCGATAGGACTCTCTCCTCGGTCGTCGTCCTCCTCCCTGATTCTCTTGGGCACAGGAGGCTCATCGTTCGGCTCGC** 87346
WF-Teratorn GGGCTTTGGCCCTCAGTTCGCCGTACAGCCTCCTGTATTCCGCGTCGATAGGACTCTCTCCTTGGTCGTCGTCCTCCTCCCTGATTCTCTTGGGCGCAGGAGGCTCATCGTTCGGCTCGC 90225

TeratornLC199500 **CTTCGCCCTCCCAGTCCGGGAGCGAGCCGTAGCATGCTTTGACGGCGCTCGCCACGAAAAAAAGTTCGGGGATGTCCGCGGGACGCGAACCTCCCGTTTGCGTGGACATAACCGACTGCG** 87466
WF-Teratorn CTTCGCCCTCCCAGTTCGGGAGCGAGCCGTAGCATGCTTTGACGGCGCTCGCCACGAAAAAAAGTTCGGGGATGTCCGCGGGACGCGAACCTCCCGTTTGCGTGGACATAACCGACTGCG 90345

TeratornLC199500 **CCCACTTGATCCGATACCTGGTCGACGAACTCGGAAGCGCCTCGCTCGCCGACCTGCCTTTAGTAGAGTGCCTGGTCAAGCATCTGGAGCTGGTCTCCGGCCCTGACGTCTGGTCTCCTA** 87586
WF-Teratorn CCCACTTGATCCGATACCTGGTCGACGAACTCGGAAGCGCCTCGCTCGCCGACGTGCCTTTAGTAGAGTGCCTGGTCAAGCATCTGGAGCTGGTCTCCGGCCCTGACGTCTGGTCTCCTA 90465

TeratornLC199500 **GGGCACGCGCCTACGTCGGCTACGACCCTCAGGTCCACGGCTCTTTCGGGCGGTTGAGGCCCGGACCGGCCGGCCGCGTGCGGTTCCTGGGGAAGCTGCTGCCCGTTTACTGCAGGCTGT** 87706
WF-Teratorn GGGCACGCGCCTACGTCGGCTACGACCCTCAGGTCCACGGCTCTTTCGGGCGGTTGAGGCCCGGACCGGCCGGCCGCGTGCGGTTCCTGCGGAAGCTGCTGCCCGTTTACTGCAGGCTGT 90585

TeratornLC199500 **GCTTCCTCTGCGACTGTCACGGCAGGGCTCTGGATGCCGCGAGGACGCTGCGGTCTAGACTGACGGCGGCCGCCGAGTCCGGGGGATCGGCGGCGCCCCCGAGCCTCGCGTACGAGACCA** 87826
WF-Teratorn GCTTCCTCTGCGACTGTCACGGCAGGGCTCTGGATGCCGCGAGGACGCTGCGGTCTAGACTGGCGGCGGCCGCCGAGTCCGGGGGATCGGCGGCGTCCCCGAGCCTCGCGTACGAGACCA 90705

TeratornLC199500 **GGGCCCTGGAGTACCTGACGGAAAACCTCTCCTCCGCTCCACCTCCCGAGTGGGACTGCGTGCGCCGTTCCCCTGACGTTTCGGCCGAAGTGAAAATCGCGAGCGAGCTGCTCTGGTCCC** 87946
WF-Teratorn GGGCCTTGGAGTACCTGACGGAAAACCTCTCCTCCGCTCCACCTCCCGAGTGGGACTGCGTGCGCCGTTCCCCTGACGTTTCGGCCGAAGTGAAAATCGCGAGCGAGCTGCTCTGGTCCC 90825

TeratornLC199500 **ACTTGGTGCTGGTTTACGCCGGCAACCGCAGGCTGCTGCTCTTCAACGCGGAGCTGTTTTTCTCAAGGCAGCTACAGTCGACGGCCCTCTGCTCCAGCGAGCTCAGGGAGGCGCTGTTTT** 88066
WF-Teratorn ACTTGGTGCTGGTTTACGCCGGCAACCGCAGGCTGCTGCTCTTCAACGCGGAGCTGTTTTTCTCGAGGCCGCTACAGTCGACGGCCCTCTGCTCCAGCGAGCTCAGGGAGGCGCTGTTTT 90945

TeratornLC199500 **TGCACGGCAAGCATGGACCCTTCTCCCAGCGCAAAGGCCGCCGTCAGGCGCGAGGAGCGAGTATCGGCTCTGCGTGGCAGGGGCGAGACTGCCCCGTTCTTCCACCAGACCCGCGTGCAC** 88186
WF-Teratorn TGCACGGCAAGCATGGACCCTTCTCCCAGCGCAAAGGCCGCCTTCAGGCGCGAGGAGCGAGTATCGGCTCTGCGTGGCAGGGGCGAGACTGCCCCGTTCTTCCACCAGACCCGCGTGCAC 91065

TeratornLC199500 **AGCGAGGGCCTAGATCCGACAGGGGGGGAGGAATGTTGCGAGAGACCCGAGTGTCTCGACTGTGCTGGCTCGCGGTTTTGTGGCTCCTGGACCAGCACCGATGCCACAGCGAACTTGGTT** 88306
WF-Teratorn AGCGAGGGCCTAGATCCGACGGGGGGAGAGGAATGTTGCGAGAGACCCGAGTGTCTCGACTGTGCTGGCTCGCGGTTTTGTGGCTCTTGGACCAGCACCGATGCCACAGCGAACTTGGTT 91185

TeratornLC199500 **GGCTCAGTAGGCCCGTACCGTGTCGTCCCTAGCCCCGACGTGCTCGGGCGAGGCGGGGAGCCAGAGCCCCGGCGCCCTTCTCCGCAGGAGACAAACGACACCCTCAGGGCCGCCGGCAGC** 88426
WF-Teratorn GGCTCAGTAGGCCCGTACCGTGCCGTCCCTAGCCCCGACGTGCTCGGGCGGGGCGGGGAGCCAGAGCCCCGGCGCCCTTCTCCGCAGGAGACGAACGACACCCTCAGGGCCGCCGGCAGC 91305

TeratornLC199500 **TGCGTGGTGGACCTGAACGGCTCCGTGCTCCTCCACGTGCTTCACGTGAACACGGTGGGCAACCGCAAGTTCAGCGCCGTGGTGTACTCGGAAAGCCTCAGGCGCTACGCCCTTCAGCAG** 88546
WF-Teratorn TGCGTGGTGGACCTGAACGGCTCCGTGCTCCTCCACGTGCTTCACGTGAACACGGTGGGCAACCGCAAGTTCAGCGCCGTGGTGTACTCGGAGAGCCTCAGGCGCTACGCCCTTCAGCAG 91425

TeratornLC199500 **TCCAACACGACGAAAACCAAGCTCGTGAGCAGGCTCCGCTTCGTCTCGGACAAGAAGAAGCTCAGCTCCCGACACGGCTACAAGAACACCCGCATCGAGTTCTCGTCCAACCCCAGCCGC** 88666
WF-Teratorn TCCAACACAACAAAAACCAAGCTCGTGAGCAGGCTCCGCTTCGTCTCGGACAAGAAGAAGCTCAGCTCCCGACACGGCTACAAGAACACCCGCATCGAGTTCTCGTCCAACCCCAGCCGC 91545

TeratornLC199500 **AGCGCGAACTACACGTCCAACGCGAAGCACGCGCTGCAGCCCTACTTCAGGGGGCTGTTCGAGACCGTGTCCATGGCCCAGGGCACCTGCGTGGTCTCCGCTAAGCCCTTCGAGGTGAAC** 88786
WF-Teratorn AGCGCGAACTACACGTCCAACGCGAAGCACGCGCTGCAGCCCTACTTCAGGGGGCTGTTCGAGACCGTGTCCATGGCCCAGGGCACCTGCGTGGTCTCCGCTAAGCCCTTCGAGGTGAAC 91665

TeratornLC199500 **AACGAAAACCTCACGAGCTACATGTACGGCGTGGCGAACCCCGCGAGCGTGGTGATTCGCATGAAAGACACCAAAAAGGACTTTCACACTCCCTTCTTCTTCTCCTCGGTAAACGTGACG** 88906
WF-Teratorn AACGAAAACCTCACGAGCTACATGTACGGCGTGGCGAACCCCGCGAGCGTGGTGATTCGCATGAAAGACACCAAGAAGGACTTTCACACTCCCTTCTTCTTCTCCTCGGTAAACGTGACG 91785

TeratornLC199500 **GGCGAGGAGAACAGGCACCAACTTTTCTCCTCCATGTCCATCAATTCGCAGAGGGTGAAGAGCGCGGCCGAAGAACTCCCCCGGGCCCTGGCATCCGTGGCCTCGCTCGTGGAGCTCCAC** 89026
WF-Teratorn GGCGAGGAGAACAGGCACCAACTTTTCTCCTCCATGTCCATCAATTCGCAGAGGGTGAAGAGCGCGGCCGAAGAACTCCCCCGGGCCCTGGCCTCCGTGGCCTCGCTCGTGGAGCTCCAC 91905

TeratornLC199500 **GGCCACTTCAACAGACTTTGCTGCGAGCTGGCGCACGAGCGAGGTGGCACTCTCACGACCAGGGAAAAGGTCTGCCTGGTGTACCTCACGGAGATGTACTTTTTGCGAGTCAGGGACCGG** 89146
WF-Teratorn GGCCACTTCAACAGACTTTGCTGCGAGCTGGCGCACGAGCGAGGTGGCACTCTCACGACCAGGGAAAAGGTCTGCCTGGTGTACCTCACGGAGATGTACTTTTTGCGAGTCAGGGACCGG 92025

TeratornLC199500 **AACCACCCCACGCGCTGGCCCGGCTACAGGGAGGCGCTGAGGGGCTCATTCGCCGGCCGCTACGGGAGCCGGCTGACGGCGTTTCAGGACAGAGTCCTCTCCAACTACTACCAGAAAAAC** 89266
WF-Teratorn AACAACCCCACGCGCTGGCCCGGCTACAGGGAGGCGCTGAGGGGCTCATTCGCGGGCCGCTACGGGAGCCGGCTGACGGCATTTCAGGACAGAGTCCTCTCCAACTACTACCAGAAAAAC 92145

TeratornLC199500 **CCAGAGTTCGTCCACCTGGTCCCGTGGCTCCTGGACAACCCGAGCGTGCGGCACCGGGAGATGCTCTTGGCCCACGAAGCCTACCTGAGAGAGGAGGGAAAAGAGGCCGCTCGAGCAGAG** 89386
WF-Teratorn CCAGAGTTCGTCCACCTGGTCCCGTGGCTCCTGGACAACCCGAGCGTGCGGCACCGCGAGATGCTCTCGGCCCACGAAGCCTACCTGAGAGAGGAGGGAAAAGAGGCCGCTCGAGCAGAG 92265

TeratornLC199500 **CTGATAGCGGCCGTGGTCTGCCACTGCGAAAACGACCTGACCCGCCGCCCCCCGGAGGGCTTCCCTGACCTGCTGGACGTCGACTTCTCGCGACTCGGCTACAGTGAACTCACGCTGAGG** 89506
WF-Teratorn CTGATAGCGGCCGTGGTCTGCCACTGCGAAAACGACCTGACCCGCCGCCCCCCGGAGGGCTTCCCTGACCTGCGGGACGTCGACTTCTCGCGACTCGGCTACAGTGAACTCACGCTGAGG 92385

TeratornLC199500 **CACGCGCTGGACAGGTCCACCAAGGCCAGCGACGACGCTCGCCCAGGCGGTCGGGCTAAATGACCGGCGCGGTCCCCACGATAGCGAGCCCCCC-TACCATGACGACCGAAGTGAGCGGC** 89625
WF-Teratorn CACGCGCTGGACAGGTCCACCAAGGCCAGCGACGACGCTCGCCCAGGCGGTCGGGCTAAATGACCGGCGCGGTCCCCACGATAGCGAGCCCCCCCTACCATGACGACCGAAGTGAGCGGT 92505

TeratornLC199500 **GGCACCGAACTCGGGAGCCTCCCTCGAGCTTCCATCCTCAGGGTCTGGGAGATAGTGAACCGGCACTACGAATCCACCTTGAGCTCGCTCGAACGAGAGGACACCTCTCGTAGAGCCGCG** 89745
WF-Teratorn GGCACCGAACTCGGGAGCCTCCCTCGAGCTTCCATCCTCAGGGTCTGGGAGATAGTGAACCGGCACTACGAATCCACCTTGAGCTCGCTCGAACGAGAGGACACCTCTCGTAGAGCCGCG 92625

TeratornLC199500 **GCTCGGGTGAGCTCGCTGGCGAAGGAGCTGCGTGACCTGACCGCAGCCTGCACGGAGAGGCTCGACCGCGTGGAAGCTCGCGTCAAGGAACTCAAGGCCCTCGAGTTCGGCGGAGTTCGC** 89865
WF-Teratorn GCTCGGGTCAGCTCGCTGGCGAAGGAGCTGCATGACCTGACCGCAGCCTGCACGGAGAGGCTCGAACGCGTGGAAGCTCGCGTCAAGGAACTCAAGGCCCTCGAGTTCGGCGGAGTTCGC 92745

TeratornLC199500 **AACGAGCAGCCCGACGGAGCGCGGGCCGGCGAGGCGCTCGGCGGAGAGTGGCACTTTGTCTACACGCGCCGAGGAGATGTGTTCGGATACATGAGGCTGATCCCCGAAACGCTCTCGCGG** 89985
WF-Teratorn AACGAGCGGCCCGACGGAGCGCGGGCCGGCGAGGCTCTCGGCGGAGAGTGGCACTTTGTCTACACGCGCCGAGGAGATGTGTTCGGATACATGAGGCTGATCCCCGAAACGCTCTCGCGG 92865

TeratornLC199500 **CTCCCCTCGGACGCGGACGACACGAGCCTCTGCGGCGGGCAGAGGGCCGAGTTCACGCTCCCGACACAGGTGGCGGCGGAGGCCCGTCGCGTCCCCACGGGCCGGTCCGGGAGCGCCCGA** 90105
WF-Teratorn CTCCCCTCGGACGCGGACGACACGAGCCTCTGCGGCGGGCAGAGGGCCGAGTTCACGCTGCCGACACAGGTGGCGGCGGAGGCCCGTCGCGTCCCCACGGGCCGGTCCGGGAGCGCCCGA 92985

TeratornLC199500 **GGACGAGCGGAGTGCCCCCGCTGCGGCGAGTCTCTGGATCTCTCGGAGTTCTGGAGGCACGCGGTGTATGAGATCGCAGAGAGGGGAGCCAGGGTTCCTGCGCGCCGCCCCCCGGTTCGT** 90225
WF-Teratorn GGACGAGCGGAGTGCCCCCGCTGCGGCGAGTCTCTGGATCTCTCGGAGTTCTGGAGGCACGCGGTGTATGAGATCGCAGAGAGGGGAGCCAGGGTTCCTGCGCGCCGCCCCCCCGTTCGT 93105

TeratornLC199500 **TCAGACGGTCCGCGACCGAGGTGGACCTTGCAAGAGGGCGAAGGTTGGGACGAGCCCATTTACAGGCCCAGAGACCTGACGAGCACGTGCGAAATCGGCGAGTGCCTCTCCCAGCTGAAC** 90345
WF-Teratorn TCAGACGGACCGCGACCGAGGTGGACCTTGGAAGAGGGCGAAGGTTGGGACGAGCCCATTTACAGGCCCAGAGACCTGACGAGCACGTGCGAAATCGGCGAGTGCCTCTCCCAGCTGAAC 93225

TeratornLC199500 **TTGAAACACTTTCTCTCGGACGATGAGTTCTCCGGGGCCCTGGACCGCGCTTTCGAACTCTTGGCTCTAAGGTTGAGATCTTAA**CGGAGACCACAAAGACGACTCGGCTGTTGACAAAAA 90465
WF-Teratorn TTGAAACACTTTCTCTCGGACGATGACTTCTCCGGGGCCCTGGACCGCGCTTTCGAACTCTTGGCCCTGAGGTTGAGATCTTAACGGAGACCACAAAGACGACACGGCTGTTGGCAAAAA 93345

TeratornLC199500 AAAAAAGGTTGAAAGATGAAGAGAAGAATCGAAAGGAAGAAAAAAAAAAA-GAAAAAGAAACGGTTGTGTGCACGCAGAAAGTTGTTGACAGCGAACACATCTGAATAAAAATAAAAGTTG 90584
WF-Teratorn AAAAAAGGTTGAAAGATTAAGAGAAGAATCGAAAGGAAGAAAAAAAAAAAGAAAAAGAAACGGTTGTGTGCACGCAGAAAGTTGTTGACAGCGAACACATCTGAATAAAAATGAAAGTTG 93465

TeratornLC199500 TGAGAGAACAAACACAGTGCTGTTTCATTGTTTTATTGCGTCAAGGTTTCATAAGCATACACAATGAAGTATGTTCAGGTGTTACAGTTCCACTGTGACCGTGAGGGCTTTGGGTTGCTC 90704
WF-Teratorn TGAGATAACAAACACAGTGCTGTTTCATTGTTTTATTGCGTCAAGGTTTCATAAGCATACACAATGAAGTATGTTCAGGTGTTACAGTTCCACTGTGACCGTGAGGGCTTTGGGTTGCTC 93585

TeratornLC199500 AAGGCCGGTTGATGTTCACGAAGCGGGTGCTCAACGGTCAATCGTGCCTGGTA-GCGAACAGGAAGGCTTCGACTGTCAAAGCAGAGCCACCGGGGACAAAGGGTCGGAGAGGGGAAGCT 90823
WF-Teratorn AAGGCCGGTTGATGTTCACGAAGCAGGTGCTCAACGGTCAATCGTGCCTGGTACGCGAACAGGAGGGCTTCGACTGTCAAAGCAGAGCCACCGGGGACAAAGGGTCGGAGAGGGGAAGCT 93705

TeratornLC199500 TGGACGAAACATTTTGTATTTTCGAAAAAGGAAGACTTTTTCTGTCGGGAAATCTCAGAGTGGGGCAATGAGCTGTAAAAGAACAACAACAAATAGTGGTTGAAAGCAGACGTGTGCAAA 90943
WF-Teratorn TGGACGAAACATTTTGTATTTTCGAAAAAGGAAGACTTTTTCTGTCGGGAAATCTCAGAGTGGGGCAATGAGCTGTGAAAGAACAAAAACAAATAGTGGTTGAAAGCAGACGTGTGCAAA 93825

TeratornLC199500 CGAAAAGCAACGAGAGAAGAAAACTGAAAGTTGTTTAATTGCCTCAATGTCTCAAGCCTTCAAAATAAAGTACGTTCAGGTGTGACAAAGACTTTTTTTTGTCCTTTTATTTGTTCGGAG 91063
WF-Teratorn CGAAAAGCAACGAGAGAACAAAACTGAAAGTTGTTTCATTGCCTCAATGTCTCAAGCCTTCAAAATAAAGTACGTTCAGGTGTGACAAAGACTTTTTTTTGTCCTTTTATTTGTTCGGAG 93945

TeratornLC199500 ATTTCCGAGGTACTTTTCGCGAAGAAAGTGGTCAAAGGCAAACGCTTGCGAATAAAATTGTACGAGAGCTGTTTCATTTGTTTTCTATCGTCTCGAATTTCTGCGAAATGATATTCCGGC 91183
WF-Teratorn ATTTCCGAGGTACTTTTCGCGAAGAAAGTGGTCAAAGGCAAACGCTTGCGAATAAAATTGTACGAGAGCTGTTTCATTTGTTTTCTATCGTCTCGAATTTCTGCGAAATGATATTCCGGC 94065

TeratornLC199500 ACAACAGTTTGACCGTGACCGTCGTCTGCCAGTAGGTGCAGCCCAGAGCGTGCAGATCCCGGCGCACGCAGAGAGCGAATCGAGATTCCGGACTCCCGAGGACGCTCCGATTTTGACCCG 91303
WF-Teratorn ACAACAGTTTGACCGTGACCGTCGTCAGCCAGTAGGTGCAGCCCAGAGCGTGCAGATCCCGGCGCACGCAGAGAGCGAATCGAGATTCCGGACTCCCGAGGACGCTCCGATTTTGACCCG 94185

TeratornLC199500 CTTATTTACAAGCCCAGAGACCTGTTGATTACCCGCGAAATCGACGAGAGCCGCTTCCACCGGCCACCATGTTCTCATACTCCTACAAGCCGAGCGCAGCCTTCGCCTCGGCTCCGCTCG 91423
WF-Teratorn CTTATTTACAAGCCCAGAGACCTGTTGATTACCCGCGAAATCGACGAGAGCCGCTTCCACCGGCCACCATGTTCTCATACTCCTACAAGCCGAGCGCAGCCTTCGCCTCGGCTCCGCTCG 94305

TeratornLC199500 GAGCCCGTCCCTCCGCCGCAACGCCCACAGTCAGCGGTGTGGCGAGACAGGCCGAGAAAGATCTCCGCTCGGAGAACGAGGGGGTCGACGGCGCGGCGGTCGGGCCTTCTCGGGCGGCTC 91543
WF-Teratorn GAGCCCGTCCCTCCGCCGCAACGCCCACAGTCAGCGGTGTGGCGAGACAGGCCGAGAAAGATCTCCGCTCGGAGAACGAGGGGGTCG---------------GGCCTTCTCGGGCGGCTC 94410

TeratornLC199500 CTGGGATCTACGAGACCCTTGCGAGCGATGAAGTGGTCACGCTCGACTGCCTGACTGGGGTGGTGCGATCTGAGCGGGGCTGCACGTTCGTGCAGTACATCCCGACGGAGGAGCACAAGC 91663
WF-Teratorn CTGGGATCTACGAGACCCTCGCGAGCGATGAAGTGGTCACGCTCGACTGCCTGACTGGGGTGGTGCGATCTGAGCGGGGCTGCACGTTCGTGCAGTACACCCCGACGGAGGAGCACAAGC 94530

TeratornLC199500 TGCGCGCCGGGCGGAAGAGCGCCCCAGTGGAACAGATCAACAGGTTCCTCAGGTCG**TTAGGGGACAGCGGCGTGCCAGCTGAAACCTCCTTGATTGCGCTCTGCCCGTTGGAGCCGGGCC** 91783
WF-Teratorn TGCGCGCCGGGCGGAAGAGCGTCCCAGTGGAACAGATCAACAGGTTCCTTAGGTCGTTAGGGGACAGCGGCGTGCCAGCTGAAACCTCCTTGATTTCGCTCTGCCCGCTGGAGCCGGGCC 94650

TeratornLC199500 **GCCGGAAGACCCTGTGGATAAGGGTAGCGCTGGAGGAAGTGCAAGAACTCAGCCTGGAAGCCCTGTGGATGCTCAAGGGCGAAGCGCGGCTTCACGCGGAGAGGACCAGCCTCGTCGTGT** 91903
WF-Teratorn GCCGGAAGACCCTGTGGATAAGGGTAGCGCTGGAGGAAGTGCAAGAACTCAGCCTGGAAGCCCTGTGGATGCTCAAGGGCGAAGCCCGGCTTCACGCGGAGAGGACCAGCCTCGTCGTGT 94770

TeratornLC199500 **GGCTGGAGTCGAGCGCGCTGGGTTCCAGCTCGCTGAGCCCGACGGCCGAACGAACCCTGTGCCCCGCATCCGAAAGGGACGCGGAACGTCTCGCCGCTCTCGCCGAAGAGGGCCTCCTGA** 92023
WF-Teratorn GGCTGGAGTCGAGCGCGCTGGGTTCCAGCTCGCTGAGCCCGACGGCCGAACGAACCCTGTGCCCCGCATCCGAAAGGGACGCGGAACGTCTCGCCGCTCTCGCCGAAGAGGGCCTCCTGA 94890

TeratornLC199500 **GCCCCTGTGAGAGAGGAGAGCGGCGCGAAGAGGGCTCGAGTCGAGAGGCCGCCCGCGTCTCGACACAA**CGAGACGAGTCGGGGCCCGGACGGTCCGGTCTGAAGCGCGTGAGGGTCAGAA 92143
WF-Teratorn GCCCCTCTGAGAGAGGAGAACGGCGCGAAGAGGGCTCGAGTCGAGAGGCCGCCCACGTCTCGACACAACGAGACGAGTCGGGGCCTGAACAGTCCGGTCTGAAGCGCGTGAGGGTCAGAA 95010

TeratornLC199500 CGGGACTGGTTGGCGACTGCCCAACACACAGCCTACCCGAGAATTCGAGAGACGGCCAGCCGTGCAGGAGATCGAAAAGACAAATACGACGGCGGGCGCAGTCCTCCGGGGTAGGTTTCA 92263
WF-Teratorn CGGGAGAGGTTGGCGACTGCCCAACACACAGCCTACCCGAGAACTCGAGAGACGGCCAGCCGTGCAGGAGATCGAAAAGACAAATACGACGGCGGGTGCAGTCCTCCGGGGTAGGTTTCA 95130

TeratornLC199500 TCAGAAAAAGAATCAAAGCTACAAAGACAGCCGAGAAGAGGGTCGAAGCGGAGTCGACGTTTGCTCGATTCGGACGGGAGAACTCCACGATCTCTCCTGAAGAGACTTCATCGCTTCCTC 92383
WF-Teratorn TCAGAAAAAGAATCAAAGCTACAAAGGCAGCCGAGAAAAGGGTCGAAGCGGAGTCGACATTTGCTCGATTCGGACGGGAGAACTCCACGATCTCTCCTGAAGAGACTTCATCGCTTCCTC 95250

TeratornLC199500 CTCCTCCTCCGCCTCAGCCTCCCGTTCGTCCTCACGATCCTGTTCCCCGTCACGGCACCGTCAAGGCCCCCACGCGCAAAGCGCAGGATCGGGCCGAAAGAGAGCTCGGGGCTTTGTCCC 92503
WF-Teratorn CTCCTCCTCCGCCTCAGCCTCCCGTTCGTCCTCACGATCCTGTTCCCCGTCACGGCACCGTCAAGGCCCCCACGCGCAAAGCGCAGGATCGGGCCGAAAGAGGGCTCGGGGCTTTGTCCC 95370

TeratornLC199500 TCTCGTCTCGGCGAGTAGAAAGGCAGCCCCGGGCGCTTGGCGGAGAGTCTCCACCGTGTTCGGTTCCCAGTTTCGGACAGGCCAGAGAGACCAGTCTGTCGCCTTCCGGGGAGAGAGTCC 92623
WF-Teratorn TCTCGTCTCGGCGAGTAGAAAGGCAGCCCCGGGCGCTTGGCGGAGAGTCTCCACCATGTTCGGTTCCCAGCTTCGGAGAGGCCCGAGAGACCAGTCTGTCGCCTTCCGGGGAGAGAGTCC 95490

```
TeratornLC199500  GTGTCTCGTCGTCTCATCAACGCCCGGTGCCCACGGCGGTTTCGGCCGAAGCTTCTCGGGGCTGCGGGCTGATGTAGAGCGCGGACGGGTCGTGCGGAGCACCCGGGACCCGCTCTTCCG  92743
WF-Teratorn       GTGTCTCGTCGTCTCATCAACGCCCGGTGCCCACTGTGGTTTCGGCCGAAGCTTCTCGGGGCTGCGGGCTGATGTAGAGCGCGGACGGGTCGTGCGGAGCACCCGGGACCCGCTCTTCCG  95610

TeratornLC199500  TCGTGACCTCTTCGCAGCTTTTCACGTCAGAGGGCCTGAGGCCGAGTTTGTTCGCCAACAGCAGGCCGTAGGCAGGCGGGTGCAGGGAATCCTTTGTGAGCATGATATCGCTCACGGAGA  92863
WF-Teratorn       TCGTGACCTCTTCGCAGCTTTTCACGTCAGAGGGCCTGAGGCCGAGTTTGTTCGCCAACAGCAGGCCGTAGGCAGGCGGGTGCAGGGAATCCTTTGTGAGCATGATATCGCTCACGGAGA  95730

TeratornLC199500  GGGACCCGTCTGGCTTTCTCCCCGACAGGCTGTCGGCGAACGTGGCACCGGACGTGTTGAGGAGAGCGACCAGGGCGCTGTGGAAGATGTGCACGGGGGATCCGGACTCCAGTTCTCTCA  92983
WF-Teratorn       GGGACCCGTCTGGCTTTCTCCCCGACAGGCTGTCGGCGAACGTGGCACCGGACGTGTTGAGGAGAGCGACCAGGGCGCTGTGGAAGATGTGCACGGGGGATCCGGACTCCAGTTCTCTCA  95850

TeratornLC199500  CGCCGTCCCTGTGGTTGTACTGTATGGTGTTGTGTCTCCTGTTGAAGTGGCTGAATCCCGACAAGGCGCTACGCCAAAAATTTGTCCAAATGCTCAGAGTTTCCTCCAGGTGAGCAACGC  93103
WF-Teratorn       CGCCGTCCCTGTGGTTGTACTGTATGGTGTTGTGTCTCCTGTTGAAGTGGCTGAATCCCGACAAGGCGCTACGCCAAAAATTTGTCCAAATGCTCAGAGTTTCCTCCAGGTGAGCAACGC  95970

TeratornLC199500  TGACAGAGGACGGGTAGCTGCACACCAGCCTCACGTCCGAGCCGAGCTCGGTCACGGGATCCCGGTCTCCTTCGGCCCCGTCCTCCCGCTCGGCGAGTCGACGTTCCAGACGCCTGGACA  93223
WF-Teratorn       TGACAGAGGACGGGTAGCTGCACACCAGCCTCACGTCCGAGCCGAGCTCGGTCACGGGATCCCGGTCTCCTTCGGCCCCGTCCTCCCGCTCGGCGAGTCGACGTTCCAGACGCCTGGACA  96090

TeratornLC199500  CGTTCTCCGCCATCCTCAGCAGCCCGGGATCCTCGCACTCGGACCTGAGCTTTTCCAGAGCCGCGCGAGGACGAGAGGTGACCTCGGTCAGAAATCCGCGCACCAGCAGGACCTCTTTGC  93343
WF-Teratorn       CGTTCTCCGCCATCCTCAGCAGCCCGGGATCCTCGCACTCGGACCTGAGCTTTTCCAGAGCCGCGCGAGGACGAGAGGTGACCTCGGTCAGAAATCCGCGCACCAGCAGGACCTCTTTGC  96210

TeratornLC199500  CTTCGGATATCATGGAGGCCATCATCTTGCTGAGGCTGTGGGACACGCTGGCGGACTTGTGCTTGGACACCGAGGCGCACGCGCCCGACTGCAGCAAGGAGTCGGGCAGCTCGGCGTCGT  93463
WF-Teratorn       CTTCGGATATCATGGAGGCCATCATCTTGCTGAGGCTGTGGGACACGCTGGCGGACTTGTGCTTGGACACCGAGGCGCACGCGCCCGACTGCAGCAAGGAGTCGGGCAGCTCGGCGTCGT  96330

TeratornLC199500  TCAGCAGTCCCAGCATGTCCGTCTGTTGGTTGCAGGCGAGGGCTCGCGACAGGTTGTCCAGCAAAGGTCTGTCGGTCGCGTTGCCTACCACCGAGATGCCAGCGTCCCGCTGCTCGGCCC  93583
WF-Teratorn       TCAGCAGTCCCAGCATGTCCGTCTGTTGGTTGCAGGCGAGGGCTCGCGACAGGTTGTCCAACAAAGGTCTGTCGGTCGCGTTGCCTACCACCGAGATGCCAGCGTCCCGCTGCTCGGCCC  96450

TeratornLC199500  TCGAGGAGAGCATCACGTCTGCCACGCGCGCCAGCGACGACCGATCGGCCCCCGCGCTGGTGTACACGTGCCTCGCCGCGGCCTCGTTCAGACCCCTGGTTATCCTGCCGAAAAGCGTCC  93703
WF-Teratorn       TCGAGGAGAGCATCACGTCTGCCACGCGCGCCAGCGACGACCGATCGGCCCCCGCGCTGGTGTACACGTGCCTCGCCGCGGCCTCGTTCAGACCCCTGGTTATCCTGGCGAAAAGCGTCC  96570

TeratornLC199500  TCCAGTTCTTGACGGTGTCGTGAGCCCTCTCTGCCAAGGAGGAAAAGGAAGGACCCGAAGCCGACCAGCTGCTGTCCCACACCGACGCTCGTCTGTAGAGGACCCCGTCCACCGCCGCAG  93823
WF-Teratorn       TCCAGTTCTTGACGGTGTCGTGAGCCCTCTCCGCCAAGGAGGAAAAGGAAGGACCCGAAGCCGACCAGCTGCTGTCCCACACCGACGCTCGTCTGTAGAGGACCCCGTCCACCACCGTGG  96690

TeratornLC199500  TGACGAGAGCGGTGCGCTTGTCGTAGAAGGCCAAGCACGAGGAGGCTCTGCACTCGGGCACCACCGGGCACCGTCCGTGTTCGAAGGTCTTCATCACACCCCATAGAGTCACCACAGACA  93943
WF-Teratorn       TGACGAGAGCGGTGCGCTTGTCGTAGAAGGCCAAGCACGAGGAGGCTCTGCACTCGGGCACCACCGGGCACCGTCCGTGTTCGAAGGTCTTCATCACACCCCATAGAGTCACCACAGACA  96810

TeratornLC199500  TCTTCAGCTCTGTCTCCTTCACGGGGTTCCGAACGAAAGCCTCGCGCTCTCCCCTCACCCAGTCCGCGACTATCTTACACACCACGGCCGTCGGCGCGCAGGGCACAGACGCCATCATCA  94063
WF-Teratorn       TCTTCAGCTCTGTCTCCTTCACGGGGTTCCGAACGAAAGCCTCGCGCTCTCCCCTCACCCAGTCCGCGACTATCTTACACACCACGGCCGCCGGCGCGCAGGGCACCGACGCCATCATCA  96930

TeratornLC199500  GCAACTTGGCTGTCTCCGCCAGACTGTCCAAGGGAACTTCGAGCACGCGAGGAAGCGCCTCGTCGTGAGCCTCTTTCTCTTGGATCGCTTCGGCCGCGGCCCCGGGAGGGCTGAGGCCGT  94183
WF-Teratorn       GCAACTTGGCTGTCTCCGCCAGACTGTCCAAGGGAACTTCGAGCACGCGAGGAAGCGCCTCGTCGTGAGCCTCCTTCTCTTGGATCGCTTCAGCCGCGGCCCCGGGAGGGCTGAGGCCGT  97050

TeratornLC199500  CCGGCGACACCACAAACAGGCGTCCGGCACTACGGGGCGGGGCCGCTTCGGCATCTTTCCTGCAGCGCTTGCTGGGGCGCGCGTCCGGCCCGCCGGAGCCCGAAACCATGTTGATCACGG  94303
WF-Teratorn       CCGGCGACACCACAAACAGGCGTCCGGCACTACGGGGCGGGGCCGCTTCGACATCTTTCCTGCATCGCTTGCTGGGGCGCGCGTCCGGCCCGCCGGAGCCCGAAACCATGTTGATCACGG  97170

TeratornLC199500  CAGCTTTGCCAGACAGGCCGAGGCCGAGGGATCTGATTTTCTTCCACGGCAGGAAAAGGCAAGGCACGAAGCCCGCCAGCACGGTCCAGCGGATCATGTTTCTCAGGGTGCTTTTGATGT  94423
WF-Teratorn       CAGCTTTACCAGACAGGCCGAGG------GATCTGATTTTCTTCCACGGAAGGAAAAGGCAAGGCACGAAGCCCGCTAGCACGGTCCAGCGGATCATGTTTCTCAGGGTGCTTTTGATGT  97284

TeratornLC199500  GCCCGCAGCAGGGGCGGTTGGCCACGTAGTGCCCGACGAGATAGCGGACCAGGCACGCGGCGCTCTCGCTGTTGCTGCCCAGCACGTTGCACAGCAGCAACTCCACCGACAGGGCCGGGC  94543
WF-Teratorn       GCCCGCAGCAGGGGCGGTTGGCCACGTAGTGCCCGACGAGATAGCGGACCAGGCACGCGGCGCTCTCGCTGTTGCTGCCCAGCACGTTGCACAGCAGCAACTCCACTGACAGGGCCGGGC  97404

TeratornLC199500  TGAAAGTGTTCACGATCTCCTCTATCACCACGGCCAGCTGCGGATCCTCGCTCACAAAAGCCCAAGATGCGACGTCCCTGTGTAGCGGCAACAGGTCGTAGCACTGCTCGTAGCAGTCCC  94663
WF-Teratorn       TGAAAGTGTTCACGATCTCCTCTATCACCACGGCCAGCTGCGGATCCTCGCTCACAAAAGCCCAAGACGCGACGTCCCTGTGTAGCGGCAACAGGTCGTAGCACTGCTCGTAGCAGTCCC  97524

TeratornLC199500  AGCGCGCCTCTCCGCGGGGGCGATTACGGTTGCGTGCACGATGAGCGCAACGGCCGTTCCTGAAGTCCTCTAGAACCCTCGCGTCCCCTCTGTCCGCCAAGGCGCACTCGGCCAAGGCCG  94783
WF-Teratorn       AGCGCGCCTCTCCGCGGGGGCGATTACGGTTGCGTGCACGATGAGCGCAACGGCCGTTCCTGAAGTCCTCTAGAACCCTCGCGTCCCCTCTGTCCGCCAAGGCGCACTCGGCCACGGCCG  97644

TeratornLC199500  CCTCCCGGACGTCGAGCGGTTCTTCGTCCGCCTTCATCGTCACAATTCTGTTCGACTCGACTTTCGGACGGTAACCGGGAGGGGTTAATGAGAATCCGCTCCTCAGCTCCTCACCCGTAC  94903
WF-Teratorn       CCTCCCGGACGTCGAGCGGTTCTTCGTCTGCCTTCATCGTCACAATTCTGTTCGACTCGACTTTCGGACGGTAACCGGGAGGGGTTAATGAGAATCCGCTCCTCAGCTCCTCACCCGTAC  97764
```

TeratornLC199500 TCAGCTGCTCCCGCCGCGTCTTCGACCCCTCTCGTCGGCC**ATGTTCTGCGTACTGTTCAAGGCGCTCTGCTGCCTTTGCGGCAACGGGCGGACGGCGCGCGGCCCTAGCGGGTACCGTTA** 95023
WF-Teratorn TCAGCTGCTCCCGCCGCGTCTTCGACCCCTCTCGTCGGCCATGTTCTGCGTACTGTTCAAGGCGCTCTGCTGCCTTTGCGGCAACGGGCGGACGGCGCGCGGCCCTAGCGGGTACCGTTA 97884

TeratornLC199500 **TGAGGTCCTCGGAGACAGCGGCGACGAGTACGGCGTCGAGGAAGGCGACGAGGCGGACGAAGCCGAACCTCGCCTCAGGGACTTCGCGCTGACGCGCAAGAGAGCTATCGGCCTGACCAG** 95143
WF-Teratorn TGAGGTCCTCGGAGACAGCGGCGACGAGTACGGCGTCGAGGAAGGCGACGAGGCGGACGAAGCCGAACCTCGCCTCAGGGACTTCGCGCTGATGCGCAAGAGAGCTATCGGCCTGACCAG 98004

TeratornLC199500 **GTTGAAACTGAAAGCTATCAACTCCGCCTCGGACGTGTCCGTGGCGGAGAGGCTCTTGGAACGCCACGGTCAAAATGACCCAAGCTTACTAACTTACGAGGGCAGGTGCCGCGAGGAAAT** 95263
WF-Teratorn GTTGAAACTGAAAGCTATCAACTCCGCCTCGGACGTGTCCGTGGCGGAGAGGCTCTTGGAACGCCACGGTCAAAATGACCCAAGCTTACTAACTTACGAGGGCAGGTGCCGCGAGGAAAT 98124

TeratornLC199500 **GCTGGCGGCTTTGAGCAAGCTCCCACCTTACGCGCACGGGACCCAGATCGACAGACCTTACAGGATAACGCCAGCCTCGAGCATCGGAAGCGAACGCAACGGAGCGATCGTGTGA**GCGCA 95383
WF-Teratorn GCTGGCGGCTTTGAGCAAGCTCCCACCTTACGCGCACGGGACCCAGATCGACAGACCTTACAGGATAACGCCAGCCTCGCGCATCGGAAGCGAACGCAACGGAGCGATCGTGTGAGCGCA 98244

TeratornLC199500 AGGGTCCCTTTCGCCGAAGAGGGGGGGGGGTGAAAAGAAAGAGAAAAGGAAAAAAGAAAAAGAAAGAAAAAAGAAATAAATCACC**ATGAGCGTGGTCAAGACTGACCCCAGCGTTTACTC** 95503
WF-Teratorn AGGGTCCCTTTCGCCGAAGAGGGGGGGGGG----------GTGAAAAGGAAAAAAGAAAAAGAAAGAAAAAATAAATAAATCACCATGAGCGTGGTCAAGACTGACCCCAGCGTTTACTC 98354

TeratornLC199500 **GCTGAGATCTCTGGAGGCTGCCAAACGACTGGTCAAAGCGCCTGGGTTTCTGGCCGACTCGACCGGGGCGTTCTCCGAGCTCGTGGCGGAGAACGTGATCAGGGCGTCCGCGGCCAACAC** 95623
WF-Teratorn GCTGAGATCTCTGGAGGCTGCCAAACGACTGGTCAAAGCGCCTGGGTTTCTGGCCGACTCGACCGGGGCGTTCTCCGAGCTCGTGGCGGAGAAGGTGATCAGGGCGTCCGCGGCCAACAC 98474

TeratornLC199500 **CTCGGTGTCGGCCGACGCGGTCGTGATGATACTGCAGGCCCACATGGGAGTGTCTCTCTTCAAGCAGCAGAAGGACACGAGCCGTTTGCTGGCGGGCTTCGTGTACAAGATGTTGATCCC** 95743
WF-Teratorn CTCGGTGTCGGCGGACGCGGTCGTGATGATACTACAGGCCCACATGGGAGTGTCTCTCTTCAAGCAGCAGAAGGACACGAGCCGTTTGCTGGCGGGCTTCGTGTACAAGATGTTGATCCC 98594

TeratornLC199500 **GGACAAGTCCGTGCCCACGGGCACGAGGTTCACGAGGCGTTACAAGATCACGGATCAGCGAGAGGCCCAGCGAGGTACCTACGACACGGTGGGAAACGGGGTTCGGACCAGGGAGTACAG** 95863
WF-Teratorn GGACAAGTCCGTGCCCACGGGCACGCGGTTCACGAGGCGTTACAAGATCACGGATCAGCGAGAGGCCCAGAGAGGTACCTACGACACGGTGGGAAACGGGGTTCGGACCAGGGAGTACAG 98714

TeratornLC199500 **CCAGGTGTTTCAGCACACGTCCAAAGTGGTGACCATGCAGTTTGGATTTCTGCTCGGCATCACGGTGCCTTCCGTGGCGGAGGAGCTGATACGCAGGGAGCTGGAGCAGGCCAGGACTTC** 95983
WF-Teratorn CCAGGTGTTTCAGCACACGTCCAAAGTGGTGACCATGCAGTTTGGATTTCTGCTCGGCATCACGGTGCCTTCCGTGGCGGAGGAGCTGATACGCAGGGAGCTGGAGCAGGCCAGGACTTC 98834

TeratornLC199500 **GTGGGAACTCACCAAGACCATAGAGGTCCTGGAGTACTGCGCGGCTCAGCCCACGCTCACGGACAGGATCTGCCAGAAACTCGCATCCCTCCCCAGAGCGGAGAAGCTGGACTCTGTGCT** 96103
WF-Teratorn GTGGGAGCTCACCAAGACCATAGAGGTCCTGGAGTACTGCGCGGCTCAGCCCACGCTCACGGACAGGATCTGCCAGAAACTCTCATCCCTCCCCAGAGCGGAGAAGCTGGACTCTGTGCT 98954

TeratornLC199500 **GACGATGGCCGAGCTCATGTGCGGCACCGTGAACGCGCATCCGGATTCTTTTGTGTGCGTGGTGGAGAACGCTCGCAGGGTCCTCTGCAGCGACAGGCCAGAGGTCCTCGTCATGGGAGA** 96223
WF-Teratorn GACGATGGCCGAGCTCATGTGCGGCACCGTGAACACGCATCCGGATTCTTTCGTGTGCGTGGTGGAGAACGCTCGCAGGGTCCTCTGCAGCGACAGGCCAGAGGTCCTCGTCATGGGAGA 99074

TeratornLC199500 **CACCCTCTTCAGGGTGGTGTCCCAAGACTCCAGTGCCGTCAGGACGCGCATCGCTTGCGAAAACGTGAAGCTCGCGCCCGAGTTCTGCGTCTACACCGCCGCCGACGAGAAGTCGCGGGG** 96343
WF-Teratorn CACCCTTTTCAGGGTGGTGTCCCAAGACTCCAGCGCCGTCAGGACGCGCATCGCTTGCGAAAACGTGAAGCTCGCGCCCGAGTTCTGCGTCTACACCGCCGCCGACGAGAAGTCGCGGGG 99194

TeratornLC199500 **CGACGGGCTCCACGAGTTCACCATGGAGTTGGTCAACAAACGCGAGCTGGGCGACGGGGACCTGGCTTTCGCGGACGACTCCAGCTTCAGGGACCAGGTCACGGAGATGCCGGTGAACCA** 96463
WF-Teratorn CGACGGGGTCCACGAGTTCACCATGGAGTTGGTCAACAAACGCGAGCTGGGCGACGGGGACCTGGCTTTCGCGGACGACTCCAGCTTCAGGGACCAGGTCACGGAGATGCCGGTGAACCA 99314

TeratornLC199500 **CATTCTGATCGGCGGCGGGCGTTCGGTCGGCATTCCCATCGCGCTGGTGGACGGCGTGCAGATGGAACACGGATGCGGGAGGGAGACGTCCACTGAGCACAAAGCTTTCAGCAACACCCA** 96583
WF-Teratorn CATTCTGATCGGCGGCGGGCGTTCGGTCGGCATTCCCATCGCGCTGGTGGACGGCGTGCAGATGGAACACGGATGCGGGAGGGAGACGTCCACGGAGCACAAAGCTTTCAGCAACACCCA 99434

TeratornLC199500 **GGGTTTCAAGTGGGAGTACTTCACGGTGGGCTGCACGGCTCCGGCGCTCGGCCAGATATCGCCGTCCAAAGTGCTCTCTGCCACTGACTCCGAAGAGGTCGGCTACAAGCAGCTGGCCCA** 96703
WF-Teratorn GGGTTTCAAGTGGGAGTACTTCACGGTGGGCTGCACGGCTCCGGCGCTCGGCCAGATATCGCCGTCCAAAGTGCTCTCTGCCACTGACTCCGAAGAGGTCGGCTACAAGCAGCTGGCCCA 99554

TeratornLC199500 **GCGATCGCCCTCGGTCTCGTACCTGCACCAGTACGACGGCGCCGTGGTCTGCGTGACGCTGAGGGAGCTGCACCGCCTCGGCAGTCCTCGAGCTCTCTTCGGCAGGGAGCGCAGGGACAA** 96823
WF-Teratorn GCGATCGCCCTCGGTATCGTACCTGCACCAGTACGACGGCGCCGTGGTCTGCGTGACGCTGAGGGAGCTGCACCGCCTCGGCAGTCCTCGAGCTCTCTTCGGCAGGGAGCGCAGGGACAA 99674

TeratornLC199500 **GCTGCAGGGGGACTTGAACGCCTTCATGAGGAACGGACCCAGAGGGATCGCCGTGGGACGCATGGCGCAGTTGGCGCACAGCGATGACTGGTACCAGAAGATCAAGAACCCCGCGTCGCT** 96943
WF-Teratorn GCTGCAGGGGGACTTGAACGCCTTCATGAGGAACGGACCCAGAGGGATCGCCGTGGGACGCATGGCGCAGTTGGCGCACAGCGATGACTGGTACCAGAAGATCAAGAACCCCGCGTCGCT 99794

TeratornLC199500 **GCTGGAGTTCGACGTTCGAAACATGGTTGTGCCCGAAGCCCCGGACGACGCCTCTCACGGATCGGCCGCCGTCGAGCCCGTGCGCTGGGTCGTCAGGCCCAGGGTGGCCCTGTTCAACAC** 97063
WF-Teratorn GCTGGAGTTCGACGTTCGAAACATGGTTGTGCCCGAAGCCCCGGACGACGCCTCTCACGGATCGGCCGCCGTCGAGCCCGTGCGCTGGGTCGTCAGGCCCAGGGTGGCCCTGTTCAACAC 99914

TeratornLC199500 **CTCCGCCATGAACAGCCTCTCCAACGAGATCGCGCAGATCGGAAGCGAGCTCCTCGCCCGCTTCGGAGACAGCGAGCACGCCAAGACCGTGACCCTCAACCTCACCTTTCTCAGCAAAGC** 97183
WF-Teratorn CTCCGCCATGAACAGCCTCTCCAACGAGATCGCGCAGATCGGAAGCGAGCTCCTCGCCCGCTTCGGAGACAGCGAGCACGCCAAGACCGTGACCCTCAACCTCACCTTTCTCAGCAAAGC 100034

TeratornLC199500 **CCTGGATTACCGCGGTGGTGCGCAGCGCATGCTGGAGATGAGAGGCAAGATTCTCGCGAACCAGCCCATCCGTGACTTAGAAGTGGATGGGTGGACGAGGGGCCGCCTGTTCTGGTCCTG** 97303
WF-Teratorn CCTGGATTACCGCGGTGGTGCGCAGCGCATGCTGGAGATGAGAGGCAAGATTCTCGCGAACCAGCCCATCCGTGACTTAGAAGTGGATGGGTGGACGAGGGGCCGCCTGTTCTGGTCCTG 100154

TeratornLC199500 **GATGGCGGACCCCATATGCGCCGTGTACATGCGAAAGGCGCTCGGGGCCAAGGAGATCGGCGCCAATGCGCCTGATTTCACCGTGGAGGAACGGGAGGTCCTGGTGCAGCTCGAGGGGTT** 97423
WF-Teratorn GATGGCGGACCCCATATGCGCCGTGTACATGCGAAAGGCGCTCGGGGCCAAGGAGATCGGCGCCAATGCGCCTGATTTCACCGTGGAGGAACGGGAGGTCCTGGTGCAGCTCGAGGGGTT 100274

TeratornLC199500 **GATAACCGGAGTGCTGGCACTGTTTAGAAACGTGGTGGGGGACAGGGCGAACTCTGTCGTGGCCTCCATGCCCTGCTGCTTCAGCCACCGGCGCTACGCGGCCATGGATGGGTCCCAGGC** 97543
WF-Teratorn GATAACCGGAGTGCTGGCACTGTTTAGAAACGTGGTGGGGGACAGGGCGAACTCTGTCGTGGCCTCCATGCCCTGCTGCTTCAGCCACCGGCGCTACGCGGCCATGGATGGGTCCCAGGC 100394

TeratornLC199500 **CACGGTCAACGAACGCGGATATCCCGTCTGGTTCACCTTGGACGACATGGACTACTGCAACCTGATGTATTGGATCATCCTACCGAGCCTGGGGGCGAGGGTGTATCGCCGGCAACCCGT** 97663
WF-Teratorn CACGGTCAACGAACGCGGATATCCCGTCTGGTTCACCTTGGACGACATGGACTACTGCAACCTGATGTATTGGATCATCCTACCGAGCCTGGGGGCGAGGGTGTATCGCCGGCAACCCGT 100514

TeratornLC199500 **CGTCGAAAACGAAGCCCACGCGGTGCGCGACCCGCTGAGGTACAGGGACGGGGAGCCGAACCTCTCTCTCGAGCTGGGCATCGACGGCAACCTAGGATTGACCGTCGCCCGTCTGACCGG** 97783
WF-Teratorn CGTCGAAAACGAAGCCCACGCGGTGCGCGACCCGCTGAGGTACAGGGACGGGGAGCCGAACCTCTCTCTCGAGCTGGGCATCGACGGCAACCTAGGATTGACCGTCGCCCGTCTGACCGG 100634

TeratornLC199500 **GGAATGCGTCTTCTCCAGGGTGGCGCCCGAGGCCTTCGCCGAAGACGCCAGCGAGCAGTTCGAGCCCACGGCCTCTCGCCACGTCTCCTGCACGGACCTGGGATTCCACAGGAACTCTGT** 97903
WF-Teratorn GGAATGCGTCTTCTCCAGGGTGGCGCCCGAGGCCTTCGCCGAAGACGCCAGCGAGCAGTTCGAGCCCACGGCCTCTCGCCACGTCTCCTGCACGGACCTGGGATTCCACAGGAACTCTGT 100754

TeratornLC199500 **GGTCACGTACATGTGGTCGATAAGACTCAGGAAGATCTCCGGCATCGGAAACGACCTGGCCAAAGCGCTCCTCGGCATGCACTACAGCACCCTGCTCACTCACGACGTGGTCAGGGATCA** 98023
WF-Teratorn GGTCACGTACATGTGGTCGATAAGACTCAGGAAGATCTCCGGCATCGGAAACGACCTGGCCAAAGCGCTCCTCGGCATGCACTACAGCACCCTGCTCACTCACGACGTGGTCAGGGATCA 100874

TeratornLC199500 **CTACGACACCGTCTACTACTCCGGCTTGTCCTACCTGATGTTCAGGGGAGTGAGGTTCACGGGCTGCGGAGTGTCCCTGATGCAGCAAAAGTCGGCTCTGTACGCGCTGGGCACGCAGGT** 98143
WF-Teratorn CTACGACACCGTCTACTACTCCGGCTTGTCCTACCTGATGTTCAGGGGAGTGAGGTTCACGGGCTGCGGAGTGTCCCTGATGCAGCAAAAGTCGGCTCTGTACGCGCTGGGCACGCAGGT 100994

TeratornLC199500 **GCTGTGCAACCCGACGGCGAACAACACTCACCAGGTGTGCACCGTGAACCAGTACTGCGAGTCGGTGGTGATACCGGACACCCTGGAGTCGCCCGGGCTTTACATCCCCAACCTGTACAT** 98263
WF-Teratorn GCTGTGCAACCCGACGGCGAACAACACTCACCAGGTGTGCACCGTGAACCAGTACTGCGAGTCGGTGGTGATACCGGACACCCTGGAGTCGCCCGGGCTTTACATCCCCAACCTGTACAT 101114

TeratornLC199500 **GAAAGACGTGCGAGGGGGAGACGTGCACCTGGAGGGGCGCGGAGGGCCGTACGACATGGTGATCGCGGACGCCTTCAACGGCTTCTGCCCGGAGTCCGACAGCGCTTGCGGCTACCGCAG** 98383
WF-Teratorn GAAAGACGTGCGAGGGGGAGACGTGCACCTGGAGGGGCGCGGAGGGCCGTACGACATGGTGATCGCGGACGCCTTCAACGGCTTCTGCCCGGAGTCCGACAGCGCTTGCGGCTACCGCAG 101234

TeratornLC199500 **GCCGTACATCTTCACCACCGGCCGGTCCCAGAGCCTGCAGGGACTCACGGGCGAGGTGCCCTTGGATCCGCTCATGCGCTCGGAGGCCTACACCTGCATGCCCACTCTGCTCAGGCCCTT** 98503
WF-Teratorn GCCGTACATCTTCACCACCGGCCGGTCCCAGAGCCTGCAGGGACTCACGGGCGAGGTGCCCTTGGATCCGCTCATGCGCTCGGAGGCCTACACCTGCATGCCCACTCTGCTCAGGCCCTT 101354

TeratornLC199500 **CAGCTCGGCGCTCAGGTCGCGAGGCAGACGCAGAGGCGACGGCGGAGAGGACAGCAACCACAGCGTCTTCTGCGACGCCCTGCACGCGGCGGAGTTCGAAAAGAGCGTGCTGGATCACGT** 98623
WF-Teratorn CAGCTCGGCGCTCAGGTCGCGAGGCAGACGCAGAGGCGACGGCGGAGAGGACAGCAACCACAGCGTCTTCTGCGACGCCCTGCACGCGGCGGAGTTCGAAAAGAGCGTGCTGGATCACGT 101474

TeratornLC199500 **GGGGGACCTCTTCAAGGACACCACGGAGAGAGCGAATGGGAACAAGCGGGCCACGGACATCATGCAGCCCTCGCTCGGGGTGGCTTTCTGCGGTACTTACAGCATCGGGAAGACCTCGGA** 98743
WF-Teratorn GGGGGACCTCTTCAAGGACACCACGGAGAGAGCGAATGGGAACAAGCGGGCCACGGACATCATGCAGCCCTCGCTCGGGGTGGCTTTCTGCGGTACTTACAGCATCGGGAAGACCTCGGA 101594

TeratornLC199500 **CAGGAAGCTCATGAGCGCCTGCGCCGCGACGGGCGGCGAGCCCCTCGTCGACAGGCTCGCGCCCAGAATCAGCGGCACGGGCCTGTTCGCTTGCAACCCCATGCAGACCACCGCTCGGTT** 98863
WF-Teratorn CAGGAAGCTCATGAGCGCCTGCGCCGCGACGGGCGGCGAGCCCCTCGTCGACAGGCTCGCGCCCAGAATCAGCGGCACGGGCCTGTTCGCTTGCAACCCCATGCAGACCACCGCTCGGTT 101714

TeratornLC199500 **GATAGCTCCCATATTCAGCAGGATATTCGAGAGCCCCAACCATGGTCGCAGGCTGTTCTTGTGA**GGGAGGAAACGGAACGTAAAACGGGCGAAGCGCGCGGGAGAAAGGAGAGAGAAGAG 98983
WF-Teratorn GATAGCTCCCATATTCAGCAGGATATTCGAGAGCCCCAACCATGGTCGCAGGCTGTTCTTGTGAGGGAGGAAACGGAACGTAAAACGGGCGAAGCGCGCGGGAGAAAGGAGAGAGAAGAG 101834

TeratornLC199500 AAAGCAGGGAGGAAGG**ATGCCTCGGGTCAGGGGGCCCGGCTTAGCGCGCAAGGTGGCCCGCGGACTCTGGGCCTTTGCTGAATTGGTGCTTTTGTGCGGGGTCAATTTCTTCGTGACCCT** 99103
WF-Teratorn AAAGCAGGGAGGAAGGATGCCTCGGGTCAGGGGGCCCGGCTTAGCGCGCAAGGTGGCCCGCGGACTCTGGGCCTTTGCTGAATTGGTGCTTTTGTGCGGGGTCAATTTCTTCGTGACCCT 101954

TeratornLC199500 **GTACATGCACCCAGTTGACCGCGAGCGCGCCATCGCCTTGATTAGGAAGAGGGAGGAGGCCGAAGTGGACGCCGCCATCAACACAGTGGCCAGGTCCATCACGGGCCTGGACGCGAACAT** 99223
WF-Teratorn GTACATGCACCCAGTTGACCGCGAGCGCGCCATCGCCTTGATTAGGAAGAGGGAGGAGGCCGAAGTGGACGCCGCCATCAACACAGTGGCCAGGTCCATCACGGGCCTGGACGCGAACAT 102074

TeratornLC199500 **AGAAATGATCAAGGGGATCATCGGCTCCGAGTCATGTGACAGGGCCGTGACCGCCGAGTACGTGAACGAGCTGCGCTCAGCCGAGACTCAGAGGGACCAGCTCCGCTACATGCTCATGGC** 99343
WF-Teratorn AGAAATGATCAAGGGGATCATCGGCTCCGAGTCATGTGACAGGGCCGTGACCGCCGAGTACGTGAACGAGCTGCGCTCGGCCGAGACTCAGAGGGACCAGCTCCGCTACATGCTCATGGC 102194

TeratornLC199500 **GCTCAGGGGCCGGATCGATGCTGTCAGGCTGGTGGGCGCGGTGGACACTCTGTCAGACAGCGTGGGGCTTCAACTCAACGCTTCGGCCAAGATAAACAGGACGGTGCTCGCCTTGAACGT** 99463
WF-Teratorn GCTCAGGGGCCGGATCGATGCTGTCAGGCTGGTGGGCGCGGTGGACACTCTGTCAGACAGCGTGGGGCTTCAACTCAACGCTTCGGCCAAGATAAACAGGACGGTGCTCGCCTTGAACGT 102314

TeratornLC199500 **CCACAGGGACCTCGACGACATGAGGGAACCGGTGTCTAAAATGACGGTGGAAGATGCCAGCGGGGCTTACGAAGACGGCCATCTTACTCAGAGGCACCCTCACTGGCTCGTGTGA**CCGCC 99583
WF-Teratorn CCACAGGGACCTCGACGACATGAGGGAACCGGTGTCTAAAATGACGGTGGAAGATGCCAGCGGGGCTTACGAAGACGGCCATCTTACTCAGAGGCACCCTCACTGGCTCGTGTGACCGCC 102434

TeratornLC199500 CTTAAACTGGGTAGTCCCGCTACGGTTCTCTGATGGGGAACGCATATAAATTCCTACCGGAAGGTCTGGGTTGTTGGGGTTTCAGGGCTTTTCTTTTCTTTTCTTTCTTAGTGTACCTGC 99703
WF-Teratorn CTTAAACTGGGTAGTCCCGCTACGGTTCTCTGATGGGGAACGCATATAAATTCCTACCGGAAGGTCTGGGTTGTTGGGGTTTCAGGGCTTTTCTTTTCTTTTCTTTCTTAGTGTACCTGC 102554

TeratornLC199500 CCTTTGTATAAAGTGTGTCTCTTTTGCTGTGCTGCCTTTTTTGTTTTGTTTTCAAATGCAACAGTGTTTGACTGTCTGTGTGTGTTGATCTTGTAACAAATTCTGAATAAAATAATTTCT 99823
WF-Teratorn CCTTTGTATAAAGTGTGTCTCTTTT---------------TTGTTTTGTTTTCAAATGCAACAGTGTTTGACTGTCTGTGTGTGTTGATCTTGTAACAAATTCTGAATAAAATAATTTCT 102659

TeratornLC199500 GAAAACACAACTGCTTGGCTTGTTTCCATATTCTGTTTGCCAAGAAATGTAGAGAGAAGATGTGAGACTACAGTAGGCGTACTTCAATCAACAGCAATATGATTTTATTCAAAGTACAAA 99943
WF-Teratorn GAAAACACAACTGCTTGGCTTGTTTCCATATTCTGTTTGCCAAGAAATGTAGAGAGAAGATGTGAGACTACAGTAGGCGTACTTCAATCAACAGCAATATGATTTTATTCAAAGTACAAA 102779

TeratornLC199500 AGGAAAATAAGTCTTACATTTCAAGAGGGTACAGTTCATGTGATTGGATTGCATGCTGAGGGTACAGTTCATGTGATTGGATTGCATGCTATGAAAAAGAAATGCACGAGGTAGAAAGAC 100063
WF-Teratorn AGGAAAATAAGTCTTACATTTCAAGAGGGTACAGTTCATGTGATTGGATTGCATGCTGAGGGTACAGTTCATGTGATTGGATTGCATGCTATGAAAAAGAAATGCACGAGGTAGAAAGAC 102899

TeratornLC199500 AACTACGCAAGCCTTCGCTGTTTCAATGGCAGTCACAAGGAATAACATCTGCTTGTCGAAAGAAACCTCTGAGATAACTGTGGTAAAGAAAACAAATTGTAAAAGAGTAAAGCAAAACA 100183
WF-Teratorn AACTACGCAAGCCTTCGCTGTTTCAATGGCAGTCACAAGGAATAACATCTGCTTGTCGAAAGAAACCTCTGAGATAACTGTGGTAAAGAAAACAAATTGTAAAAGAGTAAAGCAAAACA 103019

TeratornLC199500 CTTCGATGACAATTCCTATGCAATCATGTTCCACACACCTGTTTTTTTCAAAAAAAAAATATAATATAAAAAAATGATGATGAAATAATGTGAACATTGTGAGCAAAAATGT**TCATACGT** 100303
WF-Teratorn CTTCGATGACAATTCCTATGCAATCATGTTCCACACACCTGTTTTTTTTTAAAAAAAAATATAATACAAAAAAATGATGATGAAATAATGTGAACATTGTGAGCAAAAATGTTCATACGT 103139

TeratornLC199500 **AGACATTTTTGTCACCTTCGTAGCAAATGGGTCGGGTCAGCGTATTTCCAGTCTCCGAGTTACGGTGGCAGTCGCTGAACTCCTCTGCCACGTTCATGTAAAGCGTCTGGGATTGGCAGC** 100423
WF-Teratorn AGACATTTTTGTCACCTTCGTAGCAAATGGGTCGGGTCAGCGTATTTCCAGTCTCCGAGTTACGGTGGCAGTCGCTGAACTCCTCTGCCACGTTCATGTAAAGCGTCTGGGATTGGCAGC 103259

TeratornLC199500 **AAGACAGCGGCCTGTAGTCGAACAGAGCGGCCGCAGCTGCGTTTTTAGCCGAAAACACATCTGCGTCCAAGGCGAAGGGTCGTACCCTTTTGTGTACGTAGCTGGGATCCGCTTTCTCCA** 100543
WF-Teratorn AAGACAGCGGCCTGTAGTCGAACAGAGCGGCCGCAGCTGCGTTTTTAGCCGAAAACACATCTGCGTCCAAGGCGAAGGGTCGTACCCTTTTGTGTACGTAGCTGGGATCCGCTTTCTCCA 103379

TeratornLC199500 **TTCCGCTTCGGGCGGCAAGCTCAGCGATGGCTTGAACGATGGTGGGTCCATCGAGCGGCTTGCAGTCCAAGACCACGTGGATGTCCCTTTCAACGCGCCCGCCGCCTTCCCTTGAGCGGT** 100663
WF-Teratorn TTCCGCTTCGGGCGGCAAGCTCAGCGATGGCTTGAACGATGGTGGGTCCATCGAGCGGCTTGCAGTCCAAGACCACGTGGATGTCCCTTTCAACGCGCCCGCCGCCTTCCCTTGAGCGGT 103499

TeratornLC199500 **ACTGCACGGTGAGCCACGTGCGCAGGAGGAAAATCATCCTCTTGAACTTGACCTCGGACACCGTCTCCCTCAAAGAACTGTCGATCAGTGGGAGGACTCTCATGCTCCCACCCTGGCACA** 100783
WF-Teratorn ACTGCACGGTGAACCACGTGCGCAGGAGGAAAATCATCCTCTTGAACTTGACCTCGGACACTGTCTCCCTCAAAGAACTGTCGATCAGTGGGAGGACTCTCATGCTCCCACCCTGGCACA 103619

TeratornLC199500 **CTCTGGTGGTGCACCACGCTGCCAAGACGATCTGTTCCTTGAACACGAAACAGCACCAGTGAAAGTAATGCGTGCCCATGTCCACGTCGAACAGCCGCGGCAGGTGCACGTCGGGCCCGA** 100903
WF-Teratorn CTCTGGTGGTGCACCACGCTGCCAAGACGATCTGTTCCTTGAACACGAAACAGCACCAGTGAAAGTAATGCGTGCCCATGTCCACGTCGAACAGCCGCGGCAGGTGCACGTCGGGCCCGA 103739

TeratornLC199500 **TGGCCGCCTGGGTGCTCTCGGGGCACCATTTCCCCACTAGCACGCGGTCCCAAAACCCCGGGATGTGAGGCATGCAAACGTATCTGTAGGCATCATACTCAGGCAAGAGCTTATTTTCCC** 101023
WF-Teratorn TGGCCGCCTGGGTGCTCTCGGGGCACCATTTCCCCACTAGCACGCGGTCCCAAAACCCCGGGATGTGAGGCATGCAAACGTATCTGTAGGCATCATACTCAGGCAAGAGCTTATTTTCCC 103859

TeratornLC199500 **AGTAGGTGAAAGACATGTCGTTTGCTGGATTCTTTATCAGGGTCTGGAACGCTGCGTAAAAAAAAAAAAGAGAGAGAGAAATCAGAGAGAGGAAAGTCAGACTCTTGAAGCTGTGCAA**GG 101143
WF-Teratorn AGTAGGTGAAAGACATGTCGTTTGCTGGATTCTTTATCAGGGTCTGGAACGCTGCGTAAAAAAAAAAAAGAGAGAGAGAAATCAGAGAGAGGAAAGTCAGACTCTTGAAGCTGTGCAAGG 103979

TeratornLC199500 TCGAGGTG**CTAGAGGGTGTGCCTCTTACCTCGCTGAAGTTCCTCGTCACTTGGCGACCCTTCGGCCAACATCCTTTCAACGCATTCCTCTTTTTCGATCCTCGTCATGAAGCGGTCGCTA** 101263
WF-Teratorn TCGAGGTGCTAGAGGGTGTGCCTCTTACCTCGCTGAAGTTCCTCGTCACTTGGCGACCCTTCGGCCAACATCCTTTCAACGCATTCCTCTTTTTCGATCCTCGTCATGAAGCGGTCGCTA 104099

TeratornLC199500 **CCTTCTCGCTCCAGTTCCCAAGCTCCTACAGCCTCCATCTCCGGCATGTCAGCCAGCGCGTTTGCCGTGGCGTTCACCATTCCGTTCTCGGAGACGACCCAGTGCTTTCGGCAGCCTCCC** 101383
WF-Teratorn CCTTCTCGCTCCAGTTCCCAAGCTCCTACAGCCTCCATCTCCGGCATGTCAGCCAGCGCGTTTGCCGTGGCGTTCACCATTCCGTTCTCGGAGACGACCCAGTGCTTTCGGCAGCCTCCC 104219

TeratornLC199500 **TGCGCTCTGATGCACAGCGTGCACAGGACGCATCCGCAGAGGCACTGCACGGGTCGATTGAGTTGCAGGAGATACCCGCAGCACACCGGCGCACGGTTGAAAACCCGTCTGCATGATGGG** 101503
WF-Teratorn TGCGCTCTGATGCACAGCGTGCACAGGACGCATCCGCAGAGGCACTGCACGGGTCGATTGAGTTGCAGGAGATACCCGCAGCACACCGGCGCACGGTTGAAAACCCGTCTGCATGATGGG 104339

TeratornLC199500 **CAAGTGATGAGATCTTCGAGCGTGGATAGGGAGACGCCGGGGATGCCGTCCCAAGGGCGCGGCCCCGCGCTGCCCTCTGCGCTCCAGCACGCAAGCTCCTCTTCTGCGGGCGTCCTGGGC** 101623
WF-Teratorn CAAGTGATGAGATCTTCGAGCGTGGATAGGGAGACGCCGGGGATGCCGTCCCAAGGGCGCGGCCCCGCGCTGCCCTTTGCGCTCCAGCACGCAAGCTCCTCTTCTGCGGGCGTCCTGGGC 104459

TeratornLC199500 **TCGCGTCGGTTGGACTCCACGGAGAGACAGAGCCCTGGAGACTCACTCTGCAGCGCTGCGCCGGGCTCGTCTTTCGGCGGCTCGACGTCAGGCTGCGAGCTCGGGCGTCCGTCCCC-TCC** 101742
WF-Teratorn TCGCGTCGGTTGGACTCCACGGAGAGACAGAGCCCTGGAGACTCACTCTGCAGCGCTGCGCCGGGCTCGTTTTTCGGCGGCTCGACGTCAGGCTGCGAGCTCGGGCGTCCGTCCCCCTCC 104579

TeratornLC199500 **ATGGTCGAGATCTTCGTCACCCGTCTCGGGTGGCGGAAGTAGAGAGCCTGGAAGCTCAATCGTCTCCGCGCAAGTCGAAGCGACCCGAGGAGGAACTTGGTAACCGTTTGGGGGTTCCGC** 101862<br>
WF-Teratorn ATGGTCGAGATCTTCGTCACCCGTCTCGGGTGGCGGAAGTAGAGAGCCTGGAAGCTCAATCGTCTCCACGCAAGTCGAAGCGACCCGAGGAGGAACTTGGTAACCGTTTGGGGGTTCCGC 104699

TeratornLC199500 **CGCGCGCGGCTCGATGCAATCGGGGTACGTGATCAAGTTGATGTCGTCGTACGCTTTCGTGATGACGCTCTCGCACAGCATGGCGCTGGCTTCTTCCTCCAGCGTACCGTACTCCTCGTC** 101982<br>
WF-Teratorn CGCGCGCGGCTCGATGCAATCGGGGTACGTGATCAAGTTGATGTCGTCGTACGCTTTCGTGATGACGCTCTCGCACAGCATGGCGCTGGCTTCTTCCTCCAGCGTACCGTACTCCTCGTC 104819

TeratornLC199500 **CTCAGTCGAAGTCAA**ATAATATCTGGAGGGGTTTCTGTAACGCACCCACAGTGACACAAAAGACAACTTGTGAATATCACCT**TCAAACCATGCCTCCCAGGATCGACGGGGATACCCACG** 102102<br>
WF-Teratorn CTCAGTCGAAGTCAAATAATATCTGGAGGGGTTTCTGTAACGCACCCACAGTGACACAAAAGACAACTTGTGAATATCACCTTCAAACCATGCCTCCCAGGATCGACGGGGATACCCACG 104939

TeratornLC199500 **GCTGGCCGCTGATGCTCCAGATGGCAGTGTGCAGCTCTTCGTCCATTCGCTTCGCGACCTCGACGCCTTCCTCGGACTCCGGGACGTCGATCTCCCCCGGGCGGAGCGAGCGTTCGCTCT** 102222<br>
WF-Teratorn GCTGGCCGCTGATGCTCCAGATGGCAGTGTGCAGCTCTTCGTCCATTCGCTTCGCGACCTCGACGCCTTCCTCGGACTCCGGGACGTCGATCTCCCCCGGGCGGAGCGAGCGTTCGCTCT 105059

TeratornLC199500 **CCGCGCCTCGGTCTTGCTCCGTGGCGTCCACTGGGCTAGGCGGAGGCGCCGGAGTGGGCTCGACGTGGTTTGCTTTCCGCAGGGCTTCGAGTCCAACGCAGATTGCGCCTCCCGGTACCG** 102342<br>
WF-Teratorn CCGCGCCTCGGTCTTGCTCCGTGGCGTCCACTGGGCTAGGCGGAGGCGCCGGAGTGGGCTCGACGTGGTTTGCTTTCCGCAGGGCTTCGAGTCCAACGCAGATTGCGCCTCCCGGTACCG 105179

TeratornLC199500 **GCTGTCGCCCTCCATCCGCGAGGGCCTCGCGACACCCCGCGCTTTCAGCACCAGGACACAGAGGCTCGCCGCGAGCCTCTTCCGCCTCGTCTTCGGTTTCCTCCGGAGAGGCCGCCTCGT** 102462<br>
WF-Teratorn GCTGTCGCCCTCCATCCGCGAGGGCCTCGCGACACCCCGCGCTTTCAGCACCAGGACACAGAGGCTCGCCGCGAGCCTCTTCCGCCTCGTCTTCGGTTTCCTCCGGAGAGGCCGCCTCGT 105299

TeratornLC199500 **TTTCTACGCGCCCCCGACTCTTGTCGTCTTCGTCCGAGGAGGAGGCGCTGACATCAGATGGACTGCACGCGCGCTCTTTGGCCTCGTACTCGGCCGTGCGAATTTCGCCCTTCTCTTCTT** 102582<br>
WF-Teratorn TTTCTACGCGCCCCCGACTCTTGTCGTCTTCGTCCGAGGAGGAGGCGCTGACATCAGATGGACTGCACGCGCGCTCTTTGGCCTCGTACTCGGACGTGCGAATTTCGCCCTTCTCTTCTT 105419

TeratornLC199500 **CGCCCTCGCTGTACTCGCCTTCGCTGGAGCTGTCTTCCTTTGTCGAACAACACCCTAGTCTCCAGGGCCTGGTCGCGGTGTCTCGCTCCGTGCGACGGCCTCTCCGCGAGGACGACGATT** 102702<br>
WF-Teratorn CGCCCTCGCTGTACTCGCCTTCGCTGGAGCTGTCTTCCTTTGTCGAACAACACCCTAGTCTCCAGGGCCTGGTCGCGGTGTCTCGCTCCGTGCGACGGCCTCTCCGCGAGGACGACGATT 105539

TeratornLC199500 **GGGGGTTCGCGCGGTCAAAATTGCAACCGGCGTGAGGCACAGCCCAGCGAACTGCCTTTCGCCTGCCGTCGTGAGCTTCACCGACATCCACGCCAAAGTGCGACGGTTCTTCACCAGTGG** 102822<br>
WF-Teratorn GGGGGTTCGCGCGGTCAAAATTGCAACCGGCGTGAGGCACAGCCCAGCGAACTGCCTTTCGCCTGCCGTCGTGAGCTTCGCCGACATCCACGCCAAAGTGCGACGGTTCTTCACCAGTGG 105659

TeratornLC199500 **GCCCCTCTGGCTCGGTCCTCCTTTTCACGAGCCGAATCTCGGATTTAGTCCTGCGAGGGCTGAGGCTCGGGTCCCGCTTGCGCTTGATCCTCTCGCTCCGGTGATATTTGCCGTCTCCTT** 102942<br>
WF-Teratorn GCCCCTCTGGCTCGGTCCTCCTTTTCACGAGCCGAATCTCGGATTTAGTCCTGCGAGGGCTGAGGCTCGGGTCCCGCTTGCGCTTGATCCTCTCGCTCCGGTGATATTTGCCGTCTCCTT 105779

TeratornLC199500 **TTCGCACGCAACCGCCCGCAGGGTTTGTCCTTCTGCGACGAGGTGCAAATTCAGGGTCGAGCTCCTTTCCAACCTGCCTCTTGAGTGTCGGGTAGAGGGGAGGCTCGCGGTCGAGACGGA** 103062<br>
WF-Teratorn TTCGCACGCAACCGCCCGCAGGGTTTGCCCTTCTGCGACGAGGTGCAAATTCAGGGTCGAGCTCCTTTCCAACCTGCCTCTTGAGTGTCGGGTAGAGGGGAGGCTCGCGGTCGAGACGGA 105899

TeratornLC199500 **CGCATCCGAGGGCCCGCTCGCGTCCGCTTCCTCGGTAACGATCGCGGACTCCATCCCTCTGGTCGCGGCGGGAGATCTCGCCCTCGGCCCACTCGGGCGTCCAGTCGAAGCGCTTTCTTT** 103182<br>
WF-Teratorn CGCATCCGAGGGCCCGCTCGCGTCCGCTTCCTCGGTAACGATCGCGGACTCCATCCCTCTGGTCGCGGCGGGAGATCTCGCCCTCGGCCCACTCGGGCGTCCAGTCGAAGCGCTTTCTTT 106019

TeratornLC199500 **CTCTCCTTCTCTTCTCCGTTTCTTGCCTTCTCCTTTGCTGCAGGTTGCGCAGGGGAGAAGGCGTAGTGTATCTGCGTTGGCCCCCGCCGATCCTTCTTTGTTCGGGCCTGGCCTTCCTCT** 103302<br>
WF-Teratorn CTCTCCTTCTCTTCTCCGTTTCGTGCCTTCTCCTTTGCTGCAGGTTGCGCAGGGGAGAAGGCGTAGTGTATCTGCGTTGGCCCCCGCCGATCCTTCTTTGTTCGGGCCTGGCCTTCCTCT 106139

TeratornLC199500 **GCATCTTTTCGTTCTCGTATCCAGCCGAAATCACCCTCAT**TGCACCGCGGGCTTCGTTAGAACAGAGAGCCGAGTTCCACCCAGATGCCGAGGGGCGAATCTCTAGTGTTCCTGTACGAA 103422<br>
WF-Teratorn GCATCTTTTCGTTCTCGTATCCAGCCGAAATCACCCTCATTGCACCGCGGGCTTCGTTAGAACAGAGAGCCGAGTTCCACCCAGATGCCGAGGGGCGAATCTCTAGTGTTCCTGTACGAA 106259

TeratornLC199500 CGAACGCTGGGGTGTGAGCTCAGTCCTCTGTGAGCAAACTTGGCAGCACTTATATAGCCCCTCCTCGACAGTCTCGAAATCTCTGCAACAGAAGAAACTCTCACCTTCCTCACTTTCATG 103542<br>
WF-Teratorn CGAACGCTGGGGTGTGAGCTCAGTCCTCTGTGAGCAAACTTGGCAGCACTTATATAGCCCCTCCTCGACAGTCTCGAAATCTCTGCAACAGAAGAAACTCTCACCTTCCTCACTTTCATG 106379

TeratornLC199500 CCAAAACAAAACTCATCACTATCAGTGAGTCTGACCTTCCCCCAGAGTGCAAGAACTCCGCTCGCAGCAAGACGTCTCTGGGTTGGACGAGATCCGTCAGAGTCAGGCCGATGTTTACAG 103662<br>
WF-Teratorn CCAAAACAAAACTCATCACTATCAGTGAGTCTGACCTTCCCCCAGAGTGCAAGAACTCCGCTCGCAGCAAGACGTCTCTGGGTTGGACGAGATCCGTCAGAGTCAGGCCGATGTTTACAG 106499

TeratornLC199500 CGTCTGACCGATGTGAACAGCGTCCGTCCGACAGCCACTCGCCTCCTCCGACCAGGCTTAAAAACGGCCGAGTGCAGTTCATAGAATACGACCGATGCTCGCAGTCACTGTCCTCCCTCC 103782<br>
WF-Teratorn CGTCTGACCGATGTGAACAGCGTCCGTCCGACAGCCACTCGCCTCCTCCGACCAGGCTTAAAAACGGCCGAGTGCAGTTCATAGAATACGACCGATGCTCGCAGTCACTGTCCTCCCTCC 106619

TeratornLC199500 TCGACCTTCCCGCGAAGCTCAAAAAGTCAGACCGGCATCTCGCACGCCAAGCATCCACGGTGGCATCAATCCGCTGTTAATCGGAGCAGCCCAGACATTCGAAACACAAAAAGTTCTGGG 103902<br>
WF-Teratorn TCGACCTTCCCGCGAAGCTCAAAAAGTCAGACCGGCATCTCGCACGCCAAGCATCCACGGTGGCATCAATCCGCTGTTAATCGGAGCAGCCCAGACATTCGAAACACAAAAAGTTCTGGG 106739

TeratornLC199500 CTCAGAAAAGAAAACTGGTAGGCTCAGGGGAAGGTAGATAATGTTTGGGAGCAATAGGAAAACAGGAGAGGCAGGGAGTAAGACATGAAGGTGGAGACTTGGGAGGAGCAGTGCTCAGGG 104022<br>
WF-Teratorn CTCAGAAAAGAAAACTGGTAGGCTCAGGGGAAGGTAGAGAATGTTTGGGAGCAATAGGAAAACAGGAGAGGCAGGGAGTAAGACATGAAGGTGGAGACTTGGGAGGAGCAGTGCTCAGGG 106859

| | | |
|---|---|---|
| TeratornLC199500 | GAAGGTGGAGAATGTTTGGGACCAATAGGAAAAAAGGAGAGGCAGGGAGTAAAAGATGAAGGTGGAGACTTGGGAGGAGCAGTGGAAGCCCATGCTCTATGAAGGTGGGGGGAGAAAAGA | 104142 |
| WF-Teratorn | GAAGGTGGAGAATGTTTGGGACCAATAGGAAAAAAGGAGAGGCAGGGAGTAAAAGATGAAGGTGGAGACTTGGGAGGAGCAGTGGAAGCCCATGCTCTATGAAGGTGGGGGGAGAAAAGA | 106979 |
| TeratornLC199500 | GAGCAGAGTGGTGTTGGTGACCATAAAATTACTTCTGATCAGCGCAACTGCTTGTGCACGCAGATGTTTTTTTTT----GTTTTGTTTTTTTTGTTTTGTCAATTCACGCTTGCAGAAGC | 104258 |
| WF-Teratorn | GAGCAGAGTGGTGTTGGTGACCATAAAATTACTTCTGATCAGCGCAACTGCTTGTGCACGCAGATGTTTTTTTTTTTTTGTTTTGTTTGTTTTGTTTTGTCAATTCACGCTTGCAGAAGC | 107099 |
| TeratornLC199500 | CAAGCTTACAAACCTCTCTCTTTCCCACTCCGGAGATCGGCAGGAATAAAGCACAGGGTGCACTGCCGTGAACGTGTGAACACATTTTTTGG---------------------------- | 104350 |
| WF-Teratorn | CAGGCTTACAAACCTCTCTCTTTCCCACTCCGGAGATCGGCAGGAATAAAGCACAGGGTGCACTGCCGTGAACGTGTGAACACATTTTTTGGGTTTTTTTTTTTGACAAAGAAAATGTG | 107219 |
| TeratornLC199500 | ------------------------------------------------------------------------------------------------------------------------ | 104350 |
| WF-Teratorn | TAGTGTATTTTCCCAGAGATGAAGCGCAAGCTGTGTTCGGTGGCAAAACACTTCAAAGCAACGAGAGCATTTTTGAACGAGCTCCTTTATTTCGCACTCAAACGCACAGTGTCCATCTCA | 107339 |
| TeratornLC199500 | ------------------------------------------------------------------------------------------------------------------------ | 104350 |
| WF-Teratorn | GAGCGCACCGAGCTTTTCAAACAGTTTCACGACAAGATACATCTGCTTATATCCAAAGGATGGTGCACATGCAACACAGACAAAATGAAATACACATGAAAACCCACTACGCAAACTACA | 107459 |
| TeratornLC199500 | ------------------------------------------------------------------------------------------------------------------------ | 104350 |
| WF-Teratorn | TTTCTCTTTCCCACTCCGGAGATCGGCAGGAATAAAGCACAGGGTGCACTGCCGTGAACGTGTGAACACATTTTTTGGTTGTTTTTTTTTTGACAAAGAAAATGTGTAGTGTATTTTCCC | 107579 |
| TeratornLC199500 | ------------------------------------------------------------------------------------------------------------------------ | 104350 |
| WF-Teratorn | AGAGATGAAGCGCAAGCTGTGTTCGGTGGCAAAACACTTCAAAGCAACGAGAGCATTTTTGAACGAGCTCCTTTATTTCGCACTCAAACGCACAGTGTCCATCTCAGAGAGCACCGAGCT | 107699 |
| TeratornLC199500 | ------------------------------------------------------------------------------------------------------------------------ | 104350 |
| WF-Teratorn | TTTCAAACAGTTTCACGACAAGATAGATCTGCTTCTATCCAAAGGATGGTGCACATGCAACACAGACAAAATGAAAAAAAAAAAATACACATTCGATCCAACTACACAAACTACATTTCTC | 107819 |
| TeratornLC199500 | -----------------------------------------------------------------------------TTTTTTTTTTGACAAAGAAAATGTGTAGTGTATTTTCCCAGAG | 104393 |
| WF-Teratorn | TTTCCCACTCCGGAGATCGGCAGGAATAAAGCACAGGGTGCACTGCCGTGAACGTGTGAACACATTTTTTGGGTTTTTTTTTTTTTTGACAAAGAAAATGTGTAGTGTATTTTCCCAGAG | 107939 |
| TeratornLC199500 | ATGAAGCGCAAGCTGTGTTCGGTGGCAAAACACTTCAAAGCAACGAGAGCATTTTTGAACGAGCTCCTTTATTTCGCACTCAAACGCACAGTGTCCATCTCAGAGCGCACCGAGCTTTTC | 104513 |
| WF-Teratorn | ATGAAGCGCAAGCTGTGTTCGGTGGCAAAACACTTCAAAGCAACGAGAGCATTTTTGAACGAGCTCCTTTATTTCGCACTCAAACGCACAGTGTCCATCTCAGAGAGCACCGAGCTTTTC | 108059 |
| TeratornLC199500 | AAACAGTTTCACGACAAGATACATCTGCTTCTATCCAAAGGATGGTGCACATGCAACACAGACAAAATGAAAAAAAAAA-TACACATTCGATCCAACTACGCAAA**CTACAGCATCTTAGT** | 104632 |
| WF-Teratorn | AAACAGTTTCACGACAAGATACATCTGCTTCTATCCAAAGGATGGTGCACATGCAACACAGACAAAATGAAAAAAAAAAAATACACATTCGATCCAACTACACAAACTACAGCATCTTAGT | 108179 |
| TeratornLC199500 | **CCTCCTCAGCTTTGCGGATCTGGGGATCGCAGGCCCTGCTCCGCCTCCCCCTCCGCAGTACACCAAGTAGATGATCCCAGCAGAAAGCGCTCCGCACCAGGTCGTGAGAAGGCCCATCGA** | 104752 |
| WF-Teratorn | CCTCCTCAGCTTTGCGGATCTGGGGATCGCAGGCCCTGCTCCGCCTCCCCCTCCGCAGTACACCAAGTAGATGATCCCAGCAGAAAGCGCTCCGCACCAGGTCGTGAGAAGGCCCATCGA | 108299 |
| TeratornLC199500 | **AATCCACAAAGCCGAGCTGCTGGCGAACCCCGATAGCCAGCCTCCCAAAGAGACTCTCCCTGCCACGATCTCGTTGTACTTTGCGTCAAAATCAGTCAGGTCTCTGGTGATGTTGTTCAG** | 104872 |
| WF-Teratorn | AATCCACAAAGCCGAGCTGCTGGCGAACCCCGATAGCCAGCCTCCCAAAGAGACTCTCCCTGCCACGATCTCGTTGTACTTTGCGTCAAAATCAGTCAGGTCTCTGGTGATGTTGTTCAG | 108419 |
| TeratornLC199500 | **GTTTTGACGCACGGACAACAGGTTCGACGTCGTGTTCACGGCTCGCAGCAGCTCCATTTCAGGAGTCCCGGGCAGACTGTCTTTCAGCAGGTCCGCGAACGACACCGGCCCGTCCGATAT** | 104992 |
| WF-Teratorn | GTTTTGACGCACGGACAACAGGTTCGACGTCGTGTTCACGGCTCGCAGCAGCTCCATTTCAGGAGTCCCGGGCAGACTGTCTTTCAGCAGGTCCGCGAACGACACCGGCCCGTCCGATAT | 108539 |
| TeratornLC199500 | **GTGCGGCCTGGAGAAAGGCTTCAGCGGGGGGGATTCGATCTCGACCGTGACCGACCCCTCCATCCAAGTTACCTTCTTGTCGCGGTAGCAGATGTTTCTGTGGCACTCTCCGCACACGGT** | 105112 |
| WF-Teratorn | GTGCGGCCTGGAGAAAGGCTTCAGCGGGGGGGATTCGATCTCGACCGTGACCGACCCCTCCATCCAAGTTACCTTCTTGTCGCGGTAGCAGATGTTTCTGTGGCACTCTCCGCACACGGT | 108659 |
| TeratornLC199500 | **CTTGCATTCCGAGGGGTCCTCGACGCACCTCCTGTACCTTTCGTGAAGCTCGGGCCTCCTGCAGTACTTTCCGTGGCACTCGGAGGGCTCGTAGAACCGTCCGTCTATGAGCCAGACATC** | 105232 |
| WF-Teratorn | CTTGCATTCCGAGGGGTCCTCGACGCACCTCCTGTACCTTTCGTGAAGCTCGGGCCTCCTGCAGTACTTTCCGTGGCACTCGGAGGGCTCGTAGAACCGTCCGTCTATGAGCCAAACATC | 108779 |
| TeratornLC199500 | **GTAGTCTGGCCAACAGGGAACGCCATCGGTGCTGATTTTGGGCATTGGTATGATCCCTCGGACCACGGCGCGCTCCACGGCGTTCGGCAACGAGTGCACGATGAACAGCTTTCGATTTCT** | 105352 |
| WF-Teratorn | GTAGTCTGGCCAACAGGGAACGCCATCGGTGCTGATTTTGGGCATTGGTATGATCCCTCGGACCACGGCGCGCTCCACGGCGTTCGGCAACGAGTGCACGATGAACAGCTTTCGATTTCT | 108899 |
| TeratornLC199500 | **GTACAGGGCCGCGCCCACGGTCGGGAACTCGGACAGACCCGGGTGATCGCGCATCACGGTCATGCCACTGGGGCAGCTCCCCGAGCGCTTGGAGGCGATTTGCCTCAGACAGTTTTCCAC** | 105472 |
| WF-Teratorn | GTACAGGGCCGCGCCCACGGTCGGGAACTCGGACAGACCCGGGTGATCGCGCATCACGGTCATGCCACTGGGGCAGCTCCCCGAGCGCTTGGAGGCGATTTGCCTGAGACAGTTTTCCAC | 109019 |
| TeratornLC199500 | **CCCCCTGGCCATGAACTTGGTGTGCATGGCAGCCTGCGTCATCTGATGGGTCACGCTTTGCATCTGAGCGTACCACATCTGGTAGCTGGCCATCACGGAGAACTTGACGTTGGTGTCTCG** | 105592 |
| WF-Teratorn | CCCCCTGGCCATGAACTTGGTGTGCATGGCAGCCTGCGTCATCTGATGGGTCACGCTTTGCATCTGAGCGTACCACATCTGGTAGCTGGCCATCACGGAGAACTTGACGTTGGTGTCTCG | 109139 |

TeratornLC199500 **CAGGGCTGCCTCGGTGTTGCGCTTCATCGCGTCCCTGAGGAGTCTAAAGTTCTTGTTGGCGATCTCGGTGTTGCGCTTCATCTTCCTCTCCTGTTCGTCTATCCTCCCGTCTATCAGCGC** 105712
WF-Teratorn CAGGGCTGCCTCGGTGTTGCGCTTCATCGCGTCCCTGAGGAGTCCAAAGTTCTTGTTGGCGATCTCGGTGTTGCGCTTCATCTTCCTCTCCTGTTCGTCTATCCTCCCGTCTATCAGCGC 109259

TeratornLC199500 **CACGTTGGCGTCGAGCCGGTTGCTCACCTCCACCAGTTGCCCAGCGACATTCGTGAACGCGTTCTTCACGGAGTCCATCTGGACCTCCAAGCGATCCACTCTCTGAGACACGTGCAATGC** 105832
WF-Teratorn CACGTTGGCGTCGAGCCGGTTGCTCACCTCCACCAGTTGCCCAGCGACATTCGTGAACGCGTTCTTCACGGAGTCCATCTGGACCTCCAAGCGATCCACTCTCTGAGACACGTGCAATGC 109379

TeratornLC199500 **CAGACCCAGGGCAGCGCCTCCGAGCAACGAGCCCAGGAACCTCTTGTGCCTCGTGCGGCGTCGCTCGGGCCCGTCGAGTGCCGGGAAGGGTGCATTGTCGGGCAGGGCGTACGCAGAGGC** 105952
WF-Teratorn CAGACCCAGGGCAGCGCCTCCGAGCAACGAGCCCAGGAACCTCTTGTGCCTCGTGCGGCGTCGCTCGGGCCCGTCGAGTGCCGGGAAGGGTGCATTGTCGGGCAGGGCGTACGCAGAGGC 109499

TeratornLC199500 **CTTGCGCTCCTGCGCGGCCAACTCCTCCAGACCCTCTCTCATCAGGTCTCCGGCCCTGCGGGCATCGTCTGTGATCTCGTGCGCTGAATCGCACAGCTTGCCGTAGTCTCTCCGGCAGCC** 106072
WF-Teratorn CTTGCGCTCCTGCGCGGCCAACTCCTCCAGACCCTCTCTCATCAGGTCTCCGGCCCTGCGGGCATCGTCTGTGATCTCGTGCGCTGAATCGCACAGCTTGCCGTAGTCTCTCCGGCAGCC 109619

TeratornLC199500 **GTCATCACCTCCGCAGACCGAGATCAGGCAGGGCTTGAGGGCGCTCACGTCGTAGTTCACGCAGCTCACGTTCAGGTTGATGTAGTCCAGGCCCGGGGTGAAGTTGGACACGAAGTGAAT** 106192
WF-Teratorn GTCATCACCTCCGCAGACCGAGATCAGGCAGGGCTTGAGGGCGCTCACGTCATAGTTGACGCAGCTCACGTTCAGGTTGATGTAGTCCAGGCCCGGGGTGAAGTTGGACACGAAGTGAAT 109739

TeratornLC199500 **GGCTTCCGCCGGTTTCACGATCACCCCGCGATCGTGCTCCACCACCGGCCTGTCCAACGCGTAGCTGTCGTCCACGGAGACTAGATTCAGGCCCTGGCTGCTCTCTCCCGTGTTGTGCGC** 106312
WF-Teratorn GGCTTCCGCCGGTTTCACGATCACCCCGCGATCGTGCTCCACCACCGGCCTGTCCAACGCGTAGCTTTCGTCCACGGAGACTAGATTCAGGCCCTGGCTGCTCTCTCCCGTGTTGTGCGC 109859

TeratornLC199500 **GGTGCACACCCTGCTCACGGGCGGAGCGTACTCGACGAACCTTTTCGCGTCCAGGATCCCCACAGTGTAGTGATTGGGATCGGTCGTGGGCACCATGGTGTTGATGTTTATCGCTACCTG** 106432
WF-Teratorn GGTGCACACCCTGCTCACGGGCGGAGCGTACTCGACGAACCTTTTCGCGTCCAGGATCCCCACAGTGTAGTGATTGGGATCGGTCGTGGGCACCATGGTGTTGATGTTTATCGCTACCTG 109979

TeratornLC199500 **CGGAGGCCTGCCCGACGACTCGACCCGCTTGTCCGGTTCCCCGGGGGTGCACGACTCCATGGCGTACAAGTTCCCCACGTAGATCCTGTCGTAGCTGGTCGAGTACATCTCCCTGTGGGG** 106552
WF-Teratorn CGGAGGCCTGCCCGACGACTCGACCCGCTTGTCCGGTTCCCCGGGGGTGCACGACTCCATGGCGTACAAGTTCCCCACGTAGATCCTGTCGTAGCTGGTCGAGTACATCTCCCTGTGGGG 110099

TeratornLC199500 **CTCCTTGAACACCCCGAGCGCTCCGTGGACGGCCACGAGGCCGGGGCACACCAGAGTGTGGCTCAGACCCACGTTGTACCGAGGATCCAGGGAGCTCTTGGAACCCGACGCCGGATCTTT** 106672
WF-Teratorn CTCCTTGAACACCCCGAGCGCTCCGTGGACGGCCACGAGGCCGGGGCACACCAGAGTGTGGCTCAGACCCACGTTGTACCGAGGATCCAGGGAGCTCTTGGAACCCGACGCCGGATCTTT 110219

TeratornLC199500 **GTACGCGCTCTCCGGGACGGTGATCACGATGTCGCCCAGCTCGTCGAGCTCGGGCAGTCTTTCGTTCGTAGAGGGCAGCGGCGGCGGCGTGCGGAGCTCGGCGGACCTCGCGAACCTGGA** 106792
WF-Teratorn GTACGCGCTCTCCGGGACGGTGATCACGATGTCGCCCAGCTCGTCGAGCTCGGGCAGTCTTTTGTTCGTAGAGGGCAGCGGCGGCGGCGTGCGGAGCTCGGCGGACCTCGCGAACCTGGA 110339

TeratornLC199500 **GGGCTTC------------------GCGGCTCCGGCCTTCGCGGTGGATATGCCCGCCGCCATCTGCTGCCTCTTCCAGTCCCTCAGCCTGTGCACGGCCGAGGCGATTCCCAGTATCGC** 106894
WF-Teratorn GGGCTTCCAGCCGGACGCAGGCGTCGCGGCTCCGGCCTTCGCGGTGGATATGCCCGCCGCCATCTGCTGCCTCTTCCAGTCCCTCAGCCTGTGCACGGCCGAGGCGATTCCCAGTATTGC 110459

TeratornLC199500 **GGGCATGTCGGAACGCACGCCACGCTTGAAGTCAGAGTCCACGGGGGCAATCATGAACTGGTTCACGCCGAAGGCCACGGCCACGAGTTCGGGAAACCGCTTCAGGGTCTCGAAAGACGT** 107014
WF-Teratorn GGGCATGTCAGAACGGACGCCACGCTTGAAGTCAGAGTCCACGGGGGCTATCAAGAACTGGTTCACGCCGAAGGCCACGGCCACGAGTTCGGGAAACCGCTTCAGGGTCTCGAAAGACGT 110579

TeratornLC199500 **CGAAGACACGGTCACGTTCACCATCACTCCCCTGGCCCAGACCGTGGTGCTGGAGCCCGAGCAGCACTTGGTGTTGAGGTCTTCTTCGCACCTGAGCGCTGTGTTGGTCTCCAGGTCCGC** 107134
WF-Teratorn CGAAGACACGGTCACGTTCACCATCACTCCCCTGGCCCAGACCGTGGTGCTGGAGCCCGAGCAGCACTTGGTGTTGAGGTCTTCTTCGCACCTGAGCGCGGTGTTGGTCTCCAGGTCCGC 110699

TeratornLC199500 **TAAAGGCAGACCGCCCTCCACCTCCCACCTGCCCGCGCCCTCAGGAGCGCTGAGCTTTAGTCCTCCGGTGAGGGATAAGGTGAAGACGATCTTGTTTGCTCGGACTCCTGTCCTGAGCAG** 107254
WF-Teratorn TAAAGGCAGACCGCCCTCCACCTCCCACCTGCCCGCGCCCTCAGGGGCGCTGAGCTTCAGTCCTCCGGTGAGGGATAGGGTGAAAACGATCTTGTTTGCTCGGACTCCTGTCCTGAGCAG 110819

TeratornLC199500 **AGTCATGAGGTAGGGGACGACGGCGTAGCGCTCGTCGTCGCCGCCGTCAGTCCTCTGTCCGCTCGGGCTCACCCACGCGTTCTGGCAGTGCCTGGGATCCTTCGCGTCCGCGCGGCCCCG** 107374
WF-Teratorn AGTCATGAGGTAGGGGACGACGGCGTAGCGCTCGTCGTCGCCGCCGCCAGTCCTCTGTCCGCTCGGGCTCACCCACGCGTTCTGGCAGTGCCTGGGATCCTTCGCGTCAGCGCGGCCCCG 110939

TeratornLC199500 **CGGCGTCACGAACGCTTCGAACTCGGCCAGGTCCAAGAAGTCAGTGCTCGCCAGCAGCCCGCGAGAAGGAGGCAGTCCGAGGAACAACTTGCGGACGCCTCGTCTGTGCAGGGCGGACGC** 107494
WF-Teratorn CGGCGTCACGAACGCTTCGAACTCGGCCAGGTCCAAGAAGTCAGTGCTCGCCAGCAGCCCACGTGAAGGAGGCAGGCCGAGGAACAACTTGCGGACGCCTCGTCTGTGCAGGGCGGACGC 111059

TeratornLC199500 **GGCAAGCTTCGCAACTCTGCCGAACTGACGCCTTTCCCAGTTCGACTTGGAGCAGTTCGTCAGCCCCCACCGGCTGGCGTCCAAGAGCAAAGCCGTGGACGAGACCGAGGTGCTGACCCT** 107614
WF-Teratorn GGCAAGCTTCGCGACTCTGCCGAACTGACGCCTCTCCCAGTTCGACTTGGAGCAGTTCGTCAGCCCCCACCGGCTGGCGTCCAAGAGCAAAGCCGTGGACGAGACCGAGGCGCTGACCCT 111179

TeratornLC199500 **CGCCAGGGTTTCGGCCACGGACTCCAGGACCGCGGGGAACTCTTTTCGGTCCGGGGACACGGGGACCACGCACTCGTCAAACGGGACCTTTGCCACGACGTGGCCGGCTTGCGCCACCGT** 107734
WF-Teratorn CGCCAGGGTTTCGGCCACGGACCCCAGGACCGCGGGGAACTCTTTTCGGTCCGGGGACACGGGGACCACGCACTCGTCGAACGGGACCTTTGCCACGACGTGGCCGGCTTGCGCCACCGT 111299

TeratornLC199500 **GGCGAACAGGGCCGCGTCTAGAGCTACGGCTAAGTAGCCTCCGGCCGAGGAGTACAGAGCGCGCTCCACCGAGATGACAGTTGCGTCACACATGGAGGGTGCGACAAAGTCTCTGATATC** 107854
WF-Teratorn GGCGAACAGGGCCGCGTCTAGAGCTACGGCTAAGTAGCCTCCGGCCGAGGAGTACAGAGCGCGCTCCACCGAGATGACAGTTGCGTCACACATGGAGGGTGCGACAAAGTCTCTGATCTC 111419

| | | |
|---|---|---|
| TeratornLC199500 | **CTCGGGCGTCGGCGAGTAATCCGGGCCGCGGGCGGCCAAGTCGTCGACGCTGGAGATCTCACACATGGCCAACGGAGCGGAGGCCAGGCCGTCCCAGTTGGTCAGACCTGGCCTCGCGTG** | 107974 |
| WF-Teratorn | CTCGGGCGTCGGCGAGTAATCCGGGCCGCGGGCGGCCAAGTCGTCGACGCCGGAGATCTCGCACATGGCCAACGGAGCGGAGGCCAGGCCGTCCCAGTTGGTCAGACCTGGCCTCGCGTG | 111539 |
| TeratornLC199500 | **CTCCGAGCTGCTCACGCAGAACGTGTTGCCGTTGGATTCGAAACAGTTGTCTCCCAGCTGCTGGTGGCAGGCGCCCATCGCCTCTGCGTCCCCCAGGCCGACGTACACCAGCACGCAGGC** | 108094 |
| WF-Teratorn | CTCCGAGCTGCTCACGCAGAACGTGTTGCCGTTGGATTCGAAACAGTTGTCTCCCAGCTGCTGGTGGCAGGCGCCCATCGCCTCTGCGTCCCCAAGGCCGACGTACACCAGCACGCAGGC | 111659 |
| TeratornLC199500 | **GTAGGCCGAGTTGCGGATGGCCAGGACGTCGTCCGAGTCAGGCCACCGCCTGCTGGCCTCGGGCGCGCGCAGCGTTCTCTTCATGAGCTTGTAGGCCGCGCACTGAGCTCCCGGGTCGCA** | 108214 |
| WF-Teratorn | GTAGGCCGAGTTGCGGATGGCCAGGACGTCGTCCGAGTCAGGCCACCGCCTGCTGGCCTCGGGCGCGCGCAGCGTTCTCTTCATGAGCTTGTAGGCCGCGCACTGAGCTCCCGGGTCGCA | 111779 |
| TeratornLC199500 | **TTCGGCCGACGAGTTCGCCCCGAAGTAATAGTCTCTGTTGCGAGGGCTGGCGTGCAGCATCGCCGATATCCTGTTCAGCGGCGCGTGCGGGACCACCCGCTCGGAGCACCCGAACCCCCC** | 108334 |
| WF-Teratorn | TTCGGCCGACGAGTTCGCCCCGAAGTAATAGTCTCTGTTGCGAGGGCTGGCGTGCAGCATCGCCGATATCCTGTTCAGCGGCGCGTGCGCGACCACCCGCTCGGAGCACCCGAACCCCCC | 111899 |
| TeratornLC199500 | **CGGGACCGGGACGCACGAGGACCTGTCGAGCTTCTTCACGTAGGCTCCCAGCATCACGCCTTCCGTCTGGTACCTGGTCAAGTCGGGAGAGCGTCCGCGCGTGTCTCTCCACAGGGCGCC** | 108454 |
| WF-Teratorn | CGGGACCGGGACGCACGAGGACCTGTCGAGCTTCTTCACGTAGGCTCCCAGCATCACGTCTTCCGTCTGGTACCTGGTCAAGTCGGGAGAGCGTCCGCGCGTGTCTCTCCACAGGGCGCC | 112019 |
| TeratornLC199500 | **GATACTTCCCGGGCAATAATGCGTGCACTGGTCCCGTTCACCCTCCTGAGAGCCGATCCTGCGCAAGGCGAAGCCTCCCTGGCTGGTGCCCATGCACCAGCCGGTGTGAGCGCCACATTT** | 108574 |
| WF-Teratorn | GATACTTCCCGGGCAATAATGCGTGCACTGGTCCCGTTCACCCTCCTGGGAGCCGATCCTGCGCAAGGCGAAGCCTCCCTGGCTGGTGCCCATGCACCAGCCGGTGTGAGCGCCACATTT | 112139 |
| TeratornLC199500 | **CGTCACGG-AGATCCACGCGTAAGGAGCATACCCGTAAGGTTCGTAGTCCGCGGGCGCTTGGGTTCGGGCTCGGCAAAGAAGCCACACGGCAAGCGTGGCGACGAGCGCGAGCAT**GTCAC | 108693 |
| WF-Teratorn | CGTCAGGGGAGATCCACGCGTAAGGAGCATACCCGTAAGGTTCGTAGTCCGCGGGCGCTTGGGTTCGGGCTCGGCAAAGAAGCCAGACGGCAAGCGTGGCGACGAGCGCGAGCATGTCAC | 112259 |
| TeratornLC199500 | CACCCTTCCGAGGCGGTCGATCGTTGACCGACTCGAGAAGGTGCGCGCTAATACTCGACGCCTCCCCCCCCCCCCCCCCGGGAGGAGTCTGGAAGCCCGGCCTCGCTCT**TCACAGGAGAG** | 108813 |
| WF-Teratorn | CACCCTTCCGAGGCGGTCGATCGTCGACCGGCTCGAGAAGGTGCGCGCTAATACTCGACGCCTCCCCCCCCCCT-----GGTAGGAGTCGGGAAGCCCGGCCTCGCTCTTCACAGGAGAG | 112374 |
| TeratornLC199500 | **GGTGAGCGGGGTCGCCGTTACCCTTGTCCGCATCTGTCTGCTTGCGCTTCATGAGCGACTGGCACGTGGCGGCCAAGCCTATGAACTGCTTCACCGCGTCGTTGTGAACCTTGGCCACGG** | 108933 |
| WF-Teratorn | GGTGAGAGGGGTCGCCGTTACCCTTGTCCGCATCTGTCTGCTTGCGCTTCATGAGCGACTGACACGTGGCGGCCAAGCCTATGAACTGCTTCACCGCGTCGTTGTGAACCTTGGCCACGG | 112494 |
| TeratornLC199500 | **CCTGGTCAGGGACCAGGAAAAGTCTACCCATGTGGGTGCGCACGCGGTTCATGACGCGAACGGACGCCTGGCTCGCCAGGGAGTACGAGCCCGAAATCAGGTCCAGGGTCATGCTGCGCA** | 109053 |
| WF-Teratorn | CCTGGTCAGGGACCAGGAAAAGTCTACCCATGTGGGTGCGCACGCGGTTCATGACGCGAACGGACGCCTGGCTCGCCAGGGAGTACGAGCCCGAGATCAGGTCCAGGGTCATGCTGCGCA | 112614 |
| TeratornLC199500 | **GCACCCCGGCGACGTTCTCGTCGTCTCTCGCGCAGTGGGCCAAGTGAAGGTTGATCTGCTTCCTTGTGAAAGCGTGATGGCACGACTTCAAGTACGGAGCCACCTCCTGCGGTTCGAGCA** | 109173 |
| WF-Teratorn | GCACCCCGGCGACGTTCTCGTCGTCTCTCGCGCAGTGGGCCAAGTGAAGGTTGATCTGCTTCCTTGTGAAAGCGTGATGGCATGACTTCAAGTAGGGAGCCACCTCCTGCGGTTCGAGCA | 112734 |
| TeratornLC199500 | **CTCCGATGGAGTCGAGGGTGTTTTTGTCCAACCTCTCGCACAGGAGGCACTTGGAGCGCTCCTTTTCGTAACAGAGACTCCTGATCGAGGGGTGCTCCCCTCCGCTTATGTTCATGTGAA** | 109293 |
| WF-Teratorn | CTCCGATGGAGTCGAGGGTGTTTCTGTCCAACCTCTCGCACAGGAGGCACTTGGAGCGCTCCTTTTCGTAACAGAGACTCCTGATCGAGGGATGCTCCCCACCGCTTATGTTCATGTGAA | 112854 |
| TeratornLC199500 | **GCATGGGATACTGTTTTCTGAGAAACTGATCCGCTTCGCTGACGCTGCGCAACTCGCCCTCCGTCAAGTCAGAGGCCCCTGCCTCGGGAAACATCTGAAAGATGGCCTCGGAGGCCAT**GC | 109413 |
| WF-Teratorn | GCATGGGATACTGTTTTCTGAGAAACTGATCCGCTTCGCTGACGCTGCGCATCTCGCCCTCCGTCAATTCAGAGGCCCCGGCCTCGGGAAACATCTGAAAGATGGCCTCGGAGGCCATGC | 112974 |
| TeratornLC199500 | TCGACAAGTAGCCCGGCGAGTCATGGGACGCTTCGTTCTCGGCGCTCGACGACGACGACGGCGGGCCGTCCGGGTACCGCCCGCGTCTGCCAGCGGCCTCGTGGCCTCGATTACTAACCG | 109533 |
| WF-Teratorn | TCGACAAGTAGCCCGGCGAGTCGTGGGACGCTTCGTTCTCGGCGCTCGACGACGACGACGGCGGTCCGTCCGGGTACCGCCCGAGTCTGCAAGCGGCCTCGCGGCCTCGATTACTAACCG | 113094 |
| TeratornLC199500 | CGGAGGCCCTCCGTCCCCCGAGCGGCACCCTGGTCCTTTCCCCCTCGCCTTCCGCTCCGTGACGCGTCGACCGGC**ATGGACGGCCCGAGGCTTCACCGCGTGAAGACGGGCGGGGCCCGA** | 109653 |
| WF-Teratorn | CGGAGGCCCTCCGTCCCCCGAGCGGTACCCTGGTCTTTTCCCCCTCGCCTTCCGCTCCGCGACGCGTCGACCGGCATGGACGGCCCGAGGCTTCACCGCGTGAAGACGGGCGGGGCCCGA | 113214 |
| TeratornLC199500 | **GCTCAGAGTCCATCGTTCGCGAGCCGTGAGGAGCCCGGTGCTCCATCCAGAGGAGTCTGGGGGGTGATGGAGAGCGACAGGTCGACCGTGCGTGGGGCGCTCTTTTCTTCCACGAACCCG** | 109773 |
| WF-Teratorn | GCTCAGAGTCCATCGTTCGCGAGCCGTGAGGAGCCCGGTGCTCCATCCAGAGGAGTCTGGGGGGTGATGGAGAGCGACAGGTCGACCGTGCGTGGGGCGCTCTTTTCCTCCACGAACCCG | 113334 |
| TeratornLC199500 | **TGGTTCTGGGGTATCCCGGCGCACAGGCGATGCGTGTTCCGCACAAGCTCGGAGCGCTCGATAGCCGTGATAGACCGCGCGAGGTGTAAATGGTGCAAACCGTCCCGCGGGGAGGTGCCG** | 109893 |
| WF-Teratorn | TGGTTCTGGGGTATCCCGGCGCACAGGCGATGCGTGTTCCGCACAAGCTCGGAGCGCTCGATAGCCGTGATAGAACGCGCGAGGTGTAAATGGTGCAAACCGTCCCGCGGGGAGGTGCCG | 113454 |
| TeratornLC199500 | **CGTTTCGACCAGGACGAGTTCGAGCGGATCAGGACGAGGTCTTACTTGCGCGGGACGTTCGGCTTGTCGCCAGCCGAAGCCGGTCGTCTTTCGGAGCTGCTGGCGGAGCTTGCACTTTCG** | 110013 |
| WF-Teratorn | CGTTTCGACCAGGACGAGTTCGAGCGGATCAGGACGAGGTCTTACTTGCGCGGGACGTTCGGCTTGTCGCCAGCCGAAGCCGGTCGTCTTTCGGAGCTGCTGGCGGAGCTTGCACTTTCG | 113574 |
| TeratornLC199500 | **GCCGAGGGGAAGGGCGCCTGTCCGTTCGCGGGCACCCCGCCTTACTGCACCGTCGAGAACGTGGTTCACATCTCGGGGCCGGCGCTGGAGAGACTCATGGAGTTCGACGACGAGTTCGTC** | 110133 |
| WF-Teratorn | GCCGAGGGGAAGGGCGCCTGTCCGTTCGCGGGCACCCCGCCTTACTGCACCGTCGAGAACGTGGTTCACATCTCGGGGCCGGCGCTGGAGAGACTCATGGAGTTCGACGACGAGTTCGTC | 113694 |

TeratornLC199500 **TGGGGAGACAATCTTATCTTCCACGCGGGAGCCCGTCCACGCTCGTGCCACACTTAG**CCACTCGAAGCGTGAGCCCGCCTATATACGGCGGCGGGTTACGTCTCTCGATGAGATGGGCGC 110253
WF-Teratorn TGGGGAGACAATCTGATCTTCCACGCGGGAGCCCATCCACGCTCGTGCCACACTTAGCCACTCGAAGCGTGAGCCCACCTATATACGGCGGCGGGTTACGTCTCTCGAGGAGATGGGCGC 113814

TeratornLC199500 CTCGCGATCGGGGAGAGGAGAG**ATGCCGAGCGGGGACTGCCGAGGATCCCCCTACGTCCCCCGGACGGGACCACTGTTCTGCCTTAAGGGCCAGAGCGTGTACGTGAAGATGACCGAGGC** 110373
WF-Teratorn CTCGAGATCGGGGAGAGGAGAGATGCCGAGCGGGGACTGTCGAGGATCCCCCTACGTCCCCCGGACGGGACCACTGTTCTGCCTTAAGGGCCAGAGCGTGTACGTGAAGATGACCGAGGC 113934

TeratornLC199500 **GGACGAGAAGAGGGCCTTGCTGAGCGAGAGCGACATGACGCTGGTGGGCTACGCCGATAGCGGCGAGTGCGAGGGCGAGGAGCTGCGCTCTGACCCGTCCGTCTTCGTGCCGATCTTCTG** 110493
WF-Teratorn GGACGAGAAGAGGGCCGTGCTGAGCGAGAGCGACATGACGCTGGCGGGCTACGCCGATAGCGGCGAGTGCGAGGGCGAGGAGCTGCGCTCTGACCCGTCCGTCTTCGTGCCGATCTTCTG 114054

TeratornLC199500 **CAGAGAGTGCTACGGGACAGAGACACGGGATTACGAGGAGGCTCTGCGGGACATTCTCGCTCGCGGGACAAACTCGTTCATCGTGGGCGTCTGCAGCGTGTATGGCGCGGCCCCCGACGA** 110613
WF-Teratorn CAGAGAGTGCTACGGGACAGGGACACGGGATTACGAGGAGGCTCTGCGGGACATTCTCGCTCGCGGGACAGACTCGTTCATCGTGGGCGTCTGCAGCGTGTATGGCGCGGCCCCCGACGA 114174

TeratornLC199500 **GATTTACCCCGAATCGAGGGCCGAGTTCTCGGTGGAGAACGCGTTGTCATTCTCGGGGGCCTGCCACCCGCAGCCCGGCAAGGCCGTGTTCCTGCACGACGCGGAAAGAGACATCGGCTG** 110733
WF-Teratorn GATTTACCCCGAATCGAGGGCCGAGTTCTCGGTGGGGAACGCGTTGTCATTCTCGGGGGCCTGCCACCCGCAGCCCGGCAAGGCCGTGTTCCTGCACGACGCGGAAAGAGACATCGGCCG 114294

TeratornLC199500 **CGTAAAGTGCTACTGGCACGTTCACAGCAGGTCAGCGGACAAGACCTTCCTGTTTTGCTGCGTGGCCTGCGACGGCACAGAGTACACCGCCAGGTTGCTCTCGGAGCTGAGGAAAAACGA** 110853
WF-Teratorn TGTAAAGTGCTACTGGCACGTTCACAGCAGGTCAGCGGACAAGACCTTCCTGTTTTGCTGCGTGGCCTGCGACGGCACAGAGTACACCGCCAGGTTGCTCTCGGAGCTAAGGAAAAACGA 114414

TeratornLC199500 **GACGCTGAGTACGGCCTTCAGCTTGGGAACAGTGGGAGACAACTCCGTCGGGCCCGTCGTCGAGGAGTGCAGCCTGGTCTCGGTGCCCATGCGACCAGCCTGCTTCTGCGTGGCGCTCAC** 110973
WF-Teratorn GACGCTGAGCACGGCCTTCAGCTTGGGAACAGTGGGAGACAACTCCGTCGGGCCCGTCGTCGAGGAGTGCAGCCTGGTCTCGGTGCCCATGCGACCAGCCTGCTTCTGCGTGGCGCTCAC 114534

TeratornLC199500 **GGACAAGGACCTCGTGTCTCGGACCATGACGAGGATGGGCTTTTGCTCGGTGAAGGACAAAGCCCTCGAAGCCGGGCTGCGCCCGGACGAGGGGGGCAGCCCGTGGGAGCCGGAGGGGAC** 111093
WF-Teratorn GGACAAGGACCTCGTGTCTCGGACCATGACGAGGATGGGCTTTTGCTCGGTGAAGGACAAAGCCCTCGAAGCCGGGCTGCGCCCGGACGAGGGGGGCAGCCCGCGGGAGCCGGAGGGGAC 114654

TeratornLC199500 **CGGACACCCCGGACTCCGGGAGGGCGAGAGCTCGGAAGATGGGATCGAGGCCGTGAGGCGGCTCGCGATCAAGCAGGCGGACGAGTTGCGAGCGCTCGAGGACGTCCTGAAAGCGATCAA** 111213
WF-Teratorn CGGACACCCCGGACTCCGGGAGGGCGAGAGCTCGGAAGACGGGATCGAGGCCGTGAGGCGGCTCGCGATCAAGCAGGCGGACGAGTTGCGAGCGCTCGAGGACGTCCTGAAAGCGATCAT 114774

TeratornLC199500 **GGTACAGAACGGGGTTCTCGCGCAAAGGTTGAAATGGCCCCAGGACCCCGACACTCTGTTCACGATGCGAGGGGCGAAGGACTTCACGGAGCGCACCGGGGTGAGCAACAGACTGTGTGC** 111333
WF-Teratorn GGTACAGAACGGGGTTCTCGCGCAAAGGTTGAAATGGCCCCAGGACCCCGACACTCTGTTCACGATGCGAGGGGCGAAGGACTTCACGGAGCGCACCGGGGTGAGCAACAGACTGTGTGC 114894

TeratornLC199500 **GGGCGAGGAAGCCCGTACTAACGCCGCGACACCGGACCGAGGGGGTCGGGGGTCCGAAAAGGACATGGAGGCCCGCGAGGATCAATCGCAACAAGCTTTGTCGAGGTACGAACTGGAAGA** 111453
WF-Teratorn GGGCGAGGAAGCTCGTACTAACGCCGCGAGACCGGACCGAGGGGGTCGGGGGTCCGAAAAGGACATGGAGGCCCGCGAGGATCAATCGCAACAAGCTTTGTCGAGGTACGAACTGGAAGA 115014

TeratornLC199500 **AGTCAGGCGACTACTCTCCGCTTCGAGACGAGCCGGGGAGGAAGTGGGCCAGCGCCTGCCGCAATACGGAGTCCGTGGGGGAGATGCTCTGGCGGCACTGGGGGGGTCGACCCGGCGAGC** 111573
WF-Teratorn AGTAAGGCGACTACTCTCCGCTTCGAGACGAGCCGGGGAGGAAGTGGGCCAGCGCCTGCCGCAATACGGAGTCCGTGGGGGAGATGCTCTGGCGGCACTGGGGGGGTCGACCCGGCGAGC 115134

TeratornLC199500 **CCCGCACTACAGCTACTGCGAGGACGACGACTACGACTGCAACGAAGTCGCGTCAGGTCGTCCCCGGACGCGATCTCAGAGTGGGAAGACAAAGCCCTCACCGGTCGACCGCCTGACGGA** 111693
WF-Teratorn CCCGCACTACAGCTACTGCGAGGACGACGACTACGACTGCAACGAAGTCGCGTCAGGTCGTCCCCGGACGCGATCTCAGAGTGGGAAGACAAAGCCCTCACCGGTCGACCGCCTGACGGA 115254

TeratornLC199500 **GCAACAGAGGAGAGAGGTAGCGCTGGAGTACAACAGACAAGAGTCCGAGAGAGAGGAGGCCTCGCGGAAAAAGAGGGAGACCATGATCGAAATGATCAAAGCGAGCATGGGCGACTTGAT** 111813
WF-Teratorn GCAACAGAGGAGAGAGGTAGCGCTGGAGTACAACAGACAAGAGTCCGAGAGAGAGGAGGCCTCGCGGAAAAAGAGGGAGACCATGATCGAAATGATCAAAGTGAGCATGGGCGACTTGAT 115374

TeratornLC199500 **CAAGAGCGGCATCAAGGATGCGATGCACCAAGAGCGCGGGTTCCAGGCGACCGCTGAAGAGGTCGGGGCACCCCAGTCAGCTTTGCCCAGCGCCGAGTTGGAGACCCTAGCGAGAGACAC** 111933
WF-Teratorn CAAGAGCGGCATCAAGGATGCGATGCACCAAGAGCGCGGGTTCCAGGCGACCGCTGAAGAGGTCGGGGCACCCCAGTCAGCTTTGCCCAGCGCCGAGTTGCAGACCCTAGCGAGAGACAC 115494

TeratornLC199500 **TCCGTACAACAGGCAGCCTAGGAACTTGGACGGGCGGCGGCCTTTGCCCGAGAGCCTGTCCGTGGACGTCCCAGCGCTGCTGAAAGAACTGGAAGAGCTGCGACAGCGATCGTTCGCGAG** 112053
WF-Teratorn TCCGTACAACAGGCAGCCTAGGAACTTGGACGGGCGGCGGCCTTTGCCCGAGAGCCTGTCCGTGGACGTCCCAGCGCTGCTGAAAGAACTGGAAGAGCTGCGACAGCGATCGTTCGCGAG 115614

TeratornLC199500 **GAGCTCCGGAAAGAGCGGGGACAGCGAAGGATCGCTGCCCATGGACACGGACGACGACGCTCCCACCGGCGGAGGCGCGGAGGCCACTAGCTCCTTGCGCAGCAACGGCACAGACACGAC** 112173
WF-Teratorn GAGCTCCGGAAAGAGCGGGGACAGCGAAGGATCGCTGCCCATGGACACGGACGACGACGCTCCCACCGGCGGAGGCGCGGAGGCCACTAGCTCCTTGCGCAGCAACGGCACAGACACGAC 115734

TeratornLC199500 **GGCGGCGAAAAACAAGCAGCAGGCGGGCTACACTCTGAGGCCGTCCGGAGAGGTGCCCGGCATTTCTGACTCTCACGGAACGCTGCAAGAGTTCATTTCTGCGGTCTCGGTGCACAGATG** 112293
WF-Teratorn GGCGGCGAAAAACAAGCAGCAGGCGGGCTACACTCTGAGGCCGTCCGGAGAGGTGCCCGGCATTTCTGACTCTCACGGAACGCTGCAAGAGTTCATTTCTGCGGTCTCGGTGCACAGATG 115854

TeratornLC199500 **A**TTCCGCTTACCGGGGCCGGGACCCTGCATAACATGTTTTTGTACACATTTTCAGAATGTCTGTAGGCTTGTATGACTTGCTTTTTTATTTTACTTTTTTTTTTCCTACAAGAAATAAAG 112413
WF-Teratorn ATTCCGCTTACCGGGGCCGGGACCCTGCATAACATGTTTTTGTACACATTTTCAGAATGTCTGTAGGCTTGTATAACTTGCTTTTTTATTTTACTTTTTTTTTTCCTACAAGAAATAAAG 115974

TeratornLC199500 CAGAAAAACTCTTTGAACGCTCCACACACTGTTGTTCTCTTTCCACTTCCCGCTCGAATAAGATTAAAAAATGAACACACGTTTGCGTATAACAGCACTTGCAATCGCAATGCCTTTCCT 112533
WF-Teratorn CAGAAAAACTCTTTGAACGCTCCACACACTGTTGTTCTCTTTCCACTTCCCGCTCGAATAAGATTAAAAAATGAACACACGTTTGCGTATAACAGCACTTGCAATCGCAATGCCTTTCCT 116094

TeratornLC199500 GGGCTTCGGTCTCAGGATCTGTACGCTTTCCACTAACGACCAGCTCACGGTTCGAACGTTTTGCCAAAGTGCATC**ATGGCGAACTCGATCTGCGTTACCCTCAACCTGCCCTCTGTGACG** 112653
WF-Teratorn GGGCTTCGGTCTCAGGATCTGTACGCTTTCCACTAACGACCAGCTCACGGTTCGAACGTTTTGCCAAAGTGCATCATGGCGAACTCGATCTGCGTTACCCTCAACCTGCCCTCTGTGACG 116214

TeratornLC199500 **GTGGCTCAGGTTCAGCAACTGCAACATTCGCGTTGCGAAGAGGCTGACGACGACGACGCGGTACACACGGGCGCCTCGTCCTGTTGGAAGAGAAGCTGCGATAACATGGCCATGGGCGCT** 112773
WF-Teratorn GTGGCTCAGGTTCAGCAACTGCAACATTCGCGTTGCGAAGAGGCTGACGACGACGACGCGGTACACACGGGCGCCTCGTCCTGTTGGAAGAGAAGCTGCGATAACATGGCCATGGGCGCT 116334

TeratornLC199500 **GTCGATTTCAGTTACGTCGCATCGCAGATGATTCCTTCCACGGTAAAGCAGGCGCCTCAAGTGGGAGACCTTTGCGTGCAGCTGGCTCCCCTCGCAGCTCGCACGGAAGAGTATGCGTCG** 112893
WF-Teratorn GTCGATTTCAGTTACG-------------------------------------------------------------------------------------------------------- 116350

TeratornLC199500 **ATAGTGGATCTGTACAACACGTACTGTGCAAAGGGTAGCGCACAGATGGCCGTGAGGTGCATGCGGAGCTTCGCGAGGAGCCAAAGCGAGCTGTGCATCCCCGCGATACCGATACACATC** 113013
WF-Teratorn ------------------------------------------------------------------------------------------------------------------------ 116350

TeratornLC199500 **TTCGTGTCAGAATGGAACAGGTCCCTGAAACGGTTCCACGACATGGTCAGGTGCAACAGGGAATGGACGAACGCGCGCACAGGCGCTGCTCCCGCACAGGCTGTGAGAACCTCGCTGCTG** 113133
WF-Teratorn --------------------------------------------GGTCA----------------------------------------------------------------------- 116355

TeratornLC199500 **AACAACGTGATCACGCTGACTCTGGGGGACGTGCCTGCTCGAAGCGAGCACCTCTGGTACCTGCTACACTGCGTGATTTCCAAACCGTCGTTGGTCAAAGAGGCGCTGCAGCTCGTGGGC** 113253
WF-Teratorn ------------------------------------------------------------------------------------------------------------------------ 116355

TeratornLC199500 **TGCGTGTCCTCGTGCTCGGGCACGCTCTATCGCACGGGCGACGTGACCAACTCGACCGTTCCGTGCAAGAACGGCTCGAGCGCATCCAGGCTGTACCTGATCGCCAAGTCGGCGGCCGGC** 113373
WF-Teratorn ------------------------------------------------------------------------------------------------------------------------ 116355

TeratornLC199500 **GCGAACGACACTCGGCTGGGCGAGAACGCGGGTGCCTCTCTGGGTCTGATTTGGGCGTCGAGCGAAGACGAGCTCTTCGATCACTTTCGCAACGCGAGATGCTGCCACTACGTCCACCTG** 113493
WF-Teratorn ------------------------------------------------------------------------------------------------------------------------ 116355

TeratornLC199500 **GGGACAGAGTCCGGGAACGCGTTCAGGTGCAACGGGTACTGCGCGTGGCGCACCCACGACCGGGAGGTTTTCGAACGGATGAGCGAGCTCGGTGCAGAAGTTGCGTCGATGCCCTGCGTG** 113613
WF-Teratorn ------------------------------------------------------------------------------------------------------------------------ 116355

TeratornLC199500 **CAGGATTCGCACCTGGCCGGAAACACGGCCGACGCGGTGAGAGCAGCGTGCCAGCTGTGA**GGCGAAGGCTGACGGCCTCCGGTCCCTCGAGCTTGGGGTTTCAGACGGCGCATCTGACCG 113733
WF-Teratorn ------------------------------------------------------------------------------------------------------------------------ 116355

TeratornLC199500 ACTAACGATCGTAACGCTGCGGACGATGGTGGGGGGGGGGGGGGACTCGAGAACAGCAGCTGGAGA**ATGCTGCCGAGGACTTTCGTGCCCGATTGTGACCTGATCGCGTCGGCCGCACTG** 113853
WF-Teratorn ------------------------------------------------------------------------------------------------------------------------ 116355

TeratornLC199500 **AACAGAGGAGCGGTCTTGACCTCCGGCGTCCACCCGCTATATCTATCCCCCGTGGTGTGCTGGCTCAATCGCACGCGTGAGCGATTGCGTCGGGCCAAGCGCTCTGCAGTCTGCGCGCTG** 113973
WF-Teratorn ------------------------------------------------------------------------------------------------------------------------ 116355

TeratornLC199500 **AAAGGAGCGTCAAACGCGGCGAGCATGCTAGCCATCTTATCCCACTCCGTGAAGCTCTGGGATTACCCGACCTTGGACGCGACCGCGGCAGGTTGGTCCCTGACGGTCTGCGGGGCTCTG** 114093
WF-Teratorn ------------------------------------------------------------------------------------------------------------------------ 116355

TeratornLC199500 **ATCGCAGGCCTCGCCGTGGCCGCCAACAGTTCCAGAGCATTGGTCTCTCGAGCGGACCCCAAAGTCATCGCTCAACCGCTGCGGGTCATTCCCCTCTGGGTCAAATGCCTTCTGTCCGCA** 114213
WF-Teratorn ------------------------------------------------------------------------------------------------------------------------ 116355

TeratornLC199500 **CTTTTGTTCTCTGCCTACACGTACACCTGGATGTACTGCGCGGACCCGTCTCGTTCTGGCTGCTGGGAAGAGGGCCGTAGAGCCTGGCCGCCGCTCTGGTCTTGCACGGTCGGCTTGTTC** 114333
WF-Teratorn ------------------------------------------------------------------------------------------------------------------------ 116355

TeratornLC199500 **CTGTGGATTAATGCGGTCGGTTTGCTGGGACTGCTCAGGCAGACCCGTTTGTGCTAA**GCGAGA**ATGGGATTGTGCGCAATCGGGGCGAGGTTTGGGCTTTCTTTTTTCATCTTTTGTTCA** 114453
WF-Teratorn ---------------------------------------------------------------------------------------AGGTTTGGGCTTTCTTTTTTCATCTTTTGTTCA 116388

TeratornLC199500 **CTGATCATCCGCAACCCTCTGGAACGGGGTTTCCAGTGGATCGTAGGATACGAGACCCTACTTGCGGTGGCGGCCGTAACATCGGTAGCGTCCTGTGTGTGCACGAAGTTCGCCTGCCTG** 114573
WF-Teratorn CTGATCATCCGCAACCCTCTGGAACGGGGTTTCCAGTGGATCGTAGGATACGAGACCCTACTTGCGGTGGCGGCCGTAACATCGGTAGCGTCCTGTGTATGCACGAAGTTCGCCTGCCTG 116508

TeratornLC199500 **ATCGAAACGCACAGACGCATGCAACACCTGGCCCTCTCAAACACGAGCAAAGGGGAAACTGTGGCAGTGTACAGGGAGAAGAAGCTGTCGACGGTCTCCCCTCAGGACGACGACGCGGCT** 114693
WF-Teratorn ATCGAAACGCACAGACGCATGCAACACCTGGCCCTCTCAAACACGAGCAAAGGGGAAACTGTGGCAGTGTACAGGGAGAAGAAGCTGTCGACGGTCTCCCCTCAGGACGACGACGCGGCT 116628

TeratornLC199500 **CGCTACGGGAAGCGAGCCTTCGGGCAGCTCCAGGACACACCCTTCCACCCTCGCGACAGATACGACGCGCCAAACTCTTTCTGGTGCGCGCTGACGACTGTCGCGCTCGCCTACAACACG** 114813

WF-Teratorn CGCTACGGGAAGCGAGCCTTCGGGCAGCTCCAGGACACACCCTTCCACCCTCGCGACAGATACGACGCGCCAAACTCTTTCTGGTGCGCGCTGACGACTGTCGCGCTCGCCTACAACACG 116748

TeratornLC199500 **GCTCACATCGCGCTGACCGACGCCTCGGATCCCGAGCCCCTCGCCTGTGGCGCTCCGGCACTGGCGCTTTACTGGTGCGTGACCAGAAGCGCGCAGCTGGCGCTCTCTTGA**CAGCGTTGG 114933

WF-Teratorn GCTCACATCGCGCTGACCGACGCCTCGGATCCCGAGCCCCTCGCCTGTGGCGCTCCGGCACTGGCGCTTTACTGGTGCGTGACCAGAAGCGCGCAGCTGGCGCTCTCTTGACAGCGTTGG 116868

TeratornLC199500 CGCCGCTCGCACTCTGACCGTG**ATGGTATTTACAAACGGGACGAGGGAGAAGAGGATGATGCACCCGAGGACTCCAGTGCGCTCAGAGAGCTGTCTGGAGGTGCTCCTGGAGGTCTTTCT** 115053

WF-Teratorn CGCAGCTCGCACTCTGACCGTGATGGTATTTACAAACGGGACAAGGGAGAAGAGGATGATGCACCCGAGGACTCCAGTGCGCTCCGAGAGCTGTCTGGAGGTGCTCCTGGAGGTCTTTCT 116988

TeratornLC199500 **ATCCGCATTGGTGATCGCCAACATCGTGCTGTGGTTGCAGAGGTTGTTCTTCCAGATACACCAAAGACTGGAATAA**GGGAGAGG**ATGGGTACCGCGGGGAGTTCGGGAAGCGCCTGGTGG** 115173

WF-Teratorn ATCCGCATTGGTGATCGCCAACATCGTGCTGTGGTTGCAGAGGTTGTTCTTCCAGATACACCAAAGACTGGAATAAGGGAGAGGATGGGTACCGCGGGGAGTTCGGGAAGCGCCTGGTGG 117108

TeratornLC199500 **ACGAAACTCGCCTCTCCGAAAGACGACGGGGCTCTGCGGAAGCGCCTCGGGTTCTCGAGCCCCGAGGCTGACCCGGCATCCCTCCGATGGTGGTCCGATCTCGCTCGCGGAGGAGCATCG** 115293

WF-Teratorn ACGAAACTCGCCTCTCCGAAAGACGACGGGGCTCTGCGGAAGCGCCTCGGGTTCTCGAGCCCCGAGGCTGACCCGGCATCCCTCCGATGGTGGTCCGATCTCGCTCGCGGAGGAGCATCG 117228

TeratornLC199500 **GAGCAGCAGCGGGACCTCGGCCACCAAAGCGTAGAGGACGGGGTCGACCTGGGTTCGAAGCAGAAGCTCCAGCCCCAGATCTCCTCGAATCTGTCAGCAAGGAAATCTACCAGCAAAACA** 115413

WF-Teratorn GAGCAGCAGCGGGACCTCGGCCACCAAAGCGTAGAGGACGGGGTCGACCTGGGTTCGAAGCAGAAGCTCCAGCCCCAGATCTCCTCGAATCTGTCAGCAAGGAAATCTACCAGCAAAACA 117348

TeratornLC199500 **ATGAGCACGGACGACTTTTGGAGATTGAAGGTGGCAAAAGTCCAGGGTGGCCAAACCCTTCACCGCAGTTATGACAGAAAACAGCGAAAGAAGGAAAAGAGTCAGCCAGAGTTGGAGAAA** 115533

WF-Teratorn ATGAGCACGGACGACTTTTGGAGATTGAAGGTGGCAAAAGTCCAGGGTGGCCAAACCCTTCACCGCAGTTATGACAGAAAACAGCGAAAGAAGGAAAAGAGTCAGCCAGAGTTGGAGAAA 117468

TeratornLC199500 **CTGTTGTGA**CCGCGTTTGACAAATGCAACATGTTTGCGAGCTCACACGAGCGTGTGCAGGTGATTGTAATGAAACGAGAATAAACCAATCATTCGCTGAAACCAAAGTCTTTTTGTTGTC 115653

WF-Teratorn CTGTTGTGACCGCGTTTGACAAATGCAACATGTTTGCGAGCTCACACGAGCGTGTGCAGGTGATTGTAATGAAACGAGAATAAACCAATCATTCGCTGAAACCAAAGTCTTTTTGTTGTC 117588

TeratornLC199500 TTTTATTCGCATTGGTACACAGTACA**ATGTCACACAAGAGAGAGCATTCTTTTATTTGTTTTTTTTTAAAAAAAAAAGAAAACACCACCACCAAAGAGACCACCACTCCCAAGGCCACC** 115773

WF-Teratorn TTTTATTCGCATTGGTACACAGTACAATGTCACACAAGAGAGAGCATTCTTTTATTTGTTTTTTTTA-AAAAAAAAAAGAAAACACCACCACCAAAGAGACCACCACTCCCAAGGCCACC 117707

TeratornLC199500 **CTTCACCACACACACCCACACCCCTACACACCATCCACCTCCCGGCAAAGCCGACCCACTCGGAGTTTAAACTAAAAAGACAGAGGGAAACGGAGAAGTTCGACACCGTTGCGACCCGCA** 115893

WF-Teratorn CTTCACCACACACACCCACACCCCTACACACCATCCACCTCCCGGCAAAGCCGACCCACTCGGAGTTTAAACTAAAAAGACAGAGGGAAACGGAGAAGTTCGACACCGTTGCGACCCGCA 117827

TeratornLC199500 **GACCGATCAACAAGCAAACGGGCATGCTCCGCAAATATTCGCAGAGAAGTCTTCTGGCCATCTGGGCTCCTCGATCGGCTCCGCCGTCGCCAGCCGTTCGAGAAAGCGAAATGATCGATC** 116013

WF-Teratorn GACCGATCAACAAGCAAACGGGCATGCTCCGCAAATATTCGCAGAGAAGTCTTCTGGCCATCTGGGCTCCTCGATCGGCTCCGCCGTCGCCAGCCGTTCGAGAAAGCGAAATGATCGATC 117947

TeratornLC199500 **GGGCTGCGCTTTCTTCTCGGACGCAGCCGAGCCAGTCTCCAGAATGCTCGCCCTCGTTGCTGCAAGGTGTCGTCCTTCTCCCCTTCTGCAAAAAAGTGGAAGCGGGAATTCAGAAAACTA** 116133

WF-Teratorn GGGCTGCGCTTTCTTCTCGGACGCAGCCGAGCCAGTCTCCAGAATGCTCGCCCTCGCTGCTGCAAGGTGTCGTCCTTCTCCCCTTCTGCAAAAAAGTGGAAGCGGGAATTCAGAAAACTA 118067

TeratornLC199500 **CTACTGAGACAGACCTCGAGCAGGGATCCGATCTGGACATGACCCCAGAGCTGGACCTATCTCAGCACAATCTTCCGAGCACTTACAAAAATACGAGCGAGAGCCACAAAGTGCAGGGGG** 116253

WF-Teratorn CTACTGAGACAGACCTCGAGCAGGGATCCGATCTGGACATGACCCCAGAGCTGGACCTATCTCAGCACAATCTTCCGAGCACTTACAAAAATACGAGCGAGAGCCACAAAGTGCAGGGGG 118187

TeratornLC199500 **GTTATCTCCCTCACCTCCAGGAAAAGGCTGAGAAGAAGATAAAGAAAACCACCCGCGAGAGCCAAAACTCGCAGACCTCTTCTACCGAGCCGTGCGGCCTTCGGAGCCGAGAACCGGACG** 116373

WF-Teratorn GTTATCTCCCTCACCTCCAGGAAAAGGCTGAGAAGAAGATAAAGAAAACCACCCGCGAGAGCCAAAACTCGCAGACCTCTTCTACCGAGCCGTGCGGCCTTCGGAGCCGAGAACCGGACG 118307

TeratornLC199500 **AAGTGGAAGCTCTTATCTCGGGGCCTCTCGTAAAAGCGGTGAAGATAGTCGTGTTCCATCTCCTCACCACCTGTTTGAGCAAGGTCCGTAAAGAAGAGTGCTACGGCTGTCAGGTCGACC** 116493

WF-Teratorn AAGTGGAAGCTCTTATCTCGGGGCCTCTCGTAAAAGCGGTGAAGATAGTCGTGTTCCATCTCCTCACCACCTGTTTGAGCAAGGTCCGTAAAGAAGAGTGCTACGGCTGTCAGGTCGACC 118427

TeratornLC199500 **ACCCCAGTCAGATGCAGCACCCCTGTCTGTTCGTAATCGAAGAGGGCTTTGTTTACTCCAACTTTCCTGCTCTGGTTAAGAAACTGCTGACTCCTGGATATCTTCCAGCCGTCAAAACGT** 116613

WF-Teratorn ACCCCAGTCAGATGCAGCACCCCTGTCTGTTCGTAATCGAAGAGGGCTTTGTTTACTCCAACTTTTCTGCTCTGGTTAAGAAACTGCTGACTCCTGGATATCTTCCAGCCGTCAAAACGT 118547

TeratornLC199500 **TCCTGTCCATGCGCAACATCGTAGAGAAGGACGAGAGGATCCGAGCAGTGACCGACGCTCTGCTTTACGAACTGAGACACAGCGCTTTGAACTCACACGGTGATCTCGTGTCGACTGTGT** 116733

WF-Teratorn TCCTGTCCATGCGCAACATCGTAGAGAAGGACGAGAGGATCCGAGCAGTGACCGACGCTCTGCTTTACGAACTGAGACACAGCGCTTTGGACTCACACGGTGATCTCGTGTCGACTGTGT 118667

TeratornLC199500 **ACGAAGGACTGACAGAGGCCGAGGAGATTGATTTCGAACAGCTGGAAGTTGTGTCTAAGGTTTGGGAAGGTGATCTTTCCGCCTGTCCCAATTGTTTAATTTGA**AGAAGAGATGCGAAAA 116853

WF-Teratorn ACGAAGGACTGACAGAGGCCGAGGAGATTGATTTCGAACAGCTGGAAGTTGTGTCTAAGGTTTGGGAAGGTGATCTTTCCGCCTGTCCCAATTGTTTAATTTGAAGAAGAGATGCGAAAA 118787

TeratornLC199500 ACACTGTGATGGCTTTCACAGTAAAAGTTTGTGTGTAACCTGTGGGGGGAAAAAAAAAAAAAAAACATCTCTCACAAGATGAAATGTGAAAACAAAGAAATAAAATTCTGTCTTGAAAGA 116973

WF-Teratorn ACACTGTGATGGCTTTCACAGTAAAAGTTTGTGTGTAACCTGTGGGGGGGAAAAAAAAAAAAA--CATCTCTCACAAGATGAAATGTGAAAACAAAGAAATGAAATTCTGTCTTGAAAGA 118905

TeratornLC199500 AGTGTGTGCTTCATCGTTCATCAAGTCAGACAAACCATTAGCCGTCAGACCTATATCTGGAGCCAGACAATTCTGCGAACCGGCTGCAAGAATTACATCTGAAGCCAGTCCTCGCAACTA 117093
WF-Teratorn AGTGTGTGCTTCATCGTTCATCAAGTCAGACAAACCATTAGCCGTCAGACCTATATCTGGAGCCAGACAATTCTGCGAACCGGCTGCAAGAATTACATCTGAAGCCAGTCCTCGCAACTA 119025

TeratornLC199500 AAAAAAAAAAAA--------CATTCCTCGTACTAGCAAGGCCAGAAGCCTTCGCTTGAAAAGGACAGGGTTCAGACTGCGTTCAGAGTTTCATTACCCGGTCCAACCAGGCCGCACGGT 117205
WF-Teratorn AAAAAAAAAAAAAAAAAAAACATTCCTCGTACTAGCAAGGCCAGAAGCCTTCGCTTGAAAAGGACAGGGTTCAGACTGCGTTCAGAGTTTCATTACCCGGTCCAACCAGGCCGCACGGT 119145

TeratornLC199500 GGAATACGGTCAAGACGAATGTGGTGTGATTATCTATGCCAACGTTTAAAACAGTCACGTACCAGAATGCCTATGTTTAAAACAGTCACGTACCAGAATAGGATCGTTGTGGAACGAGAA 117325
WF-Teratorn GGAATACGGTCAAGACGAATGTGGTGTGATTATCTATGCCAACGTTTAAAACAGTCACGTACCAGAATGCCTATGTTTAAAACAGTCACGTACCAGAATAGGATCGTTGTGGAACGAGAA 119265

TeratornLC199500 TAAATCAGTGACTCAAACCAAGGTGTGTTGTCTTTTATTCACATCAGCACAATTTCATACACGAGAGAACATCTGTTCCACAAGCAATTTCAACAAGCAAACTGCATGATTGTGCACAAG 117445
WF-Teratorn TAAATCAGTGACTCAAACCAAGGTGTGTTGTCTTTTATTCACATCAGCACAATTTCATACACGAGAGAACATCTGTTCCACAAGCAATTTCAACAAGCAAACTGCATGATTGTGCACAAG 119385

TeratornLC199500 CAAAGCTGTCTTTCTGGGTCCGAGGTAACAGTCGTCCTCCCCTCTTCGTGCGGGACTCCTTCCGTCTCTGCTG**CTACAACAGCCACCTATGCATGTCGAGTTGAGCCACGAACCTATCCC** 117565
WF-Teratorn CAAAGCTGTCTTTCTGGGTCCGAGGTAACAGTCGTCCTCCCCTCTTCGTGCGGGACTCCTTCCGTCTCTGCTGCTACAACAGCCACCTATGCATGTCGAGTTGAGCCACGAACCTATCCC 119505

TeratornLC199500 **CAGTCCGGATCGAGCTGGCGATGTGCCCTTTCCGAGCTCGTAACACCTGTCCTCCCGCGAAACCCTGCACACTTTCCACGACGGTCATCGCGTGGCCGATCACAGTCTTGAGGAATCGCT** 117685
WF-Teratorn CAGTCCGGATCGAGCTGGCGATGTGCCCTTTCCGAGCTCGTAACACCTGTCCTCCCGCGAAACCCTGCACACTTTCCACGACGGTCATCGCGTGGCCGATCACAGTCTTGAGGAATCGCT 119625

TeratornLC199500 **TCGCCTCTTGGTCCTGCACTAGCCGAGCTGCCAGGTTCGAGTCGAATCTGTCGTTCACCAAAGACGCGGCCGCGTTTCGTGTGACAATGTCCACTCGATGGTAGTCTGCTCCGTTGGCCA** 117805
WF-Teratorn TCGCCTCTTGGTCCTGCACTAGCCGAGCTGCCAGGTTCGAGTCGAATCTGTCGTTCACCAAAGACGCGGCCGCGTTTCGTGTGACAATGTCCACTCGATGGTAGTCTGCTCCGTTGGCCA 119745

TeratornLC199500 **GGAGGGTCATTATGGCCAGCACGCTCTGGGGGTGGTAAGGGGAGACCAGCTGGAACACGTCGAAGTCACCGTCATGCTTGAGCGCCTGCACGACCTCCTCCACGGGATCTCCTCCGCAAT** 117925
WF-Teratorn GGAGGGTCATTATGGCCAGCACGCTCTGGGGGTGGTAAGGGGAGACCAGCTGGAACACGTCGAAGTCACCGTCATGCTTGAGCGCCTGCACGACCTCCTCCACGGGATCTCCTCCGCAAT 119865

TeratornLC199500 **CGCAATTTACGCGTCTGGTCGCGAGCAGACCGCCCCAGATGCCCTTGTTCACGTTGAGATCGTAGCCCGGGTCCCCCTCGACTCCGAGCTGCGCCGCCACGTCTTTTCTGTCCGGGAGCT** 118045
WF-Teratorn CGCAATTTACGCGTCTGGTCGCGAGCAGACCGCCCCAGATGCCCTTGTTGACGTTGAGATCGTAGCCCGGGTCCCCCTCGACTCCGAGCTGCGCCGCCACGTCTTTTCTGTCCGGGAGCT 119985

TeratornLC199500 **GAACAGAGAACCTGTCGCCCTGTTTCAACGTTTCGTGGCCCGTGTTGATCAGCTCCACTTCGTACAAGTGGGGAGCGTTCAGGAACGGTCTGTCCACGATGGAGCTGCTCGCGCAGCACA** 118165
WF-Teratorn GAACAGAGAACCTGTCGCCCTGTTTCAACGTTTCGTGGCCCGTGTTGATCAGCTCCACTTCGTACAAGTGGGGAGCGTTCAGGAACGGTCTGTCCACGATGGAGCTGCTCGCGCAGCACA 120105

TeratornLC199500 **CGGCCACCTCCTTGAGGCGGCCAACGGATTCGAGGCTGAGGCCGTGTAGAGTGGGAACGTAGCGGTTGCTGTCTTTGACAAAGTGGAACAGGCCCGGAGGGAGGTAGTTTTTCGAGGAGT** 118285
WF-Teratorn CGGCCACCTCCTTGAGGCGGCCAACGGATTCGAGGCTGAGGCCGTGTAGAGTGGGAACGTAGCGGTTGCTGTCTTTGACAAAGTGGAACAGGCCCGGAGGGAGGTAGTTTTTCGAGGAGT 120225

TeratornLC199500 **CGTACTTGTAAGCTCCGTAGGGGACAGACTGGCTGCGAGGCTTGCTCTCGCTCAAGCCGTGTGTGTAAGCGCGGATCGGCTCGTGTATCCTGGTCGACGACAT**CTGCGTTTGCGGCTGAA 118405
WF-Teratorn CGTACTTGTAAGCTCCGTAGGGGACAGACTGGCTGCGAGGCTTGCTCTCGCTCAAGCCGTGTGTGTAAGCGCGGATCGGCTCGTGTATCCTGGTCGACGACATCTGCGTTTGCGGCTGAA 120345

TeratornLC199500 AACGAAAAAAAAAAAAAAAAAAA-CAAGCAACGACAACAA**CTAGTCCGCGATCGGACGACAGAGAAACTCGAGCGCTCGCAAAGGCTCTCTGCAGGTGGGCGCGTGCGGGGCGTTGGGCG** 118524
WF-Teratorn AACGAAAAAAAAAAAAAAAAAAAACAAGCAACGACAACAACTAGTCCGCGATCGGACGACAGAGAAACTCGAGCGCTCGCAAAGGCTCTCTGCAGGTGGGCGCGTGCGGGGCGTTGGGCG 120465

TeratornLC199500 **CTTTCAGGACGATCAGAGCCGCGGCGAGCACCTCCAGTAGCACAGGACCCTCTTCTACGTGAACTCTGAGCAGCCTGACCGTAAGCATGTTGAGATTCTCCTCCAAGAGCTCCCCGAATG** 118644
WF-Teratorn CTTTCAGGACGATCAGAGCCGCGGCGAGCACCTCCAGTAGCACAGGACCCTCTTCTACGTGAACTCTGAGCAGCCTAACCGTAAGCATGTTGAGATTCTCCTCCGAGAGCTCCCCGAATG 120585

TeratornLC199500 **TCCGCGAGAACCACGGGCACCACTTGGTCACGATGTAAGATCTCAAATCACACCGGGACTCCGCTCGAGGGGGCTCCGGCTTCTCCACCGAGCCGTCCTCTCCTCTCGTTCGGGGGTTCG** 118764
WF-Teratorn TCCGCGAGAACCACGGGCACCACTTGGTCACGATGTAAGATCTCAAATCACACCGGGACTCCGCTTGAGGGGGCTCCGGCTTCTCCACCGAGCCGTCCTCTCCTCTCGTTCGGGGGTTCG 120705

TeratornLC199500 **TCGCGCGAGCCGCGACCCGGCTCGGCCCTCGAGCGGCGGACTTTGGCGAAGCGAGGAGTATCAGCTCTTCCAGCGTGAGTTCGTGCCACAT**GGTTCGAGCGAAAGGTCGGTCGCGACGAC 118884
WF-Teratorn TCGCGCGAGCCGCGACCCGGCTCGGCCCTCGAGCGGCGGACTTTGGCGAAGCGAGGAGTATCAGCTCTTCCAGCGTGAGTTCGTGCCACATGGTTCGAGCGAAAGGTCGGTCGCGACGAC 120825

TeratornLC199500 CGTAAAGAAGAGGAGATTTCTGGCTAGGCCGAGGGC**TCAGGCCGAGTAGTTCATCTCCACGCGTTTGCGGTAGGCCACTAGCCTCTCGTCGTAGAAACCCACCCGACCGACAGACTCTGC** 119004
WF-Teratorn CGTAAAGAAGAGGAGATTTCTGGCTAGGCCGAGGGCTCAGGCCGAGTAGTTCATCTCCACGCGTTTGCGGTAGGCCACTAGCCTCTCGTCGTAGAAACCCACCCGACCGACAGACTCTGC 120945

TeratornLC199500 **GAAGTTGCAGTTTGTCATCTGCAGCTTTTCTCTCCGAGTGACTCTGCTGAGCGCCACGTACACCGATGGTTTTATAGGGGACATGGAAAAGTTTCGCGGCGGGTGGTACAGCACCTCTCC** 119124
WF-Teratorn GAAGTTGCAGTTTGTCATCTGCAGCTTTTCTCTCCGAGTGACTCTGCTGAGCGCCACGTACACCGATGGTTTTATAGGGGACATGGAAAAGTTTCGCGGCGGGTGGTACAGCACCTCTCC 121065

TeratornLC199500 **GCTTATCGTACAGCCCTGACAGTCGTAGGTGTTCATGGCCAGCATGCAGGCCAACGGGTGCGTGGTCACCGTCCAGTTTTCGCAGTGCCACTTGGAAGGTCCCACTTTGTAAAGACGACC** 119244
WF-Teratorn GCTTATCGTACAGCCCTGACAGTCGTAGGTGTTCATGGCCAGCATGCAGGCCAACGGGTGCGTGGTCACCGTCCAGTTTTCGCAGTGCCACTTGGAAGGTCCCACTTTGTAAAGACGACC 121185

TeratornLC199500 **GGTCCTGTCTCCTCTCACGTAGAACTCGTAAGCCTGGGTCTCTTTCCGGAACGCCATCCCGCTGATCAGGCCTTCCACTTCGAGCGGCGAAACCGGGTCCCCGGCGAGGTTGGCCGCCGG** 119364
WF-Teratorn GGTCCTGTCTCCTCTCACGTAGAACTCGTAAGCCTGGGTCTCTTTCCGGAACGCCATCCCGCTGATCAGGCCTTCCACTTCGAGCGGCGAAACCGGGTCCCCGGCGAGGTTGGCCGCCGG 121305

TeratornLC199500 **CGAAAGGGCTTTACGAGGCACCGGTCCGTCCTCTACGGGCTGTATGCCGTTGTGGTCCATGTGCCAAAGCATCCTGACCTTGCAACCGACCCAGAGCTCGGGGTTCCTGAGGTAGTCCCT** 119484
WF-Teratorn CGAAAGGGCTTTACGAGGCACCGGTCCGTCCTCTACGGGCTGTATGCCGTTGTGGTCCATGTGCCAAAGCATCCTGACCTTGCAACCGACCCAGAGCTCGGGGTTCCTGAGGTAGTCCCT 121425

TeratornLC199500 **TTCGCGTGGGCGCTGGCCGCCCTTGAAAAACACCCTGGGCATCGCAAAGAGTTTGGGTAAAGCCTGAGTCGCCTGCTGAAGGACGTTGTACCACTGGGCGGGTCCCTTGCTCGATGATCC** 119604
WF-Teratorn TTCGCGTGGGCGCTGGCCGCCCTTGAAAAACACCCTGGGCATCGCAAAGAGTTTGGGTAAAGCCTGAGTCGCCTGCTGAAGGACGTTGTACCACTGGGCGGGTCCCTTGCTCGATGATCC 121545

TeratornLC199500 **GTGCTTGGTCACGCTCAGGAACACCTCCGCCTTTCGTGTTGCCCTGGCGGCGTAAGCCTTTGTGTAGGCAGCGAGCTGTTGGTCCTGGTGGAAGATGCGAGTCTTGTGAGGCTCGTGCTC** 119724
WF-Teratorn GTGCTTGGTCACGCTCAGGAACACCTCCGCCTTTCGTGTTGCCCTGGCGGCGTAAGCCTTTGTGTAGGCAGCGAGCTGTTGGTCCTGGTGGAAGATGCGAGTCTTGTGAGGCTCGTGCTC 121665

TeratornLC199500 **GGGGTCCATCGCCTGTCTCTCGCTGACGAGCAGCTGACTCCTGAAAATGGTCCGGGACTGCTGCGTCACCGTGTTGAACTGGAGGTAGGAGAGTGTCTCGGCGTAGCCCGGGTCGTCGAA** 119844
WF-Teratorn GGGGTCCATCGCCTGTCTCTCGCTGACGAGCAGCTGACTCCTGAAAATGGTCCGGGACTGCTGCGTCACCGTGTTGAACTGGAGGTAGGAGAGTGTCTCGGCGTAGCCCGGGTCGTCGAA 121785

TeratornLC199500 **CTGCCGCCTGTTGACAAAGAGCGGGGTGGAGCAGCACAGCAGCGTTTCCATGCGACCGGTCTCCCAGAGGAGCGGGCCCGAACCGGCTGGCTGCAACTGGGACACGCTGCCCGCCAGGAT** 119964
WF-Teratorn CTGCCGCCTGTTGACAAAGAGCGGGGTGGAGCAGCACAGCAGCGTTTCCATGCGACCGGTCTCCCAGAGGAGCGGGCCCGAACCGGCTGGCTGCAACTGGGACACGCTGCCCGCCAGGAT 121905

TeratornLC199500 **TATCAGAGGACCCGCGTCCGGACCGTAGAACAGGGCGAGTATTCTGAGCATCCTCTCCAGAAGCCTGGCCGTCATGAGCCCGTACTCGTCCACGACCAGCACGTTCGTGCCCAAGAACGG** 120084
WF-Teratorn TATCAGAGGACCCGCGTCCGGACCGTAGAACAGGGCGAGTATTCTGAGCATCCTCTCCAGAAGCCTGGCCGTCATGAGCCCGTACTCGTCCACGACCAGCACGTTCGTGCCCAAGAACGG 122025

TeratornLC199500 **AGGCGCGTCCGCCAAGGCGCTGCGGCCTCTCAGCCGAGCGAAGATCCTGCTGCAGGCGGAGCAGCTCTCTGGCCTCGGCAGCTCGCAGGAGTGTTTGTCGACGGCCCGGTCGACCTTCGA** 120204
WF-Teratorn AGGCGCGTCCGCCAAGGCGCTGCGGCCTCTCAGCCGAGCGAAGATCCTGCTGCAGGCGGAGCAGCTCTCTGGCCTCGGCAGCTCGCAGGAGTGTTTGTCGACGGCCCGGTCGACCTTCGA 122145

TeratornLC199500 **CCGGCGGAACGCCCGGAAGGCGAGGATCGCACCCAGGTAGCATCTCCTGTAGCGCTCGACAAAGGCGTCCGCTTCGAGCGAATCGTCCAGCAGATCCTTTTTGAGACCCATCATCTTGTG** 120324
WF-Teratorn CCGGCGGAACGCCCGGAAGGCGAGGATCGCACCCAGGTAGCATCTCCTGTAGCGCTCGACAAAGGCGTCCGCTTCGAGCGAATCGTCCAGCAGATCCTTTTTGAGACCCATCATCTTGTG 122265

TeratornLC199500 **AAACGTTTGCACCTTGTCCAGATGACCGGACTCCATCAGCGTTCCTGCCGCCTTGTTGGTCATGGCGCTCACTGCGACGCTGATCCTGAGGGCTTTCAGCTTTTTCTCCAGCAGACCGAT** 120444
WF-Teratorn AAACGTTTGCACCTTGTCCAGATGACCGGACTCCATCAGCGTTCCTGCCGCCTTGTTGGTCATGGCGCTCACTGCGACGCTGATCCTGAGGGCTTTCAGCTTTTTCTCCAGCAGACCGAT 122385

TeratornLC199500 **CGCGTACGATTTCCCGCAGCCCGCGTCTCCGCTCAGCAAAGCCACTTTCAGGCGATCGCCTCGAAACATGTGCAACTTCACCATGCGTCGAACCCACTCCTCCTGGCAGTATCCCGAAAG** 120564
WF-Teratorn CGCGTACGATTTCCCGCAGCCCGCGTCTCCGCTCAGCAAAGCCACTTTCAGGCGATCGCCTCGAAACATGTGCAACTTCACCATGCGTCGAACCCACTCCTCCTGGCAGTATCCCGAAAG 122505

TeratornLC199500 **AGAGAGCAAAGCCCGAGCCGCGGGGTCCTCGTCGGCGACGCGCGGTTCCTCTCTGGCGCGCTTGCGGGCACCGGGCGCGCCAAATCCCTCGGCCGGAGAGACGTCGACCCCCGGCTCGTC** 120684
WF-Teratorn AGAGAGCAAAGCCCGAGCCGCGGGGTCCTCGTCGGCGACGCGCGGTTCCTCTCTGGCGCGCTTGCGGGCACCGGGCGCGCCAAATCCCTCGGCCGGAGAGACGTCGACCCCCGGCTCGTC 122625

TeratornLC199500 **CAT**CTGTCTGTGCGGGGGCTGCGCAGTGTGACCG**TCAGAGAAACGGCTCGATCTGCACCCGCTTCGCGAGGAGCGAGCTTCGCTCTGGGCGGTGCGGAGCGGCAACCCGAAGCTGCTCGG** 120804
WF-Teratorn CATCTGTCTGTGCGGGGGCTGCGCAGTGTGACCGTCAGAGAAACGGCTCGATCTGCACCCGCTTCGCGAGGAGCGAGCTTCGCTCTGGGCGGTGCGGAGCGGCAACCCGAAGCTGCTCGG 122745

TeratornLC199500 **CTCGATCAAACCTCGAGGTCCTCGAGCCTCCGTTCCCGCCACTTAGTTCTCCTGGGTCTCTCTGTCTGCGTTTGATGAAGGCAGACAGAGAACGGTCAATGTTCAAGGAGTCTCGAGCGC** 120924
WF-Teratorn CTCGATCGAACCTCGAGGTCCTCGAGCCTCCGTTCCCGCCACTTAGTTCTCCTGGGTCTCTCTGTCTGCGTTTGATGAAGGCAGACAGAGAACGGTCAATGTTCAAGGAGTCTCGAGCGC 122865

TeratornLC199500 **CGAGGGATCCGTCCTCCTCCCGTTGCTTTGGAGAAAAGCTCGCGAAACGGGGGCTTTCTATGCCTTCCAGGACGGACCTGAGTACGCCGTTCCTCGCGCGGTTCGAACCCCTGTGGCGAT** 121044
WF-Teratorn CGAGGGATCCGTCCTCCTCCCGTTGCTTTGGAGAAAAGCTCGCGAAACGGGGGCTTTCTATGCCTTCCAGGACGGACCTGAGTACGCCGTTCCTCGCGCGGTTCGAACCCCTGTGGCGAT 122985

TeratornLC199500 **CGCTGTGTTCCTCAGGCAGCCCCGCGTCGACCGGGACCGTGACGGCGCCCAGAGACTCGCGAGCCTGCGCGGCTCTGCCGCGAACACAGAGAAAAGACTCGGATGCCCTGGCGACCGTTT** 121164
WF-Teratorn CGCTGTGTTCCTCAGGCAGCCCCGCGTCGACCGGGACCGTGACGGCGCCCAGAGACTCGCGAGCCTGCGCGGCTCTGCCGCGAACACAGAGAAAAGACTCGGATGCCCTGGCGACCGTTT 123105

TeratornLC199500 **CTCTGAACAGCGTCACGCTCGAGGAGTACAGGATCTGGCTCAGTTTCACGGCCACGGCGGTGACGCCGCCCTTCCCCGTGCTGTTGAGCACGGTCAACAGCCCGGCCAGCAGCTCGACGA** 121284
WF-Teratorn CTCTGAACAGCGTCACGCTCGAGGAGTACAGGATCTGGCTCAGTTCCACGGCCACGGCGGTGACGCCGCCCTTCCCCGTGCTGTTGAGCACGGTCAACAGCCCGGCCAGCAGCTCGACGA 123225

TeratornLC199500 **AGGGAGCCTGCGCCCCGTACTCGAACAGGAGCCAGGCGAAGTCTTCCGTGTCCACCTTGTCCGTGCACCTGTAGCCGTTGGCGTAGACGCGGGGCCTCGTGAAGCCAAAGTGTTTCGCGT** 121404
WF-Teratorn AGGGAGCCTGCGCCCCGTACTCGAACAGGAGCCAGGCGAAGTCTTCCGTGTCCACCTTGTCCGTGCACCTGTAGCCGTTGGCGTAGACGCGGGGCCTCGTGAAGCCAAAGTGTTTCGCGT 123345

TeratornLC199500 **ACCACTCCGGCTCTTTCTGAGAGAGCGCGATGTTGACGTTTTCCAGCAGTATGCTGAACCCCGCCGATCGCCTCTCGCCCATCCTCATCTCGAACAGTTTCTGCAACACCCCGACCGGCG** 121524
WF-Teratorn ACCACTCCGGCTCTTTCTGAGAGAGCGCGATGTTGACGTTTTCCAGCAGTATGCTGAACCCCGCCGATCGCCTCTCGCCCATCCTCATCTCGAACAGTTTCTGCAACACCCCGACCGGCG 123465

TeratornLC199500 **TCACGGACTCCCCGGGTCTGAACATGCGATCTTCGGGCAGAGACACCGCCAAGAAATGCCTGCCGACCCAGTCCAGGTGAGGATGAGAGCGAGGCGATTCGCTTTTACCCGCCTCGCCTT** 121644
WF-Teratorn TCACGGACTCCCCGGGTCTGAACATGCGATCTTCGGGCAGAGACACCGCCAAGAAATGCCTGCCGACCCAGTCCAGGTGAGGATGAGAGCGAGGCGATTCGCTTTTACCCGCCTCGCCTT 123585

TeratornLC199500 **CCGCGCTCCGGGCGGCTTTCTCCCTCAGCTCGATCTCGAGGGCTTTGCGGTCTTCCAAAGCGCGGTGCAGAGCTTGTTTGGCCTCCTCGAGATCCCTCCTGCAAACCTCCAGCTCGTAGG** 121764
WF-Teratorn CCGCGCTCCGGGCGGCTTTCTCCCTCAGCTCGATCTCGAGGGCTTTGCGGTCTTCCAAAGCGCGGTGCAGAGCTTGTTTGGCCTCCTCGAGATCCCTCCTGCAAACCTCCAGCTCGTAGG 123705

TeratornLC199500 **CCATGCGGACGACAGCCTCCTCGCCCGAGCGCCCCGCTCGCGCGAGGTTTCCGCTCGACTCTCGAACGCCTGACCGCAGGTCGACACGTTCATGGAGCGCTCCTCTATCCCGCTCCCTTT** 121884
WF-Teratorn CCATGCGGACGACAGCCTCCTCGCCCGAGCGCCCCGCTCGCGCGAGGTTTCCGCTCGACTCTCGAACGCCTGACCGCAGGTCGACACGTTCATGGAGCGCTCCTCTATCCCGCTCCCTTT 123825

TeratornLC199500 **CCGTTCTGTCCAGAGCGCCCTTCAGCTCTGCGATCTCGCGTCTCATCGAAGCGATCATGTTCAT**GTGAGAACACCGGTATTCGCAAGCGTCCTCATCTTCGGATTCTTCTCTACCCTGCT 122004
WF-Teratorn CCGTTCTGTCCAGAGCGCCCTTCAGCTCTGCGATCTCGCGTCTCATCGAAGCGATCATGTTCATGTGAGAACACCGGTATTCGCAAGCGTCCTCATCTTCGGATTCTTCTCTACCCTGCT 123945

TeratornLC199500 CTCCCTCGCCGCGATCTCTCTTCCGCCTGCGCGAGACCGTTGATCGTGTGAGCATGGCGATTCCTTTTGCGAGTGCGATTTTGGATTCCGAAGGATCGACCGGGGTGAAGTGTCTTGCTG 122124
WF-Teratorn CTCCCTCGCCGCGATCTCTCTTCCGCCTGCGCGAGACCGTTGATCGTGTGAGCATGGCGATTCCTTTTGCGAGTGCGATTTTGGATTCCGAAGGATCGACCGGGGTGAAGTGTCTTGCTG 124065

TeratornLC199500 TGTTATGTCAAAAGTCTGTTTGTATACTGTGCTATGACCAAAAACTGCTCTGATGCTAGTGGGGCCCGCGACAGAGACGCTGACCTGCGCAACGTTTTTCATTCCGTTGTGTCCGTTTGA 122244
WF-Teratorn TGTTATGTCAAAAGTCTGTTTGTATACTGTGCTATGACCAAAAACTGCTCTGATGCTAGTGGGGCCCGCGACAGAGACGCTGACCTGCGCAACGTTTTTCATTCCGTTGTGTCCGTTTGA 124185

TeratornLC199500 TGAAAAATAAAGTCGATCGCGACCGCTGCCGCCCTGTTGGACAGCTTCTCTGTGTGAAACAAACCAGTTTGCTCGAAAGGCCTGGGCGCTGCCAAAGACCCCTCTAACCTTCTCGCGGTC 122364
WF-Teratorn TGAAAAATAAAGTCGATCGCGACCGCTGCCGCCCTGTTGGACAGCTTCTCTGTGTGAAACAAACCAGTTTGCTCGAAAGGCCTGGGCGCTGCCAAAGACCCCTCTAACCTTCTCGCGGTC 124305

TeratornLC199500 GCACATCTGACTGTAACAAGGCAGGCTCGCACACAAACACGCACACCAGATCGGCTCCCCTCGTGAACACACGGCTCACGTCCAGGGCGCTGACGGGACGGCGATGGCCATGAAGCAGAA 122484
WF-Teratorn GCACATCTGACTGTAACAAGGCAGGCTCGCACACAAACACGCACACCAGATCGGCTCCCCTCGTGAACACACGGCTCACGTCCAGGGCGCTGACGGGACGGCGATGGCCATGAAGCAGAA 124425

TeratornLC199500 GGAACCCCCGGGTCGTGTGATGACAAAAAAAAAA-TCACAGTGTAAAATCAAGGTGGATTTTCTGTCCCATTTTTTTTTTAACTTTTATTTTACAGCCAGTTAAAACCTCACAGAACATT 122603
WF-Teratorn GGAACCCCCGGGTCGTGTGATGACAAAAAAAAAAAATCACAGTTAAAATCAAGGTGGATTTTCTGTCCCATTTTTTTTTAAACTTTTATTTTACAGCCAGTTAAAACCTCACAGAACATT 124545

TeratornLC199500 TGTGATAGCTTCAGGTGTGGTACTGGAAGGCCAACTCTTTCTCAATTCCGCTGCGCTCTGGGCAGTTTCGGTTCCATCTCTGGGTTGGATCCCTAGGACATGTTGTAATAAAAAAGTCAT 122723
WF-Teratorn TGTGATAGCTTCAGGTGTGGTACTGGAAGGCCAACTCTTTCTCAATTCCGCTGCGCTCTGGGCAGTTTCGGTTCCATCTCTGGGTTGGATCCCTAGGACATGTTGTAATAAAAAAGTCAT 124665

TeratornLC199500 ACAAAAAACAATTCCTGTGTTCATTTGGATTTTCTGTCCTGTTTTTA------TTTTTATTTTATTTTTT-ATTTTACAGCCAGTGAAAATCTCACAGAACATTCGCAATTGCTTCACGC 122836
WF-Teratorn ACAAAAAACAATTCCTGTGTTCGTTTGGATTTTCTGTCCTGTTTTTTTTTTATTTTTTATTTTATTTTTTTATTTTAAAGCCAGTGAAAATCTCACAGAACATTCGCAATTGCTTCACGC 124785

TeratornLC199500 GAAGAGCTAGATGACCGACTTTCTCTCTCCTTGCTGTTGAGCTGTGGGCCGTTTCAGTTCCCTCTCCGGGCTGGCTCCGCTTTTATGCCTGACAGCTCGATTGAGAAAACCTTTCCAAAT 122956
WF-Teratorn CAAGAGCTAAATGACCGACTTTCTCTCTCCTTGCTGTTGAGCTGTGGGTCGTTTCAGTTCCCTCTCCGGGCTGGCTCCGCTTTTATGCCTGACAGCTCGATTGAGAAAACCTTTCCAAAT 124905

TeratornLC199500 GCAACGTTCCAAATGTAAAGCAACCCCAAGAGAGCCCAGCCCAAGAGCGTTCCGATCACCTCCCCTCTGACAGCACATGAAGCTCCGTGGAGAGGCTCAGGTTTGGGTGAAGGGAACTGC 123076
WF-Teratorn GCAACGTTCCAAATGTAAAGCAACCCCAAGAGAGCCCAGCCCAAGAGCGTTCCGATCACCTCCCCTCTGACAGCACATGAAGCTCCGTGGAGAGGCTCAGGTTTGGGTGAAGGGAACCGC 125025

TeratornLC199500 ACGAGCGGCTCTGCGTGTAGTCACACAACAGAGCTCTGTGTCATGGGGGG-CCGACGCAGAGGAAGGCTGGAGCCTTTCGCTGAGACGAGTCACACTCCACGGTGCTCTGCGCCCTGTCG 123195
WF-Teratorn ACGAGCGGCTCTGCGTGTAGTCACACAACAGAGCTCTGTGTCATGGGGGGGCCGACGCAGAGGAAGGCTGGAGCCTTTCGCTGAGACGAGTCACACTCCACGGAGCTCTGCGCCCTGTCG 125145

TeratornLC199500 AAAGGGGCAGACTCAGAGGTGGGCTGCAATAGCTTCAGATGTGGGGCTGGAAAGCCAACTCTTTGTTCATTCCGCTGAGCTGTGGACCTTTTCGGTTCCATCCCCGGGTTGGGTCCCTTT 123315
WF-Teratorn AAGGGGGCAGACTCAGAGGCGGGCTGCAATAGCTTCAGATGTGGGGCTGGAAAGCCAACTCTTTGTTCATTCCGCTGAGCTGTGGACCTTTTCGGTTCCATCCCCGGGTTGGGTCCCTTT 125265

TeratornLC199500 ACTCCAGTAGCTTTGGACTGGGGGAACTGACAGGCATAAGGCTGATGCGGACCTGTTGAGAGGGTGGAGGTTGAAGGGTGGTCAAGATATTTGGAGGTGACATTACAAGAGGTAAAGTGG 123435
WF-Teratorn ACTCCAGTAGCTTTGGACTGGGGGAACTGACAGGCATAAGGCTGATGCGGCCCTGTTGAGAGGGTGGAGGTTGAAGGGTGGTCAAGATATTTGGAGGTGACATTACAAGAGGTAAAGTGG 125385

TeratornLC199500 TTAAAGATTATCTGAAATGAATGGAGGGTGGGGATAGTGAGAGGGAGCAGCAGAAACATAATCCTGGTAAAAGGGGGGACAGGGGTGGTCATCATTTTGGGACAAACATAGGCGTGTGTG 123555
WF-Teratorn TTAAAGATTATCTGAAATGAATGGAGGGTGGGGATAGTGAGAGGGAGCAGCAGAAACATAATCCTGGTAAAAGGGGGGACAGGGGTGGTCATCATTTTGGGACAAACATAGGCGTGTGTG 125505

TeratornLC199500 ATCCTGAGTAAAGGGACCTGGGTGTGAAAAATCCTGACTGAACCTATGTGGGTGAACCGGATCTGGGGAATAGCTTGAACCTGTGACCTGGCCTGGGTGAG**GTGGCGACTGGTACAAGCT** 123675
WF-Teratorn ATCCTGAGTAAAGGGACCTGGGTGTGAAAAATCCTGACTGAACCTATGTGGGTGAACCGGATCTGGGGAATAGCTTGAACCTGTGGCCTGGCCTGGGTGAGGTGGCGACTGGTACAAGCT 125625

TeratornLC199500 **TGAACCTGTGGCCTGGCCTGGGTGAACCGGATCTGGGAAATAGCTTGAACCTGTGGCCTGGCCTGGGTAAGGTGGTGGCTGGTACGAGCTTGAACCTGTGGCCTGGTCTGGGTAAGGTGG** 123795
WF-Teratorn TGAACCTGTGGCCTGGCCTGGGTGAACCGGATCTGGGAAATAGCTTGAACCTGTGGCCTGGCCTGGGTAAGGTGGTGGCTGGTACGAGCTTGAACCTGTGGCCTGGTCTGGGTAAGGTGG 125745

| | | |
|---|---|---|
| TeratornLC199500 | **TGGCTGGTACAAGCTTGAACCTGTGGCCTGGCCTGGGTGAACCGGATCTGGGAAATAGCTTGAACCTGTGGCCTGGCCTGGGTGAACCGGATCTGGGAAATAGCTTGAACCTGTGGCCTG** | 123915 |
| WF-Teratorn | TGGCTGGTACAAGCTTGAACCTGTGGCCTGGCCTGGGTGAACCGGATCTGGGAAATAGCTTGAACCTGTGGCCTGGCCTGGGTGAACCGGATCTGGGAAATAGCTTGAACCTGTGGCCTG | 125865 |
| TeratornLC199500 | **GCCTGGGTAAGGTGGTGGCTGGTACGAGCTTGAACCTGTGGCCTGGTCTGGGTGAGGTGGTGGCTGGTACGAGCTTGAACCTGTGGCCTGGTCTGGGTGAGGTGGTGGCTGGTACGAGCT** | 124035 |
| WF-Teratorn | GCCTG-------TGGT-------------------------------------------------------------------------------------------------------- | 125874 |
| TeratornLC199500 | **TGAACCTGTGGCCTGGTCTGGGTCAGGTGGTGGCTGGTACGAGCTTGAACCTGTGGCCTGGTCTGGGTCAGGTGGTGGCTGGTACGAGCTTGAACCTGTGGCCTGGTCTGGGTCAGGTGG** | 124155 |
| WF-Teratorn | ------------------------------------------------------------------------------------------------------------------------ | 125874 |
| TeratornLC199500 | **TGGCTGGTACGAGCTTGAACCTGTGGCCTGGTCTGGGTGAGGTGGTGGCTGGTACGAGCTTGAACCTGTGGCCTGGTCTGGGTCAGGTGGTGGCTGATACGATTGTCTTCTTGCAAGAAA** | 124275 |
| WF-Teratorn | ------------------------------------------------------------------------------------------------------------------------ | 125874 |
| TeratornLC199500 | **TTGTGTGA**CCATTGTCATCATCTTCATTTCGTGGGCGTGCTGGGCCGCTTGCAATTTCAGCTGTAGCTCTGTGTCATCTGCCTGAGACAGGGCTTCTTTGATTGTCCTAGAAAAAACATC | 124395 |
| WF-Teratorn | ------------------------------------------------------------------------------------------------------------------------ | 125874 |
| TeratornLC199500 | CATTGTGGATTCTAGCTTGTTTTTGGGCGTTTGACGGGTGGTAGGTGCACCTGTAAAATGACAAATGTTTTACATAATGAGCTATTTCTTTGTTGAGTGCTTTATCAATTACTACATTAT | 124515 |
| WF-Teratorn | ------------------------------------------------------------------------------------------------------------------------ | 125874 |
| TeratornLC199500 | GAGTATTGCCCAATAACATTATAGCTGTATTGCTGTAATATTGATTTTTATGTAAATAAAAAAAAAGTACTTAACACTGTGAATGCAAATGAAATGACCTACTTGTAAAAGACAAAATGA | 124635 |
| WF-Teratorn | ------------------------------------------------------------------------------------------------------------------------ | 125874 |
| TeratornLC199500 | GTATGACTAACTGTGATCACGTTTGTCATGAGTCTATGATAATAAACATAGAAATGCAATATATAAAATGGAAGTAATAGCCTATAGTGTATTATATAATATTGAAGCATTGTTTGAACT | 124755 |
| WF-Teratorn | ------------------------------------------------------------------------------------------------------------------------ | 125874 |
| TeratornLC199500 | ATAAATACCTGTTAGCTATCAATCATACTTTGTTATGAATGATGGTAGTTGTAATATTTTAATTGCTTACATATTTTTCTAATCTGTCGAATACTAAGACAACTTTTAAATTGCTATACT | 124875 |
| WF-Teratorn | ------------------------------------------------------------------------------------------------------------------------ | 125874 |
| TeratornLC199500 | ATTATAAATACAATAAAATAACATTAACAACATAAACGAAGAT**TTAAAATAAATTCCGTTTGCCCTTACCTGTACTGTTACTGTTCTCAAACTCCGGCTCTTCACTGTCCACCGATGACG** | 124995 |
| WF-Teratorn | ------------------------------------------------------------------------------------------------------------------------ | 125874 |
| TeratornLC199500 | **GCACATCATCAACTCTGGTGGTATCCAGTAATTCCGGTGGTCGGAAACTTGGCCTGTCACCCAAAATGCCTTCCATTTTATCATAGAATAGGAATGTTTCACAGCTGTTTCCACTCTTAT** | 125115 |
| WF-Teratorn | -------------------------------TTCCGGTGGTCGGAAACTTGGCCTGTCACCCAAAATGCCTTCCATTTTATCATAGAATAGGAATGTTTCACAGCTGTTTCCACTCTTAT | 125963 |
| TeratornLC199500 | **TATTGTGGTCTTTTACCGATCTGTATTTGTATTTCAAGTTTTTGATCTTTTGCCGGCAATCTAGCCACGAGCGCTTAAAACCTTGCCTCCCCATATTTGTTGAAATGTCTTCAAACAAGA** | 125235 |
| WF-Teratorn | TATTGTGGTCTTTTACCGATCTGTATTTGTATTTCAAGTTTTTGATCTTTTGCCGGCAATTTAGCCACGAGCGCTTAAAACCTTGCCTCCCCATATTTGTTGAAATGTCTTCAAACAAGA | 126083 |
| TeratornLC199500 | **ATTTGTTATGATAATGATCCTGCAATTCCGTCTGCATTGTCTCGTCGCCCCACAGACTTAAGAGGCATTTAACCTCATCATCGCTCCACCTTGTGGACTTTCCGACCTCCAT**CACTCTTG | 125355 |
| WF-Teratorn | ATTTGTTATGAAAATGATCCTGCAATTCCGTCTGCATTGTCTCGTCACCCCACAGACTTAAGAGGCATTTAACCTCATCATCGCTCCACCTTGTGGACTTTCCGACCTCCATCACTCTTG | 126203 |
| TeratornLC199500 | TGGCACACGCAAATGCACACAGTCGATTCGTTCCTGATTTATTTCCACATATGAGAGAGGCCTGTATCCGACCAGCCTGCCTCCTGAGCGTGCCTCTGTCGATTCGTTTCTGATTTATTT | 125475 |
| WF-Teratorn | TGGCACACGCAAATGCACACAGTCGATTCGTTCCTGATTTATTTCCACATATGAGAGAGGCCTGTATCCGACCAGCCTGCCTCCTGAGCGTGCCTCTGTCGATTCGTTTCTGATTTATTT | 126323 |
| TeratornLC199500 | CCACATATGAGAGAGGCCTGTATCCGACCAGCCTGCCTCCTGAGCGTGCCTCTCGCGACAGACGTGGAGCTCCGAGGCTGTGTGGCTCTGGTTGCGAAAGGCTCCAGCCTGCCCCCACCG | 125595 |
| WF-Teratorn | CCACATATGAGAGAGGCCTGTATCCGACCAGCCTGCCTCCTGAGCGTGCCTCTCGCGACAGACGTGGAGCTCCGAGGCTGTGTGGCTCTGGTTGCGAAAGGCTCCAGCCTGCCCCCACCG | 126443 |
| TeratornLC199500 | AGAGGAAGTCTCCAGCCTGCCACTGAGTTGGCCTCTCCTGACAGACTCAGCTGTGTGGCTCGCGTTGCGTAAGGCTGCAGCCTGCCTTTGACTTGGTCCCGCCTGACACAACACTCCACG | 125715 |
| WF-Teratorn | AGAGGAAGTCTCCAGCCTGCCACTGAGTTGGCCTCTCCTGACAGACTCAGCTGTGTGGCTCGCGTTGCGTAAGGCTGCAGCCTGCCTTTGACTTGGTCCCCCCTGACACAGCACTCCACG | 126563 |
| TeratornLC199500 | CAGAGCCGCTCGTGCTTTTGACTTTGACAAACCCCGCCTTTCCGTCGAGCTTCTCGTTCCGTCCGACGGGAGCGGATCGGCCTGCGGGCCGCTCTCAGACGATGTAGGATCCAGCCTGCC | 125835 |
| WF-Teratorn | CAGAGCCGCTCGTGCTTTTTACTTTGACAAACCCCGCCTTTCCGTCGAGCTTCTCGTTCCGTCCGACGGGAGCGGATCGGCCTGCGGGCCGCTCTCAGACGATGTAGGATCCAGCCTGCC | 126683 |
| TeratornLC199500 | TCTCGGTCTGCTCCCCCGCAGGCACGGGCCGCGTGTGTGCCTCAGACGCGAAAGGCTCCAGCCTTCCTCTGCTTCGGCCCCCCATGACA----GCTCTGTGGTGTGACTCGCCGCAGAGC | 125951 |
| WF-Teratorn | TCTCGGTCTGCTCCCCCGCAGGCACGGGCCGCGTGTGTGCCTCAGACGCGAAAGGCTCCAGCCTTCCTCTGCTTCGGCCCCCCATGACACAGAGCTCTGTGGTGTGACTCGCCGCAGAGC | 126803 |
| TeratornLC199500 | TGCTTGTGCTTTTCCGCTTAGGCCCGCTCAGGAGGCAGGCTGGAGACTTCCTCTCGGTGGGGGCAGGCTGGAGCCTTTCGCTGGCACGACTCACACTCCACGGAGCACTGCGTCTGTCAC | 126071 |
| WF-Teratorn | TGCTTGTGCTTTTCCGCTTAGGCCCGCTCAGGAGGCAGGCTGGAGACTTCCTCTCGGTGGGGGCAGGCTGGAGCCTTTCGCTGGCACGACTCACACTCCACGGAGCACTGCGTCTGTCAC | 126923 |

TeratornLC199500 GAGAGGCCCGCTCAGGGGCAGGCTGGAGACTTCCTCTCAGACAAGGAACACTCCACGGAGCTCTGCGTCCTGTCAGGAGAGGCAGACTCAGAGGCGGGTTGCAATTGCTTCAGATGTTTA 126191
WF-Teratorn GAGAGGCCCGCTCAGGGGCAGGCTGGAGACTTCCTCTCAGACAAGGAACACTCCACGGAGCTCTGCGTCCTGTCAGGAGGGGCAGACTCAGAGGCGGGTTGCAATTGCTTCAGATGTTTA 127043

TeratornLC199500 TTTTTATTTTTTATTTTTATTTTTGTTTTACAGCTAGAGAAAAACCTTTCCCAAAACGAGACAAAGCTCGGCGGGGTCTCTGAGCCTGCTCTGAAGCATCTCTGTTCCCCGTTGTACGCC 126311
WF-Teratorn TTTTTATTTTTTATTTTTATGTTTGTTTTACAGCTAGAGAAAAACCTTTCCCAAAACGAGACAAAGCTCGGCGGGGTCTCTGAGCCTGCTCTGAAGCATCTCTGTTCCCCGTTGTACGCC 127163

TeratornLC199500 CACAAAAACAAAACACAAATCTGGGTTTGATATTGTTTCGCTTATTCGTGTCTTTATTTTGTATATTTTACAGCCTGTGACAACTACAAAGAATAATTGCAAAAGTT**TCAGGTGTTCGAC** 126431
WF-Teratorn CACAAAAACAAAACACAAATCTGGGTTTGATATTGTTTCGCTTATTCGTGTCTTTATTTTGTATATTTTACAGCCTGTGACAACTACAAAGAATAATTGCAAAAGTTTCAGGTGTTCGAC 127283

TeratornLC199500 **CCACTCTTGGTGCTCGTAGACCCAATTCTTCTGACCGACTGGGGGGTCCTCATGGACAGATACAGGGCTGGGGTGGTGATGCATTGCATGTTCCTTGCGGCCAAGCTCACGTTGGTCACC** 126551
WF-Teratorn CCACTCTTGGTGCTCGTAGACCCAATTCTTCTGACCGACTGGGGGGTCCTCATGGACAGATACAGGGCTGGGGTGGTGATGCATTGCATGTTCCTTGCGGCCAAGCTCACGTTGGTCACC 127403

TeratornLC199500 **AGGAATAATTTCAGCTGATTGTGTACCAGGGTTTTAGTGAGAGAGTAGTAAATGCTGACAGCGATCCAGGGAGTCCAGCAGCAAACAAAAACTATGACAGTGAGCAATGAAAGGATGTAG** 126671
WF-Teratorn AGGAATAATTTCAGCTGATTGTGTACCAGGGTTTTAGTGAGAGAGTAGTAAATGCTGACAGCGATCCAGGGAGTCCAGCAGCAAACAAAAACTATGACAGTGAGCAATGAAAGGATGTAA 127523

TeratornLC199500 **GATTTTGCAAGGGGAAGCTGTTCTCTCTTTAAGTGAGCCGACCCCCTTCTAACATGTCTTCGCACAGTGAAGAAGATTGCGCCGTAACAAGCAGCCAGAATGATGCAGGGTGCGATAAAA** 126791
WF-Teratorn GATTTTGCAAGGGGAAGCTGTTCTCTCTTTAAGTGAGCCGACCCCCTTCTAACATGTCTTCGCACAGTGAAGAAGATTGCGCCGTAACAAGCAGCCAGAATGATGCAGGGTGCGATAAAA 127643

TeratornLC199500 **CAGATGGTCATCCTGATCCACGTGATGACGAACCTTTCGCTCGGGGAATAGGTGTCAGCGCACAGCGTAATCAGACCGACTTTGAAGGTTTCGTGATTGGATATGTTCACCAGAGCTGAC** 126911
WF-Teratorn CAGATGGTCATCCTGATCCACGTGATGACGAACCTTTCGCTCGGGGAATAGGTGTCAGCGCACAGCGTAATCAGACCGACTTTGAAGGTTTCGTGATTGGATATGTTCACCAGAGCTGAC 127763

TeratornLC199500 **ACGGAGGGAGGAATGGCACACACCAGGGACAGGAAAGCTCTCCTCGCCCACTCTCCTTTCACCCTCTTAAGAGAGGAGAAGGCAGCTTTTCTGAGGACCACCCTGTTGTATCTCTCTACA** 127031
WF-Teratorn ACGGAGGGAGGAATGGCACACACCAGGGACAGGAAAGCTCTCCTCGCCCACTCTCCTTTCACCCTCTTAAGAGAGGAGAAGGCAGCTTTTCTGAGGACCACCCTGTTGTATCTCTCTATA 127883

TeratornLC199500 **CTGATGCAGACCATCATGAGTACCCCGTATTGCAGGCAGAA**CTGTGGAGGAAACAAATACGCAAACAAAAAAAAAACCAAAACAGGTAATGAGGAGGGATTGAAGAAAGCAGAGGTTTTT 127151
WF-Teratorn CTGATGCAGACCATCATGAGTACCCCGTATTGCAGGCAGAACTGTGGAGGAAACAAATACGCAAACAAAAAAAAAAACAAAACAGGTAATGAGGAGGGATTGAAGAAAGCAGATGTTTTT 128003

TeratornLC199500 CATTTGTCCGGATAGAGAAAAGAACAGGAAAAAGCAAAGGCGACTCAC**AGTTATCACCCCCTTCGTGGCTTTGCAGACGACATATCCGAAGATCCATTCTTTGGCCATGTTCTGCATGAC** 127271
WF-Teratorn CATTTGTCCGGATAGAGAAAAGAACAGGAAAAAGCAAAGGCGACTCACAGTTATCACCCCCTTCGTGGCTTTGCAGACGACATATCCGAAGATCCATTCTTTGGCCATGTTCTGCATGAC 128123

TeratornLC199500 **CCACAGCGGCGTCGCGGACAATGTGCCTATGTCGCAGAGCCCGACGACCACGAAAAAGTAATCGCAGCGCGAACGAGCCTTCGGAGGTCTGCACAGCCGTAATCCCAGGTTTATCATGCA** 127391
WF-Teratorn CCACAGCGGCGTCGCGGACAATGTGCCTATGTCGCAGAGCCCGACGACCACGAAAAAGTAATCGCAGCGCGAACGAGCCTTCGGAGGTCTGCACAGCCGTAATCCCAGGTTTATCATGCA 128243

TeratornLC199500 **AACGTTTACGATCAAGCCCAGCACCATCACCGCAAAGTGAAGCATGGGTACCACACTGAACGCGATCGAAGAGTGAATCTCAGAGGACGCGTTGGTTGGTGACGGGCCAGTAGCATTGTA** 127511
WF-Teratorn AACGTTTACGATCAAGCCCAGCACCATCACCGCAAAGTGAAGCATGGGTACCACACTGAACGCGATCGAAGAGTGAATCTCAGAGGACGCGTTGGTTGGTGACGGGCCGGTAGCATTGTA 128363

TeratornLC199500 **CGTTGTCCTTTTGCCAAACAT**CGTCGCAGCTGACCTGAGGGACAGTGAACACTAGCACAGACAGAGAGACGTACGCTTGTGGTGCTGAAATAAGCTCCTCTCTCCCTAGCTCCAAGCCCC 127631
WF-Teratorn CGTTGTCCTTCTGCCAAACATCGTCGCAGCTGACCTGAGGGACAGTGAACACTAGCACAGACAGAGAGACGTACGCTTGTGGTGCTGAAATAAGCTCCTCTCTCCCTAGCTCCAAGCCCC 128483

TeratornLC199500 CGACTTAGACTGCCTTGCTGTATGGACATCCAACGGCGAGCCACACTCCCCCCACCTCACTCAGTCTCCGAACGGAGCAAACGGAGTACGGCCGACTCAGCTCGGAGTCGGACCTCAGCG 127751
WF-Teratorn CGACTTAGACTGCCTTGCTGTATGGACATCCAACGGCGAGCCACACTCCCCCCACCTCACTCAGTCTCCGAACGGAGCAAACGGAGTACGGCCGACTCAGCTCGGAGTCGGACCTCAGCG 128603

TeratornLC199500 TGTGGAGCCAACTAGTTGGCTAACTCTAGTTCCATAGCTGCTCAAAGTCTTGCGCTCTTGTCTAGGCGATTTGTGTGCCTCTTGCCTCAGTCGAGACAAGCGAAATCTCGATAACCTATT 127871
WF-Teratorn TGTGGAGCCAACTAGTTGGCTAACTCTAGTTCCATAGCTGCTCAAAGTCTTGCGCTCTTGTCTAGGCGATTTGTGTGCCTCTTGCCTCAGTCGAGACAAGCGAAATCTCGATAACCTATT 128723

TeratornLC199500 GTGCGAGCGCTGGCTGTGGCCCTTAGAAACCTGGAACTCGATCTTGAGCACACTTTACACTCATC**ATGGAAGTTACTCAGCAGACGTACGTGTCGTGTCCCGCATTGGACTTTGGTTTGG** 127991
WF-Teratorn GTGCGAGCGCTGGCTGTGGCCCTTAGAAACCTGGAACTCGATCTTGAGCACACTTTACACTCATCATGGAAGTTACTCAGCAGACGTACGTGTCGTGTCCCGCATTGGACTTTGGTTTGG 128843

TeratornLC199500 **CAGCGACAGACACGAAAGACCTCAGAAAAGGCTTGGAGGATCAGCTTCGCCTTCTCGACCTGTTCGACTCAAGCGCAGGACTTGACCGCTTTCTGTTGTCATCTTCCGGCTTGTTTTGGG** 128111
WF-Teratorn CAGCGACAGACACGAAAGACCTCAGAAAAGGCTTGGAGGATCAGCTTCGCCTTCTCGACCTGTTCGACTCAAGCGCAGGACTTGACCGCTTTCTGTTGTCATCTTCCGGCTTGTTTTGGG 128963

TeratornLC199500 **AGTACCGAAATCGCAGGTCAGAGGCTGCTGGCTGGTTCGCGGACAGTATAAGGCGACTGAACGTCGACACCTCAGACGTTCGTTCACTGTTTTTGGGCTCGTTGAACGTCATGGACAGAT** 128231
WF-Teratorn AGTACCGAAATCGCAGGTCAGAGGCTGCTGGCTGGTTCGCGGACAGTATAAGGCGACTGAACGTCGACACCTCAGACGTTCGTTCACTGTTTTTGGGCTCGTTGAACGTCATGGACAGAT 129083

TeratornLC199500 **ACGTCGCCACGCGTGACACTGAGTCCGAGCTCGTGGAGCTTTTGACGAACTCGGAAAATGTCCGAGCGGCCTGCTTCTCGCTCTCTGCGAAGATGCACGCCACGTGCTCTCAAATCGATC** 128351
WF-Teratorn ACGTCGCCACGCGTGACACTGAGTCCGAGCTCGTGGAGCTTTTGACGAACTCGGAAAATGTCCGAGCGGCCTGCTTCTCGCTCTCTGCGAAGATGCACGCCACGTGCTCTCAAATCGATC 129203

TeratornLC199500 **TGGACGACATCTGCGTCCGCAATCGCGTCTGGGCCAACTCCACCGTCAAATCCCAGCCGGGGCCGAACCTGACGGTCGCCGACGCATCCTCTACTGGTGAGGGGATACTGAGAGCGGAGA** 128471
WF-Teratorn TGGACGACATCTGCGTCCGCAATCGCGTCTGGGCCAACTCCACCGTCAAATCCCAGCCGGGGCCGAACCTGACGGTCGCCGACGCATCCTCTACTGGTGAGGGGATACTGAGAGCGGAGA 129323

TeratornLC199500 **AGGACGTATTGATCGCTCTCGGAGGAGCTGCGTTCCCTGTGAGGGAAGAGGACATTGTGCGTCTCTTGGTGAGGTTTCTTGCGCTGAAGCGTTCTAAGCGCGTTGCATTCCAAGAGCCGC** 128591
WF-Teratorn AGGACGTATTGATCGCTCTCGGAGGAGCTGCGTTCCCTGTGAGGGAAGAGGACATTGTGCGTCTCTTGGTGAGGTTTCTTGCGCTGAAGCGTTCTAAGCGCGTTGCATTCCGAGAGCCGC 129443

TeratornLC199500 **TGAGCGCGCTCCGAGAGCCACTAAGCGAGCTCCGACCAACTGACGTCGAGCCGCTTGCAGCGCCGATCTCTCGAGCCGTAGATTTGCTAACCAGGGCGGCTTTGGACATTGAGCTTAAGA** 128711
WF-Teratorn TGAGCGCGCTCCGAGAGCCACTAAGTGAGCTCCGACCAACTGACGTCGAGCCGCTTGCAGCGCCGATCTCTCGAGCCGTAGATTTGCTAACCAGGGCGGCTTTGGACATTGAGCTTAAGA 129563

TeratornLC199500 **CGCGAGTGAGCACCTGGCAGCTCGCGGCCGTGTGCTCTTTAATTGGGTTCCTGGATGTCAGCGGAGCTTCGCCGGACGAGCGAGAGCTCGGCGAGGGTGGGCTGATGGCGCTGTTCGATA** 128831
WF-Teratorn CGCGAGTGAGCACCTGGCAGCTCGCGGCCGTGTGCTCTTTAATTGGGTTCCTGGATGTCAGCGGAGCTTCGCCGGACGAGCGAGAGCTCGGCGAGGGTGGGCTGATGGCGCTGTTCGATA 129683

TeratornLC199500 **GCCTGCGGCCGGGATCATTGACGCAATCCATCATCCGCGAGATCGAGGCTTCATTGAACTCACGCTGTTTGGTGTGTGGATCGGGGGACCTCCATCTGCTCGTGGCTAGGAGCCTGCTGA** 128951
WF-Teratorn GCCTGCGGCCGGGATCATTGACGCAATCCATCATCCGCGAGATCGAGGCTTCATTGAACTCACGCTGTTTGGTGTGTGGATCGGGGGACCTCCATCCGCTCGTGGCTAGGAGCCTGCTGA 129803

TeratornLC199500 **GCGTCTCGAAAAAGACTTGTTCGAAGTGCCGGAAGGGTAAAGCTCGAGGTAAAACTCCCCATGCCCGTGGACCCAAAGTCCGCGGAGAGACATCTACTACCCATCATTCGACCAACTCCC** 129071
WF-Teratorn GCATCTCGAAAAAGACTTGTTCGAGGTGCCGGAAGGGTAAAGCGCGAGGTAAAACTCCCCATGCCCGTGGACACAAAGTCCTCGGAGAGACATCTACTACTCATCATTCGACCTACTCCC 129923

TeratornLC199500 **CTAGGGCTGCGTTGCTCACGTGCCACGAGACGATGATGGGCTCCCAGTGA**AAATGCGCTCGTGGACTTTGGTGGAGCTTGGA------GTCCTTTTTTTT-GCACTGTATTTTTGTGTGT 129184
WF-Teratorn CTAGGGCTGCGTTGCTCACGTGCCACGAGACGATGATGGGCTCCCAGTGAAAATGCGCTCGTGGACTTTGGTGGAGCTTGGACACCGAGTCCTTTTTTTTTGCACTGTATTTTTGTGTGT 130043

TeratornLC199500 GTGTGTGGGTGTGTGTGAGTGTGTCTGTTGTGTACCCGAGCAACAGATGTAGAAAAAAAAAA----CGTGTCCCTGCTTCACAGAGACTCTCTTCCAATGCGAAACCTGCCTCTTCACCT 129300
WF-Teratorn GTGTGTGTGTGTGTGTC----TGTCTGTTGTGTACCCGAGCAACAGATGTAGAAAAAAAAAAAACAACGTGTCCCTGCTTCACAGAGACTCTCTTCCAATGCGAAACCTGCCTCTTCACCG 130159

TeratornLC199500 TAACCGTCATCGATGCTCCGTCCGACGAGTAGGCAGTATAAAATGAAAGTGACCCAGAGAATGGAATGGATACTGAGGAGGAACCGCGTCGTGTCTCAGTTTCTGTAGAGAAG**ATGGCAG** 129420
WF-Teratorn TAACCGTCATCGATGCTCCGTCCGACGAGTAGGCAGTATAAAATAAAAGTGACCCAGAGAATGGAATGGATACTGAGGAGGAACCGCGTCGTGTCTCAGTTTCTGTAGAGAAGATGGCAG 130279

TeratornLC199500 **GAAAGGAGGAAAATTGGGTGGACAGACTTCAATCCTTCGCAGATCAGCGAAATTTAACAGTCGTGTACGACACTATAATTTCAAGAGACGACGACTTCGAGATGTACAAATGTAGTGCGC** 129540
WF-Teratorn GAAAGGAGGAAAATTGGGTGGACAGACTTCAATCCTTCGCAGATCAGCGAAATTTAACAGTCGTGTACGACACTATAATTTCAAGAGACGACTTCTTCGAGATGTACAAATGTAGTGCGC 130399

TeratornLC199500 **GGCTTTTCGCTGAGGGCAGCGTCAAGGAAGATTCCCTCGCCCTAGGCATCGGACCTCAAAAGAGGTTGGCAAAATTAGCCGCAGCCAAAGCTTTGCTATTGGGGGTGCTGAAAGAATCCG** 129660
WF-Teratorn GGCTTTTCGCTGAGGGCAGCGTCAAGGAAGATTCCCTCGCCCTAGGCATCGGACCTCAAAAGAGGTTGGCAAAATTAGCCGCAGCCAAAGCTTTGCTATTGGGGGTGCTGAAAGAATCCG 130519

TeratornLC199500 **CTGCAGACAACGCCAAGCTCCCGTCACTGGAGGCTACGAAAGATGAATGGGAATCGAAGCTTGTCGCTAAGTTCGCAAGTTTGCCCACTGCGGAGAACTACGTCGGCCGACTGCAA**GTGC 129780
WF-Teratorn CTGCAGACAACGCCAAGCTCCCGTCACTGGAGGCTACGAAAGATGAATGGGAATCGAAGCTTGTCGCTAAGTTCGCAAGTTTGCCCACTGCGGAGAACTACGTCGGCCGACTGCAAGTGG 130639

TeratornLC199500 GTATCCTATTCACGTGACCCAATGCCGCTGTTTGTGCATGAGAAAAAAAAAAA---CTCTCTTTGCGTTCACAG**GAATTATGCCAGAGGTACAACTGGCCTTTTCCCACCTACAACTTCG** 129897
WF-Teratorn GTATCCTATTCACGTGACCCAATGCCGCTGTTTGTGCATGAGAAAAAAAAAAAAAAGCTCTCTTTGCGTTCACAGGAATTATGCCAGAGGTACAACTGGCCTTTTCCCACCTACAACTTCG 130759

TeratornLC199500 **AGGAACTCACACAAGACCAGTGGGTTCCGTCCGGCCGATTTCGTTGCTTCTGCTCAGCATTTGCGATCTTTTTAGACGGAGCGACCAAGGTGGATGTGCAAGCGACAG**GTAACAGCGATG 130017
WF-Teratorn AGGAACTCACACAAGACCAGTGGGTTGCGTCCGGCAGATTTCGTTGCTTCTGCTCAGCGTTTGCGATCTTTTTAGACGGAGCGACCAAGGTGGATGTGCAAGCGACAGGTAACAGCGATG 130879

TeratornLC199500 TGACACTGTGATTTGTTTTCCCTCAAGTTTTTGGGTTTCAACTAG**GGCCGTTGTCTTACACAGGGACCGGTGAGACCAAGAAGGCAGCTAAGGCCGCGTCGGCAATGGGTGCGCTGGACG** 130137
WF-Teratorn TGACACTGTGATTTGTTT-CCCTCCAGTTTTTGGGTTTCAACTAGGGCCGTTGTCTCACACAGGGACCGGTGAGACCAAGAAGGCAGCTAAGGCCGCGTCGGCAATGGGTGCGCTGGATG 130998

TeratornLC199500 **TCGTGAGACTTCTCGTCCCACTGGTTGAGAAAACCTAG**AGGGGAGGTTCGCCCCAATGTATTCAGAGTCTTCTCCGCATCACAACTCAACTCTGTCTCAGCCGCCGCTCGTGGACCGGCC 130257
WF-Teratorn TCGTGAGACTTCTCGTCCCACTGGTTGAGAAAACCTAGAGGGGAGGTTCGCCCCAATGTATTCAGAGTCTTCTCCGCATCACTACTCAACTCTGTCTCAGCCGCCGTTCGTGGACCGGCC 131118

TeratornLC199500 TTGCACTCGGAGTCTGGTCTTCATGGCTGCCTGCTCTGCTCATCGTAGTGGCATACCTTTTGTGTGTTTTCTGCCCCTATGGCAAGAAGCGCAGACGATACGGCAAGGTGTCGTCTGTTG 130377
WF-Teratorn TTGCACTCGGAGTCTGGTCTTCATGGCTGCCTGCTCTGCTCATCGTAGTGGCATACCTTTTGTGTGTTTTCTGCCCCTATGGCAAGAAGCGCAGACGATACGGCAAGGTGTCGTCTGTTG 131238

TeratornLC199500 CTTTGTAATTGTTGCAGGGATATGGATGAAGTTGCTTGTCCAGAATAAAACATCACTCGAGCATAATTTGGATCTGTGCATTTTTTATTGTGTCGACAGCTTGACATTGTGAACACACTA 130497
WF-Teratorn CCTTGTAATTGTTGCAGGGATATGGATGAAGTTGCTTGTCCAGAATAAAACATCACTCGAGCATAATTTGGATCTGTGCATTTTTTATTGTGTCGACAGCTTGACATTGTGAACACACTA 131358

TeratornLC199500 GTTACAATGAAAAAAAAAAAAAAGAAAAAAGAAAGAAGAAGAACTGTCTGCAGCAGGATCTCAACAACTTGACATTGTGAACACGCTAGTTACAATTAAAGAAAAACAATGAAGAACTGTC 130617
WF-Teratorn GTTACAATGAAAAAAAAAA---GAAAAAAGAAAGAAGAAGAACTGTCTGCAGCAGGATCTCAACAACTTGACATTGTGAACACGCTAGTTACAATTAAAGAAAAACAATGAAGAACTGTC 131475

TeratornLC199500 CACAGCAGGATCTCAGCAAATTGGCCTCGTTGTCTTCTGGGCTGCTCCTCCTGCTATCACTGCAGACCTGCGTAAACTAGGAGAGATAAAAGGGTACAAGTTTCAGCTGTGTGCAGCTTT 130737
WF-Teratorn CACAGCAGGATCTCAGCAAATTGGCCTCGTTGTCTTCTGGGCTGCTCCTCCTGCTATCACTGCAGACCTGCGTAAACTAGGAGAGATAAAAGGGTACAAGTTTCAGCTGTGTGCAGCTTT 131595

TeratornLC199500 CTTTTTTTTTTTATTTAACAAAGCAGACAAACAGAAGAAAGAAGGAAAAAAAAAAAAATTCTCGCACCCCTTTAGCTCTGCC**TCACAGAACCGAGGGGTATTTCTCATCGATCGGGACCTC** 130857
WF-Teratorn CTTTTTTTTT--ATTTAACAAAGCAGACAAACAGAAGAAAGAAGGAAAAAAAAAAAAATTCTCGCACCCCTTTAGCTCTGCCTCACAGAACCGAGGGGTATTTCTCATCGATCGGGACCTC 131713

TeratornLC199500 **CAGAACTTGGTCTGAGCGACTCAGGGCTGCGGGCTCCGGGCTCTCTGCCACTCTTCGCTCTGCACCCCGGCAGTTGCAGGTCTCGACGGTCTCACGCCAATTCGACAAGGTGGACGGTTG** 130977
WF-Teratorn CAGAACTTGGTCTGAGCGACTCAGGGCTGCGGGCTCCGGGCTCTCTGCCACTTTTCGCTCTGCACCCCGGCAGTTGCAGGTCTCGACGGTCTCACGCCAATTCGACAAGGTGGACGGTTG 131833

TeratornLC199500 **ACGTTCGGTCGTGGGCTGGTACACAGCACCGACCCGTTGAGCGTTGGGCTTGTAGATGTTCCCAGCGGGCCTTTTGCAGAACCTGTGCATGATGAGCACCAAGCCCAGCACCATCAAAGG** 131097
WF-Teratorn ACGTTCGGTCGTGGGCTGGTACACAGCACCGACCCGTTGAGCGTTGGGCTTGTAGATGTTCCCAGCGGGCCTTTTGCAGAACCTGTGCATGATGAGCACCAAGCCCAGCACCATCAAAGG 131953

TeratornLC199500 **CGCTGCGACCATGGTGTATGCCGCCCACTCGTTCCGTTCGTACCACCACCAGAAACTGTAATTTTCATTCACCAT**CGTCTCATTGTCCAGCGAGTGGACACGGGGCAATTAGTATACCTC 131217
WF-Teratorn CGCTGCGACCATGGTGTATGCCGCCCACTCGTTCCGTTCGTACCACCACCAGAAACTGTAATTTTCATTCACCATCGTCTCATTGTCCAGCGAGTGGACACGGGGCAATTAGTATACCTC 132073

TeratornLC199500 GGCTCAGTCTGGGTTCGCGTCACGGAAACTACGATCGCAACAAAATACCTCTGACGGCTTGCGAGTGAGAGCCGCGAGCCCAGAGCCTAACCCGGTCGCCGATCGAAACGCGTCGTCGTT 131337
WF-Teratorn GGCTCAGTCTGGGTTCGCGTCACGGAAACTACGATCGCAACAAAATACCTCTGACGGCTTGCGAGTGAGAGCCGCGAGCCCAGAGCCTAACCCGCTCGCCGATCGAAACGCGTCGTCGTT 132193

TeratornLC199500 CCGGCTGCGGCGCTAGGTTGGATCCGACCGCTGCCCCAAGTATCACCCCCTCCGAGTCATGTTTGAAAGCGGCAGTTACCCGGGCTCAACAGAGTTTCGCCTTGTCCTCCAGACCTCATC 131457
WF-Teratorn CCGGCTGCGGCGCTAGGTTGGATCCGACCGCTGCCCCAAGTATCACCCCCTCCGAGTCATGTTTGAAAGCGGCAGTTACCCGGGCTCAACAGAGTTTCGCCTTGTCCTCCAGACCTCATC 132313

TeratornLC199500 AGAACCTCGCGCTCAGCCACCCGGATAGAGGGGAGCAACGATGACGGACGACTCAAATCGCGAGCTACCTCCGGAGAAGTCTCCGGTCATCTCCGCAGCCGCGACCGAAGTTGACCGCGA 131577
WF-Teratorn AGAACCTCGCGCTCAGCCACCCGGATAGAGGGGAGCAACGATGACGGACGACTCAAATCGCGAGCTACCTCCGGAGAAGTCTCCGGTCATCTCCGCAGCCGCGAGCGAAGTTGACCGCGA 132433

TeratornLC199500 GAGCCTCGAGGCCGTCGGCACCCGAAGCGGGACGGGATCCGGAGTGAACGACCTGACCTGCTCGACTGCCAGAATTTCTGCAGATGAGAATACGACTCCCCTTCGCCCTTTGCGGGGAAG 131697
WF-Teratorn GAGCCTCGAGGCCGTCGGCACCCGAAGCGGGACGGGATCCGGAGTGAACGACCTGACCTGCTCGACTGCCAGAATTTCTGCAGATGAGAATACGACTCCCCTTCGCCCTTTGCGGGGAAG 132553

TeratornLC199500 GACGGTGCAACGCTCGCCTGCGCCGACGGAACACGATGGCGAGCGAATGGGCGGCAGTCAGAGGGAACCTTCGCCTCGGAGATTCCCGCCCTCCTCTTTTTGTACCCCTGTGGAGCAGCC 131817
WF-Teratorn GACGGTGCAACGCTCGCATGCGCCGAGGGAACGCGAGGGCGAGCGAATGGGCGGCAGTCAGAGGGAACCTTCGCCTCGGAGATTCCCGCCCTCCTCTTTTCGTACCCCTGTGGAGCAGCC 132673

TeratornLC199500 CATCCCGAGGCACGGAGTTCCTGCGTTGAAAACGAAGATTCCCCACAGCAGAAGGCGGACGAAATCTTCTGACTGCATCAAGAAAAAACTTTTCAGTCAGCGTAACGCACACCCCGTCCC 131937
WF-Teratorn CATCCCGAGGCACGGAGTTCCTGCGTTGAAAACGAAGATTCTCCACAGCAGAAGGCGGACGAAATCTTCTGACTGCATCAAGAAAAAACTTTTCAGTCAGCGTAACGCACACCCCGTCCG 132793

TeratornLC199500 GTTGTGATCACGGCGCTTGTCTTGCTTTATCCCCTAACACAATTGTTCTCTCTAGCGTTGTACGCTCACTGTTGTGAATTAAACAGCATTGAAAACCTCAGCTTGTCCTGTGTCTGTTAT 132057
WF-Teratorn GGTGTGATCACTGCGCTTGTCTTGATTTATCCCCTAACACAATTGTTCTCTCTAGCGTTGTACGCTCACTGTTGTGAATTAAACAGCATTGAAAACCTCAGCTTGTCCTGTGTCTGTCAT 132913

TeratornLC199500 TTGTATCGCATTGGGGTTTT-GTGCCTTTGTCATATATGCGAGGGGCGCTTAGGAACACACGTAGGCCAAAGCGGGGAGGAAAAGTCAGAGCCTCTAGTCCCTTAGACCGAGAGAGAAGT 132176
WF-Teratorn TTGTATCGCATTGGGGTTTTTGTGCCTTTATCATATATGCGAGGGGCGCTTAGGTACACACGTAGGCCAAAGCGGGGAGGAAAAGTCAGAGTCTCTAGTCCCTTAGACCGAGAGAGAAGT 133033

TeratornLC199500 GAAAACAGACAAAGACGCACCCAAGTGAGCTGAGAGACCACATGATTTTATTCAAGGGAGCACGTTCAACAACACACACTCATGTAGCTACATGTTCAGCACACAAGCCCAACAACTTGA 132296
WF-Teratorn GAAAACAGACAAAGACGCACCCAAGTGAGCTGAGAGACCACATGATTTTATTCAAGGGAGCACGTTCAACAACACACGCTCATGTAGCTACATGTTCAGCACACAAGCCCAACAACTTGA 133153

TeratornLC199500 GTGCGGATGTGTGTCACGGTGACAAGTGCATTGTTCGCGAGCTGTCAGGTTGACACGTTACACAGAGGGTGAAAGCGAGGCAAGACCACATTTTGGAGTTGACCGGTACTTTATTGATTT 132416
WF-Teratorn GTG--------TGTCACGGTGACAAGTGCATTTTTCGCGAGCTGTCAGGTTGACACGTTACACAGAGGGTGAAAGCGAGGCAAGACCACATTTTGGAGTTGACCGGTACTTTATTGATTT 133265

TeratornLC199500 TTTTTACATCATCATTTTTCTCTCACTTCATTTCATTGAGAAGAACACATGCCGTGAAAAGTGTGGACCTTTCAAAAATGTATAACAAGCAGTACGTGCACAAGGCCGGGCTCGTAGGGG 132536
WF-Teratorn TTTT-ACATCATCATTTTTCTCTCGCTTCATTTCATTGAGAAGAACACATACCGTGAAAAGTGTGGACCTTTCAAAAATGTATAACAAGCAGTAAGTGCACAAGGCCGGGCTCGTAGGGG 133384

TeratornLC199500 ATGAAGTCGTTAGGTTTGAGGATGAGCGATGCTGCATCAGCAAAATTTTGTTGCCGCTCAAG**TCAAAGCAGTATCAGGATACCCAGGGAGGACAATGTCAACAGAGTGAGGGACGAGACA** 132656
WF-Teratorn ATGAAGTCGTTAGGTTTGAGGATGGGCGATGCTGCATCAGCAAAATTTTGTTGCCGCTCAAGTCAAAGCAGTATCAGGATACCCAGGGAGGACAATGTCAACAGAGTGAGGGACGAGACA 133504

TeratornLC199500 **GAGGGCTTCGTCGAGCCCCCCTTGGCCCAAACTGAGGGACTGGGAGGTTTGACGATCAACGTCACGGAGCAGTTTGAAACAGAGAAACTGTACTGCTCGTTGCGATGGCCCACGTTCGCC** 132776
WF-Teratorn GAGGGCTTTGTCGAGCCCCCCTTGGCCCAAACTGAGGGACTGAGAGGTTTGACGATCAACGTCACGGAGCAGTTTGAAACAGAGAAACTGTACTGCTCGTTGCGATGGCCCACGTTCGCC 133624

TeratornLC199500 **CAGCGGCTACATTCGTACGAGCCCTCGTATCCTGTTGAAATTGGCGAGATGAGTAGAGTTGAGTTCGAGCCCAGGAACTTGTGAGTGGCGCTGATGCTCTCCTCGAGCTCGTACTCCAAG** 132896
WF-Teratorn CAGCGGCTACATTCGTACGAGCCCTCGTATCCTGTTGAAATTGGCGAGATGAGTAGAGTTGAGTTCGAGCCCAGGAACTTGTGAGTGGCGCTGATGCTCTCCTCGAGCTCGTACTCCAAG 133744

TeratornLC199500 **GCAGTTCCGTTGTCCCGGATGTAAAAGAGTCCGCGCTTGGTCCCACGTACAACCTGGTACCAGGAGAAGGCCATGAACCTTTCGGGATTCTCACGCGGACAAGTCAGTGCGACCGCACAC** 133016
WF-Teratorn GCAGTTCCGTTGTCCTGGATGTAAAAGAGTCCGCGCTTGGTCCCACGTACAACCTGGTACCAGGAGAAGGCCATGAACCTTTCGGGATTCTCACGCGGACAAGTCAGTGCGACCGCACAC 133864

TeratornLC199500 **TCTCCCCCATCCGCGCTGACATGGGTGCCGTGAGCCTCGTCCGCTTCTCTCACCTCGTGGCATGAGCGGAACGGGACCAGGGCGGTGCACGCCAACACCGTCAGAAACAGCAT**CTTCGCG 133136
WF-Teratorn TCTCCCCCATCCGCGCTGACACGGGTGCCGTGAGCCTCGACCGCTTCTCTCACCTCGTGGCACGAGCGGAACGGGACCAGGGCGGTGCACGCCAACACCGTCAGAAACAGCATATTCGCG 133984

TeratornLC199500 AAAGGCCCGGAGCTTCTCTATCCTCGTGTCTTTAGCAGAGCAGACCGCGTTGGTCGAGAGGAGAAGTGTGGCGCGGATCTTCTGAACGACGGTCTTATACCGTCCGGGCTTGAGCCTTCT 133256
WF-Teratorn AAAGGCCCGGAGCTTCTCTACCCGCGTGTCTTTAGCAGAGCAGACCGCGTTGGTCGAGAGGAGAAGTGTGGTGCGGATGTTCTGATCGACGGTCTTATACCGTCCGGGCTCGAGCCTTCG 134104

TeratornLC199500 AGCGTCGGCCTCCCGCTGCCGACAGTCTCCTCTATGGGGAATTCCGAGATGGGAAAAATGCCTCTCAACCTCATCGGACCCTGTGCCCAAACCTTGAGCCTCTCTCTCTCTCGAGCGTCC 133376
WF-Teratorn AGCGTCGGCCTCCCGCTGCCGACAGTCTCCTCTATGGGGAATTCCGAGATGGGAAAAATGCCTCTCAACCTCATCGGACCCTGTGCCCAAACCTTGAGCCTCTCTCTCTC--GAGCGCCC 134222

TeratornLC199500 ACCCGACAGCACGAAGGTGATTGCGTTCGGCTTGGTCGAGGGCAACAGGGCTCCGCTCCTCTGTGCGTTTGTCCTCGTCCTATATGTCTGCAC-CAAGCACAACTTGATTTTTTTTTTTA 133495
WF-Teratorn ACCCGACAGCACGAAGGTGATTGCATTCGGCTTGGTCGAGGGCAACAGGGCTCCGCTCCTCTGTGCGTTTGTCCTCGTCCTATATGTCTGCACGCAAGCACAACTTGATTTTTTTTTTTA 134342

TeratornLC199500 CCCTCTGGAATAAACGCAACCGTCTGTGGTATCTGACTTGTGTGAGCGTGTGTGTTTCGGTCCCACATTTACTGTTTGCAAGCTGTTAGGTTGACAAATACACAGAGGTCGAAAGCAAGG 133615
WF-Teratorn CCCTTTGGAATAAACGCAACCGTCTGTGGTATCTGACTTGTGTGAGCGTGTGTGTTTTGGTCCCACGTTTGCTGTTTGCAAGCTGTTAGGTTGACAAATACACAGAGGTCGAAAGCAAGG 134462

TeratornLC199500 CAAACCACATTTCTGAGTTGACAGGTCTTTTATTTCATTTTTTTTTACACCTTCATTTTTTCCTCTCTGACATTTCACAGAAAACGACATAAGCCATGAAACTTTAAACTCTTGGGCTGT 133735
WF-Teratorn CAAACCACATTTCTGAGTTGACAGGTCTTTTATTTCATTTTTTTTTACACCTTCATTTTTTC-TCTCTGACATTTCACAGAAAACGACATAAGCCATGAAACTTTAAACTCTTGGGCTGT 134581

TeratornLC199500 TTGCGGCGAGCGGCAGGTGTCTGAAACGTCAGCCGGATCC**TCACGGGTCTTCGAAGTACGTGGAGAGGTCTTTTTCCGAGTATCTGTCGCCCTCAAGCGGCCTGTGGTCGCAGCAGCTTG** 133855
WF-Teratorn TTGCGGCGAGCGGCAGGTGTCTGAAACGTCAGCCGGATCCTCACGGGTCTTCGAAGTATGTGGAGAGGTCTGTTTCCGAGTATCTGTCGCCCTCAAGCGGCCTGTGGTCGCAGCAGCTTG 134701

TeratornLC199500 **TCATGGACACTCGCGCCCAAGGGCAGGATTCTCCGGTTATGGCGAACACAGAGGCCCTGAAGGCTTCGAGGGCGGTGTAGTCGGCGTCCGTTATGTACTCCTGGAGCGCGTCCAGCACCA** 133975
WF-Teratorn TCATGGACACTCGCGCCCAAGGGCAGGATTCTCCGGTTATGGCGAACACAGAGGCCCTGAAGGCTTCGAGGGCGGTGTAGTCAGCGTCCGTTATGTACTCCTGGAGCGCGTCCAGCACCA 134821

TeratornLC199500 **TCCGAGCCAGGTCTTTGTTTTGCGGGTGAGGAAACCTCTCCTTGCTGAACATCCCCAACCAGGCCTTCACGTGCGGGGGCTTCACTCTTTCGAAGACGCAGGCCAGACCGTCCACCACGG** 134095
WF-Teratorn TCCGAGCCAGGTCTTTGTTTTGCGGATGAGGAAACCTCTCCTTGCTGAACATCCCCAACCAGGCCTTCACGTGCGGGGGCTTCACTCTTTCGAAGACGCAGGCCAGACCGTCCACCACGG 134941

TeratornLC199500 **CCTCCACCACATACACGCCTGGCATCCTCTTGTCGCTGTTGGTGTAGTGCTCAGGGTGATGCTGGTTGTTTTCATAGTGGTGCCTGAAGCCGATGTTGGCCATCGGAGCGGTCATGGTGG** 134215
WF-Teratorn CCTCCACCACATACACGTCTGGCATCCTCTTGTCGCTGTTGGTGTAGTGCTCAGGGTGATGCTGGTTGTTTTCGTAGTGGTGCCTGAAGCCGATGTTGGCCATCGGAGCGGTCATGGTGG 135061

TeratornLC199500 **CGTGGTCCTTGTAGGAGTCGTCTGTCTTGCTCCCCGTCGGGCTGCCGCTCAGTCTGCATCTGCGCTTGAAGCTTTCCTCCAGGAGAGCCCTTGTCATGTTCACGGCCATGATGCCGGAGT** 134335
WF-Teratorn CGTGGTCCTTGTAGGAGTCGTCTGTCTTGCTCCCCGTCGGGCTGCCGCTCAGTCTGCATCTGCGCTTGAAGCTTTCCTCCAGGAGAGCCCTTGTCATGTTCACGGCCATGATGCCGGAGT 135181

TeratornLC199500 **ACGCCACGTCTTCCACCTCGCTGGTCTTGGAGCAGTCGTGGCAGGCGATGTTTCTCACATAGTGTCGGTCGGCCAAGAGGTCCAGGGTGAGACCCCGCAGTTCGGACTCGTCGGCTCCGT** 134455
WF-Teratorn ACGCCACGTCTTCCACCTCGCTGGTCTTGGAGCAGTCGTGGCAGGCGATGTTTCTCACATAGTGTCGGTCGGCCAAGAGGTCCAGGGTGAGACCCCGCAGTTCGGACTCGTCGGCTCCGT 135301

TeratornLC199500 **CGCCCAGCCTCTTTGCCGCCCCGCTCCTGGTCGCCTGCTCGAGAGCCCTGGCGACCACAGCTCCGATCATGGTCACGACCATGTTGTGAGTGAGGAAGGCGACGTGGTGTTCTCTCACCG** 134575
WF-Teratorn CGCCCAGCCTCTTTGTCGCCCCGCTCCTGGTCGCCTGCTCGAGAGCCCTGGCGACCACAGCTCCGATCATGGTCACGACCATGTTGTGAGTGAGGAAGGCGATGTGGTGTTCTCTCACCG 135421

TeratornLC199500 **GAAGGTAGTTCGCCAAGGCATTTTCGAACTCCCTGTTCCAGGATTCTTCCGCACGAAGCACCTTCTGCTCCCTCAAAAGGGCTGCGTTGGCCATCACGTGCACCAGTTTGCATACCCCTT** 134695
WF-Teratorn GAAGGTAGTTCGCCAAGGCATTTTCGAACTCCCTGTTCCAGGATTCTTCCGCACGAAGCACCTTCTGCTCCCTCAAAAGGGCTGCGTTGGCCATCACGTGCACCAGTTTGCATACCCCTT 135541

TeratornLC199500 **CCTGATCTTGAGCGGGCAT**AGTTTCGCAGATGCTCCGCCTGGCAGCTCTCGGGAGAATTCCTTCGAGCAAGCTGTGAGCTGCCCTACTTGAGGGGCTGGTCTTGTGCGAGTACCGGTTGC 134815
WF-Teratorn CCTGATCTTGAGCGGGCATAGTTTCGCAGATGCTCCGCCTGGCAGCTCTCGGGAGAATTCCTTCGAGCAAGCTGTGAGCTGCCCTACTTGAGGGGCTGGTCTTGTGCGAGTACCGGTTGC 135661

TeratornLC199500 GTTTGGGTATCAGGGGCTCGGGTCTTTCCTCGTCATCGTTCTTGCTCGACCTCGAAGAGCGGGGTTTGGGTGTGGAGGGTTTTGAGCTGGCCATGGTTTAGCTTCTTCCCTGGAGTTCAG 134935
WF-Teratorn GTTTGGGTATCAGGGGCTCGGGTCTTTCCTCGTCATCGTTCTTGCTCGACCTCGAAGAGCGGGGTTTGGGTGTGGAGGGTTTTGAGCTGGCCATGGTTTAGCTTCTTCCCTGGAGTTCAG 135781

TeratornLC199500 CGGCTGCAATGAAGGCGGCGGTGATGTCAAATGCGGTGGTGGCAGTAGCAGGGATAAGTGTCAGTGGTTTCAATTCGTGTGGCGAAGACCCAGGGAGGACACTGTCAACAGAGCGAGGGA 135055
WF-Teratorn CGGCTGCAATGAAGGCGGCGGTGATGTCAAATGCGGTGGTGGCAGTAGCAGGGATAAGTGTCAGTGGTTTCAATTCGTGTGGCGAAGACCCAGGGAGGACACTGTCAACAGAGCGAGGGA 135901

TeratornLC199500 CC**TTAAGAGCAGGGATTGCTGAAGATCGAAGTTACGGTGCAGTTTGAAACCAAGAAAGTGTACTGCTCGCTTTGATGGCCCGCGATCGCCCAGCGGAAACAGCCGTAGGAGCTCTCGTAT** 135175
WF-Teratorn CCTTAAGAGCAGGGATTGCTGAAGATCGAAGTTACGGTGCAGTTTGAAACCAAGAAAGTGTACTGCTCGCTTTGATGGCCCGCGATCGCCCAGCGGAAACAGCCGTAGGAGCTCTCGTAT 136021

TeratornLC199500 **CCTCTCGTAAATGGCATGATGAGCAGATTTGAGCCGAGCAGAAAGTGACTGGCGCCGACGCTGGCCTTGTGGTCGTAGTTCGTGGTAATCCCGTTCTCGATGCGAAAGAGCCCGTGCTTG** 135295
WF-Teratorn CCTCTCGTAAACGGCATGATGAGCAGATTTGAGCCGAGCAGAAAGTGACTGGCGCCGACGCTGGCCTTGTGGTCGTAGTTCGTGGTAATCCCGTTCTCGATGCGAAAGAGCCCGTGCTTG 136141

TeratornLC199500 **GTCCCTTGTACATCCACGAACCAGGCGAAGCCCTTGAACTGTCCTGGATTCGGTTGTGGACAAGTCAGTGCCACCTCACACCCTTTCACATACTTCTTGTCACTGATGCCTTTAGCCTTG** 135415
WF-Teratorn GTCCCTTGTACATCCACGAACCAGGCGAAGCCCTTGAACTGTCCTGGATTCGGTTGTGGACAAGTCAGTGCCACCTCACACCCTTTCACATACTTCTTGTCACTGATGCCTTTAGCCTTG 136261

TeratornLC199500 **TCCGCTTCTTTCGCCTCGCTGCATGGGTAGAACGTGACCAGGGCACTGTAGGCTAACACCGTCAGAAACAGCAT**CTTTCCGAAAGACCAGGAGCTTCTGGGCGCGTGTGTCTCTCGTAAA 135535
WF-Teratorn TCCGCTTCTTTCGCCTCGCTGCATGGGTAGAACGTGACCAGGGCACTGTAGGCTAACACCGTCAGAAACAGCATCTTTCCGAAAGACCAGGAGCTTCTGGGCGCGTGTGTCTCTCGTAAA 136381

TeratornLC199500 GCTGACCGCGTTGGTCGAGAGGACGAGTGTGCCGCGGATCTTCTGGACGACGGTCTTATGGCGTCTGGGCTCGAGCCTTCAAGCGCCGGCCTCCCACTGCCGACAGCCTCCTCTTCGGGG 135655
WF-Teratorn GCTGACCGCGTTGGTCGAGAGGACGAGTGTGCTGCGGATCTTCTGGACGACGGTCTTATGGCGTCTGGGCTCGAGCCTTCAAGCGCCGGCCTCCCACTGCCA-----CTCCTCTTCGGGG 136496

TeratornLC199500 TGTTCCGGGATGGGGGGAAGCATCCCA**TCAGCACTTCATCTTCACGATCACCAATACCAGCACGGGGCAGGTCACGGTCGACAAAAAGAGAGCCACCGAGGCTGCAGCGGTTCCGCGGAC** 135775
WF-Teratorn TGTTCCGGGATGGGGGGAAGCATCCCATCAGCACTTCATCTTCACGATCACCAATACCAGCACGGGGCAGGTCACGGTGGACAAAAAGAGAGCCACCGAGGCTGCAGCGGTTCCGCGGAC 136616

TeratornLC199500 **AGCCGCCTTCTCGCACTGCCGCTGCCTTAGTGAACGCTCGTCCGCCTTGGACTCCAGTCTCTTGGCGTAACCCTTGAAGGTGGCGTTCAGCGTCTCGACGGCTCGAGTCACGGACACCCC** 135895
WF-Teratorn AGCCGCCTTCTCGCACTGTCGCTCCCTCAGTGAACGCTCGTCCGCCTTGGACTCCAGTCTCTTGGCGTAACCCTTGAAGGTGGCGTTCAGCGTCTCGACGGCTCGAGTCACGGACACCCC 136736

TeratornLC199500 **TCGGTTGAGGAGATCGAGGGCTGCGTTGGCTCGGGCTCGCATCTCGAGGACAGTCGCTCTTTCGTCCTCACCTCTCGCGAGGTGTGCTCCTGAGAGGGTCTGGGGTTCGCCTGCCACCAG** 136015
WF-Teratorn TCGGTTGAGGAGATCGAGGACTGCGTTGGCTCGGGCTCGCATCTCGAGGACAGTCGCTCTTTCGTCCTCACCTCTCGCGAGGTGTGCTCCTGAGAGGGTCTGGGGTTCGCGTGCCACCAG 136856

TeratornLC199500 **TATCTCTCTCGACCTACCCCTCCAGCTCTCTGCCTCGACCCCCTTCGTCAGGGTGTACGAGTTGAAATGGCTGTCGTAGCAGAGTGGCCCGAAACACGCAGACCCGCACACCAATGGACA** 136135
WF-Teratorn TATCTCTCCCGACCTGCCCCTCCAGCTCTCTGCCTCGACCCCCTTCGTCAGGGTGTACGAGTTGAAATGGCTGTCGTAGCAGAGCGGCCCGAAGCACGCAGACCCGCACACCAATGGACA 136976

TeratornLC199500 **ATCTTGGCTTATGTCGCTGCCACGCTTCACTTGGCACGCGGATAGCGTGTCTCCGTCGACCCGTAGCGAGTCGCACAGGCCTCTCCGGTCGCATCTGGGTTCCCTGTAGCTGGAGCGCGA** 136255
WF-Teratorn ATCTTG-------------CCACGCTTCACTTGGCACGCGGATAGCGTGTCTCCGTCGACCCGTAGCGAGTCGCACAGGCCTCTCCGGTCGCATCTGGGTTCCCTGTAGCTGGAGCGCGA 137083

TeratornLC199500 **GTCGTTCACCATGCCCCTGTCCAGCCAGCACGTCCGATTTCCCGAGGACGAGTCCACAAAGGCCATGTACCAGCGGGACAGCAGCACGGGTTCCCACTTTTTTCTCTCGACGTCCAGCGC** 136375
WF-Teratorn GTCGTTCACCATGCCCCTGTCCAGCCAGCACGTCCGATTTCCCGAGGACGAGTCCACAAAGGCCATGTACCAGCGGGACAGCAGCACGGGTTCCCACTTTTTTCTCTCGACGTCCAGCGC 137203

TeratornLC199500 **CAGCCCACCGTGGGTCGAACGAGAGAGGATCGGACGAGCGGCTGTGGGAACGTCCGTGTGAAACTCTGAGGGCGATCCGGGCGCGCAGCCCGTCATGTCGGCCCTCATCAGGCCCTGCGC** 136495
WF-Teratorn CAGCCCACCGTGGGTCGAACGAGAGAGGATTGGACGAGCGGCTGTGGGAACGTCCGTGTGAAACTCTGAGGGCGATCCGGGCGCGCAGCCCGTCATGTCGGCCCTCATCAGGCCCTGCGC 137323

TeratornLC199500 **GCAAACCGCCAGTCTCTCTCGAGCCCCGACCAACGCAAAGGTGACGGCGTTCAGCTGCGCCGAGAGCAGCAGGGCTCCGCTCCTCCACGCGTTCATCCTCGCAGCGAAGGCGCTACCGCT** 136615
WF-Teratorn GCAAACCGCCAGTCTCTCTCGAGCCCCGACCAACGCAAAGGTGACGGCGTTCAGCTGCGCCGAGAGCAGCAGGGCTCCGCTCCTCCACGCGTTCATCCTCGCAGCGAAGGCGCTACCGCT 137443

TeratornLC199500 **CAGCTGAGAATCGTCGTCGTTCTCGTAGTCTCCCTCGCGCTTTTTCTGAAATTTTTCCTCTGTGGCCTTGGCGGCCGCCCGGGCCTCTTCTATCAAAGACCTGTGGTAGCCCATCACGTC** 136735
WF-Teratorn CGGCTGAGAATCGTCGTCGTTCTCGTAGTCTCCCTCGCGCTTTTTCTGAAATTTTTCCTCTGTGGCCTTGGCTGCCGCCCTGGCCTCTTCTATCAAAGACCTGTGGTAGCCCATCACGTC 137563

TeratornLC199500 **CACCGTCGCCGACGGCTGACTTCCCGAGTCGAGGTCCGAGCCCAGGTGCGCGTCGGCCAGAGCCTCGAGGTCCTTGAAGGCCGCCTGCAGCTCCCTCGCTGTCCGATACTGGCTGCGGAC** 136855
WF-Teratorn CACCGTCGCCGACGGCTGACTTCCCGAGTCGAGGTCCGAGCCCAGGTGCGCGTCGGCCAGAGCCTCGAGGTCCTTGAAGGCAGCCTGCAGCTCACTCGCTGTCCGATAGTGGCTGCGGAC 137683

TeratornLC199500 **GGATGTGGACAAGGCGACGTCCGCGTCCCGCCAGCTACCAGAGCCGCAGAGGACGTCGCAGCCCAGCCGCTCGCCGTCGTCGGGCAGGCCTTGGCAAGAGACTTTGGCGCACTCCGGATA** 136975
WF-Teratorn GGATGTGGACAAGGCGACGTCCGCGTCCCGCCAGCTACCAGAGCCGCAGAGGACGTCGCAGCCCAGCCGCTCGCCGTCGTCGGGCAGGCCTCGGCAGGAGACTTTGGCGCACTCCGGATA 137803

TeratornLC199500 **GTCCGAGGGAACGAAGAAAGTCAGATTTGAATTGAGGTAGAACTGGTACGAGAGGAACGTCTGACTGTCTCCCGGTCGCAGTATCTTCCACCTCCCGGGTCTGAGTATGACCCTTGGATC** 137095
WF-Teratorn GTCCGAGGGAACGAAGAAAGTCAGATTTGAATTGAGGTAGAACTGGTACGATAGGAACGTCTGACTGTCTCCCGGTCGCAGTATCTTCCACCTCCCGGGTCTGAGTATGACCCTGGGATC 137923

TeratornLC199500 **TCGAGGCGCAGAGGGCCCGCTTTCCTCCGAGCGAGTCGCCTCGGGACCCGGAGTCACGCGAAACAGAGTCAGGGCGACCGTAGAGAGAAGGCACGCAGCCCTTGTCGGTCGTTTCATTTC** 137215
WF-Teratorn TCGAGGCGCAGAGGGTCCGCTTTCCTCCGAGCGAGTCGCCTTGGGACCCAGAGTCACGCGAAACAGAGTCAGGGCGACCGTAGAAAGAAGGCACGCAGCCCTTGTCGGTCGTTTCATTTC 138043

TeratornLC199500 **AGTTTCCTGTACCTGGGAGTCCGAGACCGAGAGCAGACTGCGCAAAGGGCCCCCATCGCGGCGGCATTGATTAATACGGAGACAGCCACCGCAGTCAAGCACAACGCGTAGGAGCGCTCC** 137335
WF-Teratorn AGTTTCCTGTACCTGGGAGTCCGAGACCGAGAGCAGACTGCGCAAAGGGCCCCCATCGCGGCGGCATTGATTAATACGGAGACAGCCACCGCAGTCAAGCACAACGCGGAGGAGCGCTCC 138163

TeratornLC199500 **ATTGCCAGGGCTCGACGCTCGGACACGACGCTTTCCAGTGCGTCGTTCTTTCGAGCCCGCCTCTCGAGCAGCGAAGCTTCGGAGCCCAGGCTCGCCAGACTTTGGTTGGCTTGTCGAAGG** 137455
WF-Teratorn ATTGCCAGGGCTCGGCGCTCGGACACAACGCTTTCCAGTGCGTCGTTCTTTCGAGCCCGCCTCTCGACCAGCGAAGCTTCGGAGCCCAGGCTCGCCAGACTTTGGTTGGCTTGTCGAAGG 138283

```
TeratornLC199500 CAGTCCAGGAGCTGGGAGAGAGGGGAGCTCGCCTCGCCGCAGTGGCGCAGCGCGAGCGACTCGTAGGCCAGCAGGGCGAGAGCTCGGGCTTCCTGGGCGTAGTCCCTCGACGCGGAGCGG 137575
WF-Teratorn      CAGTCCAGGAGCTGGGAGAGAGGGGAGCTCGCCTCGCCACAGTGGCGCAGCGCGAGCGACTCGTAGGCCAGCAGGGCGAGAGCTCGGGCTTCCTGGGCGTAGTCCCTCGACGCGGAGCGG 138403

TeratornLC199500 ACCCGCTCTCCGGGCCTCGTGAAAAACCCTGGGAAGCTCTGCGGGCCGTCCCCGGGCCCCTCTGAGCCGAAGGAGAAGGGCCGCAAATTGCAAGGAACTGCGGCACGCACGGCTCCGTGG 137695
WF-Teratorn      ACCCGCTCTCCGGGCCTCGTGAAAAACCCGGGGAAGCTCTGCGGGCCGTCCCCGGGCCCCTCTGAGCCGAAGGAGAAGGGCCGCAAATTGCAAGGAACTGCGGCACGCACGGCTCCGTGG 138523

TeratornLC199500 TCCGCTCGTAACAGGATCCTGAGCAGGATGTCGTCTCCGTAGTCGACCGCGATCCGCGCCGCCACGGGGTACCCCGTCGCTCCCTCTGGCCAGTCTCCTTCCGCAGAGTCGGCCAGTTCC 137815
WF-Teratorn      TCCGCTCGTAACAGGATCCTGAGCAGGATGTCGTCTCCGTAGTCGACCGCGATCCGCGCCGCCACGGGGTACCCCGTCGCTCCCTCTGGCCAGTCTCCTTCCGCAGAGTCGGCCAGTTCC 138643

TeratornLC199500 AGGCACGAACCGCTTTGCTCGCCAAAGCACTCGAGTAGCAGACTCAGCTGGTCGCGTGCCCTCCAGGGCCAGGAGACGGCTGGTCGTGTCCTTTCCGAAGCCCCATCGCCCGACAGCCAC 137935
WF-Teratorn      AGGCACGAACCGCTTTGCTCGCCAAAGCACTCGAGTAGCAGACTCAGCTGGTCGCGTGCCCTCCAGGGCCAGGAGACGGCTGGTCGTGTCCTTTCCGAAGCCCCATCGCGCGACAGCCAC 138763

TeratornLC199500 TCTTCCTCTCGCCTCGTGAGAACCGGTGGAGGTAACCACCGCTTAGAGTCCATCAGTTTTCGGAGACGGGCCTTGGTCACTGGCTTTGTTGCCGAGGGCACGGACAGTTCCAATTTCACT 138055
WF-Teratorn      TCTTCCTCTCGCCTCGTGAGAACCGGTGGAGGTAACCACCGCGTAGAGTCCATCAGTTTTCGGAGACGGGCCTTGGTCACTGGCTTTGTTGCCGAGGGCACGGACAGTTCCAATTTCACT 138883

TeratornLC199500 CTCGTTCGAGCCTCGCGTACTCGTCGGGAGCAGCATTCAAGCTCGCCGTGGTCGCCCTGCGGTCTTCGATGGCAAAGCGGGCTCTCCATGCTCCGCGCTCTCGAGACCTGCGTCCGGAGT 138175
WF-Teratorn      CTCGTTCGAGCCTCGCGTACTCGTCGGGAGCAGCATTCAAGCTCGCCGTGGTCGCCCTGCGGTCTTCGATGGCAAAGCGGGCTCTCCATGCTCCGCGCTCTCGAGACCTGCGTCCGGAGT 139003

TeratornLC199500 CTGGCGTGCCCCTGCCGTTTGGCCTCGTGAACGTAGATCCTGTCCGCCTCGACGAACGCCACGAGTTTCCCTCCCGAGGTCAGTTTCACAACCTGCGCCCCGACAGGCACGGACTCTGCG 138295
WF-Teratorn      CTGGCGTGCCCCTGCCGTTTGGCCTCGTGAACGTAGATCCTGTCCGCCTCGACGAACGCCACGAGTTTCCCTCCCGAGGTCAGTTTCACAACCTGCGCCCCGACAGGCACGGACTCTGCG 139123

TeratornLC199500 ATCTCTGCCTGCTCTCGCAGCAGCTGGGCGCCGCCACCCTTCAGAGGATCACCGATGTTCATGCGAGAGGATAGGCCCAGTCTCAAGCCGGGAACACCCGGGATCGTCACGAAGCCTTTG 138415
WF-Teratorn      ATCTCTGCCTGCTCTCGCAGCAGCTGGGCGCCGCCACCCTTCAGAGGATCACCGATGTTCATGCGAGAGGATAGGCCCAGTCTCAAGCCGGGAACACCCGGGATCGTCACGAAGCCTTTG 139243

TeratornLC199500 GCCAGCGGAGCGTTCTCTGCAGCTCGCTGAAAAAATATCCCGCTCTGTAGCTTGTATAGGCCGTCGACCTCGACGTGGGACGAGCCCCCGTCGTCGAACGAGGAGCTCCTCGTGTACCTG 138535
WF-Teratorn      GCCAGCGGAGCGTTCTCTGCAGCTCGCTGAAAAAATATCCCGCTCTGTAGCTTGTATAGGCCGTCGACCTCGACGTGGGACGAGCCCCCGTCGTCGAACGAGGAGCTCCTCGTGTACCTG 139363

TeratornLC199500 TAGTGCAGGGTGCCGGAGCGTCGGGTGGCGGAGTCGATCGCTGTGCTGAGCTCTGCAAACAGCGAACTTCCCTCCGGACGCCCGAACGTGTGGTCCAGGTATCCCAGGGAGTACCGGCTC 138655
WF-Teratorn      TAGTGCAGGGTGCCGGAGCGTCGGGTGGCGGAGTCGATCGCTGTGCTGAGCTCTGCAAACAGCGAACTTCCCTCCGGACGCCCGAACGTGTGGTCCAGGTATCCCAGGGAGTACCGGCTC 139483

TeratornLC199500 ACCCCCCACCTCGCCATGTCCGAGACGATGTCCCGCAGCTCACCCGCCGAATACATACGATCGGTTGAGCTCTTTTTTCCCATGTGTCTGTTGCGCATCCTCATTGCTGAGCATTTTTCG 138775
WF-Teratorn      ACCCCCCACCTCGCCATGTCCGAGACGATGTCCCGCAGCTCACCCGCCGAATACATACGATCGGTTGAGCTCTTTTTTCCCATGTGTCTGTTGCGCATCCTCATTGCTGAGCATTTTTCG 139603

TeratornLC199500 TCCGGCTCGTAGCGGGCCGTCGGCAGCGTCACAGCGATCCCCGCTCGGTTGTTTGACCTGCGGCTGAGCTCCAGGAAAACGGACGGCCCCGTCAGGCCAAGCGGCGCTTCGCCAGCGATC 138895
WF-Teratorn      TCCGGCTCGTAGCGGGCCGTCGGCAGCGTCACAACGATCCCCGCTCGGTTGTTTGACCTGCGGCTGAGCTCCAGGAAAACGGACGGCCCCGTCAGGCCAAGCGGCGCTTCGCCAGCGATC 139723

TeratornLC199500 GACTCGACCGAGTTGTGCTTCTCCGGGGGCCAGCGCCCGGGTTCAGTCCACTTGCAGGGGCCCTCCAGTTTTTCCATCCAGGAATCCAGGTCCAGCAAGGCGCCATGCTGAGCCGCCACC 139015
WF-Teratorn      GACTCGACCGAGTTGTGCTTCTCCGGGGGCCAGCGCCCGGGTTCAGTCCACTTGCAGGGGCCCTCCAGTTTTTCCATCCAGGAATCCAGGTCCAGCAAGGCGCCATGCTGAGCCGCCACC 139843

TeratornLC199500 TCCCCGGCTTCCCGCAAGGCCGCTTCGGCCTCCTCGGCCCGACTCGTTTCGAGCCACAGGGCCGGCTTCCCGGGGGTCCGGGATCCTTTCTCTGCGACCTCCGCGGGCCTCAGACCCGTG 139135
WF-Teratorn      TCCCCGGCTTCCCGCAAGGCCGCTTCGGCCTCCTCGGCCCGACTCGTTTCGAGCCACAGGGCCGGCTTCCCGGGGGTCCGGGATCCTTTCTCTGCGACCTCCGCGGGCCTCAGACCCGTG 139963

TeratornLC199500 GGGCACAACGCGTACACGGTGCCTCGCTCTTGACACATGGTGCAGTTCCGCCCGTGGGGCGCCGTGCGCCCACACTCGAGCAGAGGCCGTCTCCCGTGCCTTCGGTGCAGCATCCTCCCG 139255
WF-Teratorn      GGGCACAACGCGTACACGGTGCCTCGCTCTTGACACATGGTGCAGTTCCGCCCGTGGGGCGCCGTGCGCCCACACTCGAGCAGAGGCCGTCTCCCGTGCCTTCGGTGCAGCATCCTCCCG 140083

TeratornLC199500 TCCTCCCCGTACAGCGTCCCGATCTCAGAGCACTCGGAGTAAGTTTTGGTGCAAGCGTAGCGCGTCGCGTTATCGTTCTTCGCCGACATGAGGCGCGCGATGGTCTTCCCGGACATTGAG 139375
WF-Teratorn      TCCTCCCCGTACAGCGTCCCGATCTCAGAGCACTCGGAGTAAGTTTTGGTGCAAGCGTAGCGCGTCGCGTTATCGTTCTTCGCCGACATGAGGCGCGCGATGGTCTTCCCGGACATTGAG 140203

TeratornLC199500 GGCATGACGTCCACGCGTGACCCGTACATGTCGCGCACGACTCCGAAGTAGGCCCCGAACCCGGGTTCACGCGGGTCCGCGTCCAACACGGCGAACTTCAGGTCTGGGCAGCCGCCCAAT 139495
WF-Teratorn      GGCATGACGTCCACGCGTGACCCGTACATGTCGCGCACGACTCCGAAGTAGGCCCCGAACCCGGGTTCACGCGGGTCCGCGTCCAACACGGCGAACTTCAGGTCTGGGCAGCCGCCCAAT 140323

TeratornLC199500 AACCAATCCAGCACGGCGGCTGTCTCCTCGACTCTCAGGTCTCTGCTCCTGGGCTCGGGTCGAACGAGCCTCACTCCTATCGCGCCTCCCCATCCTTGACGACTCATCTCGCGCAGGGTC 139615
WF-Teratorn      AACCAATCCAGCACGGCGGCTGTCTCCTCGACTCTCAGGTCTCTGCTCCTGGGCTCGGGCCGAACGAGCCTCACTCCTATCGCGCCTCCCCATCCTTGACGACTCATCTCGCGCAGGGTC 140443

TeratornLC199500 TCGTTGGCCCCGGCAGGCACTCTCGGGCCGGTCTCGACCTGCTTCTGTCTCCTTGCGCTCGTTTGTCGTGCGGGGAGACGGAGGTAGGGACCGCCTTGTCGTCCCGCGTCGTTGTGCAGC 139735
WF-Teratorn      TCGTTGGCCCCGGCAGGCACTCTCGGGCCGGTCTCGACCTGCTTCTGTCTCCTTGCGCTCGTTTGTCGTGCGGGGAGACGGAGGTAGGGACCGCCTTGTCGTCCCGCGTCGTTGTGCAGC 140563
```

TeratornLC199500 **AGCACGGGTACCTCGAGCAGCATCACGTCACAGCTCTTGACGTCCAGAGGCGCGTTCGGCGAGTCGCGCCTGGGGGCTGGGGCGCCGTCAAACACGGCGCTTGTGGCCTGGCAGAGGCGC** 139855
WF-Teratorn AGCACGGGTACCTCGAGCAGCATCACGTCACAGCTCTTGACGTCCAGAGGCGCGTTCGGCGAGTCGCGCCTGGGGGCTGGGGCGCCGTCAAACACGGCGCTTGTGGCCTGGCAGAGGCGC 140683

TeratornLC199500 **TCGGTCTTGCGCGTTGTCGTCCCCCACAGCGTCCTACCCTGTGCTGCGCCGATGGCGAACTGCCGGCCGGACACCTCGAGCGTCCAGTCGTTGCCTCCGAGTCCGTGGTAAGCGTGCCTG** 139975
WF-Teratorn TCGGTCTTGCGCGTTGTCGTCCCCCACAGCGTCCTACCCTGTGCTGCGCCGATGGCGAACTGCCGGCCGGACACCTCGAGCGTCCAGTCGTTGCCTCCGAGTCCGTGGTAAGCGTGCCTG 140803

TeratornLC199500 **AGGCCGATCAGGGCGCACTCGGGCGAGCAGGGGCCCGCGGGGGAGCTGTACCGCAGGAGGTCCTTCGTGCAGTTGCAGCCGGGGCTCTGCTGCTGCTGCGGGTTCGCGACGGTACGCCAC** 140095
WF-Teratorn AGGCCGATCAGGGCGCACTCGGGCGAGCAGGGGCCCGCGGGGGAGCTGTACCGCAGGAGGTCCTTCGTGCAGTTGCAGCCGGGGCTCTGCTGCTGCTGCGGGTTCGCGACGGTACGCCAC 140923

TeratornLC199500 **TCCAAGAGGGCCACCGCTAAGAGCAGCCGCACTTGCCAACTGGGAAACATGGCCCCCTGTTGTTTTTTTTTTTTTTTTTTTTTTAGTTTTCAA**TGGTTCGCTTTACATGGCCCCCTGTTGT 140215
WF-Teratorn TCCAAGAGGGCCACCGCTAAGAGCAGCCGCACTTGCCAACTGGGAAACATGGCCCCCTGTTGTTTTTATTTTTTTTTTTTT---AGTTTTCAATGGTTCGCTTTACATGGCCCCCTGTTGT 141040

TeratornLC199500 TTTTATTTTATTTTTTTTTAGTTTTCAATTGTTCGCTTTCCGTCGCCGAGGTCGGAGTTCCTCTCGACTAACGGGGGATAAGTTAATTCACCTAACGAGCTCGCAGGCTCGCGGAGCC**AT** 140335
WF-Teratorn TTTTATTTTATTTTTTTT-AGTTTTCAATGGTTCGCTTTCCGTCGCCGAGGTCGGAGTTCCTCTCGACTAACGGGGGATAAGTTAATTCACCTAACGAGCTCGCAGGCTCGCGGAGCCAT 141159

TeratornLC199500 **GGCGCTGGACCAGGGATTTCCGTCGTCGCGACGCTGTGTGTCGAAACCCAAGACGCTCCTGAGTTTGTTCGACGGGACCGGCAGCATATGCAGGCCTTTCGTTGCCGCCGGGTGGACGGT** 140455
WF-Teratorn GGCGCTGGACCAGGGATTTCCGTCGTCGCGACGCTGTGTGTCGAAACCCAAGACGCTCCTGAGTTTGTTCGACGGGACCGGCAGCATATGCAGGCCTTTCGTTGCCGCCGGGTGGACGGT 141279

TeratornLC199500 **GCGTAGGCTCGACATGGACGGGAGACACGGGGCTGATATAGTGACGGACGTAAGGCAGTGGGACCCGAGCAGGGACTGGCGGGGGCCCGCGCCCGACGTCGTTTTCGCTGGGCCTCCGTG** 140575
WF-Teratorn GCGTAGGCTCGACATGGACGGGAGACACGGGGCCGATATAGTGACGGACGTAAGGCAGTGGGACCCGAGCAGGGACTGGCGGGGGCCCGCGCCCGACGTTGTTTTCGCTGGGCCTCCGTG 141399

TeratornLC199500 **CGAAAACTACAGCTTGGCCCGGACCAGAGCCTGCACGCCCAGGGACCTGAGACTTGCTGACTCGCTGGTGACCAAGACGCTGGAAATCATAGAACACTTTCATGGACAAAACCCTCGCAT** 140695
WF-Teratorn CGAAAACTACAGCTTGGCCCGGACCAGAGCCTGCACGCCCAGGGACCTGAGACTCGCTGACTCGCTGGTGACCAAGACGCTGGAAATCATAGAACACTTTCATGGACAAAACCCTCGCAT 141519

TeratornLC199500 **GCTGTACTTTGTCGAGAACCCGGATTCCAGCATGCTATGGAGGCGGTGGGTTTCTCACCGGCTGTACGAGGACCCCGGCAAGCACTTTCTCAGCTCCTTTGTCGAGTACATGGACGCGAA** 140815
WF-Teratorn GCTGTACTTTGTCGAGAACCCGGATTCCAGCATGCTATGGAAGCGCTGGGTTTCTCACCGGCTGTACGAGGACCCCGGCAAGCACTTTCTCAGCTCCTTTGTCGAGTACATGGACGCGAA 141639

TeratornLC199500 **AAGGGGCGCCAAGAGCTACAAGTACCTGACTTGGGTCAAGCGCAAAAGCTTCACCTCGGGCTGCAGAGACAGTCGCATCGTCAGGCTGGACTACTGCATGTACGGCACCAAGTACCGAAA** 140935
WF-Teratorn AAGGGGCGCCAAGAGCTACAAGTACCTGACTTGGGCCAAGCGCAAAAGCTTCACCTCGGGCTGCAGAGACAGTCGCGTCGTCAGGCTGGACTACTGCATGTACGGCACCAATTACCGAAA 141759

TeratornLC199500 **GAGGACGAGGCTGATGACCAACAGCCCCTTCGAGGCCTCGCTGTGCAAAAAGAACTGCTCGGGTTTCAGAGAGCGCAAGCACGCGGAAGCGGCTCAGCGGGGACCCTACGACAGGGGCGC** 141055
WF-Teratorn GAGGACGAGGCTGATGACCAACAGCCCCTTCGAGGCCTCGCTGTGCAAAAAGAACTGCTCGGGTTTCAGAGAGCGCAAGCACGTGGAAGCGGCTCAGCGGGGACCCTACGACAGGGGCGC 141879

TeratornLC199500 **TAAGGGCGAGGGGCACTCCCTGGACGAGCTTCACGCCTATCCCGCCCGACTGGCGGACGCGATATTCACACACGTCAGCCGGGCCGAGTGGCGCGTACGACAGGGACTGAGCCCGTCGAC** 141175
WF-Teratorn TAAGCGCGAGGGGCACACCCTGGACGAGCTTCACGCCTATCCCGCCCGACTGGCGGACGCGATATTCACACACGTCAGCCGGGCCGAGTGGCGCGTACGACAGGGACTGAGCCCGTCGAT 141999

TeratornLC199500 **TGCCGAGCCCGTCTTGAGACCGGTGAAGCGCCAGAAGAATGTGTGA**TGTGAATGAGTGGATGAGATATGCTGACATGGTGTATCTTGGGGGCCCATACTGACCGTGCGATCGTGGGTCAT 141295
WF-Teratorn TGCCGAGCCCGTCTTGAGACCGGTGAAGCGCCAGAAGATTGTGTGATGTGAATGAGTGGATGAGATATGCTGACATGGTGTATCTTGGGGGCCCATACTGACCGTGCGATCGTGGGTCAT 142119

TeratornLC199500 TTCAAAAGCTCCAACTAAAAACCAACAGGCTCTGTGTCTCCTGTCTTGAGATGTCTGGGCGATTGTACACATACACTCTTTTGCTTGCTGCTTCAGTTTCTGCTGAGCGTTCCAAATAAA 141415
WF-Teratorn TTCAAAAGCTCCAACTAAAAACCAACAGGCTCTGCGTCTCCTGTCTTGATATGTCTGGGCGATTGTACACATACACTCTTTTGCTTGCTGCTTCAGTTTCTGCTGAGCGTTCCAAATAAA 142239

TeratornLC199500 GCTCTGACTAGAAAACAGCACCCTCTGTGATTCTTGTCTAACCTCGTCTGTTTTTATTTTTTTTT----ATTTTTTGTTCACAGGGTCAATGCATTTGCATCCAAAGTGCAAACCTAGGC 141531
WF-Teratorn GCTCTGACTAGAAAACAGCACCCTCTGTGATTCTTGTCTAACTTCGTCTGTTTTTATTTTTTTTTTTTAAATTTTTTGCTCACAGGGTCAATGCATTTGCATCCAAAGTGCAAACCTAGGC 142359

TeratornLC199500 GAAATAAAATGTTGACACTCGAAAAACAAGCAAAAAAAAAAAAAAGCAACACATATAAGACTCAAATGGGGTTTGGTGCAAACTTTATTATTTCACACATATGTATTGATCCATTTGTAA 141651
WF-Teratorn GAAATAAAATGTTGACACTCGCAAAACAAGCAAAAAAGAAAAA--GCAACACATATAAGACTCAAATGGGGTTTGGTGAAAACTTTATTATTTCACACATATGTATTGATCCATTTGTAA 142477

TeratornLC199500 GACACTGATTTTCAAAGCAGAGAAACAGAACAGAAAAG-----AAAAAAAAAAAACGGAAAAAAAAGCAGCGGATGTCAAACAACCCCAGTTTCCCCCAGTAAGTCTCTTAAGAGCATCA 141766
WF-Teratorn GACACTGATTTTCAAAGCAGAGAAACAGAACAGAAAAGGAAAAAAAAAAAAAAAACGGAAAAAAAA-CAGCGGATGTCAAACAACCCCAGTTTCCCCCAATAAGTCCCTTAAGAGCATCA 142596

TeratornLC199500 GTTACAAGCAGGTCTGACCCGTCTTCCTTCC**TCAAGCCACAAAACATTTGGCTTTGGTGTGTGAGTCAAAGACCAGGGCACAAAGCTTGGTCCTGGAATGTACCCTTCCGTGAATGGGGC** 141886
WF-Teratorn GTTACAAGCAGGTCTGACCCGTCTTCCTTCCTCAAGCCACAAAACATTTGGCTTTGGTGTGTGAGTCAAAGACCAGGGCGCAAAGCTTGGTCCTGGAATGTACCCTTCCGTGAATGGGGC 142716

TeratornLC199500 **ACGGGTAGGTCTCAGACTTTTCGTCGGCTTGCACGTACACCAGCCCCGATCCGGTCGTCCTCAGCGTCACCGGCACGCCCCACTGATCTGACAGTCTCCGAGCCTCGCTTTCTGCCACCG** 142006
WF-Teratorn ACGGGTAGGTCTCAGACTTTTCGTCGGCTTGCACGTACACCAGCCCCGATCCGGTCGTCCTCAGCGTCACTGGCACGCCCCACTGATCGGACAGTCTCCGAGCCTCGCTTTCTGCCACCG 142836

| | | |
|---|---|---|
| TeratornLC199500 | **CTCGATCCGGGGCCCCTGGTCGCTCCATTTCGGGACCAGGCCAGATTCTCTCCGCCCCGACTGCGCACTCGAAGCGTGCGCCGAGCGACGTCACGTCGGGAGCGATGGGGTTAGAGGAGA** | 142126 |
| WF-Teratorn | CTCGATCCGGGGCCCCTGGTCGCTCCATTTCGGGACCAGGCCAGATTCTCTCCGCCCCGACTGCGCACTCAAAGCGTGCGCCGAGCGACGTCACGTCGGGAGCGATGGGGTTAGAGGAGA | 142956 |
| TeratornLC199500 | **GGCCCACGGAGAGAGCGCAGGGGTCCTCGAAGCGAGGATCCTCTAAGTCTTCTCGAGCGTTGAAGGTGTCCACGTACTCGAAAGCGAAGCCGTCCTTGCTGCAGTTTGGAAGTCTCACGC** | 142246 |
| WF-Teratorn | GGCCCACGGAGAGAGCGCAGGGGTCCTCGAAGCGAGGATCCTCTAAGTCTTCTCGAGCGTTGAAGGTGTCCACGTACTCGAAAGCGAAGCCGTCCTTGCTGCAGTTTGGAAGTCTCACGC | 143076 |
| TeratornLC199500 | **ACTTTCTGCAAGAATACGGAGCTTTGTCGATCGCCGAGCGCCTGCGCGAGCCGGCGTTCGGGTCCTCGGTCCTCAGCCATCCCGACCCGGACACCCTCCGCTCACCGACCGCCCTTCTGA** | 142366 |
| WF-Teratorn | ACTTTCTGCAAGAATACGGAGCTTTGTCGATCGCCGAGCGCCTGCGCGAGCCGGCGTTCGGGTCCTCGGTCCTCAGCCATCCCGACCCGGACACCCTCCGCTCACCGACCGCTCTTCTGA | 143196 |
| TeratornLC199500 | **GATCGCACAGCCCGCACTGAGACTGGTCGTCCGGGTATGAGAGCACAGAGTAGCGCAGGGACCTTCTCTCCATCTCTCTCAGCAGCCGCCCGTAAACTTCGATCCCATCGAAGCTGCACA** | 142486 |
| WF-Teratorn | GATCGCACAGCCCGCACTGAGACTGGTCGTCCGGGTATGAGAGCACAGAGTAGCGCAGGGACCTTCTCTCCATCTCTCTCAGCAGCCGCCCGTAAACTTCGATCCCATCGAAGCTGCACA | 143316 |
| TeratornLC199500 | **GTTCGACGGGCAGGTACCACAGGATCCTCATGCTGAGCTTAGCCTTGGCCGTGGACACCCTCTGGTACGCTGCGAACTTGCCCAGCGATCGGCACAGTTCCCCCGAAAGCTCGGGGTTTC** | 142606 |
| WF-Teratorn | GTTCGACGGGCAGGTACCACAGGATCCTCATGCTGAGCTTAGCCTTGGCCGTGGACACCCTCTGGTACGCTGCGAACTTGCCCAGCGATCGGCACAGTTCCCCCGAAAGCTCGGGGTTTC | 143436 |
| TeratornLC199500 | **CCTCGACCTCCGCGCGCTTCACTCCGGCCAGCCCGAGCACAGATTCAGCGGCGACGAGGACGAGGCTCGAGACTACGTTCTGACACAGGCGAGCCTTGGTGGCTCGGTCGGTCCTGTTCG** | 142726 |
| WF-Teratorn | CCTCGACCTCCGCGCGCTTCACTCCGGCCAGCCCGAGCACAGATTCAGCGGCGACGAGGACGAGGCTCGAGACTACGTTCTGACACAGGCGAGCCTTGGTGGCTCGGTCGGTCCTGTTCG | 143556 |
| TeratornLC199500 | **AAAACTTCTCGCTGCATCCCCGGCAGCACTTGAGGTCCAGGTCCAAGTTCAGCCTGCTGACGGGTCTGCCCGGCAGGAAGACCTCGTTGTAACAAGGAGGCGGTCGGTCGCAAGGCGACG** | 142846 |
| WF-Teratorn | AAAACTTCTCGCTGCATCCCCGGCAGCACTTGAGGTCCAGGTCCAAGTTCAGCCTGCTGACGGGTCTGCCCGGCAGGAAGACCTCGTTGTAACAAGGGGGCGGTCGGTCGCAAGGCGACG | 143676 |
| TeratornLC199500 | **AGGAGGATGATCTTGCGATCAGGGTGCTTAGGTGGCCTCTGGAGGTTACGGTGTACCTGCGCGAAACCTTCCCCGTGTCGCGATCAACCGGGTCCTCCATGAGGAGCACAGGGTTTATCG** | 142966 |
| WF-Teratorn | AGGAGGATGATCTTGCGATCAGGGTGCTTAGGTGGCCTCTGGAGGTTACGGTGTACCTGCGCGAAACCTTCCCCGTGTCGCGATCAACCGGGTCCTCCATGAGGAGCACAGGGTTTATCG | 143796 |
| TeratornLC199500 | **ATCCCCAGTGACAGCGCGCGGGCAAGGGACCCAGGGGTCCCGAGGCTCGTTTCAGGCAGTAGTCGACCAGGTCGTTGAAGCGTAGCTCCGCGCCCGTGAACACAGTGAACGCGTAAGGAA** | 143086 |
| WF-Teratorn | ATCCCCAGTGACAGCGCGCGGGCAAGGGACCCAGGGGTCCCGAGGCTCGTTTCAGGCAGTAGTCGACCAGGTCGTTGAAGCGTAGCTCCGCGCCCGTGAACACAGTGAACGCGTAAGGAA | 143916 |
| TeratornLC199500 | **AGCCCTTCGGGCCGTCCGTCAGCGTTCCGGACAGATTGTTGAAGGCGGCATAGATCTCCCCGACCCCGTGAGCGATTCCCGGATCGACTGCGGCCACGTCTTCGTCCTCGCCGCTCGCGA** | 143206 |
| WF-Teratorn | AGCCCTTCGGGCCGTCCGTCAGCGTTCCGGACAGATTGTTGAAGGCGGCATAGATCTCCCCGACCCCGTGAGCGATTCCCGGATCGACTGCGGCCACGTCTTCGTCCTCGCTGCTCGCGA | 144036 |
| TeratornLC199500 | **GGTCCTCGCCATCATTGCCGCCGACCTCCTCTTCGCGGGAGCTCCCGTTTCCTCCGCTAATAGGGGTCTCCCCCTCAGCCTCTGCGCTCCCGGGCTCTTCGACGTCGAGTTTGTCACCTT** | 143326 |
| WF-Teratorn | GGTCCTCGCCATCATTGCCGCCGACCTCCTCTTCGCGGGAGCTCCCGTTTCCTCCGCTAATAGGGGTCTCCCCCTCAGCCTCTGCGCTCCCGGGCTCTTCGACGTCGAGTTTGTCACCTT | 144156 |
| TeratornLC199500 | **CTGTCTCGTCGACAGAGCCGTCCCGAGCCTCGCCCTTTCGGTGGTCCTCCGCAGGGTCCTCGTCTTCTTCGTAGTACCTCCATCGAGAAAACGAAAAGCGGTCCGAACCCACCAGTGGTA** | 143446 |
| WF-Teratorn | CTGTCTCGTCGACAGAGCCGTCCCGAGCCTCGCCCTCTCGGTGGTCCTCCGCAGGGTCCTCGTCTTCTTCGTAGTACCTCCATCGAGAAAACGAAAAGCGGTCCGAACCCACCAGTGGTA | 144276 |
| TeratornLC199500 | **CACCGTGCAGAGTGACGTGCTCCTCCGCACTACGGTCGAGCAGCCCCCCTGGCGCCACCATGTCCTCCACCTCGGGCGGCCCGGACCGGACCTCGGACGTGTAGTGCAGGCTCAAAGCTC** | 143566 |
| WF-Teratorn | CACCGTGCAGAGTGACGTGCTCCTCCGCACTACGGTCGAGCAGCCCCCCTGGCGCCACCATGTCCTCCAACTCGGGCGGCCCGGACCGGACCTCGGACGTGTAGTGCAGGCTCAAAGCTC | 144396 |
| TeratornLC199500 | **GACGGAACAGGGGGTAATGGTGCGTCACGTCGTACTGCCTGCGGAGCATCTCCTTCACGGAGCGCGCAAACAAGACCTGAAGCCGGGCTCCGAGCCTTGAGTCCTCGCCCTTCTCGTCCT** | 143686 |
| WF-Teratorn | GACGGAACAGGGGGTAATGGTGCGTCACGTCGTACTGCCTGCGGAGCATCTCCTTCACGGAGCGCGCAAACAAGACCTGAAGCCGGGCTCCGAGCCTTGAGTCCTCGCCCTTCTCGTCCT | 144516 |
| TeratornLC199500 | **CGGGCTTCTGCGCGTAGCAATTCAGGAAGAGAGAACAGTCCACTCCTTTGGACAGGCACAGTCTCGTGAGCTCGTCCTGGCGCGCCCGCGCACTGAAGGACGCCCGCGTGCGTCTGTGTGAA** | 143806 |
| WF-Teratorn | CGGGCTTCTGCGCGTAGCAATTCAGGAAGAGAGAACAGTCCACTCCTTTGGACAGGCACAGTCTCGTGAGCTCGTCCTGGCGCGCCCGCGCACTGAAGGACGCCCGCGTGCGTCTGTGTGAA | 144636 |
| TeratornLC199500 | **AGCGCACATCTCGATCGAGGGCGCAGGCTCTGCCAGAGCACCCGAGCAGGACGAGGTCTACGAGTGCTTGCACGACAATGCTCGCGCAGCTCTCAAAGTTTCGATCCAATATCCTCGCCG** | 143926 |
| WF-Teratorn | AGCGCACATCTCGATCGAGGGCGCAGGCTCTGCCAGAGCACCCGAGCAGGACGAGGTCTACGAGTGCTTGCACGACAATGCTCGCGCAGCTCTCAAAGTTTCGATCCAATATCCTCGCCG | 144756 |
| TeratornLC199500 | **AGCATTTGTCTCCGGGGCGCAGGTTCACAGCTTCGGACGACGCGTTCCACACGTTCCACTCATTCTCCTCAGATCCTCTTGTCAGGTAGCAGACGGGTTTGTACACGCGGTCAGGCGAGG** | 144046 |
| WF-Teratorn | AGCATTTGTCTCCGGGGCGCAGGTTCACAGCTTCGGACGACGCGTTCCACACGTTCCACTCCTTCTCCTCAGATCCTCTTGTCAGGTAGCAGACGGGTTTGTACACGCGGTCAGGCGAGG | 144876 |
| TeratornLC199500 | **GCCTCTGAGTTCCTCCCAGAGGACCAGAGCGCTGCTGCTGCTGCTGCTCTGCCGTGTTCACCCTTCTTTTCGCACGCTTCAT**CGTGTGACTGACGTTCTGCAAGAGGGTGCGATGTTGCT | 144166 |
| WF-Teratorn | GCCTCTGAGTTCCTCCCAGAGGACCAGAGCGCTGCTGCTGCTGCT---CTGCCGTGTCCACCCTTCTTTTCGCACGCTTCATCGTGTGACTGACGTTCTGCAAGAGGGTGCGATGTTGCT | 144993 |
| TeratornLC199500 | GATATTCTTTTCCTACTTTT-TTTTTTCCTTCTCTTTCGGTAACAATCCACAAAGTGAAATTCGTCTCAGATTCTCGGGCCCGACGGGGACGAGGCGCAGGGTGAGGCTCGCTCGCGCCT | 144285 |
| WF-Teratorn | GATATTCTTTTCCTACTTTTATTTTTTCCTTCTCTTTCGGTAACAATCCACGAAGTGAAATTCGTCTCAGATTCTCGGGCCCGACGGGGACGAGGCGCAGGGCGAGGCTCGCTCGCGCCT | 145113 |

TeratornLC199500 AGAGAACTCGCCTCCGAAGCTCGGTGACCCG**TCACAGAACCGAATTTAGGAGGGTGTCTTGTAGAGCGCCCGCCAGCACTCTTATTTTGTGATAACAGCGAGGCCCGTGCTCCCCTGGGC** 144405
WF-Teratorn AGAGAACTCGCCTCCGAAGCTCGGTGACCCGTCACAGAACCGAATTTAGGAGGGTGTCTTGTAGAGCGCCCGCCAGCACTCTTATTTTGTGATAACAGCGAGGCCCGTGCTCCCCTGGGC 145233

TeratornLC199500 **AGCGGCCGCACATGTAGCACCACAGTTTCTCGGACAACGTTACGTATAACCCGTGGGTGAGATGAACTTTCAGAGAAATGTCCGTCCCGGCCGGCACCCACACAGCCTGAGGGTGAGCGA** 144525
WF-Teratorn AGCGGCCGCACATGTAGCACCACAGTTTCTCGGACAACGTTACGTATAACCCGTGGGTGAGATGAACTTTCAGAGAAATGTCCGTCCCGACCGGCACCCACACAGCCTGAGGGTGAGCGA 145353

TeratornLC199500 **TTAGGCCTCTAGTGAACCTGGCTTGGCCCAACTCAATGGCGATGCGCTCAGGTGTGTAGTTTGAAGTCAGGACGGGGCAAGCGCCTCCCGAGACGAAAGCGTGCTGGTTCAGATAGGTCT** 144645
WF-Teratorn TTAGGCCTCTAGTGAACCTGGCTTGGCCCAACTCAATGGCGATGCGCTCAGGTGTGTAGTTTGAAGTCAGGACGGGGCAAGCGCCTCCCGAGACGAAAGCGTGCTGGTTCAGATAGGTCT 145473

TeratornLC199500 **CGCCTGGCGCCGCAACGACCACGTCCATGCCGCAGGAGAAACATCGACTGTGCCCCGTGGGCTCGGCGGGAGCACGGCCTTGGGCGGCCTGCGAAATACCCGTTTCATCGTTCTCTCTGG** 144765
WF-Teratorn CGCCTGGCGCCGCAACGACCACGTCCATGCCGCAGGAGAAACATCGACTGTGCCCCGTGGGCTCGGCGGGAGCACGGCCTTGGGCGGCCTGCGAAATACCCGTTTCATCGTTCTCTCTGG 145593

TeratornLC199500 **GCATTCCCAGCCGTACCATTTCTTGCGCAAAATACGAATTGTCGTCCGGGTGGACGTCTCGGCCTCCATTTGTGCTGTTCAACCACTCGAGCCTGCGCGACATGGAGACGTCAAAGTACA** 144885
WF-Teratorn GCATTCCCAGCCGTACCATTTCTTGCGCAAAATACGAATTGTCGTCCGGGTGGACGTCTCGGCCTCCATTTGTGCTGTTCAACCACTCGAGCCTGCGCGACATGGAGACGTCAAAGTACG 145713

TeratornLC199500 **GGTCTAGGCCGATCGGCACGTACACGCCGTCCCTGCGGACGATGTTGCTGTAGACCGTAGGTTCGTCTCCCTGTCGTCGCTCCAT**CCCCTGGACTAGTTTGACACCTGATGCGAGCCTGG 145005
WF-Teratorn GGTCTAGGCCGATCGGCACGTACACGCCGTCCCTGCGGACGATGTTGCTGTAGACCGTAGGTTCGTCTCCCTGTCGTCGCTCCATCCCCTGGACTAGTTTGACTCCTGATGCGAGCCTGG 145833

TeratornLC199500 CTGCAAAATGTTTGTGAAACAGAATCCTGGAGCCTTATAAAGAGTCACTTACCTGGCACAGATGTCTGAAGTTGGTG**TTACGAAATAAAATGAAAGGAGAGAGAGAGAGAAAGAAAAAAGCA** 145125
WF-Teratorn CTGCATAATGTTTGTGAAACAGAATCCTCGAGCCTTATAAAGAGTCACTTACCTGGCACAGATGTCTGAAGTTGGTGTTACGAAATAAAATGAAAGGAGAGAGAGAGATAAAGAAAAAAGCA 145953

TeratornLC199500 **AAGACAGCAAGAAGAAGGAGCGAGGTGACAGAGAATGAGATAGATAGGGACGGACGAGGAGGACTAGTAAAAGAGTGCCCACTCACGTCGTTGCTGCTGGACAGAAAGTGAGCGAAGGAC** 145245
WF-Teratorn AAGACAGCAAGAAGAAGGAGCGAGGTGACAGAGAATGAGATAGATAGGGACGGACGAGGAGGACTAGTAAAAGAGTGCCCACTCACGTCGTTGCTGCTGGACAGAAAGTGAGCGAAGGAC 146073

TeratornLC199500 **GAGCCCTTGTACCCGGGATCGAATCTGTACACCCGGTTCTCGGGGCTTCCGTGAACGCTCACGGTCCTCTCGGTCTGAAGGAAGGATGGGAGTCTCTGGAGTTCCTCCATGGCGGCTTCC** 145365
WF-Teratorn GAGCCCTTGTACCCGGGATCGAATCTGTACACCCGGTTCTCGGGGCTTCCGTGAACGCTCACGGTCCTCTCGGTCTGAAGGAAGGATGGGAGTCTCTGGAGTTCCTCCATGGCGGCTTCC 146193

TeratornLC199500 **ACCCCTAGCCTCAGTGTGGTGCACATGTGTTTGTTCTGAGCCTGCACCATGACGTTGATGTCCAAGTGCAAGAGCACCGCCGCCAGTATCAGCCTCATCGCCAGGCTCTTGCCGCACTGC** 145485
WF-Teratorn ACCCCTAGCCTCAGTGTGGTGCACATGTGTTTGTTCTGAGCCTGCACCATGACGTTGATGTCCAAGTGCAAGAGCACCGCCGCCAGTATCAGCCTCATCGCCAGGCTCTTGCCGCACTGC 146313

TeratornLC199500 **CTCGGGGCGAGGCACAGGGTGTACGGGTTGGCGTATCGAGCAAACTCCTTTTCCACTTCGTGCAGGTGGCTGCGCGCCGGGTCCTGGTAATCGTAGCTCTCGGAACCCAGAGGCGCGAGG** 145605
WF-Teratorn CTCGGGGCGAGGCACAGGGTGTACGGGTTGGCGTATTGAGCAAACTCCTTTTCCACTTCGTGCAGGTGGCTGCGCGCCGGGTCCTGGTAATCGTAGCTCTCGGAACCAAGAGGCGCGAGG 146433

TeratornLC199500 **TTCAGCGTGCTCAGCAGGGCGTGCTTGTGCCTGAAGAGCTCGTCTCCGAACTGTGCCTCGGTGACGCCCAGCACGACTCCTCTCAAGAGCTCCATCTGAAAGGGCTCCATGCGCATGCCG** 145725
WF-Teratorn TTCAGCGTGCTCAGCAGGGCGTGCTTGTGCCTGAAGAGCTCGTCTCCGAACTGTGCCTCGGTGACGCCCAGCACGACTCCTCTCAAGAGCTCCATCTGAAAGGGCTCCATGCGCATGCCG 146553

TeratornLC199500 **TTCGTTTGGAAATATTCAAACACCCGTTCTACCATGCGCCGAAATCGATACATGAGGCGATCGGAGGCGTCCCCTTCTAGATTTCCAGAGGCCCGTATCTCGCGCACCCTCGCCTTCGCG** 145845
WF-Teratorn TTCGTTTGGAAATATTCAAACACCCGTTCTACCATGCGCCGAAATCGATACATGAGGCGATCGGAGGCGTCCCCTTCTAGATTTCCAGAGGCCCGTATCTCGCGCACCCTCGCCTTCGCG 146673

TeratornLC199500 **CCTTCCAAACCCAGCTCGCGAATCAAGACGGCTTGCGCCACTCGGTCTGTCATCAGCCGATTGTATTCCGAGACCCTCTCCCGTTCGGAGCGCCTGCTCTCCTCCTCGACCTGATCCTTG** 145965
WF-Teratorn CCTTCCAAACCCAGCTCGCGAATCAAGACGGCTTGCGCCACTCGGTCTGTCATCAGCCGGTTGTATTCCGAGACCCTCTCCCATTCCGAGCGCCTGCTCTCCTCCTCGACCTGATCCTTG 146793

TeratornLC199500 **TAGAGCCTGAGCCGTACGCAGTCCACAGCCACAGCCAAAGGCACCGGCGGCCTGGAGCCAGGCACGGGATCGCGCACGATTTCTCTTCGCAACCAAAGGATGTCATCAGGGTCTGGTGAC** 146085
WF-Teratorn TAGAGCCTGAGCCGTACGCAGTCCACAGCCACAGCCAAAGGCACCGGCGGCCTGGAGCCAGGCACGGGATCGCGCACGATTTCTCTTCGCAACCAAAGGATGTCATCAGGGTCTGGTGAC 146913

TeratornLC199500 **AAAGCTCTCCACGGAGCTCTCGACAGGCCAGGGCGCGCGAGGCGCAATGGCCTAGGTCCACGCGTGAGCTCGCTGCTCGAGGAGAGGGCAGCGGGTGAGGCGCTGTGTCCGGCGGAGCTC** 146205
WF-Teratorn AGAGCTCTCCACGGGGCTCTCGACAGGCCAGGGCGCGCTAGGCGCAATGGCCCAGGTCCACGCGTGAGCTCGCTGCTCGAGGAGAGGGCAGCGGGTGAGGCGCTGTGTCCGGCGGAGCTC 147033

TeratornLC199500 **ATCGCGCACAGTGCCTGGGCTCGCCGAACCCGCGACGCAGAAAGACGTCCCTGCAAGGCCGCCGTGAGCGAGTTCGTGCTGCCGCTCAAAATCCACTTCAGGTACTGCTGGCTCTTCTCC** 146325
WF-Teratorn ATCGCGCACAGTGCCTGGGCTCGCCGAACCCGCGACGCAGAAAGACGTCCCTGCAAGGCCGCCGTGAGCGAGTTTGTGCTGCCGCTCAAAATCCACTTCAGGTACTGCTGGCTCTTCTCC 147153

TeratornLC199500 **GGGATTCCCGTCAGCGATGCGGCGCGACCGAGAGACGCGTGCCTGCCGGGAGTTTTAACAGATCTCGAAAGAGTTCGCTTCAACCGCGTAATGATGTCGGGAGACGGCGATCCCGAATTC** 146445
WF-Teratorn GGGATTCCCGTCAGCGATGCGGCGCGACCGAGAGACGCGTGCCTGCCGGGAGTTTTAACAGATCTCGAAAGAGTTCGCTTCAACCGCGTAATGATGTCGGGAGACGGCGATCCCGAATTC 147273

TeratornLC199500 **TGCTGCGAGGGAAGGCTCTCGGACGTCTTCGGGATGAAATCTTTCTGGCTCACCACCGACGGAAGGTTGGTCGTCGTCGACTCGCCTTTCGAAGACGCCAAGTTGTCTCTGCTGAGGCAC** 146565
WF-Teratorn TGCTGCGAGGGAAGGCTCTCGGACGTCTTCGGGATGAAATCTTTCTGGCTCACCACCGACGGAAGGTTGGTCGTCGTCGACTCGCCTTTCGAAGACGCCAAATTGTCTCTGCTGAGGCAC 147393

| | | |
|---|---|---|
| TeratornLC199500 | **CGCAATCTCACGGACGCCCTGGCTCTCTGGGAAGTGCAGCCCGAGCTGTGCTCCCGGCTCACCGAGGACTCGAGCGCTGCGCACTCAGCCTTGATCGGGCTTTGCGGGGACGCGTGCCGC** | 146685 |
| WF-Teratorn | CGAAATCTCACGGACGCCCTGGCTCTCTGGGAAGTGCAGCCCGAGCTGTGCTCCCGGCTCACCGAGGACTCGAGCGCTGCGCACTCAGCCTTGATCGGGCTTTGCGGGGACGCGTGCCGC | 147513 |
| TeratornLC199500 | **CTGAGCCGAGGCCGCTTCTACAACCTGTACGCCAGGTACCCTTTGCCCGTGGAGAAGCAAAGAGGCGGCCTCGCTTCCGCACCTCCCCAGAATCGTAAGGCCCCCGTCGAGGAATGCGAC** | 146805 |
| WF-Teratorn | CTGAGCCGAGGCCGCTTCTACAACCTGTATGCCAGGTACCCTTTGCCCGTGGAGAAGCAAAGCGGCGGCCTCGCTTCCGCACCTCCCCAGAATCGTAAGGCCCCCGTCGAGGAATGCGAC | 147633 |
| TeratornLC199500 | **GTCTCCGCTCGAGAGATACGTTCTGGTCAGGAGTTGCTCTCGATCCTCGCACCTCTCTACGCCACGTATTTCGCCCTGGTGGAGGGGGACCCGGCTTTGGATGAGCTCTGGCGCAGAGTG** | 146925 |
| WF-Teratorn | GTCTCCGCTCGAGAGATACGTTCTGGTCAGGAGTTGCTCTCGATCCTCGCACCTCTCTACGCCACGTATTTCGCCTTGGTGGAGGGGGACCCGGCTCTGGATGAGCTCTGGCGCAGAGTG | 147753 |
| TeratornLC199500 | **GATCTCCTCGGGGCGAGGTGGCTGGTTCGAGAAGAGTTTTCGGAGCTGAGCCTCTCCGTCAGGCTGCGCGACAGTCGGTTCGCGAGATTGGCGTTCGCCTGCGCTTTCTTTCTCTCCGAC** | 147045 |
| WF-Teratorn | GATCTCCTCGGGGCGAGGTGGCTGGTTCGAGAAGAGTTTTCGGAGCTGAGCCTCTCCGTCAGGCTGGGCGACAGTCGGTTCGCGAGATTGGCGTTCGCCTGCGCTTTCTTTCTCTCCGAC | 147873 |
| TeratornLC199500 | **TGTTGA**GCATGGCAGGTCTTTTGGGCGCAGAAACCGCCGCTCATTAACCCTGGACGCTGACCCGCGAAGGCCGCGGTGATCAGCTCTCGCTCCACCGAAGAC**ATGGACCCTCCCAAGTTC** | 147165 |
| WF-Teratorn | TGTTGAGCATGGCAGGTCTTTTGGGCGCAGAAACCGCCGCTCATTAACCCTGGACGCTGACCCGCGAAGGCCGCCGTGATCAGCTCTCGCTCCACCGAAGACATGGACCCTCCCAAGTTC | 147993 |
| TeratornLC199500 | **GAAGACAAGGAAGAAACGATTAGGTCCATGCTCAGCACCCGCTTTGGCAGCTACCGCGTGGTCTTCAAGCAAGCTGCGTGCGACGTGTGGGTGCGAGGCAGAGATTGCACGGTCCAGTAC** | 147285 |
| WF-Teratorn | GAAGACAAGGAAGAAACGATTAGGTCCATGCTCAGCACCCGCTTTGGCAGCTACCGCGTGGTCTTCAAGCAAGCTGCGTGCGACGTGTGGGTGCGAGGCAGAGATTGCACGGTCCAGTAC | 148113 |
| TeratornLC199500 | **CAGGTGCGCAAGCGCCCGGACGGGGCCGGGTGGTACGCCTACTCGAAATCCGAGTCCAGAATATTGGGGGAGTCGGCGGAAGACGTGATAAAGCTGTGCGTGAAGGACACGATTTCCGAA** | 147405 |
| WF-Teratorn | CAGGTGCGCAAGCGCCCGGACGGGGCCGGGTGGTACGCCTACTCGAAATCCGAGTCCAGAATATTGGGGGAGTCGGCGGAAGACGTGATAAAGCTGTGCGTGAAGGACACGATTTCCGAA | 148233 |
| TeratornLC199500 | **CAGGACGAACGGGAGGACGAGGTGGTGGGGAAGGAGGACAACGAGGATGAGAACGAGGCGTCGCGGCACGACTACGAAGTGATACGCAAGCTCTTGGAGCGCGAGAAAGACAGCCGGCCC** | 147525 |
| WF-Teratorn | CAGGACGAACGGGAGGACGAGGTGGTGGGGAAGGAGGACAACGAGGATGAGAACGAGGCGTCGCGGCACGACTACGAAGTGATACGCAAGCTCTTGGAGCGCGAGAAAGACAGCCGGCCC | 148353 |
| TeratornLC199500 | **CAGGGAGAGACGATCCCGGACGTCGGGGACTATCGAAGGGCCGGGGGGTGGTTGAACTCGGGGCGGAGATGCGAGGATGGCGACCGATCGACCGAGGACGACGAGAGTGGCGATGAGCGA** | 147645 |
| WF-Teratorn | CAGGGAGAGACGATCCCGGACGTCGGGGACTATCGAAGGGCCGGGGGGTGGTTGAACTCGGGGCGGAGATGCGAGGATGGCGACCGATCGACCGAGGACGACGAGAGTGGCGATGAGCGA | 148473 |
| TeratornLC199500 | **GACCGCTTGCGGGAAAGCTTTGGCAACCTCAGGCTCGGGGCAACGGATCGTGTGAAAGTCCAGCACCGCCATGCCGCTCGCCCCGTCAGCCCGAGTCTGGCTAACTGGGACCTCGACCCA** | 147765 |
| WF-Teratorn | GACCGCTTGCGGGAAAGCTTTGGCAACCTCAGGCTCGGGGCAACGGATCATGTGAAAGTCCAGCACCGCCATGCCGCTCGCCCCGTCAGCCCGAGTCTGGCTAACTGGGACCTCGACCCA | 148593 |
| TeratornLC199500 | **TCCGAAAGACGCTGGCCTGGCTTCCCCCCTCCAGTCCGTCGAGACCTGAAACCCGTGGAGCGCCTCTTCGCCAGGGGAATCCGTTCGATCTATTGTGGTGACGAGGAAGTCACGGATTCG** | 147885 |
| WF-Teratorn | TCCGAAAGACGCTGGCCTGGCTTCCCCCCTCCAGTCCGTCGAGACCTGAAACCCGTGGAGCGCCTCTTCGCCAGGGGAATCCGTTCGATCTATTGTGGTGACGAGGAAGTCACGGATTCG | 148713 |
| TeratornLC199500 | **AAAGGTCCATCGGAGTACGACAGCGAAGAGGATGGATACGCGTGGGGCTGCCGAGATAACAAGTCCTGA**GCGTCGCAGCAAGCACCCGTACAAGCGCAAGGCAACGGGGCTGAATTTAGA | 148005 |
| WF-Teratorn | AAAGGTCCATCGGAGTACGACAGCGAAGAGGATGGATACGCGTGGGGCTGCCGAGATAACAAGTCCTGAGCGTCGCAGCAAGCACCCGTACAAGCGCAAGGCAACGGGGCTGAATTTAGA | 148833 |
| TeratornLC199500 | AATTTAGAGTGTGTGTAATAATTTTTTTTTTAATTTTATTTCATTTTCTACCCATTTTCATCGGAACCACTTTACACCTCAGAGCTCCGGTCTATGTTCCGTGGAGATGAACGAGGTCGG | 148125 |
| WF-Teratorn | AATTTAGAGTGTGTGTAATAATTTTTTTTTTAATTTTATTTCATTTTCTACCCATTTTCATCGGAACCACTTTACACCTCAGAGCTCCGGTCTATGTTCCGTGGAGATGAACGAGGTCGG | 148953 |
| TeratornLC199500 | ACGAGCCTTTCCCGTCTACTTGAGCCCATAAGCAGCATAAGAGCGAAGAGACGAAGCCTCGGTCTTGTCTACTTGTCTGTTTCTGGAGCGTACACTCACACGAAAGCAGTCTGATACAAC | 148245 |
| WF-Teratorn | ACGAGCCTTTCCCGTCTACTTGAGCCCATAAGCAGCATAAGAGCGAAGAGACGAAGCCTCGGTCTTGTCTACTTGTCTGTTTCTGGAGCGTACACTCACACGAAAGCAGTCTGATACAAC | 149073 |
| TeratornLC199500 | AAGCCCATCCGCCACTGCAGCA**ATGTCAACCACTCCAGAACAAGACTTGTACTCCTCGGATGTGACGCGCACCAGGGCATCGTCCTTCACGGTCACAACGGACACTGAATCGTTCACCAC** | 148365 |
| WF-Teratorn | AAGCCCATCCGCCACTGCAGCAATGTCAACCACTCCAGAACAAGACTTGTACTCCTCGGATGTGACGCGCACCAGGGCATCGTCCTTCACGGTCACAACGGACACTGAATCGTTCACCAC | 149193 |
| TeratornLC199500 | **GGCCCGCAGTCCCGAAGGAGACGGTGTTGTTCCCGTATTCTCGCCCAGAGAGATGGCCCGCTCGGAGCATGACGGCGATTCGGGCCTTGAGGAGACCGAGGCCGAACCGGACAGCGTCCT** | 148485 |
| WF-Teratorn | GGCCCGCAGTCCCGAAGGAGACGGTGTTGTTCCCGTATTCTCGCCCAGAGAGATGGCCCGCTCGGAGCACGACGGCGATTCGGGCCTTGAGGAGACCGAGGCCGAACCGGACAGCGTCCT | 149313 |
| TeratornLC199500 | **TTACCTCCAGCTCTCGTCGAACCCTGGGATCGCAGAGCCGCGGATCGATTTCGAGCCCCCGTCCGATTCATCGGGAAGCGAGATCTACAGCTCCTCCGATGACGATGAGGAAGACAGCAG** | 148605 |
| WF-Teratorn | TTACCTCCAGCTCTCGTCGAACCCTGGGATCGCAGAGCCGCGGATCGATTTCGAGCCCCCGTCCGATTCATCGGGAAGCGAGATCTACAGCTCCTCCGATGACGATGAGGAAGACAGCAG | 149433 |
| TeratornLC199500 | **CATAGAAAATTATGATCGCTACGACCCAGCACTCCGGTGGCACCACCCAGACGTTGTAGACCACATGCGAGGACTGGGCCTGGCCATCGACGAAGAAGGATACTGGTTCGATCCGGGTTC** | 148725 |
| WF-Teratorn | CATAGAAAATTATGATCGCTACGACCCAGCACTCCGGTGGCACCACCCAGACGTCGTAGACCACATGCGAGGACTGGGCCTGGCCATCGACGAAGAAGGATACTGGTTCGATCCGGGTTC | 149553 |
| TeratornLC199500 | **CTTCCCCCTCTTCTCTGAGCCTAAAGGCGTCGTGTTTGAAGACGACGATTGA**AGTTCTGCTTGCTGTGTGTGTGTCTGTTCACATTTTCGCTACCATCTTTATCTTATGAGACTCAATAA | 148845 |
| WF-Teratorn | CTTCCCCCTCTTCTCTGAGCCTAAAGGCGTCGTGTTTGAAGACGACGATTGAAGTTCTGCTTGCTGTGTGTGTGTCTGTTCACATTTTCGCTACCATCTTTATCTTATGAGACTCAATAA | 149673 |

```
TeratornLC199500  AGGTTCTGCTATCTGATAACCAGTGTGCTTGGGTTTGTCGAGAGCTCCCGACGCTCGTGCAGTGAAAGGTTTAGGGGGAGCATGGGTTCGAAACCCTAACGCCGGGAGTTTGAAATGGGG  148965
WF-Teratorn       AGGTTCTGCTATCTGATAACCAGTGTGCTTGGGTTTGTCGAGAGCTCCCGACGCTCGTGCAGTGAAAGGTTTAGGGGGAGCATGGGTTCGAAACCCTAACGCCGGGAGTTTGAAATGGGG  149793

TeratornLC199500  GACATAAAATGAAATAAAGAAACGTGACCACCGGCAACAGCAAGCGTTTGAGCCTTTTCCCTTTTCATTTTATTGAGGTGAAACAAAACAAAACACAGCTGCATTTCACGTACAATGAAC  149085
WF-Teratorn       GACATAAAATGAAATAAAGAAACGTGACCACCGGCAACAGCAAGCGTTTGAGCCTTTTCCCTTTTCATTTTATTGAGGTGAAACAAAACAAAACACAGCTGCATTTCACGTACAATGAAC  149913

TeratornLC199500  CTCTCCGCATATTCTCACGGAGCTCTCTCCCCCGTGAATCTCGTCTCTTGTGTGGCAAAGGAGCAGAAACGCGGAGAGCACGTCGTCCCTGGTGTATTCGCCCCCCTTGGACTTTTTGCC  149205
WF-Teratorn       CTCTCCGCATATTCTCACGGAGCTCTCTCCCCCGTGAATCTCGTCTCTTGTGTGGCAAAGGAGCAGAAACGCGGAGAGCACGTCGTCCCTGGTGTATTCGCCCCCCTTGGACTTTTTGCC  150033

TeratornLC199500  TCCGGTGACCACGGTCCCCCCGCTGCCCTTGGCTGACTCCTTGATCGAGACGGCAGCCATTTCCTCCCTCAGTTTGCCCAAGAGCCGCTTCCCGTGCTCTGCGTGCAACCTGTCGGCCAA  149325
WF-Teratorn       TCCGGTGACCACGGTCCCCCCGCCGCCCTTGGTTGACTCCTTGATCGAGACGGCAGCCATTTCCTCCCTCAGTTTGCCCAAGAGCCGCTTCCCGTGCTCTGCGTGCAACCTGTCGGCCAA  150153

TeratornLC199500  CCTGCTGTCGGGTAGGGAGATGGTGTCTCGGATCTCCAGAGCTCGAGCGGTCGCGTCCGTGGCCAAATTGATACCGCTCCTCGACCTCCTGTACATGTGCACGGCCATGGAGGAGCCGGT  149445
WF-Teratorn       CCTGCTGTCGGGTAGGGAGATGGTGTCTCGGATCTCCAGAGCTCGAGCGGTCGCGTCCGTGGCCAAATTGATACCGCTCCTCGACCTCCTGTACATGTGCACGGCCATGGAGGAGCCGGT  150273

TeratornLC199500  TCTCGTGCTGGAGCCGGCGCCCCGAGAGACGCCCAGGGTGGCTTCCACGGCTCTGCCCTTCCCCACAGCCTCGGCCAGCACGGCCCAGCCGAGGGACATGACGCACTCGCAGTAGCCCAG  149565
WF-Teratorn       TCTCGTGCTGGAGCCGGCGCCCCGAGAGACGCCCAGGGTGGCTTCCACGGCTCTGCCCTTCCCCACAGCCTCGGCCAGCACGGCCCAGCCGAGGGACATGACGCACTCACAGTAGCCCAG  150393

TeratornLC199500  ATCGTCCCTCTCTATGGCGGACACGGTGCTGAGCACCATGGCCAGTTTGGTCTTGGCACTGACGTGTTTCCCCAGCATGTACCTGGAGTTGACCTGGGACCACTTGCTGTAAAAGAGAAC  149685
WF-Teratorn       ATCGTCCCTCTCTATGGCGGACACGGTGCTGAGCACCATGGCCAGTTTGGTCTTGGCACTGACGTGTTTCCCCAGCATGTACCTGGAGTTGACCTGGGACCACTTGCTGTAAAAGAGCAC  150513

TeratornLC199500  CCTGAAGCCAGCGCGTGCCAGAGCGTGTTCGAGCAGCTCCTTCAGGTCGGCCAGCGCGTTCACGTAGGTGTTTTGCTCAAACACCACAACCACCGGCACCTCTTTCCAGTGCGCTGAGGG  149805
WF-Teratorn       CCTGAAGCCAGCGCGTGCCAGAGCGTGTTCGAGCAGCTCCTTCAGGTCGGCCAGCGCATTCACGTAGGTGTTTTGCTCAAACACCACAACCACCGGCACCTCTTTCCAGTGCGCTGAGGG  150633

TeratornLC199500  CAAAAGGACTTTGAGCAGCCTGACGTGAGCGCTCAAAAGGGCCGCGTACACCCTGGTGACTTGATCTAGGTCACGTATCTCCAGCTCGTCCAGTGACATGATCTGCAAAGAGACGGGGGA  149925
WF-Teratorn       CAAAAGGACTTTGAGCAGCCTGACGTGAGCGCTCAATAGGGCCGCGTACACCCTGGTGACTTGATCTAGGTCACGTATCTCCAGCTCGTCCAGTGACATGATCTGCAAAGAGACGGGGGA  150753

TeratornLC199500  GAAAGCAGGTTTAGGTACTTTGTGGCGAAAAACGAATTGCACATAGGAAACCGCTGGCTGCTCAACACGATCACATTCCGCTTCTTAATCATCATCATCATCGTCATCAT---TATCATC  150042
WF-Teratorn       GAAAGCGGGTTTAGGTACTTTGTGGCGAGAAACGAATTGCACATAGGAAACCGCTGGGCGCTCAACACGATCACATTCAGCTTCTTAATCATCATCATCATCATCATCATCATTATCATC  150873

TeratornLC199500  ATCATCATCACTTGTTGCAATATCACCCTTCGAGTAAGACGAAGGTTACGCTTTGTCAACATGTCTTTCTGTCCCGCGTGCCGCAGGGCTCGAAGACAACTCTTGAGCCTCATGCGCATG  150162
WF-Teratorn       ATCATCATCACTTGTTGCAATATCACCCTTCGAGTAAGACTAAGGTTACGCTTTGTCAACATGTCTTTCTGCCCCGCGTGCCGCAGGGCTCGAAGACAACTCTTGAGCCTGATGAGCATG  150993

TeratornLC199500  CCGTGGGAGCTGCTGCTCTCCCGGTACGAGTTCATGGTAGCCCTCGCTTCGTTCCGCCGCAACATGGACCTGTTCGTCGAGATGGACATCGGGCAACTGAGGACGGCGTACAGTCAGGTG  150282
WF-Teratorn       CCGTGGGAGCTGCTACTCTCCCGGTACGAGTTCATGGTAGCCCTCGCTTCGTTCCGCCGCAACATGGACCTGTTCGTCGAGATGGACATCGGGCAACTGAGGACGGCGTACAGTCAGGTG  151113

TeratornLC199500  GTGTCGTTCTCGCCCGCGAGAATACTGGTGACAGACTCGCTCGTCTCTCATTACGTGCTGAGTTCGAGGAACGAGCACTCGGGGCGAGAAGCTCTGGCTTCGGCCCCGCGCGTCCCTGTG  150402
WF-Teratorn       GTGTCGTTCTCGCCCGCGAGGATACTGGTGACAGACTCGCTCGTCTCTCATTACGTGCTGAGTTCGAGGAACGAGCACTCGGGACGAGAAGCTCTGGCTTCGGCCCCGCGCGTCCCCGTG  151233

TeratornLC199500  GAGCTTTCGGAGGCAGTGACCCTTGCCCTCGGAAACAGCGTGCCGAGACGTTTTGTGCACGATTGCATGAGCCAGCATGAGTGGCGTGGGCATGAGGAGATGGTTCTCTCTGCCTTCGCC  150522
WF-Teratorn       GAGCTTTCGGAGGCAGTGACCCTTGCCCTCGGAAACAGCGTGCCGAGACGTTTTGTGCACGATTGCATGAGCCAGCATGAGTGGCGTGGGCACGAGGAGATGGTTCTCTCTGCCTTCGCC  151353

TeratornLC199500  TCTGCAGAGTCTGTGCGCGGCGGGCCCCTTAAGTACCTGGCCCTGGTAAAGGCCAGCTCCCGGAACGTGGCCTCTCTGCACACGGTGGTGGCGCTCGCCATTACGGGCTGGCTGATGGAG  150642
WF-Teratorn       TCTGCAGAGTCTGTGCGCGGCGGGCCCCTAAAGTACCTGGCCCTGGTAAAGGCCAGCTCCCGGAACGTGGCCTCTCTGCACACGGTGGTGGCGCTCGCCATTACGGGCTGGCTGATGGAG  151473

TeratornLC199500  CTGTCGGGGCATAGCGCGGGGAGTTGCAAGTGCGGGGTCTGGGAACAGGCCCTGGAGCTCATCTCGTGCTCCGTGCCAGATGCGGAGACCTCCCTCAGCGTGCAGGACGAGAACCGCGTA  150762
WF-Teratorn       CTGTCGGGGCATAGCGTGGGGAGTTGCGAGTGCGGGGTCTGGGAACAGGCCCTGGAGCTCATCTCGTGCTCCGTGCCAGACGCGGAGACCTCCCTCAGCGTGCAGGACGAGAACCGCGTG  151593

TeratornLC199500  AGCGCGTACATCTGCACCCTGATGGACGTGAAGCCCGGGCCCGAGGAGCGAGCTCTTCCCGGGCCGGTTCCCGATTACGGGTACGTTTTTAGAGATCTTCTGGTGTCCGGTTTAGGGGGG  150882
WF-Teratorn       AGCGCGTACATCTGCACCCTGATGGACGTGAAGCCCGGGCCCCAGGAGCGAGCTCTTCCCGGGCCGGCTCCCGATTACGGGTACGTTTTTAGAGATCTTCTGGTGTCCGGTTTAGGGGGG  151713

TeratornLC199500  ACAGAGGAGGGAGGCGCCGATCGTGGGCGGGAGACCTACCACTAGCTTGCCTCCAGCGATCTCCACGGCGCAACAAACTCCGGCG-----------------------------------  150967
WF-Teratorn       ACAGAGGAGGAAGGCGCCGATCGTGGGCGGGAGACCTACCACTAGCTTGCCTCCAGCGATCTCCACGGCGCAACACACTCCGGCGCAGGACCTGGAACCTCCCCCGAAGGTTGGGTCCAG  151833

TeratornLC199500  ------------------------------------------------------------------------------------------------------------------------  150967
WF-Teratorn       ACACAAGTAAGCCTTGGTCACGATCGATCGGGCCACGTCTTCCAGGGCCCAACAGTTGTTGGACAGGAACCTGTTGATGACGGAAGCCTCGAACGGGACTGTGGTGTCGGTCTTGATGTC  151953
```

TeratornLC199500 ------------------------------------------------------------------------------------------------------------------------ 150967
WF-Teratorn CTCGGACGAAGTGCTGCTGCTGCCTGTGACCTCGCTCTCGAAACCCTTGGGTGAGACCATGTTCATGAGCTCCTTGAGCTGCCCGTCCACCGAAATGTGTTGCGGGCAGTAGACGCCGAG 152073

TeratornLC199500 -------------------------------------------------------------------------------------------------------------TTCAGCCAAGG 150978
WF-Teratorn GCAGCAGCAGGTGAGCCCCGGATCCCTTTCGTGGTATGCGCATTTGAAACTCTGAGACACCACGTGGTAAGCGGGGCCTCCCCGGTCGTCGCGGATGTCCGCGACCTTGTTCAGCCAAGG 152193

TeratornLC199500 CGTGTTGTTGACGTGGCTGGTGATGTGAACCTGTTTCCGTCCTCTGAACGCGATGTTGGGCAGGACGGCCAGAAGCACCTTGGAATTCACGAAGCCTACTTCGTCCACCAGGATGAGGTC 151098
WF-Teratorn CGTGTTGTTGACGTGGCTGGTGATGTGAACCTGTTTCCGTCCTCTGAACGCGATGTTGGGCAGGACGGCCAGAAGCACCTTGGAATTCACGAAGCCTACTTCGTCCACCAGGATGAGGTC 152313

TeratornLC199500 GGCGCTCTGTCCCCTGCTCGCCTGTGAAGAGAGGCAATAAAAATTGATAAGGAACAAAGAAAAGGGAAAGGAGAAGAACACAGAGAAAGGAAATACGAAGGGAGAGGAGGAGAATCGAGA 151218
WF-Teratorn GGCGCTCTGTCCCCTGCTCGCCTGTGAAGAGAGGCAATAAAAATTGATAAGGAACAAAGAAAAGGGAAAGGAGAAGAACACAGAGAAAGGAAATACGAAGGGAGAGGAGGAGAATCGAGA 152433

TeratornLC199500 GACGCTAAGGGAAATACGAAGGGAGAGGAGGAGAATCGAGAGACGCTAAGGGAAGCGCCG**ATGGACCGCTGGCCTTCAGAATTAATAGACGATCCGCAGTGGGTGAGCCACTGCGTGGAG** 151338
WF-Teratorn GACGCTAAGGGAAATACGAAGGGAGAGGAGGAGAATCGAGAGACGCTAAGGGAAGCGCCGATGGACCGCTGGCCTTCAGAATTAATAGACGATCCGCAGTGGGTGAGCCACTGCGTGGAG 152553

TeratornLC199500 **GAGGAATGCGCTTTTGACGACCGATACATCCCGGGCTGTTACGTATGCTCAGCGCACCTCAGCAAGCACTCGTGCTCGGGATTCGACTGCGCCCTCGTGCGGGCTATGGCCGCCCGAGTT** 151458
WF-Teratorn GAGGAATGCGCTTTTGACGACCGATACATCCCGGGCTGTTACGTATGCTCAGCGCACCTCAGCAAGCACTCGTGCTCGGGATTCGACTGCGCCCTCGTGCGGGCTATGGCCGCCCGAGTT 152673

TeratornLC199500 **TCTTACGCCTCGGCGTCGGGGGAATTTTCCTGCCCCGTGGTGGGGCTAGCCGCATTTCGCAGGGAGGACCCTGACTTCAGGGGGCCCTGGTACTCTGGGACGGAGGCGACGGTGGGCTAC** 151578
WF-Teratorn TCTTACGCCTCGGCGTCGGGGGAATTTTCCTGCCCCGTGGTGGGGCTAGCCGCATTTCGCAGGGAGGACCCTGACTTCAGGGGGCCCTGGTACTCTGGGACGGAGGCGACGGTGGGCTAC 152793

TeratornLC199500 **GGAGCATTGGAAGTTGACCCGCGGACGGGCGGACCTCGCGGAGGTCCGTGGGTCGAAGACCAGTGCGGCGTATCGGCGCGGGTGCTGAAATCCTGCCTGCCGGCCATGATTGCACTTTTC** 151698
WF-Teratorn GGAGCATTGGAAGTTGACCCGCGGACGGGCGGACCTCGCGGAGGTCCGTGGGTCGAAGACCAGTGCGGCGTATCGGCGCGGGTGCTGAAATCCTGCCTGCCGGCCATGATTGCACTTTTC 152913

TeratornLC199500 **TCCACCGTCCTCACGCGGGAATTTGCAGACGAGTACCGGACCGTTGAGCGCGGAGAGGGTCAAGTCCCGTCGCAGCGCTGCCGATGGACCTCTCGAGTGTTCAGGCTCGCGGGCTGTATC** 151818
WF-Teratorn TCCACCATCCTCACGCGGGAATTTGCAGACGAGTACCGGACCGTTGAGCGCGGAGAGGGTCAAGTCCCGTCGCAGCGCTGCCGATGGACCTCTCGAGTGTTCAGGCTCGCGGGCTGTATC 153033

TeratornLC199500 **GAGTTCCGGACCGTCGGCGAGCGGGTCCGAGCGGACCTAGGAGATCTGGGAACGGTGAGGGACGCCGCTCTCAAAAGCGCAGCCGCCGTGTCCACCCTTTTCTGCTCACCCGAAGCTAGG** 151938
WF-Teratorn GAGTTCCGGACCGTCGGCGAGCGGGTCCGAGCGGACCTAGGAGATCTGGGAACCGTGAGGGATGCCGCTCTCAAAAGCGCAGCCGCCGTGTCCACCCTTTTCTGCTCACCCGAAGCTAGG 153153

TeratornLC199500 **ATCAAGCGCTTCGTGAAGAAAATGGTGAACAAGCAAAGCGACAAGCACTCGAGGAGAGACGGCTTTTACACTCTGGTGATCATGGCCTCTCTTTGCTTCTTCCCTTACTTAAAACTCGCC** 152058
WF-Teratorn ATCAAGCGCTTCGTGAAGAAAATGGTGAACAAGCAAAGCGACAAGCACTCGAGGAGAGACGGCTTTTACACTCTGGTGATCATGGCCTCTCTTTGCTTCTTCCCTTACTTAAAACTCGCC 153273

TeratornLC199500 **GTGCTGAGCGAAATGTTCGACAACAGGACCCTGGTGCTCATGTTTTTCGACAAGACCCTCGCCAGGCACAACCAGTGGTCCATGCTCTCTACCAACGTCCACCTGCACGTGAAACGGCCT** 152178
WF-Teratorn GTGCTGAGCGAAATGTTCGACAACAGGACCCTGGTGCTCATGTTTTTCGACAAGACCCTCGCCAGGCACAACCAGTGGTCCATGCTCTCTACCAACGTCCACCTGCACGTGAAACGGCCT 153393

TeratornLC199500 **CAGTTTCAGAGAGCGCTCGCCGACGCCGACCCGGGTAGACTCCTGCCACTGGTCGTGAGGGCCGCCTTGGACGACTACGTTTCTCTATCGGGTCAGAGCAGCGAGCTGAAATGCCTCCCG** 152298
WF-Teratorn CAGTTTCAGAGAGCGCTCGCCGACGCCGACCCGGGTAGACTCCTGCCACTGGTCGTGAGGGCCGCCTTGGACGACTACGTTTCTCTATCGGGTCAGAGCAGCGAGCTGAAATGCCTCCCG 153513

TeratornLC199500 **GCCACAGTAAGGGCCGCGAGCCAGTCCGAGCCTCGCTCGGAGAAAATGGAAACGCGCTCCGATGCCGCTCCCGAGGCAAAAGGGGGATGCCCTCTCCGGCGGGCTTTTACGAGGTCCACA** 152418
WF-Teratorn GCCACAGTAAGGGCCGCGAGCCAGTCCGAGCCTCGCTCGGAGAAAATGGAAACGCGCTCCGATGCCGCTCCCGAGGCAAAAGGAGGATGCCCTCTCCGGCGGGCTTTTACGAGGTCCACA 153633

TeratornLC199500 **GAGGCAGGGTCGAGGAGAAGAGAATCCGCGTGA**GCGACGAGTCGCCCTCGTCGCAGCGATCCAGACAAATCCGCGTGGGATGACGACATGCTACCTTTCGTCCGATGTCTTCGCATCTCA 152538
WF-Teratorn GAGGCAGGGTCGAGGAGAAGAGAATCCGCGTGAGCGACGAGTCGCCCTCGTCGCAGCGATCCAGACAAATCCGCGTGGGATGACGACATGCTACCTTTCGTCCGATGTCTTCGCATCTCA 153753

TeratornLC199500 CGCTGAACATTGTACCTCGTGGCTGTTGTTTCTCTGCTGTGTTGTACCTACGCTACCTTCAATAAAATCTCCAACGAAAATGTGTCTGATGTGTTGGTTTGCATGCCTTGCCATCTTTTC 152658
WF-Teratorn CGCTGAACATTGTACCTCGAGGCTGTTGTTTCTCTGCTGTGTTGTACCTACGCTACCTTCAATAAAATCTCCAACGAAAATGTGTCTGATGTGTTGGTTTGCATGCCTTGCCATCTTTTC 153873

TeratornLC199500 TACCCCTTCATCCCCCCTTAAGAAAAGAACGGAAAAAAAAACAAAACAGCGCTTGTTCAAATTCAGGTTTGGTGTATCGGCTTTATTTTCCCAACCATGTCACCCCAATGATACAAAGAA 152778
WF-Teratorn TACCCCTTCATCCCCCCTTAAGAAAAGAACGGAAAAAAAAACAAAACAGCGCTTGTTCAAATTCAGGTTTGGTGTATCGGCTTTATTTTCCCAACCATGTCACCCCAATGATACAAAGAA 153993

TeratornLC199500 ACGATGAAACGAAAGACAACGAAACAAAAAACAAAACAAAACAACAAACATAGAACCAGCAACTCGTACACGTTCCACAGCAGCACAAAGACACAGGAGGGCC**TTAGACATCCCCATATC** 152898
WF-Teratorn ACGATGAAACGAAAGACAACGAAACAAAAAACAAAACAAAACAACAAACATAGAACCAGCAACTCGCACACGTTCCACAGCAGCACAAAGACACAGGAGGGCCTTAGACATCCCCATATC 154113

TeratornLC199500 **TCTTCTCTTTCAGTTTCCTCCCTTTCTTTCGCTTGGAGCACAGAGGTCTCTTCCATTCCACCCCTTTGCGCCTCGCGCGTTTCGCCTCTCTCGTCTCCACGTGGCCGGACGCGTCCTGCT** 153018
WF-Teratorn TCTTCTCTTTCAGTTTCCTCCCTTTCTTTCGCTTGGAGCACAGAGGTCTCTTCCATTCCACCCCTTTGCGCCTCGCGCGTTTCGCCTCTCTCGTCTCCACGTGGCCGGACGCGTCCTGCT 154233

TeratornLC199500 **CCCGTAGGGCCGCGCTCTCCAGAGGCAGCTCTAAGAATCGCAGAGCACGATCCGATTTGAGCTTCTGCAGCGCCTCCCTCTTCCACTGGAGGTTGACCAGGGCGGTCAACGGCGCGTGTC** 153138
WF-Teratorn CCCGTAGGGCCGCGCTCTCCAGAGGCAGCTCTAAGAATCGCAGAGCACGATCCGATTTGAGCTTCTGCAGCGCCTCCCTCTTCCACTGGAGGTTGACCAGGGCGGTCAACGGCGCGTGTC 154353

TeratornLC199500 **CGAATCGCACCAAAGCGTCGTACGAGCTTTCTGGGTTGGCCCTCGCCAGCCTATCGGCGAAAAATCTCACAAACGCGATCTCGCATCGCAAGCAGTTCCGCGATGCGCTCGAAGGGCACG** 153258
WF-Teratorn CGAATCGCACCAAAGCGTCGTACGAGCTTTCTGGGTTGGCCCTCGCCAGCCTATCGGCGAAAAATCTCACAAACGCGATCTCGCATCGCAAGCAGTTCCGCGATGCGCTCGAAGGGCACG 154473

TeratornLC199500 **TGGGCTGAGGCCACGGGTTGGGCCGAGTCCGATCCTCTTTGTCCTCTCGCAGACAGCCTGAATCCAACGTGGTCGTCATCAAACACGTAGATCCCGTCCCGAGACAGTTCCTCGGGCGAG** 153378
WF-Teratorn TGGGCTGAGGCCACGGGTTGGGCCGAGTCCGATCCTCTTTGTCCTCTCGCAGACAGCCTGAATCCAACGTGGTCGTCATCAAACACGTAGATCCCGTCCCGAGACAGTTCCTCGGGCGAG 154593

TeratornLC199500 **TCTCCCATGAGCACGTATCCATGAGGAAGCTTGCGCAGTGTGCAGTACGATGTGATGTGAGCGAACAGCCAGACCAGAGAAGCGATCTTTCTGTTATAGGGCGTGGGTCTTGTCGTGTGT** 153498
WF-Teratorn TCTCCCATGAGCACGTATCCATGAGGAAGCTTGCGCAGTGTGCAGTACGATGTGATGTGAGCGAACAGCCAGACCAGAGAAGCGATTTTTCTGTTATAGGGCGTGGGTCTTGTCGTGTGT 154713

TeratornLC199500 **TTTAGCTGTCTGTCCACTCCCTCGCTCAAAGTCGCTCCGATCAGCGGAACCTTCGCTACCCGAGACACGCTCCTCACCAGGCTCCACGAGGTCTCCACCGACAGCCCCGACGCCACCAGG** 153618
WF-Teratorn TTTAGCTGTCTGTCCACTCCCTCGCTCAAAGTCGCTCCGATCAGCGGAACCTTCGCTACCCGAGACACGCTCCTCACCAGGCTCCACGAGGTCTCCACCGACAGCCCCGACGCCACCAGG 154833

TeratornLC199500 **TGCTTGAACAGAACGATCCTCTCGGCCGTCCCGTTGTACCTGTCCGTGTATCTGCGGACAGGGTCCAGCGAGGCCATGCACACGTATGAGAGCAGTTTGTCATTGGTACCCGTTGAGGCA** 153738
WF-Teratorn TGCTTGAACAGAACGATCCTCTCGGCCGTCCCGTTGTACCTGTCCGTGTATCTGCGGACAGGGTCCAGCGAGGCCATGCACACGTATGAGAGCAGTTTGTCATTGGTACCCGTTGAGGCA 154953

TeratornLC199500 **ACGGAGGCCAAAGAGAGCTTGAGCTGCTCCACCTCGAGGCCCGCTCGCGTGCGATAGTCCTCCCTCGCCGGGTCGGGGCAACGCGTCGCCAGGAGCCTGGCCCTTTCTCGGCTCCTGTTC** 153858
WF-Teratorn ACGGAGGCCAAAGAGAGCTTGAGCTGCTCCACCTCGAGGCCCGCTCGCGTGCGATAGTCCTCCCTCGCCGGGTCGGGGCAACACGTCGCCAGGAGCCTGGCCCTTTCTCGGCTCCTGTTC 155073

TeratornLC199500 **AGCGAACCGTAAAAGCGGTGCAGCCAGCGCACGGGATGGGTGCTCAGGTACCTCGGGGCTTCCAGCTCCTCCAGCGTGCGCACGTACGATCGGAGCTGATGAACCACTTCTTTCCCTAGG** 153978
WF-Teratorn AGCGAACCGTAAAAGCGGTGCAGCCAGCGCACGGGATGGGTGCTCAGGTACCTCGGGGCTTCCAGCTCCTCCAGCGTGCGCACGTACGATCGGAGCTGATGAACCACTTCTTTCCCTAGG 155193

TeratornLC199500 **GCTATCTGTTTGTCCATCTCGTCCCGCCGCTTGATCGACTCCAGCTGCTCGAGGAACCTCACGGACCTCGGGCCACCACCCGTACCTTCCGCGGGCCGCTCCGTCCCGGGAAACGCGCCT** 154098
WF-Teratorn GCTATCTGTTTGTCCATCTCGTCCCGCCGCTTGATCGACTCCAGCTGCTCGAGGAACCTCACGGACCTCGGGCCACCACCCGTACCTTCCGCGGGCCGCTCCGTCCCGGGAAACGCGCCT 155313

TeratornLC199500 **CCGCTCGGCCTGGCCCCGGCCACCTCCCCGAAGCCTTCGCTGTTGCCCAAGTCCAGCGCGGCCGACGCGGCAGAGTCCGGGTCGACGGGGGGCTCCGCGCCCTCGTGCCTCAGCGTGAAG** 154218
WF-Teratorn CCGCTCGGCCTGGCCCCGGCCACCTCCCCGAAGCCTTCGCTGTTGCCCAAGTCCAGCGCGGCCGACGCGGCAGAGTCCGGGTCGACGGGGGGCTCCGCGCCCTCGTGCCTCAGCGTGAAG 155433

TeratornLC199500 **TGGCAGCAGGAAAACATTCTCCTGGTTTCTTCCCAAGTGTCCAGGGTAATGTCGGTTTCGAAATAGCACTCAGGGCACTCCTTGGTGTCTGCGACCAGCCTCGGAATCATTGGGTAGCGC** 154338
WF-Teratorn TGGCAGCAGGAAAACATTCTCCTGGTTTCTTCCCAAGTGTCCAGGGTAATGTCGGTTTCGAAATAGCACTCAGGGCACTCCTTGGTGTCTGCGACCAGCCTCGGAATCATTGGGTAGCGC 155553

TeratornLC199500 **AGCACCAACGCGAAAAAGTCCCTGAGAGAGTCCAGCGCGAGCAACGACTCTTCGGTCGAGATTGTGCCGAACTCCCTGCTCAGCCTCGCGTACACCTCCAGGGCTTCGGAGAACGTGAGG** 154458
WF-Teratorn AGCACCAACGCGAAAAAGTCCCTGAGAGAGTCCAGCGCGAGCAACGACTCTTCGGTCGAGATTGTGCCGAACTCCCTGCTCAGCCTCGCGTACACCTCCAGGGCTTCGGAGAACGTGAGG 155673

TeratornLC199500 **GGTTGGCCCGGCCGCCTGGCGATGGCTTCTCCTTCTTCTATGCCTCGCACCGCGAGCCACAGGGTGAACTCGTTCACTCGCTCTCCGCCCGGGTCCGCTGGGGTCGAGGCGGCCGTGCGG** 154578
WF-Teratorn GGTTGGCCCGGCCGCCTGGCGATGGCTTCTCCTTCTTCTATGCCTCGCACCGCGAGCCACAGGGTGAACTCGTTCACTCGCTCTCCGCCCGGGTCCGCTGGGGTCGAGGCGGCCGTGCGG 155793

TeratornLC199500 **GGCTCGGAGAAAGTCCTCGCGAATTCTAGCAGCGTTGCGTCCGCGTGGCCCAGCAGGCGCGCCGCACAGGCTCTGTCGTACAGCTCGGCCACGGACCTCCGAGCCGTCACGGCTCTCGCT** 154698
WF-Teratorn GGCTCGGAGAAAGTCCTCGCGAATTCTAGCAGCGTTGCGTCCGCGTGGCCCAGCAGGCGCGCCGCACAGGCTCTGTCGTACAGCTCGGCCACGGACCTCCGAGCCGTCACGGCTCTCGCT 155913

TeratornLC199500 **TCATCTCTGGAGCCATCAGGGCTTCCAGGGCGCCCGGAACCAGCGCGTTCGCGACGAGCCGGTGACCTTGCCACTCGGCCAGACAAACCATGAGTCCTATCCATCGCGGGTCCCCCGAAC** 154818
WF-Teratorn TCATCTCTGGAGCCATCAGGGCTTCCAGGGCACCCGGAACCAGCGCGTTCGCGACGAGCCGGTGACCTTGCCACTCGGCCAGACAAACCATGAGTCCTATCCATCGCGGGTCCCCCGAAC 156033

TeratornLC199500 **ACGACGGGTGCTTCACGTCTGACCTGAGCCCCGCCTGAGGCCAGAGGGTTGTTAGTTGAGCGAACGTGGCCGCTCCGCTCTCTGAGAGGAACCGCCTGGGTTCGCCGTAGACGGGTCGCG** 154938
WF-Teratorn ACGACGGGTGCTTCACGTCTGACCTGAGCCCCGCCTGAGGCCAGAGGGTTGTTAGTTGAGCGAACGTGGCCGCTCCGCTCTCTGAGAGGAACCGCCTGGGTTCGCCGTAGACGGGTCGCG 156153

TeratornLC199500 **CCCCGTAGCTCGACGAGGACGGCGATGCGGCCAACTCGCGGTCCAATCTCTTGGGCAGTTCTTCGAACTCGTAGGCCCCTGCCTTCGTCACGATCACGCAACGGGAGAGCTCTCGGGCGC** 155058
WF-Teratorn CCCCGTAGCTCGACGAGGACGGCGATGCGGCCAACTCGCGGTCCAATCTCTCGGGCAGTTCTTCGAACTCGTAGGCCCCTGCCTTCGTCACGATCACGCAACGGGAGAGCTCTCGGGCGC 156273

TeratornLC199500 **GCTTCAGCATCTCCCCTCTGCCTCCGCGCACCACCAGCGTGACACCGAACCGTCCCGGGGGCGCGTCCGTCGATAGCACGTTGCCCAGCGCCAGAGCGCAAAGGACCAGGTCCAAGATCT** 155178
WF-Teratorn GCTTCAGCATCTCCCCTCTGCCTCCGCGCACCACCAGCGTGACACTGAACCGTCCCGGGGGCGCGTCCGTCGATAGCACGTTGCCCATTGC-AGAGCGCAAAGGACCAGGTCCAAGATCT 156392

TeratornLC199500 **TATCCCCCGCCGGGTCTGCCGATCCGTGCTCAAACACTGGCCTGAGGCCGCAGCCGAACGTCAAAACGGTGTGAACGTGATACGTCAGGGGTGAAAACGAACAGGGTCTGTAATAGCGTA** 155298
WF-Teratorn TATCCCCCGCCGGGTCTGCCGATCCGTGCTCAAACACTGGCCTGAGGCCGCAGCCGAACGTCAAAACGGTGTGAACGTGATACGTCAGGGGTGAAAACGAACAGGGTCTGTAATAGCGTA 156512

```
TeratornLC199500  CGAGCTTGCCTCTTTCCGTCAGATCGAAATGCCTTTTCCAGGCCGGGTTGGGAGCGGGTGCGAAAAAAACGTGGCCCTCCACGGCAGGGACGTCTGGAACCAATTCGATTTCCAGTCGAG  155418
WF-Teratorn       CGAGCTTGCCTCTTTCCGTCAGATCGAAATGCCTTTTCCAGGCCGGGTTGGGAGCGGGTGCGAAAAAAACGTGGCCCTCCACGGCAGGGACGTCTGGAACCAATTCGATTTCCAGTCGAG  156632

TeratornLC199500  AGTTCAACGTGAGGATACTCGAAAGGCCTCTCTCTCGAAAGTCGCACTTTACTCTCAGGGGCGCGGGCGTTGGTACCGCAGGGTCTCCTTCTTCTCCTTCATCCACAGCTACCCCTTCTA  155538
WF-Teratorn       AGTTCAACGTGAGGATACTCGAAAGGCCTCTCTCTCGAAAGTCGCACTTTACTCTCAGGGGCGCGGGCGTTGGTACCGCAGGGTCTCCTTCTTCTCCTTCATCCACAGCTACCCCTTCTA  156752

TeratornLC199500  GATCTTGGTCTTCTTCTTCTTCTTCTTC---ATCCTCCTCCTCTTCTACCCGTTCGTCTTCATCGTTTTCAGCTCTCTCGGTCTGCACCTGTGCATTTCCCGCCTCTGCATTGAGCGTCA  155655
WF-Teratorn       GATCTTGGTCTTCTTCTTCTTCTTCTTCTTCATCCTCCTCCTCTTCTACCCGTTCGTCTTCATCGTTTTCAGCTCTCTCGGTCTGCACCTGTGCATTTCCCGCCTCTGCATTGAGCGTCA  156872

TeratornLC199500  GGGAGGCAGCTCGGCGCCTCATCTCGACGAGCGACGCCTTTGTGTCCACCGAATGCCAAAACCTGTCTCGCTTGAGCACGAGTCGACCGTCTGGGCTCCTCTGCCATCTCGGAGGGCTCG  155775
WF-Teratorn       GGGAGGCAGCTCGGCGCCTCATCTCGACGAGCGACGCCTTTGTGTCCACCGAATGCCAAAACCTGTCTCGCTTGAGCACGAGTCGACCGTCTGGGCTCCTCTGCCATCTCGGAGGGCTCG  156992

TeratornLC199500  AACGTAGACGGGGAAGCCATTCGCAAGGTCCTCGAGCTGGCGCTTCAAGCGGCGCTTCGGTTCCCGACTCCCACCTGAACTTCAGCTCGGCCACCTTTGAGCAAAACGATCCTGGCTGTC  155895
WF-Teratorn       AACGTAGACGGGGAAGCCATTCGCAAGGTCCTCGAGCTGGCGCTTCAAGCGGCGCTTCGGTTCCCGACTCCCACCTGAACTTCAGCTCGGCCACCTTTGAGCAAAACGATCCTGGCTGTC  157112

TeratornLC199500  TCTTTGACCACGGCGAGGGGTCCTTCCAGGTCTCGGGGCCTCGTATCCCCCCAGGCACTCCGTTCATTTCGCAGAAGGCCATGAGCGAATCTTTGTTGCGCATTCGCAGAGTCTCCTGCT  156015
WF-Teratorn       TCTTTGACCACGGCGAGGGGTCCTTCCAGGTCTCGGGGCCTCGTATCCCCCCAGGCACTCCGTTCATTTCGCAGAAGGCCATGAGCGAATCTTTGTTGCGCATTCGCAGAGTCTCCTGCT  157232

TeratornLC199500  GCTCCTCCGATGGTTCGGCGTCCTCGGGTGCGCTGACGAGCACGTGCAGGCCGGGATCGCCCCGTATCGCTGCGGCCCTTAGGGTCCTGGGCTCGGAATGCAGCACCAAGTTGCCAGAGT  156135
WF-Teratorn       GCTCCTCCGATGGTTCGGCGTCCTCGGGTGCGCTGACGAGCACGTGCAGGCCGGGATCGCCCCGTATCGCTGCGGCCCTTAGGGTCCTGGGCTCGGAATGCAGCACCAAGTTGCCAGAGT  157352

TeratornLC199500  GTCTCACCGGTATGACCCCTTTGGACCTTTCGACCAGAAGCGCGATAGCATCGTCCTCCCGCGGGCCGTCCGGACGTGACACCAGGTACCCCGTCCTTCTGCTCCTCAGCACGTCCAAGT  156255
WF-Teratorn       GTCTCACCGGTATGACCCCTTTGGACCTTTCGACCAGAAGCGCGATAGCATCGTCCTCCCGCGGGCCGTCCGGACGTGACACCAGGTACCCCGTCCTTCTGCTCCTCAGCACGTCCAAGT  157472

TeratornLC199500  CCCTCGGCAGCGCGTGGGTCCAGCGACAGGCGAAAGTCTGCGAGTCTGCCTCCGGGGTGTACACAGTTACGGCGCGACTCCTGAAGGGGCCCGTGGCTCCCGCGAACCAAGCCACCGCGC  156375
WF-Teratorn       CCCTCGGCAGCGCGTGGGTCCAGCGACAGGCGAAAGTCTGCGAGTCTGCCTCCGGGGTGTACACAGTTACGGCGCGACTCCTGAAGGGGCCCGTGGCTCCCGCGAACCAAGCCACCGCGC  157592

TeratornLC199500  GCTGCAGGTAACCGTACAGACCGCGTCTCGAGCATCTGATCTCCAGCTCTGCGTGAGTGTTCCCCGAGAGCATGGCTCTCGCGAGCGAGTCGGCGAACTCTCCGGCCGAGCTCACCTCGG  156495
WF-Teratorn       GCTGCAGGTAACCGTACAGACCGCGTCTCGAGCATCTGATCTCCAGCTCTGCGTGAGTGTTCCCCGAGAGCATGGCTCTCGCGAGCGAGTCGGCGAACTCTCCGGCCGAGCTCACCTCGG  157712

TeratornLC199500  CCCGAAAGCTCCAGCTCTCGAGCTCCTCTTCGCCGTCGGCCTCCGGCTCGCGGCCGTCGTCCTCGGGAACCGGCTTCGCGTACCACCTTCCGGGCTCCGCTATTTCGATCAGGCTGGAGA  156615
WF-Teratorn       CCCGAAAGCTCCAGCTCTCGAGCTCCTCTTCGCCGTCGGCCTCCGGCTCGCGGCCGTCGTCCTCGGGAACCGGCTTCGCGTACCACCTTCCGGGCTCCGCTATTTCGATCAGGCTGGAGA  157832

TeratornLC199500  TGGGCATGCGAGATTCGCGCTGAAACCGACTGGTGCCTCTATTCGTGACGACGCAGAGGCCTCTGGTCCACCTTCTTTCCGAGGGGCACGTCAAAACGCGCGATCGCAGCACGTCGAACC  156735
WF-Teratorn       TGGGCATGCGAGATTCGCGCTGAAACCGACTGGTGCCTCTATTCGTGACGACGCAGAGGCCTCTGGTCCACCTTCTTTCCGAGGGGCACGTCAAAACGCGCGATCGCAGCACGTCGAACC  157952

TeratornLC199500  CTCTCCACTCTCTTGGGTTCAGGGCCCTGGCGTGCCCGTCTGCCATCTCCACACGCCCGAGGCCCTCGCCGGGGCGGCCGTGGTGCGTGAACTCCTCCGGCGGAGCCCCGCAGAGCCATC  156855
WF-Teratorn       CTCTCCACTCTCTTGGGTTCAGGGCCCTGGCGTGCCCGTCTGCCATCTCCACACGCCCGAGGCCCTCGCCGGGGCGGCCGTGGTGCGTGAACTCCTCCGGCGGAGCCCCGCAGAGCCATC  158072

TeratornLC199500  TGACGATCTCGACCCCTGCGCCCTCCGGGCCCGCGGACCTGGAGCTCCGGGACACCGGGTAGTCTCGCTCCCAGTCGCTCGGGCGCGAGGTCTCGGCCGGGACTTTGGCAAAAGTCCACA  156975
WF-Teratorn       TGACGATCTCGACCCCTGCGCCCTCCGGGCCCGCGGACCTGGAGCTCCGGGACACCGGGTAGTCTCGCTCCCAGTCGCTCGGGCGCGAGGTCTCGGCCGGGACTTTGGCAAAAGTCCACA  158192

TeratornLC199500  GAGCGTCCTTCAGTCTTTCCCAGGAGCTCTCGCTCGTGGCGTAGCCCTCCAACGGATCCCTGCGCCCCCGACGCATGAACATCTTGTCGATGGCGGACGCTTTGTCCAGGCACTTGATGC  157095
WF-Teratorn       GAGCGTCCTTCAGTCTTTCCCAGGAGCTCTCGCTCGTGGCGTAGCCCTCCAACGGATCCCTGCGCCCCCGACGCATGAACATCTTGTCGATGGCGGACGCTTTGTCCAGGCACTTGATGC  158312

TeratornLC199500  GTGCCCTGGCCTCGAGCGGCTCGGGAGCAGGCCACGCGGGGTCTTCGTTCGTCAGCGCGTCGGTCCGGAACAGCGAACAGTTCTCCGCGACCACGGCCACGGACTCCAGCCTCTGAACCA  157215
WF-Teratorn       GTGCCCTGGCCTCGAGCGGCTCGGGAGCAGGCCACGCGGGGTCTTCGTTCGTCAGCGCGTCGGTCCGGAACAGCGAACAGTTCTCCGCGACCACGGCCACGGACTCCAGCCTCTGAACCA  158432

TeratornLC199500  ATGGGCCTTTGCCCCTCAGGTAGAACAGGGGAGACTCAGAAGCGAACGCCACCGTGTGCCCGTCTATCACGGACATGCCGGGGCGTCTGGCGGACGCCGTCGACTCGGGTCTCGAGGCCC  157335
WF-Teratorn       ATGGGCCTTTGCCCCTCAGGTAGAACAGGGGAGACTCAGAAGCGAACGCCACCGTGTGCCCGTCTATCACGGACATGCCGGGGCGTCTAGCGGACGCCGTCGACTCGGGTCTCGAGGCCC  158552

TeratornLC199500  CTTCGCTCGCCAGCCTAAGAAAGCTCCAGTCCCCCAGGCTCGTCGAAACTCTGCCCGGGAAGGTCTCGACCCTCTCGAACACCAGCGCGGGTCCCAGCGTCTCAGAGTACGCCCTCAGGA  157455
WF-Teratorn       CTTCGCTCGCCAGCCTAAGAAAGCTCCAGTCCCCCAGGCTCGTCGAAACTCTGCCCGGGAAGGTCTCGACCCTCTCGAACACCAGCGCGGGTCCCAGCGTCTCAGAGTACGCCCTCAGGA  158672

TeratornLC199500  GGAGGTCTTCCTCATCCCCCCGAGGCAGCAGCTCGCTCCCGGGGCAGAGCTCGGAAGCGCTCAGGGGTCTGGGCTCGCAGCACCTGGCCAGGAGCGGAGCCAGCCTGTGGTCCCGGGAGC  157575
WF-Teratorn       GGAGGTCTTCCTCATCCCCCCGAGGCAGCAGCTCGCTCCCGGGGCAGAGCTCGGAAGCGCTCAGGGGTCTGGGCTCGCAGCACCTGGCCAGGAGCGGAGCCAGCCTGTGGTCCCGGGAGC  158792
```

```
TeratornLC199500  AGAGGAAGGCTCCGATTGTGTTGACCCTGAGGCTGCGCGAGTGGCGGACCTCTGCGTCCGTTCCCCTGGTCCAGGCAGACGCACCCCTGTGCCTGCTCAGCGTGAACAGCTTGCAGGCCG  157695
WF-Teratorn       AGAGGAAGGCTCCGATTGTGTTGACCCTGAGGCTGCGCGAGTGGCGGACCTCTGCGTCCGTTCCCCTGGTCCAGGCAGACGCACCCCTGTGCCTGCTCAGCGTGAACAGCTTGCAGGCCG  158912

TeratornLC199500  GGTGGCAGGTCAGTCTGCAGCTCTCGGCCAGGTGGCCCAGGAGAGCGTGCGCCGGGGGCAGAAGGCCCTGTCCGTGCAGGACGAACTCGTAGTCGAGCATCACGTCACCCCGGCTCATCT  157815
WF-Teratorn       GGTGGCAGGTCAGTCTGCAGCTCTCGGCCAGGTGGCCCAGGAGAGCGTGCGCCGGGGGCAGAAGGCCCCGTCCGTGCAGGACGAACTCGTAGTCGAGCATCACGTCACCCCGGCTCATCT  159032

TeratornLC199500  CTGGCACGCAAGGGTGCACGGCTCTGTAGAAGCGAGTCAGGCAGGCGCTCACGTCAAAGTCCTTCTCTGCCCCCCTCGTCGCCAGGACTCTCGCGCAGTGGCCCAGCATCACAGCCTCAT  157935
WF-Teratorn       CTGGCACGCAAGGGTGCACGGCTCTGTAGAAGCGAGTCAGGCAGGCGCTCACGTCAAAGTCCTTCTCTGCCCCCCTCGTCGCCAGGACTCTCGCGCAGTGGCCCAGCATCACAGCTTCAT  159152

TeratornLC199500  CCAGGCAGGCTTCGTACGCAGAGCCGCACTCTCCGGGGCACCTGTCGGCCCTGTCCAGCGCGAAGCGCTTAGGGCGCGCGTTCGCCGCCAGGTTCAGGTTGTATTCACGCAGGGGATGGG  158055
WF-Teratorn       CCAGGCAGGCTTCGTACGCAGAGCCGCACTCTCCGGGGCACCTGTCGGCCCTGTCCAGAGCGAAGCGCTTAGGGCGCGCGTTCGCCGCCAGGTTCAGGTTGTATTCACGCAGGGGATGGG  159272

TeratornLC199500  AGAGGGCGTCCTCCGAGACGCTCCCGCTCGACTCCTTCTCGGCCCTCGCCACCTCGACCGTGGGGAAGAACTCTCCGTCCCCGATGGCGCTGTGGCACTGTCGGTCCGCCGCCGGCGACC  158175
WF-Teratorn       AGAGGGCGTCCTCCGAGACGCTCCCGCTCGACTCCTTCTCGGCCCTCGCCACCTCGACCGTGGGGAAGAACTCTCCGTCCCCGATGGCGCTGCGGCACTGTCGGTCCGCCGCCGGCGACC  159392

TeratornLC199500  TGAGCCACGAACGCGTGTGTGGGGGCTCTCCTCCACACGGGCCTCGAGCGAAAGGCTCGTCTACGGCCAGCTCCAGCTCGCAACGCCCGGCCGTCCGGCGGGAAGATCTGAAAATGCGCG  158295
WF-Teratorn       TGAGCCACGAACGCGTGTGCGGGGGCTCTCCTCCACATGGGCCTCGAGCGAAAGGCTCGTCTAAGGCCAGCTCCAGCTCGCAACGCCCGGCCGTCCGGCGGGAAGATCTGAAAATGCGCG  159512

TeratornLC199500  ACAGCCTGCGCTCCTCGTAACCCCTGGCCTGATCGCTGCGCACGGCCCGTTCCAGGCTGGACGCGGCGTCTGCTGTGAGCGGGAAGCGCTCCAGTTCGTGGCCGGAGCGGCTCCAGATAC  158415
WF-Teratorn       ACAGCCTGCGCTCCTCGTAACCAATGGCCTGATCGCTGCGCACGGCCCGTTCCAGGCTGGACGCGGTGTCTGCTGTGAGCGGGAAGCGCTCCAGTTCGTGGCCGGAGCGGCTCCAGATAC  159632

TeratornLC199500  GTTCCCTCCCTGTCTCCGGGGCGTAATTCAGCTGAAAGTAGCAGTCGGGCGGCGGGATAGTCGAGGAGGCGGCGATCTCGCGCAAGTCCTCGGTGAGCCTCGTGCAACGGGCGCCACCTG  158535
WF-Teratorn       GTTCCCTCCCTGTCTCCGGGGCGTAATTCAGCTGAAAGTAGCAGTCGGGCGGCGGGATAGTCGAGGAGGCGGCGATCTCGCGCAAGTCCTCGGTGAGCCTCGTGCAACGGGCGCCACCTG  159752

TeratornLC199500  ATGCGCGGGAGCCCGGGCGCACGCCCTGCGTGTCGGGAATCTTCAGGAGGGCCCTGTGACGGCCGCGTCTCGAGTGAAACACCCCGCAGACCCTCAGTATGTTCAAGTCGGCGGGCCCCA  158655
WF-Teratorn       ATGCGCGGGAGCCCGGGCGCACGCCCTGCGTGTCGGGAATCTTCAGGAGGGCCCTGTGACGGCCGCGTCTCGAGTGAAACACCCCGCAGACCCTCAGTATGTTCAAGTCGGCGGGCCCCA  159872

TeratornLC199500  GCGTGCGTATGCCTATCGTGACCCACTTGAAACGAGAGCTCGGCCCGAGCCCCCTCCCGAAAAGGACGATGTGGCCCGTGGTCCTCTCGCAGATCGACGGAGAATCCACGCGGGCCGGCG  158775
WF-Teratorn       GCGTGCGTATGCCTATCGTGACCCACTTGAAACGAGAGCTCGGCCCGAGCCCCCTCCCGAAAAGGACGATGTGGCCCGTGGTCCTCTCGCAGATCGACGGAGAATCCACGCGGGCCGGCG  159992

TeratornLC199500  GATCTGCCTCTTTGATGCCCCTGGCGTCCGAGAGCGGCGCCCTGCTCGGGGTGCCCCGCAGCAGAGCCAGGTCCCCATCCAAATCCTCGTGCATCGGGCGCTCGGGCGACTCGGACGGCC  158895
WF-Teratorn       GACCTGCCTCTTTGATGCCCCTGGCGTCCGAGAGCGGCGCCCTGCTCGGGGTGCCCCGCAGCAGAGCCAGGTCCCCCTCCAAATCCTCGTGCATCGGGCGCTCGGGCGACTCGGACGGCC  160112

TeratornLC199500  CAAGACTCGCTAGGCGAGGACAGGTGCCAGGTACGTGCTCTGACACGATCTGCCTGAGCTTCGACAGAATGGAGCAGTCGCTCGCCTCGAAGCGAATGCTGCTTTCCCTGGAACCTCGTT  159015
WF-Teratorn       CAAGACTCGCTAGGCGAGGACAGGTGCCAGGTACGTGCTCTGACACGATCTGCCTGAGCTTCGACAGAATGGAGCAGTCGCTCGCCTCGAAGCGAATGCTGCTTTCCCTGGAACCTCGTT  160232

TeratornLC199500  CGAAGCCCGCGTTTTCCATGCGTAGGTAAGAAAGGACCAGTGCCCGCGCCTCTGTCCACTCCAAGGGCCCCGAGAACCATAGGCTCGTCGGTACGTCTGGGCCCGCCAGCTTCGCGAAGA  159135
WF-Teratorn       CGAAGCCCGCGTTTTCCATGCGTAGGTAAGAAAGGACCAGTGCGCGCGCCTCTGTCCACTCCAAGGGCCCCGAGAACCATAGGCTCGTCGGTACGTCTGGGCCCGCCAGCCTCGCGAAGA  160352

TeratornLC199500  CTTCCAGCGCGTACAGCACTGCCGCGCAGCTCGCGCCGTCGAGCTGAGGAGGAGCGATCACGTCCATGCCCCCGGGGATCAGACACACGGCCCGATCGAAGAGGTACGCCTCTGACCTGA  159255
WF-Teratorn       CTTCCAGCGCGTACAGCACTGCCGCGCAGCTCGCGCCGTCGAGCTGAGGAGGAGCGATCACGTCCATGCCCCCGGGGATCAGACACACGGCCCGATCGAAGAGGTACGCCTCTGACCTGA  160472

TeratornLC199500  AGCCCTGCGGCATGACACCGCTGTACCGAGGGGTCCTGTAGACACGCAACGTGGGCGTCCTGCTTGCAGACGGTGATCTAGAGTTCCCTCGGACGACCTTCGGCGAGGATATCAGCAGGC  159375
WF-Teratorn       AGCCCTGCGGCATGACACCGCTGTACCGAGGGGTCCTGTAGACACGCAACGTGGGCGTCCTGCTTGCAGACGGTGATCTAGAGTTCCCTCGGACGACCTTCGGCGAGGATATCAGCAGGC  160592

TeratornLC199500  GGGGCGCCCCTGGCGCGCCTGTCTCGATCTCCAGCACAGAGTGCCCGCTGGGAAACCATTCGGTGAGCCGTCCGCTCGGGCCTTCCGCAGCTGTCCTCGTCCTCAATCTGCTCTCGCTTT  159495
WF-Teratorn       GGGGCGCCCCTGGCGCGCCTGTCTCGATCTCCAGCACAGAGTGCCCGCTGGGAAACCATTCGGTGAGCCGTCCGCTCGGGCTTTCCGCAGCTGTCCTCGTCCTCAATCTGCTCTCGCTTT  160712

TeratornLC199500  TGCTTTCGGCCTCCCGGGTTTCCCGTGCTGTCGTCTTCCCGCCTCTGCTCACACCTGCAATGTCCTGGCGGCCCGTGCCCCTTCTGGCGTTCGGACTCGACTTGGGCACGATCACCAGAG  159615
WF-Teratorn       TGCTTTCGGCCTCCCGGGTTTCCCGTGCTGTCGTCTTCCCGCCTCTGCTCACACCTGCAGTGTCCTGGTGGCCCGTGCCCCTTCTGGCGTTCGGACTCGACTTGGGCACGATCACCAGAG  160832

TeratornLC199500  CATCCCTGAGAAGGTGCCTCAGCACGGCGACGTGGTCGTTCTCGGCGAGCACCGAGAACGCCAGTGGCGTGGAGATCACCTCCCTCAGAGCCGCAAGGCCGGAGAATGCGCCGTGGCGAC  159735
WF-Teratorn       CATCCCTGAGAAGGTGCCTCAGCACGGCGACGTGGTCGTTCTCGGCGAGCACCGAGAACGCCAGTGGCGTGGAGATCACCTCCCTCAGAGCCGCAAGGCCGGAGAATGCGCCGTGGCGAC  160952

TeratornLC199500  TTTCCGCCGTTCCGTCCAGTTCCCGTAGAAATCCCTGCCTCCCGCTGGCGCCGGGGCCTAGCGCGGGCCCCGGGAACCAAGGATCTTTCGGCTCCAGTTCCCTCAGGTCGAACGCGCGGT  159855
WF-Teratorn       TTTCCGCCGTTCCGTCCAGTTCCCGTAGAAATCCCTGCCTCCCGCTGGCGCCGGGGCCTAGCGCGGGCCCCGGGAACCAAGGATCTTTAGGCTCCAGTTCCCTCAGGTCGAACGCGCGGT  161072
```

```
TeratornLC199500  CGCCTTCTCCGGGTAGCACGCACCGCAGGGCGAACCGCGTCCCCAGCCCGCAGTCGCCCAAGGCAAGGGCGCAGAGCGTGCGGACAACCTCCTCGCGGCAGTCGGGCGAGAAGACGAGGC  159975
WF-Teratorn       CGCTTTCTCCGGGTAGCACGCACCGCAGGGCGAACCGCGTCCCCAGCCCGCAGTCGCCCAAGGCAAGGGCGCAGAGCGTGCGGACAACCTCCTCGCGGCAGTCGGGCGAGAAGACGAGGC  161192

TeratornLC199500  AAACGCGAGCCTTCCAAGAAGACACCACCTGGTAGAGCGACAGGCCATCTGAAGCGTAGATGATACCCGGGACGACCTCGGCTCCGCTCACGTCCCTCTCCTCGACCTCGTAGCCCACGC  160095
WF-Teratorn       AAACGCGAGCCTTCCAAGAAGACACCACCTGGTAGAGCGACAGGCCATCTGAAGCGTAGATGATACCCGGGACGACCTCGGCTCCGCTCACGTCCCTCTCCTCGACCTCGTAGCCCACGC  161312

TeratornLC199500  TCGTTTCTGGCCACCAGTGGGAAGAGCCGCCCGATTCGATCTCTGAGCGCTTCGAGATCACCAAAGCGGGCTCGGCCCCACCGCCTGCGAGCGCCCTGAAGGCCACGTCCGAGAACACAG  160215
WF-Teratorn       TCGTTTCTGGCCACCAGTGGGAAGAGCCGCCCGATTCGATCTCTGAGCGCTTCGAGATCACCAAAGCGGGCTCGGCCCCACCGCCTGCGAGCGCCCTGAAGGCCACGTCCGAGAACACAG  161432

TeratornLC199500  ACAGGGTCTCCCTGAACACGGCGAACGCGCTGTCTGCGGGCCGCAGCCTCCTGGGAGGACGGGGCTCCGCGAATCGCTGGATGCTCAGCAGTCTTCCATCCTCGTCGGCGGTGATCAGCG  160335
WF-Teratorn       ACAGGGTCTCCCTGAACACGGCGAATGCGCTGTCTGCGGGCCGCAGCCTCCTGGGAGGACGGGGCTCCGCGAATCGCTGGATGCTCAGCAGTCTTCCATCCTCGTCGGCGGTGATCAGCG  161552

TeratornLC199500  CACCGTCACCCAGCCTCGCAGCGTAGAGCGACGAAGCGTTCGAGATGCTCAGGGGGCTAGGCCGCAGGCGCCGCCGACGTTCCCGCTCGTACGCGTCGCGGCACCTCGACCGGTAGCCCT  160455
WF-Teratorn       CACCGTCACCCAGCCGCGTCGCATAAAGCATCGA------CGAGGAGCTCAGGGGGCTCTGGCGCAGGCGCCGCCGACGTTCCTGCTCGTATGCGTGGCGGCACCTCAAACGGTAGCCTT  161666

TeratornLC199500  CGAGCCACTGCGGATGGAGGCGGGCTCGCAGGTTTCGGCCCCGCGTCCCAAACAGGTAGGCCAGCTCGTCTTCCGAGAGTTCTGCCTTGCCGGCGCGGAGGCATGCGTACACGTAGCGGA  160575
WF-Teratorn       CGAGCCACTGCGGATGAAGGCGGGCTCGCAGGTTTCGATCCCGTGTCCCGAACAGGTAGGCCAGCTCGTCGTCCGAGAGTTCCGCTTTGCCGGCGCAGAGGCACGCGTAGATGTAGCAGA  161786

TeratornLC199500  TGTACTTATCACACATGCGGACGGGGGGCCCCGAACGGCCTCCTCCCGAAATTTTCGGAGGCGAACGGGCTTCGAGGGAGCTGCTCTCGAACACTTCTACCCCATGGACACGGGCGCGGC  160695
WF-Teratorn       TGTACTTATCACACATGCGGACGATTGGCCCCGAACGGCCCCCTCCCAAACGTTTCGGAGGCAAACGGGCTTCGAGGGAGCTGCTCCTGAAGGCTTCTACCCCATGGACATGACCGCGAC  161906

TeratornLC199500  CCCATCTTTCTGCTCCGCGTGGAGCCGGGCCGCCGGACATGCTGGCGGATCGCAGCGCACGACGCTGTTCCAGGCCGTGGAGTCTGCGTTTCGGAGGTATGAGGAAAGGGCTGCCGCTTG  160815
WF-Teratorn       ------TTTCTGCTCCGCGTGGAGCCGGGCCGCCGGACATGCCAGCGGATCGCAGCAGGCAACGCTGGTCCAGGCTGTGGAGTCTGCGTTTCGGAGGTATGAGGAAAGGGCT------CG  162014

TeratornLC199500  GAGGAAGA---TCGAACGAACGGAGCGTGAGACTAGCGGAGGGGAGCGCAGAGAGAGCAGAACCGGATGGTCCGCACCGGAGGAGGTCGAAATGAACACAGAGGACGAGTCGTTCGATCT  160932
WF-Teratorn       GAGGGAGAGCGTGGAACGCAGGGAGCGTGAGACGAGCGGAGGGGAGCGCGGCGAGCGCAGAGCCGGAAGGTTCCCACAAGACGACGTAGAAATGTACACAGAAGGCGAGTCGCTCGATCT  162134

TeratornLC199500  GGATTCTCCGTCCGAGAGGTCCTTAGTGAACTTGAGGCGCTGCGAAAGCGAAGAGAGCATGTCTGTGCTCTCTCAAGGAAGCGAAGGCGAACCCGAGGGCGAACCCGGGGACGAACGCGA  161052
WF-Teratorn       GGACTCTCCATCCGAGAGGTCCTTAGTGAACCTGAGGCGCAGCGAAAGCGAAGAGAGCATGTCTGTGCACTCTCAAGGCAGCGAAGGCGAACCTGAGGGCGAACCCGCGGACAAACGCGA  162254

TeratornLC199500  ATACAGCATTTCCTGGGTACCCGTGAGACGGGAAAGACGCTGGCGCGAAGACTCTCTGAGCGACGGACGCCTCGGTCCGGGAGATCCTCCGTGCGGGCCCGCCCGGTGCCTCAAGTGCGA  161172
WF-Teratorn       GTACAGCATTTCCTGGGTACCCGTGAGACGGGAAAGACGCCGGCGCGAAGACTCTCTGAGCGACGGAGGCCTCGGTCCGGGAGATCCACCACAAGGGCCTGCCCGGTGCCTCAAGTGCGA  162374

TeratornLC199500  AGCCTACGACGCCCACTGCAAGTCCGCTGCTGAGAGCTACAGAAGGACCTTGGGCTGCTACACGAAAACGGTGTCCGAGCTCATGTTGGGCTTCAGGACCGTGCTGGAGAGGTATCCCGA  161292
WF-Teratorn       AGCCTACGACGCCCACTGCAAGTCCGCTGCCGAGAGCTACAGACGGACCTTGGGCTGCTACACGAAAACGGTGTCCGAGCTCATGTTGGGCTTTAGAACCGTGCTGGAGAGGTATCCCGA  162494

TeratornLC199500  GCTCAGGGTCGACACGGGCGACGAACGCCTGCACTTCATGGTTCAGGCCGCGCTGGGCCGTAACAACCTGGTCCGAGAAATGGTCCGCCTCTCTGCGAGGAGGACTCACGAGCGCGCGCT  161412
WF-Teratorn       GCTGAGGGTCGACACGGGCGACGAACGCCTGCACTTCATGGTTCAAGCTGCTCTGGGCCGTAACAACCTGGTCCGAGAAATGGTCCGCCTTTCCGCGAGGAGGACTCACGATCGCACGCG  162614

TeratornLC199500  CGGGGAATCCAGGCGGGGAGAGTCCTGTCGAGAGTCCTGTCGAGAGTCTGCGGCGGCGGCGACCGTGTGCGAAGCGCTGAACTCCTGCTACACGGACGGCGAGGCGCTGCGGATGATCCT  161532
WF-Teratorn       TGGGGAGTCCGGGCAGGGAGAGTCCTGTCGAGAGTCT------GCGTC---GGCGGCGGCGACCGTGTGCGAAGCGCTGAACTCCTGCTACACGGACGGCGAGGCGCTGCGGATGATCCT  162725

TeratornLC199500  GGAGCGCTACACCGACCTGTACGTGAGGTACATGGAGGTGTGGAGGGGGGATGTGCGCCAAGATTTCCAGGGATGTCACGCCACTGGAGTTTGTGGAAATCGTGCGGGACAGGCCCGGGGT  161652
WF-Teratorn       GGAGCGCTACACCGACCTGTACGTGAGATACATGGAGGTGTGGAGGGGGGATGTGCGCAAAGATTTCCAGGGATGTTACGCCGCTGGAATTTGTGGAAATCGTGCGGGACAGACCCGGGGT  162845

TeratornLC199500  GAGCCACGGTGAGCCGTGGAGCGAGGCGAGAGCGTATGTGGGTCGCTTGCTCGTCAAGGCATCTTCGCTGGTGCCCAGGCGCGTCGGGGACGAGTGCGCCCGGACGCTCAGGATGCTGGA  161772
WF-Teratorn       GAGCCACGGCGAGCCGTGGAGCGAGGCAAGAGCGTACGTGGGTCGCTTGCTTGTCAAGGCGTCTTTGCTGGTGCCCAGGCGCGTTGGGGACGATTGCGCCCGGACGCTCAGGATGCTGGT  162965

TeratornLC199500  CCTCGAACTGTCCAGAGCCAGACCTCAGAGCTCAGAGGGTCGGGACAGGGTCCTCATAGGGCACGTGGAGAACGTGCGCGAGCGTCTCGACGAGGCGGGCTCCGGCATGTGGAGCTGGGA  161892
WF-Teratorn       CGTCGAACTTTCCCGAGCCAGACCTCGAAGCTCAGAAGGTCGGGACAGGGTCCTCGTAGGGCACGTGGAAGACGTGCGCGATCTTCTCGACGAGGCAGGCTCCGGCATGTGGAGCTGGGA  163085

TeratornLC199500  TTTTCACTCGAACGATGCCTACTTTGACATCAGGAGCGACCTGGCGGTTCTCGAGAGCCTAATCGGCATCGGCGACATCTCCGTCAAGAGCCTGCTTTACGAGAGATCCCTGCTGATCTC  162012
WF-Teratorn       TTTTCACTTGAACGACGCCTACTTTGACATCAGGAGCGACCTGGCGGTTCTCGAGAGCCTAATCGGAATCGGCGACATCTCCGTCAAGAGCCTGCTTTACGAGAGATCCCTACTGATCTC  163205

TeratornLC199500  TCAGTTGAGAAAGGTCGTGAGGAACGAAGGTTCTCTGGAGCTCAGCCTGAGCAGGGCACGCGGCGAGCTCGCGGAGGCCCGAGCGTCGATACGCCAGCTCGAGGCCGCCAGCTCCTACTC  162132
WF-Teratorn       TCAGTTGAGAAAGGTCGTGAGGAACGAAGGCTCTCTGGAGCTCAGCCTGAGCAAGGCACGCGGTGAGCTCACGGAGGCCCAAGCATCGAGGCGCCGGCTTGAGGCCGCCGGCTCCTTCTT  163325
```

| | | |
|---|---|---|
| TeratornLC199500 | **CGCGTTCTGA**AGGTGCGCGTACGGGGGTGGGACGGGACGCGGTAAGATACTTAACGCGGAGGGCGACGGGCCGAACTCG-CCATCCCCCTCACC**ATGGACGTCGACTGCGATTACTTGAC** | 162251 |
| WF-Teratorn | CGCGTTCTGAAGGTGTGCCTATGGGGGTGGGACGGGACACGGTAAGATACTTAACGCGGCCGGCAGCGGGCCCAACTCGACCACCCCCCTTACCATGGACGTCGACTGCGATTACTTGAC | 163445 |
| TeratornLC199500 | **GATCGGCGGTGTCGACGCTGAACCTGACGGTCTGATTGACTCTTGTCCGCATTGCTCCTCCGCGCGCTCCCTGCGCAGGGCATTGGTGTACCAGCCAGCGCACCAGGAATCTGACGGGGT** | 162371 |
| WF-Teratorn | GGTCGGCGGTGTCGACGCTGAACCTGACGGCCGGAATGACCCCTGTCCGCATTGCTCCTCCGCGCGCTCCCTGCGCAGGGCCCTGGTGTACAAGCCAGTGCACCAGGAATCTGATGGGGT | 163565 |
| TeratornLC199500 | **CCCGGCTTCCATTTCGGTCAGGCCGGTCACTTTCGAGGAGGCTTCTTCCGGCAGCATGGTCCCCTACGTGCTTTCGCATGAGACAAGGGACGGTCGCGGCATCGACCCGGAGGACGGGGA** | 162491 |
| WF-Teratorn | CTCGGCATCCATTTCGGTCAGGCCGGTCACCTTCGAGGAGGCTTCGTCCGGCCGAATGGTCCCCTACGTGCTGTCGCACGAGACAAGGTACGGTCGCGGCATCGACCCGGAGGACGGGGA | 163685 |
| TeratornLC199500 | **CAAGACCGCGCATTACTACACGGTGCGGAACTGCCTCGCCTCGATCCGTGGCAGCGTCCACCTGAGAGACGCGGCGGAATACGGAACCTGGCTGTCAGAGGTGGACCAGTGTGCGGCGGC** | 162611 |
| WF-Teratorn | CGAGACCGCGCATTACTACACGGTGCGGAACTGCCTCGCCTCGATCCGTGGCAGCGTCCACCTGAGAGACGCGGCGGAATACGGAACCTGGCTGTCAGAGGTGGACCAGTGTGCGGCGGC | 163805 |
| TeratornLC199500 | **ACTCGGCCGGTTGCGAAGCGCAGCTCACGTCTGCTCTCACCTGCCCGACCTGAAACGATTTGTGCTCGAGCAGAAAAATGACGATGGCCTGCTGCACCTCGCCAGCAAGTGGCTCAGGCG** | 162731 |
| WF-Teratorn | ACTCGGCCGGTTGCGAAGCGCAGCTCACGTCTGCTCTCACCTGCCCGACCTGAAACGATTTGTGCTCGAGCAGAAAAATGACGATGGCCTGCTGCACCTCGCCAGCAAGTGGCTCAGGCG | 163925 |
| TeratornLC199500 | **GGGCGAACACGCAACCGTCGGGATCGAGATGTTCGGGCCCTTTTTCGTCGATGTGGCTTTCGCGCCGGTTTTTCACAAGATCTTGAGATCTGACTGGCTCGTTATGCGCAAACGGAACTG** | 162851 |
| WF-Teratorn | GGGCGAACACGCAACCGTCGGGATCGAGATGTTCGGGCCCTTTTTCGTCGATGTGGCTTTCGCGCCGGTTCTTCACAAGATCTTGAGATCTGACTGGCTCGTTATGCGCAAACGGAACTG | 164045 |
| TeratornLC199500 | **TGGCACCTGCAACGAACTGAAACCGGGCTGGGCTCAGGAGTTCTACGACGCTCGCGAGGTCCCGCGCTCGCCAGACAGTGAGATGAGGTTCGTGGGCCACCCGACTTGTCCTTGTTGCAA** | 162971 |
| WF-Teratorn | TGGCACCTGCAACGAACTGAAACCGGGCTGGGCTCAGGAGTTCTACGACGCTCGCGAGGTCCCGCGCTCGCCAGACAGTGAGATGAGGTTCGTGGGCCACCCGACTTGTCCTTGTTGCAA | 164165 |
| TeratornLC199500 | **GCCGGGGGAGGCGAAGGACCTTGGCGTCGTGGCTCGGGCCATTGTGAGGGAGATGAGGCTCTGCGGATACACCCCTTGGTCCGCTGTGCAGGCGGTGCGGCGCTGCGAAAGCTCCATGCT** | 163091 |
| WF-Teratorn | GCCGGGGGAGGCGAAGGACCTTGGCGTCGTGGCTCGGGCCATTGTGAGGGAGATGAGGCTCTGCGGATACACCCCTTGGTCCGCTGTGCAGGCGGTGCGGCGCTGCGAAAGCTCCATGCT | 164285 |
| TeratornLC199500 | **GGAGCTGTCGTGCTCCGAGTGCCGTCGGGTGATTTTTACCCATTGCGCATTTCTCATGGAGTGCGCTGCGGTCGAGGCTTCTCGGGAACGCGACTGGCTGGTCCGCAACGAGGCCGTGAT** | 163211 |
| WF-Teratorn | GGAGCTGTCGTGCTCCGAGTGCCGTCGGGTGATTTTTACCCATTGCGCATTTCTCATGGAGTGCGCTGCGGTCGAGGCTTCTCGGGAACGCGACTGGCTGGTCCGCAACGAGGCCGTGAT | 164405 |
| TeratornLC199500 | **TCAACTGTGCTGGGTCGTTCATTCCCTCCTTCCGCTCGAGAAGCAAGCCGCGTTCCCTGCGACGGACCTTTCGGGCGCCTGCTGCGTATCGGAAAACTTTGCAGAATCAGAAGATGGGGG** | 163331 |
| WF-Teratorn | TCAACTGTGCTGGGTCGTTCATTCCCTCCTTCCGCTCGAGAAGCAAGCCGCGTTCCCTGCGACGGACCTTTCGGGCGCCTGCTGCGTATCGGAAAACTTTGCAGAATCAGAAGATGGGGG | 164525 |
| TeratornLC199500 | **TGGAGGAGAAGGAGAAATCCCCAGACGAGGAGAAGAAGGAGGGAACGGGGAAGGAAAAGAGGATTTGCCGGCTTGCGAGACGCGCCAGCACGGAGGGGCTGCTGCGATGTACGCTTGGGC** | 163451 |
| WF-Teratorn | TGGAGGAGAAAGAGAAATCCCCAGACGAGGAGAAGAAGGAGGGAACGGGGAAGGAGAAGAGGATTTGCCGGCCTGCGAGACGCGCCAGCACGGAGGGGCTGCTGCGATGTACGCTTGGGC | 164645 |
| TeratornLC199500 | **TTTTTCGGTGGCCGCGCACTGCGTCGCGTTTCAAACCTACGAGCGAAACGGCTCGCTGGATTTGGAAAAGATCGTGGTGCTGTACGCAAACGATCATCTCCCCATGCTCCTCGACGATCA** | 163571 |
| WF-Teratorn | TTTTTCGGTGGCCGCGCACTGCGTCGCGTTTCAAACCTACGAGCGAAACGGCTCGCTGGATTTGGAAAAGATCGTGGTGCTGTACGCAAACGATCATCTCCCCATGCTCCTCGACGATCA | 164765 |
| TeratornLC199500 | **GGACGCCCGGGGACAAGGCATCCTGACGACGAGCGTGTCGCCGGACCCCAGACTGTTTGAGACGGCAGCGCTGAACACTGGCTTCGTCTCTCCTGGGCTCGACCCGAGGGATCGCCTCGA** | 163691 |
| WF-Teratorn | GGACGCCCGGGGAGAAGGCATCCTGACGACGAGCGTGTCGCCGGACCCCAGACTGTTTGAGACGGCAGCGCTGAACACTGGCTTCGTCTCTCCTGGGCTCGACCCGAGGGATCGCCTCGA | 164885 |
| TeratornLC199500 | **GGAGCGGGACATGCGGGTCCCCCCCGCCGGAGAAGTGCTGTACGAGCTCCCTGCACCGAGGACATTCCGTCGCTCGGGTTCCGGGACCCCGGTCGCCCTGGCGGAGATCCCCAGTCACAC** | 163811 |
| WF-Teratorn | GGAGCGGGACATGCGGGTCCCCCCCGCCGGAGAAGTGCTGTACGAGCTCCCTGCGCCGAGGACATTCCGTCGCTCGGGTTCCGGGACCCCGGTCGCCCCGGCGGAGATCCCCAGTCACAC | 165005 |
| TeratornLC199500 | **CGAGGCCCGGGCTCAGGATCCCGAAGCGATAGCGGCCGCCATAGCGCTTCACTTTCCCAAAGTGTCGCCGTGCGCTCTCGACCGTCTGAGCCTAAAAGTGCCTCGACCCAGCTCGCTCAG** | 163931 |
| WF-Teratorn | CGAGGCCCGGGCTCAGGATCCCGAAGCGATAGCGGCCGCCATAGCGCTTCACTTTCCCAAAGTGTCGCCGTGCGCTCTCGACCGTCTGAGCCTAAAAGTGCCTCGACCCAGCTCTCTCAG | 165125 |
| TeratornLC199500 | **GGCTGACGCCAGCTTGTTCTGGCCTGTGTTTGTCAACCCCACAAGGTCCTGCCACAAGCTGTTCAAGGCCCTGGCTCTGCCCGCGTCTCCGAGTCCCGCTTACGCGCTAGAAACGTTGGC** | 164051 |
| WF-Teratorn | GGCTGACGCCAGCTTGTTCTGGCCTGTGTTTGTCAACCCCACAAGGTCCTGCCACAAGCTGTTCAAGGCCCTGGCTCTGCCCGCGTCTCCGAGTCCCGCTTACGCGCTAGAAACGTTGGC | 165245 |
| TeratornLC199500 | **GCGAGGATGGCGACGGGTGAAGGACCTCGTGCGACCAGACTCGCCTCTGCCCACCGCCTCCGGAGGGTGGGTCGCTGTGGAGCGCGAGCATTACGAACGTTATTTCGAGGGCCCCCTGGA** | 164171 |
| WF-Teratorn | GCGAGGATGGCGACGGGTGAAGGACCTCGTGCGACCAGACTCGCCTCTGCCCACCGCCTCCGGAGGGTGGGTCGCTGTGGAGCGCGAGCATTACGAACGTTATTTCGAGGACCCCCTGGA | 165365 |
| TeratornLC199500 | **CCTCTTGACGCGCGCAGGGTTGACGAGTGAGAGCGAAAAGGAGGCGATGCAAGAGGCTTTGGCGAGAGGCGTTTTGCCGTGCTTCGCGGCCGAAGGGGAAAGCTTACCCGACGCGCTTGT** | 164291 |
| WF-Teratorn | CCTCTTGACGCGCGCAGGGTTGACGAGTGAGAGCGAAAAGGAGGCGATGCAAAAGGCTTTGGCGAGAGGCGTTTTGCCGTGCTTCGCGGCCGAAGGGGAAAGCTTACCCGACGCCCTTGT | 165485 |
| TeratornLC199500 | **GAATTCCACCCTGGCGCTATCGTTCGCGGCCAGCGGGTTGAGACTCTCTGCGCCTCGTCACCTTGGAGGAGTAGAGCGGGGTCGGCGAGGAGTAGAGGCGGACGGCCGGTCCGCGGTGAG** | 164411 |
| WF-Teratorn | GAATTCCACCCTGGCGCTATCGTTCGCGGCCAGCGGGTTGAGACTCTCTGCGCCTCGTCACCTTGGAGGAGTAGAGCGGGGTCGGCGAGGAGTAGAGGCGGACGGCCGGTCCGCGATGAG | 165605 |

| | | |
|---|---|---|
| TeratornLC199500 | **AGTCACCGTGGCGGGTCTGTTGTACTTTGCGGCGTGGTACGTGCTGGTGTACGGTAACGCTCTGCACTCTCGTCCCAGCGCTCTCGAGGTTCTGGCTCGCAACCCGGGAGGAACGGACGC** | 164531 |
| WF-Teratorn | AGTCACCGTGGCGGGTCTGTTGTACTTTGCGGCGTGGTACGTGCTGGTGTACGGTAACGCTCTGCACTCTCGTCCCAGCGCTCTCGAGGTTCTGGCTCGCAACCCGGGAGGAACGGACGC | 165725 |
| TeratornLC199500 | **AGACAGGCACTTGCGTGCGGCCATGGAACTGAAGATCGACCGACGCGCCACGACAGGAGACTGGGGAGAGGACGACGTCGGCCCAGACCCGTCTTGCCGCTACTCTGTCCACGGGTACAA** | 164651 |
| WF-Teratorn | AGACAGGCACTTGCGTGCGGCCATGGAACTGAAGATCGACCGACGCGCCACGACAGGAGACTGGGGAGAGGACGACGTCGGCCCAGACCCGTCTTGCCGCTACTCTGTCCGCGGGTACAA | 165845 |
| TeratornLC199500 | **CGACCGCAGCATCGTGAGAGCCATGCTGATCGTGTTTGAGGGATTGCTCGATGCTCGAACCGTGCTGCGCGCGGCACAGCTGTACGTGCCCGAGCTCACGGCTCAGCTGCCTTTCTTCAT** | 164771 |
| WF-Teratorn | CGACCGCAGCATCGTGAGAGCCATGCTGATCGTGTTTGAGGGATTGCTCGATGCTCGAACCGTGCTGCGCGCGGCACAGCTGTACGTGCCAGAGCTCACGGCTCAGCTGCCTTTCTTCAT | 165965 |
| TeratornLC199500 | **GGCCCTAAACGTCGTTCCCTGTACGGTGAAGAGCGCGATCGAGGCACGCGACTCGTGGTTCGCTGCCCGCCTAGGCAACGCGGAGTCGTCTCCGAACGACCCGCCTTACGCCTACCTGGG** | 164891 |
| WF-Teratorn | GGCCCTAAACGTCGTTCCCTGTACGGTGAAGAGCGCGATCGAGGCACGCGACTCGTGGTTCGCTGCCCGCCTAGGCAACGCGGAGTCGTCTCCGGACGACCCGCCTTACGCCTACCTGGG | 166085 |
| TeratornLC199500 | **CGACATTGCCCACGGGATTCGCAGCAGTATGAGAGGCGCCTTAGATCGCGGCGACGAGCGGCCCGTCGCGGCTCCTCCAGCGAGCCTCGACGCTCGAGCCTCAGACCCCTGGCACGGCTG** | 165011 |
| WF-Teratorn | CGACATTGCCCACGGGATTCGCAGCAGTATGAGAGGCGCCTTAGATCGCGGCGACGAGCGGCCCGTCGCGGCTCCTCCAGCGAGCCTCGACGCTCGAGCCTCAGACCCCTGGCACGGCTG | 166205 |
| TeratornLC199500 | **CCTCCTGTCTCTGAGCGAGAGCGACGACGACTGGGGTTCTATCCTCAGTTGCGCGACGCGCGACCTGGAGAGCATGTCTGTTCAGAGCGACCCAGATTGCTGGGAAAACGAGGTGGAACC** | 165131 |
| WF-Teratorn | CCTCCTGTCTCTGAGCGAGAGCGACGACGACTGGGGTTCTATCCTCAGTTGCGCGACGCGCGACCTGGAGAGCATGTCTGTTCAGAGCGACCCAGATTGTTGGGAAAACGAGGTGGAACC | 166325 |
| TeratornLC199500 | **AGGGAACCAGTACGTCGATTCGACCGCTCGCCATCCAGGCACTGCAGACGGGACGGAACGAGTAGCGACCTGCTCTGAGAGCGCCCCTCGAGACCCCCCGGTCGCCTCCGGAGATCAAGA** | 165251 |
| WF-Teratorn | AGGGAACCAGTACGTCGATTCGACCGCTCGCCATCCAGGCACTGCAGACGGGACGGAACGAGTAGCGACCTGCTCTGAGAGCGCCCCTCGAGACCCCCCGGTCGCCTCCGGAGATCAAGA | 166445 |
| TeratornLC199500 | **CGTTCCGAGCGCGGCACTCGCGCAGCGTGACTGCCCTCGGTCGACGCTGGAAACGCTGCTGGTGGTGGACTCTCCAGATAGTCTCGGCTTTGCGCACCAAAAGGTTTTTCCCAGCGAAAC** | 165371 |
| WF-Teratorn | CGTTCCGAGCGCGGCACTCGCGCAGCGTGACTGCCCTCGGTCGACGCTGGAAACGCTGCTGGTGGTGGACTCTCCAGATAGTCTCGGCTTTGCGCACCAAAAGGTTTTTCCCAGCGAAAC | 166565 |
| TeratornLC199500 | **CGCGGACGCGAACGAATGGCCGGGCTTTGTTGGGGTGAGCTCGGTTCCGAGTCTGAGAAAGCCTTGCCTGACGAACGTGTGGAATACGACTTTCGCCAAGACGCGCAGGGACGCCGTGCC** | 165491 |
| WF-Teratorn | CGCGGACGCGAACGAATGGCCGGGCTTTGTTGGGGTGAGCTCGGTTCCGAGTCTGAGAAAGCCTTGCCTGACGAACGTGTGGAATACGACTTTCGCCAAGACGCGCAGGGACGCCGTGCC | 166685 |
| TeratornLC199500 | **GAGCTCAGAACCTGCCGTATGCAGCTCGGCGTCGCGCCCGTGCAAAACCCTGGCGGAAGCGCTCTGGCTGATGTTCGTCGGCGCGTACGGACGCCGAGAGTCGGACTCGGCTGCCCTGAC** | 165611 |
| WF-Teratorn | GAGCTCAGAACCTGCCGTATGCAGCTCGGCGTCGCGCCCGTGCAAAACCCTGGCGGAAGCGCTCTGGCTGATGTTCGTCGGCGCGTACGGACGCCGAGAGTCGGACTCGGCTGCCCTGAC | 166805 |
| TeratornLC199500 | **GGTGGAAGCCCTCGCGGTTGTTGTACACTTGGCCTCGAAAATCATCGATAGAAATTCTACCTCAGGCCTAGGGCCCTGCAAAGACATAAAGGCCCTGTGCGGCGACTTGTACGCAGCGCC** | 165731 |
| WF-Teratorn | GGTGGAAGCCCTCGCGGTTGTTGTACACTTGGCCTCGAAAATCATCGATAGAAATTCTACCTCAGGCCTAGTGCCCTGCAAAGACATAAAGGCCCTGTGCGGCGACTTGTACGCAGCGCC | 166925 |
| TeratornLC199500 | **GGAACAGTTTCTGCCCGTCGTCTCCATAGCGGCAGGTTTGTTCGGGCGCAACGTCACGGTCTGCTCGCAGGGGACCTTCTTTCACATTTGCAACAGCCATCACGGGAACTCGTGGGTGCA** | 165851 |
| WF-Teratorn | GGAACAATTTCTGCCCGTCGTCTCCATAGCGGCAGGTTTGTTCGGGCGCAACGTCACGGTCTGCTCGCAGGGGACCTTCTTTCACATTTGCAACAGCCATCACGGGAACTCGTGGGTGCA | 167045 |
| TeratornLC199500 | **GGGCTTTCTGGAATGCGACGGATCATCGTGGCAGCTGAGGGGCCTGCCGCCGCACACGACTCCAATCGAACTTCTGTTCGAGAGTGGACCGAATCACAGAGCGGGCTCGCTCCGCGCCTA** | 165971 |
| WF-Teratorn | GGGCTTTCTGGAATGCGACGGATCATCGTGGCAGCTGAGGGGCCTGCCGCCGCACACGACTCCAATCGAACTTCTGTTCGAGAGTGGACCGAATCACAGAGCGGGCTCGCTCCGCGCCTA | 167165 |
| TeratornLC199500 | **CGCGAGGCTACACGACCTCGATGTGATCGAGGTTCCTTCGGACGGTTACTGCGGATTCCACAGCGTAGCCCAGCTCCTGTTCAATGACACGGCCAGAGGGGAAACCCGCGCGTCCCTGGT** | 166091 |
| WF-Teratorn | CGCGAGGCTACACGACCTCGATGTGATCGCGGTTCCTTCGGACGGTTACTGCGGATTCCACAGCGTAGCCCAGCTCCTGTTCAATGACACGGCCAGAGGGGAAACCCGCGCGTCCCTGGT | 167285 |
| TeratornLC199500 | **GACGAGGACGGCCGAGTTTGTGTTTCTCAACGCGGAAAGGGCAGACGTCAGAGAGACGTACCTGACAGTCAACGGACCCAAAAACGTCTCTGAGCTGAAGCGACTCTTGTCTGAGGAGAA** | 166211 |
| WF-Teratorn | GACGAGGACGGCCGAGTTTGTGTTTCTCAACGCGGAAAGGGCAGACGTCAGAGAGACGTACCTGACAGTCAACGGACCCAAAAACGTCTCTGAGCTGAAGCGACTCTTGTCTGAGGAGAA | 167405 |
| TeratornLC199500 | **GAGGTGGCTCAGTGTACAGGAGGTAAAATACATGCTAGAAGCTCACGGCAAACGCTTGGGACTCCTCACGGAGACGTCGTACCCGATCTACGATAGGGCTTGCGCTCACATCGTGTGGAC** | 166331 |
| WF-Teratorn | GAGGTGGCTCAGTGTACAGGAGGTAAAATACATGCTAGAAGCTCACGGCAAACGCTTGGGACTCCTCACGGAGACGTCGTACCCGATCTACGATAGGGCTTGCGCTCACATCGTGTGGAC | 167525 |
| TeratornLC199500 | **CCGAAACCATTACGAGCCCTTGAAACCCAAAGTGTTACGATGCTGA**GCTATGCTGTTTTCTGTCAGATGTTTTTTTTTT-ATTTTTCCTTGTTGTTTCGATGAGGTGATGGAAATAAAAA | 166450 |
| WF-Teratorn | CCGAAACCATTACGAGCCCTTGAAACCCAAAGTGTTACGATGCTGAGCTATGCTGTTTTCTGTCAGATGTTTTTTTTTTTATTTTTCCTTGTTGTTTCGATGAGGTGATGGAAATAAAAA | 167645 |
| TeratornLC199500 | TCAAAGACATACCAAATTCTGTGGTGCACGACTGAATTCTTTATTTTGTGAAAAAGGAATACACTGGAAATACTGTACATGAACACAAGTACATGAACACAACCGCAATGCAGTCAGAGG | 166570 |
| WF-Teratorn | TCAAAGACATACCAAATTCTGTGGTGCACGACTGAATTCTTTATTTTGTGAAAAAGGAATACACTGGAAATACTGTACATGAACACAAGTACATGAACACAACCGCAATGCAGTCAGAGG | 167765 |
| TeratornLC199500 | GTGAGTTTCTTTGCTTCATGCGAGATAAAGTAAACTGCAAAAATGGTACATGGCGACTGCTCTCGGTAGAAGCGACACGGGTCACTACACAGTCAGAAGGAACATTCGGTACACTGCCCC | 166690 |
| WF-Teratorn | GTGAGTTTCTTTGCTTCATGCGAGATAAAGTAAACTGCAAAAATGGTACATGGCGACTGCTCTCGGTAGAAGCGACACTGGTCACTACACAGTCAGAAGGAACATTCGGTACACTGCCCC | 167885 |

```
TeratornLC199500  GTTGCCGAGTGAGAGACGGTGCGTGCGCTACAAATTGAAGCAAATTGAAGTGGATGAAACCTAATAGTACATGCTGCTTTGAGTGTCCTCTTCCCGCTGAGGGAGCCCTAGAGAGCTTTC  166810
WF-Teratorn       GTTGCCGAGTGAGAGACGGTGCGTGCGCTACAAATTGAAGCAAATTGAAGTGGATGAAACCTAATAGTACATGCTGCTTTGAGTGTCCTCTTCCCGCTGAGGGAGCCCTAGAGAGCTTTC  168005

TeratornLC199500  CCCCGCCCCGTCACCGTGCGTGTGCTCAGACCCTCTCGAGGTCCACCCGTCTCCAGCCACTCCTTCCCCGTCGAGTTCTTCGTTCTGATTGGCAAAGGCGGCGGCGAGCGCTCTGGTGAC  166930
WF-Teratorn       CCCCGCCCCGTCACCGTGCGTGTGCTCAGACCCTCTCGAGGTCCACCCGTCTCCAGCCACTCCTTCCCCGTCGAGTTCTTCGTTCTGATTGGCAAAGGCGGCGGCGAGCGCTCTGGTGAC  168125

TeratornLC199500  CAGGTCGTACCTCTCCAACGGGTTGAGCTGAGCGAGGGTCACGGGCCCGGCGACAGCGGGCTCGACCACAATCACCCGATTCCACTTGATGACTTGATTTTGTATCCGTACGACAAACGG  167050
WF-Teratorn       CAGGTCGTACCTCTCCAACGGGTTGAGCTGAGCGAGGGTCACGGGCCCGGCGACAGCGGGCTCGACCACAATCACCCGATTCCACTTGATGACTTGATTTTGTATCCGTACGACAAACGG  168245

TeratornLC199500  GCCGTAGGGTTGCACCTCGAAGCCTTGTTCCGTGGGCAGAGAAAGCATTTCGTATCGTCCGTCTCTACCCAGCACGTCCTCCAGCGGGCTTCGCGGTCCCGTGGCTCTTTCGGAAAAGCC  167170
WF-Teratorn       GCCGTAGGGTTGCACCTCGAAGCCTTGTTCCGTGGGCAGAGAAAGCATTTCGTATCGTCCGTCTCTACCCAGCACGTCCTCCAGCGGGCTTCGCGGTCCCGTGGCTCTTTCGGAAAAGCC  168365

TeratornLC199500  GCACAGCCCGTACACTTGGTGGATGAGCTCCTCGTCCTCGGGCAAAGTCCACTCTCTGAAGTAGTTTCTCATATCCCCGGCCTGGAGCCACGACACGCAGCTGCGCTCCTTACTGCGGCC  167290
WF-Teratorn       GCACAGCCCGTACACTTGGTGGATGAGCTCCTCGTCCTCGGGCAAAGTCCACTCTCTGAAGTAGTTTCTCATATCCCCGGCCTGGAGCCACGACACGCAGCTGCGCTCCTTACTGCGGCC  168485

TeratornLC199500  GCCCTTTGAGCACATCGGAGAGCCGTAGGCTGAGTGCGGGCACGCTCCGAAAACGGCGATGTACGCCCGCTTCAGCTTGTCAGCTGCCGCGTCGGGCACGTATTGTCGCAGGTTTAGCTG  167410
WF-Teratorn       GCCCTTTGAGCACATCGGAGAGCCGTAAGCGGAGTGCGGGCACGCTCCGAAAACGGCGATGTACGCCCGCTTCAGCTTGTCAGCTGCCGCGTCGGGCACGTATTGTCGCAGGTTTAGCTG  168605

TeratornLC199500  GGAGAGGGTTTCCCACACGGCGGCCAGACCTTCCAAAGCACCCCATGCGGGCCCGTTCCCGTCCCAGACGTACAGCGTGTCGACAAGGCACGCCAGCAGGTCGACCCTAAATCCCGTCTT  167530
WF-Teratorn       GGAGAGGGTTTCCCACACGGCGGCCAGACCTTCCAAGGCGCCCCATGCGGGCCCGTTCCCGTCCCAGACGTACAGCGTGTCGACAAGGCACGCCAGCAGGTCGACCCTAAATCCCGTCTT  168725

TeratornLC199500  GACCCTCCAAATGCGAGCCGGCTGTATGCGCTTCATGGTGAAGCCGAGAACCCTCAGGGAAAGGATCTGAACTCTGAGGAGGCACTGTAGCACGTGGTAGGCCAGCAGTACCTTGTCGTC  167650
WF-Teratorn       GACCCTCCAAATGCGAGCCGGCTGTATGCGCTTCATGGTGAAGCCGAGAACCCTCAGGGAAAGGATCTGAACTCTGAGGAGGCACTGTAGCACGTGGTAGGCCAGCAGTACCTTGTCGTC  168845

TeratornLC199500  GTAGAACCACGACACGAGCGACCCCTTGGAGCACCCGAGGTTCACAGACCCGGGGCTCTCGCCCACAAACACCCCGAGCGCCATTTCCTCCGTGTCTTTGGTCACGCAGTGTAGGGCGAA  167770
WF-Teratorn       GTAGAACCACGACACGAGCGACCCCTTAGAGCACCCGAGGTTCACAGACCCGGGGCTCTCGCCCACAAACACCCCAAGCGCCATTTCCTCCGTGTCTTTGGTCACGCAGTGTAGGGCGAA  168965

TeratornLC199500  GGTGCGTGCAGCAAAGGAGTGGACTTGCAGGCGCGTGTGCATGTCCAGCAGATTCACCAGAGACTCGTAGGCCATGTGAATCCTGTGTATGATCAAGCCCTGGACGTCCTGGACGGTGCA  167890
WF-Teratorn       GGTGCGTGCAGCAAAGGAGTGGACTTGCAGGCGCGTGTGCATGTCCAGCAGATTCACCAGAGACTCGTAGGCCATGTGAATCCTGTGTATGATCAAGCCCTGGACGTCCTGGACGGTGCA  169085

TeratornLC199500  CACGGCACCCAGGTAGTTGGTCTTTTTCCCGAGCACTCGAAGGAGATGTGGCGTGTCCTCCTTGCACAAGCTCGGCAAACTGTCCACGAGAATCTCCGTGAGCCTGAAACCCCGGTCCGG  168010
WF-Teratorn       CACGGCACCCAGGTAGTTGGTCTTTTTCCCGAGCACTCGAAGGAGATGTGGCGTGTCCTCCTTGCACAAGCTCGGCAAACTGTCCACGAGAATCTCCGTGAGCCTGAAACCCCGGTCCGG  169205

TeratornLC199500  CGGAGCAGAGAGGTCCACGGTGAAGTACCACTGCTCCGAGAGCTCGCAGTTGAGCACGGTGCGCTCCACCGCCGGCACGTAGGCAAACTCTGTGGGCCATGAGAGCTTCGTGCACCGGGC  168130
WF-Teratorn       CGGAGCGGAGAGGTCCACGGTGAAGTACCACTGCTCCGAGAGCTCGCAGTTGAGCACGGTGCGCTCCACCGCCGGCACGTAGGCAAACCCTGTGGGCCATGAGAGCTTCGTGCACCGGGC  169325

TeratornLC199500  GTGCCCCCGTCTCGTCATATCCCCGTCCCTGTGCGAGTACTCTCTGCCCATCCTCCACAGAAACCTTTGGCAAGTGATTGATCCGTGTACCAGGTCCAATACAGCTGGCCCCCGCTGCAT  168250
WF-Teratorn       GTGCCCCCGTCTCGTCATATCCCCGTCCCTGTGCGAGTACTCTCTGCCCATCCTCCACAGAAACCTTTGGCAAGTGATTGATCCGTGTACCAGGTCCAATACAGCTGGCCCCCGCTGCAT  169445

TeratornLC199500  TTGCTTGTCCTTTGCGCCCGGGCCCCACGCATCCCATCGCTCCGGCTTGTTGAGTCCGTGGCACGGCCTGCTGACGCAGTCCTCCTCTGAGTGACGTTTCACGGTGAGGTTGCTGATCAA  168370
WF-Teratorn       TTGCTTGTCCTTTGCGCCCGGGCCCCACGCATCCCATCGCTCCGGCTTGTTGAGTCCGTGGCACGGCCTGCTGACGCAGTCCTCCTCTGAGTGACGTTTCACGGTGAGGTTGCTGATCAA  169565

TeratornLC199500  ATCGGTTATGTCCCTCGAAACCAGGTAAGTGACGGGGTGAAAGCGGGCGCAGTCTGACAGTTGCCTCTCCCGCCGCGAGAGCCCTCTGGACCTCCATCTCATGCAATTAGCCTTCTGCAC  168490
WF-Teratorn       ATCGGTTATGTCCCTCGAAACCAGGTAAGTGACGGGGTGAAAGCGGGCGCAGTCTGACAGTTGCCTCTCCCGCCGCGAGAGCCCTCTGGACCTCCATCTCATGCAATTAGCCTTCTGCAC  169685

TeratornLC199500  CCGCTCGCGTTTGGCCGAGCTGAGACTGTCCTCCTCGCCGCGACTGTCCTCCATTCGGATGGGGCCCTGAAGCGACTGCCTATCTGCGCTTCCCGAGGGGTTGAGTCCTCCCGAAGCGAC  168610
WF-Teratorn       CCGCTCGCGTTTGGCCGAGCTGAGACTGTCCTCCTCGCCGCGACTGTCCTCCATTCGGATGGGGCCCTGAAGCGACTGCCTATCTGCGCTTCCCGAGGGGTTGAGTCCTCCCGAAGCGAC  169805

TeratornLC199500  GAGCGCCTCTTCTCCGTCGCACCGGGTCGTCTCTGGCAAGGGGTCCTCCGATTTTAGGCGAAGAAGGGTTAGAGTCGTGTCCGCACTCGTGCTCTGCTCGGGGTAAAAGGACTCTTGTAA  168730
WF-Teratorn       GAGCGCCTCTTCTCCGTCGCACCGGGTCGTCTCTGGCAAGGGGTCCTCCGATTTTAGGCGAAGAAGGGTTAGAGTCGTGTCCGCACTCGTGCTCTGCTCGGGGTAAAAGGACTCTTGTAA  169925

TeratornLC199500  GACTCTGTACACCATGTCGCTGCTCTGGAAATTGTCGATGTCAGTTCTGGCGAAGAAGAGATTCGTAGTGTTGTCTGTCACAATGCAGTCCAGGGAGCCTGCCGTGTGGAGGGAGCCTGC  168850
WF-Teratorn       GACTCTGTACACCATGTCGCTGCTCTGGAAATTGTCGATGTCAGTTCTGGCGAAGAAGAGATTCGTAGTGTTGTCTGTCACAATGCAGTCCAGGGAGCCTGCCGTGTGGAGGGAGCCTGC  170045

TeratornLC199500  CGTGTGGAGGGAGCCGGTCGAGCGAAGGGAGCTCGCCTCGTCCTCGTCGCAAAGTCCGAAGTCCTGCAGCTCGGCGAGGGCGTCCCGCTTTTCGACCTTGGCACTGGGCGAACTGTGGCA  168970
WF-Teratorn       CGTGTGGAGGGAGCCGGTCGAGCGAAGGGAGCTCGCCTCGTCCTCGTCGCAAAGTCCGAAGTCCTGCAGCTCGGCGAGGGCGTCCCGCTTTTCGACCTTGGCACTGGGCGAACTGTGGCA  170165
```

TeratornLC199500 **GCCTCTGT---CACAGTCTGTCAATCTCTCCCATTTCGCAGAGAGGAGGCCCTCGTTCACGGGGTTCACGTCCGCGTCTAGGTAGGTCACGGTCCGCGGCTTGCTCGAAGGTTCGCTCGA** 169087
WF-Teratorn GCCTCTGGGTGCGCGGCGATCCAATCTCTCCCATTCCGCAGAGAAGAGGTCCTCGTTCAGGGGGTTCACGTCCGCGTCTAGGTAGGTCACGGTCCGCGGCTTGCTCGAAGGTTCGCTCGA 170285

TeratornLC199500 **CGACCCGCGAGAGAGAGCCAGTTCCCTTGTCAGATCGCTGCTCATCTTCATAATGTGCTCCAGGCCGCAGCATTCTTCCAGCGTCCGCTGGCTCGCGGACTGGGATGCCTCTGGAAAAAG** 169207
WF-Teratorn CGACCCGCGAGAGAGAGCCAGTTCCCTTGTCAGATCGCTGCTCATCTTCATAATGTGCTCCAGGCCGCAGCATTCTTCCAGCGTCCGCTGGCTCGCGGACTGGGATGCCTCTGGAAAAAG 170405

TeratornLC199500 **CGGCTGACAGGGATCCATCGTCATGCTTTTTGTGACGTGCGGATTGCTCAGCGTGTTACTCTTGCGCTTCCTCCTGCCGGTCTTGGTTCGGGCTGGACTGGTGGCACGGGCGCGTCCCGA** 169327
WF-Teratorn CGGCTGACAGGGATCCATCGTCATGCTTTTTGTGACGTGCGGATTGCTCAGCGTGTTACTCTTGCGCTTCCTCCTGCCGGTCTTGGTTCGGGCTGGACTGGTGGCACGGGCGCGTCCCGA 170525

TeratornLC199500 **AGCCTGGCAAGACAGAATTTCCGCCAGGGTCTTACAGCGCTCCAGTGTCCTCCTCGCTGGCCCCAGCGAGGACAACTCGTCCTCCTTCAGGACCTCTATCATGCAGGCGAGCCCGGGCGC** 169447
WF-Teratorn AGCCTGGCAAGACAGAATTTCCGCCAGGGTCTTACAGCGCTCCAGTGTCCTCCTCGCTGGCCCCAGCGAGGACAACTCGTCCTCCTTCAGGACCTCTATCATGCAGGCGAGCCCGGGCGC 170645

TeratornLC199500 **CGCTGCGGCCCCGTCGCACCGCGTCAACCAGTCGAACCGCTCTCTCAAAGAGCTGCGATCCCTGGACCCGGGCCCCAGCCACGTGGCCCGCGGCCAGCGCCCCGTCTCTTCGGACCGCGG** 169567
WF-Teratorn CGCTGCGGCCCCGTCGCACCGCGTCAACCAGTCGAACCGCTCTCTCAAAGAGCTGCGATCCCTGGACCCGGGCCCCAGCCACGTGGCCCGCGGCCAGCGCCCCGTCTCTTCGGACCGCGG 170765

TeratornLC199500 **CGGTCTCCCGAGGCAATCCAGCACCGTGGACCAGAGGTGAGCGTCCGCCATGCCCGTCCTGCAAAACTCAGAGATGGCGGCCTCCGTTGTCGAGCCATCGTCCGGGAGCGCTCCGGCGCG** 169687
WF-Teratorn CGGTCTCCCGAGGCAATCCAGCACCGTGGACCAGAGGTGAGCGTCCGCCATGCCCGTCCTGCAAAACTCAGAGATGGCGGCCTCCGTTGTCGAGCCATCGTCCGGGAGCGCTCCGGCGCG 170885

TeratornLC199500 **CAGCACGCTCAGCGCCAAGTTGGAGCACGCGGCACGAAAACTGTCTTCGTCGTCGGTCAGCGCAGAGGCCCTGTAAAGCCCGGGCATGGAAACTAAGGGGAGCAGCGACATTGTCTCCCC** 169807
WF-Teratorn CAGCACGCTCAGCGCCAAGTTGGAGCACTCGGCACGAAAACTGTCTTCGTCGTCGGTCAGCGCAGAGGCCCTGTAAAGCCCGGGCATGGAAACTAAGGGGAGCAGCGACATTGTCTCCCC 171005

TeratornLC199500 **TGTGTGTCGGTGGATGACGAAGGCGCATGAGAGCTTGGAGGTGCGGCTGTGAGAGTCGTCATCCTTTGCGTTCTGGTACAGAACTCTGCAAGGATGGCGATCGGTGGAGCGGAGGTCTCT** 169927
WF-Teratorn TGTGTGTCGGTGGATGACGAAGGCGCATGAGAGCTTGGAGGTGCGGCTGTGAGAGTCGTCATCCTTTGCGTTCTGGTACAGAACTCTGCAAGGATGGCGATCGGTGGAGCGGAGGTCTCT 171125

TeratornLC199500 **CAGGGAGTCCAACACGCAGACCAGCGCTCTGCGGACCACGACGTCGCGGACCTGCTTTCCTCGGGTCTCCCTCAGCCAGCTGTGGAGCGGCAGCCACCCGGCCCCCGACGTTGTCACCAC** 170047
WF-Teratorn CAGGGAGTCCAACACGCAGACCAGCGCTCTGCGGACCACGACGTCGCGGACCTGCTTTCCTCGGGTCTCCCTCAGCCAGCTGTGGAGCGGCAGCCACCCGGCCCCCGACGTTGTCACCAC 171245

TeratornLC199500 **GTGCATGACAGTCGACGGCGCTCCTCCCGCTTGCTCCAGCGCGATGCAATCGAGCTGCGGGTAGTGCTGCCCGACCCCCGGCGAAAAGACCTGTCCTTCGTCTTTCAGCCTGACCATCCG** 170167
WF-Teratorn GTGCATGACAGTCGACGGCGCTCCTCCCGCTTGCTCCAGCGCGATGCAATCGAGCTGCGGGTAGTGCTGCCCGACCCCCGGTGAAAAGACCTGTCCTTCGTCTTTCAGCCTGACCATCCG 171365

TeratornLC199500 **CCTCGCTCTGCATTGGTTCTCGTAAGCCCTCGCGGCTTCGTGCAAGGTTTGCTCCAGGTGTTTGTTGGAAACCAACACACTCAGGACGACGGAGACGTCCCCGTGGAGCGAGGTGCGCAC** 170287
WF-Teratorn CCTCGCTCTGCATTGGTTCTCGTAAGCCCTCGCGGCTTCGTGCAAGGTTTGCTCCAGGTGTTTGTTGGAAACCAACACACTCAGGACGACGGAGACGTCCCCGTGGAGCGAGGTGCGCAC 171485

TeratornLC199500 **CTCGCGCACCCTCACCAGCGGCTTACTCTGTGTCCTTTGGCTCGTCATTTCGGGTCCGGGCGCGGCCCTCAGCTCGTGAAACAGGCAATGGTTCCTCACAAAGCCGTAGAGGGGTTCGGG** 170407
WF-Teratorn CTCGCGCACCCTCACCAGCGGCTTACTCTGTGTCCTTTGGCTCGTCATTTCGGGTCCGGGTGCGGCCCTCAGCTCGTGAAACAGGCAATGGTTCCTCACGAAGCCGTAGAGGGGTTCGGG 171605

TeratornLC199500 **CATCCTCAGCTCGAAGCGGTCGGTCGACGGGGTCCACGACCAGCTCTCCAGCTCCGAGCCCAGGTTGAAGTAGTAGGGAATCGAAACGCCTAGCCTGGTGCTCTGCTTGTCCTGATATCT** 170527
WF-Teratorn CATCCTCAGCTCGAAGCGGTCGGTCGACGGGGTCCACGACCAGCTCTCCAGCTCCGAGCCCAGGTTGAAGTAGTAGGGAATCGAAACGCCTAGCCTGGTGCTCTGCTTGTCCTGATATCT 171725

TeratornLC199500 **CCACCAGAGCTTGAGGATGCTGGCATGAAGGTGAAGCTCCTCTTCTGGGTCGTGAGGCGAGCGCCGGGATGCGTCGGGACCTAGCGCTAGAAGTAGGTGCCTGGCGACGTCCCTCAACTC** 170647
WF-Teratorn CCACCAGAGCTTGAGGATGCTGGCATGAAGGTGAAGCTCCTCTTCTGGGTCGTGAGGCGAGCGCCGGGATGCGTCGGGACCTAGCGCTAGAAGTAGGTGCCTGGCGACGTCCCTCAACTC 171845

TeratornLC199500 **GCCGGGGGATCTGTAAAACGTGTAGCGCCTCCTTTTGGCCAGGCTGACGAAGAGCATTACGAAGTACGGAGTGTGGAGGGTGGCCAAGTGTTCCAACCGCCGCAGGTTCAGCCCCACTTC** 170767
WF-Teratorn GCCGGGGGATCTGTAAAACGTGTAGCGCCTCCTTTTGGCCAGGCTGACGAAGAGCATTACGAAGTACGGAGTGTGGAGGGTGGCCAAGTGTTCCAACCGCCGCAGGTTCAGCCCCACTTC 171965

TeratornLC199500 **TTGCGGTGCGCGGTTATCGCAGCGATCTTCCACCTCCAGCACGTGGCGACACAACGATGGTTCGAAACGTATTTTCACCCCTCTGCACAGCTGACAATATTTGCACTGGCCTGAGACGAA** 170887
WF-Teratorn TTGCGGTGCGCGGTTATCGCAGCGATCTTCCACCTCCAGCACGTGGCGACACAACGATGGTTCGAAACGTATTTTCACCCCTCTGCACAGCTGACAATATTTGCACTGGCCTGAGACGAA 172085

TeratornLC199500 **AAACGAACTTAGTCCGATCGAGCCCCTTAGGTCTTGCGGGTGGTAGAAGTAATCTCTGCGCTCTGCTGCGAGGCGGTCCCGCTCCCCCGGTGGAATCTCCCACGTAGCCCGATTGCAGAG** 171007
WF-Teratorn AAACGAACTTAGTCCGATCGAGCCCCTTAGGTCTTGCGGGTGGTAGAAGTAATCTCTGCGCTCTGCTGCGAGGCGGTCCCGCTCCCCCGGTGGAATCTCCCACGTAGCCCGATTGCAGAG 172205

TeratornLC199500 **GTAACAGATGGCCTCTTCGACGCAGTCGAAGCAGTCTCGGTCGACGTGGAGTTCCAGAGCACCTGCGTGCAGTGCTGCGGCTGCCACCAGCCACTTGGAGACGCCGTCCTTTGGATCCAT** 171127
WF-Teratorn GTAACAGATGGCCTCTTCGACGCAGTCGAAGCAGTCTCGGTCGACGTGGAGTTCCAGAGCACCTGCGTGCAGTGCTGCGGCTGCCACCAGCCACTTGGAGACGCCGTCCTTTGGATCCAT 172325

TeratornLC199500 GGGCTCAGCTGCGCAACTGACAAAAAAAACATAACACAGAGGAATGGTTGGGCAGGGGTGAGAGACAGAGGAAAGGACTCAGATGATGATGATGATGATGATAATGATTATGATTTGTCA 171247
WF-Teratorn GGGCTCAGCTGCGCAACTGACAAAAAAAACATAACACAGAGGAATGGTTGGGCAGGGGTGAGAGACAGAGGAAAGGACTCAGATGATGATGATGATGATGATAATGATTATGATTTGTCA 172445

TeratornLC199500 CAGACGCGGTTAGTTTTTTTTTTTCACTGCGTTTTATAACTTTGCCTCGTCCTTTTCACTGGATCGCTTCGCGCTTCGAAGAAAAGTCTGAAAAGTCTGTTAATATCCCTCACGTGACG 171367
WF-Teratorn CAGACGCGGTTAGTTTTTTTTTTTCACTGCGTTTTATAACTTTGCCTCGTCCTTTTCACTGGATCGCTTCGCGCTTCGAAGAAAAGTCTGAAAAGTCTGTTAATATCCCTCACGTGACG 172565

TeratornLC199500 CCCGCCATGTAAGCATCACGAGCGCCCCCCTGTCATCTTGACTCGGCG**TCACAGTTTGTAAATGTTCAGCGAGGTGTCGCCGTCCAACACTGTGTGCGTTACCATCAGTCTGTACTCTCT** 171487
WF-Teratorn CCCGCCATGTAAGCATCACGAGCGCCCCCCTGTCATCTTGACTCGGCGTCACAGTTTGTAAATGTTCAGCGAGGTGTCGCCGTCCAACACTGTGTGCGTTACCATCAGTCTGTACTCTCT 172685

TeratornLC199500 **GTCTCTCGCGTGGTAGTACAGGGTGTGTTGCACGGCTATGGCGAGGCAGCAAGCCCCGTCCGTCTTCCCGAGAGCCCAGGTCCTGAACTGGTCCCGAAAGCCTTGCGGCTCACACACACG** 171607
WF-Teratorn GTCTCTCGCGTGGTAGTACAGGGTGTGTTGCACGGCTATGGCGAGGCAGCAAGCCCCGTCCGTCTTCCCGAGAGCCCAGGTCCTGAACTGGTCCCGAAAGCCTTGCGGCTCACACACACG 172805

TeratornLC199500 **GCGAGACCTGGAGCCGCTGCTGAAAGGCGTGGGCTTCAGGGTGCCCAGGGATGCCGGCAGCTCGTACCAGAAAGCGGGAGATCCTGCCCGCGGGGAGCCCATCCTGCTCGAGGCCTCTGG** 171727
WF-Teratorn GCGAGACCTGGAGCCGCTGCTGAAAGGCGTGGGCTTCAGGGTGCCCAGGGATGCCGGCAGCTCGTACCAGAAAGCGGGAGATCCTGCCCGCGGGGAGCCCATCCTGCTCGAGGCCTCTGG 172925

TeratornLC199500 **GAGCTCACCGATGGCCGCCGCGGCGAGCCCGACATCTCCCTCGAAGTACGTGTGCGCGAAGCAGGAGGCTTCACCCTCGGGTGGCTCGGGCAGAGTCCCCGAAAAACCATCCACGTCTAC** 171847
WF-Teratorn GAGCTCACCGATGGCCGCCGCGGCGAGCCCGACATCTCCCTCGAAGTACGTGTGCGCGAAGCAGGAGGCTTCACCCTCGGGTGGCTCGGGCAGAGTCCCCGAAAAACCATCCACGTCTAC 173045

TeratornLC199500 **GATGATGTAGCGCACGCCGAGACTCTCGTTTTTGTAGTCTACGCTGCGGACGGCCGCGGGCAGCACGGTGACGTACCCGAAGAGCCAGGCCAGCACATCGGACTCGATCTTCACAGGGTC** 171967
WF-Teratorn GATGATGTAGCGCACGCCGAGACTCTCGTTTTTGTAGTCTACGCTGCGGACGGCCGCGGGCAGCGCGGTGACGTACCCGAAGAGCCAGGCCAGCACATCGGACTCGATCTTCACAGGGTC 173165

TeratornLC199500 **GGGCACGTGGTTGGACCAGACCACCGCCTGCATTCCCCTGATCTTCGGAACCGCCTTCATGCAAGCGCTCATCACGTCGAAGGAGTTCCAGCGGAGGTCTATCACGCAGGCTTGCTCCGT** 172087
WF-Teratorn GGGCACGTGGTTGGACCAGACCACCGCCTGCATTCCCCTGATCTTCGGAACCGCCTTCATGCAAGCGCTCATCACGTCGAAGGAGTTCCAGCGGAGGTCTATCACGCAGGCTTGCTCCGT 173285

TeratornLC199500 **TCTCATGCCGATCAGGTGGTGCTTGCAGGCGACGGCTTTCTCGCATGACTCCGGAGACACCAGGTCGAGCGCCAGGGCTTCTCCGAAAAGCCTGTTTCCGTATTCGGGCGTGGGTGCCGT** 172207
WF-Teratorn TCTCATGCCGATCAGGTGGTGCTTGCAGGCGACGGCTTTCTCGCATGACTCCGGAGACACCAGGTCGAGCGCCAGGGCTTCTCCGAAAAGCCTGTTTCCGTATTCGGGCGTGGGTGCCGT 173405

TeratornLC199500 **GATGGCGCGCCCTATCACGTACAGGGCGCTGAAGTTGTCGCGGCCCGGGAAGAGCTCCAGGGTTATCTCCTTGGGCAGCGCACTCTCGAAGCAATGCGGCTCTTCCTCCACCGAAGCCGG** 172327
WF-Teratorn GATGGCGCGCCCTATCACGTACAGGGCGCTGAAGTTGTCGCGGCCCGGGAAGAGCTCCAGGGTTATCTCCTTGGGCAGCGCACTCTCGAAGCAATGCGGCTCTTCCTCCACCGAAACCGG 173525

TeratornLC199500 **TCCCCGTCTCTGGACACGAAAATTAAACTGGTCCACCACACAGAGGGGTCTCTCTGCGGTGCATCCGAAGTGGGATGGGCGTGATTTGTACACGTACAGTCCCGGAGGCAGGGTCCTGGA** 172447
WF-Teratorn TCCCCGTCTCTGGACACGAAAATTAAACTGGTCCACCACACAGAGGGGTCTCTCTGCGGTGCATCCGAAGTGGGATGGGCGTGATTTGTACACGTACAGTCCCGGAGGCAGGGTCCTGGA 173645

TeratornLC199500 **ACGGTCTACCTGGTAGACCAACACAGAGTCGCAGCATTTGGGGTATCTCGCGCTCCGGTTGCACATCAGCCACCTGTACGCCGCTGCCACCGGCTCCATTTCGTACGCCCACAGGACTTC** 172567
WF-Teratorn ACGGTCTACCTGGTAGACCAACACAGAGTCGCAGCATTTGGGGTATCTCGCGCTCCGGTTGCACATCAGCCACCTGTACGCCGCTGCCACCGGCTCCATTTCGTACGCCCACAGGACTTG 173765

TeratornLC199500 **TGACTGGGCTCTGGCGATTTCGCGGGAGGAGCCCTGCCCGCCCGGTTTGGTGGCGCCCCGTCGGCGGCGTTCGGGGTTCGGAACATCTGCACCGTCCCCCCCTCTGTAGCGCTGTTCCGC** 172687
WF-Teratorn GGACTGGGCTCTGGCGATTTCGCGGGAGGAGCCCTGCCCGCCCGGTTTGGTGGCGCACCGTCGGCGGCGTTCGGGGTTCGGAACATCTGCGCCGTCCCCCCCTCTGTAGCGCTGTTCCGC 173885

TeratornLC199500 **GGACGCGCTAAGCCGTTGCGCGCAAGTGCAGGGCCCTTCTTTCGTGCCATGTCGCAGCAACGCGTCCACTGCATGCCTAAAGTTGAACCTGTGGTGGGAGCAGTTGGTCGTGCCCTCTCC** 172807
WF-Teratorn GGACGCGCTAATCCGTTGCGCGCAAGTGCAGGGCCCTTCTTTCGTGCCATGTCGCAGCAACGCGTCCACTGCATGCCTAAAGTTGAACCTGTGGTGGGAGCAGTTGGTCGTGCCCTCTCC 174005

TeratornLC199500 **GAGAGACCTCAGGCTCACGCCTTCAGTTCCGGGGGCGTACAGCACGCCGCCGATCACCGGCACGGGGTCTAGCGCCAAGGCCCTGTACAGGGAGCAGCACAGCTCCCGCTTCGCTTCCAT** 172927
WF-Teratorn GAGAGACCTCAGGCTCACGCCTTCAGTTCCGGGGGCGTACAGCACGCCGCCGATCACTGGCACGGGGTCTAGCGCCAAGGCCCTGTACAGGGAGCAGCACAGCTCCCGCTTCGCTTCCAT 174125

TeratornLC199500 CGTGGACTGAGCTTCCTCGGACGGCGGAGGGGGCCACGGGTGTTACGTCAACGATGTCGCGAGGAGGGGGGGAGGCTCCTCAAGAGGGCCCTCACGACGCCTTTGCAGGGAGGGTACCCG 173047
WF-Teratorn CGTGGACTGAGCTTCCTCAGACGGCGGAGGGGGCCACGGGTGTTATGTCGACGATGTCGCGAGGAGAGGGGGAAGCTCCTCGAGAGGGCCCTCACGACGCCTTTGCAGGGAGGGTACCCG 174245

TeratornLC199500 TCGAACTGTTGATGGAGTGTAAAAGAAAGCGTCAGAAGGCCTCGGTGGAGGCCGCTCTCAAGTCTACGAATCTATACAGGAGAGTTTCCACGATCAGCTACGTCGGCGGAGCGTCCAAGG 173167
WF-Teratorn TCGAACTGTTGATGGAGTGTAAAAGAAAGCGTCAGAAGGCCTCAGTGGAGGCCGCTCTCAAGTCTACGAATCTATACAGGAGAGTTTCCACGATCAGCTACGTCGGCGGAGCGTCCAAGG 174365

TeratornLC199500 TCGGCAGAGCGAGGGTCGACAAGCGGTCGGGGTGCGCAAAGTGCCTGTCGTGTCTGGTGACCTTGGTCAAAGGCAGCAAGCAAAGGAGCAGAGCCGGCGTCAAGCTCCGATGTCCTTGCG 173287
WF-Teratorn TCGGCAGAGCGAGGGTCGACAAGCGGTCGGGGTGCGCAAAGTGCCTGTCGTGTCTGGTGACCTTGGTCAAAGGCAGCAAGCAAAGGAGCAGAGCCGGCGTCAAGCTCCGATGTCCTTGCG 174485

TeratornLC199500 ACGCCAGGGTGTGCGAGACCTGTTTCGTGAATGACCCACACTGGTACCTGACGGAGCCTCACTTCAGCAGCTGCCCGTCCCTCCGCGGGTTCATCTGTTCCGCGTGCGCCTGTTTTGCAG 173407
WF-Teratorn AAGCCAGGGTGTGCGAGACCTGTTTCGTGAATGACCCACACTGGTACCTGACGGAGCCTCACTTCAGCAGCTGCCCGTCCCTCCGCGGGTTCATCTGTTCCACGTGCGCCTGTTTTGCAG 174605

TeratornLC199500 GACTCGTCGAGAAACCATGTTGCCTCTGGATGTGCCCGGGCTGTTCGGACGGGGTCCGAGAGAAGGGCGTCTCCTGCTCGCTGTGCTCAAAACCCGTGCCTCGCGGGAAGCCATCCGCGC 173527
WF-Teratorn GACTCGTCGAGAAACCATGTTGCCTCTGGATGTGCCCGGGCTGTTCGGACGGGGTCCGAGAGAAGGGCGTCGCCTGCTCGCTGTGCTCAAAACCCGTGCCTCGCGGGAAGCCATCCGCGC 174725

TeratornLC199500 GTCGCTCCCTCTGGTGCGAAGCGGCGGATCCGGGTCGACGGGCTGCGGAGCCTCCCTCGGACGCCCCATGT**TTGGACGAGGGCAAAGTCTCAGACGGCAGGGAGCTGATCAAGGCTCGCG** 173647
WF-Teratorn GTCGCTCCCTCTGGTGCGAAGCGGCGGATCCGGGTCGACGGGCTGCGGAGCCTCCCTCGGACGCCCCATGTTTGGACGAGGGCAAAGTCTCAGACGGCAGGGAGCTGATCAAGGCTCGCG 174845

TeratornLC199500 **TCTTGGAGCGTGCGGCAGAGAGAAGACACCTGTCGTCTGCCGACCTCGCCGTCCTCTTCCCGGAGTGCGAGCCTCGGCCGGGCAGATGCTCGTGCGAGGCAAACGGCGACCCCAATCCCA** 173767
WF-Teratorn TCTTGGAGCGTGCGGCAGAGAGAAGACACCTGTCGTCTGCCGACCTCGCCGTCCTCTTCCCGGAGTGCGAGCCTCGGCCGGGCAGATGCTCGTGCGAGGCAAACGGCGACCCCAATCCCA 174965

TeratornLC199500 **TGTGCCCACGCTGCACCCCCGTGCTGGCCGTGTTGGAGCTGCTGGCCAGGAACCCGAAACCGCTCATGGCATCTTTCGCGGAGCGCGCGGTGCCGGGACGAGCCCTGGCCCGTAACACGT** 173887
WF-Teratorn TGTGCCCACGCTGCACCCCCGTGCTGGCCGTGTTGGAGCTGCTGGCCAGGAACCCGAAACCGCTCATGGCATCTTTCGCGGAGCGCGCGGTGCCTGGACGAGCCCTGGCCCGTAACACGT 175085

TeratornLC199500 **ACGACAGGGCCCTGCTCACGGAGATGCTGAATGACAGAGGGCTCCCTAGGTGCGCTAATCGCAACTCCTGCAAAGGAATGCTGGTCAACGTGCGGACGGGCCCCTTGACGACTCTGATGT** 174007
WF-Teratorn ACGACAGGGCCCTGCTCACGGAGATGCTGAATGACAGAGGGCTCCCTTGGTGCGCTAATCGCAACTCCTGCAAAGGAATGCTGGTCAACGTGCGGACGGGCCCCTTGACGACTCTGATGT 175205

TeratornLC199500 **CGCCAGAGGCTTACAAGCAATATGTTCTGGAACGAGATAAGGTGCAGCGCTCAGGCGACGGCGAGCCGAACATCGTGCCGCAGCCTTCCTGCTGCGTCCTGTGCCTGCTGTTTAATCAGT** 174127
WF-Teratorn CCCCAGAGGCTTACAAGCAATATGTTCTGGAACGAGATAAGGTGCAGCGCTCGGGCGACGGCGAGCCGAACATCGTGCCGCAGCCTTCCTGCTGCGTCCTGTGCCTGCTGTTTAATCAGT 175325

TeratornLC199500 **CCGCCGCGGTCGCGCAGATGTGCGGTTCGCAGGGCCTCCGGCTCGAAACCCCCCGCACGGACCCCGTGTACTACTTCAACGTGAAGTTCGCACCAGACGTGGGCCTGCCTGAGGTATCGG** 174247
WF-Teratorn CCGCCGCGGTCTCGCAGATGTGCGGTTCGCAGGGCCTCCGGCTCGAAACCCCCCGCACGGACCCCGTGTACTACTTCAACGTGAAGTTCGCACCAGACGTGGGCCTGCCTGAGGTATCGG 175445

TeratornLC199500 **TGAGAGACTACCAAGGGTTCCTTGTCAACTTTCAAGGTTCCGTGGGAAGCTACAAGCCCGCGTTCTACTACAACTGGAGACACATGCTGGAAATGTTACACAGAGACGCCTCTGGAGCAG** 174367
WF-Teratorn TGAGAGACTACCAAGGGTTCCTTGTCAACTTTCAAGGTTCCGTGGGAAGCTACAAGCCCGCGTTCTACTACAACTGGAGACACATGCTGGAAATGTTACACAGAGACGCCTCTGGAGCAG 175565

TeratornLC199500 **TGACCCTCATTCACCCCGGACAAAGGCAGCTTTAG**ACTCT-------TCGAGGTACGATAAGCTGTGCTCGAAAAGGGCGACTGTTTTGTAAAAGAAAAAAAAAAAAGAGAAAAGCCAAC 174480
WF-Teratorn TGACCCTCATTCACCCCGGACAAAGGCAGCTTTAGACTCTTGTCTCTTCGAGGTACGATAAGCTGTGCTCGAAAAGGGCGACTGTTTTGTAAAAGAAAAAAAAAA--GAGAAAAGCCAAC 175683

TeratornLC199500 CGTTGATCAATTAGGATTTAAAGTATGAACTCACTTACGTGGTCCTGAGCTCTAAGCTGTGCGATCCAGGTTCAGCAACACGAAGCCTTGTGAGATTCTGCAAAGTGGAAAAAGAGGGGA 174600
WF-Teratorn CGTTGATCAATTAGGATTTAAAGTATGAACTCACTTACGTGGTCCTGAGCTCTAAGCTGTGCGATCCAGGTTCAGCAACACGAAGCCTTGTGAGATTCTGCAAAGTGGAAAAAGAGGGGA 175803

TeratornLC199500 GTATGCAAAGGAAAGATTGCCCTCCTCTGTGTTGCTCTTTGCAGCGTGACAACTGGCTTTGCACGGGCCCTTGGTAGAAATCTGCCACGCTACACTGAGTCTTAATCTCTGTGGTTGGAC 174720
WF-Teratorn GTATGCAAAGGAAAGATTGCCCTCCTCTGTGTTGCTCTTTGCAGCGTGACAACTGGCTTTGCACGGGCCCTTGGTAGAAATCTGCCACGCTGCACTGAGTCTTAATCTCTGTGGTTGGAC 175923

TeratornLC199500 CTCACTGTCCTCTTTGCCCCTCCCTCTTTCCCTGCAGCGGGTGTGGGGGGGCTTACAATCAGCCTCCTCCTCCTTCAGAATGAAGGCCCCACCTTCTAACGTTTCCTCTTTTTCTTCCAG 174840
WF-Teratorn CTCACTGTCCTCTTTGCCCCTCCCTCTTTCCCTGCAGCGGGTGTGGGGGGGCTTACAATCAGCCTCCTCCTCCTTCAGAATGAAGGCCCCACCTTCTAACGTTTCCTCTTTTTCTTCCAG 176043

TeratornLC199500 TGTCCTCCTTCTCCCTCTCTGTCTGACTCTGCCTCGCTGAAGTCTGACGGGAGTGCGGCGGACCGGAAACGGAAACTAGACCAGAAGATTTGGATCACAGGCCCCTTTAAAACTTGTTCCG 174960
WF-Teratorn TGTCCTCCTTCTCCCTCTCTGTCTGACTCTGCCTCGCTGAAGTCTGACGGGAGTGCGGCGGACCGGAAACGGAAACTAGACCAGAAGATTTGGATCACAGGCCCCTTTAAAACTTGTTCCG 176163

TeratornLC199500 TGCAAACCAAGACAGGTCATAGTAACCTTACGCGAGTTTAACAAAGACTGGAGGGAAGTTGCTCCGTGAAGAGCGCGACAGTTCGAGAGAGAGAAAAATCAATCCTTCTGCC**ATGGAAAA** 175080
WF-Teratorn TGCAAACCAAGACAGGTCATAGTAACCTTACGCGAGTTTAACAAAGACTGGAGGGAAGTTGCTCCGTGAAGAGCGCGACAGTTCGAGAGAGAGAAAAATCAATCCTTCTGCCATGGAAAA 176283

TeratornLC199500 **GTACTGGAAGACCAGCTTTTGCCTGGGCAAGGGGGGCTTCGGCAAAGTGTTTCTGGGAAAGAGGCGAAGCGACGGGATTTCCGTGGCCCTGAAGATTATCCCGCTCTCGCGTACGAGATG** 175200
WF-Teratorn GTACTGGAAGACCAGCTTTTGCCTGGGCAAGGGGGGCTTCGGCAAAGTGTTTCTGGGAAAGAGGCGAAGCGACGGGATTTCCGTGGCCCTGAAGATTATCCCGCTCTCGCGTACGAGATG 176403

TeratornLC199500 **CTGCGAGGTTGATGGGACCAGTAAGCCATTCGAAGCTGTCGCCTTGGAAATGGTCTCCAAAGACCCTCCGTTTCCTGGGATCATCCGCATGCTGGACTTTTACAAGACCCCGTATAATTG** 175320
WF-Teratorn CTGCGAGGTTGATGGGACCAGTAAGCCATTCGAAGCTGTCGCCTTGGAAATGGTCTCCAAAGACCCTCCGTTTCCTGGGATCATCCGCATGCTGGACTTTTACAAGACCCCGTATAATTG 176523

TeratornLC199500 **GGTGATAGTCACGGAGAAGCCAGAGAACACGCAAGATCTGTACGAATACATCACGATCAGTAAGCGTCTGCGCGAGGACACGGCCAGAGAATTTTTAGGTCAGGTCATCCAGGCCTTGCT** 175440
WF-Teratorn GGTGATAGTCACGGAGAAGCCAGAGAACACGCAAGATCTGTACGAATACATCACGATCAGTAAGCGTCTGCGCGAGGACACGGCCAGAGAATTTTTAGGTCAGGTCATCCAGGCCTTGCT 176643

TeratornLC199500 **CCACTGTCACAGCCGCGGTGTTGTGCACATGGACGTAAAAGACGAAAACCTGCTGGTGGACCTGACGACGCTCGAGCTGAAACTCATCGACTTTGGCTCCGCCGAATTTCTTGAACCCGA** 175560
WF-Teratorn CCACTGTCACAGCCGCGGTGTTGTGCACATGGACGTAAAAGACGAAAACCTGCTGGTGGACCTGACGACGCTCGAGCTGAAACTCATCGACTTTGGCTCCGCCGAATTTCTTGAACCCGA 176763

TeratornLC199500 **CAAAGTGTACGACATGTTCCACGGGACTGCGGCATATGGGCCCCCCGAACGGCTCACACAAGGATGCTACAAAGCTCTTCCATCTGAGGTTTGGTCGTTGGGGATTCTTTTGTACGACAT** 175680
WF-Teratorn CAAAGTGTACGACATGTTCCACGGGACTGCGGCATATGGGCCCCCCGAACGGCTCACACAAGGATGCTACAAAGCTCTTCCATCTGAGGTTTGGTCGTTGGGGATTCTTTTGTACGACAT 176883

TeratornLC199500 **GGTGCACGGGGACATACCATTTGGAAACGACAGGGAGCTCCTACTGGGCGAGCTTAGATTCGAAAAAGACCTCTCTGAGGACTGCGTGGCCCTGATAAAATGGTGTTTGAGCAGGACGCC** 175800
WF-Teratorn GGTGCACGGGGACATACCATTTGGAAACGACAGGGAGCTCCTACTGGGCGAGCTTAGATTCGAAAAAGACCTCTCTGAGGACTGCGTGGCCCTGATAAAATGGTGTTTGAGCAGGACGCC 177003

TeratornLC199500 **CGAAGAGAGACCGATGCTGAAAGACATACTGGCCCACCCTTGGATGCTGAAGCGAGGCTCGGAGGCGCACGAAGGCGACGCCACGGCTGGAGCCCCGAAGCCCACGCGAGGGGATCTCCA** 175920
WF-Teratorn CGAAGAGAGACCGATGCTGAAAGACATACTGGCCCACCCTTGGATGCTGAAGCGAGGCTCGGAGGCGCACGAAGGCGACGCCACGGCTGGAGCCCCGAAGCCCACGCGAGGGGATCTCCA 177123

TeratornLC199500 **GCAATCGGAGCCCGGCTCTGCTCTGGCGGCCCGAGCCCACGATGTCGACACGCCGACCCGTCGAGGGTCCGTCTCGTCCGTGGGAAACGAAAAGCAGCTTCGTCCGGCGATGGCGACGCT** 176040
WF-Teratorn GCAATCGGAGCCCGGCTCTGCTCTGGCGGCCCGAGCCCACGATGTCGACACGCCGACCCGTCGAGGGTCCGTCTCGTCCGTGGGAAACGAAAAGCAGCTTCGTCCGGCGATGGCGACGCT 177243

TeratornLC199500 **CTGCTCCGAGGGCTTCGAATCAATCTGGAGGAGATTGCCGTCCTTCGACCGACTTCCCAGTCGTCAGGGGAGTATCGCCGCAGAGACAGGACTTACCGAGGCTCGTTGCCACTCAGAGGT** 176160
WF-Teratorn CTGCTCCGAGGGCTTCGAATCAATCTGGAGGAGATTGCCGTCCTTCGACCGACTTCCCAGTCGTCAGGGGAGTATCGCCGCAGAGACAGGACTTACCGAGGCTCGTTGCCACTCAGAGGT 177363

TeratornLC199500 **CTCGCAGAGCGGAGCTCATCGTCCGGGTCAGGGAGAGCTCCCAAAAACGGAGTAG**GATGTAGGATGTTGAACAAAATGGCTAACTGCTCATGTTGTATAGATGCCAGTTCCATGATAACT 176280
WF-Teratorn CTCGCAGAGCGGAGCTCATCGTCCGGGTCAGGGAGAGCTCCCAAAAACGGAGTAGGATGTAGGATGTTGAACAAAATGGCTAACTGCTCATGTTGTATAGATGCCAGTTCCATGATAACT 177483

TeratornLC199500 CAATAGAGCTTGTTCCCTGATGCCTACTGTACAGTTTTTTTTTTTGCCTTGTTCGAATAAAAGTCAAACTGGAACCTCAACGTTGCATATTCGCTTTCTTTTTTGCTCATGGGTTTTGT 176400
WF-Teratorn CAATAGAGCTTGTTCCCTGATGCCTACTGTACAGTTTTTTTTTT-GCCTTGTTCGAATAAAAGTCAAACTGGAACCTCAACGTTGCATATTCGCTTTCTTTTTTGCTCATGGGTTTTGT 177602

TeratornLC199500 CCCAACATGACTATTTGAAGAGATATTTTTCTAATTATTGGATACATCACACAATCTCGGTTTTTAAAGATTGAAAAATTAAAAAAAAAAATACATTGGTGTGTACATTTTTAATCGTAT 176520
WF-Teratorn CCCAACATGACTATTTGAAGAGATATTTTTCTAATTATTGGATACATCACACAATCTCGGTTTTTAAAGATTGAAAAATTAAAAAAAAAAATACATTGGTGTGTACATTTTTAATCGTAT 177722

TeratornLC199500 AATGGTCAGTATATACACGTTAGATAAAGTGATTACCACACACAAAGGTTACATGATTTTGGTCAGCATATACACAGGTTAAATGAAGTGAATACCATATACAAGGAATGGCAGCCAGAG 176640
WF-Teratorn AATGGTCAGTATATACACGTTAGATAAAGTGATTACCACACACAAAGGTTACATGATTTTGGTCAGCATATACACAGGTTAAATGAAGTGAATACCATATACAAGGAATGGCAGCCAGAG 177842

TeratornLC199500 CTGTCTGCTGTGACGTTACTTTGATGACCCAGACCTCCCGCTCGCCGTAGCACCAACAGAGTGGGGTGGAAATGAAACGGTCCAAAGAACCTTCATGCAGCTCTACTCAAAAAGAAAAAC 176760
WF-Teratorn CTGTCTGCTGTGACGTTACTTTGATGACCCAGACCTCCCGCTCGCCGTAGCACCAACAGAGTGGGGTGGAAATGAAACGGTCCAAAGAACCTTCATGCAGCTCTACTCAAAAAGAAAAAC 177962

TeratornLC199500 AAAAACAAAACAAGAGTGTGTTGGAAAAGTCAGCAGAAGTCGATGGTATTTACTTACCAAATACCAAAAGAAACAAAATTAGTTGTAGCGTTAGATACAAGGGGGCTTACAAAGAGGCTC 176880
WF-Teratorn AAAAACAAAACAAGAGTGTGTTGGAAAAGTCAGCAGAAGTCGATGGTATTTACTTACCAAATACCAAAAGAAACAAAATTAGTTGTAGCGTTAGATACAAGGGGGCTTACAAAGAGGCTC 178082

TeratornLC199500 CGACCCAAGTTTAGGGACACTGACCTGAGTTCTGAGACTCAGACCATCAGAGGAAGCTTTGGACCTTAATCTCTCACAAAAAGAGTCTCAGTGAAAATCTGCTAAAAATGTATACTGAAA 177000
WF-Teratorn CGACCCAAGTTTAGGGACACTGACCTGAGTTCTGAGACTCAGACCATCAGAGGAAGCTTTGGACCTTAATCTCTCACAAAAAGAGTCTCAGTGAAAATCTGCTAAAAATGTATACTGAAA 178202

TeratornLC199500 AAAAAAAAAAAATTGTTCCACACGCTCTCTTGTAGGTCCGTTCGACCCCAACACTAACTAAAACTAAACAGTAGCTCCGCACACTGTCTCGCGGGGTCCGTGAGACCCAACGCCGACTGG 177120
WF-Teratorn AAAAAAAAAAAATTGTTCCACACGCTCTCTTGTAGGTCCGTTCGACCCCAACACTAACTAAAACTAAACAGTAGCTCCGCACACTGTCTCGCGGGGTCCGTGAGACCCAACGCCGACTGG 178322

TeratornLC199500 AACCAAACAGTGGGTCCGCGCGCTGTCTCGCGGGGTCCGTTAGACCCCGAACAGAATTGGAACGAAACACAGGCTCTGCGCGCTGTCTCGCGGGGTCCGTGAAACCCCGTACAGAATTGG 177240
WF-Teratorn AACCAAACAGTGGGTCCGCGCGCTGTCTCGCGGGGTCCGTTAGACCCCGAACAGAATTGGAACGAAACACAGGCTCTGCGCGCTGTCTCGCGGGGTCCGTGAAACCCCGTACAGAATTGG 178442

TeratornLC199500 AACGAAACACAGGCTCCGCGCGCTGTTTGCACGGTCCGTTAAACCCCGAACAGAAGTGGAACGAAACACAGGCTCCGCGTGCTGTCTCGCGGGGTCCGTGAGACCCCGTACAGAATTGGA 177360
WF-Teratorn AACGAAACACAGGCTCCGCGCGCTGTTTGCACGGTCCGTTAAACCCCGAACAGAAGTGGAACGAAACACAGGCTCCGCGTGCTGTCTCGCGGGGTCCGTGAGACCCCGTACAGAATTGGA 178562

TeratornLC199500 GCGAAACACAGGCACCTCGCGCTGTCTCGCATGGTCCCAGCGACCTCGTCTCGATTTGGAACCAAACAGTGGCACCTCGCGCTGTCTCGCGTGGTCCGTGAGACCCCGTACAGAATTGGA 177480
WF-Teratorn GCGAAACACAGGCACCTCGCGCTGTCTCGCATGGTCCCAGCGACCCCGTCTCGATTTGGAACCAAACAGTGGCACCTCGCGCTGTCTCGCGTGGTCCGTGAGACCCCGTACAGAATTGGA 178682

TeratornLC199500 ACCAAACAGTGGCACCTCGCGCTGTCTCGCGGGGTCCGTGAGACCCCGTAGAGAGTTGGAACGAAACACAGGCTCTGCGCGCTGTCTCGCGGGGTCCGTGAGACCCCGTACAGAGTTGGA 177600
WF-Teratorn ACCAAACAGTGGCACCTCGCGCTGTCTCGCGGGGTCCGTGAGACCCCGTAGAGAGTTGGAACGAAACACAGGCTCTGCGCGCTGTCTCGCGGGGTCCGTGAGACCCCGTACAGAGTTGGA 178802

TeratornLC199500 ACGAAACACAGGCTCCGAGCGCTGTCTCCCGGGGTCCGTGAGACCCCGTACAGAATTGGAAGGAAACACAGGCTCCGCGCGCTGTCTTGCACGGTCCGTGAGACCCCGAACAGAATTGGA 177720
WF-Teratorn ACGAAACACAGGCTCCGAGCGCTGTCTCCCGGGGTCCGTGAGACCCCGTACAGAATTGGAAGGAAACACAGGCTCCGCGCGCTGTCTTGCACGGTCCGTGAGACCCCGAACAGAATTGGA 178922

TeratornLC199500 ACCAAACTGTGGCTCCGAGCGCTGTCTCGCGGGGTCCGTGAGACCCCGTACAGAG------------------------------------------------------------------ 177775
WF-Teratorn ACCAAACTGTGGCTCCGAGCGCTGTCTCGCGGGGTCCGTGAGACCCCGTACAGAGTTGGAACGAAACACAGGCTCCGCGCGCTGTCTCGCGGGGTCCGTGAGACCCCGTACAGAGTTGGA 179042

TeratornLC199500 ------------------------------------------------------------------------------------------------------------------------ 177775
WF-Teratorn ACGAAACACAGGCTCCGAGCGCTGTCTCCCGGGGTCCGTGAGACCCCGTACAGAATTGGAACGAAACACAGGCTCCGCGCGCTGTTTGCACGGTCCGTTAGACCCCGAACAGAAGTGGAA 179162

TeratornLC199500 ------------------------------------------------------------------------------------------------------------------------ 177775
WF-Teratorn CGAAACACAGGCTCCGCGCGCTGTCTCGCGGGGTCCGTGAGACCCCGTACAGAATTGGAACGAAACACAGGCTCCGCGCGCTGTCTTGCACGGTCCGTGAGACCCCGAACAGAATTGGAA 179282

```
TeratornLC199500  ------------------------------------------------------------------------------------------------------------------------  177775
WF-Teratorn       CCAAACTGTGGCTCCGAGCGCTGTCTCGCGGGGTCCGTGAGACCCCGTACAGAGTTGGAACGAAACACAGGCTCCGCGCGCTGTCTCGCGGGGTCCGTGAGACCCCGTACAGAGTTGGAA  179402

TeratornLC199500  ------------------------------------------------------------------------------------------------------------------------  177775
WF-Teratorn       CGAAACACAGGCTCCGAGCGCTGTCTCCCGGGGTCCGTGAGACCCCGTACAGAATTGGAACGAAACACAGGCTCCGCGCGCTGTTTGCACGGTCCGTTAGACCCCGAACAGAAGTGGAAC  179522

TeratornLC199500  ------------------------------------------------------------------------------------------------------------------------  177775
WF-Teratorn       GAAACACAGGCTCCGCGCGCTGTCTCGCGGGGTCCGTGAGACCCCGTACAGAATTGGAACGAAACACAGGCTCCGCGCGCTGTCTCGCACGGTCCCAGCGACCCCGTCTCGGTTTGGAAC  179642

TeratornLC199500  ------------------------------------------------------------------------------------------------------------------------  177775
WF-Teratorn       CAAACAGTTGCACCTCGGGCTGTCTCGCGTGGTCCGTGAGACCCCGTCTCGATTTGGAACCAAACAGTGGCACCTCGCGCTGTCTCGCGGGGTCCGTGAGACCCCGTACAGAGTTGGAAC  179762

TeratornLC199500  ------------------------------------------------------------------------------------------------------------------------  177775
WF-Teratorn       GAAACACAGGCTCCGCGCGCTGTCTCGCGGGGTCCGTGAGACCCCGTACAGAGTTGGAACGAAACACAGGCTCCGACCGCTGTCTCCCGGGGTCCGTGAGACCCCGTACAGAATTGGAAC  179882

TeratornLC199500  ------------------------------------------------------------------------------------------------------------------------  177775
WF-Teratorn       GAAACACAGGCTCCGCGCGCTGTTTGCACGGTCCGTTAGACCCCGAACAGAAGTGGAACGAAACACAGGCTCCGCGCGCTGTCTCGCGGGGTCCGTGAGACCCCGTACAGAATTGGAACG  180002

TeratornLC199500  ------------------------------------------------------------------------------------------------------------------------  177775
WF-Teratorn       AAACACAGGCTCCGAGCGCTGTCTCCCGGGGTCCGTGAGACCCCGTACAGAATTGGAAGGAAACACAGGCTCCGCGCGCTGTCTTGCACGGTCCGTGAGACCCCGAACAGAATTGGAACC  180122

TeratornLC199500  ----------------------------------------------------TTGGAACGAAACACAGGCTCCGCGCGCTGTCTCGCGGGGTCCGTGAGACCCCGTACAGAGTTGGAACG  177843
WF-Teratorn       AAACTGTGGCTCCGAGCGCTGTCTCGCGGGGTCCGTGAGACCCCGTACAGAGTTGGAACGAAACACAGGCTCCGCGCGCTGTCTCGCGGGGTCCGTGAGACCCCGTACAGAGTTGGAACG  180242

TeratornLC199500  AAACACAGGCTCCGAGCGCTGTCTCCCGGGGTCCGTGAGACCCCGTACAGAATTGGAACGAAACACAGGCTCCGCGCGCTGTTTGCACGGTCCGTTAGACCCCGAACAGAAGTGGAACGA  177963
WF-Teratorn       AAACACAGGCTCCGAGCGCTGTCTCCCGGGGTCCGTGAGACCCCGTACAGAATTGGAACGAAACACAGGCTCCGCGCGCTGTTTGCACGGTCCGTTAGACCCCGAACAGAAGTGGAACGA  180362

TeratornLC199500  AACACAGGCTCCGCGCGCTGTCTCGCGGGGTCCGTGAGACCCCGTACAGAATTGGAACGAAACACAGGCTCCGCGCGCTGTCTTGCACGGTCCGTGAGACCCCGAACAGAATTGGAACCA  178083
WF-Teratorn       AACACAGGCTCCGCGCGCTGTCTCGCGGGGTCCGTGAGACCCCGTACAGAATTGGAACGAAACACAGGCTCCGCGCGCTGTCTTGCACGGTCCGTGAGACCCCGAACAGAATTGGAACCA  180482

TeratornLC199500  AACTGTGGCTCCGAGCGCTGTCTCGCGGGGTCCGTGAGACCCCGTACAGAGTTGGAACGAAACACAGGCTCCGCGCGCTGTCTCGCGGGGTCCGTGAGACCCCGTACAGAGTTGGAACGA  178203
WF-Teratorn       AACTGTGGCTCCGAGCGCTGTCTCGCGGGGTCCGTGAGACCCCGTACAGAGTTGGAACGAAACACAGGCTCCGCGCGCTGTCTCGCGGGGTCCGTGAGACCCCGTACAGAGTTGGAACGA  180602

TeratornLC199500  AACACAGGCTCCGAGCGCTGTCTCCCGGGGTCCGTGAGACCCCGTACAGAATTGGAACGAAACACAGGCTCCGCGCGCTGTTTGCACGGTCCGTTAGACCCCGAACAGAAGTGGAACGAA  178323
WF-Teratorn       AACACAGGCTCCGAGCGCTGTCTCCCGGGGTCCGTGAGACCCCGTACAGAATTGGAACGAAACACAGGCTCCGCGCGCTGTTTGCACGGTCCGTTAGACCCCGAACAGAAGTGGAACGAA  180722

TeratornLC199500  ACACAGGCTCCGCGCGCTGTCTCGCGGGGTCCGTGAGACCCCGTACAGAATTGGAACGAAACACAGGCTCCGCGCGCTGTCTCGCACGGTCCCAGCGACCCCGTCTCGGTTTGGAACCAA  178443
WF-Teratorn       ACACAGGCTCCGCGCGCTGTCTCGCGGGGTCCGTGAGACCCCGTACAGAATTGGAACGAAACACAGGCTCCGCGCGCTGTCTCGCACGGTCCCAGCGACCCCGTCTCGGTTTGGAACCAA  180842

TeratornLC199500  ACAGTTGCACCTCGGGCTGTCTCGCGTGGTCCGTGAGACCCCGTCTCGATTTGGAACCAAACAGTGGCACCTCGCGCTGTCTCGCGGGGTCCGTGAGACCCCGTACAGAGTTGGAACGAA  178563
WF-Teratorn       ACAGTTGCACCTCGGGCTGTCTCGCGTGGTCCGTGAGACCCCGTCTCGATTTGGAACCAAACAGTGGCACCTCGCGCTGTCTCGCGGGGTCCGTGAGACCCCGTACAGAGTTGGAACGAA  180962

TeratornLC199500  ACACAGGCTCCGCGCGCTGTCTCGCGGGGTCCGTGAGACCCCGTACAGAGTTGGAACGAAACACAGGCTCCGACCGCTGTCTCCCGGGGTCCGTGAGACCCCGTACAGAATTGGAACGAA  178683
WF-Teratorn       ACACAGGCTCCGCGCGCTGTCTCGCGGGGTCCGTGAGACCCCGTACAGAGTTGGAACGAAACACAGGCTCCGACCGCTGTCTCCCGGGGTCCGTGAGACCCCGTACAGAATTGGAACGAA  181082

TeratornLC199500  ACACAGGCTCCGCGCGCTGTTTGCACGGTCCGTTAGACCCCGAACAGAAGTGGAACGAAACACAGGCTCCGCGCGCTGTCTCGCGGGGTCCGTGAGACCCCGTACAGAATTGGAACGAAA  178803
WF-Teratorn       ACACAGGCTCCGCGCGCTGTTTGCACGGTCCGTTAGACCCCGAACAGAAGTGGAACGAAACACAGGCTCCGCGCGCTGTCTCGCGGGGTCCGTGAGACCCCGTACAGAATTGGAACGAAA  181202

TeratornLC199500  CACAGGCTCCGCGCGCTGTCTCGCACGGTCCCAGCGACCCCGTCTCGGTTTGGAACCAAACAGTTGCACCTCGGGCTGTCTCGCGTGGTCCGTGAGACCCCGTACAGAATTGGAACCAAA  178923
WF-Teratorn       CACAGGCTCCGCGCGCTGTCTCGCACGGTCCCAGCGACCCCGTCTCGGTTTGGAACCAAACAGTTGCACCTCGGGCTGTCTCGCGTGGTCCGTGAGACCCCGTACAGAATTGGAACCAAA  181322

TeratornLC199500  CAGTGGCACCTCGCGCTGTCTCGCGGGGTCCGTGAGACCCCGTACAGAGTTGGAACGAAACACAGGCTCCGCGCGCTGTCTCGCGGGGTCCGTGAGACCCCGTACAGAGTTGGAACGAAA  179043
WF-Teratorn       CAGTGGCACCTCGCGCTGTCTCGCGGGGTCCGTGAGACCCCGTACAGAGTTGGAACGAAACACAGGCTCCGCGCGCTGTCTCGCGGGGTCCGTGAGACCCCGTACAGAGTTGGAACGAAA  181442

TeratornLC199500  CACAGGCTCCGAGCGCTGTCTCCCGGGGTCCGTGAGACCCCGTACAGAGTTGGAACGAAACACAGGCTCCGAGCGCTGTCTCCCGGGGTCCGTGAGACCCCGTACAGAGTTGGAAGGAAA  179163
WF-Teratorn       CACAGGCTCCGAGCGCTGTCTCCCGGGGTCCGTGAGACCCCGTACAGAGTTGGAACGAAACACAGGCTCCGAGCGCTGTCTCCCGGGGTCCGTGAGACCCCGTACAGAGTTGGAAGGAAA  181562
```

TeratornLC199500 CACAGGCTCCGAGCGCTGTCTCGCGGGGTCCGTGAGACCCCGTACAGAATTGGAACGAAACACAGGCTCCGCGCGCTGTCTAACACGGTCCGTGAGACCCCGAACATAAGTGGAACGAAA 179283
WF-Teratorn CACAGGCTCCGAGCGCTGTCTCGCGGGGTCCGTGAGACCCCGTACAGAATTGGAACGAAACACAGGCTCCGCGCGCTGTCTAACACGGTCCGTGAGACCCCGAACATAAGTGGAACGAAA 181682

TeratornLC199500 CACAGGCTCCGAGCGCTGTCTCGCGGGGTCCGTGAGACCCCGTACAGAATTGGAACGAAACACAGGCTCCGCGCGCTGTCTAACACGGTCCGTGAGACCCCGAACAGAATTGGAACGAAA 179403
WF-Teratorn CACAGGCTCCGAGCGCTGTCTCGCGGGGTCCGTGAGACCCCGTACAGAATTGGAACGAAACACAGGCTCCGCGCGCTGTCTAACACGGTCCGTGAGACCCCGAACAGAATTGGAACGAAA 181802

TeratornLC199500 CACAGGCTCCGCGCGCTGTTTAGCGGGG------------------------------------------------------------------------------------------- 179431
WF-Teratorn CACAGGCTCCGCGCGCTGTTTAGCGGGGCCCGTGAGACCCCGTACAGAATTGGAACGAAACACAGGCTCCGCGCACTGTGGCTCCGAGCGCTGTCTCGCACGGTCCGTGAGACCCCGAAC 181922

TeratornLC199500 -------------------------------------------TCCGTGAGACCCCGTACAGAATTGTAACCAAACAGTGGCACCTCGCGCTGTCTCGCGGGGTCCGTGAGACCCCGTAC 179508
WF-Teratorn AGAATTGGAACGAAACACAGGCTCCGCGCGCTGTTTAGCGGGGTCCGTGAGACCCCGTACAGAATTGTAACCAAACAGTGGCACCTCGCGCTGTCTCGCGGGGTCCGTGAGACCCCGTAC 182042

TeratornLC199500 AGAATTGGAACGAAACACAGGCTCCGCGCGCTGTCTCGCGGAGTCCGTGAGACCCCGTACAGAATTGGAACGAAACACAGGCTCCGAGCGCTGTCTCCCGGGGTCCGTGAGACCCTGTAC 179628
WF-Teratorn AGAATTGGAACGAAACACAGGCTCCGCGCGCTGTCTCGCGGAGTCCGTGAGACCCCGTACAGAATTGGAACGAAACACAGGCTCCGAGCGCTGTCTCCCGGGGTCCGTGAGACCCCGTAC 182162

TeratornLC199500 AGAATTGGAACGAAACACAGGCTCCGCGCGCTGTCTCGCACGGTCCCAGCGACCCCATCTCGATTTGGACCCAAACCGTGGCACCTCGCGCTGTCTCGCAGGGTCCGTGAGACCCCGTAC 179748
WF-Teratorn AGAATTGGAAGGAAACACAGGCTCCGCGCGCTGTCTCGCACGGTCCCAGCGACCCCATCTCGATTTGAACCCAAACCGTGGCACCTCGCGCTGTCTCGCAGGGTCCGTGAGACCCCGTAC 182282

TeratornLC199500 AGAGTTGGAACGAAACACAGGCTCCGCGCGCTGTCTCGTACGGTCCGTGAGACCCCGAACAGAGTTGGAACCAAACTGTGGCTCCGCGCGCTGTCTCGCGGGGTCCGTGAGACCCCGTAC 179868
WF-Teratorn AGAGTTGGAACGAAACACAGGCTCCGCGCGCTGTCTCGTACGGTCTGTGAGACCCCGAACAGAGTTGGAACCAAACTGTGGCTCCGCGCGCTGTCTCGCGGGGTCCGTGAGACCCCGTAC 182402

TeratornLC199500 AGAATTGGAACGAAACACAGGCTCCGCGCGCTGTCTCGGGGGTCCGTGAGACCCCGTACAGAATTGGAACGAAACACAGGCTCCGCGCGCTGTCTCGCGGGGTCCCAGCGACCCC----- 179983
WF-Teratorn AGAATTGGAACGAAACACAGGCTCCGCGCGCTGTCTCGGGGGTCCGTGAGACCCCGTACAGAATTGGAACGAAACACAGGCTCCGCGCGCTGTCTCGCGGGGTCCCAGCGACCCCGTACA 182522

TeratornLC199500 ------------------------------------------------------------------------------------------------------------------------ 179983
WF-Teratorn GAGTTGGAACCAAACAGTGGCACCTCGCGATGTCTCGCGGGGTCCGTGAGACCCCGTACAGAATTGGAACGAAACACAGGCTCCGCGCGCTGTCTCGGGGGTCCGTGAGACCCCGTACAG 182642

TeratornLC199500 ----------------------------------------------------------GTACAGAGTTGGAACCAAACAGTGGCACCTCGCGATGTCTCGCACGGTCCGTGAGACCCCGAACAG 180049
WF-Teratorn AATTGGAACGAAACACAGGCTCCGCGCGCTGTCTCGCGGGGTCCCAGCGACCCCGTACAGAGTTGGAACCAAACAGTGGCACCTCGCGATGTCTCGCACGGTCCGTGAGACCCCGAACAG 182762

TeratornLC199500 AATTGGAACGAAACACAGGCTCCGAGCGCTGTCTCCCGGGGTCCGTGAGACCCCGTACAGAGTTGGAACGAAACACAGGCTCCGCGCGCTGTCTCGCACGGTCCCAGCGACCCCGTACCG 180169
WF-Teratorn AATTGGAACGAAACACAGGCTCCGAGCGCTGTCTCCCGGGGTCCGTGAGACCCCGTACAGAGTTGGAACGAAACACAGGCTCCGCGCGCTGTCTCGCACGGTCCCAGCGACCCCGTACCG 182882

TeratornLC199500 ATTTGGAACCAAACAGTGGCACCTCGCGCTGTCTCGCGGGGTCCGTGAGACCCCGTACAGAGTTGGAACCATCAGTTTCTCTTTGAAAGCTCAAACACGAACCTGTTTCTGTAGATCCAT 180289
WF-Teratorn ATTTGGAACCAAACAGTGGCACCTCGCGCTGTCTCGCGGGGTCCGTGAGACCCCGTACAGAGTTGGAACCATCAGTTTTTCTTTGAAAGCTCAAACACGAACCTGTTTCTGTAGATCCAT 183002

TeratornLC199500 GACATGTGGGTTACAGTGGGCCACAGGCTGGTTTACCTTCTCCCTTTGCTGCGGATCAGCAGCACCAGTGTTGCTGGCTTGTGACTGTGGGGCAAGTAAGATTAACTGAATA**TCAAAAAA** 180409
WF-Teratorn GACATGTGGGTTACAGTGGGCCACAGGCTGGTTTACCTTCTCCCTTTGCTGCGGATCAGCAGCACCAGTGTTGCTGGCTTGTGACTGTGGGGCAAGTAAGATTAACTGAATATCAAAAAA 183122

TeratornLC199500 **AATAAGTTGTTTCAAAGAAGTTGGATTTGACGAACATCCCACGTTGACCAACCTCAGAGTTCTTCATTGTTTTGGTCCTCGCCTCGTAGTTTAGCATGTCAAACGGGTCGAGCCACTCAC** 180529
WF-Teratorn A-TAAGTTGTTTCAAAGAAGTTGGATTTGACGAACATCCCACGTTGACCAACCTCAGAGTTCTTCATTGTTTTGGTCCTCGCCTCGTAGTTTAGCATGTCAAACGGGTCGAGCCACTCAC 183241

TeratornLC199500 **CGTTGACTGTCTGTCCGGAATTGGACAGAACCAGACTGCAGGACAGCAAAAACAACCAGAGCAT**CTTCTCTGGAGACCAAATTGGTTTGAACTAACTAGCTAGTAACGGAAGAAGAAAAC 180649
WF-Teratorn CGTTGACTGTCTGTCCGGAATTGGACAGAAACAGACTGCAGGACAGCAAAAACAACCAGAGCATCTTCTCTGGAGACCAAATTGGTTTGAACTAACTAGCTAGTAACGGAAGAAGAAAAC 183361

TeratornLC199500 TATGGAAGCCGAAATTGCTTCCGGTTATTTTTTT-AAAAAAAAAAGCAGCAGAAAACCATTTAGGTTTGTTCATTTGTTTCTCTTGACCTTTCTTGATCACATAAAAAAAAA---GAAAC 180765
WF-Teratorn TATGGAAGCCGAAATTGCTTCCGGTTATTTTTTTTAAAAAAAAAAGCAGCAGAAAACCATTTAGGTTTGTTCATTTGTTTCTCTTGACCTTTCTTGATCACATAAAAAAAAAAAAGAAAC 183481

TeratornLC199500 GTAAACACAAACAGTACACTTCGAAAAGCCTCAAGGGTGATGTAGAAAAGTGAATGCTAAAGAAAAAAAAAA-TACTTGGGGTCAATCGCACCCCATGAGCCAGCGGGTGGAAGGGAACC 180884
WF-Teratorn GTAAACACAAACAGTACACTTCGAAAAGCCTCAAGGGTGATGTAGAAAAGTGAATGCTAAAGAAAAAAAAAAATACTTGGGGTCAATCGCACCCCATGAGCCAGCGGGTGGAAGGGAACC 183601

TeratornLC199500 GGAAAAGCGAGAGCCCCGGTGGTGGGCCATGAGAGAAGAGTGAATGCTAAAAAAAAAAAAAAAATACTTGGGGTCAATCGCACCCCATGAGCCAGCGGGTGAAAGGGAACCGGAAAAACGA 181004
WF-Teratorn GGAAAAACGAGACCCCCGGTGGTGGGCCATGAGAGAAGAGTGAATGCTAAAGAAAAAAAAAAAATACTTGGGGTCAATCGCACCCCATGAGCCAGCGGGTGGAAGGGAACCGGAAAAACGA 183721

TeratornLC199500 GACCCCCGGTGGTGGGCCATGAGAGAAGAGTGAATGCTAAAGAAAAAAAAAAATACTTGGGGTCAATCGCACCCCATGAGCCAGCGGGTGGAAGGGAACCGGAAAAACGAGACCCCCGGT 181124
WF-Teratorn GACCCCCGGTGGTGGGCCATGAGAGAAGAGTGAATGCTAAAGAAAAAAAAAAATACTTGGGGTCAATCGCACCCCATGAGCCAGCGGGTGAAAGGGAACCGGAAAA-CGAGACCCCCGGT 183840

```
TeratornLC199500  GGTGGGCCATGAGAGAAGAGTGAATGCTACAGAAAAAAAAAAAAAATACTTGGGGTCAATCGCACCCCATGAGCCAGCGGGTGAAAGGGAACCGGTAAAGCGAGACCCCAGGTGGTGGGCC  181244
WF-Teratorn       GGTGGGCCATGAGAGAAGAGTGAATGCTACAGAAAAAAAAAAAA--TACTTGGGGTCAATCGCACCCCATGAGCCAGCGGGTGAAAGGGAACCGGTAAAGCGAGACCCCAGGTGGTGGGCC  183958

TeratornLC199500  ATGAGAGAAGAGTGAATGCTAAAGAAAAAAAAAAAAAAAAAAACTTGGGGTCAATCGCACCCCATGAGCCAGCGCCAATCCCCTCAGTGGCCACACAAAAAAAAAAAAAAAAAAAACTTGGG  181364
WF-Teratorn       ATGAGAGAAGAGTGAATGCTAAAGAAAAAAAAAAAA-------CTTGGGGTCAATCGCACCCCATGAGCCAGCGCCAATCCCCTCAGTGGCCACACAAAAAAAAAAAAAAAAA---CTTGGG  184068

TeratornLC199500  GTCAATCGAACCCCACGAACCAGCGGGAGGAAGGTGGTTGCTAAATAAGTGGCGCCCCTGGTGGCTGAGGGCAATAAAAGAGAATGGGGTATTAGTGACCCCTAGAGGCAGGGTATAGAA  181484
WF-Teratorn       GTCAATCGAACCCCACGAACCAGCGGGAGGAAGGTGGTTGCTAAAAAAGTGGCGCCCCTGGTGGCTGAGGGCAATAAAAGAGAATGGGGTATTAGTGACCCCTAGAGGCAGGGTATAGAA  184188

TeratornLC199500  ATGAACGGTGGAAAACAAGACCCCTGGTGGTCGGGGGGTATAACTAAAATGGAGAAATGGCGGCCTCTGCTGGACGGGGACAAAAA-GAAAAGGGGGGTTTGGCGCCCTCTACCAGTCGG  181603
WF-Teratorn       ATGAACGGTGGAAAACAAGACCCCTGGTGGTCGGGGGGTATAACTAAAATGGAGAAATGGCGGCCTCTGCTGGACGGGGACAAAAAAGAAAAGGGGGGTTTGGCGCCCTCTACCAGTCGG  184308

TeratornLC199500  GGATATAGAGGGTGCCTGAAACGTACAAAAAAATTTTTTCAGGAACTCGACCCTCTTCGATTGCGGGAGGGTGTACGGGGGTGTACCTGTCTGGGGTCAATACCAAAGGGGGGTATGCGT  181723
WF-Teratorn       GGATATAGAGGGTGCCTGAAACGTACAAAAAATTTTTTTCAGGAACTCGACCCTCTTCGATTGCGGTGGGGTGTACCGGG-TGTACCTGTTTGGGGTCAATACCAAAGGGGGGTATGCGT  184427

TeratornLC199500  TTTTAAAATGACCCCCCCTCCTTACCTTTTTTTTAATCCATGCTTTTCTTTCCCCCTAAGACAGCACAAGGG                                                  181795
WF-Teratorn       TTTTAAAATGACCCCCCCTCCTTACCTTTTTTTTAATCCATGCTTTTCTTTCCCCCTAAGACAGCACAAGGG                                                  184499
```
